# Genome-enabled insights into the biology of thrips as crop pests

**DOI:** 10.1101/2020.02.12.941716

**Authors:** Dorith Rotenberg, Aaron A. Baumann, Sulley Ben-Mahmoud, Olivier Christiaens, Wannes Dermauw, Panagiotis Ioannidis, Chris G.C. Jacobs, Iris M. Vargas Jentzsch, Jonathan E. Oliver, Monica F. Poelchau, Swapna Priya Rajarapu, Derek J. Schneweis, Simon Snoeck, Clauvis N.T. Taning, Dong Wei, Shirani M. K. Widana-Gamage, Daniel S.T. Hughes, Shwetha C. Murali, Sam Bailey, Nicolas E. Bejerman, Christopher J Holmes, Emily C. Jennings, Andrew J. Rosendale, Andrew Rosselot, Kaylee Hervey, Brandi A. Schneweis, Sammy Cheng, Christopher Childers, Felipe A. Simão, Ralf G. Dietzgen, Hsu Chao, Huyen Dinh, HarshaVardhan Doddapaneni, Shannon Dugan, Yi Han, Sandra L. Lee, Donna M. Muzny, Jiaxin Qu, Kim C. Worley, Joshua B. Benoit, Markus Friedrich, Jeffery W. Jones, Kristen A. Panfilio, Yoonseong Park, Hugh M. Robertson, Guy Smagghe, Diane E. Ullman, Maurijn van der Zee, Thomas Van Leeuwen, Jan A. Veenstra, Robert M. Waterhouse, Matthew T. Weirauch, John H. Werren, Anna E. Whitfield, Evgeny M. Zdobnov, Richard A. Gibbs, Stephen Richards

**Affiliations:** Department of Entomology and Plant Pathology, North Carolina State University, Raleigh, NC 27606, USA; University of Tennessee, College of Veterinary Medicine, Virology Section, A239 VTH, 2407 River Drive, Knoxville TN 37996, USA; University of California Davis, Department of Entomology and Nematology, Davis, CA 95616, USA; Laboratory of Agrozoology, Department of Plants and Crops, Ghent University, Coupure Links 653, 9000 Ghent, Belgium; Institute of Molecular Biology and Biotechnology, Foundation for Research and Technology Hellas, Vassilika Vouton, 70013, Heraklion, Greece; Department of Genetic Medicine and Development, University of Geneva Medical School, and Swiss Institute of Bioinformatics, Geneva, Switzerland; Instute of Biology, Leiden University, 2333 BE Leiden, The Netherlands; Institute for Zoology: Developmental Biology, University of Cologne, 50674 Cologne, Germany; Department of Plant Pathology, University of Georgia - Tifton Campus, Tifton, GA 31793-5737, USA; National Agricultural Library, USDA-ARS, Beltsville, MD 20705, USA; Department of Plant Pathology, Kansas State University, Manhattan KS, 66506, USA; Department of Biology, University of Washington, Seattle, WA 98105, USA; Chongqing Key Laboratory of Entomology and Pest Control Engineering, College of Plant Protection, Southwest University, Chongqing, China; International Joint Laboratory of China-Belgium on Sustainable Crop Pest Control, Academy of Agricultural Sciences, Southwest University, Chongqing, China, Ghent University, Ghent, Belgium; Department of Botany, University of Ruhuna, Matara, Sri Lanka; Human Genome Sequencing Center, Department of Human and Molecular Genetics, Baylor College of Medicine, One Baylor Plaza, Houston, TX 77030, USA; Department of Biological Sciences, University of Cincinnati, Cincinnati, OH 45221, USA; IPAVE-CIAP-INTA, 5020 Cordoba, Argentina; Department of Biology, University of Rochester, Rochester, NY 14627, USA; Queensland Alliance for Agriculture and Food Innovation, The University of Queensland, St. Lucia QLD 4072, Australia; Department of Biological Sciences, Wayne State University, MI 48202, USA; School of Life Sciences, University of Warwick, Gibbet Hill Campus, Coventry CV4 7AL, United Kingdom; Department of Entomology, Kansas State University, Manhattan, KS 66506, USA; Department of Entomology, University of Illinois at Urbana-Champaign, Urbana, IL 61801, USA; Institute of Biology, Leiden University, 2333 BE Leiden, The Netherlands; INCIA UMR 5287 CNRS, University of Bordeaux, Pessac, France; Department of Ecology and Evolution, Swiss Institute of Bioinformatics, University of Lausanne, 1015, Lausanne, Switzerland; Center for Autoimmune Genomics and Etiology, Divisions of Biomedical Informatics and Developmental Biology, Cincinnati Children’s Hospital Medical Center, Cincinnati, Ohio, 45229, USA; Department of Pediatrics, University of Cincinnati, College of Medicine, Cincinnati, Ohio, 45229, USA

**Keywords:** Thysanoptera, western flower thrips, hemipteroid assemblage, insect genomics, tospovirus, virus-vector interactions, salivary glands, chemosensory receptors, opsins, detoxification, innate immunity

## Abstract

**Background:** The western flower thrips, *Frankliniella occidentalis* (Pergande), is a globally invasive pest and plant virus vector on a wide array of food, fiber and ornamental crops. While there are numerous studies centered on thrips pest and vector biology, feeding behaviors, ecology, and insecticide resistance, the underlying genetic mechanisms of the processes governing these areas of research are largely unknown. To address this gap, we present the *F. occidentalis* draft genome assembly and official gene set.

**Results:** We report on the first genome sequence for any member of the insect order Thysanoptera. Benchmarking Universal Single-Copy Ortholog (BUSCO) assessments of the genome assembly (size = 415.8 Mb, scaffold N50 = 948.9 Kb) revealed a relatively complete and well-annotated assembly in comparison to other insect genomes. The genome is unusually GC-rich (50%) compared to other insect genomes to date. The official gene set (OGS v1.0) contains 16,859 genes, of which ∼10% were manually verified and corrected by our consortium. We focused on manual annotation, phylogenetic and expression evidence analyses for gene sets centered on primary themes in the life histories and activities of plant-colonizing insects. Highlights include: 1) divergent clades and large expansions in genes associated with environmental sensing (chemosensory receptors) and detoxification (CYP4, CYP6 and CCE enzymes) of substances encountered in agricultural environments; 2) a comprehensive set of salivary gland-associated genes supported by enriched expression; 3) apparent absence of members of the IMD innate immune defense pathway; and 4) developmental- and sex-specific expression analyses of genes associated with progression from larvae to adulthood through neometaboly, a distinct form of maturation compared to complete metamorphosis in the Holometabola.

**Conclusions:** Analysis of the *F. occidentalis* genome offers insights into the polyphagous behavior of this insect pest to find, colonize and survive on a widely diverse array of plants. The genomic resources presented here enable a more complete analysis of insect evolution and biology, providing a missing taxon for contemporary insect genomics-based analyses. Our study also offers a genomic benchmark for molecular and evolutionary investigations of other thysanopteran species.

## BACKGROUND

Thrips are small, polyphagous and cosmopolitan insects that comprise the order Thysanoptera. Thysanoptera lies within the Paraneoptera, also commonly called the “hemipteroid assemblage” which also includes the orders Hemiptera, Psocoptera, and Phthiraptera. Among the over 7,000 reported thrips species classified into nine families with an additional five identified from fossil species [1], the plant-feeders and crop pests are the most well-characterized members of the order due to their agricultural importance. Thysanopterans present a diverse array of biological, structural and behavioral attributes, but share characteristics that are unique to insects in the order. Among these are fringed wings (Figure 1A, Adult panel) and a complex mouthcone (Figures 1B, that houses asymmetrical mouthparts composed of three stylets (Figure 1D). The paired, maxillary stylets interlock when extended during ingestion, forming a single tube, *i.e.*, food canal, that is also thought to serve as a conduit for saliva, while the single, solid-ended mandibular stylet (peg) is used to pierce substrates (its counterpart is resorbed during embryonic development [2, 3]). All the stylets are innervated, giving thrips control of stylet direction and movement in response to sensory cues [4]. Thrips also have mechano- and chemosensory structures likely governing host finding and choice. The external surface of the mouthcone supports 10 sensory pegs on each paraglossa, nine of which appear to have a dual chemosensory and mechanosensory function (sensory pegs 1-5, 7-10), and one with a mechanosensory function (sensory peg 6) (Figure 1E). In addition, internally, there are precibarial and cibarial chemosensory structures, likely important in feeding choices [4].

**Figure 1.**
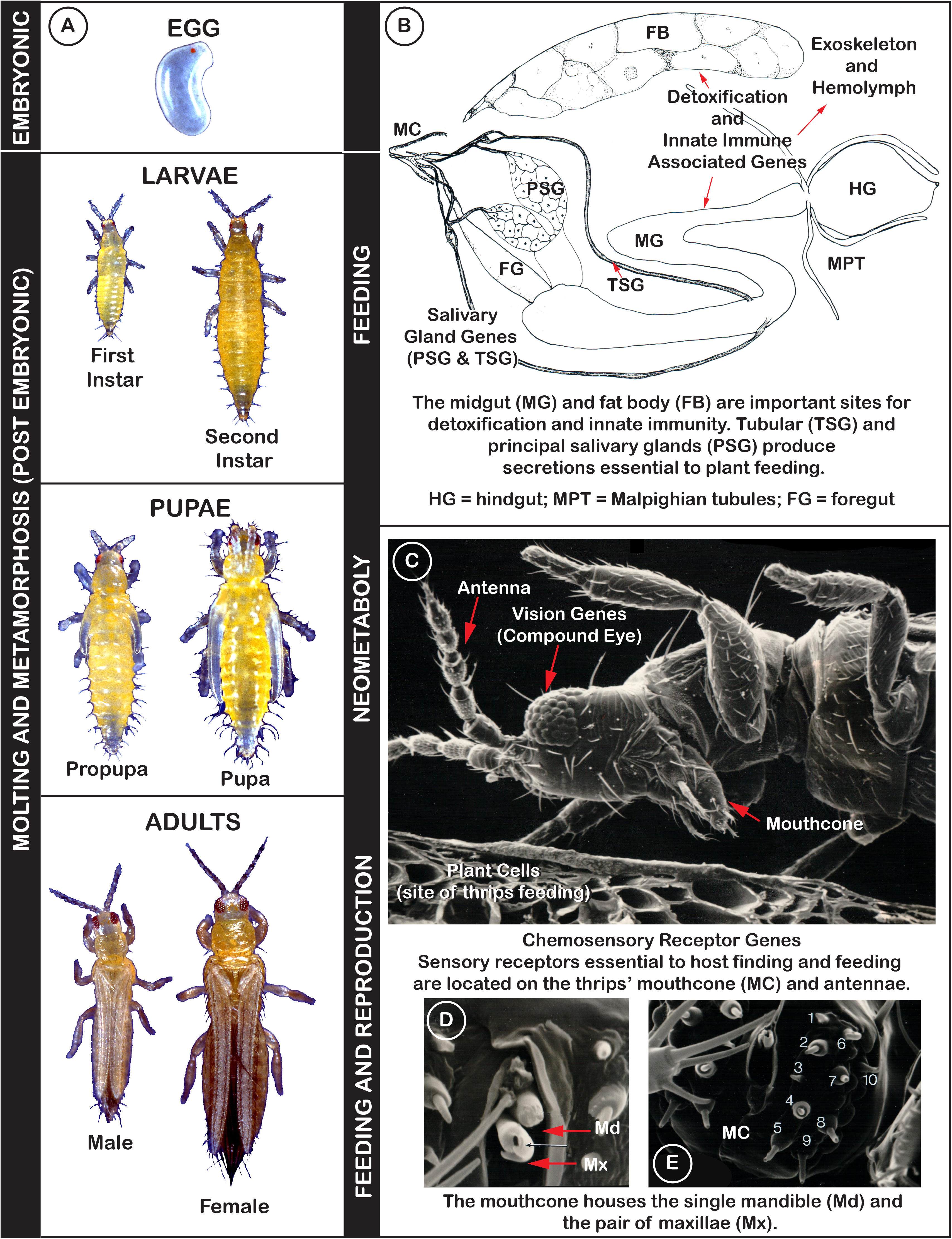
Summary of the role of curated gene sets in understanding biological processes of *Frankliniella occidentalis.* **A.** Developmental stages. Sex not determined easily prior to adulthood. Vertical bars: (left) embryonic and postembryonic (molting and metamorphosis) stages associated with gene sets are highlighted; (right) highlights that larval and adult stages feed and during their unusual metamorphosis (neometaboly), propupal and pupal stages do not; adults reproduce sexually. Modified from [141], reproduced with permission of CAB International through PLSclear. **B.** Diagram of dissected internal organs highlighting links with salivary gland, detoxification and innate immunity associated genes. Modified from [142], permission by Elsevier. **C.** Scanning electron micrograph (SEM) of adult pre-probing behavior that includes repetitive antennal waving and proleg scraping of the plant surface, followed by proleg-antennal contact, and mouthcone dabbing on the plant surface. External antennal and mouthcone sensory structures, and chemosensilla in the precibarium (inside the head above the mouthcone) (not shown), are essential to host finding, choice and feeding, highlighting the importance of sensory related gene sets. A transverse slice through plant cells most commonly fed on by thrips - epidermal, parenchymal and mesophyll. Modified from [143], permission from Springer-Verlag. **D.** SEM showing the single mandible and paired maxillae slightly extended from the mouthcone. Modified from [49], permission of Elsevier. **E.** The paraglossae at the mouthcone tip carry dual function sensory pegs (numbered on the left paraglossa) - pegs 1-5, 7-10, are mechano- and chemosensory, while sensory peg 6 is solely mechanosensory. Presence of these organs on the thrips mouthcone suggest the critical roles plant surface microtopography and chemistry play in host and feeding-site selection. Modified from [49], permission of Elsevier.

Also remarkable is their postembryonic development, referred to as “remetaboly”[5] and more recently termed “neometabolous” [6] (Figure 1A). This developmental strategy has been described as intermediate between holo- and hemimetabolous because the two immobile and non-feeding pupal stages (propupae and/or pupae) undergo significant histolysis and histogenesis, yet the emergent adult body plan largely resembles that of the larva except for the presence of wings and mature reproductive organs (Figure 1A, Adult panel).

*Frankliniella occidentalis* (suborder Terebrantia, family Thripidae, subfamily Thripinae) is a devastatingly invasive crop pest species with a global geographical distribution and an extraordinarily broad host range, capable of feeding on hundreds of diverse plant species, tissue types, fungi, and other arthropods. Additionally, this species has developed resistance to diverse insecticides with varying modes of action [7–9]. For example, on cotton, there have been 127 cases reported of field-evolved resistance to 19 insecticides belonging to six groups (modes of action) of insecticides (https://www.pesticideresistance.org, January 01 2020). The insect is haplo-diploid, *i.e.*, haploid males arise from unfertilized eggs, while diploid females develop from fertilized eggs [10]. The short reproductive cycle and high fecundity of this species contributes to its success as an invasive species.

In addition to direct damage to plants, *F. occidentalis* and other thrips vectors interact with and transmit diverse types of plant pathogens [11–14], most notoriously orthotospoviruses [15–17], to a wide array of food, fiber and ornamental crops around the globe. With regards to orthotospovirus-thrips interactions, global expression analyses of whole bodies of *F. occidentalis* [18, 19] and other thrips vectors [20, 21] indicated the occurrence of insect innate immune responses to virus infection. In addition to serving as crop disease vectors, thrips support vertically transmitted, facultative bacterial symbionts that reside in the hindgut [22, 23].

While there are numerous studies centered on thrips systematics, feeding behaviors, ecology, virus transmission biology, pest biology and insecticide resistance [24], the underlying genetic mechanisms of the complex and dynamic processes governing these areas of research are largely unknown. Here we present the *F. occidentalis* genome assembly and annotation, with phylogenetic analyses and genome-referenced transcriptome-wide expression data of gene sets centered on primary themes in the life histories and activities of plant-colonizing insects: 1) host-locating and chemical sensory perception, 2) plant-feeding and detoxification (Figure 1B**, C**), 3) innate immunity (Figure 1B), and 4) development and reproduction (Figure 1A). Analysis of the *F. occidentalis* genome highlights evolutionary divergence and host adaptations of plant-feeding thysanopterans compared to other taxa. Our findings underscore the ability of *F. occidentalis* to sense diverse food sources, to feed on and detoxify an array of natural compounds (e.g., plant secondary compounds) and agrochemicals (e.g., insecticides), and to combat and/or support persistent microbial associations. We also provide insights into thrips development and reproduction. This is the first thysanopteran genome to be sequenced, and the annotations and resources presented herein provide a platform for further analysis and better understanding of, not just *F. occidentalis*, but all members of this intriguing insect order.

## RESULTS & DISCUSSION

### Genome metrics

The assembly size of the *F. occidentalis* draft genome was determined to be 416 Mb (**Table 1**), including gaps, and is relatively close to the published genome size estimate of 345 Mb obtained by flow cytometry of propidium iodide-stained nuclei of adult males (haploid) and females (diploid) of *F. occidentalis* [25]. The assembly consists of 6,263 scaffolds (N50 = 948 Kb). One striking feature of the genome is the GC content of ∼50%, extraordinarily larger than other insects to date [26, see supplement].

**Table 1.**
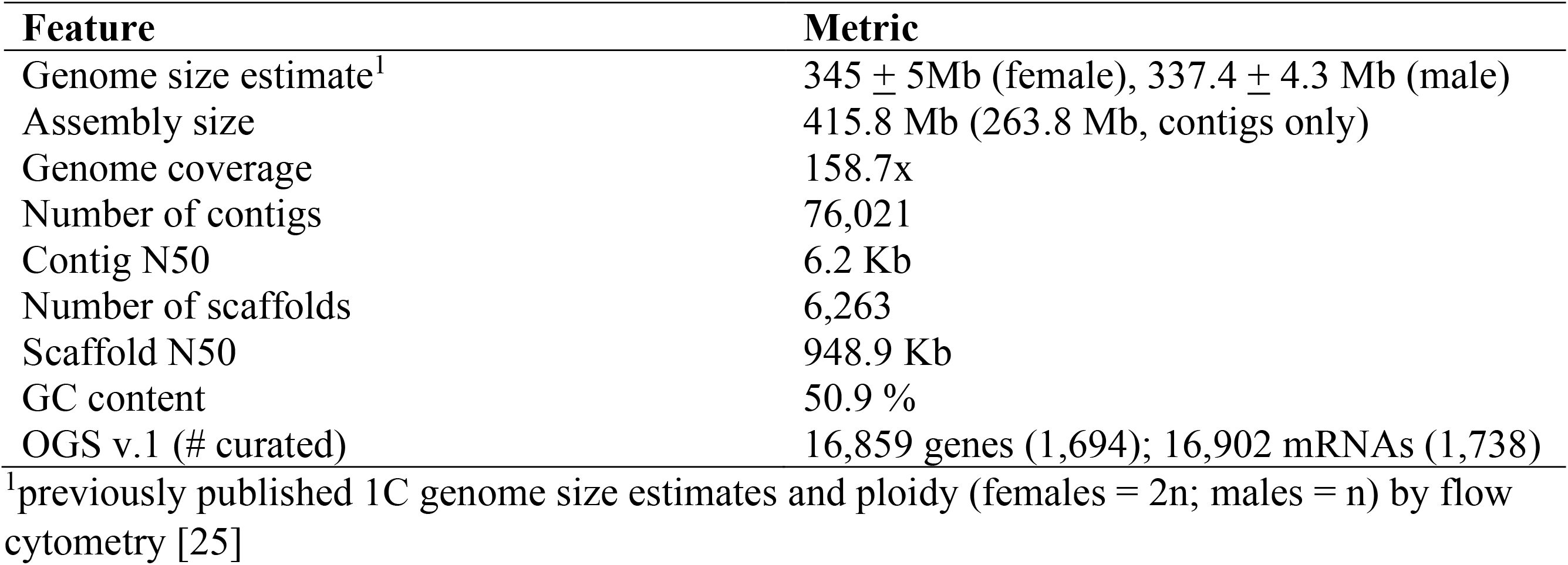
Genome metrics of *Frankliniella occidentalis*.

| Feature | Metric |
| --- | --- |
| Genome size estimate<sup>1</sup> | 345 ± 5Mb (female), 337.4 ± 4.3 Mb (male) |
| Assembly size | 415.8 Mb (263.8 Mb, contigs only) |
| Genome coverage | 158.7x |
| Number of contigs | 76,021 |
| Contig N50 | 6.2 Kb |
| Number of scaffolds | 6,263 |
| Scaffold N50 | 948.9 Kb |
| GC content | 50.9 % |
| OGS v.1 (# curated) | 16,859 genes (1,694); 16,902 mRNAs (1,738) |

<sup>1</sup>previously published 1C genome size estimates and ploidy (females = 2n; males = n) by flow cytometry [25]

### Phylogenomics with a complete and well-annotated genome assembly

Phylogenomic analysis correctly placed *F. occidentalis* (Insecta: Thysanoptera) basal to *A. pisum* and *C. lectularius* (Insecta: Hemiptera) (Figure 2A). Unexpectedly, however, the body louse *P. humanus* (Insecta: Psocodea) appears as an outgroup to all other insects, which disagrees with previous findings [27]. This discordance is most likely due to taxon sampling and would likely be resolved when more genome sequences become available from early-diverging insect lineages (e.g., Paleoptera). BUSCO assessments (see Methods) showed that the official gene set (OGS) of *F. occidentalis* is relatively complete when compared to the OGSs from other arthropods. Moreover, gene annotation (Figure 2B, left bars) in *F. occidentalis* identified 434 more complete BUSCOs, compared to the genome-based analysis (Figure 2B, right bars), resulting in reduced numbers of fragmented and missing BUSCOs equally. These findings indicate that the *F. occidentalis* gene annotation strategy successfully managed to capture even difficult-to-annotate genes.

**Figure 2.**
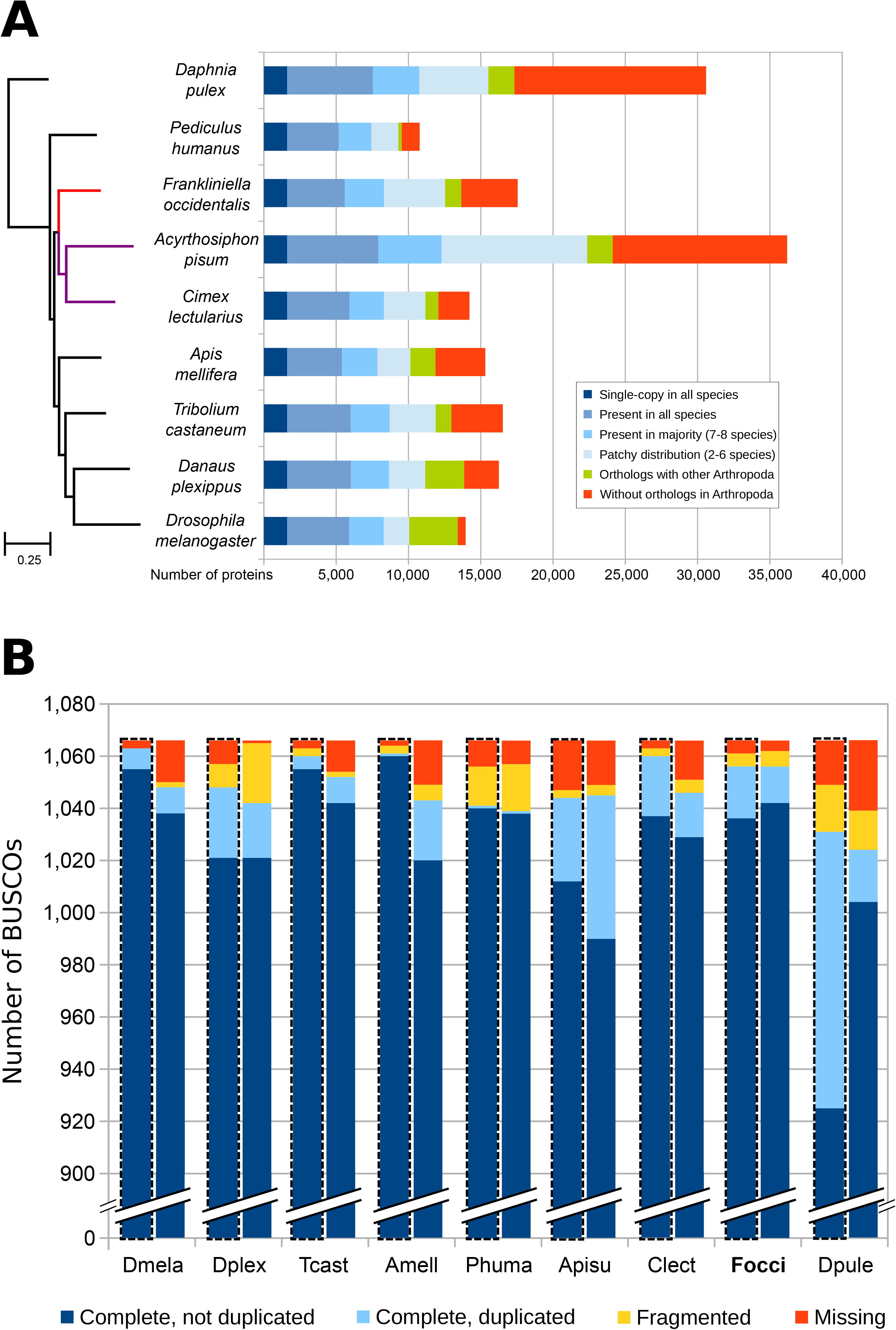
Phylogeny and orthology of *Frankliniella occidentalis* with other arthropods, with genome and gene set completeness assessments. **A.** The phylogenomic analysis was based on the aligned amino acid sequences of 1,604 single-copy orthologs and placed *F. occidentalis* (shown in red) as basal to the hemipteran species *Acyrthosiphon pisum* and *Cimex lectularius* (shown in purple). All nodes have bootstrap support of 100% and the scale bar corresponds to substitutions per site. OrthoDB orthology delineation with the protein-coding genes from the *F. occidentalis* official gene set identify genes with orthologs in all or most of the representative insects and the outgroup species, *Daphnia pulex*, as well as those with more limited distributions or with no confidently-identifiable arthropod orthologs. **B.** Assessments using the 1,066 arthropod Benchmarking Universal Single-Copy Orthologs (BUSCOs) show few missing genes from *F. occidentalis*, with better recovery than the other two hemipterans (*A. pisum* and *C. lectularius*). The *F. occidentalis* official gene set (OGS) scores better than its genome assembly, indicating that the gene annotation strategy has successfully managed to capture even difficult to annotate genes. The left bars for each species, also outlined with a dashed line show the results based on the genome, whereas the right bars show the results for the OGSs. Species names abbreviations: Dmela – *Drosophila melanogaster*, Dplex – *Danaus plexippus*, Tcast – *Tribolium castaneum*, Amell – *Apis mellifera*, Phuma – *Pediculus humanus*, Apisu – *Acyrthosiphon pisum*, Clect – *Cimex lectularius*, Focci – *Frankliniella occidentalis*, Dpule – *Daphnia pulex*.

### Assembly assessment via homeodomain transcription factor gene structure and cluster synteny

The Hox and Iro-C gene clusters that encode homeodomain transcription factors are highly conserved in bilaterian animals and in insects, respectively [28–31] and offer an additional quality appraisal for genome assembly. All single copy gene models for the expected Hox and Iro-C orthologs were successfully constructed (**Additional File 1: Section 2, Table S2.1**) and, with regards to synteny, we could reconstitute the small Iro-C cluster and partially assemble the larger Hox cluster (**Additional File 1: Section 2, Figure S2.1**), with linkage for *Hox2/3/4*, *Hox5/6/7*, and *Hox8/9/10*. All linked Hox genes occurred in the expected order and with the expected, shared transcriptional orientation, albeit with some missing coding sequence for some gene models. However, direct concatenation of the four scaffolds with Hox genes would yield a Hox cluster of 5.9 Mb in a genome assembly of 416 Mb, which is disproportionately large (3.5-fold larger relative cluster size compared to the beetle *T. castaneum* and other, *de novo* insect genomes [30, 32–34]). This finding suggests incorrect assembly of portions of these scaffolds.

Interestingly, although orthology is clear for all ten Hox genes, they are rather divergent compared to other insects. Specifically, several *F. occidentalis* Hox genes have acquired novel introns in what are generally highly conserved gene structures, and several Hox genes encode unusually large proteins compared to their orthologs, corroborating a previous, pilot analysis on unique protein-coding gene properties in this unusually GC-rich genome ([33], see supplement).

### Genome-wide analysis of transcription factors

In addition to the selected homeodomain proteins, we comprehensively identified likely transcription factors (TFs) among our entire OGS by scanning the amino acid sequences of predicted protein coding genes for putative DNA binding domains (DBDs). When possible, we also predicted the DNA binding specificity of each TF using the procedures previously described [35] (**Additional File 1: Section 1.5)**. Using this approach, we discovered 843 putative TFs in the *occidentalis* genome, which is similar to other insect genomes (e.g., 701 for *D. melanogaster*). Likewise, the number of members of each *F. occidentalis* TF family is comparable to that of other insects (Figure 3A). Of the 843 *F. occidentalis* TFs, we were able to infer motifs for 197 (23%) (**Additional file 2, Table S5**), mostly based on DNA binding specificity data from *D. melanogaster* (120 TFs), but also from species as distant as human (43 TFs) and mouse (12 TFs). Many of the largest TF families have inferred motifs for a substantial proportion of their TFs, including homeodomain/Hox (64 of 78, 82%), bHLH (30 of 36, 83%), and nuclear receptors (11 of 17, 65%). As expected, the largest gap is for C_2_H_2_ zinc fingers (only 24 of 321, ∼7%), which evolve quickly by shuffling their many zinc finger arrays, resulting in largely dissimilar DBD sequences (and hence, DNA-binding motifs) across organisms [36]. Weighted correlation network analysis (WGCNA, see **Additional file 1: Section 1.6**, supplemental methods) revealed stage-specific patterns in TF expression (Figure 3B**; Additional file 3)**. For example, Fer3, a basic Helix-Loop-Helix (bHLH) TF - previously linked to reproductive mechanisms [37] – showed increased expression in *F. occidentalis* adults compared to the larvae and propupae. In addition, multiple Hox genes exhibited increased expression in the propupae, which is consistent with their role in morphological development [38].

**Figure 3.**
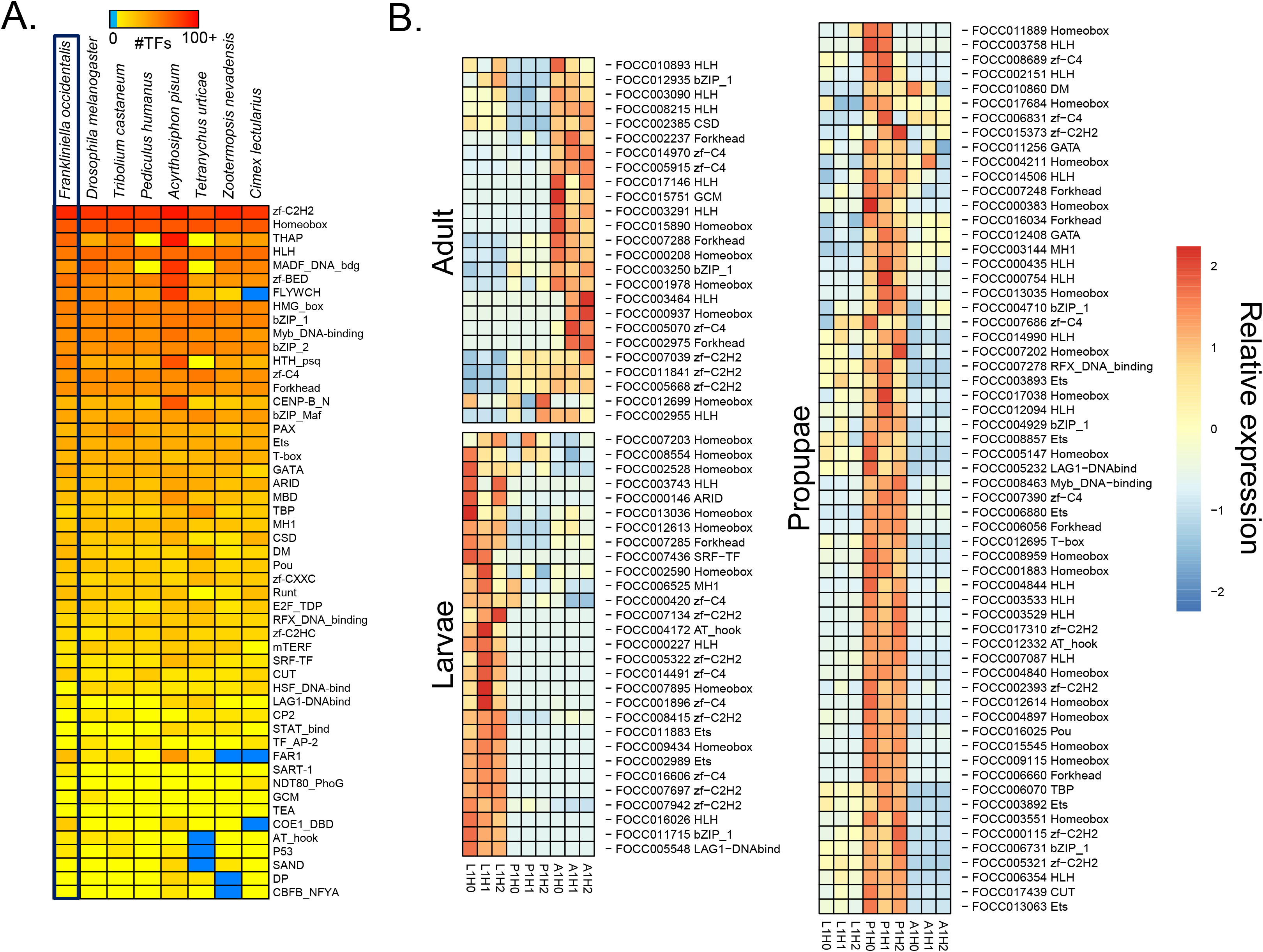
Distribution of transcription factor families across insect genomes and stage-specific expression in *Frankliniella occidentalis.* **A.** Heatmap depicting the abundance of transcription factor (TF) families across a collection of insect genomes. Each entry indicates the number of TF genes for the given family in the given genome, based on presence of DNA binding domains (DBD). Color key is depicted at the top (light blue means the TF family is completely absent) – note log (base 2) scale. Species were hierarchically clustered using average linkage clustering. *F. occidentalis* is boxed. See **Additional file 2: Table S5** for TF genes with predicted DBDs. **B.** Expression of specific TFs enriched within each developmental stage (larvae, propupae, and adult) based on data presented in **Additional file 3**.

### Genome-wide search for putative lateral gene transfers (LGTs) of bacterial origin

Once thought to be rare, LGTs from microbes into genomes of arthropods are now considered to be relatively common [39]. Although LGTs are expected to degrade due to mutation and deletion, natural selection can lead to the evolution of functional genes from LGTs, thus expanding the genetic repertoire of the recipient species [40]. We investigated candidate LGTs in *F. occidentalis* using a modification of the pipeline (**Additional File 1: Section 3.2**) originally developed by Wheeler et al [41], which has been used to identify LGTs in a number of arthropod species (e.g., [33, 42, 43]).

Three ancient LGT events were detected in the *F. occidentalis* genome (**Additional File 1: Section 3.3, Table S3.1, Figures S3.1 – S3.3**). Each of these LGTs appears to have evolved into functional genes based on maintenance of an open reading frame, transcriptional activity, and a signature of purifying selection indicated by reduced levels of non-synonymous to synonymous substitution. One LGT involves an O-methyltransferase gene likely derived from a bacterium in the Silvanigrellales or a related proteobacteria in the class Oligoflexia. O-methylases are generally characterized by the addition of a methyl moiety to small molecules and can affect many biological processes [44]. Subsequent to transfer, the gene has expanded into a three-gene family and two show transcriptional activity based on currently available RNA sequencing data. The two remaining LGTs both are glycoside hydrolases (GHs), which are a large class of proteins involved in carbohydrate metabolism [45]. One is a mannanase (GH5) which was acquired from a *Bacillus* or *Paenibacillus* based on phylogenetic analysis. This gene also subsequently underwent expansion into three paralogs in *Frankliniella*. The third ancient LGT is a levanase (GH32) that has undergone duplication subsequent to transfer. The possible origin of this gene is a bacterium in the genus *Streptomyces* or *Massilia*, although the phylogenetic reconstruction precludes a clear resolution of its source. These LGTs could be important in biological functions of *F. occidentalis*, although their actual functions remain a topic for future study. To provide support for LGT integration in the *F. occidentalis* genome, a PCR-based approach was used to confirm physical linkage between the putative LGTs and the nearest annotated thrips genes found on the same genomic scaffolds (**Additional File 1: Section 3.3, Table S3.2**). The characterization of these LTGs in different thrips life stages and their functional analyses could further help to elucidate their evolutionary role in thrips.

### Transcriptome-wide analysis of gene expression across development

We performed WGCNA using previously generated normalized read count data for non-infected first instar larvae, propupae and adult *F. occidentalis* ([19], **Additional file 3**), using methods described in **Additional file 1: Section 1.6**. The resulting network assembled co-expressed suites of genes with similar expression profiles into 35 different modules, 21 of which were significantly associated (*P* < 0.05) to one of the developmental stages (Figure 4). Eleven modules were associated with adults, nine modules for propupae, and a single module for larvae (Figure 4A). For each stage, there were ∼2,000 - 3,000 stage-associated genes. For both the propupae and larvae, there was an enrichment of Gene Ontology categories associated with metabolism and growth processes, which has been similarly seen in other systems [33]. The propupae varied from the larvae in that there was significant enrichment in processes associated with anatomical structure development, which likely underlies the structural changes associated with the larval to adult transition (Figure 4B **& C**). Adult-enriched categories were associated with protein binding, transferase activity, and nucleotidyltransferase activity (Figure 4D). This developmental analysis characterizes genes associated with progression from *F. occidentalis* larvae to adulthood.

**Figure 4.**
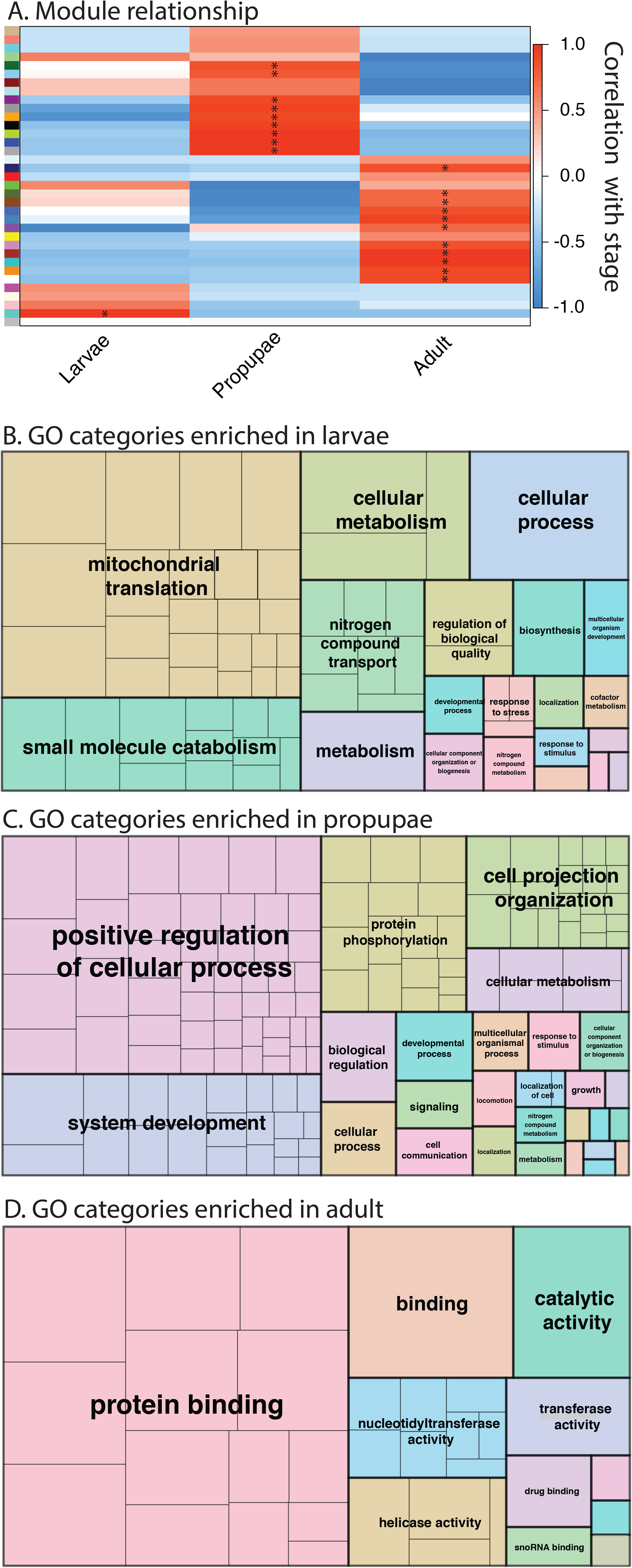
Identification of co-expressed genes and gene ontologies associated with three developmental stages of *Frankliniella occidentalis*. **A.** Weighted gene co-expression network analysis [144] was performed on a matrix of normalized read counts (FPKM values) obtained from a published *F. occidentalis* RNA-seq study involving three biological replicates of healthy first instar larvae, propupae and adults (mixed males and females) [19]. Modules were determined by the dynamic tree cutting algorithm with a minimum of 20 genes per module. Modules of co-expressed genes that exhibited the highest correlation (Pearsons coefficient, r_p_) with a developmental stage are indicated by an asterisk (*). **B.** REVIGO (<u>re</u>duce and <u>vi</u>sualize <u>g</u>ene <u>o</u>ntologies, [145]) was used to visualize specific GO terms enriched in each developmental stage; sizes of delineated blocks indicate the number of genes within each GO category. See **Additional file 3 for co-expressed genes per module**.

### Gene set annotations

Here we report on the consortium’s analysis of thrips gene sets and, in select cases, gene expression associated with four primary themes centered on interactions between phytophagous insects, plants and their environment:

i. host-locating, sensing and neural processes,
ii. plant-feeding and detoxification,
iii. innate immunity, including RNA interference and
iv. development and reproduction.

HOST-LOCATING, SENSING and NEURAL PROCESSES

### Chemosensory receptors

Chemosensation is important for most insects, including thrips, and the three major gene families of chemoreceptors, the odorant and gustatory receptors (ORs and GRs) in the insect chemoreceptor superfamily [46] and the unrelated ionotropic receptors (IRs) [47], mediate most smell and taste abilities [48]. Chemosensory organs have been described on the antennae of several thrips species, and on the mouthcone, and within the precibarium and cibarium of *F. occidentalis* [4, 49, 50]. Chemosensation plays an important role in the sequence of behaviors involved in host exploration by *F. occidentalis*. This behavioral repertoire includes surface exploration (antennal waving, presumably perceiving olfactory cues; labial dabbing, detecting surface chemistry with paraglossal sensory pegs) and internal exploration (perception of plant fluids with precibarial and cibarial sensilla) (Fig. 13 in [4]). The OR, GR and IR gene families from *F. occidentalis* were compared with those from other representative hemipteroids, specifically the human body louse *P. humanus* [51], the pea aphid *A. pisum* [52], and the bedbug *C. lectularius* [30], as well as conserved representatives from *D. melanogaster* [46, 47] and other insects (**Additional file 1: Section 4, Figures S4.1 – S4.3; Additional file 2: Table S7**). The OR family consists of a highly conserved 1:1 ortholog in most insects, the Odorant receptor co-receptor (Orco) plus a variable number of “specific” ORs that bind particular ligands. *Frankliniella occidentalis* has 84 specific OR genes, which form a divergent clade in phylogenetic analysis of the family, reflecting the divergence of thrips from other hemipteroids or Paraneoptera [27]. In addition, *F. occidentalis* has 102 GRs, including large expansions within the candidate sugar and carbon dioxide receptor subfamilies (18 and 30 genes, respectively), and a small expansion of the candidate fructose receptor subfamily to 5 genes. The remaining 49 GRs are highly divergent from those of other hemipteroids and include a recent expansion of very similar GRs perhaps involved in sensing host plant chemicals. The IR family consists of several proteins that are conserved throughout most pterygote insects including the three known co-receptors (Ir8a, 25a, and 76b) and a set of four proteins involved in perception of temperature and humidity (Ir21a, 40a, 68a, and 93a) [53]. Like other hemipteroids and most other insects, *F. occidentalis* has single orthologs of each of these seven genes. This insect species also has eight members of the Ir75 clade that is commonly expanded in insects and involved in perception of acids and amines [54]. The IR family commonly has a set of divergent proteins, some encoded by intron-containing genes, while most are intronless. *F. occidentalis* has one intron-containing gene (Ir101) with relatives in other hemipteroids, and a large divergent clade of 167 IRs including several sets of recently duplicated genes that are encoded by mostly intronless genes (the few with single introns apparently gained them idiosyncratically after expansion of an original intronless gene). This is a considerable expansion of IRs compared with the other known hemipteroids. By analogy with the divergent IRs of *D. melanogaster* that appear to function in gustation [55], these genes likely encode gustatory receptors that perhaps mediate perception of host plant chemicals and hence, host and feeding choices.

There is considerable evidence that chemosensation is important to host, feeding and oviposition choices made by *F. occidentalis*. For example, *F. occidentalis* detects pheromones and prefers specific plant volatiles [56, 57]. In choice tests with diverse tomato cultivars, adult female *F. occidentalis* preferred fully developed flowers with sepals and petals fully open to those in earlier stages of development and opening, fed preferentially on specific portions of the flower depending on tomato cultivar, and avoided specific acylsugar exudates from Type IV trichomes of tomatoes [58]. Adult females also distinguished between acylsugar molecules, different acylsugar amounts and fatty acid profiles with differentially suppressed oviposition [58–60].

### Vision genes

In contrast to their uniquely modified wings and mode of flight, thrips are equipped with the canonical pair of lateral compound eyes (Figure 1C) and three dorsal ocelli, as is typical for winged insects [61]. The success of a multitude of color and light enhanced thrips-trapping devices highlights the importance of vision for host plant recognition in this insect order [62]. For instance, female *F. occidentalis* have been found to exhibit preference for mature host plant flowers over senescent ones during dispersal within a radius of 4 meters [63]. In phototaxis assays, *F. occidentalis* displayed conspicuous peak attraction to UV (355 nm) and green (525 nm) light sources in comparison to blue (405, 470 nm), yellow (590 nm), and red 660 nm) [64]. Electroretinogram studies suggested the presence of UV, blue- and green sensitive photopigments in both sexes [64].

Compared to hemipteran genome species studied so far [30, 33, 65], the *F. occidentalis* genome contains a rich repertoire of the opsin G-protein coupled receptor subfamilies that are expressed in the photoreceptors of the insect compound eye retina. This includes singleton homologs of the UV-, and blue (B) opsin subfamilies as well as three homologs of the long wavelength (LW)-opsin subfamily (**Additional file 2: Table S8**). The latter are closely linked within a 30k region, indicative of a tandem gene duplication driven gene family expansion.

Gene tree analysis provided tentative support that the *F. occidentalis* LW opsin cluster expansion occurred independently of the previously reported LW opsin expansions in different hemipteran groups such as water striders, shield bugs, and seed bugs (**Additional file 1: Section 5, Fig S5.1**) [33, 65]. At the same time, the considerable protein sequence divergence of the three paralogs, which differ at over 140 amino acid sites in each pairwise comparison, indicated a more ancient origin of the cluster, potentially associated with elevated adaptive sequence change. Comparative searches for possible wavelength-sensitivity shifting/tuning substitutions paralleling those identified in the water strider LW opsin paralogs did not produce compelling evidence of candidate changes (not shown) [65]. Understanding the functional significance of the *F. occidentalis* LW opsin gene cluster thus requires future study.

By comparison to the differential deployment of three LW opsins in *Drosophila* [66], it seems likely that one *F. occidentalis* LW opsin paralog is specific to the ocelli, while the remaining two paralogs may be expressed in subsets of the compound eye photoreceptor cells. Overall, the presence of homologs of all three major insect retinal opsin subfamilies correlates well with the previous findings on the visual sensitivities and preferences in this species [64].

The *F. occidentalis* genome also contains singleton homologs of two opsin gene families generally expressed in extraretinal tissues and most often the central nervous system: c-opsin [67] and Rh7 opsin [68]. We failed to detect sequence conservation evidence for Arthropsins, the third extra-retinal opsin gene family discovered in arthropods [69], despite the fact that all three extraretinal opsins are present, although at variable consistency, in hemipteran species [33, 65].

### Neuropeptide signaling

Insect genomes contain large numbers of neuropeptide and protein hormones (> 40), and their receptors, many of which play significant roles in modulating sensory signals and feeding. Neuropeptides are generally encoded by small genes and occasionally evolve rapidly including the loss and duplications of these genes in different evolutionary lineages. While a number of neuropeptides are missing in several insect genomes, the genome of *F. occidentali*s still seems to have a complete set of neuropeptides (**Additional file 2: Table S10**), including all three allatostatin C-like peptides, which is a rather rare case in insects. Alternatively spliced exons encoding similar, but distinctive mature peptides are also conserved: mutually exclusive exons of ion transport peptide A and B [70] and orcokinin A and B [71]. Exceptions occurred in natalisin and ACP signaling pathways [72, 73], for which both neuropeptides and the receptors are missing in this species. A surprising finding in this genome is a second corazonin gene that encodes a slightly different version of corazonin [74]. The gene clearly arose from a duplication of the corazonin gene and it has accumulated a substantial number of changes in the sequence. The duplicated gene encoding the corazonin precursor does not contain disruptive mutations in the open reading frame and its signal peptide is expected to be functional. The transcripts were also confirmed by RNAseq evidence provided with the genome resources. Together, this evidence collectively suggests that it is unlikely to be a pseudogene.

Similar to the case of conserved gene number, the motif sequences of the putative mature peptides are also well conserved in *F. occidentalis* (**Additional file 5**). An exception in this case is found in MIP (myoinhibitory peptide or allatostatin B) [75]. While its peptide motif is highly conserved not only in insects but also in mollusks and annelids, in *F. occidentalis*, the C-terminal tryptophan is replaced by a phenylalanine and 23 of the 25 MIP paracopies of the precursor have this unusual sequence. The predicted presence of a disulfide bridge in the N-terminal of the longest pyrokinin is another unusual and noteworthy structural feature.

Receptors associated with the set of *F. occidentalis* neuropeptides and hormones were also catalogued (**Additional file 2: Table S9**). In *Drosophila* only a few neuropeptide genes have more than one receptor. However, in the *F. occidentalis* genome, there are duplicate G protein-coupled receptors (GPCR) for SIFamide, PTH, the CRF-like diuretic hormone 44 and CNMamide. These are ancestral and are generally conserved in other insect species as single copies. What is unusual in the *F. occidentalis* genome is that GPCRs for trissin, vasopressin, leucokinin and RYamide as well as the orphan GPCR moody all have local duplications, which are likely generated by recent events in this species. These recently duplicated GPCRs include receptors for neuropeptides implicated in water homeostasis: vasopressin, leucokinin, and RYamide [76–78], implying that osmoregulatory processes are tightly regulated in *F. occidentalis*.

## PLANT FEEDING

### Salivary gland-associated genes

Among piercing-sucking insects, salivation is a key component of their ability to feed on plants. Saliva may form a protective sheath for the stylets, permit intra and intercellular probing and serve as elicitors that interact with plant defense pathways in ways that may benefit the insect (reviewed in [79, 80]). While little is known about the function of *F. occidentalis* saliva, it is expected to play a key role in this insect’s capacity to feed on an extraordinarily large number of plant species and its ability to transmit viruses. To generate a comprehensive set of putative *F. occidentalis* salivary gland-associated genes, we performed comparative analyses of RNA-seq datasets (**Additional file 1: Section 1.7**) derived from salivary glands (SGs: principal salivary glands and tubular salivary glands, Figure 1B) [81] relative to the entire body. The analysis revealed 141 and 137 transcript sequences in SGs of *F. occidentalis* females and males, respectively, and 127 in a combined sex analysis that were significantly greater in abundance compared to whole-body expression. There were 123 sequences that overlapped between the three salivary gland sets (Figure 5A; **Additional file 2: Table S11**). These 123 sequences represent 83-88% of all reads mapped in salivary gland libraries and only a maximum of 14.7% of the reads from the whole-body samples (Figure 5B). Many of the SG-enriched sequences (∼68%) have less than one million reads mapped per salivary gland dataset and very few (11%) are highly expressed with over 2.5 million reads mapped per sequence (Figure 5C). The highly expressed sequences code for many unknown proteins (unknown proteins have matches in other species but have yet to be functionally characterized) or are *F. occidentalis*-specific proteins (Figure 5D). Many of the sequences with provisional functions likely have specific roles related to the breakdown of plant materials or response to the host during feeding (Figure 5E). Validation of enriched expression of select genes by real-time reverse transcriptase (RT)-PCR (**Additional file 1: Section 6, Fig S6.1**) yielded a Pearson correlation coefficient of 0.845, indicating that these results have accurately identified putative salivary gland-associated sequences. Apart from the SG-enrichment analysis, other gene models encoding digestive enzymes were pulled out of the genome; we therefore consider these likely gut-associated genes (**Additional file 2: Table S12**). The SG gene set will be very valuable in future investigations aimed at understanding the diverse diet of *F. occidentalis*, and the role of saliva as a vehicle for virus inoculation and possibly a means by which the insect manages plant defenses by its many hosts.

**Figure 5.**
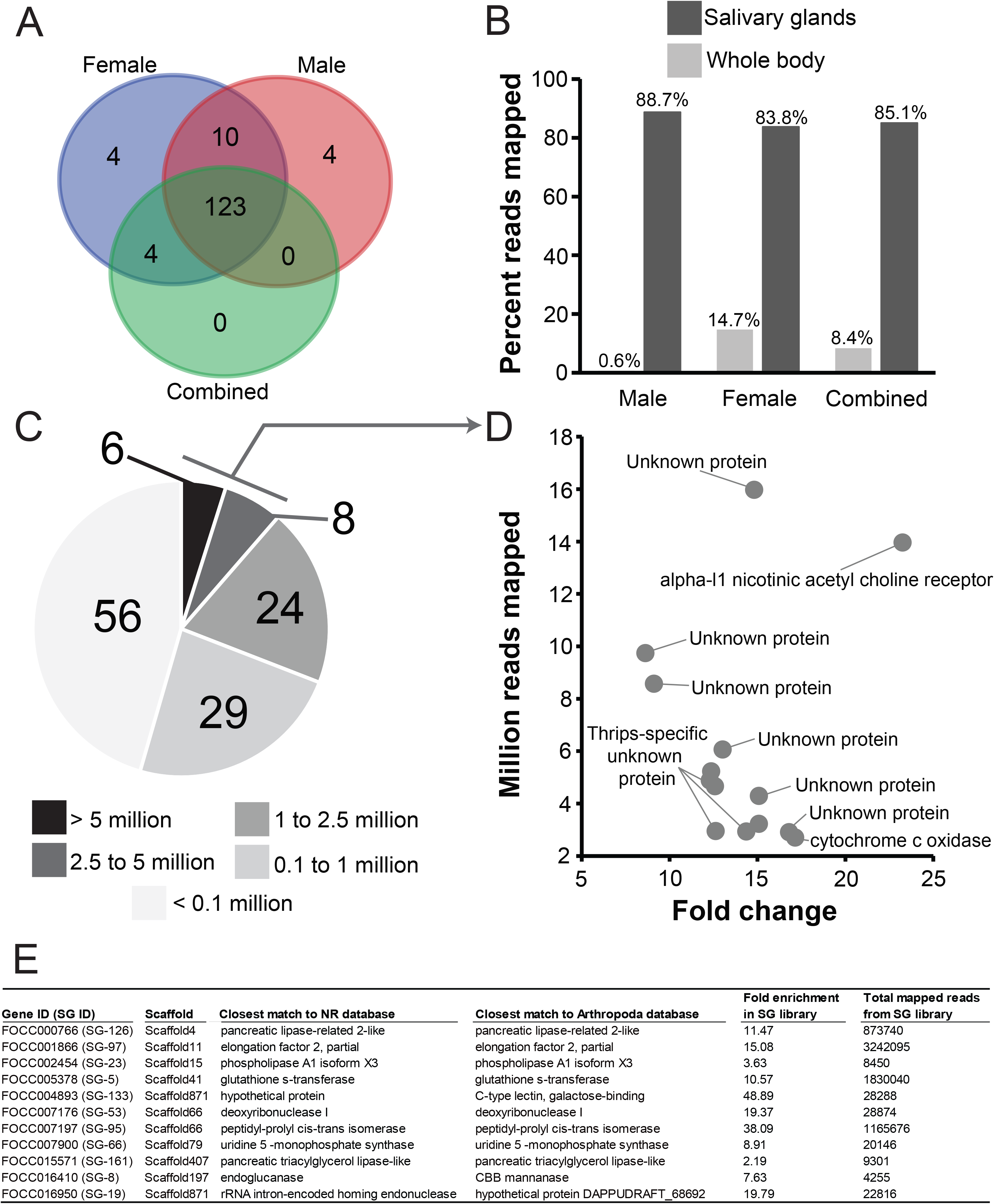
Genes/contigs with enriched expression in the salivary glands of *Frankliniella occidentalis.* RNA-seq reads generated from male and female principal and tubular salivary glands collectively [81] and whole bodies (this study) were used for the enrichment analysis. **A**. Venn diagram depicting the overlap in transcript sequences enriched in the salivary glands of males, females, and combined sexes compared respectively to whole bodies. **B**. Percent reads from salivary glands and whole-body RNA-seq datasets mapped to the putative 123 salivary gland-associated sequences. **C.** Number of reads from the female salivary gland RNA-seq dataset mapping to each of the 123 salivary gland-associated sequences. **D.** Reads mapped by fold change for 14 sequences with the highest number of mapped reads denoted in panel C. **E**. Specific sequences with functional assignment that are likely related to *F. occidentalis* feeding. See **Additional file 2: Table S11** for the salivary gland gene set.

### Detoxification

#### Cytochrome P450s

Cytochrome P450s (CYPs) are a large, ancient superfamily of enzymes identified in all domains of life and are involved in the metabolism of multiple substrates with prominent roles in hormone synthesis and breakdown, development, and detoxification [82, 83]. In agricultural systems, *F. occidentalis* has shown a propensity for developing resistance to insecticides commonly utilized to manage this species, and P450s have been specifically implicated in the detoxification of insecticides by *F. occidentalis* [84, 85]. Within the *F. occidentalis* genome, we identified and classified (CYP nomenclature) a relatively large number of P450s (130 models compiled into 89 complete genes, 23 partial sequences) of 24 different CYP families (**Additional file 2: Table S13**), where, like in other insects, the CYP4 and CYP6 gene families are overrepresented – these families include members frequently associated with the breakdown of toxic plant products and insecticides [84]. Phylogenetic analysis revealed expansions in clan CYP 3 (CYP6 gene family) and clan CYP4 within *F. occidentalis* (**Additional file 1: Section 7.1, Fig S7.1**). The majority of annotated *F. occidentalis* P450s showed relatively low amino acid identity to other insect P450s (**Additional file 1: Section 7.1, Table S7.2**), a common aspect of insect genomes [86]. Given the already described importance of P450s in insecticide resistance [84, 85], the importance of insecticides in the management of thrips species [84], and the multitude of plant defense compounds encountered during the their phytophagous lifestyle [83], knowledge of the diversity of P450s present within the *F. occidentalis* genome is likely essential for optimizing management of this important agricultural pest. The annotation of these P450 genes will enable future functional studies in *F. occidentalis* related to the detoxification of insecticidal and plant defense compounds.

#### ATP-binding cassette (ABC) transporters and carboxyl/choline esterase (CCE) genes

Forty-five and 50 putative ABC and CCE genes were annotated in the *F. occidentalis* genome, respectively (**Additional file 2: Table S15 and S16**). The number of *F. occidentalis* ABC genes is on the lower side among those reported for other insect species (**Additional file 1: Section 7.2, Table S7.3**, [87, 88]), including *Bemisia tabaci* of the Hemiptera, the sister-group of the Thysanoptera [89]. Nevertheless, we did identify a lineage-specific expansion of ABCH genes within the *F. occidentalis* genome (**Additional file 1: Section 7.2, Figure S7.6**). Lineage-specific arthropod ABCH genes were previously shown to respond to environmental changes or xenobiotic exposure [90–92] and hence these ABCH genes might have a similar function in *F. occidentalis*. In contrast to ABC genes, the number of *F. occidentalis* CCE genes is among the highest of those identified in any insect species (**Additional file 1: Section 7.2, Table S7.4)** [88]). This high number of CCEs is due to a lineage-specific expansion within the dietary/detoxification class of CCEs (**Additional file 1: Section 7.2, Figure S7.7**). Future work should confirm whether these 28 *F. occidentalis* specific CCEs are actually detoxification CCEs and whether the polyphagous nature and/or rapid development of insecticide resistance in *F. occidentalis* [93] might be related to this CCE expansion.

## INNATE IMMUNITY

### Canonical signaling pathways

Insects rely on innate immunity to respond to and limit infections by myriad microbes, viruses and parasites encountered in their environments. Here we report the annotation of genes associated with pathogen recognition, signal transduction, and execution of defense in *F. occidentalis*, and support these findings with a comparative analysis of immune-related transcripts in two other thrips vector species, *F. fusca* and *Thrips palmi*.

In total, 96 innate immune genes were curated from the genome (**Additional file 2: Table S17)**. Toll and JAK-STAT pathway members were well represented, and all but two members of the IMD pathway were located. Based on the number of different pathogen recognition receptors, *F. occidentalis* has a well-developed surveillance system – 14 PGRPs and 8 GNBPs – greatly exceeding the number reported for other insects [33, 94]. The broad plant host range and biogeography of this thysanopteran species may have expanded the repertoire of receptors capable of recognizing diverse pathogen and/or microbial-associated molecular patterns in these diverse biomes. Expansion of these surveillance systems could be due to the close contact of pupal stages with the soil environment during their development. Likewise, the melanization pathway encoded by the *F. occidentalis* genome is notably extensive compared to other insect genomes [33, 94]. The melanization pathway is triggered by the binding of pathogen recognition molecules to PGRPs and is the first line of defense in insects. Prophenoloxidase (PPO) and serine proteases are the primary players of the melanization pathway. These primary players are well represented in the *F. occidentalis* genome, with six PPOs and serine proteases, compared to the closest plant feeding hemipteran relatives that have only two PPOs each (*Acyrthosiphon pisum* and *Oncopeltus fasicatus)*.

The most striking finding is the absence of the signal transducing molecule IMD, as well as FADD, another death domain-containing protein that acts downstream of IMD to activate transcription of AMPs [95]. Absence of IMD has also been reported for the hemipteran species *Rhodnius prolixus*, *A. pisum*, *Bemisia tabaci* and *Diaphorina citri* [94, 96–99]. In *Oncopeltus*, IMD could not be identified by homology searches, but was identified by cloning the gene using degenerate primers [33]. IMD was also reported missing from the bedbug *C. lectularius* [30], but was later found using the *Plautia stali* IMD sequence as a query [100]. These findings in hemipterans illustrate that IMD sequences can be highly divergent and conclusions about their absence should be drawn with care.

It has been suggested for *A. pisum* that its phloem-limited diet and dependence on gram-negative endosymbionts accounts for a generally reduced immune repertoire and the absence of IMD [94, 98, 101]. This does not seem valid for the polyphagous, mesophyll feeding thrips. In contrast to *A. pisum*, almost all other components of the IMD signaling pathway are present in *Frankliniella*, including two Relish molecules. In conclusion, the absence of IMD in *F. occidentalis* does not seem to suggest a reduced immune repertoire, but rather a different way of mediating the response to Gram negative bacteria, possibly by Toll signaling components. In *Drosophila*, DAP-type peptidoglycans of Gram-negative bacteria moderately induce Toll signaling [102, 103]. In *Tenebrio molitor*, PGRP-SA recognizes both Gram positive and Gram negative bacteria [104]. Extensive cross reactivity of the Toll and IMD signaling pathway is the currently emerging picture from studies on other insects [100, 105, 106] and might have set the stage for multiple independent IMD losses in evolution [100].

### Comparative analysis of innate immune transcripts in three thrips vector species

Using a custom database of innate immune protein sequences (**Additional file 2: section 8.3.1**), we identified a diverse repertoire of transcripts implicating the activity of canonical humoral and cellular innate immunity from a previously assembled transcriptome of *F. occidentalis* adults [19] (**Additional file 2: Table S18**) and similarly for two other known vectors of orthotospoviruses: *F. fusca* ([20]) and *Thrips palmi* adults [21]. Comparative analysis revealed the occurrence of shared and species-specific innate immune-associated transcripts (Figure 6**; Additional file 9**). Both IMD and FADD transcripts were apparently absent (E-value cut-off = 10^−5^) in all three species which agrees with the annotation of the *F. occidentalis* genome. Relaxing the cut-off (10^−3^) resulted in weak and ambiguous matches to IMD or IMD-like sequences (**Additional file 1: Table S8.2**) of other hemipterans. Absence of transcripts encoding these two canonical genes suggests either cross-reactivity with the other immune signaling pathways or evolution of an atypical signaling pathway which is yet to be deciphered. All components of the JAK/STAT pathway were identified in all three thrips species. There appeared to be an over-representation of sequence matches to cytokine receptors in *F. occidentalis* and *F. fusca*, and while some of these may be involved in innate immunity, they likely play roles in other biological processes as well. Antioxidants, autophagy-related proteins and inhibitors of apoptosis were well represented among the three transcriptomes. Differences in the number of immune-related transcripts identified between the species should be taken with caution - different biological and experimental factors, including thrips rearing conditions, sampling strategies, and sequencing/assembly parameters may contribute to this variation.

**Figure 6.**
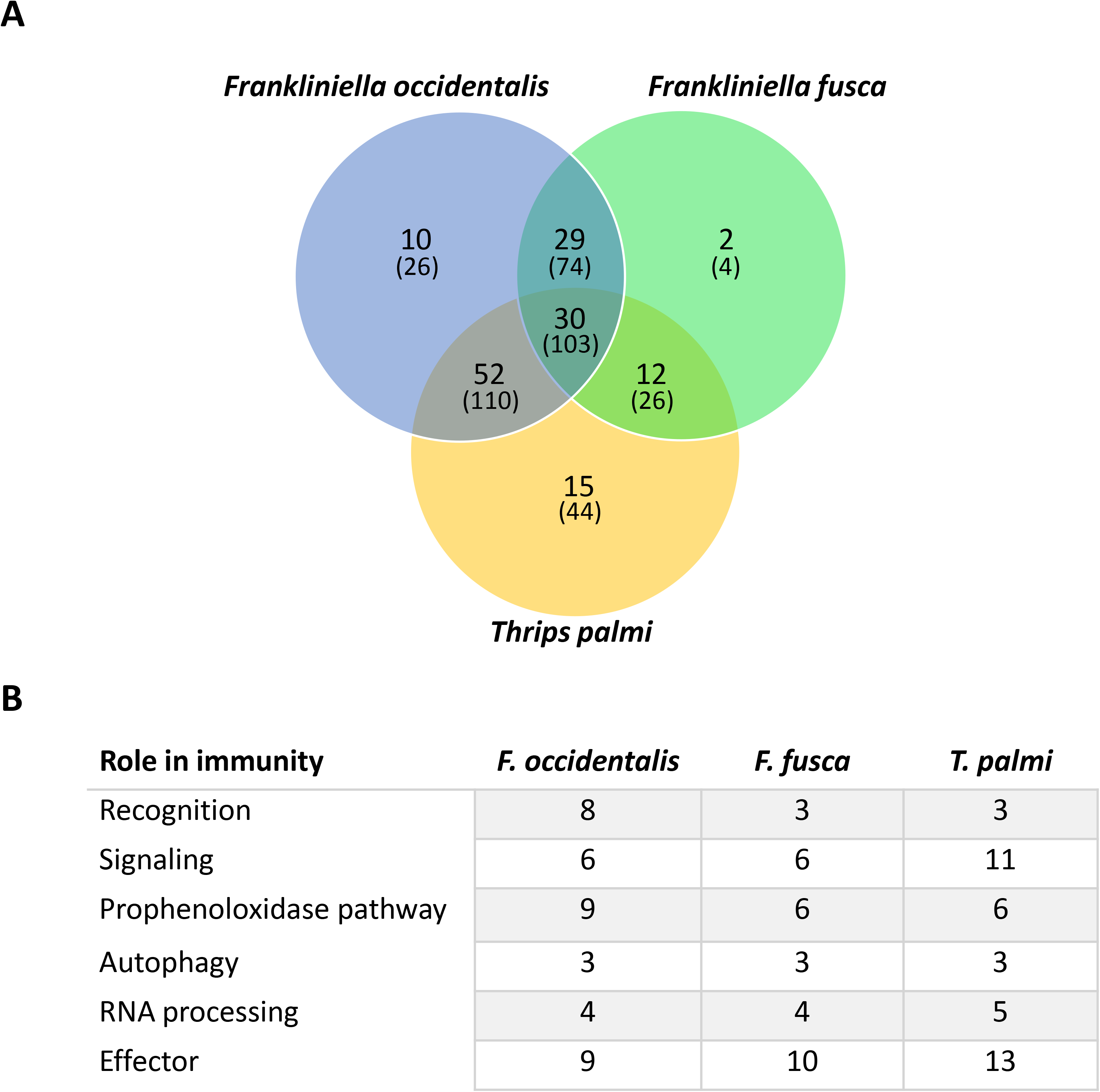
Unique and shared innate immunity-associated transcripts in three thrips vector species of orthotospoviruses. Whole-body, assembled transcriptomes obtained from published orthotospovirus-thrips RNAseq studies [19–21] were mined for putative innate-immune transcripts using an innate immune-associated protein database derived from ImmunoDb (http://cegg.unige.ch/Insecta/immunodb). (A) Venn diagram depicting overlap in orthologous clusters (bold) and transcripts (in parentheses) of innate immune-associated protein sequences in *Frankliniella occidentalis* (tomato spotted with virus), *F. fusca* (tomato spotted wilt virus) and *Thrips plami* (capsicum chlorosis virus) using Orthovenn.v2. (B) Number of transcripts classified into innate immune categories (roles) and shared across all three vector species. Sequences may fall into more than one category. See **Additional file 2: Table S17 and S18**, respectively, for innate immune genes and transcript sets; **Additional file 9** for Orthovenn outputs.

### Small RNA-mediated gene silencing pathways and auxiliary genes

The RNAi-related gene set examined in this study constitutes a group of genes that are all members of a diverse range of gene (super)families that are evolutionarily unrelated but are linked based on their roles in RNAi [107, 108]. This group includes core machinery genes for the siRNA and miRNA pathways, including several *dicer* and *argonaute* genes, *drosha*, *pasha*, *aubergine*, *loquacious* as well as several genes involved in antiviral immune response and genes encoding auxiliary proteins (*stau*, *maelstrom*, *fmr-1*, *clp-1*, *translin*, *gawky*, *prmt5*, *hen-1*, *p68 RNA helicase*, *ars2*, *egghead*). Also, the gene encoding the transmembrane channel protein *sid1* implicated in cellular uptake of dsRNA was identified in the *F. occidentalis* genome. *F. occidentalis* transmits seven described orthotospovirus species (Order *Bunyavirales*, Family *Tospoviridae*) of economic importance, including tomato spotted wilt virus (TSWV) (reviewed in [17]). These plant-pathogenic viruses are transmitted in a persistent-propagative manner by the thrips vector, i.e., retained through molts, replicating in infected tissues, and inoculative over the lifespan of the adult. In the case of *F. occidentalis*, however, virus infection does not appear to have a negative effect on thrips development or fitness [109, 110]. As RNAi is a potent innate antiviral defense in arthropods, the activities of the core cellular machinery in thrips vectors may be associated with orthotospovirus persistence.

Of the 24 RNAi-related genes queried against the genome, 23 were identified (**Additional file 2: Table S19**). One gene, *r2d2*, which encodes a co-factor of Dicer-2 and is therefore an element of the siRNA pathway, was not located. This could be due to the absence of *r2d2* in this species, extensive divergence precluding its identification using orthologs, or location in a region of the genome that was not covered by our sequencing. Using pre-existing transcriptome sequence databases for *F. occidentalis*, dsRNA-binding proteins were located, however, they did not match the r2d2 sequences used as queries. For example, in a published *F. occidentalis* EST library of first-instar larvae [111], one sequence (GT302686) was annotated as ‘tar RNA binding’ containing a predicted conserved domain indicative of double-stranded RNA binding (DSRM), matching a staufen-like homolog, while one sequence (contig01752) obtained from a 454 de novo assembled transcriptome representing mixed stages of *F. occidentalis* matched RISC-loading complex subunit tar RNA binding proteins. In the *F. occidentalis* genome sequence, one gene coding for an RNA-binding protein similar to *r2d2* was located, but it appeared to encode the very similar protein Loquacious (Loqs) and had a significant match (99.8%) to contig01752. Given their similarity, a phylogenetic tree was constructed with the four isoforms identified to be coded by this gene, clearly confirming that it is indeed the *loqs* homolog (**Additional file 1: Section 8.5, Fig S8.8**).

*r2d2* has been reported to be missing in other annotated winged and wingless arthropod genomes and transcriptomes. For example, *r2d2* is missing from the hemipteran *D. citri* [112]. A recent study on the phylogenetic origin and diversification of RNAi genes reported that the gene could not be found in the transcriptomes of any of the wingless insects investigated, and did not occur in some older orders of winged insects [113]. Furthermore, *r2d2* also seems to be missing in non-insect arthropods. In the common shrimp *Crangon crangon* for example, no *r2d2* could be found in the transcriptome [107] and data-mining of other Crustacea such as *Daphnia pulex* [113] and *Artemia franciscana* [114] and in the chelicerates *T. urticae* and *Ixodes scapularis* [113, 115] also suggested that *r2d2* is missing in those respective genomes. It has been suggested that in these arthropods and insects, the role of *r2d2* and its interaction with Dicer-2 in the siRNA pathway may have been replaced by Loqs, which serves a similar function, interacting with Dicer-1 in the miRNA pathway. In fact, the involvement of Loqs in the siRNA pathway has been reported in the fruitfly *D. melanogaster*, where four dsRNA-binding proteins interacting with Dicer enzymes have been found, one encoded by the *r2d2* gene and three by the *loqs* gene through alternative splicing. In these fruit flies, Fukunaga and Zamore [116] have shown that one of the Loqs isoforms interacts with Dicer-2 and is involved in siRNA processing. A dual role in both pathways has also been described for Loqs in *Aedes aegypti* [117]. Whether or not this is also the case in non-dipteran insects, such as *F. occidentalis*, or other arthropods is yet to be determined.

### Antioxidants

Twenty-nine putative proteins in seven families related to antioxidant capacity were identified within the *F. occidentalis* genome (**Additional file 2: Table S20**). Consequently, the suite of antioxidant proteins identified in *F. occidentalis* was largely as expected and further investigation into the antioxidant system of *F. occidentalis* will further elucidate the players. The twenty-nine antioxidant response proteins showed high homology to related proteins in other published genomes including *A. pisum*, *A. mellifera*, *Bombyx mori*, *C. lectularius*, *D. melanogaster*, *P. humanus*, and *T. castaneum*. In most comparisons, homologs in *T. castaneum* showed the highest degree of similarity followed by *A. pisum* and *P. humanus*.

## DEVELOPMENT & REPRODUCTION

### Embryonic development

#### Wnt pathway

The Wnt pathway is a signal transduction pathway with fundamental regulatory roles in embryonic development in all metazoans. The emergence of several gene families of both Wnt ligands and Frizzled receptors allowed the evolution of complex combinatorial interactions with multiple layers of regulation [118]. Wnt signaling affects cell migration and segment polarity as well as segment patterning and addition in most arthropods [119]. Surveying and comparing the gene repertoire of conserved gene families within and between taxonomic groups is the first step towards understanding their function during development and evolution.

Here we curated gene models for the main components of the Wnt signaling pathway in the *F. occidentalis* genome (**Additional file 1: Section 9.1, Table S9.1**) and confirmed their orthology by phylogenetic analysis. We found 9 Wnt ligand subfamilies, three Frizzled transmembrane receptor subfamilies, the co-receptor *arrow*, and the downstream components *armadillo/beta-catenin, dishevelled, axin*, and *shaggy/ GSK-3*. All of these genes, with the exception of the Frizzled family (three *fz-2* paralogs), were present in single copy in the assembly. Three Wnt genes, *wingless, Wnt6* and *Wnt10*, were linked on the same scaffold, reflecting the ancient arrangement of Wnt genes in Metazoa. One of the Wnt ligands, *Wnt16*, has so far only been reported in the pea aphid *A. pisum* [120], the Russian wheat aphid *Diuraphis noxia* [121] and *fasciatus* [33], suggesting that the hemipteroid assemblage (clade Acercaria) has retained a Wnt ligand that was subsequently lost within the Holometabola.

### Post-embryonic development

We curated *F. occidentalis* genes associated with molting and metamorphosis (**Additional file 2: Table S21**) and report here on enriched expression during development to support our curations. For the complete set of significantly co-expressed postembryonic genes and their normalized read counts (FPKM) for first instar larvae (L1), propupae (P1) and adults (mixed sex) of *F. occidentalis* refer to **Additional file 2: Table S22**.

The juvenile hormone (JH) and ecdysone pathway genes were determined to be generally conserved in *F. occidentalis*. The MEKRE93 pathway [122] - consisting of the transcription factors Met, Kr-h1 and E93 - was fully annotated, along with the pupal-specifying gene *Broad*. The antimetamorphic gene Kr-h1 in *F. occidentalis* was previously identified [123], and the published sequence is consistent with the genome annotation. In our dataset, Met expression was associated with L1 as expected for hemi- and holometabolous insects. Kr-h1 showed low expression and was not associated with any one stage, which is concordant with previous findings [123]. E93, the specifier for adult development that is thus expected to increase in expression during late nymph or propupae stages [122], was indeed upregulated and enriched in the P1 stage. *Broad* showed low expression in L1 as expected, however expression was exceptionally low in P1, a finding that may be explained by P1 age at time of sampling [123]. While expression across the three *F. occidentalis* stages appeared to be low overall, *Broad* expression was associated with the young adult. Three copies of xanthine dehydrogenase (*rosy*), a protein essential in mediating JH action in the developing abdominal epidermis of *D. melanogaster* [124] were identified. Of the three copies associated with *F. occidentalis*, xanthine dehydrogenase-2 was supported by expression data and was relatively more abundant in the adult stage. Finally, both Taiman, the steroid receptor coactivator (*AaFISC* in [125], *TcSRC* in [126]), and FtzF1, which serves as a physical bridge between the JH receptor machinery and ecdysone, were identified with their transcripts upregulated in the P1 stage.

Ecdysone associated genes were identified with varying levels of expression during development. Thirteen ecdysone cascade genes and coactivators and eight p450 (CYP) ‘Halloween’ genes that catalyze the biosynthesis or inactivation of 20-hydroxyecdysone (20E) were identified (**Additional file 1: Section 7.1, Table S7.2)** [127, 128]. As expected, the evolutionarily conserved developmental CYP genes showed some of the highest amino acid conservation observed among the collection of P450s from the *F. occidentalis* genome versus P450s in other insect genomes.

JH and ecdysone titers are tightly regulated via the action of biosynthetic and metabolic genes. In contrast to *D. melanogaster*, a mevalonate kinase ortholog was not identified in *F. occidentalis*. With regards to JH degradation – which is performed by JH epoxide hydrolase (JHEH) and JH esterase (JHE) - a single obvious JHEH gene was identified in contrast to three orthologs in *D. melanogaster* and showed marked upregulation and enrichment in the L1 stage. The *F. occidentalis* genome, however, carries an additional four epoxide hydrolase orthologs, any of which may have JHEH activity – all four showed expression in L1s. Notably, several of the *F. occidentalis* carboxylesterase annotations meet a “diagnostic” criterion (GQS<u>A</u>G motif; <u>A</u> replaced by S in *F. occidentalis*) of functional JHE proteins [129] (**Additional file 1: Section 9.2, Figure S9.1)**; however, based on the developmental expression profiles, only one of the putative JHE genes in the *F. occidentalis* genome is predicted as the true JHE (**Additional file 2: Table S22**). Three *apterous* (Ap) orthologs were identified, apparently the result of tandem duplications. The *apterous* mutation in *Drosophila* results in misregulated JH production, leading to female sterility. In light of this reproductive fitness cost, expression of Ap during *F. occidentalis* larval and adult life - during which JH is necessary for development and reproduction - is expected. In addition to its role in promoting JH synthesis, Ap is a homeodomain protein that establishes dorsoventral boundary in the developing wing disc and *Ap* misexpression has a range of developmental consequences on wing morphology [130]. It is therefore intriguing to ponder a role for *apterous* duplications in the context of thrips’ unique wing morphology.

As expected, most of the annotated post-embryonic genes belonged to the bHLH superfamily (**Additional file 1: Section 9.2**), transcription factors that regulate various developmental processes across all domains of life. In *F. occidentalis*, 45 bHLH-PAS/myc family members were conclusively annotated (**Additional file 2: Table S21**). This gene superfamily showed putative duplication events - three *Enhancer of split* (*E(spl)-bHLH*) paralogs, two *hairy* orthologs, two presumed paralogs of the *dimmed*, and similarly*, knot* (syn. *Collier*) - and their expression profiles may indicate stage-specific sub/neofunctionalization (**Additional file 1: Section 9.2**).

### Cuticular proteins

Sequence motifs that are characteristic of several families of cuticle proteins [131] were used to search the genome of *F. occidentalis* for putative cuticle proteins. In total, 101 genes were identified, analyzed with CutProtFam-Pred, a cuticular protein family prediction tool described in Ioannidou et al. (2014), and assigned to one of seven families (CPR, CPAP1, CPAP3, CPF, CPCFC, CPLCP, and TWDL) (**Additional file 2: Table S23**). As with most insects, the CPR RR-1 (soft cuticle), RR-2 (hard cuticle), and unclassifiable types, constituted the largest group of cuticle protein genes in the *F. occidentalis* genome (**Additional file 1: Section 10, Table S10.1**). The number of genes in the protein families CPR, CPAP1, CPAP3, CPCFC, and CPF were similar to the number in other insects [131]. However, the 10 genes in the TWDL family was greater than that found in most insect orders and is reminiscent of the expansion of this family observed in Diptera (**Additional file 1: Section 10, Figure S10.1**). Many of the genes (∼40%) were arranged in clusters of 3 to 5 genes (**Additional file 1, Table S10.2**) that were primarily type-specific. However, the sizes of gene clusters were smaller than those observed in other insects, which are typically 3 to ∼20 genes in size. Additionally, a larger portion (50-70%) of cuticle proteins is typically found in clusters in other insects - clustering of these genes could allow for the coordinated regulation of cuticle proteins and thereby facilitate the development of insecticide resistance.

### Nuclear receptors

Nuclear receptors (NRs) play important roles in development, reproduction and cell differentiation in eukaryotes. In insects, many are part of the ecdysteroid signaling cascade. Most of these NRs contain a highly conserved DNA-binding domain (DBD) and a more moderately conserved ligand-binding domain (LBD). These molecules have a very specific working mechanism, being simultaneously a transcription factor and a receptor for small amphiphilic molecules such as steroids, thyroids, vitamins and fatty acids. In this way, they allow a direct response to certain hormone stimuli by controlling gene expression without requiring a complex cellular signaling cascade. The proteins in this superfamily are categorized into six major subfamilies (NR1-NR6) based on phylogenetic relationships, with an additional subfamily (NR0) containing non-canonical NRs usually lacking either a DBD or LBD [132, 133]. All expected nuclear receptor genes (21 in total) commonly found in insect species were identified in the *F. occidentalis* genome (**Additional file 2: Table S24**). All known insect members of the NR1-NR6 subfamilies were identified including the NR2E6 and NR1J1 genes that were previously reported to be missing in the hemipteran *A pisum*, the nearest relative to thrips and the first hemimetabolous insect to have its genome sequenced [98, 134]. In the NR0 group, three receptors were identified (Egon, Knirps, and Knirps-like), as was the case with other members of the hemipteroid assemblage (*A. pisum* and *P. humanis*) and *Drosophila*. It is possible that the three NR0 genes found in the *F. occidentalis* genome are orthologous to those in Hemiptera, however, phylogenies of the arthropod NR0 genes are notoriously difficult to resolve due to the lack of semi-conserved LBD and the high divergence between these different NRs.

## REPRODUCTION

Curation and WGCNA of post-embryonic developmental genes revealed members of JH, ecdysone, and insulin signaling pathways in *F. occidentalis* that are known to be required in other insects for vitellogenesis, functioning uniquely across taxonomic lines. For instance, ecdysone and JH have opposing functions in reproductive tissue maturation in *Tribolium* and *Drosophila*. In *F. occidentalis*, there were nine adult-stage, co-expressed genes implicated in oocyte development and reproductive biology (**Additional File 2: Table S22**) - *hydroxymethylglutaryl-CoA synthase 1* and *farnesoic acid O-methyltransferase are* involved in JH biosynthesis, while the others are involved in nutritional (e.g. insulin) and steroid signaling. One oddity that begs further research is the finding that *methoprene tolerant* (Met) was not upregulated in the sampled adult stage of *F. occidentalis*, since this JH receptor has roles in oocyte maturation and vitellogenesis, as well as accessory gland development and function, and in courtship behaviors. Of two lipase-3 like annotations, one was enriched in adults, while the other was enriched in larvae. Larval expression is likely related to nutritional signaling and feeding, whereas the adult transcript is likely required for reproduction.

### Comparison of reproductive gene expression in male and female thrips

To identify male and female enriched genes, we performed a comparative RNA-seq analysis between females, males, and larvae (**Additional file 1: Section 1.8; Additional file 8: normalized fold change and annotations**). Following the *F. occidentalis*-specific analysis, specific sets were compared to previous de novo assemblies for other thysanopteran species (Figure 7). Based on these comparative analyses, 644 female-enriched, 343 male-enriched, and 181 larvae-enriched genes were identified in common among the thrips (Figure 7A**-C**). These overlapping sets for females included many factors expected to be increased in this egg generating stage, including vitellogenin and vitellogenin receptors along with other factors associated with oocyte development (Figure 7D, **Additional file 8: Table S1**). Males had enriched levels for many factors associated with sperm generation and seminal fluid production (Figure 7E**; Additional file 8: Table S2**). Many of these male-associated genes are hypothetical and not characterized, which is common for seminal proteins [135]. The larvae sets were enriched for aspects associated with growth and development, such as cuticle proteins (**Additional file 8: Table S3**). Overall, these gene expression profiles provide putative male- and female-associated gene sets for *F. occidentalis*.

**Figure 7:**
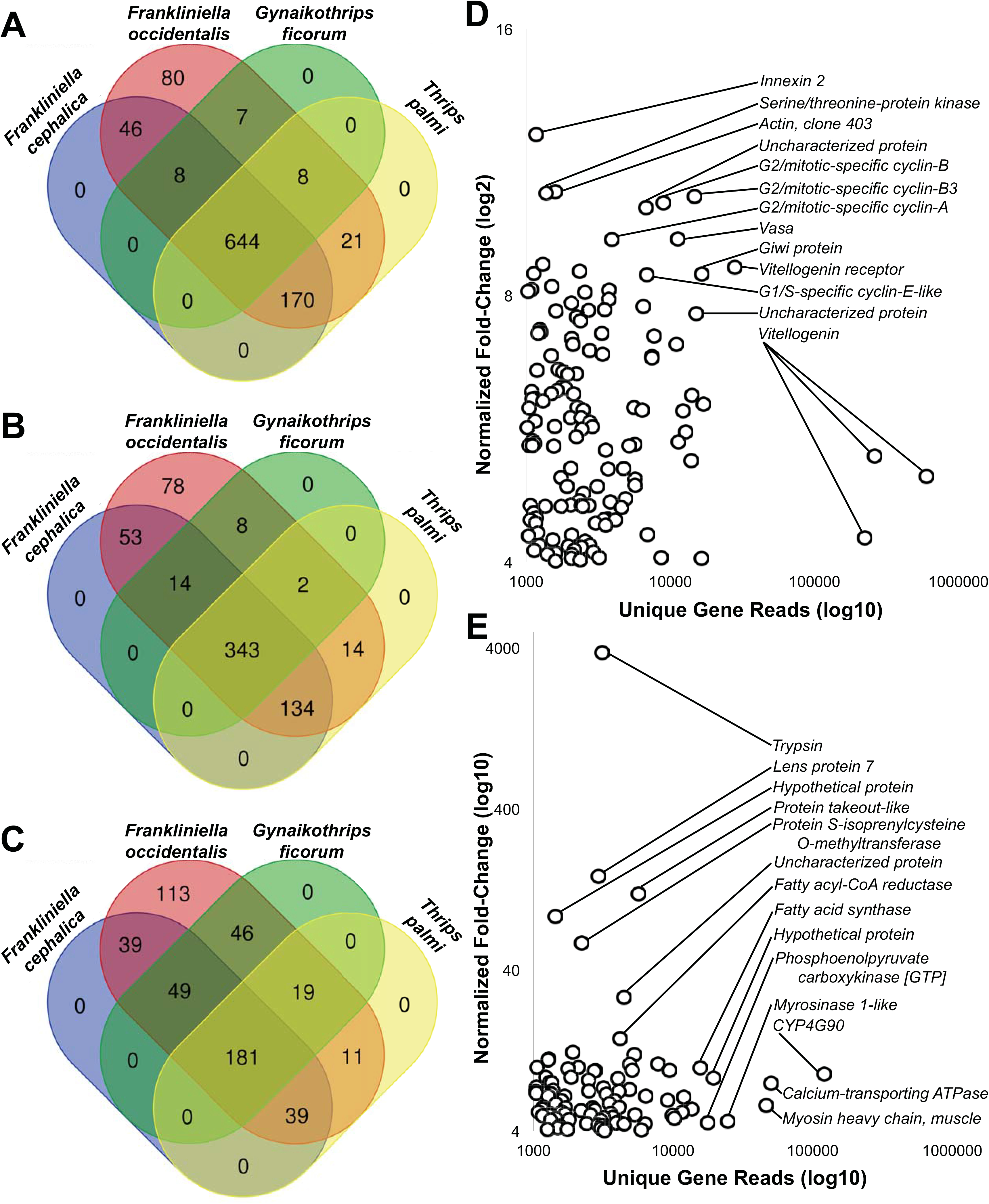
Conserved sex-specific gene expression in thrips. Genome-assembled transcripts derived from RNA-seq reads for females, males and pre-adults (larval and pupal combined) of *Frankliniella occidentalis* (this study, PRJNA203209) were compared to transcripts generated *de novo* from publicly-available RNA-seq data sets for *Frankliniella cephalica* (PRJNA219559), *Gynaikothrips ficorum* (PRJNA219563) and *Thrips palmi* (PRJNA219609). Venn diagrams depict the number of transcript sequences associated with **A**) females, **B**) males, and **C**) pre-adult stages of thrips. Highly enriched sequences (> 1000 unique reads and > 4-fold difference) conserved in female and **E**) male thrips. Labeled gene descriptions in D and E represent highly enriched gene ontology (GO) terms. See **Additional file 8** for sex-specific genes sets and associated fold change values.

## CONCLUDING REMARKS

The *F. occidentalis* genome resources fill a missing taxon in phylogenomic-scale studies of thysanopterans and hemipterans. On the ecological level, the genome will forge new frontiers for thrips genetics and epigenetics studies, genome-wide analyses of biotic and abiotic stress encountered by this pest in diverse environments, and identification of genes and gene products associated with plant-microbe/virus-thrips vector interactions. The availability of this genome may also provide a means to address the challenge of determining whether *F. occidentalis* is a single widespread, interbreeding gene pool or a series of weakly interbreeding (even non-interbreeding) gene pools (i.e., sibling species) (Mound, personal communication, [24]). From a pest management perspective, the genome provides tools that may accelerate genome-editing for development of innovative new-generation insecticides and population suppression of targeted thrips. The first look at gene annotations presented here points to unique features of this herbivorous pest and plant virus vector, such as the repertoire of salivary gland proteins and detoxification enzymes, opening new avenues of both basic and translational research for *F. occidentalis* and other thysanopteran species.

## METHODS

### Thrips rearing and genomic DNA isolation

Genomic DNA was isolated from a 10^th^ generation sibling-sibling line of *F. occidentalis* (Pergande) inbred for genome homozygosity from a lab colony (**Additional file 1: Section 1.1**). The original colony progenitor was collected from O’hau, Hawaii [136]. The gDNA samples were sent to the Baylor College of Medicine Human Genome Sequencing Center (BCM-HGSC) for sequencing.

### Genome sequencing and assembly

*Frankliniella occidentalis* was one of 28 arthropod species sequenced and assembled (**Additional file 2, 1.2.1**) as a part of a pilot project for the i5K arthropod genomes project at the Baylor College of Medicine Human Genome Sequencing Center [137]. The assembly has been deposited in NCBI (https://www.ncbi.nlm.nih.gov/assembly/GCA_000697945.4, **Additional File 2: Table S1**).

## Automated Gene Annotation Using a Maker 2.0 Pipeline Tuned for Arthropods

The 28 i5K pilot genome assemblies including *F. occidentalis* were subjected to automatic gene annotation using a Maker 2.0 annotation pipeline tuned specifically for arthropods (**Additional file 1: Section 1.2.2**). The automated gene sets are available from the BCM-HGSC website (https://www.hgsc.bcm.edu/arthropods/western-flower-thrips-genome-project) as well as the National Agricultural Library (https://i5k.nal.usda.gov/Frankliniella_occidentalis) where a web-browser of the genome, annotations, and supporting annotation data is accessible.

### RNA evidence used to support manual genome curation

Both newly obtained and published *F. occidentalis* transcriptome resources were used to aid in annotation efforts (**Additional File 1: Section 1.4**). All of the transcriptome resources were mapped to the genome for visualization by Web Apollo/JBrowser at the National Agriculture Library (NAL) i5k workspace sequence repository, data-share and curation site for all i5k projects (https://i5k.nal.usda.gov/).

### Assessment of gene set completeness and genome assembly quality

The OrthoDB v8 resource [138] was queried to find shared orthologs among *F. occidentalis* and another eight arthropods genomes; *Daphnia pulex*, *Pediculus humanus*, *Acyrthosiphon pisum*, *Cimex lectularius*, *Apis mellifera*, *Tribolium castaneum*, *Danaus plexippus* and *Drosophila melanogaster.* Custom Perl scripts were used to compute the number of genes in each category shown in Figure 2A. For the phylogenomic analysis, only single-copy orthologs were used to build a concatenated protein sequence alignment from which to estimate the phylogenetic tree using RAxML [139] (**Additional File 1: Section 1.3.2**). For evaluating the completeness of the *F. occidentalis* official gene set and genome assembly, we used Benchmarking Universal Single-Copy Orthologs (BUSCO) [140] of the Arthropoda gene set, which consists of 1,066 single-copy genes that are present in at least 90% of selected representative arthropods. BUSCO assessments were run with the default parameters (**Additional file 1: Section 1.3.1**).

### Community curation of the thrips genome

Seventeen groups were recruited from the i5k pilot project and thrips research community to manually curate MAKER-predicted models (Focc_v0.5.3) of gene sets of interest to thrips consortium members. The consortium used Web Apollo/JBrowser tools, online training, and written guidelines made available by the National Agriculture Library (NAL) i5k workspace, along with the RNA evidence described above, to locate and correct 1,694 genes, 1,738 mRNAs, and 13 pseudogenes. At the completion of the community curation period, the manually curated models were exported in gff3 format and quality-checked for formatting and curation errors, and then integrated with the MAKER-predicted gene models (**Additional File 2: Table S2**) to generate a non-redundant official gene set (OGS v1.0) (**Additional File 2: Table S3**). After removal of bacterial sequence contamination, the OGS v1.0 includes 16,859 genes, 16902 mRNAs, and 13 pseudogenes and is currently housed at NAL, AgData Commons repository (http://doi.org/10.17616/R3G051).

## AVAILABILITY OF DATA AND MATERIALS

The datasets supporting the conclusions of this article are publicly available at the NCBI, Bioproject PRJNA203209, and in the United State Department of Agriculture’s National Agricultural Library (NAL), AgData Commons repository under the specific identifiers: http://doi.org/10.15482/USDA.ADC/1503960, *Frankliniella occidentalis* genome assembly v1.0 [146]; http://doi.org/10.15482/USDA.ADC/1503959, *Frankliniella occidentalis* genome annotations v0.5.3 [147]; and http://doi.org/10.15482/USDA.ADC/1504029, *Frankliniella occidentalis* Official Gene Set OGSv1.0 [148]. Also, scaffolds, gene models and genome browser are available at the NAL i5k workspace (https://i5k.nal.usda.gov/).

## ACKNOWLEDGMENTS

We thank the staff at the Baylor College of Medicine Human Genome Sequencing Center for their contributions. We thank David Nelson (University of Tennessee, USA) for officially naming the CYP sequences generated from this study. We also thank undergraduate students Nathan Nager, Michael Cosmos, and William Crusey for assisting HMR with analysis of the chemoreceptor gene families. We thank Anita Shrestha (University of Georgia, USA, currently Vertex Pharmaceuticals, USA) and Rajagopalbab (Babu) Srinivasan (University of Georgia, USA) for sharing the *de novo*-assembled transcriptome sequences of *Frankliniella fusca* from a previously published study for use in this study. We also thank Eileen Rendahl for graphic design of Figure 1.

## FUNDING

Funding for genome sequencing, assembly and automated annotation was provided by NHGRI grant U54 HG003273 to RAG. We also acknowledge NSF grants DEB1257053 and IOS1456233 to JHW, University of Cincinnati Faculty Development Research grant and NSF grant DEB 1654417 to JBB, and Swiss NSF awards 31003A-125350 and 31003A-143936 to EMZ. RMW was supported by Swiss National Science Foundation grant PP00P3_170664. SS, WD and TVL were supported by Research Foundation Flanders (FWO) grant (G053815N). JEO, SB-M and DJS were supported by AFRI NIFA Coordinated Agricultural Project grant (2012-68004-20166) and SPR was supported by USDA NIFA grant (2018-67013-28495). The funders had no role in study design, data collection and analysis, decision to publish, or preparation of the manuscript.

## AUTHOR INFORMATION

Affiliations (see title page)

### Contributions

SR and DR conceived the project. DR coordinated the project. DR and AEW provided the biological material for sequencing, KH and BAS produced the sib-sib inbred line and extracted DNA for sequencing, and DJS extracted RNA for sequencing. SR, DMM, and RAG served key roles in program management of the sequencing project. SD, SLL, HC, HVD, HD, YH, and DMM constructed the libraries and sequenced the genome. SR, JQ, SCM, DSTH, and KCW assembled the genome and performed the automated gene predictions. MFP and CC implemented the Web Apollo-based manual curation. JHW and SC performed the bacterial scaffold detection and LGT analysis and CNTT and DW performed PCR validation of LGTs. CGCJ, MvdZ, DJS, SMKW-G, NEB, ECJ, AJR, AR, AAB, OC, IMVJ, SB-M, JBB, MTW, JEO, SS, CNTT, WD, MF, JWJ, KAP, YP, HMR, JAV, GS, TVL, DEU, AEW, and RGD participated in manually curating genes and/or supervised curators. MFP performed quality control of the manual curation and provided the OGS. PI, RMW, FAS and EMZ performed the analyses of gene set completeness and genome assembly quality. KAP and IMVJ analyzed homeodomain transcription factor gene structure and cluster synteny to assess assembly quality. MTW performed the genome-wide analysis of transcription factors. SB and JBB performed WGCNA of development with contributions from DR and DJS. JBB performed comparative analyses of RNAseq datasets to generate the SG gene set and SB-M performed real-time qRT-PCR to validate the SG analysis. SPR data-mined transcriptomes for innate immune gene transcripts for comparative analysis. CJH and JBB performed comparative analyses of sex-specific genes. DR prepared the manuscript and DR, SR, CGCJ, ECJ, AJR, AR, AAB, OC, IMVJ, SB-M, JBB, MTW, JEO, SS, CNTT, WD, MF, JWJ, KAP, YP, HMR, JAV, GS, TVL, DEU, MvdZ, AEW, JHW, SC, PI contributed to supplemental notes, including phylogenetic trees, large supplementary tables and writing of the manuscript. DR and SPR prepared the supplementary notes and DR assembled the large supplementary tables. DEU, MTW, JBB, PI, and SPR prepared primary figures.

## ETHICS DECLARATION

The authors declare they have no competing interests ADDITIONAL FILES

## ADDITIONAL FILES

**Additional file 1**: Supplementary notes, figures and small tables (PDF)

**Additional file 2**: Workbook of large supplementary tables (XLSX)

**Additional file 3**: Weighted correlation network analysis (WGCNA) modules of co-expressed gene sets of developmental stages (XLSX)

**Additional file 4**: Chemosensory receptor sequences (DOCX)

**Additional file 5**: Neuropeptide and protein hormone peptide sequences with highlighted features (DOCX)

**Additional file 6**: Complete and partial cytochrome p450 (CYP) sequences (DOCX)

**Additional file 7**: Complete ATP-binding cassette transporter (ABC) and carboxylesterase sequences (DOCX)

**Additional file 8**: Supporting tables of RNAseq expression outputs for conserved sex-specific genes in thrips species (XLSX)

**Additional file 9**: Supporting tables for Orthovenn comparison of innate immunity-associated transcripts in thrips vector species (XLSX)

## Notes

https://www.ncbi.nlm.nih.gov/assembly/GCA_000697945.4

https://i5k.nal.usda.gov/

http://doi.org/10.15482/USDA.ADC/1503960

http://doi.org/10.15482/USDA.ADC/1503959

http://doi.org/10.15482/USDA.ADC/1504029

