## Additional file 1 for "Genome-enabled insights into the biology of thrips as crop pests"

***Frankliniella occidentalis* (FOCC) genome paper**

**Additional file 1: Supplementary methods, annotation notes, small tables and phylogenetic trees contributed by the FOCC genome annotation consortium.**

The contents of this supplement provide supporting details that were too extensive to include in the primary paper. Analysis of gene sets and expression data are provided by section (see table of contents), and supplemental tables and figures are numbered sequentially by section number and referred to as such in the primary paper. The type, amount and style of information presented in each section may differ due to the priorities set by the different members of the FOCC genome annotation consortium.

Note: The gene model identifiers (FOCC) in this supplement are the original MAKER gene model identifiers PRIOR to consortium curation of the selected gene sets and reassignment of OGS v1.0 identifiers. The OGS v1.0 identifiers that correspond to the MAKER identifiers in this supplement can be found in Additional file 2: Table S2.

### 1. Primary Methods

#### 1.1. Thrips inbred line and gDNA isolation methods

*Contributed by Dorith Rotenberg, Kaylee Hervey and Brandi (Schneweis) McElhoes*

Thirty-one males and females were singly paired in small 1-oz clear plastic cups with lids fitted with thrips-proof-screen, and each cup contained a small cut segment of surface-disinfested green bean pod serving as the rearing and oviposition substrate. To reduce the likelihood of parthenogenic reproduction in subsequent generations – as unfertilized female *Frankliniella occidentalis* produce only male progeny – early second instar larvae (L2) developing from each mating pair were removed with a fine, water-moistened paintbrush and transferred as single pairs to individual cups with a fresh cut bean to develop to adulthood. Pairs that did not develop into male-female pairs were discarded from their lineage. By the $10^{th}$ generation, four inbred lines were moved to larger colony-size, 12-oz deli cups to initiate amplification of the line, of which one survived to establish a healthy, reproductive colony. Pools of adult females from this colony served as the biological material for genomic DNA isolation. Genomic DNA (gDNA) was isolated from eight, 10-mg subsamples of $CO_2$-anesthetized females (hundreds of individuals per subsample) that were flash-frozen in liquid nitrogen, pulverized by hand with Kontes pestles (DWK Life Sciences, Thermo Fisher Scientific, Inc) then processed using the OmniPrep Genomic DNA Isolation Kit (G-Biosciences, Geno Technology, Inc., Saint Louis, MO, USA) following manufacturer's instructions, with the added step of 15 min room-temperature incubation of the dissolved pellet with 1 µl of LongLife RNAse (G-Biosciences) to remove residual RNA. The concentration of gDNA determined by Nanodrop spectrophotometer (ThermoFisher Scientific Inc.) and they ranged from 48 – 89 µg of DNA. Gel electrophoresis (2% agarose gel) resolved single bands of gDNA of greater than 24 kb in size.

#### 1.2. Genome sequencing, assembly and automated gene annotation

*Contributed by Stephen Richards, Daniel S.T. Hughes, Shwetha C. Murali, Jiaxin Qu, Shannon Dugan, Sandra L. Lee, Hsu Chao, Huyen Dinh, Yi Han, HarshaVardhan Doddapaneni, Kim C. Worley, Donna M. Muzny, Richard A. Gibbs*

##### 1.2.1. Genome sequencing and assembly

The *F. occidentalis* is one of 28 arthropod species sequenced as a part of a pilot project for the i5K arthropod genomes project at the Baylor College of Medicine Human Genome Sequencing Center. As with the other i5k species, an enhanced Illumina-ALLPATHS-LG sequencing and assembly strategy for *F. occidentalis* enabled multiple species to be approached in parallel at reduced costs. For most species, including *F. occidentalis*, for the pilot we sequenced four libraries of nominal insert sizes 180bp, 500bp, 3kb and 8kb. The amount of sequence generated from each of these libraries is noted in **Additional file 2: Table S1** with NCBI SRA accessions. All the libraries were made from DNA isolated from the product of 10 generations of sibling-sibling breeding for genome homozygosity of a lab colony raised in Dorith Rotenberg's Lab (see section 1.1). The original colony progenitor was isolated in the Kamilo Iki valley on the island of O'ahu, Hawaii.

To prepare the 180bp and 500bp libraries, we used a gel-cut paired end library protocol. Briefly, 1 µg of the DNA was sheared using a Covaris S-2 system (Covaris, Inc. Woburn, MA) using the 180-bp or 500-bp program. Sheared DNA fragments were purified with Agencourt AMPure XP beads, end-repaired, dA-tailed, and ligated to Illumina universal adapters. After adapter ligation, DNA fragments were further size selected by agarose gel and PCR amplified for 6 to 8 cycles using Illumina P1 and Index primer pair and Phusion® High-Fidelity PCR Master Mix (New England Biolabs). The final library was purified using Agencourt AMPure XP beads and quality assessed by Agilent Bioanalyzer 2100 (DNA 7500 kit) determining library quantity and fragment size distribution before sequencing.

The long mate pair libraries with 3kb or 8kb insert sizes were constructed according to the manufacturer's protocol (Mate Pair Library v2 Sample Preparation Guide art # 15001464 Rev. A PILOT RELEASE). Briefly, 5 µg (for 2 and 3-kb gap size library) or 10 µg (8-10 kb gap size library) of genomic DNA was sheared to desired size fragments by Hydroshear (Digilab, Marlborough, MA), then end repaired and biotinylated. Fragment sizes between 3-3.7 kb (3kb) or 8-10 kb (8kb) were purified from 1% low melting agarose gel and then circularized by blunt-end ligation. These size selected circular DNA fragments were then sheared to 400-bp (Covaris S-2), purified using Dynabeads M-280 Streptavidin Magnetic Beads, end-repaired, dA-tailed, and ligated to Illumina PE sequencing adapters. DNA fragments with adapter molecules on both ends were amplified for 12 to 15 cycles with Illumina P1 and Index primers. Amplified DNA fragments were purified with Agencourt AMPure XP beads. Quantification and size distribution of the final library was determined before sequencing as described above.

Sequencing was performed on Illumina HiSeq2000s (Casava Version 1.8.3_V3) generating 100bp paired end reads. Reads were assembled using ALLPATHS-LG (v35218) (Gnerre et al., 2011) on a large memory computer with 1Tbyte of RAM and further scaffolded and gap-filled using in-house tools Atlas-Link (v.1.0) and Atlas gap-fill (v.2.2) (https://www.hgsc.bcm.edu/software/). The assembly has been deposited in the NCBI: Genbank assembly accession GCA_000697945.4.

##### 1.2.2. Automated Gene Annotation Using a Maker 2.0 Pipeline Tuned for Arthropods

The 28 i5K pilot genome assemblies including *F. occidentalis* were subjected to automatic gene annotation using a Maker 2.0 annotation pipeline tuned specifically for arthropods. The pipeline is designed to be systematic providing a single consistent procedure for the species in the pilot study, scalable to handle 100's of genome assemblies, evidence guided using both protein and RNAseq evidence to guide gen models, and targeted to utilize extant information on arthropod gene sets. The core of the pipeline was a Maker 2 (Cantarel et al., 2008) instance, modified slightly to enable efficient running on our computational resources. The genome assembly was first subjected to de-novo repeat prediction and CEGMA analysis to generate gene models for initial training of the ab-initio gene predictors. Three rounds of training of the Augustus (Stanke et al., 2008) and SNAP (Korf, 2004) gene predictors within Maker were used to bootstrap to a high quality training set. Input protein data included 1 million peptides from a non-redundant reduction (90% identity) of Uniprot Ecdysozoa (1.25 million peptides) supplemented with proteomes from eighteen additional species (*Strigamia maritime*, *Tetranychus urticae*, *Caenorhabditis elegans*, *Loa loa*, *Trichoplax adhaerens*, *Amphimedon queenslandica*, *Strongylocentrotus purpuratus*, *Nematostella vectensis*, *Branchiostoma floridae*, *Ciona intestinalis*, *Ciona savignyi*, *Homo sapiens*, *Mus musculus*, *Capitella teleta*, *Helobdella robusta*, *Crassostrea gigas*, *Lottia gigantean*, *Schistosoma mansoni*) leading to a final non-redundant peptide evidence set of 1.03 million peptides. RNAseq transcription data derived from adult males, females and mixed sex juveniles (**Additional file 2: Table S1**) was used judiciously to identify exon-intron boundaries but with a heuristic script to identify and split erroneously joined gene models. We used CEGMA models for QC purposes: for, of 1,977 CEGMA single copy ortholog gene models, 1,952 were found in the assembly and 1,922 in the final predicted gene set – a reasonable result given the small contig sizes of the assembly. Finally, the pipeline uses a nine-way homology prediction with human, *Drosophila* and *C. elegans*, and InterPro Scan5 to allocate gene names. The automated gene sets are available from the BCM-HGSC website (https://www.hgsc.bcm.edu/arthropods/western-flower-thrips-genome-project) as well as the National Agricultural Library (https://i5k.nal.usda.gov/Frankliniella_occidentalis) where a web-browser of the genome, annotations, and supporting annotation data is accessible.

##### 1.3. Quality assessment and phylogenomics

*Contributed by Panagiotis Ioannidis, Robert M. Waterhouse, Felipe A. Simão, Evgeny M. Zdobnov*

###### 1.3.1. BUSCO quality assessment

For evaluating the completeness of the *F. occidentalis* official gene set and genome assembly and OGS we used the Benchmarking Universal Single-Copy Orthologs (BUSCO) (Simão et al., 2017). More specifically, we used the Arthropoda gene set, which consists of 1,066 single-copy genes that are present in at least 90% of selected representative arthropods. BUSCO assessments were run with the default parameters.

###### 1.3.2. Phylogenomic analysis

For the phylogenomic analysis, only the single-copy orthologs were used to build a concatenated protein sequence alignment from which to estimate the phylogenetic tree using RAxML (Stamatakis, 2006). Briefly, a multiple sequence alignment was performed using muscle (Edgar, 2004) for each orthologous group separately. Then, the resulting alignments were trimmed using trimAl (Capella-Gutierrez et al., 2009) with parameters “-w 3 -gt 0.95 -st 0.01”. The trimmed alignments were concatenated using the “seqret” program from the EMBOSS suite (Rice et al, 2000). This concatenated alignment was used to build the phylogeny using RAxML 7.6.6 with the PROTGAMMA model of amino acid substitutions and 100 bootstrap replicates.

##### 1.4. RNA evidence used to support manual curation

*Contributed by Derek Schneweis, Dorith Rotenberg and Diane Ullman*

As mentioned above, BCM-HGSC prepared RNAseq libraries for Illumina HiSeq sequencing from RNA extracted from three different samples comprised of pools of insects: pre-adults (L1, L2, P1, P2), males, and females derived from a lab colony. Using methods described previously (Badillo Vargas et al., 2012), total RNA was isolated using Trizol (manufacturer) followed by rigorous removal of DNA using Turbo DNA-free kit (Applied Biosystem). In addition, Trinity de novo-assembled contigs (Schneweis, 2017, PhD Thesis) from Illumina HiSeq reads (DNA Core Facility, University of Missouri, USA) were obtained from larval, pupal and adult stages of thrips (+/-TSWV) (NCBI Bioproject PRJNA454326) to support manual correction of difficult to locate or fragmented cytochrome p450 (CYP) gene models of interest. Published data sets were also used: de novo-assembled contigs from 454 sequencing reads for mixed stages of TSWV-infected and noninfected individuals (Badillo-Vargas et al., 2012) and de novo-assembled contigs from

Illumina RNAseq reads for salivary glands of adult females and males (Stafford-Banks et al., 2014, NCBI TSA accession GAXD00000000.).

#### 1.5. Identification of transcription factors

*Contributed by Matthew T. Weirauch*

We identified likely transcription factors (TFs) by scanning the amino acid sequences of predicted protein coding genes for putative DNA binding domains (DBDs), and when possible, we predicted the DNA binding specificity of each TF using the procedures described in Weirauch et al. (2014). Briefly, we scanned all protein sequences for putative DBDs using the 81 Pfam (Finn et al., 2010) models listed in Weirauch and Hughes (2011) and the HMMER tool (Eddy, 2009), with the recommended detection thresholds of Per-sequence Eval < 0.01 and Per-domain conditional Eval < 0.01. Each protein was classified into a family based on its DBDs and their order in the protein sequence (e.g., bZIPx1, AP2x2, Homeodomain+Pou). We then aligned the resulting DBD sequences within each family using clustalOmega (Sievers et al., 2011) with default settings. For protein pairs with multiple DBDs, each DBD was aligned separately. From these alignments, we calculated the sequence identity of all DBD sequence pairs (i.e. the percent of AA residues that are exactly the same across all positions in the alignment). Using previously established sequence identify thresholds for each family (Weirauch, M. T. et al., 2014) we mapped the predicted DNA binding specificities by simple transfer. For example, the DBD of FOCC004943-FA is 98% identical to the *Drosophila melanogaster* mirr protein. Since the DNA binding specificity of mirr has already been experimentally determined, and the cutoff for Homeodomain family of TFs is 70%, we can infer that FOCC004943-FA will have the same binding specificity as mirr.

#### 1.6. Weighted correlation network analysis

*Contributed by Joshua Benoit and Sam Bailey*

Weighted correlation network analysis (WGCNA) is a clustering method that places transcripts into colored modules based on similar patterns of expression across samples (Langfelder and Horvath, 2008). The resulting modules are used to form a correlation network to describe the correlation between genes in RNA-seq data and determine the relationship between these groups of genes and external traits. WGCNA was performed on multiple subsets of the *F. occidentalis* RNA-seq normalized read counts (FPKM values) from a published study (Schneweis et al., 2017) set to determine the role of certain gene groups in the developmental stages. The subsets used for analysis were created by excluding samples to accentuate the differences between developmental stage. Genes of zero variance were filtered out from the RNA-seq data in preparation for WGCNA. A scale-free topology threshold of 0.8 was used to identify the proper soft power of 16 for analysis. Adjacency matrix was calculated for signed network construction. Modules were determined by the dynamic tree cutting algorithm with a minimum of 20 genes per module. After relating modules to external sample traits, modules with the highest Pearson correlation coefficient were selected for further analysis.

#### 1.7. Identification of salivary gland enriched genes

*Contributed by Joshua Benoit*

Identification of the salivary gland specific genes from *F. occidentalis* was conducted based on methods developed by Telleria et al. (2014), and Ribeiro et al. (2016) with modifications. RNA-seq data sets were acquired from whole males and whole female, SRA Accession = SRX897632 and SRX897634, respectively, sequenced to assist in gene prediction for the genome. RNA-seq datasets from the salivary glands of males and females, SRA Accession: SRS549985, SRS549981, SRS549977, SRS549984, SRS549980, SRS54997, were previously published by Stafford-Banks et al. (2014). The goal with these datasets is to identify specific gene/contigs enriched within the salivary glands of male and female thrips by comparing tissue expression to whole body expression.

First, RNA-seq datasets were individually mapped to predicted genes from the *F. occidentalis* genome project using CLC Genomics based upon settings previously described[1] with the exception that transcripts per million (TPM) was used as a proxy for gene expression. Fold changes were determined as the TPM for the salivary gland RNA-seq sets divided by the TPM for whole body datasets. The Baggerly's test (t-type test statistic) (Baggerly et al., 2003) followed by a false discovery rate (FDR) at 0.05 (Benjamini and Hochberg, 1995) was used to identify genes with significant enrichment in the salivary glands. Enriched genes were removed, and mapping and expression analyses were repeated to ensure low expressed genes were not missed. In addition, the RNA-seq data sets were mapped to the transcriptome previously generated from the salivary gland RNA-seq datasets (Stafford-Banks et al., 2014), enriched contigs were identified as before, and a second mapping and analysis following removal of initially enriched contigs was utilized to identify low expressed salivary gland-enriched contigs. This secondary analysis was conducted to identify transcripts that were not predicted in the genome or may have not been present on assembled scaffold.

Contigs and genes were compared to reduce overlap to a combined final SG-enriched sequence set was generated. Briefly, blastn comparison was utilized to match sequences and only the longest sequence was retained if 100% matched was noted. After merging the non-overlapping predicted genes and contigs, the SG-enriched was searched (BLASTx) against multiple NCBI non-redundant proteins databases including those for arthropods, hemipterans, viral, bacterial, plant, *Drosophila*, and the complete nr set with an expectation value of (e-value) of at least 0.001. For transcripts with a blast hit with an e-value above 0.001, the identification was based upon the best match that included previously assigned biological function (lipase, cellulase, etc.). Following this process, transcripts with enriched expression in the salivary glands of male, females, and combined (males and females) relative to the entire body were compared to determine those that overlap between each set.

##### 1.8. Identification of sex-specific genes

*Contributed by Christopher Holmes and Joshua Benoit*

Sex- and stage-specific RNA-seq analyses were performed similarly to Benoit et al. (2018) and Schoville et al. (2018). RNA-seq sets were developed from individual replicates of female, male, and pre-adult stages (larval and pupal) stages of *F. occidentalis* as a part of the western flower thrips genome project (NCBI Bioproject: PRJNA203209). Additional thrips were sequenced as a part of the 1KITE project (*Frankliniella cephalica*, PRJNA219559; *Gynaikothrips ficorum*, PRJNA219563; *Thrips palmi*; PRJNA219609). *F. occidentalis* RNA-seq sets were used to determine differentially expressed transcripts between females, males, and nymphs. RNA-seq data for the other thrips (*F. cephalica*, *G. ficorum*, *T. palmi*) were used for identification of conserved sex- and stage-specific genes between thrips species.

The *F. occidentalis* RNA-seq set was assessed with FastQC and trimmed with CLC Genomics (CLC Bio). Reads were permitted to match up to five locations with only two mismatches and required at least 90% similarity at 70% of transcript length. Transcript levels were normalized to transcripts per million (TPM) and significant enrichment was determined with a Baggerly's test (beta-binomial distribution statistic) followed by Bonferroni correction (at 0.01) and a two-fold difference between samples. Gene identification was obtained through BLASTx searches against an NCBI non-redundant protein arthropod database with an expectation value (e-value) of at least 0.001. RNA-seq sets for the other thrips were BLAST searched against the finalized female, male, and nymph *F. occidentalis* sets with an e-value of $1.00 \times 10^{-20}$ or less. Venn diagrams (http://bioinformatics.psb.ugent.be/webtools/Venn/) were used to represent the number of shared genes in thrips females, males, and nymphs. Fold-change and number of unique gene reads from the conserved female and male thrips genes were used to identify highly enriched genes (> 1000 unique reads, > 4-fold difference) in female and male thrips**.**

### 2. Homeodomain transcription factor gene clusters (Hox, Iro-C) and Synteny

*Contributed by Kristen A. Panfilio and Iris M. Vargas Jentzsch*

#### 2.1. Abstract

The Hox and Iroquois Complex (Iro-C) gene clusters encode highly conserved homeodomain transcription factors with essential roles in development. The Hox cluster is conserved across the Bilateria [1], and the Iro-C is found throughout the Insecta [2-4]. Annotation of the genes in these clusters provides an indicator of draft genome quality and an opportunity to assess synteny among species. In *Frankliniella occidentalis* we could construct single copy gene models for all expected orthologs. In terms of synteny, we could reconstitute the small Iro-C and there is partial assembly of the larger Hox cluster.

#### 2.2. Results and Discussion

##### 2.2.1. Cluster reconstruction

We were able to find and annotate gene models for all ten Hox cluster genes, split across four different scaffolds (**Figure S2.1A**). All linked Hox genes occurred in the expected order and with the expected, shared transcriptional orientation. While these findings would suggest that the current draft assembly is correct but simply incomplete, a note of caution arises from an assessment of estimated cluster size. Assuming direct concatenation of these four scaffolds, the Hox cluster would span a region of 5.9 Mb in a genome with a total size of 415 Mb, which is disproportionately large (3.5-fold larger relative cluster size compared to previously analyzed i5k pilot species and the beetle *Tribolium castaneum*) and suggests incorrect assembly of the non-Hox portions of some of these scaffolds. For example, the scaffold regions upstream of *Hox1/labial* and of *Hox4/Deformed* are surprisingly large. (Note that while the Hox genes are numbered in ascending order in the 5' to 3' direction, the genes are in fact transcribed on the opposite strand. Hence, "upstream" of *Hox1* refers to the genomic region between *Hox1* and *Hox2*.)

Assembly limitations are also manifest in that only partial gene models could be created for *Ultrabithorax* and *Abdominal-B*, where the missing coding sequence includes the highly conserved homeobox, which encodes the key functional domain of the DNA-binding homeodomain (**Figure S2.1B**).

For the small Iroquois Complex, clear, single copy orthologues of both *iroquois* (*iro*) and *mirror* (*mirr*) are indeed linked in the current assembly (**Figure S.2.1C**). As expected by conservation, the genes occur in the same transcriptional orientation, with *iro* upstream of *mirr*, and with no intervening non-Iro-C genes. Also, unlike the Hox cluster estimation, the *Frankliniella* Iro-C is conserved for the ratio of cluster size to genome size, despite gaps on the relevant scaffold.

##### 2.2.2. Gene structure and protein coding sequence divergence in thrips

Although all ten *Frankliniella* Hox genes could be identified and their orthology is clear, they are in some features rather divergent compared to other insects (*Zootermopsis nevadensis, Cimex lectularius, Oncopeltus fasciatus, Pediculus humanus corporis, Tribolium castaneum, Anoplophora glabripennis,* and *Drosophila melanogaster*). Specifically, *Focc-zerknüllt* encodes the largest protein among these orthologs (439 aa compared to a mean of 300 aa), while *Focc-Antennapedia* and *Focc-abdominal-A* encode larger proteins than all other species except for *Drosophila* (~14% larger than the mean, excluding *Drosophila*). Meanwhile, three of the *Frankliniella* Hox genes – *Deformed*, *fushi tarazu*, and *abdominal-A* – have uniquely acquired additional introns that interrupt coding sequence exons in what are otherwise highly conserved gene structures across the Insecta (>300 myr divergence time). While the gene locus is incomplete, the partial model for *Abdominal-B* has also acquired additional introns, which interrupt exons that encode the 5' UTR.

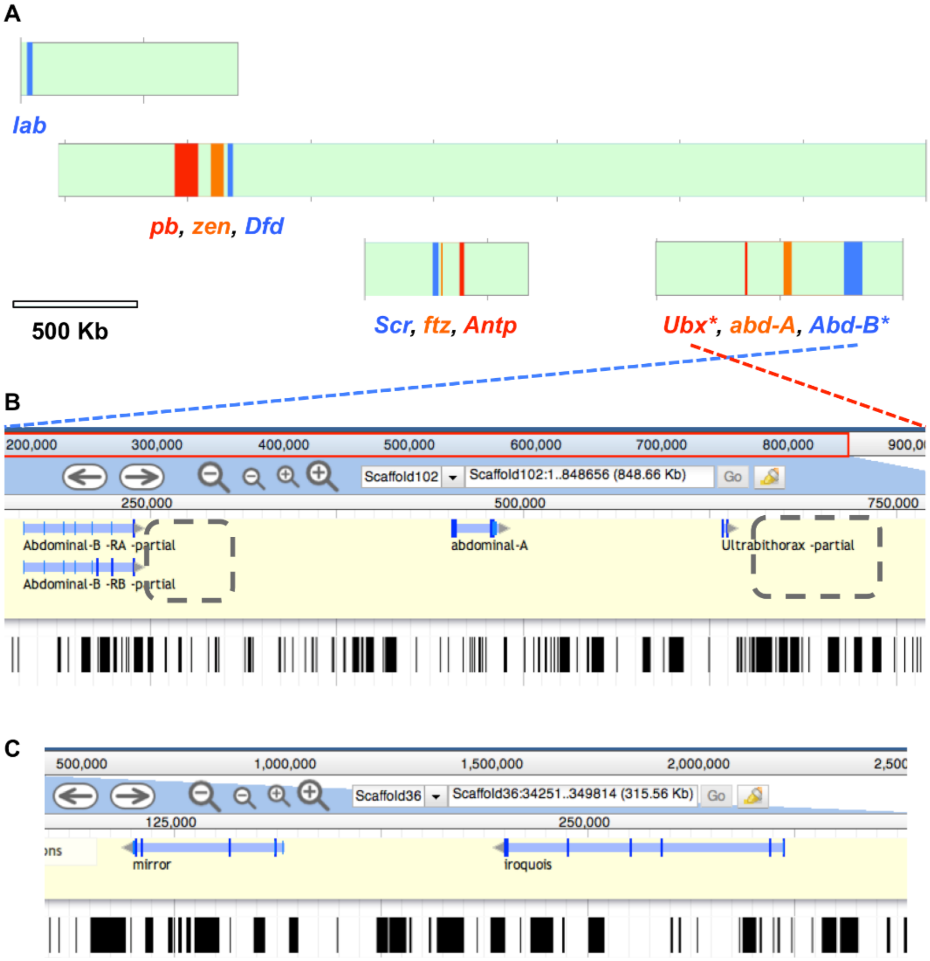

**Figure S2.1. *Frankliniella* Hox and Iro-C clusters.** (A) Scaffolds containing Hox genes are shown to scale (green), with Hox gene loci indicated. Asterisks indicate incomplete gene models. (B) Genome browser screenshot for the posterior Hox genes. Dashed gray boxes indicate where missing coding sequence exons would be expected, and gaps in the assembly are indicated below by black bars. (C) Genome browser screenshot of the Iro-C.

**Table S2.1.** Positional information for the annotated homeobox genes. Incomplete gene models are marked with an asterisk (*).

| Gene | Scaffold: start..end | Locus length (nt) | Protein length (aa) | Number of CDS exons |
|---|---|---|---|---|
| *labial* | Scaffold124:24063-47398 | 23,336 | 368 | 2 |
| *proboscipedia* | Scaffold11:2956730-3052532 | 95,803 | 733 | 4 |
| *zerknüllt* | Scaffold11:2852821-2905256 | 52,436 | 439 | 2 |
| *Deformed* | Scaffold11:2814925-2836826 | 21,902 | 447 | 3 |
| *Sex combs reduced* | Scaffold161:275917-299792 | 23,876 | 352 | 2 |
| *fushi tarazu* | Scaffold161:312168-313263 | 1,096 | 255 | 3 |
| *Antennapedia* | Scaffold161:385185-404098 | 18,914 | 367 | 2 |
| *Ultrabithorax** | Scaffold102:633776-637749 | 3,974 (partial) | 219 (partial) | 2 |
| *abdominal-A* | Scaffold102:452477-485096 | 32,620 | 394 | 5 |
| *Abdominal-B** | Scaffold102:165158-239410 | 74,253 (partial) | 238 (partial) | 1 or 3 (two isoforms) |
| *iroquois* | Scaffold36:225,058..310,105 | 85,048 | 806 | 7 |
| *mirror* | Scaffold36:113,167..159,432 | 46,266 | 591 | 5 |

### 3. Lateral Gene Transfers

*Contributed by John H. Werren, Sammy Cheng, Clauvis N.T. Taning, Dong Wei and Guy Smagghe*

#### 3.1. Abstract

Lateral gene transfers (LGT) can contribute to adaptations in insects especially by including the ability to utilize plant resources. We have identified three very interesting LGTs in the *F. occidentalis* genome, which have the hallmarks of ancient lateral gene transfers into the insect genome that have subsequently undergone gene family expansions, and that show some signatures of having evolved function in the insect. These include two carbohydrate metabolism genes (mannanase and levanase) of bacterial origin, and an O-methyltransferase gene derived from bacteria.

#### 3.2. Methods

##### 3.2.1. Bacterial Scaffold Detection Method

Bacterial scaffolds in *F. occidentalis* genome were identified using a modified nucleotide- based pipeline developed by Wheeler et al. (2013) and as described previously (Panfilio et al., 2019). Briefly, one kilo base pair DNA fragments from each scaffold were searched for bacterial homologs against an in-house bacterial database containing 2,100 bacterial species using BLASTn algorithm (**Additional File 2: Table S.25**). The bacterial database was masked for low complexity regions by NCBI Dustmasker (Morgulis et al. 2006), and similarity matches of bitscore above 50 were retained. To accurately determine the bacterial scaffolds, parameters including the number of bacterial matches per scaffold, proportion of the scaffold covered by bacterial matches, and total hit width (encompassing the distance between the leftmost and rightmost bacterial match in the scaffold) were considered. Candidate bacterial scaffolds were called based on a cut-off of ≥ 0.40 proportion bacterial hit width as this criterion with manual curation of the sequences. It should be noted, however, that represented in this set could be larger lateral gene transfers in relatively small scaffolds, as the latter cannot be readily identified without flanking eukaryotic sequences and/or further manual curation. The procedure for detecting bacterial scaffolds was performed twice with modifications – once during the early stages of the curation process (2015) using an 'older' method (Wheeler et al., 2013) and a second time more recently as described above using this 'new' method after the curation process ended and OGS v1.0 was submitted for publication. As such, only the first round of contaminating bacterial scaffolds were removed from OGS v1.0.

##### 3.2.2. Lateral Gene Transfer Detection Method

Bacterial scaffolds to detect lateral gene transfers were identified using the method described in the section 3.2.1. Analysis was limited to scaffolds more than 100 kb due to the need for flanking sequences to properly evaluate candidate LGTs. Additionally, positive bacterial hits sharing conserved genes with eukaryotes in a custom-built eukaryotic databases (ftp://ftp.hgsc.bcm.edu/I5K-pilot/LGT_analysis/ All_species_genomes/ lgt _finder_blastn_database_directories/) were filtered out. To focus on the more likely LGT candidates, we examined each bacterial match with bitscore >75 that also showed a bitscore = 0 in a reference eukaryote database. Fragments flanking the 1 kbp positive hits were examined and combined for LGT analysis.

Manual curation was conducted on candidates surviving the initial filtering steps. Each candidate was searched with BLASTn to the NCBI nr/nt database. Sequences homologous to insect genes with the exception of a possible LGT in the common ancestor of closely related insects were removed. In cases where the matches to other insects were sporadic, the candidate was retained, as our experience has indicated that these can be independent LGTs into different lineages. The region was additionally searched with BLASTx to the NCBI nr/nt database. Sequences with no hits to multiple insect proteins, were identified as an LGT candidate. The best bacterial match to the candidate was called using the NCBI nr and protein databases, and flanking genes within the scaffold were examined to determine whether they are eukaryotic or bacterial. We examined whether the LGT region was associated with an annotated gene within the insect genome, and if RNA sequencing data showed evidence of transcriptional activity in the LGT region. In a few cases, polymerase chain reactions were also conducted using primers that bridge the LGT candidate and flanking eukaryotic-like sequences (see Section 3.2.3).

##### 3.2.3. PCR of LGT Candidate flanking regions

The overlapping primers used to verify whether the potential LGTs were present in the thrips genome were designed using DNAMAN 7.0 (Lynnon Biosoft, Quebec, Canada) (**Table S3.2**). The PCR was carried out in a 20 μL reaction volume containing 4 μL 5x Phusion HF Buffer or 5x Phusion GC Buffer (depending on the difficulty in amplifying the target fragment), 0.4 μL 10 mM dNTPs, 1.0 μL 10 mM each primer, 0.6 μL DMSO, 0.2 μL Phusion DNA High-Fidelity DNA Polymerase (Thermo Scientific), 1.0 μL genome DNA template, and 11.8 μL ddH2O. The cycling program was set at: 98 °C for 30 sec, and then 34 cycles of 98 °C for 10 s, and 72 °C for ~2 min (30 sec/kb), followed by a final extension step of 72 °C for 10 min and 4 °C hold. The PCR products were checked on a 1.0% gel after electrophoresis and then purified using a Wizard SV Gel and PCR Clean-Up System (Promega). The purified fragments were cloned into pGEMT Easy vector (Thermo Scientific) and transformed into DH5α competent cells. The positive transformants were cultured for plasmid purification using the E.Z.N.A. Plasmid Mini Kit I, (V-spin) (Omega Bio-tek) and then sent for sequencing by LGC genomics (https://www.lgcgroup.com). Overlapping sequences obtained from the different amplification reactions were then re-assembled and used to verify if the LGT was indeed present in the thrips genome.

##### 3.2.4. Phylogenetic analysis and nucleotide sequence evolution of putative LGTs

The most promising LGT candidates were further analyzed by evaluating phylogenetic relationships and conducting branch specific synonymous and non-synonymous rate analysis with homologous references from NCBI to detect signatures of stabilizing or directional selection.

For phylogenetic analysis of protein sequences, the top proteins with the strongest bitscores (up to 50-60 proteins) to the translated region of the *F. occidentalis* proteins were aligned using muscle in MEGA. Protein alignments were assessed for misalignment and large in-del regions were removed. Phylogenetic trees were constructed by RAXML protein model "LG" with 1500 bootstrap replications. Outgroups were identified by identifying the bacterial species closest to the LGT in the constructed phylogeny and taking the bacteria protein and blasting to NCBI's nr protein database restricted to the bacteria's taxonomic group (e.g. family or order). Another phylogenetic tree including proteins within different genera from the bacteria taxonomic family was constructed.

To characterize nucleotide sequence evolution of the LGT and related sequences (e.g. synonymous and non-synonymous substitution rates), the translated nucleotide sequence were identified using the protein query from representative species throughout the protein tree and by comparing the similarity to the NCBI nucleotide database restricted to the specific species. The sequences obtained were then aligned using MUSCLE and large in-del regions were removed. A pruned version of the protein tree was created with the same representative sequences obtained for the nucleotide alignment. HYPHY's BUSTED and BUSTEC were both performed to validate the open reading frame using the nucleotide aligned sequences and the pruned protein tree. The branch specific non-synonymous (dN) and synonymous (dS) were calculated using PAML CODEML free-model rate on the same alignment and the protein tree topology.

##### 3.2.5. Potential Bacterial Scaffold Detection

There were 102 potential bacterial contaminating scaffolds located in the genome sequence (**Additional file 2: Table S26**). The two largest contaminating bacterial scaffolds (scaffold 122 and scaffold 83) corresponded to parts of two bacterial associate genomes of *F. occidentalis* previously described in Facey et al (2015), BioProject PRJNA234511: Bfo1 ((SAMN03389132 ID: 3389132) and Bfo2 (SAMN03389135 ID: 3389135) respectively. Nucleotide sequences from this NCBI BioProject were found using targeted searches.

#### 3.3. Results and Discussion

Growing evidence suggests that such LGTs can contribute to adaptations in insects, including the ability to utilize plant resources (Wybouw et al., 2016). We have focused our attention on three very interesting LGTs in the *F. occidentalis* genome, which have the hallmarks of ancient lateral gene transfers into the insect genome that have subsequently undergone gene family expansions with evolved functions in insects. These include two carbohydrate metabolism genes, mannanase and levanase; and an O-methyltransferase gene all derived from bacteria.

##### 3.3.1. O-methyltransferase

O-methyltransferase is involved in methylation of small molecules and is known to affect diverse biological processes in bacteria, plants and animals, including cell signaling and catalytic activities (Liscombe et al 2012). Originally found on scaffold 147, an O-methyltransferase also showed negligible similarity within insecta compared to the bacteria at both nucleotide and protein level. The sister group consisted of a bootstrap score of 85, which may suggest that Silvanigrellales bacterium could be a potential sister group to the O-methyltransferase. However, the bacterial source of the O-methyltransferase gene remains obscure, likely due to incomplete sampling of associated bacteria in insects. It clusters most closely with an O-methyltransferase gene sequence from the Silvanigrellales bacterium RF1110005 (bootstrap value 86, **Figure S3.1**) isolated from Lake Sanaru Japan; related sequences also come from environmental samples, such as a Bdellovibrionales bacterium assembled from a metagenomic sample obtained from the groundwater in Utah, USA.

Two other copies of the O-methyltransferase were found and all three can be seen to cluster together on the tree which indicates gene duplication events after transfer from the bacterium. Out of the three copies of O-methyltransferase, none of the terminal branches after gene duplication show significant purifying selection compared to either a neutral or positive selection background. However, one copy (XP_026277179.1) shows significant positive selection. Nevertheless, they have shown maintenance of the open reading frame and dn/ds ratios indicative of purifying selection (**Table S3.1**).

##### 3.3.2. Mannanase

Mannanase in bacteria hydrolyzes the endo- β1,4 glycosidic bond in carbohydrates (Wang et al, 2013). The mannanase was originally found on scaffold 197 and homology searched showed high similarity to bacterial references at both nucleotide and protein level. There were two other copies of the gene indicating subsequent gene duplications, based on their phylogenetic position (**Figure S3.2**). The *F. occidentalis* mannanase proteins are clearly embedded among bacterial sequences, although the actual sister group cannot be confidently identified due to low bootstrap support for adjacent bacterial sequences in the tree. The BUSTEC test for all three mannanase genes showed significant purifying selection p-value of 0.0066, 0.0000, 0.0016 for XP_026276666.1, XP_026285291.1, and XP_026285289.1 respectively, with no significant directional selection indicated for any of the branches (**Table S3.1**). The dN dS values are consistent with maintenance of the open reading frame during purifying selection.

##### 3.3.3. Levanase

Levanase (GH32) is involved in sugar metabolism (Wanker, 1991). Initially, scaffold 31 and scaffold 54 indicated a levanase similar bacterial genes with a high homology to bacterial nucleotide references. Protein blasts of the thrips levanase LGT also gives very strong matches to bacterial proteins, particularly Streptomyces and Massilia genus. Matches to insect references are sporadically distributed indicating likely independent LGTs. In addition, the other insect candidates match to different bacterial sources than the levanase found in the thrips, suggesting that they are likely derived from independent LGTs. For example, an ancient LGT of a bacterial levanase derived from genus *Bacillus* has been found in Lepidoptera has been described (Sun et al. 2013). The data suggest that levanases may be prone to retention and functional evolution after lateral transfer.

Reconstruction of the phylogenetic tree for the thrips levanase reveals that the two LGTs cluster together in the same clade, and therefore are paralogs that duplicated after the ancestral LGT event (**Figure S3.3**). The phylogenetic reconstruction after removal of bacterial gene duplicates suggests the sister group to be Streptomyces *sp*, however, the origin of the LGT cannot be unambiguously resolved at this time. BUSTEC test on the 2 LGT duplicates showed both have undergone significant purifying selection with p-value of 0.0000 and 0.0002 for XP_026287851.1 and XP_026273122.1 respectively. None showed significant positive selection on the BUSTED test. The dN/dS are provided in **Table S3.1.**

#### Figure S3.1: Phylogenetic analysis of *Frankliniella occidentalis* O-methyltransferase

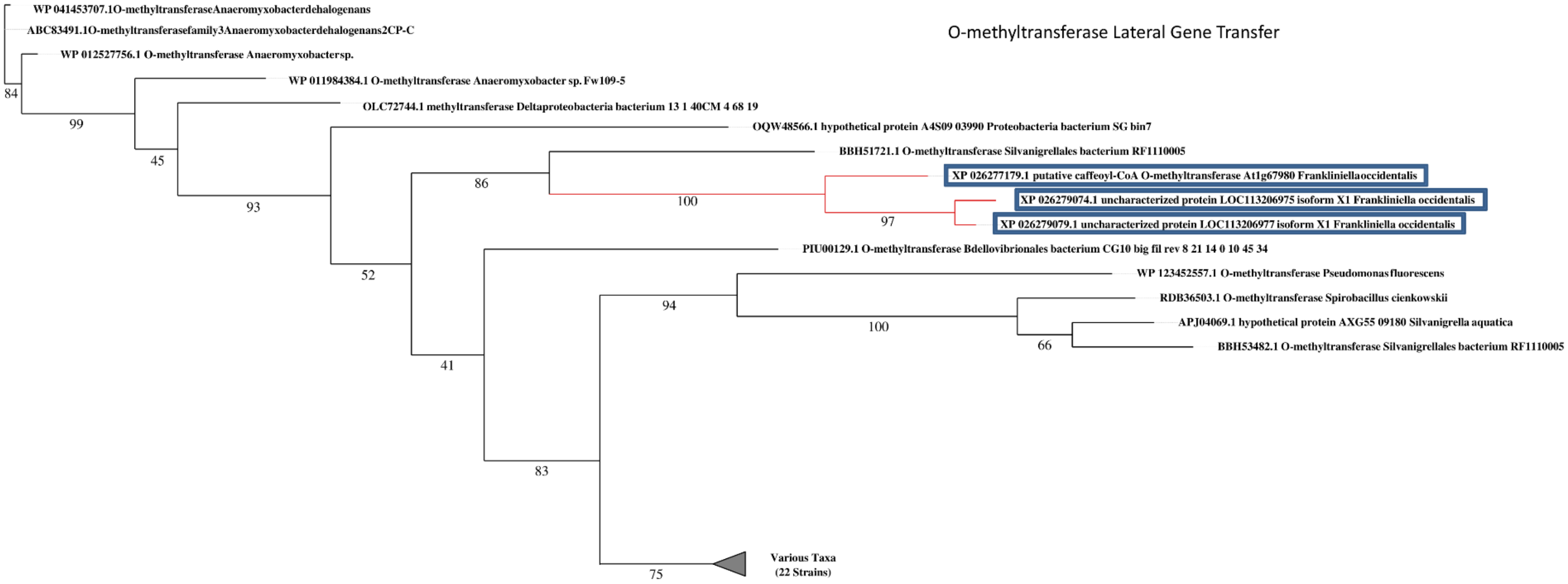

Maximum likelihood phylogeny generated using RAXML "PROTGAMMAILG" protein model at 1500 bootstraps for the o-methyltransferase including bacterial sequences obtained from NCBI blastp hits of the Frankliniella occidentalis LGT protein. Collapsed nodes contain indicated bacteria, and accession numbers are provided for reference. Ascension number for other various taxa: PIR17430.1, OGS05852.1, PIR39089.1, WP 015589357.1, WP 075138886.1, RMG88810.1, RMF77243.1, PJF39726.1, WP 119068776.1, RME13388.1, RPI81709.1, PYP20167.1, KPK66093.1, WP 078760385.1, WP 084171608.1, WP 036660971.1, WP 082067168.1, ODB56458.1, WP 007905237.1, WP 069624822.1, WP 036660971.1, WP 052282472.1.

#### Figure S3.2: Phylogenetic analysis of *Frankliniella occidentalis* mannanase

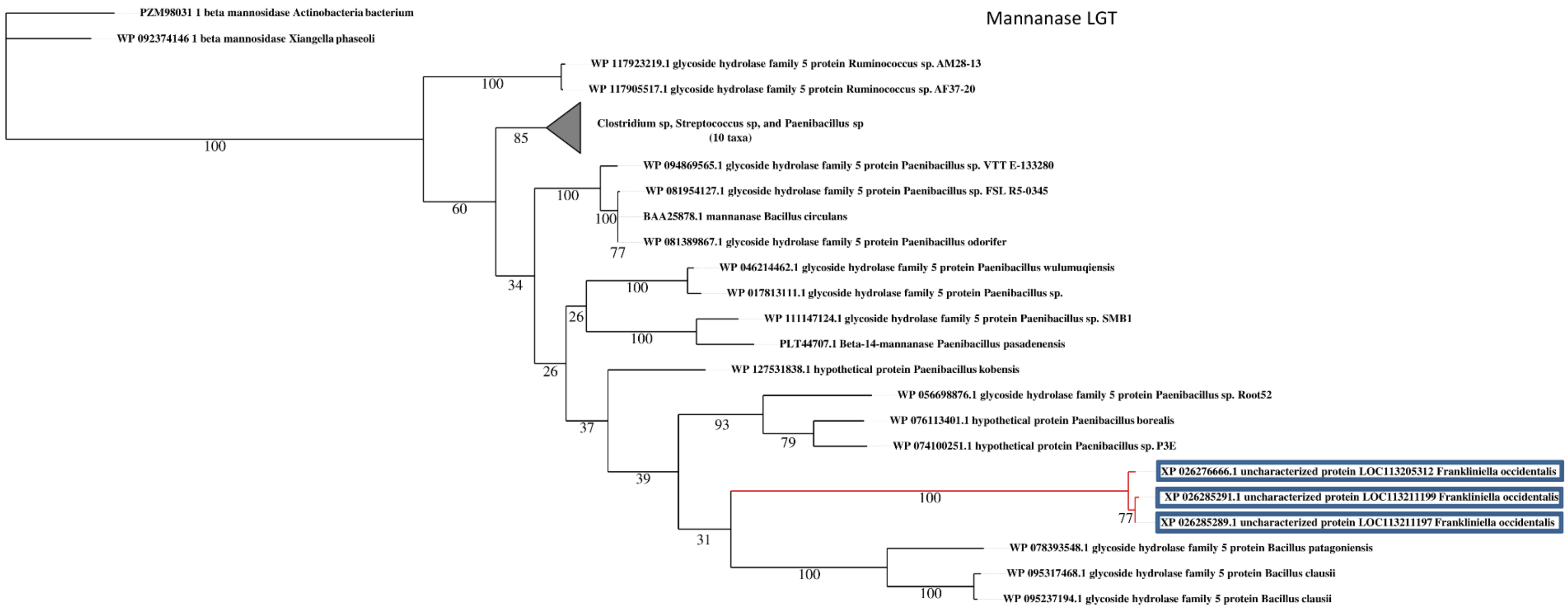

Maximum likelihood phylogeny generated using RAXML "PROTGAMMAILG" protein model at 1500 bootstraps for the protein mannanase including bacterial sequences obtained from NCBI blastp hits of the Frankliniella occidentalis LGT protein. Collapsed nodes contain indicated various bacterial taxa, and accession numbers are provided for reference. Ascension number for other bacterial species. Clostridium sp, Streptococcus sp, and Paenibacillus sp: WP 003428823.1, WP 027635375.1, WP 074561201.1 WP 012961352.1, WP 128211511.1, PWV88428.1 WP 081376921.1, WP 103049993.1, WP 127198212.1, WP 042138990.1.

#### **Figure S3.3: Phylogenetic analysis of *Frankliniella occidentalis* Levanase**

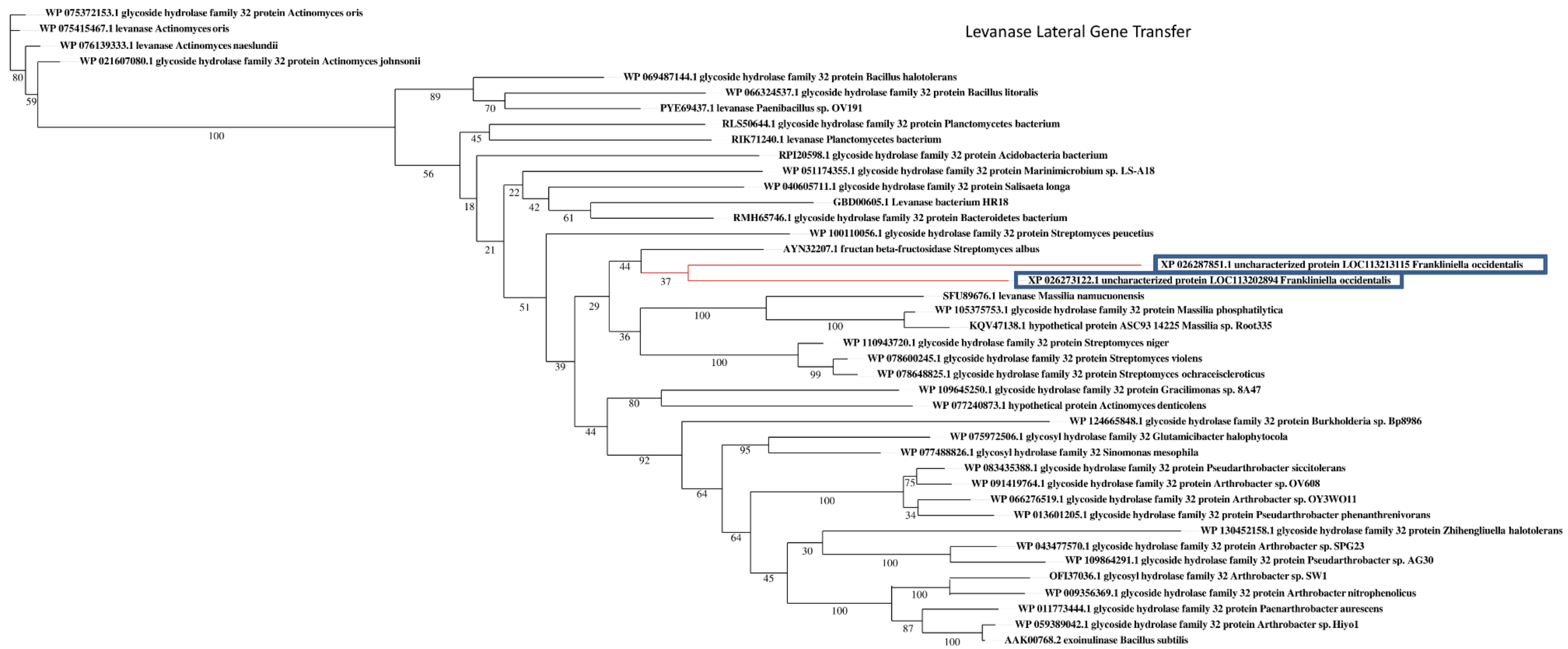

Maximum likelihood phylogeny generated using RAXML "PROTGAMMAILG" model at 1500 bootstraps for glycoside hydrolase family 32 protein levanase including bacterial sequences obtained from NCBI blastp hits of the Frankliniella *occidentalis LGT protein.*

#### Table S3.1: Lateral Gene Transfers identified in the *Frankliniella occidentalis* genome

| Scaffold | NCBI protein sequence accession | dN/dS | dN | dS | BUSTEC purifying, base branch combined (P-value) | BUSTED positive, base branch combined (P-value) | Expression (FPKM)[1] | | |
|---|---|---|---|---|---|---|---|---|---|
| | | | | | | | L1 | P1 | Adult |
| **Levanase (glycoside hydrolase)** | | | | | | | | | |
| **Branch leading to LGT event** | | 999.000 | 0.0727 | 0.0001 | | | | | |
| **Scaffold54** | XP_026287851.1 | 0.0667 | 0.2869 | 4.3031 | Yes (0.0000) | No | 0.68 | 1.50 | 119.2 |
| **Scaffold31** | XP_026273122.1 | 0.0389 | 0.1753 | 4.5068 | Yes (0.0002) | No | 9.17 | 0.16 | 9.94 |
| **Mannanase (endoglucanase)** | | | | | | | | | |
| **Branch leading to LGT event** | | 0.1185 | 1.2311 | 10.3877 | | | | | |
| **Scaffold96** | XP_026276666.1 | 0.1971 | 0.0051 | 0.0261 | Yes (0.0066) | No | 1.1 | 0.04 | 0.88 |
| **Scaffold197** | XP_026285289.1 | 0.0001 | 0.0000 | 0.0048 | Yes (0.0016) | No | 0 | 0 | 0.08 |
| **Scaffold197** | XP_026285291.1 | 0.1639 | 0.0035 | 0.0212 | Yes (0.0000) | No | 11.77 | 0.05 | 3.90 |
| **O-methyltransferase family 3** | | | | | | | | | |
| **Branch leading to LGT event** | | 0.2998 | 0.3820 | 1.2741 | | | | | |
| **Scaffold147** | XP_026279074.1 | 0.1874 | 0.0147 | 0.0783 | No | No | 0.04 | 0 | 0 |
| **Scaffold147** | XP_026279079.1 | 0.0465 | 0.0049 | 0.1052 | No | No | 2.00 | 2.26 | 8.34 |
| **Scaffold3388** | XP_026277179.1 | 0.1438 | 0.0677 | 0.4704 | No | Yes (0.000) | 31.01 | 1.80 | 52.86 |

[1]Average normalized read counts (FPKM) across four biological replications (data obtained from Schneweis et al., 2017)

#### Table S3.2: Primer sequences that bridge the LGT candidate and flanking eukaryotic-like sequences.

| Scaffold | Distance from neighboring thrips gene (kb)[1] | PCR Primer ID | Fragment size (kb) | Primer sequence (5′ → 3′) | Amplification result |
|---|---|---|---|---|---|
| O-methyltransferase family 3 | | | | | |
| Scaffold147 | 80 kb | 147.51 | 6.5 | TGTCTCTGAAACTGCCCA | positive |
| | | 147.31 | | ATGTAATGCCCGTCGTGT | |
| | | 147.52 | 1.66 | CCAGTATTACAGTCGGGTTCTT | positive |
| | | 147.32 | | CTTGGCGTTCGTAAGTGAA | |
| Mannanase/endoglucanase | | | | | |
| Scaffold197 | 35 kb | 197.F1 | 4.80 | ATGTTCGTTGAGGTGGTGC | negative |
| | | 197.R1 | | TTGGTTGGTCAGAGGATGC | |
| | | 197.F2 | 3.67 | AAGAAGAAGGCAAAGAGCG | positive |
| | | 197.R2 | | TAGACCAACCCGTCAATGA | |
| Levanase (Glycoside hydrolase) | | | | | |
| Scaffold31[2] | 300k | 31.F1 | 4.56 | GAGCAGCCCTATCAGTGTT | positive |
| | | 31.R1 | | GATGGCGTAGATTTCAGAGA | |
| | | 31.F2 | 3.65 | CTCGCTTATCAGGGTCTTTAC | positive |
| | | 31.R2 | | CTTCGTTCGTTCACTCATTC | |
| | | 31.F3 | 3.63 | ACCAGTGAGGTCAACAATGA | positive |
| | | 31.R3 | | TGATTAGATGTGCGAGGAAC | |
| Scaffold54 | 45k | 54.F1 | 3.09 | CGGACCTAACTTCAGGAACA | positive |
| | | 54.R1 | | CGTGTGATGTGTGCGAAT | |
| | | 54.F2 | 6.09 | GGTTTATCGGCACTGACCA | negative |
| | | 54.R2 | | CCTTTACTCGTCTTCCACCCT | |
| | | 54.F3 | 2.87 | CACACACACACACACCTCAT | positive |
| | | 54.R3 | | GTGCTCTGCTTGAAGTTCC | |

[1]Linkage between the putative LGT and neighboring scaffold sequence (linker) that putatively ‘connects’ the LGT to the nearest Frankliniella occidentalis (FOCC) gene model indirectly, and vice versa; however, the distances between the LGT and nearest FOCC thrips gene on the scaffold was too long to confirm direct linkage between the LGT and gene model.

[2]Linkage confirmed between LGT and nearest FOCC gene (zinc carboxypeptidase-like), complete confirmation

### 4. Chemosensory receptors

*Contributed by Hugh M. Robertson*

#### 4.1. Methods

The three major chemoreceptor gene families were manually annotated as for several other hemipteroids (Smadja et al. 2009; Kirkness et al. 2010; Terrapon et al. 2014; Mesquita et al. 2015; Benoit et al. 2016; Panfilio et al. 2017; Armisen et al. 2018) and many other insects and other arthropods using the Apollo genome browser at the i5k Workspace@NAL (Poelchau et al. 2014). Sensitive TBLASTN searches (E = 1000 and Word Size = 2) with other hemipteroid proteins as queries were used to identify loci and gene models built using a combination of the V0.5.3 gene models as well as Augustus and Snap models, supported by some spliced RNAseq reads, and knowledge of the expected gene structures supported by splice site predictions from the Splice Site Prediction by Neural Network webserver at the Berkeley Drosophila Genome Project website (http://www.fruitfly.org/seq_tools/splice.html). Iterative searches with each newly identified gene/protein were employed to search exhaustively for additional members of each family. Encoded proteins, including conceptual translation of pseudogenes using Z for stop codons and X for frameshifts and intron boundary mutations, were aligned with those from other hemipteroids and *D. melanogaster* representatives using ClustalX v2.0 (Larkin et al. 2007), and gene models were refined in light of these alignments. The final alignments included the families from the human body louse *Pediculus humanus* (Kirkness et al. 2010), the pea aphid *Acyrthosiphon pisum* (Smadja et al. 2009), and the bedbug *Cimex lectularius* (Benoit et al. 2014). The IR family was not described in these publications for the first two species, but was partially described in Croset et al. (2010) with additional genes and refinement of gene models provided in Terrapon et al. (2014). Additional available protein sets for these three families from the assassin bug *Rhodnius prolixus* (Mesquita et al. 2015), the milkweed bug *Oncopeltus fasciatus* (Panfilio et al. 2017) and a waterstrider *Gerris buenoi* (Armisen et al. 2018) do not contribute to the diversity of receptors known from heteropterans beyond those of the bedbug, so were not included in this analysis. Representatives of conserved proteins from *D. melanogaster* (Robertson et al. 2003; Benton et al. 2009), as well as a few other endopterygote species, were including in the GR and IR analyses for comparison. The alignments were trimmed to remove regions with mostly gaps using TrimAl v1.4 (Capella-Gutierrez et al. 2007), with the “gappyout” option for the OR and GR families that have generally uniform lengths, and the “strict” option for the IRs, which vary enormously in the length and sequence of their N-terminal regions, effectively removing most of this from the alignment. Phylogenetic analysis was performed using maximum likelihood analysis with PhyML v3.0 (Guindon et al. 2010) at the ATGC webserver (http://www.atgc-montpellier.fr/phyml/). The resultant trees were prepared with FigTree v1.4.2 (http://tree.bio.ed.ac.uk/software/figtree/) and Adobe Illustrator.

#### 4.2. Results and Discussion

##### 4.2.1. Gene modeling difficulties

This draft genome assembly for *F. occidentalis* has many gaps as well as some misassemblies. These caused considerable difficulties for modeling of these chemoreceptor genes, especially for sets of closely related genes as well as divergent genes that are not well enough expressed to have RNAseq reads in the available datasets, all of which are from whole animals. RNAseq support allowed resolution of some misassemblies, as well as discovery of exons missing in gaps, which were manually built using raw genome reads from the Sequence Read Archive (SRA) at the National Center Biotechnology Information. These three gene families were initially manually annotated in 2015, and revisited in 2018 with assistance from additional RNAseq information in the SRA. Nevertheless, many gene models remain incomplete, especially in large recent expansions such as the IR102-268 gene set. Examples of these difficulties are exemplified by the IR family detailed below, where among the first 15 intron-containing genes just four are intact in the assembly, (gene name suffixes are F – assembly repaired, J – model joined across scaffolds, N – N-terminus unidentified).

Ir8aF – Assembly has an 18bp deletion removing the front of the last exon, repaired with RNAseq and raw genomic reads

Ir21aJ – N-terminal and C-terminal exons on the ends of a large scaffold and a 3kb contig.

Ir25aF – Three internal exons missing, built from RNAseq and raw genomic reads

Ir40a – Intact.

Ir68aJ – N-terminal and C-terminal exons on the ends of two large scaffolds.

Ir76bF – Just three exons present, flanked by large gaps – RNAseq used to find four C-terminal exons not in assembly, plus single N-terminal exon misassembled downstream in next contig after a gap, supported by spliced RNAseq reads.

Ir93aJ – Mostly in a large scaffold, but N-terminal exons in a 19kb scaffold that must belong in a short gap within the large scaffold.

Ir75aNJ – Most exons are in a 16kb scaffold that apparently belongs in a gap in a much larger scaffold that has the final two exons. N-terminus missing in gap, but no RNAseq to help find it.

Ir75bJ – N-terminal and C-terminal exons are in middles of two large scaffolds, so some kind of misassembly, connection supported by spliced RNAseq reads.

Ir75cJF – Three central exons are in a 1kb scaffold that belongs in a gap in a much larger scaffold, and two exons in this larger scaffold are in inverted order.

Ir75d – Intact.

Ir75e – Intact.

Ir75fN – In a 16kb scaffold but N-terminal exon unidentified.

Ir7gNJ – Four central exons are in a 1kb scaffold that belongs in a gap in a much larger scaffold, but N-terminus not identified in absence of RNAseq.

Ir75hNJ – Joined across two scaffolds on basis of RNAseq. Can't find N-terminal exon.

Ir101 – Intact.

Ir102-243 – Largely mostly intronless genes, but 52 (one third) have one or both ends missing in gaps, presumably because they are so similar to each other the assembly was unable to build them completely. There are many more fragments not included in the named genes.

Ir244-268 – Three-exon genes, and 4 have parts missing in assembly gaps.

##### 4.2.2. The OR family

The OR family is unique to insects (Missbach et al. 2014; Brand et al. 2018; Robertson 2019) and usually consists of a single Orco gene and a set of "specific" ORs that mediate specificity and sensitivity of most of insect olfaction (Joseph and Carlson 2015). The expected single conserved Orco gene has the first two exons in a separate 1.6 kb scaffold that can confidently be placed in a gap within 1.4Mbp Scaffold63 based on both sequence conservation and spliced RNAseq reads. Amongst the 84 "specific" ORs, 43 are intact full-length genes (15 of which required repair of the assembly and one was joined across two scaffolds). Of the remaining 41 genes, just two are apparent pseudogenes, while the rest have parts missing in gaps, or might also be pseudogenes, but are assumed to be intact in the genome. As shown by the tree in **Figure S4.1**, all of these thrips ORs form a distinctive species-specific clade, commensurate with the generally rapid sequence divergence of ORs in insects and the phylogenetic divergence of thrips from other hemipteroid orders represented here. Like all other hemipteroid insects to date, no ligand specificities are known for any of the specific ORs, and their enormous divergence from those of endopterygote insects with known ligand specificities preclude any inferences of ligand specificity and hence specific roles in thrip biology, however they are inferred to mediate the specificity and sensitivity of most thrip olfaction, in particular sensing host plant volatiles (Tuelon et al. 1993; Koschier et al. 2000; de Kogel and Koschier 2002; Mainali and Lim 2011; Cao et al. 2014; Silva et al. 2016) as well as pollen for food (Abdullah et al. 2014). They likely also mediate perception of alarm (Teerling et al. 1993; de Brujin et al. 2006), aggregation (Hamilton et al. 2005), and sex pheromones (de Kogel and van Deventer 2003; Kirk and Hamilton 2004; Olaniran et al. 2013).

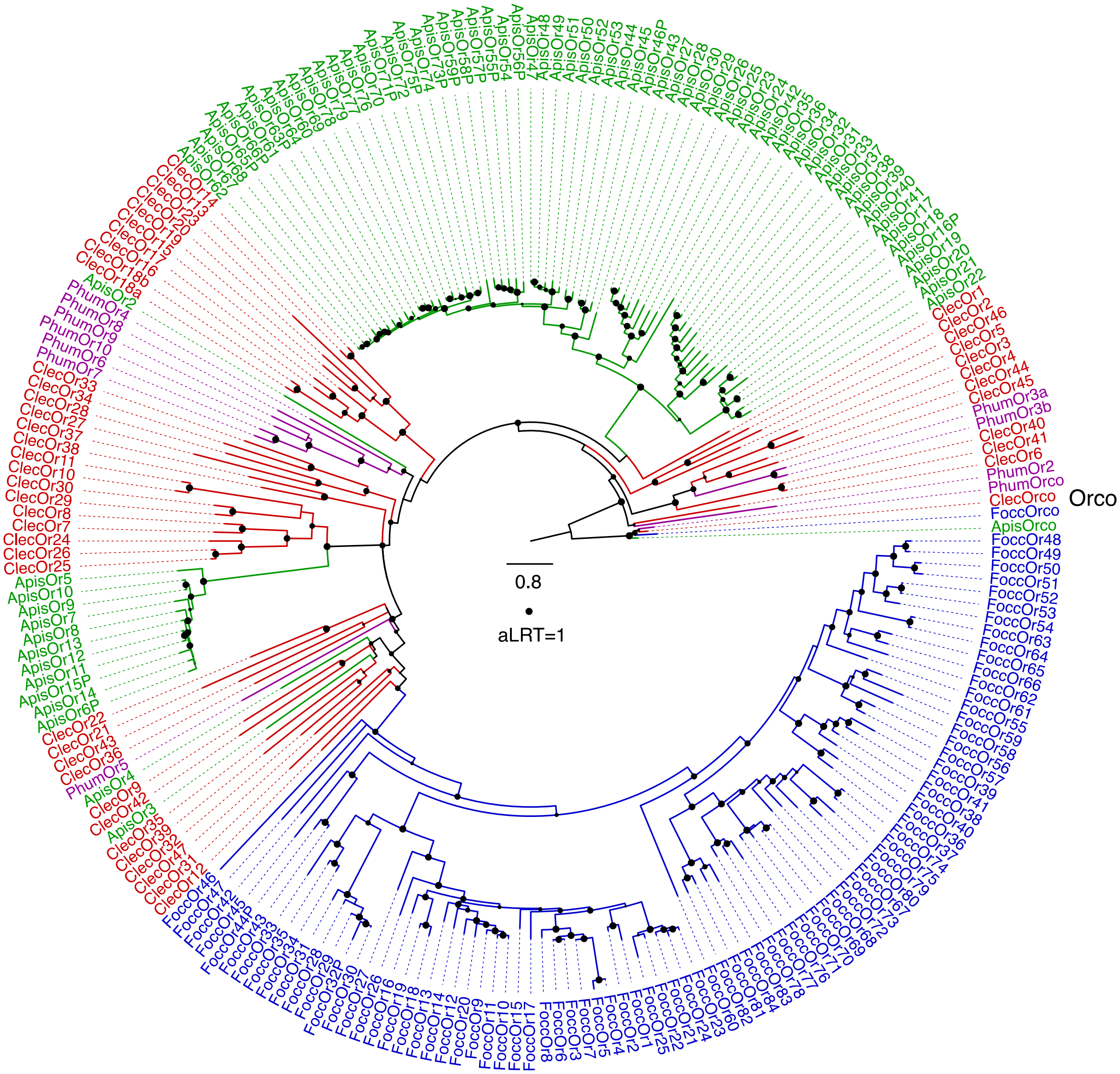

**Figure S4.1. Phylogenetic relationships of the odorant receptors of hemipteroid insects.** The Orco lineage was declared the outgroup to root the tree, based on its basal position in the OR family in analyses of the entire insect chemoreceptor superfamily (Robertson et al. 2003; Missbach et al. 2014). The scale bar is substitutions per site, and the filled dots are approximate likelihood ratio test (aLRT) values from PhyML ranging from 0-1. The list of FOCC OGS v1.0 models for OR genes are found in **Additional file 2: Table S7.**

##### 4.2.3. The GR family

The GR family is far older than the OR family, which evolved from within it (Robertson et al. 2003; Missbach et al. 2014; Brand et al. 2018), and has evolved multiple divergent lineages since its origin in basal animals (Robertson 2015, 2019; Saina et al. 2015; Eyun et al. 2017). The most prominent of these are the sugar and carbon dioxide receptor subfamilies, which are distantly related to each other. This thrips has a considerable expansion of genes in each of these two subfamilies, with 18 candidate sugar receptors and 30 members of the carbon dioxide receptor subfamily. The sugar receptors are characterized by a glutamic acid (E) after the TY in the conserved TYhhhhhQF motif of the transmembrane 7 domain replacing a normally hydrophobic amino acid (h in the motif) (Kent and Robertson 2009). This position, by inference from the three-dimensional structure of Orco (Butterwick et al. 2018), is alongside the ion channel of the receptor tetramer. Of the 18 FoccGrs in the sugar receptor subfamily 16 have this TYE, the two exceptions being TYA and TYI in Gr37/38, respectively. It is unclear how this expansion of sugar receptors, compared for example with two in honey bees (Jung et al. 2015), 8 in *D. melanogaster* (Robertson et al. 2003), 8-13 in mosquitoes (Kent and Robertson 2009), and 16 in the flour beetle *Tribolium castaneum* (Richards et al. 2008), might be involved in their utilization of flowers as host plants, in part because we have yet to fully understand how the 8 Drosophila sugar receptors are deployed to sense diverse sugars (Fujii et al. 2015).

The large expansion of 30 genes in the carbon dioxide receptor subfamily is comparable to a similar expansion of this subfamily in the dampwood termite *Zootermopsis nevadensis* (Terrapon et al. 2014) and the German cockroach *Blattella germanica* (Robertson et al. 2018), but not all are expected to be involved in perception of this gas. FoccGr1-3 are most closely related to the Gr1-3 lineage of carbon dioxide receptors in endopterygote insects (Robertson and Kent 2009) (**Figure S4.2**), and so might indeed detect this gas, while the others presumably represent the larger subfamily from which the carbon dioxide receptors evolved, a subfamily that is at least as old as odonates (Ioannidis et al. 2017).

A distinct lineage of GRs has evolved to detect the sugar fructose, exemplified by the Gr43a protein in *D. melanogaster* (Miyamoto et al. 2012). Like most insects, this lineage is represented in other hemipteroids by a single gene, however this thrips has five genes (Gr49-53), while the lineage is expanded up to 10 genes in *T. castaneum* (Richards et al. 2008). This gene lineage evolved from within a far larger evolutionary assemblage of GRs, most of which in *D. melanogaster* are implicated in detecting “bitter” compounds, typically from plants (Weiss et al. 2011).

The remaining 49 GRs in this thrips are highly divergent from the other hemipteroid GRs, perhaps consistent with a similar role of detecting “bitter” plant defensive compounds. They form three clades in the phylogenetic analysis (**Figure S4.2**), the largest consisting of 40 genes. The latter includes a recent expansion of GR54-67. This is not quite a complete catalog of the GR family in this genome, because despite successfully repairing the assembly for 22 genes, 23 genes remain incomplete models with exons missing in gaps, while two are clear pseudogenes. At least 8 more fragments that might well represent intact genes in the genome were detected, but not named and analyzed as they could not be built into reasonable length models.

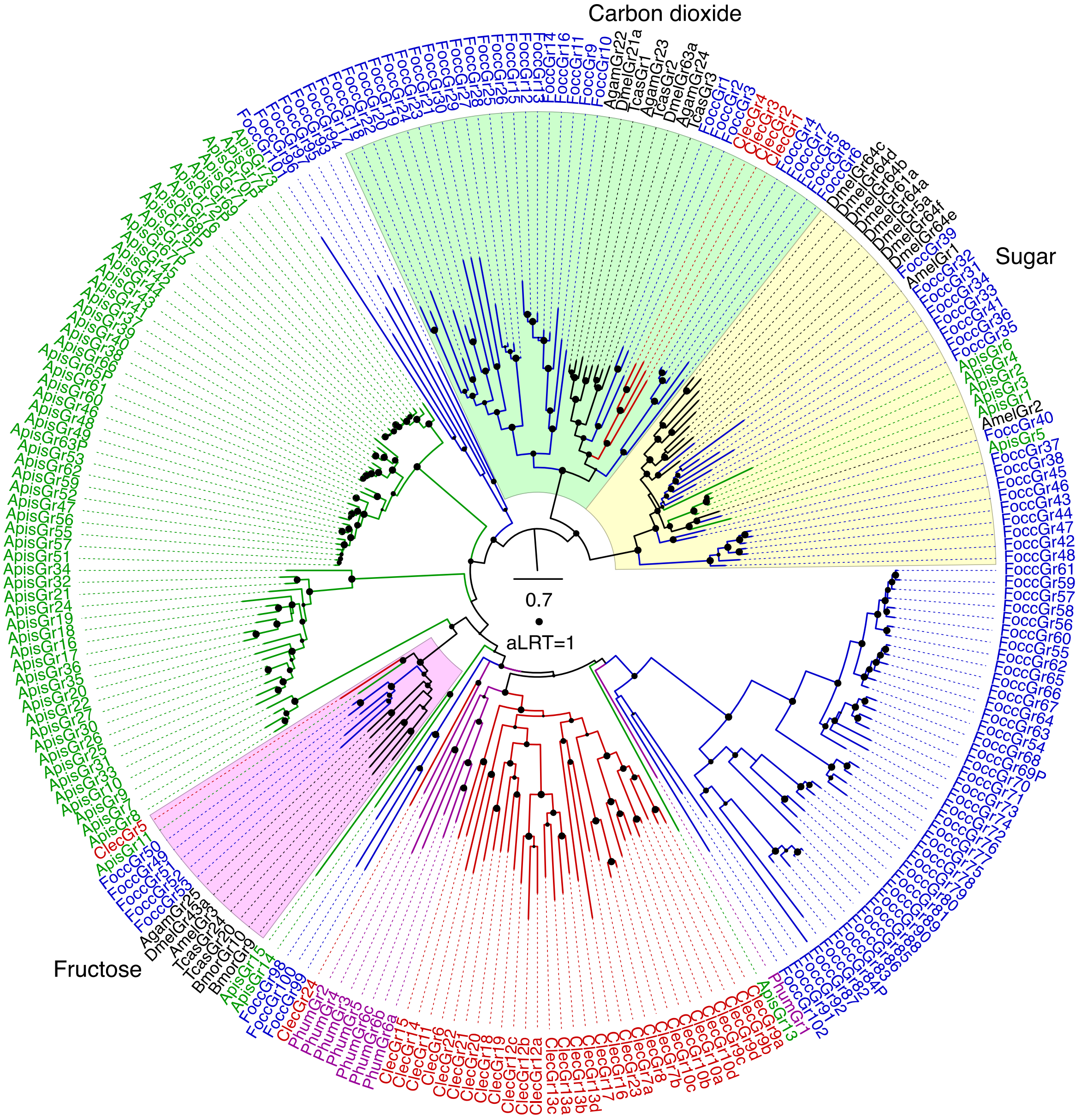

**Figure S4.2. Phylogenetic relationships of the gustatory receptors of hemipteroid insects.** The sugar and carbon dioxide receptor subfamilies were together declared the outgroup to root the tree, based on their position in phylogenetic analysis of the GR family throughout animals (Robertson 2015). The three major subfamilies are highlighted with background colors. Other details as for Figure S4.1. The list of FOCC OGS v1.0 models for GR genes are found in **Additional file 2: Table S7.**

##### 4.2.4. The IR family

The ionotropic receptor family is a divergent lineage of the ancient family of ionotropic glutamate receptors (Benton et al. 2009) and evolved in protostomes (Croset et al. 2010; Eyun et al. 2017). The family has been reviewed recently (Rytz et al. 2015; Rimal and Lee 2018). In most insects there are two highly conserved co-receptors, named Ir8a and 25a for their *D. melanogaster* orthologs, with lengths and sequences very similar to the glutamate receptors. A third co-receptor is the far shorter Ir76b protein. In addition, *D. melanogaster* has four proteins expressed in antennae that mediate perception of temperature and humidity in conjunction with some of the coreceptors, Ir21a, 40a, 68a, and 93a (Knecht et al. 2016). Most insects have single orthologs for each of these seven genes, and this thrips is no exception (**Figure S3.3**). In addition there is a clade of receptors known as the Ir75 clade with seven members in *D. melanogaster*, most of which are involved in perception of acids and amines (Prieto-Godino et al. 2019), that is also commonly expanded in other insects, usually independently of the fly expansions. *F. occidentalis* has 6 members of this clade, all independently duplicated not only from the fly genes, but from the relatives in other hemipteroids (Figure S3). As described above, just three of these 14 genes were intact in the genome assembly, the remainder requiring repairs to the assembly to encode full-length proteins. Many insects including more basal lineages like termites (Terrapon et al. 2014) and cockroaches (Robertson et al. 2018) also have relatives of the Ir41a and related proteins in *D. melanogaster*, however this thrips does not.

The remaining IRs in *D. melanogaster* form a divergent grouping, most of which fall into a large clade called the IR20a clade, and these have been implicated in gustation in both larvae and adults (Koh et al. 2014; Stewart et al. 2015; Sanchez-Alcaniz et al. 2018). In other insects these divergent IRs typically fall into two groups, a group with several introns and an “intronless” clade, and post Croset et al. (2010), these genes have been given numbers from 101 upwards (e.g. Terrapon et al. 2014; Robertson et al. 2018) to avoid confusion with the *D. melanogaster* genes which are numbered up to 100a because they were named for their cytological locations (Benton et al. 2009). This thrips has just one of the multiple-intron genes, Ir101, compared with a handful in the other hemipteroids (whose sequences for *P. humanus* and *A. pisum* were updated in Terrapon et al. 2014). The remaining “intronless” IRs form a greatly expanded thrips-specific clade of at least 167 genes, many of which are partial models with parts missing in assembly gaps, and there are many more small fragments in the assembly that might represent intact genes in the genome. Eight of these are clearly pseudogenic. This expansion is comparable to one of 93 “intronless” genes in *Z. termopsis* (Terrapon et al. 2014), and 755 in *B. germanica* (Robertson et al. 2018). A comparable large expansion of IRs was found in the deer tick *Ixodes scapularis* (Josek et al. 2018). A few of these genes do have one and sometimes two introns, however these have been idiosyncratically gained at different locations after the expansion of the clade from an intronless ancestor. By analogy with *Drosophila* flies, these IRs are likely to function in gustation, and like the divergent GRs, might be involved in perception of diverse host plant chemicals.

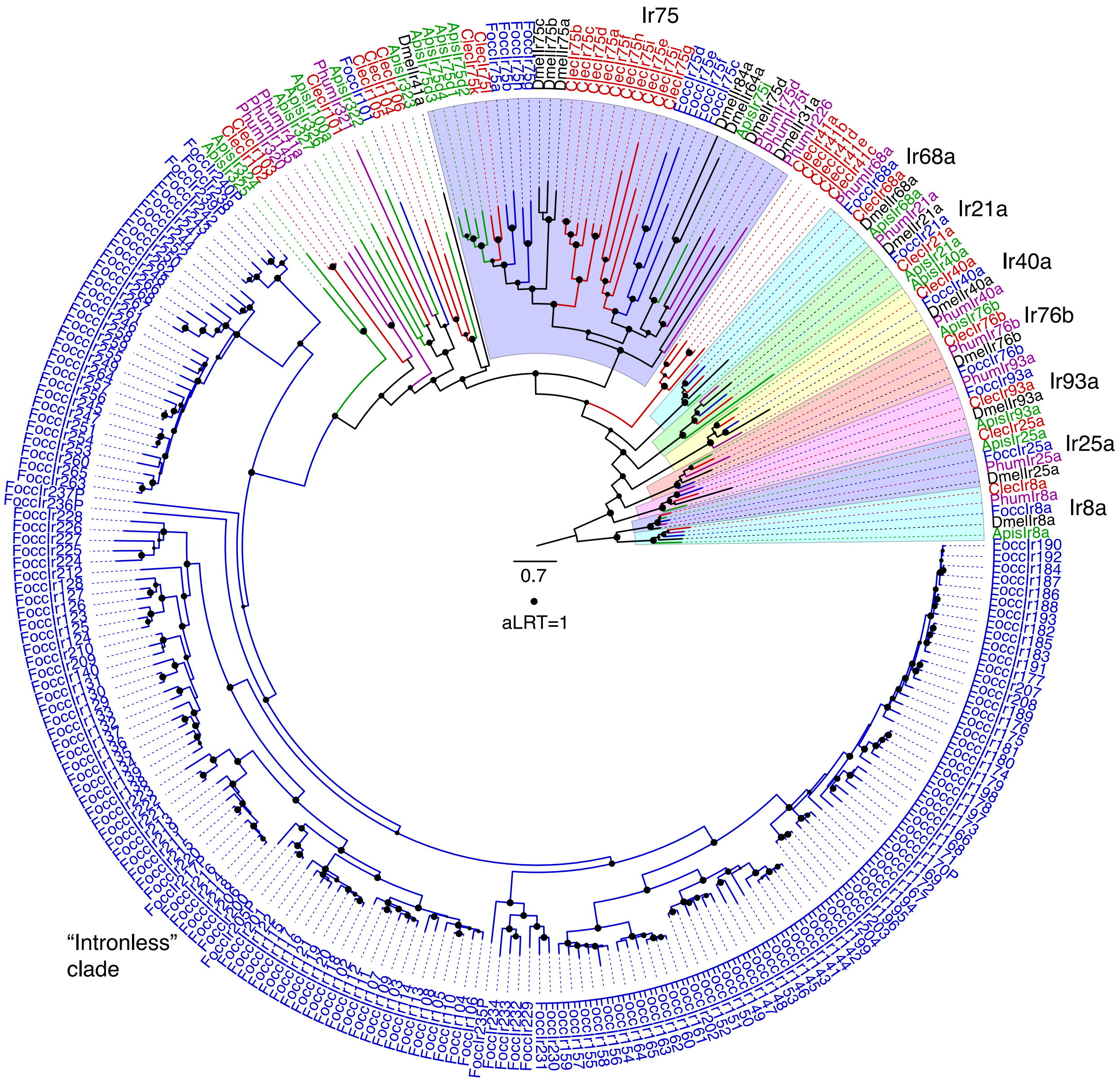

**Figure S4.3. Phylogenetic relationships of the ionotropic receptors of hemipteroid insects.** The Ir8a and 25a lineages were together declared the outgroup to root the tree, based on their close relationship to the ancestral glutamate receptors (Croset et al. 2010; Terrapon et al. 2014; Eyun et al. 2017). The major conserved lineages are highlighted with background colors. Other details are similar to Figure S4.1. The list of FOCC OGS v1.0 models for IR genes are found in **Additional file 2: Table S7.**

### 5. Vision genes

*Contributed by Markus Friedrich and Jeffery W. Jones*

#### 5.1 Background

Vision supports many aspects of insect biology and is mediated by a variety of light-sensing receptor proteins. In thrips, vision has been shown to be involved in plant host finding. This capacity is most likely mediated by homologs of the opsin gene family, expressed in the photoreceptors of the moderately sized pair of compound eyes in thrips. Opsins constitute a deeply conserved class of light-sensitive protein G-protein coupled receptors. Three major subfamilies have been identified across the animal kingdom: Ciliary opsins (c-opsins), rhabdomeric opsins (r-opsins) and RGR/Go opsins (Cronin & Porter, 2014). R-opsins have most extensively diversified in insects (Cronin & Porter, 2014; Feuda, Marlétaz, Bentley, & Holland, 2016; Hering et al., 2012), resulting in paralog groups with differential wavelength sensitivity maxima. This includes the long wavelength-sensitive subfamily (LWS-opsin), a blue or short wavelength sensitive opsin (SWS-B-opsin), and the ultraviolet-short wavelength sensitive opsin (SWS-UV-opsin), all of which are mainly expressed in the photoreceptors of the peripheral visual system, i.e. the compound eyes and the median eyes. In addition, the r-opsin subfamily also includes paralog groups whose members have been found to be predominantly expressed in non-retinal tissues: Rh7 opsins and Arthropsins (Eriksson, Fredman, Steiner, & Schmid, 2013; Ni, Baik, Holmes, & Montell, 2017). In some species, the extraretinal r-opsins are complemented by the presence of homologs of the c-opsin subfamily in insects, which are likewise expressed in non-retinal tissues (Velarde, Sauer, O. Walden, Fahrbach, & Robertson, 2005). Further light-sensitive proteins expressed in non-retinal tissues includes cryptochromes and photolyases (Porter, 2016).

#### 5.2 Methods

We searched the *F. occidentalis* genome draft in the i5k Workspace by tBLASTn using the protein sequences of characterized light sensitive genes from *Drosophila* and the red flour beetle *Tribolium castaneum* as queries (Altschul, Gish, Miller, Myers, & Lipman, 1990; Poelchau et al., 2014). For *Drosophila*, this included (6-4)-photolyase (phr6-4: CG2488), cryptochrome (cry: CG3772), Rhodopsin-1 (Rh1: CG4550), Rhodopsin-2 (Rh2: CG16740), Rhodopsin-3 (Rh3: CG10888), Rhodopsin-5 (Rh5: CG5279), Rhodopsin-6 (Rh6: CG5192), and Rhodopsin-7 (Rh7: CG5638). For *Tribolium*, this included cryptochrome 2 (cry2: XP_008200421) and ciliary opsin (c-opsin: XP_001816446). Orthology relationships of candidate homologs for scrutinized by reciprocal BLAST (Wall, Fraser, & Hirsh, 2003). For members of the opsin gene family, subfamily relationships were explored by gene tree reconstruction. Protein sequences were aligned with Clustal Omega (Sievers et al., 2011). Ambiguous alignment regions were filtered using Gblocks applying least stringent setting (Castresana, 2000). Bootstrapped maximum likelihood tree was estimated with RAxML as implemented on the CIPRES platform (Miller, Pfeiffer, & Schwartz, 2010; Stamatakis, 2014).

#### 5.3 Results and Discussion

Our searches detected seven opsin genes, two cryptochromes, and a singleton homolog of phr6-4 in the *F. occidentalis* genome (**Additional File 2: Table S8**). The opsin gene homologs represented five homologs of subfamilies expressed in the peripheral visual system, which included singleton homologs of each the UV- and B-opsin subfamily, complementing three tandem-duplicated LW opsins on scaffold 18. Gene tree analysis revealed that the *F. occidentalis* LW opsin cluster represents an independent expansion of LW opsins in relation to the LW opsin clusters found in hemipteran species (**Fig. S5.1**) (Armisen et al., 2018; Sparks et al, *in review*; Panfilio et al., 2019). In addition to these opsins that are most likely expressed in the peripheral visual system, we detected singleton homologs of c-opsin (Velarde et al., 2005) and the Rh7 opsin (Ni et al., 2017). We failed to detect sequence conservation evidence for Arthropsins in *F. occidentalis* (Eriksson et al., 2013), although this opsin gene family has been found in a variety of hemipteran species (Armisen et al., 2018; International Aphid Genomics Consortium, 2010; Panfilio et al., 2019).

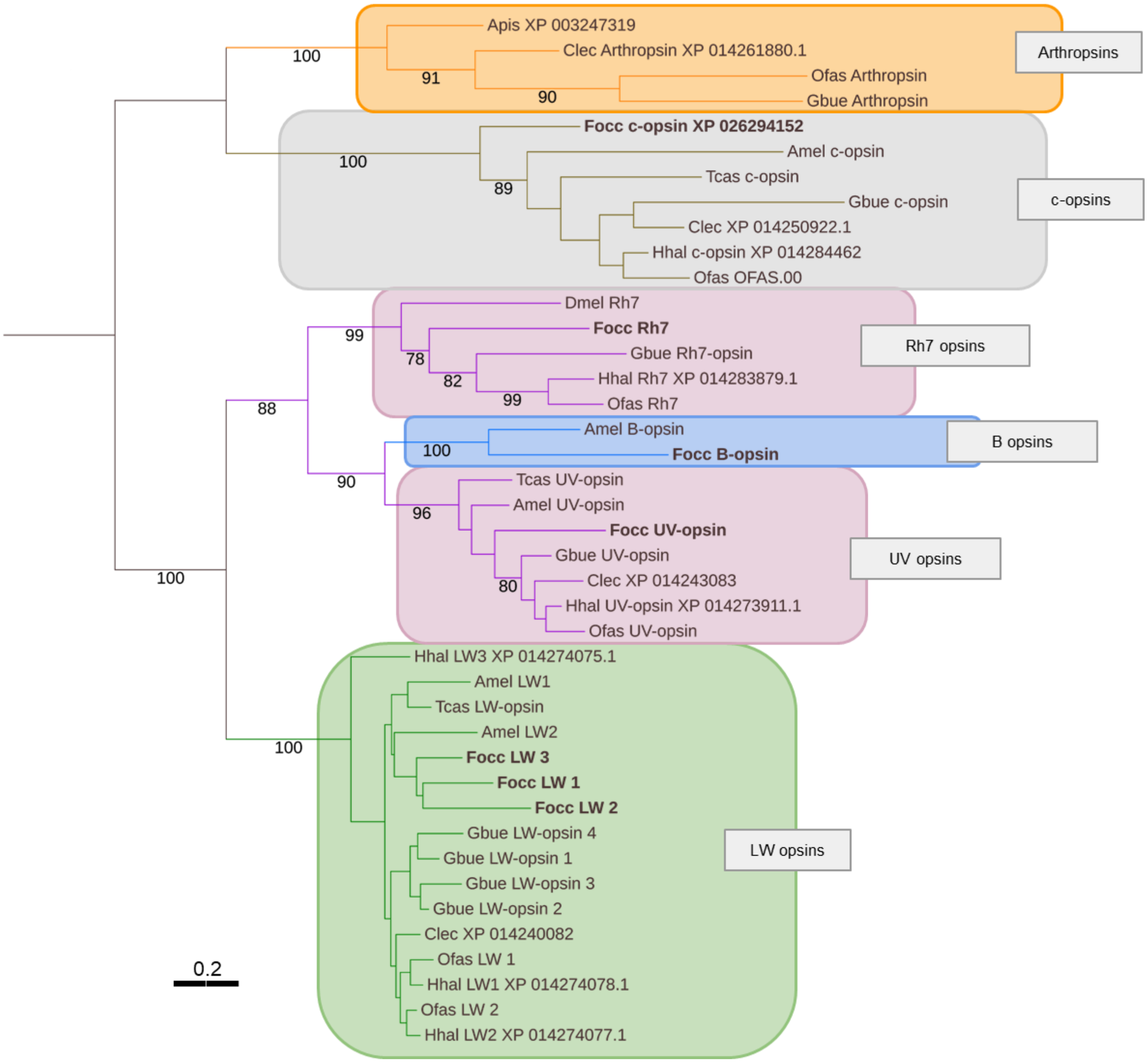

**Figure S5.1: Nonparametric bootstrap maximum likelihood tree of opsin genes from hemipteran and other arthropod species.** Species abbreviations: Amel = *Apis mellifera*, Apisum = *Acyrthosiphon pisum*, Clec = *Cimex lectularius*, Dmel = *Drosophila melanogaster*, Focc = *Frankliniella occidentalis*, Gbue = *Gerris buenoi*, Hhal = *Halyomorpha halys*, Ofas = *Oncopeltus fasciatus*, Tcas = *Tribolium castaneum*. Sequences and alignment available on request. Numbers at branches represent non-parametric bootstrap support higher than 75. Scale bar corresponds to 0.2 substitutions per site.

### 6. Validation of salivary gland genes

*Contributed by Sulley Ben-Mahmoud, Joshua Benoit, Dorith Rotenberg and Diane Ullman*

#### 6.1. Abstract

The salivary gland (SG) of the Western flower thrips (WFT), *Franklinella occidentalis* (Pergande) is critical to the insect's competence in transmitting viruses in the genus *Tospovirus,* including the type member, *Tomato spotted wilt virus* (TSWV). Sequencing of the WFT genome, provides an exciting opportunity to reveal, at the molecular and proteomic level, how salivary gland components function in insect-plant and insect-virus interactions, including identification of effectors and proteins involved in virus replication, retention and transmission. Comparison of RNAseq data from WFT salivary glands and whole bodies identified 123 genes expected to be enriched in the salivary gland (**Additional file 2: S11**; **see this supplement, section 1.5**). We present evidence validating that four of these genes, FOCC009158 (F9158), FOCC007547 (F7547), FOCC003700 (F3700), and a pancreatic triacylglycerol lipase-like gene FOCC003652 (F3652), are enriched in the SG of adult female WFT, relative to the insect head or body. These genes are predicted to be secretory proteins, and F3652 is likely to function as a plant membrane digestive enzyme. These findings are an important step towards elucidating organ-specific functions of genes in the WFT.

#### 6.2. Background

*Frankliniella occidentalis* (Pergande), commonly known as the Western flower thrips (WFT) is a very important pest of a large number of economically important crops worldwide, including tomatoes, peanuts, lettuce, peppers and tobacco. Sequencing the WFT genome opened several multiple avenues of research, including the insect's role as a vector of *Tomato spotted wilt virus* (TSWV), at the sub-organismal level. The thrips have primary and tubular salivary glands (PSG and TSG, respectively) that play key roles in virus replication, movement within the insect and competence to inoculate the virus into a plant (Montero-Astua et al., 2016). The TSG play a role in virus movement from the midgut to the PSG. Furthermore, infection of the PSG is required for inoculation to occur. Hence, understanding gene expression in the salivary glands is highly relevant to a mechanistic understanding of virus replication and inoculation.

In 2013, the WFT sialotranscriptome (Stafford-Banks et al., 2014) became available, providing a foundational tool to understand the interaction of WFT with plants hosts and TSWV. Via a comparative analysis of RNAseq reads from the sialotransciptome, a whole body transcriptome and WFT genome sequence databases, we were provided with a select list of candidate transcripts (N=38) that were enriched in the SGs of male and female adults. RNAseq reads may not always map to a single gene (Li and Dewey, 2011) and given this limitation, it is currently still necessary to validate the abundance of these genes using a different technique. We selected qRT-PCR for this purpose. The verification of such enriched and/or differentially expressed genes, takes us closer to gaining a better understanding of plant-virus-vector interactions, and importantly yield potential targets for novel strategies to manage crop losses due the TSWV-transmission by WFTs.

#### 6.3. Methods

##### 6.3.1 Thrips rearing and salivary gland dissections

*Frankliniella occidentalis* adult females were obtained from a colony originally collected from Hawaii, and maintained on green bean pods (Ullman et al., 1992) at the Ullman laboratory at University of California, Davis, California. Thrips were age-regulated to ensure that only adult females approximately 48 hours, post-eclosion were dissected into the following treatment groups: head, body, and salivary glands (PSG and TSG combined). Thrips salivary glands were surgically removed as previously described (Stafford-Banks et al., 2014). All thrips dissections were conducted at the same time of day in a consistent 2-hour window and had access to green bean pods up until they were selected to be dissected. This was done to minimize potential variations due to their circadian rhythms or nutritional status. To ensure that enough total RNA was obtained for subsequent qRT-PCR assays, tissues from 5 individual thrips were pooled into the three treatment groups. For each respective replicate, the salivary gland, head, and carcass were always obtained from the same female individuals.

##### 6.3.2. Total RNA isolation and cDNA synthesis

Total RNA, eluted in 12 µL ultra-pure water, was extracted from thrips tissues using the Arcturus® PicoPure ® RNA isolation kit (Life Technologies, USA) by following the manufacturers recommendations. The yield, and purity of extractions was determined using the Nanodrop ND-1000 (Thermo Scientific, DE, USA). For qRT-PCR, 2 ng of total RNA was used as template to synthesize first strand cDNA (10 µL reaction volume; 45◦C, 60 min, followed by a denaturation step at 85◦C for 10 minutes) with 0.5 µM gene-specific primers (see **Table S5.1**) to amplify respective targets in single reactions with the Verso™ cDNA synthesis Kit (Thermo Scientific, DE, USA). The cDNA synthesis kit comes with a reagent to remove DNA contamination. Nonetheless, during the optimization of qRT-PCRs, controls were included to ensure that possible contribution of DNA contamination to Cq values was negligible. Quantitative RT-PCR reactions were optimized to select only primer pairs that demonstrated high PCR efficiencies over a wide linear dynamic range.

##### 6.3.3. Quantitative RT-PCR of Salivary Gland Enriched Genes

Quantitative RT-PCRs were performed on a CFX96™ real-time PCR detection system (Bio-Rad Laboratories, CA) using their SsoAdvanced™ Universal SYBR® Green Supermix. Reaction mixtures consisted of 0.2 µM of forward and reverse gene-specific primers, 2 µL first strand cDNA, in a final reaction volume of 10 µL. Reaction conditions were as follows: 95◦C for 30 sec, 40 cycles of 95◦C for 10 sec, 55◦C for 10 sec, and 60◦C for 25 sec, concluded with melt curve analysis to verify the Tm product. Each reaction was run in three technical replicates, and each plate was run with two putative SG-enriched genes, alongside the actin (Act) and cytochrome oxidase 1 (COX) reference genes. Primers to the actin reference gene were obtained from Boonham et al., (2002), whereas COX primers were designed to amplify a 80 bp sequence of the cytochrome oxidase subunit I gene, GenBank: gi|355398713 (Wang et al., 2014). Additionally, all three treatment groups of each replicate were run on the same plate to avoid plate to plate differences. The abundance of SG enriched transcripts were generated the $2^{-\Delta Cq}$ method (Livak and Schmittgen, 2001), normalized to the geometric mean of Act/COX. Raw transcript abundance were log transformed prior to 1-way ANOVA followed by Tukey's multiple comparison tests with the GraphPad Prism software.

**Table S6.1. Quantitative RT-PCR Primers: gene, annotation, sequence, amplification efficiency (E), and amplicon size of qRT-PCR product for the Transcript Abundance of SG enriched genes.**

| Gene | Maker Name | Annotation | Primer ID | Sequence (5' - 3') | E (%) | qRT-PCR Amplicon (bp) |
|---|---|---|---|---|---|---|
| Actin | F0CC008191 | FoccTmpA008191-RA | FoccAct1_f | GGTATCGTCCTGGACTCTGGTG | 98 | 69 |
| | | | FoccAct1_r | GGGAAGGGCGTAACCTTCA | | |
| Cytochrome Oxidase subunit 1 | FOCC005842 | FoccTmpM005842-RA | FocCOX_f | CGTTACCAGTTTTAGCAGGAG | 102 | 80 |
| | | | FocCOX_r | TCCTCTCGGATCAAAGAAGG | | |
| F9158 | FOCC009158 | Putative SG Protein 21 | Foc26Srs3_f | CCACTGAAGACCTGACTGAT | 107 | 109 |
| | | | Foc26Srs3_r | ATAACGTGTTCCTTGGGAGT | | |
| F7547 | FOCC007547 | Putative SG Protein 22 | Foc7547_f | GAACTGTGACCATGTCTGTG | 102 | 80 |
| | | | Foc7547_r | CTTGGTGGTGATGTTCTTGG | | |
| F3700 | FOCC003700 | Putative SG Protein 23 | Foc3700b_f | AGAGGAAAAGAAGGACAGCG | 103 | 80 |
| | | | Foc3700b_r | CAGAAGAGAGGGGATTGCTG | | |
| F3652 | FOCC003652 | Pancreatic triacylglycerol lipase-like | Foc3652a_f | CGGCGGCATATTAGGAATTG | 100 | 148 |
| | | | Foc3652a_r | CTCAATGGCGTTGATGTACG | | |

##### 6.4. Results and Discussion

Examining the differences between the mean transcript abundance of respective SG enriched genes, F9158, F7547, F3700 and F3652, between the head, body, and salivary gland showed that F3700 was only reliably detected in the SG, but not the head or the body. For two of the remaining three SG enriched genes: F9158 and F3652, the transcript abundance in the head, and body were not statistically different (**Figure S6.1A and S6.1D**). Furthermore, the transcript levels in the salivary gland were significantly higher in the salivary gland ($P < 0.001$). F7547 was differentially expressed in the head, body, and salivary gland in increasing order ($P < 0.05$).

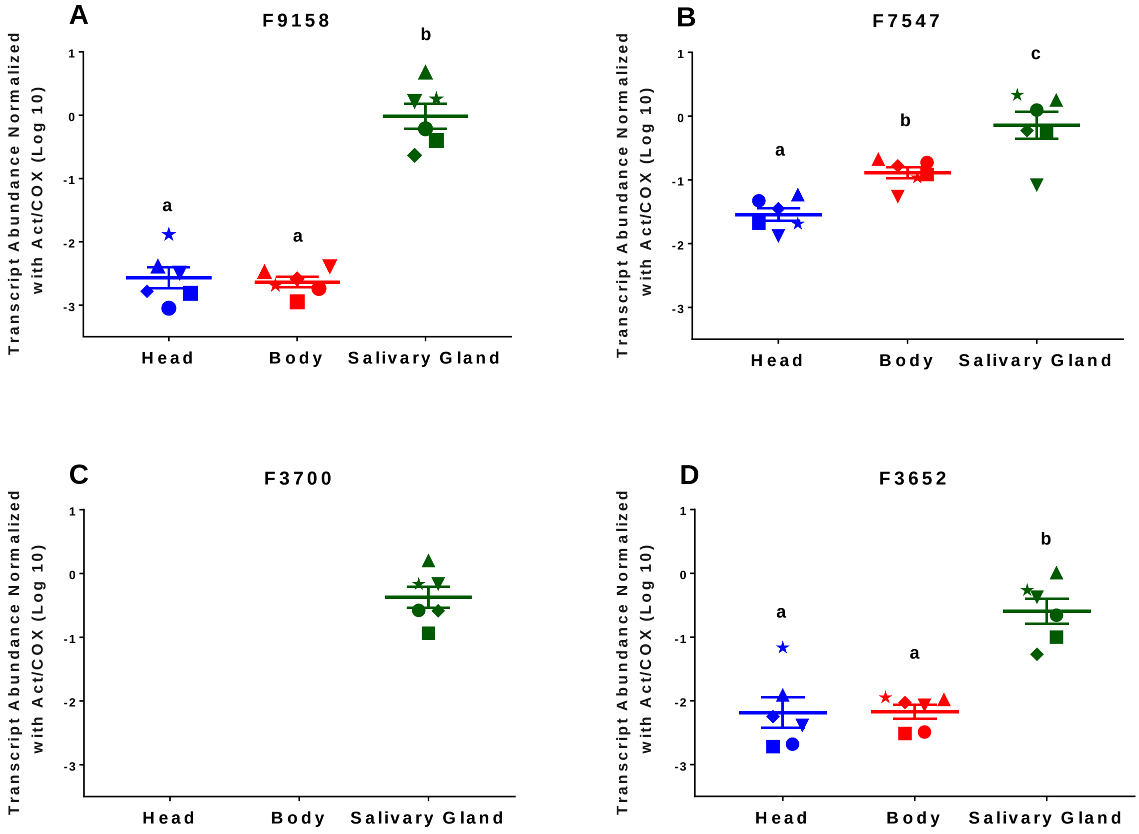

**Fig. S6.1. Transcript abundance of SG enriched genes in the head, body and salivary glands of female adult *Frankliniella occidentalis.*** The Transcript Abundance (normalized with actin (Act) and cytochrome oxidase I (COX)) of SG enriched genes with maker model ID: FOCC009158 (F9158), FOCC007547 (F7547), FOCC003700 (F3700), and FOCC003652 (F3652) was determined for six replicates each of total RNA extracted from the head, body and salivary glands of pools of five 48-hour old female adult WFTs. Symbols (replicates) of the same shape for all SG enriched gene represent the same group of 5 female adults. Lines represent the mean of the six replicates ± the standard error of the mean (SEM). One-way ANOVA: Tukey's multiple comparison tests, revealed significant differences in the abundance in the SG relative to the head, and body, in all four genes.

These data support our comparative analysis of the sialotransciptome, whole body transcriptome and WFT genome sequence databases, providing strong additional evidence that F9158, F7547, F3700 and F3652 are enriched, or more highly expressed in the salivary gland than other parts of the thrips body. For every respective biological replicate (pool of 5 female adults), and for every gene quantified in this study, the transcript abundance value (normalized against actin and cytochrome oxidase) was highest in the salivary gland. In the case of F3700, there was negligible expression in the head, and the body as the qRT-PCR primers designed to measure this gene failed amplify the expected product in these treatment groups. To control for possible starvation effects, we gave thrips access to green bean pods *ad libitum*, but we were not certain if each individual was fully fed before it was dissected.

Of the candidate 38 SG-enriched genes, at least 10 of these genes may be secreted proteins using the publicly available predictive tools: SignalP4.1, Phobius, and MultiLoc (Käll et al., 2007; Petersen et al., 2011; Pierre et al., 2006). We currently do not know the function of the proteins encoded by F9158, F7547 and F3700 in the salivary glands. It is remarkable that the protein prediction tools above suggest that they could be secreted proteins as they have signal peptides at the N-terminal region of their sequences. Thus, it is possible that they may act as plant effectors. Based on the prediction with Phobius, F3652, which shares the highest similarity to the pancreatic triacylglycerol lipases, is possibly a transmembrane protein, even though the other predictors suggest the presence of a secretory signal peptide. Interestingly, a pancreatic-lipase was also found in the SG of the mosquito *An. Stephensi* (Valenzuela et al., 2003) although its exact role is not known. Notably, this mosquito species is known to feed on plant-sugars. Members of the pancreatic lipase gene family are predicted to hydrolyze galactolipids (Aoki et al., 2007) which make up the bulk of photosynthetic membranes (Dörmann, 2013), and may help the insects to access plant sugars. F3652 may play a similar role for the thrips as it feeds on plants.

This study makes a strong case for future candidate-gene studies. The SG-enriched genes may enable entry and egress of TSWV components into/out of the SG. Furthermore, the secreted proteins may function to help the WFT circumvent plant defenses, and TSWV to colonize plant tissues (Hogenhout et al., 2009). There are other candidate genes to be pursued made possible by the sequencing of the WFTs genome. High-throughput screening methods need to be developed to suit the WFT, but we are closer to that possibility now than ever before.

#### Appendix 1

**F9158 Transcript Abundance normalized with Act/COX**

| Rep | Head | Body | Salivary Gland |
|---|---|---|---|
| 09 | 0.000894 | 0.001830 | 0.612203 |
| 10 | 0.001534 | 0.001131 | 0.399703 |
| 11 | 0.004130 | 0.003380 | 4.792270 |
| 15 | 0.001650 | 0.002600 | 0.232090 |
| 13 | 0.003190 | 0.004040 | 1.650770 |
| 16 | 0.013020 | 0.002050 | 1.808580 |

**F7547 Transcript Abundance normalized with Act/COX**

| Rep | Head | Body | Salivary Gland |
|---|---|---|---|
| 09 | 0.046940 | 0.187340 | 1.248440 |
| 10 | 0.020970 | 0.121480 | 0.565910 |
| 11 | 0.058550 | 0.212110 | 1.785100 |
| 15 | 0.035260 | 0.165920 | 0.592280 |
| 13 | 0.013220 | 0.054850 | 0.082880 |
| 16 | 0.020430 | 0.108540 | 2.145150 |

#### F3700 Transcript Abundance normalized with Act/COX

| Rep | Head | Body | Salivary Gland |
|---|---|---|---|
| 09 | 0.000000 | 0.000000 | 0.263549 |
| 10 | 0.000000 | 0.000000 | 0.115600 |
| 11 | 0.000000 | 0.000000 | 1.599431 |
| 15 | 0.000000 | 0.000000 | 0.260358 |
| 13 | 0.000000 | 0.000000 | 0.685641 |
| 16 | 0.000000 | 0.000000 | 0.681229 |

#### F3652 Transcript Abundance normalized with Act/COX

| Rep | Head | Body | Salivary Gland |
|---|---|---|---|
| 09 | 0.046940 | 0.187340 | 1.248440 |
| 10 | 0.020970 | 0.121480 | 0.565910 |
| 11 | 0.058550 | 0.212110 | 1.785100 |
| 15 | 0.035260 | 0.165920 | 0.592280 |
| 13 | 0.013220 | 0.054850 | 0.082880 |
| 16 | 0.020430 | 0.108540 | 2.145150 |

### 7. Detoxification genes

#### 7.1. Cytochrome P450s

*Contributed by Jonathan Oliver, Derek Schneweis, Dorith Rotenberg and Anna Whitfield*

##### 7.1.1. Abstract

Cytochrome P450s (P450s, CYPs) are a large superfamily of enzymes. P450 enzymes have been identified in all domains of life where they are involved in the metabolism of multiple substrates with prominent roles in hormone synthesis and breakdown, development, and detoxification (Feyereisen 1999; Heidel-Fischer and Vogel 2015). In agricultural systems, *F. occidentalis* has shown a propensity for developing resistance to insecticides commonly utilized to manage this species, and P450s have been specifically implicated in the detoxification of insecticides by *F. occidentalis* (Cifuentes et al. 2012; Yan et al. 2015). Within the *F. occidentalis* genome, a relatively large number of P450s were identified, including numerous members of CYP families frequently associated with the breakdown of toxic plant products and insecticides (Cifuentes et al. 2012).

##### 7.1.2. Results and Discussion

P450s in the genome of *F. occidentalis* were identified based upon similarity to known P450s in other insect species. Initially, 130 P450 gene models were annotated across 88 different scaffolds (**Additional file 2: Table S13**), with clustering of P450 genes on some scaffolds as has been noted to occur in other insect genomes including *D. melanogaster* and *T. castaneum* (Zhu et al. 2013; Chung et al. 2009). Based upon alignments with other insect protein orthologs and *de novo* transcriptome evidence from RNAseq (Schneweis, 2017, Phd Thesis), these 130 CYP gene models were determined to represent at least 89 unique CYP gene sequences (some fragmented across different scaffolds) as well as 23 additional partial sequences. P450 genes were assigned to families and named on the basis of overall amino acid sequence identity using the blastp function – with P450s showing greater than 40% amino acid identity receiving the same number designation (CYP family) and greater than 55% identity receiving the same letter designation (CYP subfamily) (Nelson et al. 1998) through comparisons with annotated CYPs from the genomes of other insects (including *Drosophila melanogaster*, *Pediculus humanus corporis*, *Tribolium castaneum*, *Acyrthosiphon pisum*, *Cimex lectularius*, *Zootermopsis nevadensis*, and *Diaphorina citri*). Characterization of the unique P450s from *F. occidentalis* indicated that representatives of at least 24 different CYP families are present in the genome (**Table S7.1**). Overall, more than 40% of the total number of CYP genes annotated in the *F. occidentalis* genome were assigned to the CYP4 and CYP6 gene families. Phylogenetic characterization of *F. occidentalis* P450s versus those found in other insect species has been completed (**Figure S7.1**), and phylogenetic analysis suggested expansion within the CYP 3 and CYP 4 clans of *F. occidentalis*. The majority of annotated *F. occidentalis* P450s showed relatively low identity to other insect P450s. This is in agreement with the findings of Scott and Wen (2001) that the majority of P450s in insect genomes show very limited amino acid identity (30-50%) to P450 genes in other insect species.

All told, a diverse array of P450s were annotated within the *F. occidentalis* genome. Given the already described importance of P450s in insecticide resistance (Cifuentes et al. 2012; Yan et al. 2015), the importance of insecticides in the management of thrips species (Cifuentes et al. 2012), and the multitude of plant defense compounds encountered during the thrips' phytophagous lifestyle (Heidel-Fischer and Vogel 2015), knowledge of the diversity of P450s present within the *F. occidentalis* genome is likely essential for optimizing management of this important agricultural pest. The annotation of these P450 genes will enable future functional studies in *F. occidentalis* related to the detoxification of insecticidal and plant defense compounds.

**Table S7.1** Family assignments for annotated *F. occidentalis* P450 genes.

| Clan | Family | Total | Complete | Partial |
|---|---|---|---|---|
| 2 | CYP15 | 3 | 3 | 0 |
| 2 | CYP18 | 1 | 1 | 0 |
| 2 | CYP303 | 1 | 1 | 0 |
| 2 | CYP304 | 3 | 3 | 0 |
| 2 | CYP305 | 1 | 1 | 0 |
| 2 | CYP306 | 1 | 1 | 0 |
| 2 | CYP307 | 2 | 2 | 0 |
| 3 | CYP6 | 26 | 20 | 6 |
| 3 | CYP3652 | 1 | 1 | 0 |
| 3 | CYP3653 | 2 | 2 | 0 |
| 3 | CYP3654 | 1 | 1 | 0 |
| 4 | CYP4 | 20 | 16 | 4 |
| 4 | CYP3655 | 15 | 11 | 4 |
| 4 | CYP3656 | 2 | 2 | 0 |
| 4 | CYP3657 | 9 | 8 | 1 |
| 4 | CYP3658 | 4 | 3 | 1 |
| 4 | CYP3659 | 2 | 1 | 1 |
| 4 | CYP3660 | 1 | 1 | 0 |
| 4 | CYP3661 | 3 | 0 | 3 |
| 4 | Unassigned | 1 | 0 | 1 |
| mitochondrial | CYP301 | 2 | 2 | 0 |
| mitochondrial | CYP302 | 2 | 2 | 0 |
| mitochondrial | CYP314 | 1 | 1 | 0 |
| mitochondrial | CYP315 | 1 | 1 | 0 |
| mitochondrial | CYP3118 | 7 | 5 | 2 |
| | **Totals** | **112** | **89** | **23** |

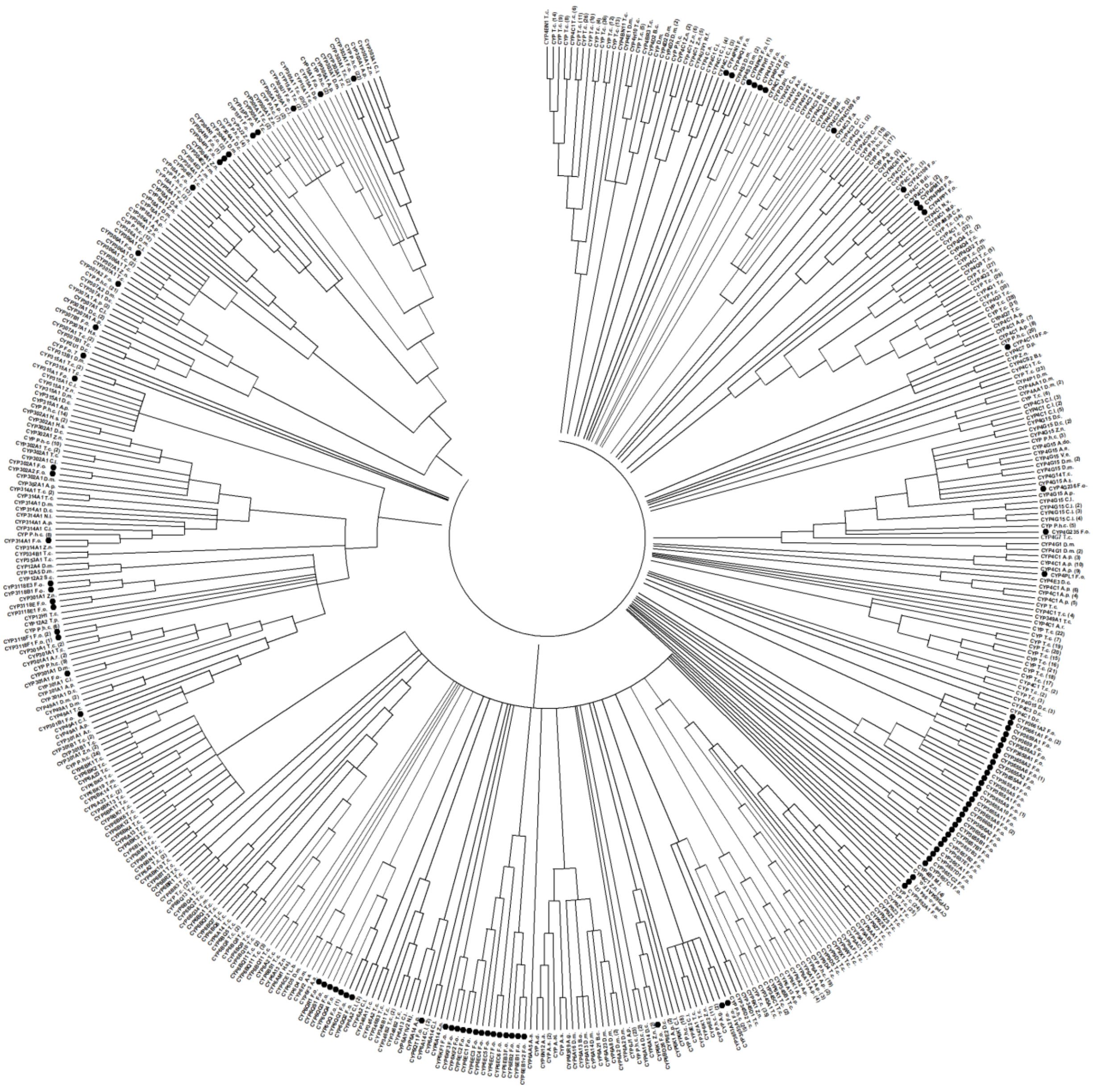

**Figure S7.1** Amino acid phylogenetic tree showing all *F. occidentalis* P450 genes alongside the most similar P450 sequences from other insect species found in Genbank ( https://www.ncbi.nlm.nih.gov/genbank/) as of 8-25-16. Black circles indicate the sequences from *F. occidentalis*. The evolutionary history was inferred using the Neighbor-Joining method (Saitou and Nei 1987). The bootstrap consensus tree inferred from 1000 replicates is taken to represent the evolutionary history of the taxa analyzed (Felsenstein 1985). Branches corresponding to partitions reproduced in less than 50% bootstrap replicates are collapsed. (Felsenstein 1985). The evolutionary distances were computed using the Poisson correction method (Zuckerkandl and Pauling 1965) and are in the units of the number of amino acid substitutions per site. The rate variation among sites was modeled with a gamma distribution (shape parameter = 5). The analysis involved 485 amino acid sequences All ambiguous positions were removed for each sequence pair. There were a total of 1421 positions in the final dataset. Evolutionary analyses were conducted in MEGA X (Kumar et al. 2018). The FOCC OGS v1.0 gene models and other insect species P450 Genbank accessions included in this tree are listed in **Additional file 2: Table S14**.

***Cytochrome P450s in development***

A steroid hormone, 20-hydroxyecdysone (20E), and juvenile hormone (JH) are known to play essential roles in the insect growth and development process, and the biosynthesis pathway for 20E includes several conserved P450s (Iga and Kataoka 2012). The P450 genes responsible for the synthesis of 20E include CYP307A1/A2, CYP306A1, CYP302A, CYP315A1, and CYP314A1. Given the importance of 20E in development, knockouts of these genes in *D. melanogaster* has been shown to produce striking phenotypes and has led these P450s to be called "spook/spookier", "phantom", "disembodied", "shadow", and "shade", respectively. In addition, CYP18A1 is known to be a key enzyme involved in the inactivation of 20E and is essential for metamorphosis in *D. melanogaster* (Guittard et al. 2011). In the *F. occidentalis* genome, corresponding homologs for each of these genes were identified (**Table S7.2**). Not unexpectedly, these evolutionarily conserved P450 genes showed some of the highest amino acid conservations observed among the P450s from the *F. occidentalis* genome versus P450s in the genomes of other sequenced insects; nonetheless, these identities were modest, ranging from 33-71%.

**Table S7.2.** P450 genes known to be involved in 20E biosynthesis or inactivation, their location within the *F. occidentalis* genome and their identity at the amino acid level versus orthologs in the genomes of other insect species.

| P450 gene (Dm name)[z] | Locations of *F. occidentalis* P450 20E pathway orthologs | Amino acid identity versus other insect species[zy] | | | | | | |
|---|---|---|---|---|---|---|---|---|
| | | *Phc* | *Ap* | *Cl* | *Dc* | *Zn* | *Tc* | *Dm* |
| CYP302a1 (Disembodied) | Scaffold54:583491-586227 | 45% | 45% | 41% | 45% | 47% | 42% | 45% |
| | Scaffold117:731027-733573 | 45% | 43% | 41% | 45% | 46% | 45% | 43% |
| CYP306a1 (Phantom) | Scaffold261:103212-126374 | 47% | 36% | 46% | 33% | 50% | 45% | 44% |
| CYP307a1/a2 (Spook/Spookier) | Scaffold2:2913680-2929983 | 59% | 51% | 45% | 52% | 60% | 61% | 49% |
| | Scaffold144:498693-502969 | 44% | 50% | 54% | 46% | 41% | 52% | 36% |
| CYP314a1 (Shade) | Scaffold820:12383-33648 | 63% | 58% | 59% | 58% | 67% | 44% | 48% |
| CYP315a1 (Shadow) | Scaffold156:439070-441178 | 42% | 35% | 38% | 36% | 49% | 44% | 37% |
| CYP18a1 | Scaffold261:97159-101980 | 71% | 55% | 59% | 58% | 69% | 66% | 60% |

[z]Organism used for comparisons are as follows: *Dm = D. melanogaster, Tc = T. castaneum, Phc = Pediculus humanus corporis, Ap = A. pisum, Dc = D. citri, Cl = C. lectularius, Zn = Z. nevadensis*

[y]The full *F. occidentalis* P450 protein sequence was compared to the most similar P450 protein sequence from each respective genome.

In addition to having important roles in the 20E pathway, CYPs are also known to be involved in other aspects of development. CYP301A1 is conserved within the genome of other insects and has been shown to play a key role in the formation of adult cuticle in *D. melanogaster* (Sztal et al. 2012). In *F. occidentalis*, two P450s (Scaffold74:870617-877535 and Scaffold74:876952-880140) (**Additional file 2: Table S13**) corresponding to CYP301A1 showed relatively high similarity (47-68%) to CYP301A1 orthologs in other insect genomes. It would be suspected that these CYPs may also be involved in thrips development in a similar manner to that observed in other insect species.

#### 7.2 ATP binding cassette and Carboxylesterase genes

*Contributed by Wannes Dermauw, Simon Snoeck, and Thomas Van Leeuwen*

##### 7.2.1. Methods

###### *7.2.1.1 ATP-binding Cassette Transporter (ABC) gene family*

The ABC genes of *F. occidentalis* were identified as previously described (Dermauw et al., 2013a; Sturm et al., 2009). Briefly, highly conserved nucleotide binding domains (NBD) of *D. melanogaster* ABC proteins were used as queries in tBLASTn searches (E-value threshold <$E^{-5}$) against the genome sequence assembly of *F. occidentalis* (Altschul et al., 1990). *F. occidentalis* ABC gene models were refined or created based on homology and available RNA-seq data. Full-length ABC genes and those genes with a sequence length larger than 70% of the average sequence length of full-length ABC genes per ABC subfamily, were considered as putative ABC genes. All *F. occidentalis* ABC gene sequences that have been annotated in this study can be found in **Additional file 2: Table S15**. Assignment of *F. occidentalis* ABC proteins to the different ABC subfamilies (A-H) was assessed by a BLASTp search against *D. melanogaster* ABC protein sequences and the NCBI protein database (Altschul et al., 1990). In addition, we also performed a phylogenetic analysis of the nuclear binding domains (NBDs) of all annotated *F. occidentalis* ABC genes (encoded by either putative or incomplete/pseudo ABC genes). NBDs were extracted from the ABC protein sequences of *F. occidentalis* using the ScanProsite facility (de Castro et al., 2006) and the Prosite profile PS50893. The alignment of the NBDs of *F. occidentalis*, *Drosophila melanogaster*, *Daphnia pulex* and *Homo sapiens* was performed with MAFFT v7 with 1000 iterations and the options “E-INS-I” and “reorder” (Katoh et al., 2002). A phylogenetic analysis was performed on the Cipres web portal (Miller et al., 2010) using RAxML v8 HPC2-XSEDE (Stamatakis, 2014) with the automatic protein model assignment algorithm using maximum likelihood criterion and 250 bootstrap replicates. The LG likelihood with empirical base frequencies was chosen as the best scoring model by RAxML. We also performed a phylogenetic analysis for the ABCA, ABCC, ABCG and ABCH subfamilies, using the same methodology as described above except that complete ABC protein sequences were used instead of NBDs and more arthropod species were included (*Anopheles gambiae*, *Apis mellifera*, *Bemisia tabaci*, *Bombyx mori*, *Lygus hesperus*, *Laodelphax striatellus*, *Plutella xylostella*, *Tribolium castaneum* and *Tetranychus urticae*) (Dermauw and Van Leeuwen, 2014; Hull et al., 2014; Qi et al., 2016; Sun et al., 2017; Tian et al., 2017). In addition, only *F. occidentalis* ABC proteins encoded by putative ABC genes (see above for definition) were included, except for FoABCC-12, FoABCG-18 and FoABCH-06 (<70% average length), being highly conserved in other insects. In all four analyses, The LG model with empirical base frequencies was chosen as the best scoring model by RAxML. All phylogenetic trees were visualized, optimized and mid-point rooted with MEGA6 (Tamura et al., 2013) and edited in Corel-DRAW Home & Student x7.

###### *7.1.1.2 Carboxyl/cholinesterase (CCE) gene family*

*D. melanogaster* and *Acyrthosiphon pisum* CCE protein sequences were used as queries to perform tBLASTn (E-value threshold <E $10^{-5}$) searches against the genome sequence assembly of *F. occidentalis.* CCE gene models were refined or created based on the homology and available RNA-seq data. Full length CCE genes and incomplete genes that had a sequence length larger than 70% of the average sequence length of complete CCEs were considered as putative CCEs and

were included in a phylogenetic analysis. All *F. occidentalis* CCE gene models that have been annotated in this study can be found in **Additional file 2: Table S16**. CCE sequences of *F. occidentalis*, *Drosophila melanogaster*, *Apis mellifera*, *Acyrthosiphon pisum* and *B. tabaci* MEAM1 (Chen et al., 2016; Claudianos et al., 2006; Ramsey et al., 2010) were aligned using MAFFT v7 with 1000 iterations and the options "l-INS-i" and "reorder" (Katoh et al., 2002). Only sequences larger than 400 AA were retained for further analysis (~ 70% length average CCE *F. occidentalis*). The Cipres web portal was used to perform the phylogenetic analysis (Miller et al., 2010) using RAxML v8 HPC2-XSEDE (Stamatakis, 2014) with the automatic protein model assignment algorithm using maximum likelihood criterion and 250 bootstrap replicates. The LG likelihood with fixed base frequencies was chosen as the best scoring model by RAxML. The resulting tree was visualized, midpoint rooted and optimized with MEGA6 (Tamura et al., 2013) and edited in Corel-DRAW Home & Student x7.

##### 7.2.2. Results and Discussion

###### *7.2.2.1 ABC gene family*

The ABC protein family is one of the largest protein families and present in all kingdoms of life. The majority functions as primary active transporters, hydrolyzing ATP to transport substrates across membranes Some ABC proteins, however, are receptors or are involved in translation. ABC proteins are divided into eight (A to H) groups in Metazoa, of which the ABCB full transporters [also named P-glycoproteins, P-gps, multidrug resistance proteins (MDRs)], ABCCs [also named multi drug resistance associated proteins (MRPs)] and ABCGs have been linked to xenobiotic resistance (Dermauw and Van Leeuwen et al. 2014). We annotated 45 putative ABC genes in the genome of *F. occidentalis* (**Table S7.3**) This number is similar to those found in *D. melanogaster* (Dermauw and Van Leeuwen et al. 2014) but less than those identified in other insect genomes, including those of *B. tabaci* and *L. hesperus* (Hull et al., 2014; Tian et al., 2017), member of the Hemiptera, a sistergroup of the Thysanoptera (Johnson et al., 2018). For those ABC proteins that are considered as conserved in metazoan species (ABCB half transporters, ABCDs, ABCE, ABCFs, FoABCC-04, FoABCG-11 and FoABCG-11; Dermauw and Van Leeuwen, 2014; **Table S7.3, Figure S7.2, Figure S7.4 and Figure S7.5**), we found orthologues in *F. occidentalis*. In many cases, we also detected *F. occidentalis* orthologues of ABC proteins that are conserved across most insects (FoABCA-01, FoABCG-01, FoABCG-02, FoABCG-04, FoABCG-05, FoABCG-08, FoABCG-13, FoABCG-21, FoABCH-01, FoABCH-03 and FoABCH-06 (Dermauw and Van Leeuwen, 2014), **Figure S7.3, Figure S7.5, Figure S7.6**). We also found a clear *F. occidentalis* ortholog (FoABCC-12) for the *D. melanogaster* sulfonylurea receptor (sur) within the ABCC subfamily (**Figure S7.4**). In contrast to *T. castaneum* and *T. urticae* we did not identify lineage specific expansions within the ABCC subfamily, a family well known for their role in multidrug resistance (Schinkel and Jonker, 2012). However, similar to *T. urticae*, *D. pulex*, *P. xylostella* and *B. tabaci* (belonging to the Hemiptera, a sister-group of the Thysanoptera) we identified a lineage specific expansion of ABCH genes within the *F. occidentalis* genome (**Figure S7.6**). The physiological functions of lineage-specific ABCH expansions remain, however, largely uncharacterized. Differential expression of ABCH genes has been reported for *T. urticae* females in diapause (Bryon et al., 2013) and some ABCH genes were overexpressed in insecticide/acaricide resistant strains of *T. urticae* (Dermauw et al., 2013b) and *P. xylostella* (Qi et al., 2016) suggesting that these lineage-specific ABCHs in *F. occidentalis* might play a role in response to environmental change or exposure to xenobiotic compounds.

**Table S7.3 - Gene numbers in ATP-binding cassette (ABC) gene subfamilies of nine arthropod species and *Homo sapiens****

| Species | Order | A | B-FT | B-HT | C | D | E | F | G | H | Total |
|---|---|---|---|---|---|---|---|---|---|---|---|
| *Homo sapiens* | Mammalia: Monotremata | 12 | 4 | 7 | 12 | 4 | 1 | 3 | 5 | 0 | 48 |
| *Daphnia pulex* | Crustacea: Cladocera | 4 | 2 | 5 | 7 | 3 | 1 | 4 | 24 | 15 | 65 |
| *Tetranychus urticae* | Arachnida: Acari: Trombidiformes | 9 | 2 | 2 | 39 | 2 | 1 | 3 | 23 | 22 | 103 |
| *Drosophila melanogaster* | Insecta: Diptera | 10 | 4 | 4 | 14 | 2 | 1 | 3 | 15 | 3 | 56 |
| *Apis mellifera* | Insecta: Hymenoptera | 3 | 3 | 4 | 9 | 2 | 1 | 3 | 15 | 3 | 43 |
| *Tribolium castaneum* | Insecta: Coleoptera | 10 | 2 | 4 | 35 | 2 | 1 | 3 | 13 | 3 | 73 |
| *Plutella xylostella* | Insecta: Lepidoptera | 15 | 7 | 7 | 21 | 3 | 1 | 3 | 19 | 6 | 82 |
| *Bombyx mori* | Insecta: Lepidoptera | 6** | 5 | 4 | 15 | 2 | 1 | 3 | 13 | 3 | 52 |
| *Lygus hesperus**** | Insecta: Hemiptera: Heteroptera | 11 | 3 | 3 | 12 | 2 | 1 | 3 | 19 | 11 | 65 |
| *Bemisia tabaci* Q | Insecta: Hemiptera: Sternorrhyncha | 8 | 0 | 3 | 6 | 2 | 1 | 3 | 23 | 9 | 55 |
| ***Frankliniella occidentalis*** | **Insecta: Thysanoptera** | **3** | **1** | **4** | **11** | **2** | **1** | **3** | **14** | **6** | **45** |

* numbers were derived from Dermauw & Van Leeuwen 2014 and references therein, Hull et al. 2014, Qi et al. 2017 and Tian et al. 2017
** a BLASTp analysis of *B. mori* ABCA proteins against the current *B. mori* annotation (January 2019) in the NCBI database revealed that the old *B. mori* ABCA gene models (BGIBMGA accessions) were incorrect and that some old models should be merged or updated, resulting in only six *B. mori* ABCA genes, instead of nine as previously reported
*** based on transcriptomic data

##### *7.2.2.2 CCE gene family*

The carboxyl/cholinesterase (CCE) enzyme family catalyzes the hydrolysis of carboxylesters and plays role in many biological processes, such as neuron signaling, development and detoxification of xenobiotics, including insecticides (Claudianos et al., 2006; Després et al., 2007; Oakeshott et al., 2005). Within the CCEs 13 clades can be distinguished, which in turn can be grouped into 3 classes: the dietary/ detoxification enzymes (clades A–C), the pheromone/hormone processing enzymes (clades D–G) and the neurodevelopmental CCEs (clades I–M, the majority being non catalytic esterases) (Claudianos et al., 2006; Oakeshott et al., 2005). We annotated 50 putative full-length CCE genes and 16 incomplete/pseudogenes in the *F. occidentalis* genome (**Table S7.4; Additional file 2: Table S16**). This number is similar to what is found in *B. tabaci* MEAM1 (51) and *T. castaneum* (49) but higher than those in *D. melanogaster* (35) and *Acyrthosiphon. pisum* (29) (**Table S7.4**, (Chen et al., 2016; Oakeshott et al., 2010; Ramsey et al., 2010; Yu et al., 2009)).

**Table S7.4 - Number of carboxyl/choline esterase (CCE) genes in insect species from different insect orders*, ****

| CCE classes and clades | *Dm* | *Bm* | *Am* | *Tc* | *Ap* | *Bt* MEAM1*** | ***Fo*** |
|---|---|---|---|---|---|---|---|
| Detoxification/Dietary | 13 | 55 | 8 | 26 | 5 | 7 | **28** |
| Pheromone/hormone processing | 8 | 8 | 5 | 11 | 17***** | 17**** | **7******* |
| Neuro/Developmental (total) | 14 | 13 | 11 | 12 | 7 | 11 | **15** |
| Clade H - Glutactins | 4 | 0 | 0 | 1 | 0 | 1 | **2** |
| Clade J - Acetylcholinesterase | 1 | 2 | 2 | 2 | 2 | 2 | **2** |
| Clade K - Gliotactin | 1 | 1 | 1 | 1 | 1 | 1 | **1** |
| Clade L - Neuroligins | 4 | 6 | 5 | 5 | 3 | 6 | **7** |
| Clade M - Neurotactin | 2 | 2 | 1 | 2 | 0 | 0 | **1** |
| Clade I - Uncharacterized CCEs | 2 | 2 | 2 | 1 | 1 | 2 | **2** |
| Total | 35 | 76 | 24 | 49 | 29 | 36 (51) | **50** |

*numbers were derived from Ramsey et al. 2010, Yu et al. 2009, Oakeshott et al. 2010, Chen et al. 2016 and this study
**abbreviations: *Dm, Drosophila melanogaster*, *Bm, Bombyx mori*, *Am*, *Apis mellifera*, *Tribolium castaneum*, *Acyrthosiphon pisum, Bt, Bemisia tabaci* and *Fo*, *Frankliniella occidentalis*
*** assignment of *B. tabaci* MEAM1 CCEs [having a clear BLASTp hit with arthropod CCEs in the NCBI database and being larger than 400 amino acids (Bta09993/Bta08457 and Bta00353/Bta03441/Bta04865/Bta05217/Bta06364/Bta08029/Bta08783/Bta08786/Bta09351, Bta09992/Bta10442/Bta11683/Bta12234 did not fulfill the first and second criterium, respectively): 36 out of 51 CCEs in Chen et al. 2016] to different classes and/or clades was based on Figure S7.7; the total number of *B. tabaci* CCEs is (51) shown between brackets
**** *F. occidentalis* and *B. tabaci* MEAM1 CCEs that did not cluster into the Detoxification/Dietary class nor the Neuro/Developmental clades in Figure S7.7 were designated as members of the Pheromone/hormone processing class
***** only 13 *Acyrthosiphon pisum* CCEs of the Pheromone/hormone Processing class were larger than 400 AA and were included in the phylogenetic analysis shown in Figure S7.7.

Based on a phylogenetic analysis with CCEs from *D. melanogaster*, *A. mellifera*, *A. pisum and B. tabaci*, the *F. occidentalis* CCEs could be assigned to the different CCE classes and/or clades (Claudianos et al., 2006; Oakeshott et al., 2005): 28 within the dietary/detoxification class, 7 within the pheromone/hormone processing class and 15 within the neurodevelopmental class. In several cases (mainly within the neurodevelopmental class) clear orthologous relationships were identified between *F. occidentalis* CCEs and those of other insect species (**Figure S7.7**) [FoCCE-28 and FoCCE-57 (clade J, AChE), FoCCE-12 (clade K, gliotactin), FoCCE-04, FoCCE-13, FoCCE-06/FoCCE-15, FoCCE-19/FoCCE-25 and FoCCE-39 (clade L, neuroligins), FoCCE-34 (clade M, neurotactins), FoCCE-03 and FoCCE-35 (clade I)]. Clade J contains CCE genes coding for acetylcholinesterase (AChE), a key enzyme in the central nervous system and in which mutations are known to confer organophosphate and/ or carbamate resistance. Contrary to *D. melanogaster*, most insects have two AChEs and it was hypothesized that the two genes were derived from an old duplication before the split of the Arthropoda (Huchard et al., 2006). In line with hypothesis, two AChEs [FoCCE-57, ortholog of Ace1, and FoCCE-28, ortholog of Ace2] could be identified in the *F. occidentalis* genome. Furthermore, we found three *F. occidentalis* CCEs (FoCCE-16, FoCCE-18, FoCCE-47) that clustered with high bootstrap support with *A. mellifera* GB15327 and GB10820, previously characterized as juvenile hormone esterase-like enzymes (Mackert et al., 2008) (**Figure S7.7**). Finally, in line with other species included in our analysis, we found a lineage-specific expansion of *F. occidentalis* CCEs within the dietary/detoxification enzyme class. Such expansions have also been reported for other species (**Table S7.4, Figure S7.7** (Yu et al., 2009)), but except for the herbivorous *B. mori*, this is the largest expansion of dietary/detoxification CCEs reported for an insect species (Table S7.4). Future work should confirm whether *F. occidentalis* CCEs are indeed detoxification CCEs and whether their expansion might be related to the polyphagous nature and/or fast resistance development of *F. occidentalis* (Jensen, 2000).

##### 7.2.3. Summary

Forty-five and 50 putative ABC and CCE genes were annotated in the *F. occidentalis* genome, respectively. The number of *F. occidentalis* ABC genes is on the lower side among those reported for other insect species (**Table S7.3**, (Tian et al., 2017; Xie et al., 2018) including *Bemisia tabaci* of the Hemiptera, the sister-group of the Thysanoptera (Johnson et al., 2018). Nevertheless, we did identify a lineage-specific expansion of ABCH genes within the *F. occidentalis* genome (**Table S7.3, Figure S7.6**). Lineage-specific arthropod ABCH genes were previously shown to respond to environmental changes or xenobiotic exposure (Bryon et al., 2013; Dermauw et al., 2013b; Qi et al., 2016) and hence these ABCH genes might have a similar function in *F. occidentalis*. In contrast to ABC genes, the number of *F. occidentalis* CCE genes is among the highest of those identified in insect species (Table S7.4, Xie et al., 2018). This high number of CCEs is due to a lineage-specific expansion within the dietary/detoxification class of CCEs (**Figure S7.7**). Future work should confirm whether these 28 *F. occidentalis* specific CCEs are actually detoxification CCEs and whether the polyphagous nature and/or fast resistance development of *F. occidentalis* (Jensen, 2000) might be related to this CCE expansion.

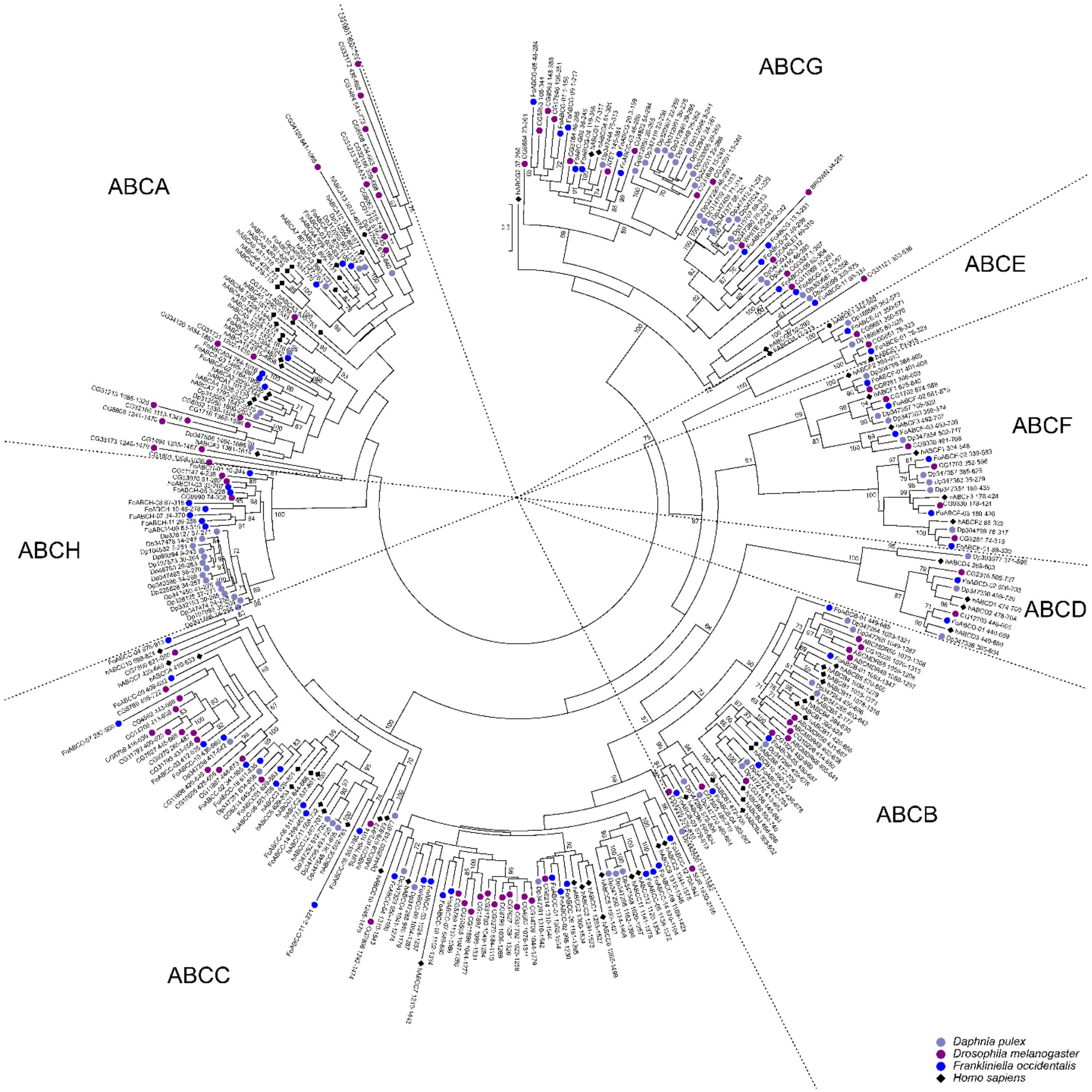

**Figure S7.2 - Maximum likelihood phylogenetic analysis of the NBDs of ABC proteins.** Maximum likelihood phylogenetic analysis of the NBDs of ABC proteins of *Daphnia pulex*, *Drosophila melanogaster*, *Frankliniella occidentalis* and *Homo sapiens.* The scale bar represents 0.5 amino-acid substitutions per site. Numbers behind the accession ID or name of an ABC genes indicate the position of the NBD in the ABC protein sequence. The different metazoan ABC protein subfamilies (A-H) are labelled.

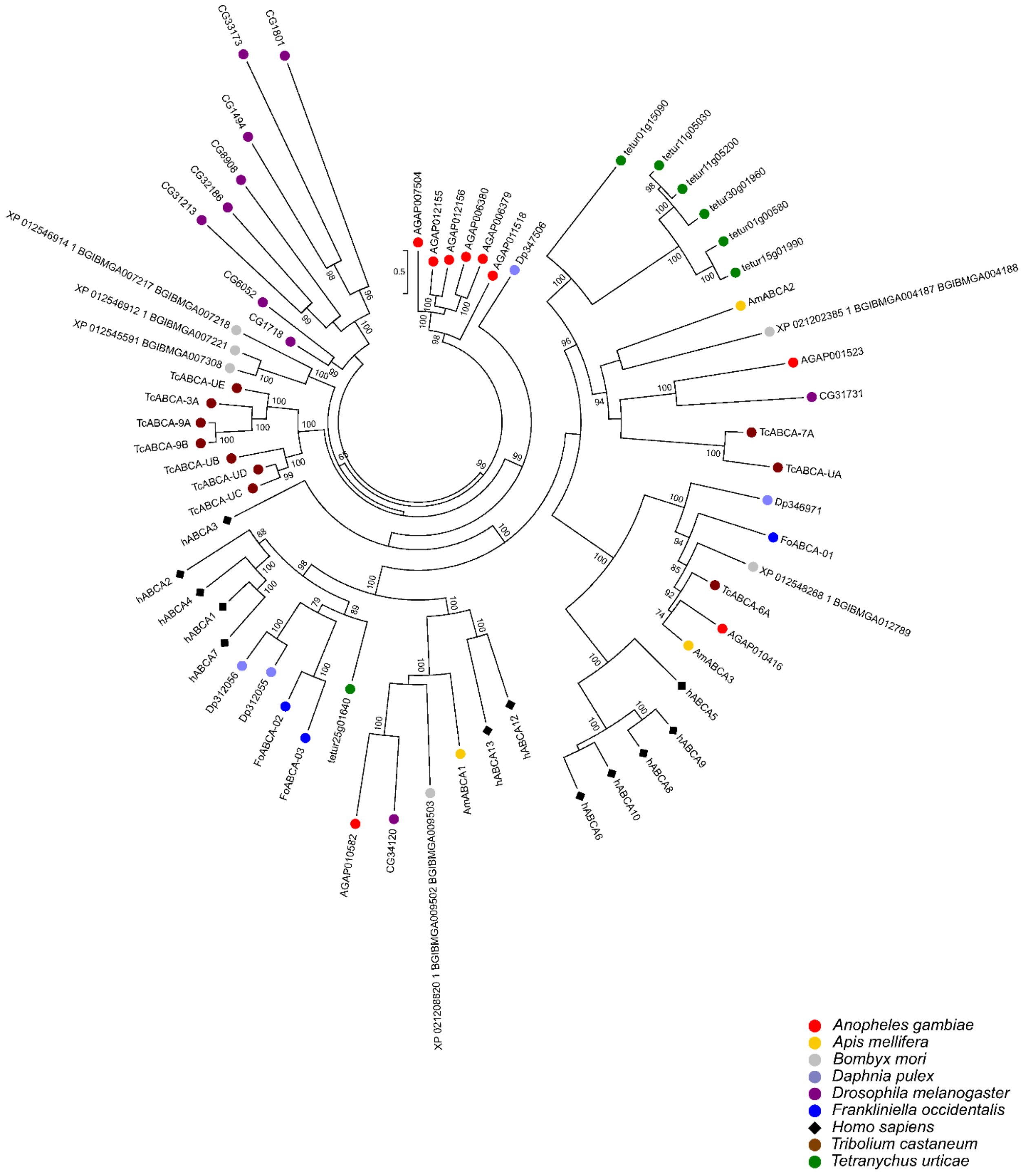

**Figure S7.3 - Maximum likelihood phylogenetic analysis of arthropod ABCA proteins.** Maximum likelihood phylogenetic analysis of the ABCA proteins of *Anopheles gambiae*, *Apis mellifera*, *Daphnia pulex*, *Drosophila melanogaster*, *Frankliniella occidentalis*, *Homo sapiens*, *Tribolium castaneum* and *Tetranychus urticae.* The scale bar represents 0.5 amino-acid substitutions per site.

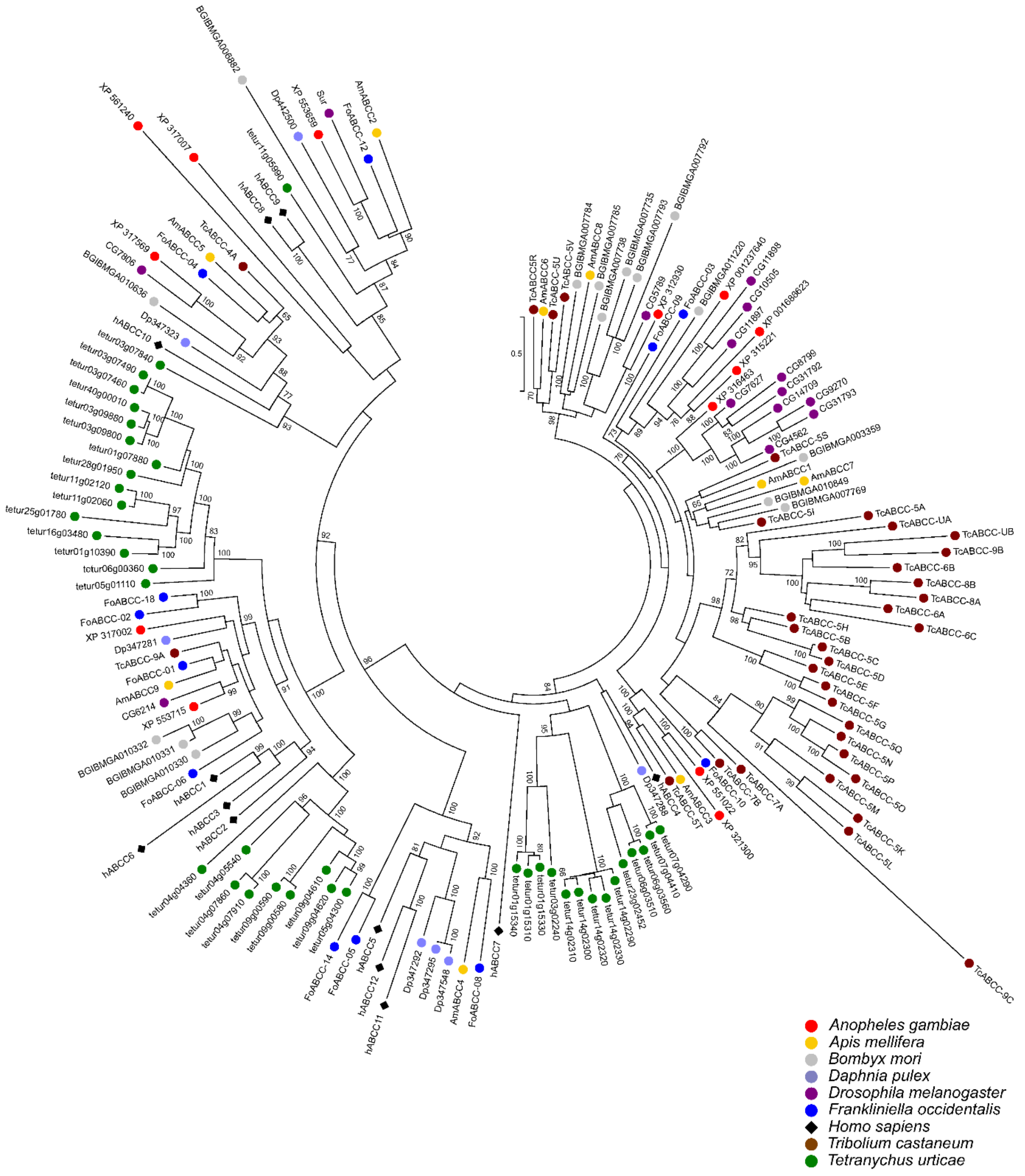

**Figure S7.4 - Maximum likelihood phylogenetic analysis of arthropod ABCC proteins.** Maximum likelihood phylogenetic analysis of the ABCC proteins of *Anopheles gambiae*, *Apis mellifera*, *Bombyx mori*, *Daphnia pulex*, *Drosophila melanogaster*, *Frankliniella occidentalis*, *Homo sapiens*, *Tribolium castaneum* and *Tetranychus urticae.* The scale bar represents 0.5 amino-acid substitutions per site.

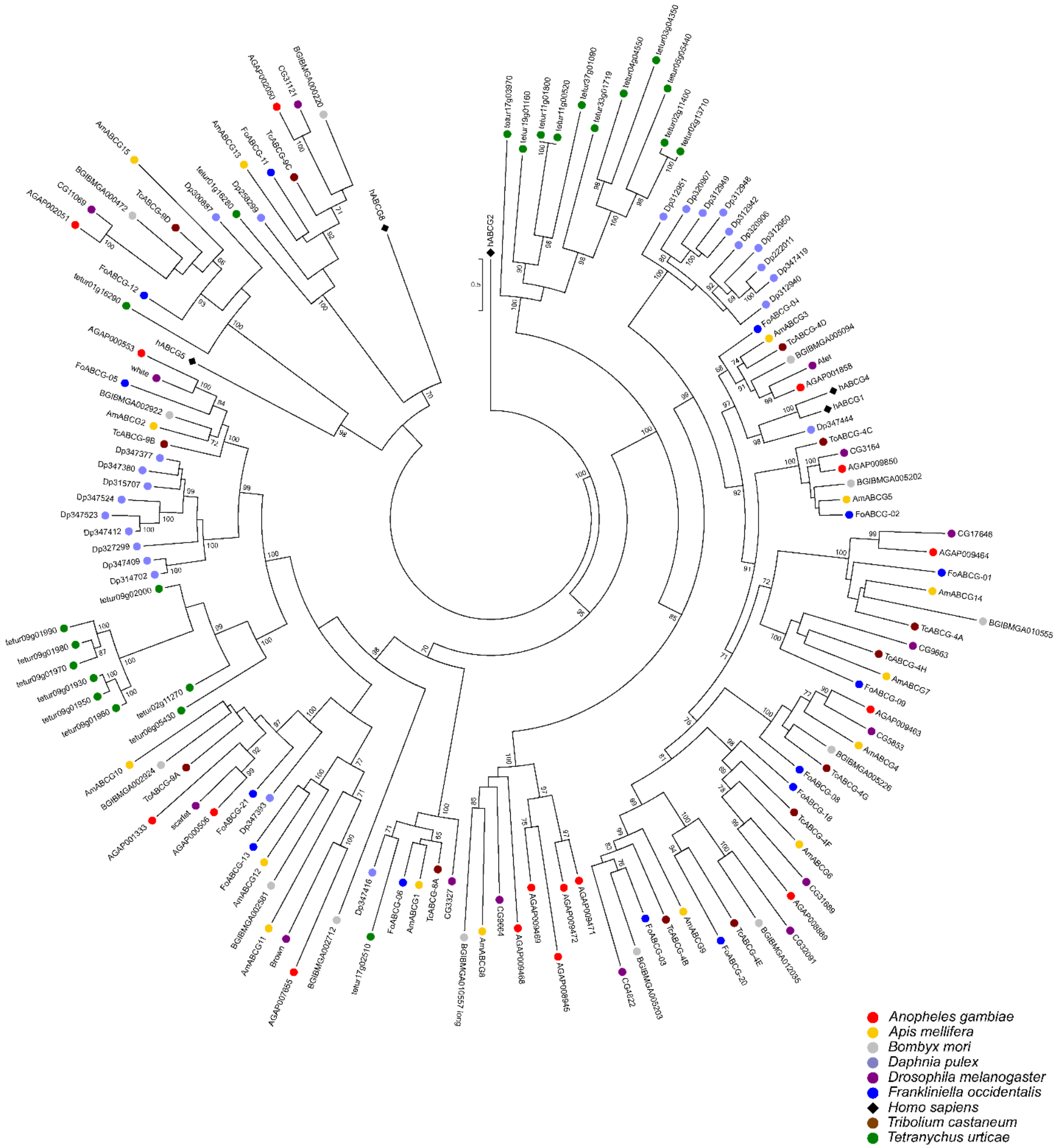

**Figure S7.5 - Maximum likelihood phylogenetic analysis of arthropod ABCG proteins.** Maximum likelihood phylogenetic analysis of the ABCG proteins of *Anopheles gambiae*, *Apis mellifera*, *Bombyx mori*, *Daphnia pulex*, *Drosophila melanogaster*, *Frankliniella occidentalis*, *Homo sapiens*, *Tribolium castaneum* and *Tetranychus urticae.* The scale bar represents 0.5 amino-acid substitutions per site.

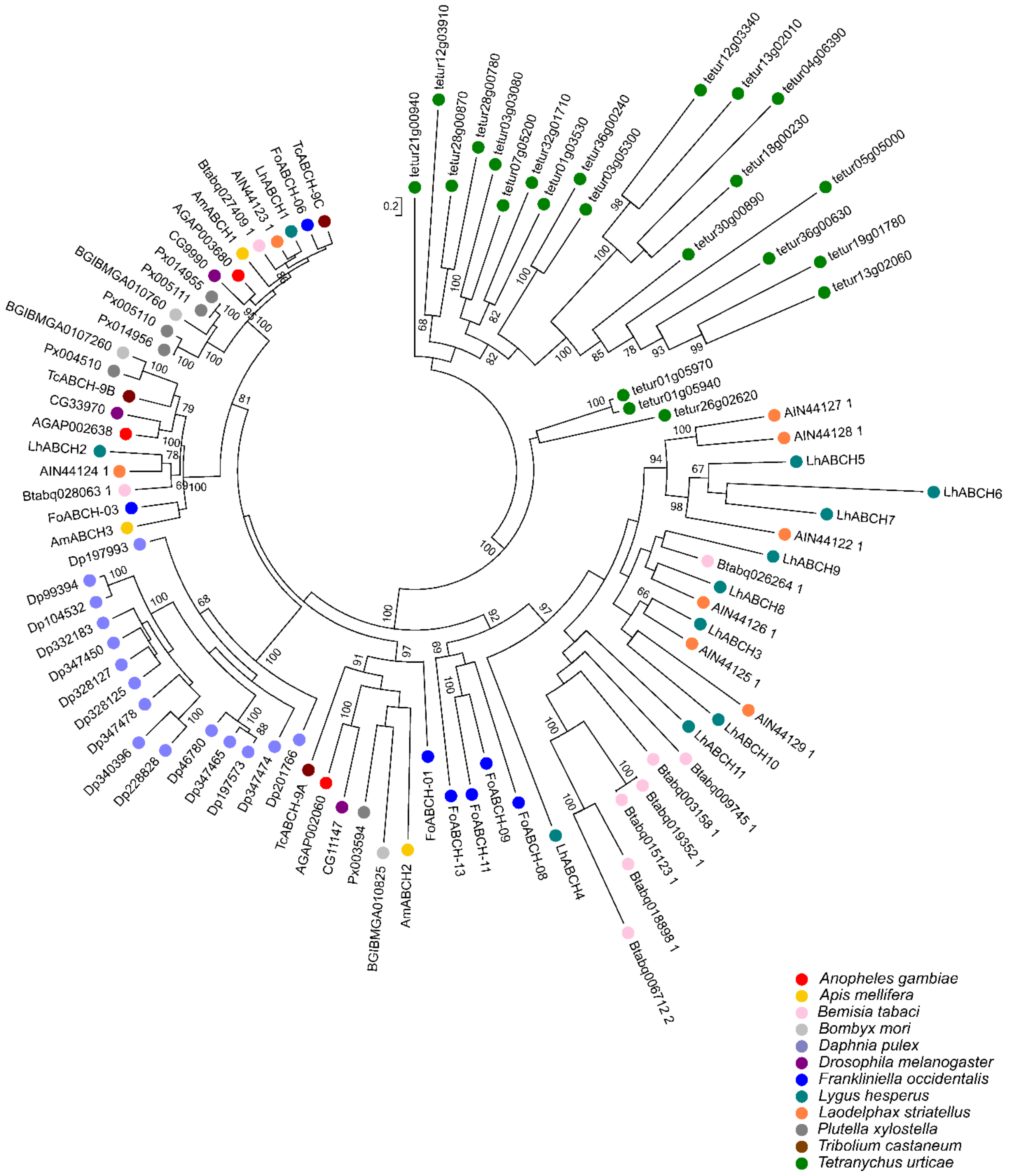

**Figure S7.6 - Maximum likelihood phylogenetic analysis of arthropod ABCH proteins.** Maximum likelihood phylogenetic analysis of the ABCH proteins of *Anopheles gambiae*, *Apis mellifera*, *Bemisia tabaci*, *Bombyx mori*, *Daphnia pulex*, *Drosophila melanogaster*, *Frankliniella occidentalis*, *Lygus hesperus*, *Laodelphax striatellus*, *Plutella xylostella*, *Tribolium castaneum* and *Tetranychus urticae.* The scale bar represents 0.2 amino-acid substitutions per site.

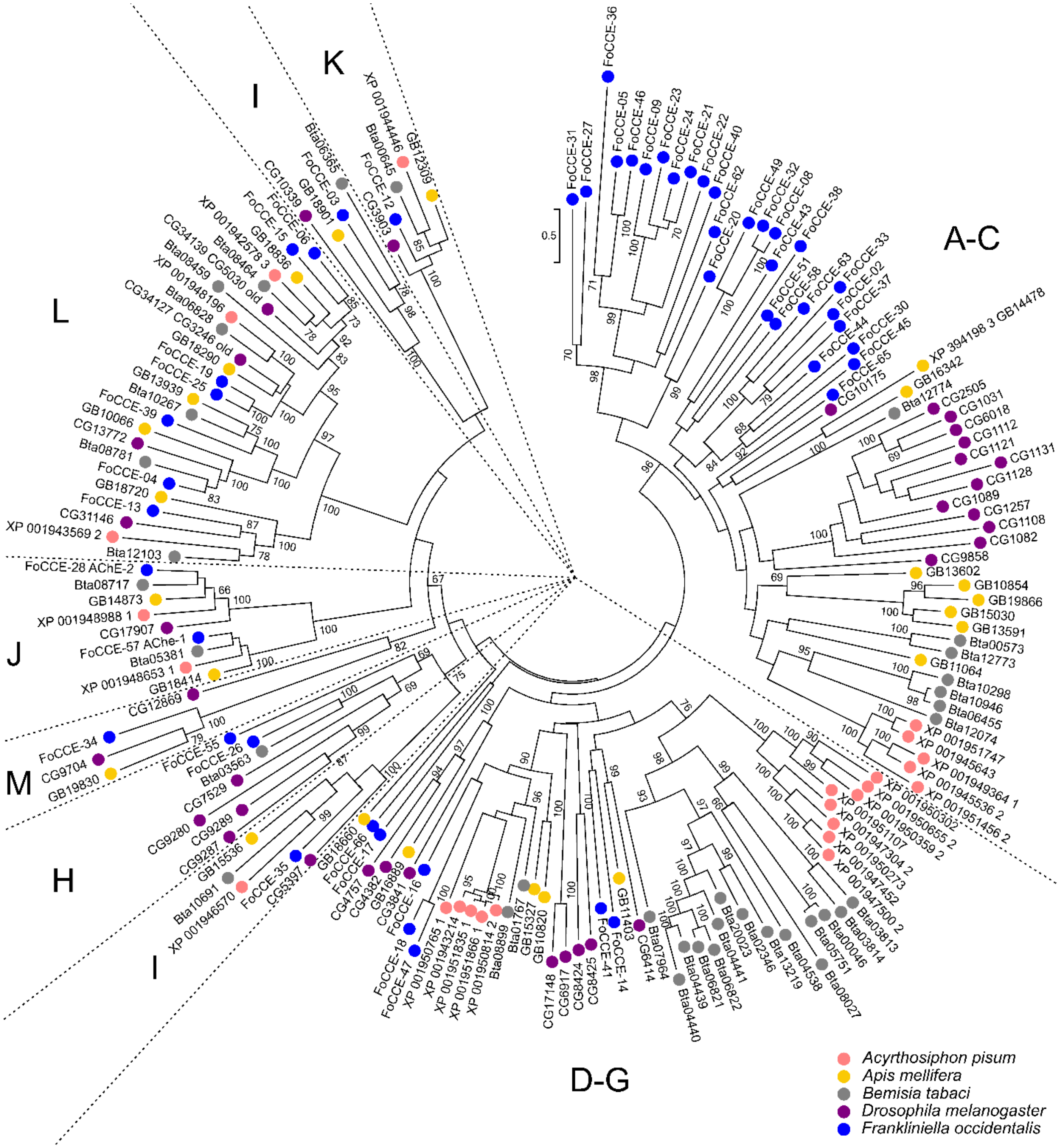

**Figure S7.7 - Maximum likelihood phylogenetic analysis of arthropod CCEs.** Maximum likelihood phylogenetic analysis of the CCEs of *Acyrthosiphon pisum*, *Apis mellifera*, *Bemisia tabaci*, *Drosophila melanogaster* and *Frankliniella occidentalis.* The CCEs clustered into classes and/or clades (Claudianos et al., 2006): A-C (dietary class), D-G (hormone/semiochemical class), H (glutactin and like enzymes), I (uncharacterized CCEs), J (AChEs), K (gliotactins), L (neuroligins), M (neurotactins). The scale bar represents 0.5 amino-acid substitutions per site.

Johnson, K.P., Dietrich, C.H., Friedrich, F., Beutel, R.G., Wipfler, B., Peters, R.S., Allen, J.M., Petersen, M., Donath, A., Walden, K.K.O., Kozlov, A.M., Podsiadlowski, L., Mayer, C., Meusemann, K., Vasilikopoulos, A., Waterhouse, R.M., Cameron, S.L., Weirauch, C., Swanson, D.R., Percy, D.M., Hardy, N.B., Terry, I., Liu, S., Zhou, X., Misof, B.,

Robertson, H.M., Yoshizawa, K., 2018. Phylogenomics and the evolution of hemipteroid insects. Proc. Natl. Acad. Sci. 115, 12775–1278. https://doi.org/10.1073/pnas.1815820115

Katoh, K., Misawa, K., Kuma, K., Miyata, T., 2002. MAFFT: a novel method for rapid multiple sequence alignment based on fast Fourier transform. Nucleic Acids Res. 30, 3059–3066. https://doi.org/10.1093/nar/gkf436

Mackert, A., do Nascimento, A.M., Bitondi, M.M.G., Hartfelder, K., Simões, Z.L.P., 2008. Identification of a juvenile hormone esterase-like gene in the honey bee, *Apis mellifera* L. - Expression analysis and functional assays. Comp. Biochem. Physiol. - B Biochem. Mol. Biol. 150, 33–44. https://doi.org/10.1016/j.cbpb.2008.01.004

Miller, M.A., Pfeiffer, W., Schwartz, T., 2010. Creating the CIPRES science gateway for inference of large phylogenetic trees, in: 2010 Gateway Computing Environments Workshop, GCE 2010. IEEE, New Orleans, LA, USA, pp. 1–8. https://doi.org/10.1109/GCE.2010.5676129

Oakeshott, J.G., Claudianos, C., Campbell, P.M., Newcomb, R.D., Russell, R.J., 2005. Biochemical genetics and genomics of insect esterases, in: Comprehensive Molecular Insect Science. pp. 309–381. https://doi.org/10.1016/B0-44-451924-6/00073-9

Oakeshott, J.G., Johnson, R.M., Berenbaum, M.R., Ranson, H., Cristino, A.S., Claudianos, C., 2010. Metabolic enzymes associated with xenobiotic and chemosensory responses in *Nasonia vitripennis*. Insect Mol. Biol. 19, 147–163. https://doi.org/10.1111/j.1365-2583.2009.00961.x

Qi, W., Ma, X., He, W., Chen, W., Zou, M., Gurr, G.M., Vasseur, L., You, M., 2016. Characterization and expression profiling of ATP-binding cassette transporter genes in the diamondback moth, *Plutella xylostella* (L.). BMC Genomics 17, 760. https://doi.org/10.1186/s12864-016-3096-1

Ramsey, J.S., Rider, D.S., Walsh, T.K., De Vos, M., Gordon, K.H.J., Ponnala, L., MacMil, S.L., Roe, B.A., Jander, G., 2010. Comparative analysis of detoxification enzymes in *Acyrthosiphon pisum* and *Myzus persicae*. Insect Mol. Biol. 2, 155–164. https://doi.org/10.1111/j.1365-2583.2009.00973.x

Schinkel, A.H., Jonker, J.W., 2012. Mammalian drug efflux transporters of the ATP binding cassette (ABC) family: An overview. Adv. Drug Deliv. Rev. https://doi.org/10.1016/j.addr.2012.09.027

Stamatakis, A., 2014. RAxML version 8: a tool for phylogenetic analysis and post-analysis of large phylogenies. Bioinformatics 30, 1312–3. https://doi.org/10.1093/bioinformatics/btu033

Sturm, A., Cunningham, P., Dean, M., 2009. The ABC transporter gene family of *Daphnia pulex*. BMC Genomics 10, 170. https://doi.org/10.1186/1471-2164-10-170

Sun, H., Pu, J., Chen, F., Wang, J., Han, Z., 2017. Multiple ATP-binding cassette transporters are involved in insecticide resistance in the small brown planthopper, *Laodelphax striatellus*. Insect Mol. Biol. 26, 343–355. https://doi.org/10.1111/imb.12299

Tamura, K., Stecher, G., Peterson, D., Filipski, A., Kumar, S., 2013. MEGA6: molecular evolutionary genetics analysis version 6.0. Mol. Biol. Evol. 30, 2725–2729. https://doi.org/10.1093/molbev/mst197

Tian, L., Song, T., He, R., Zeng, Y., Xie, W., Wu, Q., Wang, S., Zhou, X., Zhang, Y., 2017. Genome-wide analysis of ATP-binding cassette (ABC) transporters in the sweetpotato whitefly, *Bemisia tabaci*. BMC Genomics 18, 330. https://doi.org/10.1186/s12864-017-3706-6

Xie, W., Yang, X., Chen, C., Yang, Z., Guo, L., Wang, D., Huang, J., Zhang, H., Wen, Y., Zhao, J., Wu, Q., Wang, S., Coates, B.S., Zhou, X., Zhang, Y., 2018. The invasive MED/Q *Bemisia tabaci* genome: a tale of gene loss and gene gain. BMC Genomics 19, 68. https://doi.org/10.1186/s12864-018-4448-9

Yu, Q.Y., Lu, C., Li, W. Le, Xiang, Z.H., Zhang, Z., 2009. Annotation and expression of carboxylesterases in the silkworm, *Bombyx mori*. BMC Genomics 10, 533. https://doi.org/10.1186/1471-2164-10-553

### 8. Innate Immune genes

*Contributed by Chris G. C. Jacobs, Maurijn Vander Zea, Jonathan Oliver and Swapna Priya Rajarapu*

#### 8.1. Abstract

Immune responses of insect vectors play a critical role during virus infection and thus are important biological phenomenon to study. Similar to hemipterans, *F. occidentalis* genome encodes canonical immune pathway members except immunodeficiency gene. In addition, *F. occidentalis* possess higher numbers of pathogen recognition proteins relative to other insect species.

#### 8.2. Background

Insects, including thrips, encounter a wide variety of bacteria ranging from beneficial to highly pathogenic, along with fungi, viruses and parasites. To combat unwanted microbes, insects rely on innate immune defense mechanisms which have been best studied in *Drosophila* (Lemaitre and Hoffmann, 2007; Ligoxygakis, 2013). The most prominent response is the humoral or systemic response which involves the secretion of massive amounts of antimicrobial peptides (AMPs) by the fat body into the haemolymph (Ganesan et al., 2011). Local immune responses by epithelia include the production of AMPs and the generation of Reactive Oxygen Species (ROS) by the enzyme Duox (Dual Oxidase) (Davis and Engstrom, 2012; Ferrandon, 2013). Finally, haemocytes execute the cellular response including encapsulation of larger invaders such as parasites, local melanization reactions using phenoloxidase enzymes, and phagocytosis using Scavenger receptors, receptors of the Nimrod family, Dscam, and ThioEster containing Proteins (TEPs) (Vlisidou and Wood, 2015).

Invading microbes are recognized by PeptidoGlycan Recognition Proteins (PGRPs) and Gram Negative Binding Proteins (GNBPs) that activate the two main immune signaling pathways, the Toll and the IMD pathway (Ferrandon, 2013; Ganesan et al., 2011; Lemaitre and Hoffmann, 2007; Ligoxygakis, 2013). Fungi and Gram positive bacteria activate Toll signaling, whereas Gram negative bacteria activate IMD signaling. Intracellular signaling eventually leads to the nuclear localization of the NF-κBs Relish (for the IMD signaling pathway) and Dorsal/Dif (for the Toll signaling pathway), upregulating the transcription of effector genes. Although these two pathways are also activated upon viral infection (Kingsolver et al., 2013; Mussabekova et al., 2017), the main antiviral defense depends on RNAi mechanisms (see following section). In this section we report on the *Frankliniella* signal transduction, microbe recognition and immune effector genes.

##### 8.3.1. Identification of innate immune-associated genes from the transcriptomes of three orthotospovirus vector species: *F. occidentalis, F. fusca* and *Thrips palmi*

A genome-enabled, transcriptome assembly representing TSWV-infected and non-infected adult *F. occidentalis* (PRJNA454326, Schneweis et al., 2017) and two *de novo* assembled transcriptomes, one representing mixed stages of TSWV-infected and non-infected *F. fusca* (PRJNA385691, Shrestha et al., 2017) and the other representing CaCV-infected and non-infected adult *Thrips palmi* (PRJNA498538, Widana Gamage et al., 2019) were annotated against a custom made database of arthropod innate immunity genes downloaded from ImmunoDB (http://cegg.unige.ch/Insecta/immunodb) using blastx algorithm in local BLAST+ v. 2.8.1 with an E value cut-off of $10^{-5}$. Blast annotations were filtered to retain annotations with highest bit score, lowest E-value and longest alignment length.

##### 8.3.2 Comparison of innate immunity genes of *F. occidentalis, F. fusca* and *T. palmi*

Transcripts encoding immune related genes in *F. occidentalis, F. fusca* and *T. palmi*. were translated in six frames using TransDecoder (Version5.5.0, Haas, B. and Papanicolaou, A., 2016) with a minimum peptide length of 100 amino acids. Translated transcriptomes were annotated against UniProt database downloaded on April 9th 2019 using Blastp algorithm with an e-value cut-off of $10^{-5}$. Redundant translated proteins were removed by k-mer analysis to develop an initial training set for constructing Markov Model to discriminate between coding and non-coding regions. Predicted proteins homologous to proteins in UniProt database were retained for further analysis. Orthologous innate immunity genes between these three species were identified by Orthovenn2, a web-tool to identify orthologous and paralogous genes, with a pairwise sequence similarity cut-off of $10^{-5}$ and an inflation of 1.5 to define orthologous cluster structure (Xu et al 2019). Orthologous clusters were analyzed by UniProt search and GO Slim for functional annotation.

#### 8.4. Results and Discussion

##### 8.4.1. Immune genes in *F. occidentalis* genome

We have annotated 96 immune genes (**Additional file 2: Table S17**). The most striking finding is the absence of the signal transducing molecule IMD itself. Absence of IMD has also been reported for the Hemipteran species *Rhodnius prolixus, Acyrthosiphon pisum, Bemisia tabaci* and *Diaphorina citri* (Arp et al., 2016; Chen et al., 2016; Consortium, 2010; Gerardo et al., 2010; Zhang et al., 2014). In *Oncopeltus*, IMD could not be identified by homology searches, but was found by classical cloning with degenerate primers (Panfilio et al., 2019). IMD was also reported missing from the bedbug *Cimex lectularius* (Benoit et al., 2016), but was later found using *Plautia stali* IMD as query (Nishide et al., 2019). This illustrates that IMD sequences can be highly divergent and conclusions about their absence should be drawn with care. However, we could not find *Frankliniella* IMD using any of the available Hemipteran IMD sequences as query.

For *A. pisum*, it has been suggested that its relation to Gram negative endosymbionts and its rather sterile diet of phloem sap account for a generally reduced immune repertoire and the absence of IMD (Altincicek et al., 2008; Consortium, 2010; Gerardo et al., 2010). This does not seem valid for the mesophyll feeding Western flower thrips. In contrast to *A. pisum*, almost all other components of the IMD signaling pathway are present in *Frankliniella*, including two Relish molecules (**Additional file 2: Table S17**). In addition, the thrips gene content rather suggests well developed immune competence, particularly concerning the recognition molecules. We found 14 PGRPs (verus. none in *A. pisum*, 2 in *Oncopeltus*, 6 in *Tribolium* and 13 in *Drosophila*), and 8 GNBPs (vs 2 in *A. pisum*, 1 in *Oncopeltus*, 3 in *Tribolium* and 3 in *Drosophila*) (Gerardo et al., 2010; Panfilio et al., 2019). The broad range of host plants bringing the thrips into contact with a wide variety of pathogens might require this large number of pathogen recognition genes.

Concerning the effector genes, the melanization pathway seems notably extensive. We found 6 prophenoloxidases vs. 2 in *A. pisum* and 3 in *Oncopeltus*, *Tribolium* and *Drosophila* (Gerardo et al., 2010; Panfilio et al., 2019).

In conclusion, the absence of IMD in *Frankliniella* does not seem to suggest a reduced immune repertoire, but rather a different way of mediating the response to Gram negative bacteria, possibly by Toll signaling components. In Drosophila, DAP-type peptidoglycans of Gram-negative bacteria moderately induce Toll signaling (Leone et al., 2008; Leulier et al., 2003). In *Tenebrio molitor*, PGRP-SA recognizes both Gram positive and Gram negative bacteria (Park et al., 2006). Extensive cross reactivity of the Toll and IMD signaling pathway is the currently emerging picture from studies on other insects too (Nishide et al., 2019; Yokoi et al., 2012a; Yokoi et al., 2012b) and might have set the stage for multiple independent IMD losses in evolution (Nishide et al., 2019).

##### 8.4.2. Comparison of innate immunity genes (transcripts) of *F. occidentalis*, *F. fusca* and *T. palmi*

The occurrence and number of innate immunity-associated transcripts varied across the three species (**Table S8.1; Additional file 9**). Among the pathogen recognition molecules, fibrinogen related proteins were found only in *F. occidentalis* transcriptome. Proteins spaetzle and tube encoding transcripts were not found in *T. palmi*; the myeloid differentiation primary response protein and TNF receptor associated-protein 2, important players in Toll pathway, were not found in *F. fusca* transcriptome. Interestingly, Fas-associated Death Domain (FADD), immunodeficiency (IMD) and TGF-β-activated kinase 1 (TAK1) proteins of IMD pathway were absent in all the three species. Components of JAK/STAT were found in all the three species with eight and four transcripts of cytokine receptors found in *F. occidentalis* and *F. fusca*. A single antimicrobial peptide, defensin, was found only in *F. occidentalis* and *F. fusca*. Peroxidasins, autophagy related protein and inhibitors of apoptosis have been well represented among the three transcriptomes.

Six frame translation of *F. occidentalis*, *F. fusca* and *T. palmi* resulted in 994, 296, 737 predicted protein sequences with more than 100 amino acids in length respectively. Among these, 97.38 % of *F. occidentalis*, 99.32% *F. fusca* and 89.96% *T. palmi* were retained to predict coding regions of the transcripts. Further, the start position of these translated proteins was redefined based on the blastp alignment to the homologous sequences in UniProt using position weighted matrix approach and thus refining 49, 16 and 8 start position for *F. occidentalis*, *F. fusca* and *T. palmi* respectively. Together with coding sequence prediction and start position refinement, 296 proteins were predicted for *F. occidentalis* transcriptome, 184 proteins were predicted for *F. fusca* transcriptome and 240 proteins were predicted from *T. palmi* transcriptome. Homology based transcript prediction resulted in approximately one predicted protein per transcript.

**Table S8.1: Tabulation of innate immunity-associated transcripts associated putatively with recognition, signaling and execution of defense in *F. occidentalis*, *F. fusca* and *Thrips palmi.***

| **Immune components** | Number of Transcripts | | |
|---|---|---|---|
| ***Pathogen recognition molecules*** | *F. occidentalis* | *F. fusca* | *T. palmi* |
| C-type lectins | 7 | 3 | 6 |
| Fibrinogen Related proteins | 2 | 0 | 0 |
| Peptidoglycan recognition proteins | 7 | 5 | 1 |
| 1,3-β-D-glucan-binding protein | 2 | 1 | 3 |
| ***Signaling cascades*** | | | |
| *Toll Pathway* | | | |
| Toll receptors | 2 | 2 | 2 |
| Spaetzle | 3 | 2 | 0 |
| Tube | 1 | 1 | 0 |
| Pelle | 1 | 1 | 1 |
| MyD88 | 1 | 0 | 1 |
| CACT | 1 | 1 | 1 |
| TRAF6 | 2 | 0 | 0 |
| *IMD pathway* | | | |
| CASPAR | 1 | 1 | 1 |
| FADD | 0 | 0 | 0 |
| IKKB-ird5 | 2 | 2 | 2 |
| IMD | 0 | 0 | 0 |
| TAK1 | 1 | 1 | 1 |
| TAB | 2 | 0 | 0 |
| *JAK/STAT* | | | |
| DOME | 8 | 4 | 1 |
| HOP | 1 | 1 | 1 |
| STAT | 3 | 1 | 2 |
| ***Response*** | | | |
| Hexamerin | 1 | 1 | 0 |
| Antimicrobial Peptides | 1 | 1 | 0 |
| Lysozymes | 3 | 2 | 3 |
| Peroxidasins | 8 | 4 | 5 |
| Antioxidant enzymes | 9 | 5 | 1 |
| Clip-domain serine proteases | 6 | 3 | 2 |
| Autophagy | 9 | 6 | 6 |
| Prophenoloxidase | 3 | 1 | 3 |
| Inhibitors of Apoptosis | 5 | 5 | 4 |

The canonical genes participating in humoral and cellular immune responses of insects were all identified in the three species except for few members of Toll and IMD signaling pathway. Absence of transcripts encoding core IMD pathway genes is similar with other insect vectors such as *Diaphorina citri* (Asian citrus psyllid) and *Acyrthosiphon pisum* (pea aphid) (Arp et al 2016, Gerardo et al., 2010). IMD pathway is triggered in defense to gram negative bacteria and viruses.

These core genes may be entirely missing or may be highly divergent from other species. A blastx alignments using a lower e-value cut-off ($10^{-3}$) with other recently determined divergent IMD or IMD-like partial sequences from *Cimex lecticularis* (XP_014246002) and *Oncopeltus fasciatus* (Panfilio et al., 2019), respectively, revealed weak alignments ranging in lengths of 20-70 bp (**Table S8.2**). In addition to IMD, absence of transcripts encoding FADD suggests either absence of this downstream process or evolution of an atypical signaling pathway which is yet to be deciphered.

**Table S8.2:** Blast similarity of *F. occidentalis* transcripts to IMD/IMD-like amino acid sequences from *Cimex lecticularis and Oncopeltus fasciatus*

| FOCC transcripts | Transcript Length (bp) | Subject | % identity | Alignment Length (aa) | E-value | bitscore | FOCC transcript top match to NCBI nr database (blastx) |
|---|---|---|---|---|---|---|---|
| CUFF.473.2 | 8131 | *C. lecticularis* | 31.0 | 58 | 5.46E-04 | 30.0 | hypothetical protein mediator of rna polymerase ii transcription subunit 14-like |
| CUFF.5189.3 | 9814 | *C. lecticularis* | 38.6 | 44 | 5.56E-04 | 30.4 | integrator complex subunit 10 isoform x1 |
| FOCC008130-RA | 1287 | *C. lecticularis* | 22.2 | 45 | 7.85E-04 | 26.9 | hypothetical protein |
| CUFF.3730.2 | 2801 | *O. fasciatus* | 36.2 | 58 | 3.27E-04 | 29.3 | uncharacterized protein |
| FOCC011660-RA | 2404 | *O. fasciatus* | 36.2 | 58 | 2.76E-04 | 29.3 | uncharacterized protein |
| FOCC003613-RA | 4359 | *O. fasciatus* | 30.0 | 70 | 6.92E-04 | 28.9 | hypothetical protein/staphylococcal nuclease domain-containing protein |
| CUFF.2047.2 | 12389 | *O. fasciatus* | 30.0 | 70 | 0.002 | 28.9 | hypothetical protein/staphylococcal nuclease domain-containing protein |
| CUFF.3730.1 | 2719 | *O. fasciatus* | 36.2 | 58 | 3.96E-04 | 28.9 | uncharacterized protein |

In addition to IMD, absence of transcripts encoding FADD suggests either absence of this downstream process or evolution of an atypical signaling pathway which is yet to be deciphered.

All components of JAK/STAT pathway were identified in all the three thrips species. However, transcripts homologous to cytokine receptor were over-represented in *F. occidentalis* and *F. fusca* with eight and four cytokine receptor transcripts respectively. Multiple sequence alignment of these transcripts showed they are not identical indicating that these are different transcripts. Cytokine receptors are involved in many other signaling pathways and multiple transcripts encoding these receptors in *F. occidentalis* and *F. fusca* could be involved in other biological functions including innate immunity.

##### 8.5 RNAi pathway genes

*Contributed by Olivier Christiaens, Clauvis N.T. Taning, Guy Smagghe*

RNA interference (RNAi) is a post-transcriptional gene silencing mechanism that is present in most eukaryotic organisms. In insects, three distinct RNAi pathways have been identified, namely the small interfering RNA (siRNA), the micro RNA (miRNA) and piwi-interacting RNA (piRNA) pathways. Through these pathways, RNAi is involved in gene expression regulation, protection of the genome against transposons and it is also an important element of the antiviral defense system. The siRNA and miRNA pathways work in a similar way. In the siRNA pathway, longer double-stranded RNA (dsRNA), which can either be of endogenous or exogenous (eg. viral dsRNA) origin is processed by an RNase III enzyme called Dicer-2 into smaller siRNA pieces, which are typically around 21-23bp long. These siRNAs are then taken up in the RNA induced silencing complex (RISC), which is a protein complex also containing an Argonaute enzyme (Ago-2). One of the two strands of the siRNA will be removed from the complex and the remaining single-stranded small RNA will guide the RISC complex to its complementary mRNA and bind to it through base pair binding. This will eventually result in cleaving the mRNA by Ago2, preventing further translation to protein. In the miRNA pathway, pri-miRNA is the initial precursor of the functional miRNA. In the nucleus of the cell, this stem-loop RNA structure, typically a few hundreds of nucleotides long, is processed by the enzyme Drosha into one or more pre-miRNA molecules, also containing a hairpin structure. After transport into the cytoplasm, this pre-miRNA is processed into functional miRNAs by Dicer-1. After this step, the pathway follows a similar process as the siRNA pathway, where the mRNA is eventually destroyed by Ago-1. In insects, both the siRNA and miRNA typically have a separate set of Dicer and Argonaute enzymes, in contrast to for example nematodes which only have one Dicer enzyme. The piRNA pathway finally is an entirely different process, which is still not fully understood. Piwi-interacting RNAs form the largest class of non-coding RNAs in animal cells and are involved in silencing of transposons, epigenetic methylation and play a role in the regulation of genetic elements in germ line cells (references). Similar to the other RNAi pathways, different piwi-interacting proteins, including a Piwi-Argonaute (Ago-3) and Aubergine, guide the piRNAs to their target sequence, for example a transposon, leading to its destruction.

RNA interference (RNAi) is a post-transcriptional gene silencing mechanism which is present in most eukaryotic organisms. Through several pathways, RNAi is involved in gene expression regulation, protection of the genome against transposons and it is also an important element of the antiviral defense system. The RNAi-related genes constitutes a group of genes that are all members of a diverse range of gene (super)families which are not evolutionarily related, but are linked based on their involvement in RNAi (Swevers et al., 2013; Christiaens et al., 2014). This group includes core machinery genes for the siRNA and miRNA pathways, such as the *dicer* and *argonaute* genes, as well as several genes involved in antiviral immune response and a number of genes encoding auxiliary proteins. The Western Flower thrips *Frankliniella occidentalis* is a known vector for several economically important plant viruses and is considered the primary vector for tospoviruses, including the Tomato Spotted Wilt Virus (TSWV). These viruses are of great economic importance since they can cause substantial damage to a very large range of food and ornamental plant species. Also, it is known that these viruses are so-called propagating viruses, meaning they can replicate in insect tissues. As RNAi is an important pathway in the innate innate immune response to viruses, knowledge on the machinery in thrips could be of importance.

**Figure S8. Phylogenetic trees of *F. occidentalis* RNAi pathway genes.**

#### 1. Ago 1, 2 and 3

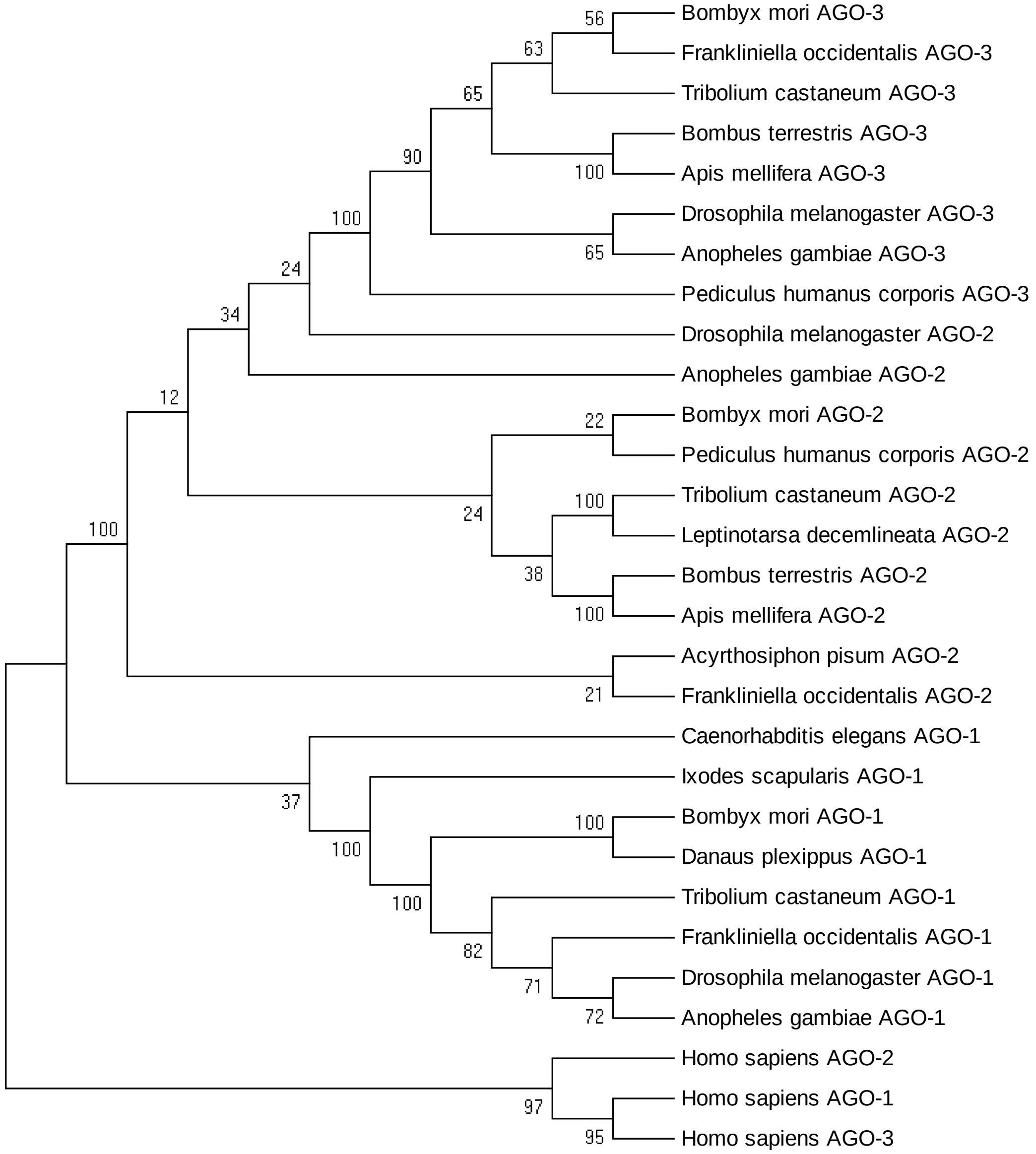

#### 2. Ago 1

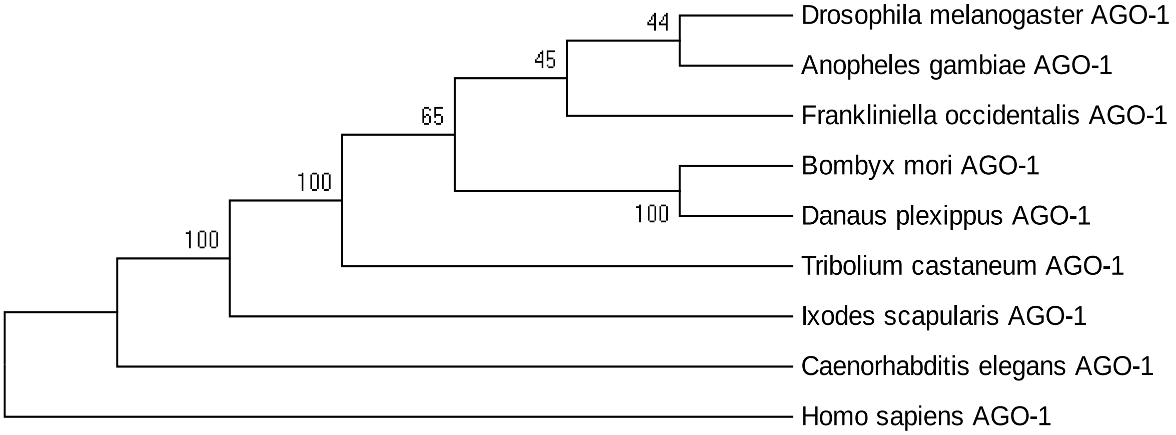

44
45
65
100
100
100
Drosophila melanogaster AGO-1
Anopheles gambiae AGO-1
Frankliniella occidentalis AGO-1
Bombyx mori AGO-1
Danaus plexippus AGO-1
Tribolium castaneum AGO-1
Ixodes scapularis AGO-1
Caenorhabditis elegans AGO-1
Homo sapiens AGO-1

#### 3. Ago 2

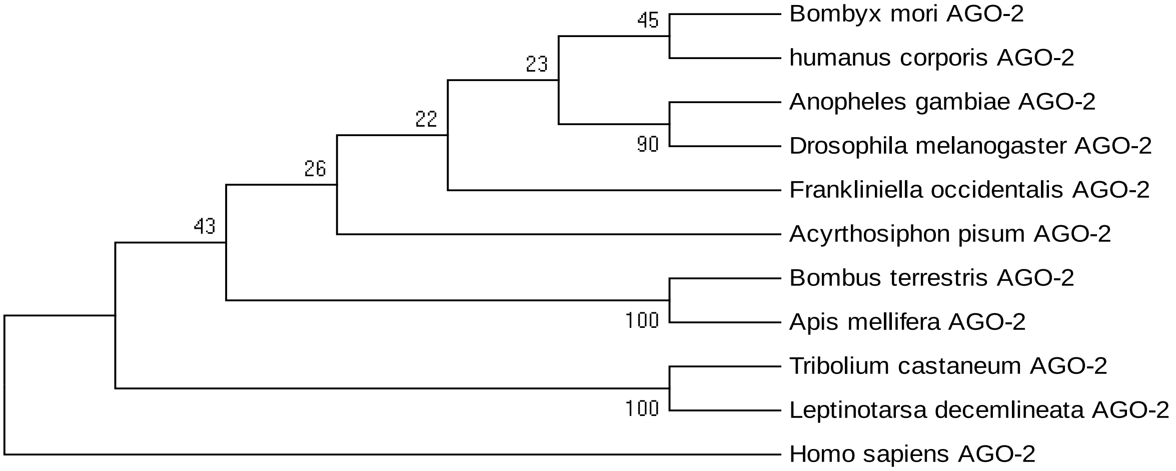

45
23
22
26
43
90
100
100
Bombyx mori AGO-2
humanus corporis AGO-2
Anopheles gambiae AGO-2
Drosophila melanogaster AGO-2
Frankliniella occidentalis AGO-2
Acyrthosiphon pisum AGO-2
Bombus terrestris AGO-2
Apis mellifera AGO-2
Tribolium castaneum AGO-2
Leptinotarsa decemlineata AGO-2
Homo sapiens AGO-2

#### 4. Ago 3

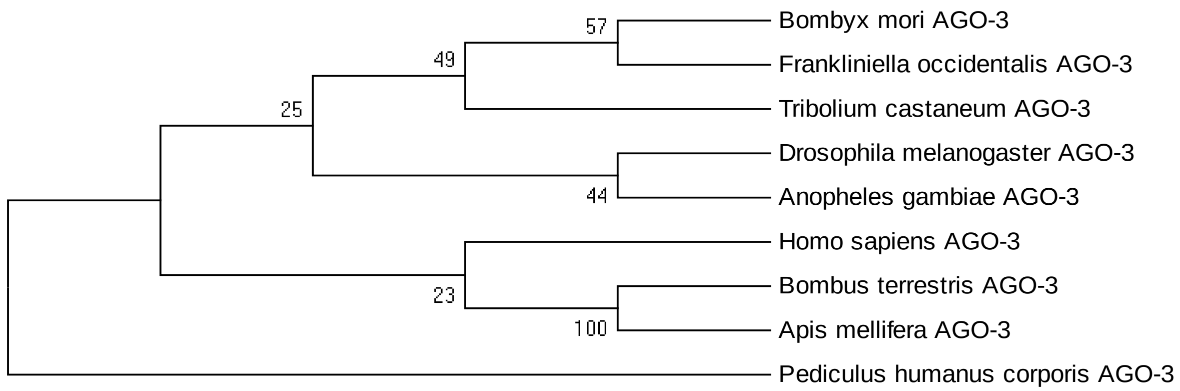

#### 5. Dicer 1, 2

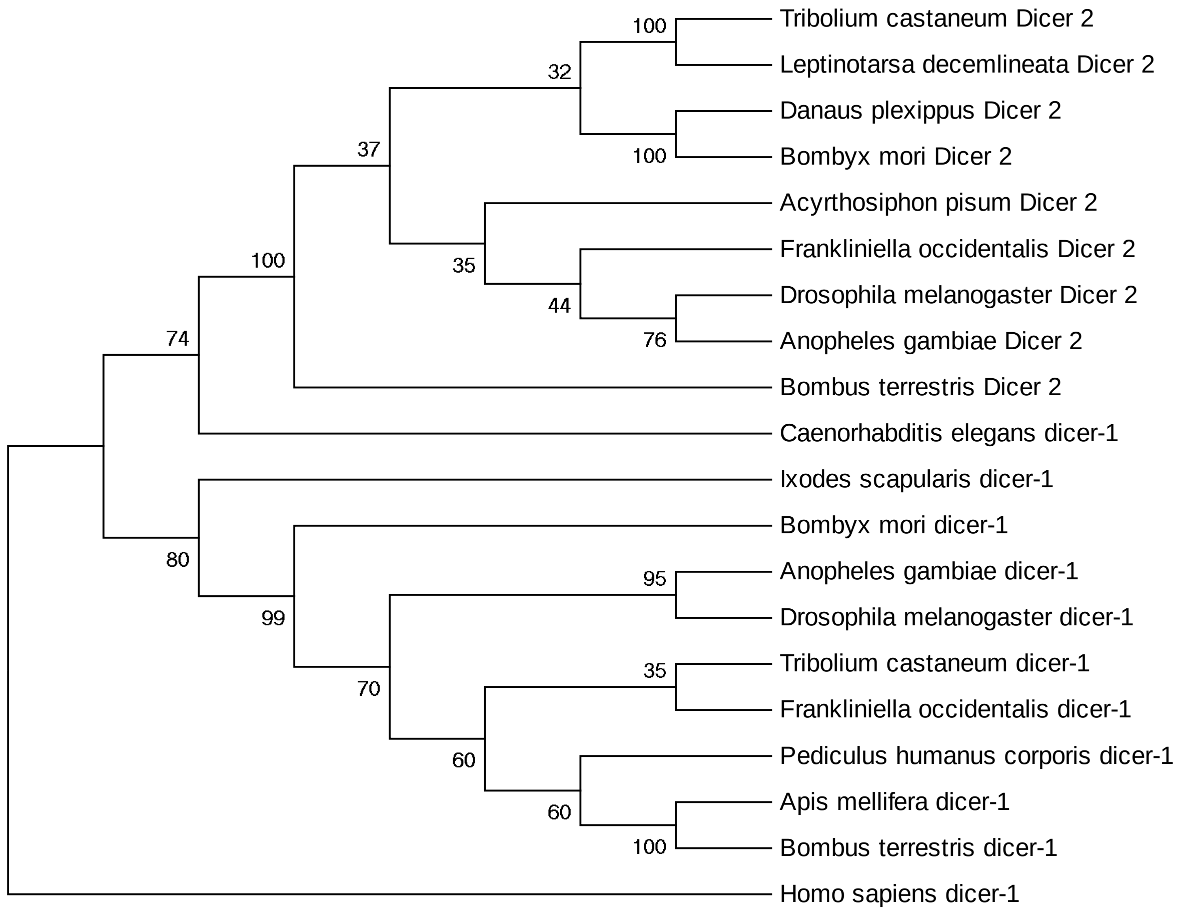

#### 6. Dicer 1

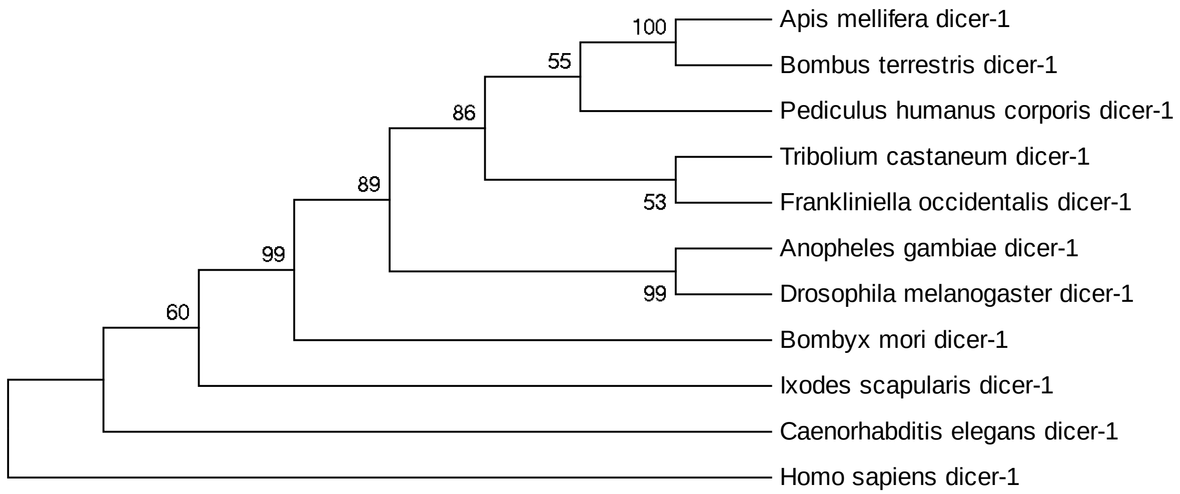

#### 7. Dicer 2

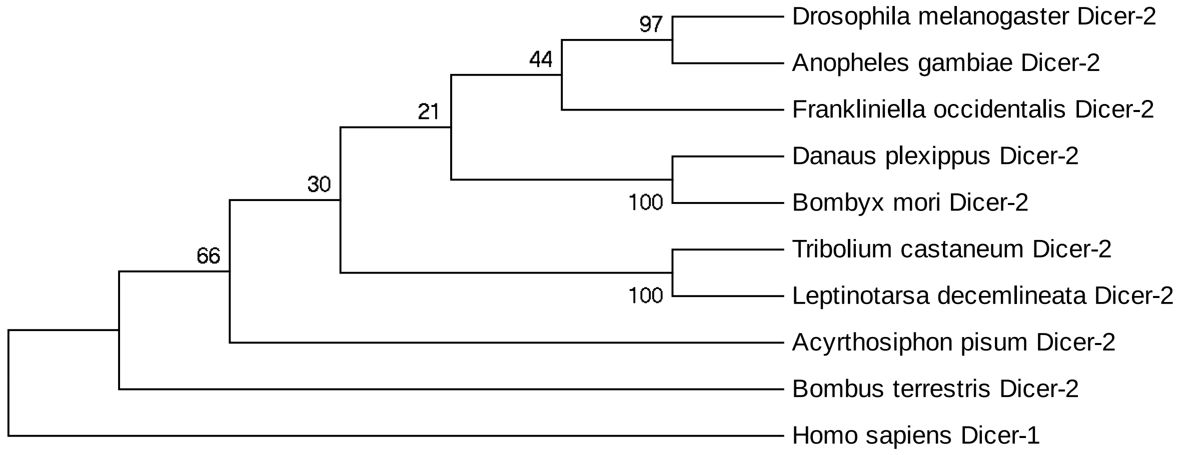

#### 8. **Loquacious and r2d2** (maximum likelihood analysis)

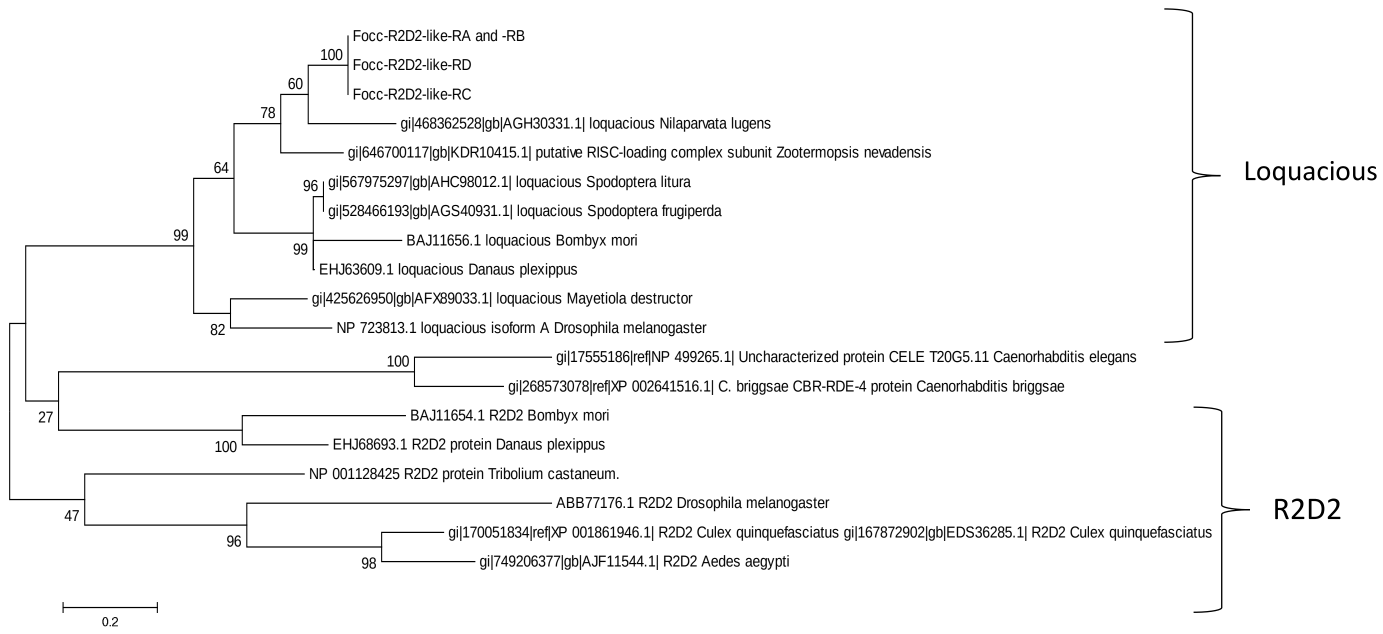

### 9. Embryonic and Post-embryonic genes

#### 9.1 *Wnt* Signaling Pathway

*Contributed by Iris Vargas Jentzsch and Kristen A. Panfilio*

##### 9.1.1. Abstract

The Wnt pathway is a signal transduction pathway with fundamental regulatory roles in embryonic development in all metazoans. The emergence of several gene families of both Wnt ligands and Frizzled receptors allowed the evolution of complex combinatorial interactions with multiple layers of regulation [1]. Wnt signaling affects cell migration and segment polarity as well as segment patterning and addition in most arthropods [2]. Surveying and comparing the gene repertoire of conserved gene families within and between taxonomic groups is the first step towards understanding their function during development and evolution.

Here we curated gene models for the main components of the Wnt signaling pathway, and confirmed their orthology by phylogenetic analysis. We found 9 Wnt ligand subfamilies, three Frizzled transmembrane receptor subfamilies, the co-receptor *arrow*, and the downstream components *armadillo*/*beta-catenin*, *dishevelled*, *arrow*, *axin*, and *shaggy*/ *GSK*-3. All of these genes, with the exception of the Frizzled family (three *fz-2* paralogs), were present in single copy in the assembly. Three Wnt genes, *wingless*, *Wnt6* and *Wnt10*, were linked on the same scaffold, reflecting the ancient arrangement of Wnt genes in Metazoa.

The thrips Wnt ligand repertoire is comparable to that of other insects and adds to the observations of reduction in ligand diversity in the lineage leading to insects (compare 9 ligands in thrips with 17 total ligand classes known in other animals). Nevertheless, the proposal of gene losses needs to be done with caution when dealing with draft assemblies from second generation sequencing, which is the case for most recently published genomes.

##### 9.1.2. Methodology

Protein sequences for *Wnt* ligands as well as receptors and downstream components (*armadillo*/ *beta-catenin*, *dishevelled*, *frizzled*, *arrow*, *axin*, *shaggy*/ *GSK*-3) from *Drosophila melanogaster*, *Tribolium castaneum*, *Acyrthosiphon pisum* and *Oncopeltus fasciatus*, were retrieved from NCBI, and used to perform standalone tblastn searches on the *Frankliniella occidentalis* scaffolds with a maximum e-value of $1e^{-10}$. Hits from all species together were ordered by scaffold and start position, and for each group of overlapping or closely adjacent hits from multiple orthologous queries, the putative gene name was identified by blasting back the hit sequence against GenBank, with a taxonomic restriction to Arthropoda accessions. The query sequences with the best hits (lowest e-value) for each gene were then used to identify the model to be curated, by doing a tblastn search into the *Frankliniella* scaffolds from the Blast instance at the National Agricultural Library (https://i5k.nal.usda.gov/legacy_blast). The Blast results were visualized in the Web Apollo instance for *Frankliniella* (https://apollo.nal.usda.gov/fraocc/selectTrack.jsp), where the corresponding automated annotation models were edited. To confirm orthology, we then Blasted the edited *Frankliniella* models back into GenBank. Homology, intron/exon boundary assessments, and protein sequence completeness were identified by manual inspection and correction of protein alignments generated with Clustal Omega (http://www.ebi.ac.uk/Tools/msa/clustalo/).

The numbering (subfamily assignment) for *Wnt and fz* orthologs was assigned based on the corresponding vertebrate homolog (the naming of *Drosophila* orthologs was changed accordingly), based on phylogenetic analyses done at http://www.phylogeny.fr/.

Possible gene loci duplications were identified by performing tblastn searches on the scaffolds using the protein sequences of completed annotation models as queries, and then re-blasting the resulting hit sequences into GenBank for Arthropoda hits.

##### 9.1.3. Results and Discussion

A total of 26 models for the main Wnt signaling genes were curated on the *F. occidentalis* assembly (**Table S9.1**). There were starting automatic predictions for all annotated genes, except for *frizzled-2a –part 1*, because this is one isolated exon, and the rest of the model was found on another scaffold. All automated models were very accurate, even in the absence of RNA-seq support. Only the beginning of the genes proved more difficult to predict, as we had to edit the start codon or add additional start exons in most of the cases. Most of the models were present on individual scaffolds, and three models were split into two parts on different scaffolds (*disheveled*, *Wnt7* and *WntA*).

All Wnt pathway genes, except for the *frizzled* receptor subfamilies, were found as single copy genes. The *armadillo* ortholog had very good RNA-seq support, and strongly conserved sequence compared to homologs. This model, however, contains a non-canonical splice site at the 3' end of the fifth exon. Given that the mapped RNA-seq reads strongly support the authenticity of the splice site, this could potentially be a sequencing error. Similarly, the models for both *disheveled* isoforms have a non-canonical splice site at the 3' end of exon 4, with otherwise strong RNA-seq and orthology support for this splice site.

We identified 9 *Wnt* gene subfamilies in the *F. occidentalis* assembly, all with single copy genes: *wingless/Wnt1*, *Wnts* 5-8, 10-11, 16 and *WntA*. *Wnt16* has so far only been reported in the pea aphid *Acyrthosiphon pisum* [3], the Russian wheat aphid *Diuraphis noxia* [4] and *Oncopeltus fasciatus,* suggesting that the hemipteroid assemblage (clade Acercaria) has retained a Wnt ligand that was subsequently lost within the Holometabola. Meanwhile, phylogenetic analysis of the protein sequences supports our designation of the *Frankliniella* WntA ortholog, which forms a well-supported clade with WntA proteins from hemipteran species and the beetle *Tribolium castaneum,* with Wnt7 a closely related outgroup. Surprisingly, Wnt4 accessions in GenBank for several species (the hymenopterans *Acromyrmex echinator, Harpegnathos saltator, Camponotus floridanus*; the termite *Zootermopsis nevadensis*; and the lepidopteran *Papilio xuthus*) also fell within our WntA clade. However, Wnt4 is not known from *Tribolium* or several other well-characterized species such as *Drosophila*. We therefore caution that these Wnt4 accessions were likely wrongly named during automated orthology assignments, and that they in fact represent additional WntA orthologs.

Three *Wnt* models were clustered on scaffold 178, showing the same gene order previously observed in *Tribolium castaneum* and *Drosophila melanogaster*: *wingless*- *Wnt6*-*Wnt10* [6]. The transcriptional orientation of *Wnt6* was inverted with respect to the other two genes, as is the case in *Tribolium* [5]. This gene arrangement still resembles the arrangement of *Wnt* genes observed in *Nematostella*, reflecting the ancient arrangement of *Wnt* genes in metazoans [7].

The *Wnt* gene repertoire observed in *F. occidentalis* is very similar to the one in *Tribolium castaneum* [6], both having 9 Wnt subfamilies, with the difference that *Tribolium* lacks *Wnt16*, while *Frankliniella* lacks *Wnt9*. The general trend is for insects to have fewer Wnt subfamilies compared to more basally branching arthropods like the water flea *Daphnia pulex* and basal metazoans like *Nematostella vectensis*, with 12 and 13 *Wnt* gene subfamilies, respectively [5]

Of the four ancient *frizzled* (*fz*) receptor subfamilies that are expected to be present in the common ancestor of arthropods [8], we found three in *F. occidentalis*: *frizzled*, three *frizzled-2* paralogs, and one *frizzled-4*. Two of these models, *frizzled*, and *frizzled-2a*, were split across two scaffolds, but the complete coding sequence could be found. For the third *frizzled* ortholog, *frizzled-2c*, the N-terminal region of about 70 amino acids was missing, reflected in a small gap (default size of only 50 bp) in the genome assembly in this region. *fz3* is also missing in *Tribolium castaneum* [9] and *Oncopeltus fasciatus* [10], which have three ( *fz*, *fz2* and *fz4*) and two *fz* families (*fz* and *fz2*), respectively. This would suggest that the loss of *fz3* preceded the split giving rise to Holometabola, but *Drosophila* does have a *fz3* ortholog with highly divergent sequence [11], suggesting that “missing” genes are in some cases not recognized due to the rates of molecular evolution relative to the taxonomic sampling currently available.

**Table S9.1.** Positional information for the annotated genes. Incomplete gene models are marked with an asterisk (*).

| Gene | Scaffold: start..end | Locus length (nt) | Protein length (aa) | Number of CDS exons |
|---|---|---|---|---|
| *axin* | Scaffold44:1704986..1711094 | 6,109 | 901 | 11 |
| *armadillo* | Scaffold972:16955..24338 | 7,384 | 817 | 12 |
| *arrow* | Scaffold115:195674..221689 | 26,016 | 1642 | 19 |
| *dishevelled-RA* | Scaffold47:1281711..1294311 | 12,601 | 765 | 15 |
| *dishevelled-RB* | Scaffold47:1281711..1294311 | 12,601 | 625 | 13 |
| *frizzled -part 1 of 2* | Scaffold2979:1031..2872 | 1,842 (partial) | 432 (partial) | 2 |
| *frizzled -part 2 of 2* | Scaffold95:720586..722997 | 2,412 (partial) | 171 (partial) | 1 |
| *frizzled-2a -part 1 of 2* | Scaffold281:127057..127287 | 231 (partial) | 77 (partial) | 1 |
| *frizzled-2a -part 2 of 2* | Scaffold151:643883..646320 | 2,438 (partial) | 628 (partial) | 3 |
| *frizzled-2b* | Scaffold169:473303..478491 | 5,189 | 689 | 4 |
| *frizzled-2c** | Scaffold508:48411..50728 | 2,318 (partial) | 579 (partial) | 3 |
| *frizzled-4* | Scaffold353:41436..50224 | 8,789 | 344 | 2 |
| *shaggy-RA* | Scaffold47:670824..677545 | 6,722 | 504 | 8 |
| *shaggy-RB* | Scaffold47:670824..678020 | 7,197 | 504 | 8 |
| *wingless* | Scaffold178:79250..116897 | 37,648 | 426 | 3 |
| *Wnt5* | Scaffold63:925819..935398 | 9,580 | 292 | 5 |
| *Wnt6* | Scaffold178:68659..69825 | 1,167 | 276 | 4 |
| *Wnt7 -part 1 of 2* | Scaffold2000:1837..5876 | 4,040 (partial) | 84 (partial) | 2 |

| | | | | |
|---|---|---|---|---|
| *Wnt7 -part 2 of 2* | Scaffold58:125442..130075 | 4,634 (partial) | 304 (partial) | 3 |
| *Wnt8-RA* | Scaffold22:1034670..1050363 | 15,694 | 402 | 6 |
| *Wnt8-RB* | Scaffold22:1034670..1050363 | 15,694 | 401 | 6 |
| *Wnt10* | Scaffold178:22725..28783 | 6,059 | 468 | 7 |
| *Wnt11* | Scaffold62:66421..68509 | 2,089 | 408 | 4 |
| *Wnt16* | Scaffold8:1141931..1213443 | 71,513 | 420 | 7 |
| *WntA* | Scaffold1:1219341..1266713 | 47,373 | 358 | 6 |
| *wntless* | Scaffold19:514011..515993 | 1,983 | 1088 | 2 |

#### 9.2 Molting and Metamorphosis

*Contributed by Aaron Baumann*

##### 9.2.1. Juvenile hormone esterase (JHE)

Numerous carboxylesterase genes were identified and annotated. Of these, none were given the JH esterase title. BLAST searches using *D. melanogaster* JHE protein sequence pull up 56 putative hits at threshold e<0.00001. Three carboxylesterase annotations meet a "diagnostic" criterion of containing the GQSAG motif characteristic of JH esterase proteins: **12120**, **4777**, and **4770** in which A is replaced with S (GQSSG). It is not unreasonable to propose that any of these could function as a JHE but this notion needs to be experimentally tested. *D. melanogaster* has two JHE proteins, JHE and JHEdup, suggesting independent duplications during insect evolution. It is worth noting that JHEdup contains the motif GHSAG rather than GQSAG.

Phylogenetic reconstruction identifies **13106** and **3184** as putative JHE orthologs.

A sequence identity matrix using trimmed protein sequences (**Fig. 9.1**; top diagonals represent gaps; bottom diagonals represent % identity) suggests greater identity between *D. melanogaster* JHE and **3184** and **12120**. There was more than 45% identity between *D. melanogaster* JHE and 3148 or 12120 vs. ~39% for 4777 and 47770. Likewise, *Rhodnius*, *Glossina*, and *T. castaneum* JHE each share highest sequence identity with **3184** and **12120**, relative to other putative *Frankliniella* JHE proteins. Thus, **3148** and/or **12120** (which also shares the GQSAG motif) seems the likeliest candidate JHE.

**Figure S9.1**. Amino acid identity matrix for putative juvenile hormone esterase sequences (Maker transcript IDs without FOCC prefix) located in the F. occidentalis genome.

| | | 1 | 2 | 3 | 4 | 5 | 6 | 7 | 8 | 9 | 10 | 11 | 12 | 13 | 14 | 15 | 16 |
|---|---|---|---|---|---|---|---|---|---|---|---|---|---|---|---|---|---|
| Dm_JHEdup | 1 | | 1 | 1 | 1 | 2 | 2 | 1 | 1 | 1 | 2 | 2 | 1 | 1 | 1 | 1 | 3 |
| 3184 | 2 | 42.15 | | 0 | 0 | 1 | 1 | 0 | 0 | 0 | 1 | 1 | 0 | 0 | 0 | 0 | 2 |
| 12120 | 3 | 39.67 | 41.32 | | 0 | 1 | 1 | 0 | 0 | 0 | 1 | 1 | 0 | 0 | 0 | 0 | 2 |
| 4770 | 4 | 38.84 | 35.95 | 39.67 | | 1 | 1 | 0 | 0 | 0 | 1 | 1 | 0 | 0 | 0 | 0 | 2 |
| 4777 | 5 | 35.54 | 37.60 | 41.74 | 50.00 | | 2 | 1 | 1 | 1 | 2 | 2 | 1 | 1 | 1 | 1 | 3 |
| Tcas_JHE_iso1 | 6 | 47.11 | 43.80 | 40.91 | 40.91 | 38.02 | | 1 | 1 | 1 | 2 | 2 | 1 | 1 | 1 | 1 | 3 |
| Gmors_JHE | 7 | 52.07 | 42.15 | 41.74 | 38.43 | 39.26 | 53.72 | | 0 | 0 | 1 | 1 | 0 | 0 | 0 | 0 | 2 |
| Dmoj_168370 | 8 | 54.13 | 43.39 | 45.04 | 38.43 | 38.02 | 55.37 | 70.25 | | 0 | 1 | 1 | 0 | 0 | 0 | 0 | 2 |
| Cu_quinque_JHE | 9 | 49.59 | 42.15 | 42.98 | 37.60 | 40.50 | 58.68 | 60.74 | 61.57 | | 1 | 1 | 0 | 0 | 0 | 0 | 2 |
| Dm_JHE | 10 | 51.65 | 45.45 | 45.87 | 39.26 | 39.26 | 52.89 | 66.94 | 79.75 | 59.92 | | 2 | 1 | 1 | 1 | 1 | 3 |
| 17251 | 11 | 42.98 | 41.74 | 43.39 | 39.26 | 41.32 | 50.41 | 47.11 | 47.11 | 50.00 | 47.93 | | 1 | 1 | 1 | 1 | 3 |
| Rhodnius_JHE | 12 | 44.63 | 45.87 | 39.26 | 35.12 | 36.78 | 47.93 | 43.39 | 50.00 | 46.28 | 48.76 | 43.39 | | 0 | 0 | 0 | 2 |
| 11633 | 13 | 42.15 | 45.87 | 47.11 | 41.32 | 44.21 | 45.87 | 45.04 | 46.69 | 45.04 | 48.76 | 50.83 | 43.80 | | 0 | 0 | 2 |
| 7334 | 14 | 42.56 | 44.63 | 45.87 | 43.80 | 41.74 | 42.56 | 43.80 | 45.87 | 45.04 | 46.69 | 44.21 | 47.93 | 59.50 | | 0 | 2 |
| 17068 | 15 | 42.15 | 43.39 | 48.76 | 41.74 | 44.63 | 47.11 | 45.04 | 46.69 | 46.28 | 46.69 | 48.76 | 45.87 | 54.13 | 50.41 | | 2 |
| 13106 | 16 | 42.98 | 45.45 | 42.98 | 40.08 | 40.91 | 48.76 | 46.28 | 45.87 | 47.52 | 48.76 | 44.21 | 47.52 | 49.17 | 46.69 | 51.24 | |

##### 9.2.2. bHLH PAS and bHLH Myc family member proteins

In total, 45 orthologs were annotated for the following bHLH-PAS/myc family members (others were already dealt with by other team members, including the E(spl)-bHLH orthologs): 48 related to 3; absent MD neurons and olfactory sensilla (amos); achaete (ac); atonal (ato); atonal-like; clock; clockwork orange (putative); cycle; daughterless; deadpan; dimmed; dimmed-like;

dysfusion-like; extra macrochaetae; grainyhead (putative); hairy; hairy (putative dup); Helix Loop Helix Protein 3B; helix-loop-helix protein 11; HLH54F; knot; knot-like (putative); knot-like (putative); max-interacting protein (putative); max-like protein; mitf-like; MLX interacting protein (putative); mnt; Myc; nautilus (putative); Olig family (oli); PAS domain-containing protein; period; scleraxis; similar; single-minded; spineless; Sterol regultory element-binding protein (SREBP/HLH-106); tango (tgo); target of Pox-n (tap); taxi; net; trachealess; twist (putative; tcf15-likehomolog);usf-like1. The **nautilus** annotation; FOCC0016897 may not be complete since this sequence occupies the 5'-most space on the scaffold and there may be additional coding or noncoding exons that were not resolved.

##### 9.2.3. bHLH super family protein

***Clock:*** the *Clock* annotation is split across two gene models, FOCC003627 and FOCC003628, which are separated by a run of NNNNN. Names were therefore given as clock (partial) to each model. According to alignments with *Drosophila melanogaster* clock,

**In addition to several gene losses (or independent gains in more diverged insects), there were several duplication events within this gene super family:**

***E(spl)-bHLH:*** three Enhancer of split paralogs were identified: FOCC004628, 4632,and 4635. I included a fourth protein, tom (FOCC004629), which is also a member of the enhancer of split complex but does not share sequence identity with the E(spl)-bHLH proteins. Shown below are *F. occidentalis* E(spl)-bHLH protein sequences aligned against Drosophila melanogaster E(spl)mBeta-HLH sequence. 4628 and 4632 are likely products of the most recent E(spl)-bHLH duplication in *F. occidentalis*.

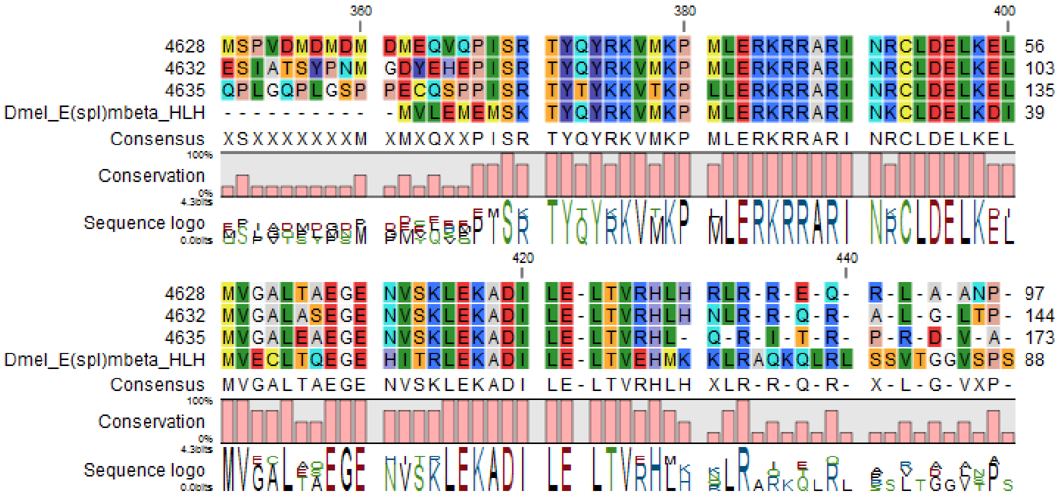

***hairy (h):*** Two *hairy* orthologs were identified, **13163** (8 introns) and **1872** (2 introns). **1872** is likely a retrotransposed copy of **13163**, as evidenced by its paucity of introns, and was thus annotated as “hairy (dup).” A multiple alignment (below) suggests that the *hairy* ortholog in *Drosophila* is likely the direct ancestor to hairy (dup) in *F. occidentalis.* The ancestral paralog is either lost in Drosophila or annotated with a unique identifier (as is often the case with *Drosophila* paralogs; see *FTZ-F1* and *HR39*, *Met* and *gce*, etc.). The ancestral paralog is thus either lost in *Drosophila* or the homolog is annotated with a unique identifier (such as the case *FTZ-F1* and *HR39* in **Boulanger et al., 2012**).

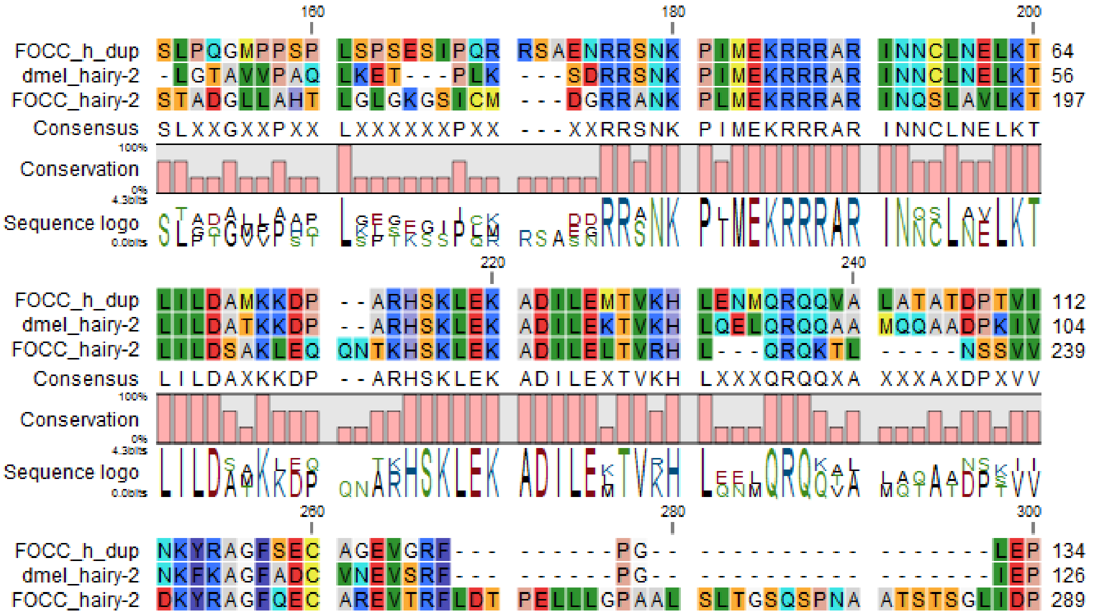

***dimmed:*** two annotations were created that are presumed paralogs of the dimmed bHLH proteins: *dimmed* (14220; 4 introns) and *dimmed-like* (12611; 5 introns). Below shows an alignment of the HLH region of these proteins aligned with *D. melanogaster* dimmed.

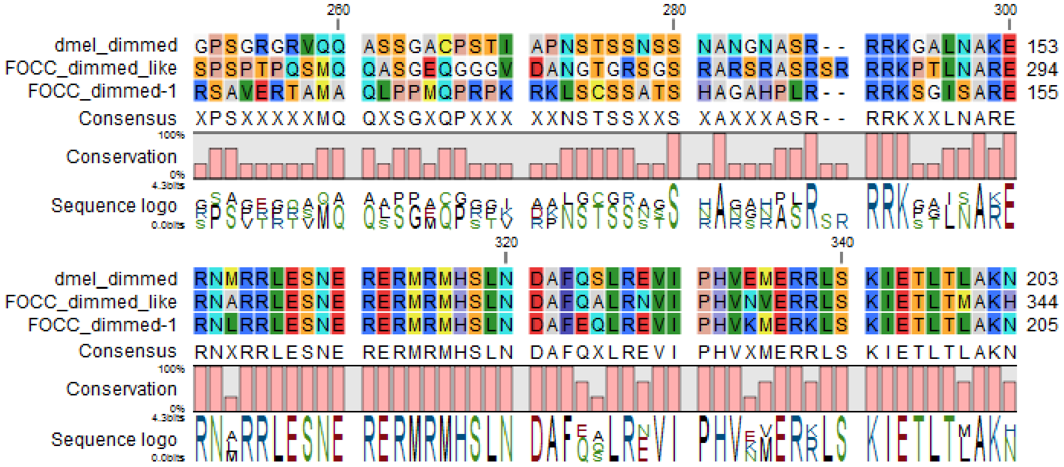

***Knot (syn. collier):*** *knot* (7498, 7499 models merged) and *knot-like* (7501).

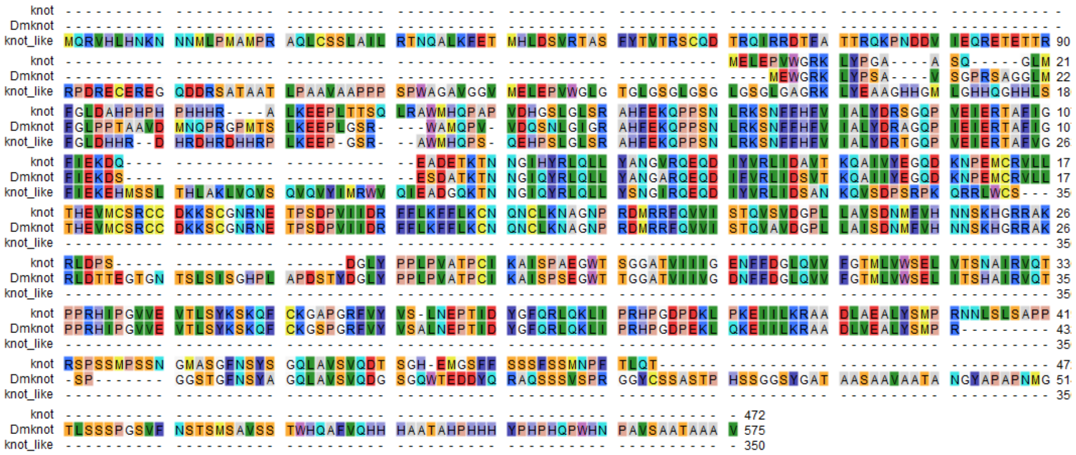

### 10.Cuticular Proteins

*Contributed by Andrew J. Rosendale, Andrew Rosselot, and Joshua Benoit*

#### 10.1. Results

Sequence motifs that are characteristic of several families of cuticle proteins (Willis, 2002) were used to search the genome of *Frankliniella occidentalis* for putative cuticle proteins. 101 genes were identified, analyzed with CutProtFam-Pred, a cuticular protein family prediction tool described in Ioannidou et al. (2014), and assigned to one of 7 families (CPR, CPAP1, CPAP3, CPF, CPCFC, CPLCP, and TWDL) (**Table S10.1**). Many of the genes (~40%) were arranged in clusters of 3 to 5 genes (**Table S10.2**) that were primarily type specific. However, the sizes of gene clusters were smaller than those observed in other insects, which are typically 3 to ~20 genes in size. Additionally, a larger portion (50-70%) of cuticle proteins is typically found in clusters in other insects. Clustering of these genes could allow for the coordinated regulation of cuticle proteins and thereby facilitate the development of insecticide resistance.

As with most insects, the CPR RR-1 (soft cuticle), RR-2 (hard cuticle), and unclassifiable types, constituted the largest group of cuticle protein genes in the *Frankliniella* genome. The number of genes in the protein families CPR, CPAP1, CPAP3, CPCFC, and CPF were similar to the number in other insect (Willis, 2002). However, the 10 genes in the TWDL family was greater than that found in most insect orders, and is reminiscent of the expansion of this family observed in Diptera (**Fig. S10.1**).

**Table S10.1. Number of genes identified as putative cuticle proteins per family in the genome of *Frankliniella occidentalis***

| CPR[a] | | | | | | | | | |
|---|---|---|---|---|---|---|---|---|---|
| RR-1 | RR-2 | Uncl | CPAP1 | CPAP3 | CPF | CPCFC | CPLCP | TWDL | Total |
| 18 | 19 | 27 | 14 | 6 | 3 | 2 | 2 | 10 | 101 |

[a]Sequences that scored above the assigned cutoffs for the RR-1 and RR-2 models were classified as the corresponding type, whereas sequences with scores below the assigned cutoffs but above 0 were characterized as “unclassified” (Uncl). For more information, see Ioannidou et al. (2014).

**Table S10.2 Clusters of genes coding cuticle proteins in the genome of *Frankliniella occidentalis***

| | Scaffold # | # Genes | Family | Length (Kbp) | Density (Kbp/gene) |
|---|---|---|---|---|---|
| 1 | 127 | 5 | TWDL | 80 | 15.9 |
| 2 | 25 | 5 | CPAP3 | 149 | 29.8 |
| 3 | 322 | 5 | CPR RR-1 | 97 | 19.5 |
| 4 | 52 | 5 | CPR RR-2 | 111 | 22.3 |
| 5 | 111 | 4 | CPR RR-1/CPR Uncl | 108 | 27.0 |
| 6 | 6 | 4 | CPR RR-2/CPR Uncl | 47 | 11.6 |
| 7 | 13 | 3 | CPR RR-2 | 52 | 17.2 |
| 8 | 2 | 3 | CPF | 26 | 8.6 |
| 9 | 47 | 3 | TWDL | 31 | 10.4 |
| 10 | 94 | 3 | CPAP1 | 155 | 51.5 |

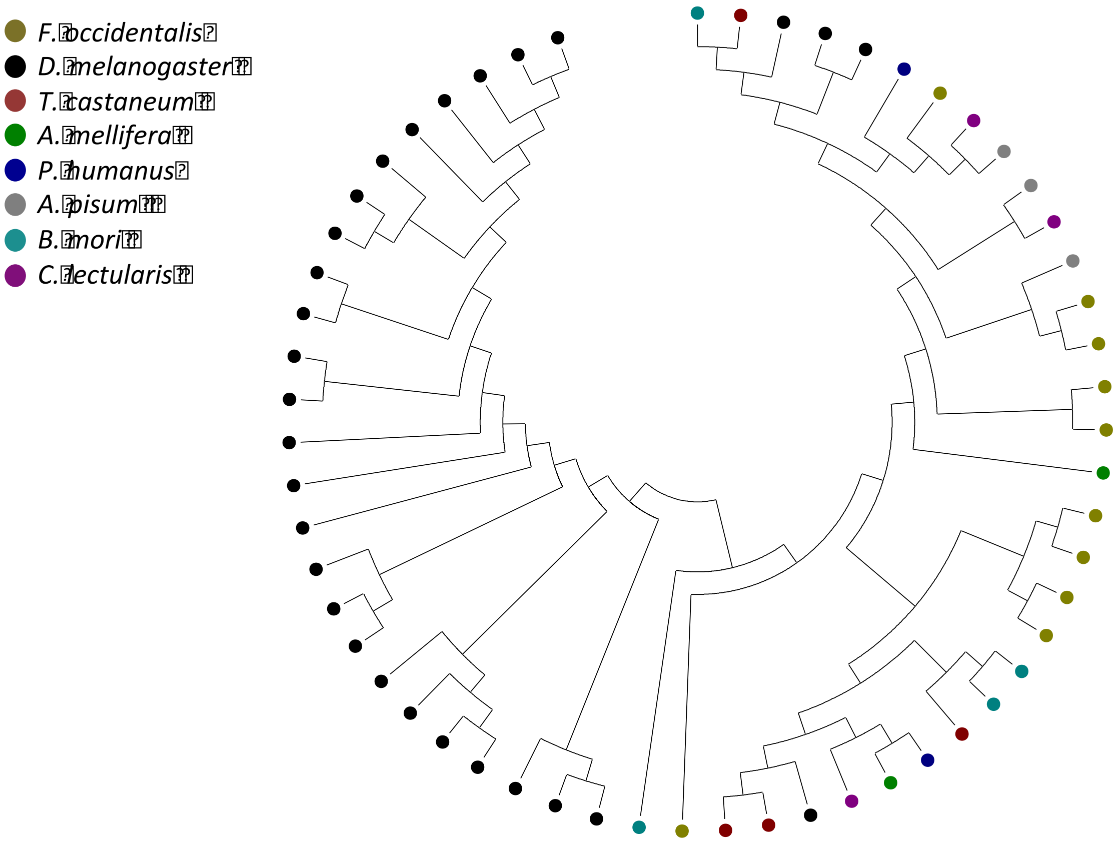

**Figure S10.1.** Phylogenetic tree demonstrating relationship of TWDL genes from *Frankliniella occidentalis, Drosophila melanogaster, Tribolium castaneum, Apis mellifera, Pediculus humanus corporis, Acyrthosiphon pisum, Bombyx mori, Cimex lectularis*. *F. occidentalis* showed a greater number of TWDL genes than other insects, with the notable exception of Dipterans such as *D. melanogaster.* The tree was constructed using the neighbor-joining method in MEGA6 with Poisson correction and bootstrap replicates (2,000 replicates).
