## Additional file 4 for "Genome-enabled insights into the biology of thrips as crop pests"

**Additional file 4: Chemosensory receptor protein sequences of *Frankliniella occidentalis* as part of the FOCC Genome Paper**

*Contributed by Hugh Robertson*

**All *F. occidentalis* chemoreceptor proteins in FASTA format.** Gene/protein names are followed by suffixes, which are not part of the name, indicating issues with the model: C – C-terminal exons/regions missing in gaps; F – assembly repaired; I – internal exons/regions missing in gaps; J – model joined across two scaffolds, N – N-terminal exons/regions missing in gaps, P – pseudogene

**85 FoccOrs**

>FoccOrcoJ

MGQKNQYRGLVADLMPNIRAMQITGLYCMEYHAENSAMQRLMRKAYSLFHVTMMTLGYATLVAFLLTESYNVEDWAAHTVTTLFFLHSLCRTFWFMSNTKRFYRMLDCWNTTNSHPMFAENDARHHQQTLQRQRRLLMQLIALTVVSVMCWIGITFVAQPVRNVPDPENENATITVEVPQLMVYWLLVMLGQANMYDVLLANVVVHACGQLRHLKEILRPLMELSAAFSPASGIAPSNGKIFTTQAVFSAKHNLVMPDMMDLDYRNMFNNARGQANLIQGQLPPVSVNGDRPPSVEDVNPLPMTNTLTVAKNTDSMAVKEEVFVRSAIKYWVERHKQLVKFVSYISLAFGGELLLHMLNATVSDFDVFACSTAGYVLYSMAQIYLFCIHGNELIEESSTVMEAAYSCQWYDGSEEVKTFVQIVCQQCQKNLVVSGARFFTVSLDLFASVLGAMVTYFMVLIQLN

>FoccOr1

MRLIEWMSWHCLEAGPLPRLVFDVMSSKLSGIFMGLCTAFVCASGMDRLGSFETAKVAIMNISSCYSANASLAVVRQGRLSSVLQELQSLVQEAEAWPGDKKWAERTLRGVAMLSTMTVSFAVLVVASLCADMGLADDPQLSLWPFVPHAGPWGAPCGRLFWGCWTLLAVFTICYVVASFWLCAAAAAGLHDALGLRLLAPGGATAEEVRDVVATHRKLRLAVLELTEYAAGKLVHVLSSSFCSSLFATVLVLGGDASAATLSLLVPAFVVFLPLAHYSQEMSDASLSLARSAYHASTDGVEHREARALLLVMLAAAKTPALRCRGLGRFGRASMRHVFNQFYTAVNTLGSKE

>FoccOr2F

MRLSEWMSWLSLETGPLQRLLFDVLSSRLSGMFMGLCAVFMTGTALQRLDSLETARAPISNLSSCYSSSACLAVIRQGRLSALLLELKSLVQAAEAGRGEKKWTDRTLRGVTMFATVTVPMAVLVFASLCADMLLADEPQLPLWPFVPHAGAWGVSCGRMFWGCWFLLAVCSVFYTVTALCLCAAAAAGLHDALALRLLVQGGATPEEVRDVVRIHMHLRSTVLDLTDYFAGNLVHILASSFCNSLFATVQVVGGDANATTLALLIPVVNVFLPLSHFSQKMSDASRSLARSAYHAATDGVELREARALLLVMLAAGRPPALRCRGLGRFNRASARNVFRQFYAAVNMLGPKH

>FoccOr3I

MRLSVHLSWLCLEQHPLVWVLRAVIKSRLRHETLNLVDFFAANLVHILASSFANTFLAVVQLLINDVTNTTYVMLLPVVVIFLQLSYFSQELSDSSLTLKRCAYAAATGGAVTVAEARALQLVILAASRPPALGCRGLGRLSLANAGRALRQAYSLVNVLSSKL

>FoccOr4C

MQLSVYLSWLCLEQHPLLRVLRAVLKSRLPMTLMGLSSIFEGTVALLQTTARTTVPAVSVASDFYSAYSCCAVACQPRFWNILSEVKRLVQVMEDIPGEKVWADSTCGHIAIVCKGFFATAVLMLGCVLADMLLSDEPMLSLWPFVPHVGSWGVLVGRLFWGISTTVCGATLSHVLVTLMCVTTTVTGMHHALAQRLLDSTGMTPEVVRDVVRLHQQLRHETLNLVDFFATNLVHILACSFVDAVLSVVQ

>FoccOr5NC

RHALMQTTARTAVPAVSVASDLYSAFSCSVVARQPRFWNILSEMRRLVQVMEDIPGEKAWVDSTRGHIAILSKGFFANAFLVLVCVLADMLLSDKPMLSLWPFVPHAGSWGVLVGRLFWGTSATVCVATLSHVLVTLMCVTTTVTGMHHALARRLLDSTGMTPEVVRDVVRLHQQLRHETLNLVDFFAVNLVHILASSFANTFLAVVQ

>FoccOr6N

LRHETLNLVDFFAANLVHILASSFANTFLAVVQLLINDVTNTTYVMLLPVVVIFLQLSYFSQELSDSSLTLKRCAYAAATGGAVTVAEARALQLVILAASRPPALGCRGLGRLSLANAGRALRQAYSLVNVLSSKL

>FoccOr7F

MQLSVYLSWLCLEQHPLLRVLRAVLKSRLPMTFMGLSCILEGIVALMQTTARTTLPAVSAASDFYSAYSCCVVARQPRFWNILSEVKRLVQVMEDIPGEKVWADSARRNIAVIFKSVFVTAVVMLGCILADMLLSDEPMLSLWPFVPHAGSWGLLVRSMFWGAASTVCCVTLFHVLVPLICVTTTVTAMHHALAQHLLDSTGKTPEVVRDVVRLHQQLRHETLNLVDFFATNLVHILACSFVDAVLSVVQLLINDVTNTTYVLLLPVVVIFLQLSYFSQELSDSSLALKRCAYAAATGGAVTLAEARALQLVILAASRPPALGCRGLGRLSLANAGRALRQAYSMVNVLSSKF

>FoccOr8

MRLSVHLSWLCLEQRPLLRVLRAVLQSGLPDRFMGFCCILEGVYALLQSSARAAMPPVSSLSDLYSAYACYAVTRQSRFWNILSEMKRLVQIMEDTPGEKVWIDSTRGQIAVIYRGNCSNATMIMVAVLADMLLSDHPLVSLWPFVPHAGSWGVLVGRLFWGAGVFVTVVSIWYMSVILTFVTATAAGMHMALAQRLLDSTATPGLGVGDVVRDVVRDVVRDVVMVHQQLRHETLNLVDFFAANLVHILASSFVNTFLSVVQVLNGDVTITTYAFAFPPLIVVIFLQLSYFSQELSDSSLALKRSAYAVATSGAVTLAEARALQLVILAADRPLALNCRGLGRLSLANAGRALRQFYSVVNVLGSKF

>FoccOr9

MRLSVYFSWMCLEGGHLQRFMLKAFRSVLLWLYLGLCVVLAWCKVLQKSVSFQTTTKTLSCFSSCYSGFVALTVIRQTRFWAMLHEIKSLVQDLEEDEAAEKVWVARTRRIIVVFSCQTVTLSIFTAASVVLHFTISGDDTMVYLWPFVPYAGSWGVYVARLVTGVGVSLIVLTISFVVITLGCTTSAATGMHHALAQGLLLEAATPEVVGNAVALHQRLRRVTLDMTDFFAANLAHVMASSFFHSSAALIQVLASGVMTSTTVFLLLRVVVVFLQLSYLSQELSDSSLLLQGSAYRAATSRGTGPAEARALVLVMLAASRTPALSCKGLGRLSLASAGRAFRQLYSVVNVLKSRY

>FoccOr10F

MRLRVYFSWMCLEGGHLQRFIRKVLRSRLLWLYLGLCVVLSWSKMLQTFASFQATMKTLSGLSSIYSGFVAFRVVRQTRFWAMLHEMKSLIQVLEEEDGAAEKVWVARTRRTIVVFSCQSITLSIAMGASIVLHFTFSGDESMVFLLWPFMPYTGSWGVYVARLVTGVCVLLILLSILFVVIVLGCTTAAATGMHHALAQGLLAAATPEVVGNAVALHQRLRTVTLDMTDFFAANLAHLMASSFFHSSAALIQVLASGVITSTTVFQLFRVVGVFLQLSYLSQELSDSSLHLGGSAYRAATGSGTGLEEARALVLVMLAASRRPALSCKGLGRLSLASAGHAFRQLYSVVNVLKSRY

>FoccOr11C

MRLSVYFSWMCLEGGHLQRFIREVLRSRLLWLYLGLCVVLSWCKVLQTFASFQVAMKALSGFSSCYSGFVAFTVVRHTRFWAMLHEMRSLIQVLEEDGAAEKVWVARTRRTIVVFSCQSITLSMSMAASIAFHFTFSGDDTMVFLWPFVPYAGSLGVYVARLVTGVGVLLILLGILFVVIVLGCTTAAATGMHHALAQGLLAAATPEVVGNAIALHQRLRNVTLDMTDFFAANLAHLLASSFFHSSAALIQ

>FoccOr12N

PLWLFLRLCIGLAVCKVLQESDSYRAAEKPLLALTSCYSANVALTVIRQMRYWNILREIKCLAEVLEDDLAEEVWVARTRRNIILFSSINLTFGISLGVSITLRFIISGDNTLLLLWPFVPYTGSWGLTIAQLVTGVCCPLLSITISIVVTALGCTTSTVTGLHHALAQCLIATATPEAVRNGAKLHQRLRRVTLDLTEFFAGNLAHVMASSFSHSSLAIVQVIASQEMTSTTFFQLFRVLVVFLWLSFLSQELSDSSLRLQRSAYLAATGSGTALAEARALNLVMLAASRPPALTCKGLGTLSLSSAGHVLRQLYSVINVLKKQN

>FoccOr13

MRLYVYLSWMCLGEGPLQRFVLRIFKSRLFLGLPASGTIATAVEQTSSLQTATRPISILGSCYSAFASLAMMRRERFWNMMHDMKSLVQIMEAAPGEQAWVAGTRKSILRFAAGNLMLWLATVGSALVGAVLHPERVVPSWPFVPLAGTLGFYIARLSYIVSVFVITVPIGFVVTSLGCVTATATGLHHGLAQSLDTATSPEKVRDVVKVHQQLRHITVDLTDLCADNLGHILVSSFCHSLIAILQVLANDTTSFTFAQLLRVVIVFLQLSHLSQELSDASLVLQTAAYQAASGGSVSLSEARALHLAMLTSSRQPALTCKGLGRLSLANAGKAFNHLYSVVNMLGPKVRGP

>FoccOr14

MRLSGYLSWLCLEEGRVRRCLFRVCTARTTALYMGACVLIAATIAVRKSTSVQAAMMPITTLSSCYSGFVCLSIFRDVRFWTILQELKRLVEVMEADVCEEVWVARTRQRIVAFSLETMALAMVTAVNVLAHSFRSDDSLLPLWPFVPYAGPLGFPLARLFWGCALCTILLSINYLVIALGCATSTATGLHHALAQRLTSTASSEVLRDAVIRHQRLRHLVMDLTDFFAGQLAPVMASSFCHSLFATLQVLSNDLTTTTYTQILRVLVVFIQLSYLSQELSDASLSLHTAAYHVASSDSVSLPEERALILVMLTASRAPALRCRGLGQLSLSSAGKVLRQFYSVVNVLGGQRY

>FoccOr15

MVRLSDYLSWLCVESGRGRLHRIVLTMLRSRLSGFFAGLGCVLQFCVVLRQFETIQSVTNPICNLSSCYSGYASLAVMRQTRFWAILHKIRSLVQTMESYPGDKGWEDHTRREIRRFFAGIIVMAVFISGSVWLSLILTGTSILPLWPFVPYAGPFGVYLAWLFYDVVSVLIILGAISFVVTALVCVTCTAAGLHYALAQRIATNETPEVVKDVVLMHQQLRDVARDLITLFDGNLAHIMASSFCNSLFATIQVLANDVTSSTFALLLPVVVIFLQLSHYSQELFDSSLKLQRAAFHAATGGTVAFLEARALNLVILVAGTPPTLHCKGLGQLSLADAGKAFRQFYSTVSVLGPKY

>FoccOr16

MRPSVYLSWLCLEEGRLQRFLFTMFRCAATALFMALCSVSLLAAILLRPETLRADMIPVCVLSSCHSTFMALAMMRGGRLWPILHEMTRLVQVFEDAPGDKVWVARTRQHGVRFCTYQLAMCAFIIPSTIVNMALTGVPFIALWPLQPHAGAWGLRVAHLVSVAVLLVMTVALPFMNMIVTCFTGAATGLHHALARRLDELAEAEAMTPEAVRDLVLAHERLRRLTLDLTDFFATNLTHLLVCPFCNSLVATVELVSAGNAVSVHTIGLLLPVVVIFLQLSYFSQELSDASLALQWSAFRLATGGGVALAEVRALRLVMLAAGRQPALSCKGLGRLSLEAARLAFRQFYSTVNVL

>FoccOr17

MRLSVYLSWVFLENVPVQRFLFKLFTSRVSGMFMGLNSVFMIITALQNSSTLQTVLLPVSNVFDCYSAFTCLSVSRQARFWSILRKILRLVNVMEATLGEKVWMDNTKRIIGMFCATTLSYAIFVFSSLWTELFLSGTSLYISWPFEPYLGSWGEHVGQVYYNFVHLSLVCLCVVWMYTAVGCATCTSAGLHHALAQYLISNVTPRTTENAVKMHQQLRLVTLDMVDFFAGNLAKIMAHSFCNAVIATIQVLANEVSSGTFVNLIAVSLLFTSLCFFSQMLSDASLALQTCAYRAATSGVQLPEAHALALVMLAAGRPPALRCRGLGRLNLAAAGDAFRRLYSVISVLGTKY

>FoccOr18I

MQLHQYLSLLLLETGPLQRSLLRALRSRMIRRGRERAILHEMVGLVRAIEDGPGDKEWVGRARRMNNVFGAVLLTCVTSTVVTVCVDMAVTGEPFFALWPLVPGLGAWGRRAAAVLSAVAVPHIAVGYLYLFVGLVWFISIATWLHHALALRLVSDPCPEVVGEVVAHHRRLRRLALAMTDLFAGNLAYIHAGSAVNSVLVTMELLAHGLTPATVVLLLTLIANFLQLSYFSQELSDASLRLRGAAYRAAVGGSVRLPDAKALALVALTAGRPPAVSFRGLGRLSLANAGTALRQFYSVINVLGPRFS

>FoccOr19NC

LQESDIFTLGQEWVGRADLMSCFSAWWSFAMIRRGRERAILVEMVGLVRAIEDGPGDKEWVGCARRMNNVFAAVLLTCAASAVVTVCVDMAVTGQPFLALWPLVPGLGSWGRRAAAVLSAVAVPHIAVSFLYMFIGLVWFTSIATGLHHALARRLDSDPSTEVVGEVVAQHQRLRRLALAMTDLFARNLAYIQAGSAVNSVLVTME

>FoccOr20NF

VLWLHLGSCVVLTVCNVLQKASPFSVQTTMKQMCSFSSCYSGMVALTVIRQPRFWTILHEMKRLVEVLDDDPSEKVWVVRTRRNIVIFSYQTLTLLVFTAISVSLHLAFSGDDDAIVPDWPFVPYAGSWGVQIAKLLTGLGMLSITLSLSFLVTALGCTPSAATGLHHALAQGLLAAATPRVVGDVVMLHQRLRRASLDLTDFFAGNLAHVMASAFVHSSVAIIQVMNSEVLTSTILLLLLRVLMVFLGLSYLSQELSDSSLRLEGSAFRAATDSKTRLAEARALVLVMLAASRPPALSCKGLGWLSLPSAKEALQQLYSVINVLKSKDFGGKTTKTE

>FoccOr21

MQLTFSLSWLSLEESQLQRFVLLVMRSRLPLSFTILFLAFVCADTVRQSATSVQAILQSINNFFFGYSMLQCVLVGRHPRLWVLMRELRSLVQAIKADPGEKPWVVHTRRTSVVLSAISTTYVVVFYVHMCQDTLFSEEPFVPLWPFVPYAGAWGALFGRLFYGACMLVWIPTSVCVIIALILSLAVPAGLHHALAQRLRVTDAPGRTLDVVQDVVAKHQRLRLVTHGLTRYFSAKLAHIQIASFLTTIFAFISVICGDVNFTTAIFLLPILNDFLQFSYLSQELSDSSASLARSAYHAATGGSVSLPEARALALVMMTASATPALRCRGVGRLSLATAGKAMSKLWKVISLLRDKL

>FoccOr22

MQLSVSLSWLCLEEGQLQRLLLHVLRNRLTLSFTILFGAFVCVDTLRQSATSVQSLLQSISNIGFGYSMLQVVLVVRHPRLWVMLRELRCLVKVIEADPGEKWVVHTKRTIVVLSTVSTTYIVVFFIHMCLDTLFSEEPVVPLWPFVPHAGAWGGLFGRLFYGACMLIWLPPGLCVILALIISTAVPAGLHHALAQRLEFTVASSKALDVVQDVVAKHRRLRLLTHGLTGYFSAKLAHIQCASFISTVFALISVVCDDVSFTMVLLLLPIVNDFLQFSYLSQELSDSSVSLATSAYYAATGGCVSLPEARALALVMRTASTTPALRCRGVGRLSLAAAGEALGKLWKVVSLLRDKL

>FoccOr23

MQLTFSLSWLSLEESQQQRLVFLVMRSRLPLSFAILFLAFVCLDTVRQSATSVQSLLQSTSNFFIGYSTLQCVLVTRHPRLWVLMRELRSLVQVMKADPGEKPWAVHTRRTSVVLSAISTTYIVVLYVQMCLDLLFSEEPFVPLWPFVPYAGAWGALFGRLFYVVCILVWLPTTVCLVVVLILSLAVPTGLYHALAQRLRVTDAPGRTLDLVQDVVAKHRRLRLVTHGLTRYFSAKLAHMQIASFITTIFAFISVICGDVSFTTAVFLFPIVSDFLQYSYLSQELSDSSASLARSAYHAATGGSVSLPEARALALVMLTASATPALSCRGVGRLSLAAAGEALGKVWKVVSLLRDKL

>FoccOr24N

TWVVHTRRTSVVLSAVSTTYIVVLYVQMCLDTLLSEETFAPLWPFVPYAGAWGALFGRLFYGAVWIVWIPNVVCFMVALVLSLAVPTGLHHALAQRLRVTDAPGRTLDLVQDVVAKHRRLRLVTHGLTGYFSAKLAHIQIASFITSIFAFINVLCGDVSFTTAVFLFPIVSDFLQYSYLSQELSDSSASLARSAYHAATGGSVSLPEARALALVMLTASATPALRITGVGRLSLAAAGEAMGKVWRVVSLLRDKL

>FoccOr25N

VTLAFTVLFTAFVCMDILRRPVTSIPALLLSVSHVIVGYAMLQLVFLVRHPGLWAQMRDLSQLVEAIEADPGGKLWVVNARRTIMVLCTVSITYNVFFFTHVTADALFAEEPVVPLWPFVPHAGAWGTHLGRLFYGACLVAWLPPSVVFTVTLILGTAVPAGLHHALARRLDDSVSPSSVRAVVAKHMRLRQLAQGLTDFYSTRLAHIQVASFFYAVFGLINVLEDDVSFTTVNFLFPLVNDFLQFSYLSQELSDSSASLAEAAYYASIGGSVGLHEAHALGLVMRIASSSPALHCRGLGRLSLAAAGNALGKTWKMVSVLRDKL

>FoccOr26

MSLTNYLSWQLQEIRPFQRFLYLVLSSRIFGYWTVLLTLAVAVCIPSYPSLREATLSILWTESFSSTIVGLGLMRRPHFWDMIRRMQELVAAALADDDDDRVGTVGKRSQEKWAVLMRRNVIWSSGALLCVGTMIIVSMWLRVLVTLKPLCAIWPLYPHAGPAAEPFASLFTATFIALPVCCVGFLVAALSFLTCSAAGLHHALAQRLTESRAPKDTYRVIAMHQKVRQLSIEFTEFFDGSMAQLMAGPIGNSLLATLQVLANDVDVNTFSQIFVVVVTLLQVSYLSQELSDSSYALRRVAYQVGVEARTLAEARALRLVILVAGRTPALSCKGLGRLSLPAAGRVIREFYSAVSVLGPRYNK

>FoccOr27I

MAAWRVGGATLWWLLPGRGSLQGSRLSALPGLLLSVVCVVMSAPPHVFLHALVCCVVAALIAMHRELAHRLEQADDAQAILTVVKQHQRLHQLTHGLCDGIADILMHFLFSSFCNAIACTLQILASESDSYTAVGFTLLIVTFLPLASLSQELSDTCLYLRGAASRAALSSSSAATRRSLLMVMVAAGRPPSLYCKGLSPLNLRAAGSAIKSWYSVVSVLATRY

>FoccOr28F

MAVSRKLQTWRQFIAPQAPLSASERWYHCFYGSIASCFILGLSFLLIISGVIHKPTSNGIDVSCGYYSDFYGQVFLILQKDRIASFLTDLADVVQEIESESRTSLKVALQQADRSIRRNTIYFSVFYVLLLCSMLPESIMTGISHAPMWPQLPEPLGNWVSVVLFCGSMTTGAGLYYMMWALIFSVLITLTALIRTLSLQLQVAASKREVIDVIKYHQRVKDLSQNFEMFFATHLMHLLASSFLVPLTCTIKGITSKFEPIILTGGSMLCGVFLPLCYRSQELIDASWCFREASYHCVLKMNQHGSLKDLQQSFVLVILAASRPCKLSVRAFGRLDLENAGNAVQCWYSFVNVLLKVTQQTKN

>FoccOr29F

MTVSQKLLTWRRLIAPRALHSCSEKWFQCLYNSFATSITVGLALLLFIGGFIQEPSFNGIDVTFGYYSNLYGHVFMMVKGGRIESFLRDLADVVKEIESKASTPVKAELQRADQSIRRNVIYFCTFYALLLCSMLPESIITGISHAPMWPQLPGALGDWSSVVLLCSSMITGAGLYFVMWALVFSVLITLTALLHAIALQLELAASKREVIDLIKYHQRVKDLSRRFETLFATHLMHLLASSFLVPLTCTIKGITSKFEPIILTGGSMLCGVFLPLCYRSQELIDASWCFREASYHCVLKMNQHGAPKDIQQSFVLIILAASRPCKMHVQAFGRLDLENARNAVQGWYSIFNVLLNVTQKKQV

>FoccOr30IJ

MLKMTVSQKLLTGRRFIAPRELHSCSEKWFQCLYNSFATGSTVGLALLLYIGGFIHEPSINGISVAFGYSSNLYVYVFMMVQRVHIESFLRDLADVVKEIESKASTPVKWALVFSVLITLTALLHAIALQLELAASKREVIDLIKYHQRVKDLSRRFETIYATHLMYLLASSFLVPITLTIKGITSKFEPIILTAGSLLCGVFLPLCYRSQELIDASWCFREASYHCVLKMNQRGAPKDIQQSFVLIILAASRPCKMHVQAFGRLDLENARNAVQGWYSIFNVLLNVTQKKQV

>FoccOr31J

MAVSQKLLTWRQLALSEEPLSSSAKWFARFFSSKVSCLVHGLSYLFIICGVIHSPTSNGIDAVFGYYSDFYGQIFLILQRDNIRSFIKDLADVVQKIEFDASPPVKVVLHHADRSIHRIVMYFSAFYTVILCSLLPESIMTGKSHGPMWPQAPGAIGNWLSAVLFCASLTTGAGVYFTLWTLICSVLITLSALIRALDLQLGLAASKREVKDLIKCHQRIRDLSQSFEILFAGHLMHLLASSFLVPLLCTFKVVNSEFEPIILSGGALLFGVFVPVCYKSQALIDASFSFRETAYHTVLRMNQRGTPIEVPQYFVLVILAASRPCKLSVRAFGRLDLENAGNALRCWYSFVQILLNATQKTHLMLSSAGSLL

>FoccOr32P

MLKMTVSQKLLTGLRFIAPRELHGYSEKWFQCLYNSFATGLTVGLALLLYIGGFIHEPSINGISVALGYSTNLYVYVFMMVQRVHIESFLRDLADVVKEIESKASTPVKAELQRADQSIRRNVIYFCTFYALMLCSVLPESIITGSSFAPMWPQLPGALGDWSSAVFLCSSMITGPGLYFVMPLCYRSQELIDASWCFREASYHCVLKMNQRGAPKDIQQSFVLIILAASZPCKMHVQAFGRLDLENAGNAVRGWYSIVNVLLNVTQKKQV

>FoccOr33

MTVTGPLGTWLWLLRTKRPLDIRQWLWASCVGSKLLMLVANTMLLLAAIQAPSMKEAAAGTLMCCGFYSCNSAHAVYLLRRPRARDMLARILRVAGAIEEEACENGKVVLTKATRSIRRMSVIQLTYMGGMVCSVWAHVLVGGRPYAPMWPPPPLPAPWAERAVGYFEMAAMLLGCLAYFTLITLFSCIVMALTGLYQALSLRVETSQRRGQVLRLIELHQQLNRISREVELFFADIIAHLMVAILVVPLVATLQVVFNVVDALTFMSSSILVSVFLPMSAVSQSLTDASASLSRSAYSSAFSEVANAPSFPLADTTSPSPRALLLVMVSASRPARISLKGLGPVSLSTARAALRVWYQWGNMLVSVAR

>FoccOr34NF

KSELIRASQMNRRMTRLQVCFGSAMVGSVWLQVCLTGRAYSPIWPMPESELGQLFVGVVQMITMLVGCTGFHAAGVLLGTLTTSMTGMFGALSSCLEKLLESTKKTHVIRVVRLHQELCVVSRQVEALFADSMMHYLLCCFSVPLISTLQVLRNDLAFLTFTGAFINLAVFMPIAVLSQRLSDASIAVRSGAYTAAMTVRGAETVPVRQVLQLVLLEAGRPACLSIKGLGPLTLSSAGNALRAWFSFTNVLIKVG

>FoccOr35NF

KSELIRASKLDRKMTRLQVYFGSAMVGSVWLQVCLTGRAYCPIWPMPESELGQLLVGVVEMITMLVGCTGFYAAGALIGTLTSSMIGMFSVLSTSLGTQLETTKRTDVLRVVRLHQDLLVVSRQVEVLFSDSIMHYMLCCFSVPLISTLQVLQNDLAFLTFTGAFINLAVFMPLAVLSQRLSDASISVRAGAYSAAMTVRGEETVPVRQALQLVLLAAGRPACLSFKGLGPLTLSSAGNALRAWYSFTNVLVKVG

>FoccOr36

MGVESVSDFFAYWLRQHDKSAPKTLRERFWLSCRGTGVKWAPFTGLLVMTLSGLDTIMTRTSLLDAGPGIRFLFGMYACTTAVYFNAFKQEETLAMLRQLSWVAVQVEGGELGGQPDPDTQAMLRRTATSSSRFIRGGEIWSCFVMISIGLPVALTGKPANPTWPPPDTPAQWWSLLVLHCLSTVFLPAAYFSLFPLITSLFRQCSALEHAIAKLLERAHTFQQVRDATLLSSELNHGFAMLNGLFHGILSHILQSTLFIPLFAAYEMLNGQFDGFLIAVIPLNFAFFLPLCLAGEGTKSAALAEAAYNGSWLEGDHNCRVLRLQVMQRAARPNQITSREMGPINLRLFQEALKSWFSFLNFMLNTL

>FoccOr37

MVYETATDYLAHWEAHFDKSIPKPPWKRLWITCHGTGVKWAVLVGVAVFVLASLDCIMTRTSLLDAGPTLRFLSGIYSCTAGVYIWAFKRQEALAALRQLSWVARRIEAGAGGALGARLDLDDQETLRRAVVRASRYLRGSNIWAICVQFSIMVPVAFTGRTVNPAWPPPRTPTAWWALFTLHSFSTLFLPIAFFPMAAFTTGIVSVCSAMSLAIGSELRKARTPQQVRDAALLSSEFNKGFSMLTNLFHGFILHHLQITLLIPLFACYEMMHGQFDEFLVVLVPINVLIFIPLCFAGQAMMDASEAMSYSAYSGSWLESDRFCRILRLQVMLRSARPQQLTSQELGTVDLKLCQNTLKSWFSFLNFMINTL

>FoccOr38

MKLSGYLERRLEEVCPPKGARLSRRAFNVLQGTRVSVVVSGAVALLAIVDCFTKESYLAASLGIRFLTGILSCIVAQGVYVYRREGMRRVLGDMAAVAQRLEANADDDAQASMARAARRCARLQRLYGAYSVMVVASVLFPTLSSGSLSMPVWPRPEHTPEPRVAAAAALASQVITGTCCPIAFYTLVGLMVVTQQACGALYRAAGAEVQAVRAPEALLDCARLHGRLSAAAALLDRLTSDVMVLLLSAVLLLPIQGMYDIVQGHIDAYLLSSFPIILVVFVPICISGQGLSDASELVGLRAYQGRWPAEAPEHRRVRAFVMLRASRPACITCRGLGALGLPVCESVLKSWFSYLQMLLNLSNKTAHATD

>FoccOr39

MKLSTYLTRLAKELCPPQRRLLSFAFKIIQGTKVSIVMTVIVSLLIAAASCTAVSYLEVSPNIRFISGLMAASVSQGVYVFRRQDVQEIVGCMASLAKQAEYSTHVDKGLTRVARRCMMLQRFFTIYSFLVAFTLLSPLASGGLPFPTWPHPDTWSYPHVTLAVVLVSQTIASVCAGFAYFTLWGLMLATQEACTALFVTIGTMVEEAVTPEDLLVCVRLQVRISTAATLLNRVTSDVMVFLLSAIMVVPIQATYEFLQGNVDFFLLLSCPIIFVVFVPLCYSGQDLTDASRRVGQKAYTADWVAGCPAYRRTCVLVMLRAARPALITCRGFGTMGLPFCKSALKSWFSYLNTLVGISS

>FoccOr40N

LSIASNIIVVSLAIMDCLTKNTYLAASLGTRFVTGIVGCSTAQIFYIFRRAALGAALERMSVVAKDLEALQNPFISAELERVAAQCYRLRRIVLAYEGMVVVSILLPTLLAGQLSMPTWPRPEDTAAPRLVYAAMFTVQLITGTSCPIAFYSFVGLLGSAHMACAALFRGAGTAVATARGTVELRVVTQLHAVLCESASLLDAVTSAGLPMLFVGVLALPIQGTYDAVQGQVDGYLLSSAPIIIVVFVPLCYSAQAMSDASAEVGLSAYRGAWPAEDARARFIRTMVMLRATRPAQFTCKGLGAVGLPACQSVLRSWFSYLQMLLNFAKDGG

>FoccOr41IF

MKLSWYLDQFQKKICPEKGMLRNRAFNMTQGTKVSLVVSVVVALLAVADGIAAKSYLDASLNIRVVSAILSAVFAQGVFVYRRANMGQIIGGMMAVAQEIETNVNLDTQALLEHVAKRCRRLHRVYTAYSVMVCQTVLFPTLTTGNLSLPVWPKPQSFPQPRVAMAVMIAFQATGTKLKEARTSDDLRACVRLHIQLTQTASLLDHLVSDVMVVILSTVLLLPIQATYEFVLGYVDAFSVSSLPIIFVVFVPMCYSGQGLLDASEVVGLQVYAREWLCETPASRRVCTIVMLRASRPARITCRGFGAMGLPLCKSVLKSWFSYLNTLVKFSTKTASELV

>FoccOr42N

AARLVMTDPVLCAQPAVCLTLGLMCMVLTGVGVMGRTLGSQGLVSSLVDMVYAVSLLGCAMSQLFFAQLAPYAAVRGEVVLHFNDPRCVGLLSFGSMKNRKDVQAMLVRLHEVATELEHDLEGPEHRQERDEELESVARLHGELNSVATVQDALFSSYLPFYLLPPLGGTALATCGFLYGAFSSNFMSLVPQIFIIFMPLCLCGDDLQQTSGETLSLCGYSGNWLDWPPRARRLLHCVMARGTRPQLVYVKAFGHLDRKACLSVLKTWFSFLQTLTNLSGISPDGH

>FoccOr43

MTGQRRAWPTPTPRRRPRHTSAAMLLSREVYVWERSLNGETARARVALWIWRKYSAGTAGFLLARRTASFVSLMRHVRLLAEDVEDAVGEVRREVQEATGRCAELVRFVRRAFTVYMTSGMSMIVTQQIVAGPPAPTIGEERSHAHVLVHINETIKWMFYAVSYFALWVLLATTLLAFNGMYNVLERRMSRAHGYRQVLAAARLHHDLLSLQAELRDVTFGPMLHVLAGALVVPLINTYQVLRGVANVFSLLTFSTLPFVFGTLSLLADGIEVRSERFSLEALSGSDVLGEPLAASKLRLMMIARANGHHGRSSGGFSMRGFGPLNRPAAFSTLHRWYQYLQLLMGSS

>FoccOr44P

MPWRCGDECPSSTLRPHIAMLLSREVLVWERSLNGATFRARAALWIWRKYSAGTAGFLLARRTASIATLMNHVRLLAEAIEAAKQDVVEATARCAWLVRFVRRAFTVYMTTGMSMIVVQQVVAGPPAPTVGEQRPHFLVHVNETVKWTFHALSYFALWVLLATVLLALNGMYNVLVLRMERAHGYAQVLAAACLHNQLLALQAELKDMTFQPMLHVLAGHLVXLLTFSTLPLVFGTLSLLADGLELRSERFALEVLSGADVLGEPLAASKLRLMMVTRAHGFQGLPSGFSLRGFGPLNKRAAFSALHRWYQYLQLLMGSS

>FoccOr45N

REKQKLLFVLLNDAEQRPSDGVCRPAAIGPGPGTHNLCTRPQRSARVVDGRAVGERLTRTGALGPVRLCGSGTSMQEARLQARLEAHQRVSRMVLASWWPRRSRRGARPEEPPTTHSELCEWVRIVDGAGLSYLLLMIMMNTVLSVWGMMICLNGVSRITISLAESFKWSCGATYFFSMWLLILTLLMFMETQYALLSVRLRTAKGLREVAECVERMSEMMQFENDMRAIFFTGVLHMLAGALLVPLINTYQLLFGPRTMFSALTLSTLPLVYGSMSLLGDRLALGCDEVGFAAYSGDWPDEEPRVRRLRVMLMQSCSRGAPFAVRGLGDFTRGAAYHVLKTWYQYLNLLISTVA

>FoccOr46

MDGDELSLYEESEDLDAVVDDEESDFDGDDEDVHKEPAYMVSRYVRRARNVLMGAWGKHPHGQVIHVLIAATTAPLQRPLAVLQASLSFVLAVRHVLNRDVIALMPLIIDFVYLSGVYILSRSTPTFVPAFVAWLERAEVVITELEENVTQEHVRELRMRMRLGRTLFIFSFFNMGSLALAIVLTPLVTKTAFFGVLPHAGAYGTAFFAEYAVLTYTLLPMALSIVLKQCCHAEGGTKIVAMYRMLHNDMLAARTPFLSAVYWGGNTAVMVCALQTLFDGPTTRGVLTGITAVMSYAWLCVLGEKLARGSGGVVSAVHRCEWLYLEPRARRVVVFLLKRAQRCDSVRPRFVGDMTFNFFMNAVITWYQEVQFVYQ

>FoccOr47

MGTSGGHALRLVRGPSALIPRPRGLSAEEGSPWSLQGLWILVHGGNLTAALSALLGVQCLLDAAVHLGGEANRSTQDAMNAAIVLCGACYQRFWARDRHLVRGLLADLLRLTRRLEGLALCAGPDDPQEVALEAERAARVRRTARLLSLCRSAMILDVPVALVFMVTWPLISVRLTFPNDHLPGVGSAGELAGELGVGGLWFGLQYALQSAAISCSFFCFAGLQVVHVTLALVSAALLRTLAQMVQVARTTRQLYAALDDHRELLQPLFHIHCGALGFALLATTALVKGHHDLYVLGLWPIAFAYTTVLGLEGDLVQESSELMLHAAFAGHWQEGARRFRRDALFVMLRATKPVSIRAAWLGNLSLPTLHSLLSSWYSFTQLFIRLS

>FoccOr48

MDKYLTEWLVHMGDRLEGTQRPTAKILQLLRESWIGGLLSLSLLMVSVAETLSPSEQPFIRRLSVRNAFPCCSSISAFIILTKRKDTILAAVRHARDAARVLCTPGETQTALTANVRWSRFLLWRLFPLYGVAVYVMVQVAMFYDTSSVNAWLRKYSEVEAFGLLKTPFEVASLACCMTSLYTVLFAELAIHNTTATLMEQLGRRLAYYRGCENTSRSVRLHVSLLAACRHANAMFGLFLPLYLLGHFALSMLSTVVAVRSSASGEVDLHALSCLPYLLVFFVPMCIAGQRVHDASYGLKDKAYAGPWLEENVQARRSRLLVMASCSRPANFTVPGVGTLDLPTCRKGFRLWFQFVQVLINLQ

>FoccOr49N

IAGLLIFSLLLVSVVDTLSPNEQPLTRRLSVRNTFGCCSSMSSVIILRKRKDTILAALRHARDTARILCVPGETQTALTAYVRWSRLVLWRLYPLYAVAVCIMVLIGIFYETSSVHAWLRKYSEAFGLMKSPFEIACLACCITSLYTVLFAELVIHTSTATLMEHLGQRLAYYRGCGNTTRSVRLHVSLLAACRHANAMFGLFLPLYLLGSLALSLLSTVAVVRSSASGEVDVHALSCLPYLFVFFVPMCIAGQRVHDASYGLKDKAYAGPWLEENVQARRSRLLVMASCSRPANFTVPGVGTLDLPTCRKGFRLWFQFVQVLINLQ

>FoccOr50N

MEQYLSEWLQHMGDQCNAPQRPLVKILKLLRESLDALNTLVRRSRFVLWRIYPQYAAAVYVAVLLAMLLDGSPNFNLWLYHHSMVLWIIKWLCAVASMSCAYVTVFTVMFVELAIHTSAAALMEQVGHRLSQKRGMYETVRSVRIHVMLLAACQQANAMFGLLLPLYHLGSFAISLLSTVSVVRGGVTDDLYATACASYIFAFFVPMCFAGQRVQDAGTSLSTRGPWPEEDITARRARLQIMVSCGRPVRFMVPGMPTLNLPTCRKGFRSWFQFIQVLINIQSAK

>FoccOr51N

MEEYLSEWMQHMGDRRSTPARPMLKILRLLRESWDKLNSSVHRSRLVLWRVYPLYAATVNVVVLLSMLLEASPNFDLWLYHRSTVLWLMKWLLGVMSLSCCFVVVFTVMFVELAIHTSAAALLEQVGHRFSQKQGKYETVRSVRIHVMLLAACQQANAMFGLLLPLYLLGSYAVSVLSTVSVVRGGVTSDMYGMVCTSYIFAFFGPMCFAGERIQDAALGLSTRGPWPEEDVCTRRTRLQIMVCCARPARFMVPGMPVMNLPTCRQGVRSWFQFIQVLINIQSRK

>FoccOr52F

MDEWLHHLGEENSAQRSKAVILLRHLRESSLSFFATVLLTLLAGVDTAALIARGDPLSVRVPVRTFVAAYSCVIAQFILRKRKQVIIGTLSRLKAASRQITALLTNPGSKDALRREVRHSRLICWAILPAYAFIVNAGVLLTMLQDVDSPVNKWIRQKAEAVEWWQPWGPTALPYLKWLFEIYSVSCCNVCLFTLLLVEQSLHVACVVLFSEVSWRFEFWRLNKHNGSHMSAFAKLHTSVLASCSEVNSVFGLLLPQYLLVSVVMSMLSTVALILSDGAPDGFASTCALYLFVFFVPMCIVGQQIEDASRTVSVAAYRGPWLEENTATRRLRLIVMASPPARFKVAGVGSLNRPTCRRVMRSWFQFVQVLVNLRKR

>FoccOr53F

MDEWLHHLGEENSAQRSKAVILLKHLRESSLSFFATVLLTLLAGVDTAALVARGDPLSVRVPVRTFVAAYSCVIAQFILRKRKQVIIGTLSRLKAASRQITALLTNPGSKDALRGEVRLSRLICWPILPAYAFIVKAGVLLTMLQDVDSPVNKWIRQKAEAVEWWQPWGPTALPYLKWLFEVYSVSCCNVCLFTLLLVEQSLHVACVVLFAEVSWRFEFWRLKKHNGSHMSAFAKLHTSVLASCSEVNSVFGLLLPQYLLVSVVMSMLSTVALILSDGAPDGFASTCALYLFVFFVPMCIVGQQIEDASRTVSVAAYRGPWLEENTATRRLRLIVMASPPARFKVAGVGSLNRPTCRRVMRSWFQFVQVLVNLRKR

>FoccOr54F

MEEWLNHLEKDGTRRPSTAKRLLQHMRESTTSRVCVVLVTLLAAVDTLALGFDGESLSVRVPVRTFVAGCSCVTAQFILIHRKRVILNAFHRVRAAARQLPYLLDNPNSKGKITREVRYSRFLHWRVAPLYASIVDAGVLFTMLQDIGSPINSWIRQNAEGILWSQLSLHQLLRYSKWLFEVYSVSCCIACLFTLLLVEQALHSACAVLFSEVAYRFQRCRMARRQGNAIFNRIARLHISVLATCRDVNLAFGLLLPSYLLVSVLMSLLSTVSIILADGTPDFFALSCFSYLFIFFVPMCLAGQRLEDTGECVAQGTYRGPWLEENALGRRQRLLVMGSRRPNFSVAGVGRLNRPTCRRVLRSWFQFVQFLVNIRKR

>FoccOr55

MSSPALTLLLLRWQSSLRNKPTQPWLQRANSRLRRSKWSQALILAWMVRCAADMVLARDAEAIRVPLRYFTGMYACVFVVRFFRRRGSKCSAILFQLATASAQLEETGCPETKELLVRSARILRWTGVFYTVFRTTQIFGVIQSSVVYHSDSAAAAKLDSYSSNLTTIVYILNATAMVVGLVAFYQMQTLVYVLFGSCAVLWRALGYECERSTAGGSGPLCRLAALHALLLQAARSSRLLLAGLLFQWLPVILVLPLLATVELIVAFKVDLLALTCFTVIGTIFVPVCITGQMLEDTSAALSDRLYNGPWVEESWKTRRLRLQFMAGATRRANLAVQGLGSLNAPTCLEAIRRWFGFINVLLNLQY

>FoccOr56N

WAVSLIVLWLLRLAADVILADSSAAQRIPLRQLLGIYSCLWGAVFFMCRTRAALEALAPMLAVAGELERWGAAETKGLLAAGALRLRRVSHFYYFMGLTQVAGITKTVYENPDSAVSRLLESNWRHGRAVVWSLYVAHTAALLLGIYAYYTMLAVLFSTFGSCSALFRALGAESGCAARRGAAAVAFLAGQHARLLDAARRARLLFADLFAHIVSALLTLPLLGTAELLVNIKDVDSLTLATFAVIATIFMPMCLAGQELEDSSAAVRERLYEGPWLQEDAATRRARAHFMLGAGCRARMAFGGLGSLNAASCLHVVNRWFSLLNVYINTNARHTAEPPPPL

>FoccOr57NC

WSVACCALWLLRLAADIALAAGPDARRLPVRQFSGIYSCTWSLLFFVWRGRDARAALDPMVAVSAQLESQGGAETKEQLAASSRRLHRVALFYYVLGLAQVAGVTHTVYSGDSAVARLVLVLPHGRLVNGLLYAAHTLAVLSGLFSYYTMLAVLFTTFGSCSALFRGLGAESARTARRGQGAVLFLARHHVRLVEAARRSRLLFADLFIHIVSTLLILPLLGTVELIVAVKKVDSLTVATFAALTTTFVPICLA

>FoccOr58F

MGTDTPLYSLLEKWRMELDGSAPQLWKQALFQIRRCKWMAPMFVVFVARSTVDLILAHSLAGIQTPLQHVANVYSAMWALFFFADFSAQCSGVLQRLAHVSIKLEHGKDAATKAMLARAARRLRVVGHVHTVLQWLHVVVGSFTILVAQTRHSKVSALLRTTALGAFLDAVLSAVQMLVLFVAVIPFYRVQALAYVLFASAAVLYRGLGAVSASTRSTGEIALRHVAAMHSPLLAAASETRKVFEVPLLRLVLCAVVLPLFGTVELAVSLRHATTGKFAVTIFRAIANLFIPICFTGQLLEDASGALRSQLYAAPWPEERPPARRLRAALMLGAGAQARVELRGLGRLNAATCLEVLQRWFSVVNVLINAQGGMD

>FoccOr59N

WSLVLTVIFAARWSVEVLLASNSETRRVPILLLSGSYSVLLLFEYGSSLVFLSTTARFLNGMRRFYTGFAMVQFFGVMHTTYGYRDVSPIVALLTSVSPLLAWLLYVVNTLGVMCALAAFYETQTYVFLLFSSCAFLWRALGVECEQGGKASSLKSCTEIHAGLVSSARYLRGLLASLVAQWTCTVLLLPFKATVDLITDFRVDIVALTCFTDILTMFLPACLVGQALRDSTESLRTGLYFAPWLEESGRARRDRARFMLGADCKPRLVALSGLLTFNAPFFLDVIHHWFGILNVFLSLHSNKANL

>FoccOr60

MQRPLLELLREWRLEVGVDARVASRRHRIAVSLRGTPFTASVSVLVGLLALLDVLGASGGGLQMNALRVLLALCSCLPAQVFFMRRMPQARRVLDHMIGAATVLETVATPADQAEMSRTASRGRLVARACAAYVLLGTVTILCVLLPDDGGSSVVERLDGPLRHAVLACVKLAVAAACTSCLVAYYTCYALLFVLYSGVASLLGAVEGCAQQAAGTRTLRRIVWAHGQAVMGASQVDAMFRDLHPFYLFAILTLPLLDTAQFVSKRKADLMTPATIPAILGVFLPICLGGQRVRDRNTAACWGFYCGPWPEEGRKPRRLRAQAMAVATKDRSVTMHGFGALDRAACVRAASTWFRFLQILLSLQ

>FoccOr61N

LAFMAFCTMCCVSAYYALLAFILGIHIACANLYKELGKQFQIDSGVAETTQLAMLHANLKETVQLLDEMLGWLYPHFLFAGTAMPLLCSAKVIISKGQTDSFALGTVPLILLVFITQCLVGQSLEDASRALPDSCYFGPWLQETSPKRKGRLLIMLLAHSHAVLRVRAFGKLNRPTCVNILKSWFSYLQVLLKFGS

>FoccOr62N

VSTVSLAVTLGQAGVDMVLADDIASVKVSRRLFISTTSCLVAQRSLAAEVRRLKGLWRFYTWFFLVTLVGTLHSMVVDRHSAVTRWMYAAGGSALRYSRLVLDFYTMLAFLAAYYRLLHLMLTLYRLSSALYRELGRRLRAAGGAHHLRWAVALHARLDATWRSTRAMLGLLLPLYLIVPLLLPLFSSAELVVFGLQAADPVALATCPLIVVAFVPQCLAGQRLLDASTGVALSAYCGPWLEEAVPERRIRLMVIDTRPQGVPVPGLGGLSRPTCLNAMRKWFQYVQVLLNFDSSL

>FoccOr63N

ILLIIRQVIEAFGSVCGDVCHFTLLAYVLVIFASCSLLLREVGRVAQQTGKVSDVAAMHVQLVLGCSRLNAAFGAYLPHIAVAPSVIPLLSTVTIIMSAEKADMYAAGSFPAVLFYLAPLCAVGDMLVAGSDSFCFSAYCGPWLEEGPATRRCRWLLMSAPTMNLSVPGMVSVNRPTFRKAARTWFQFLQVLIELKA

>FoccOr64N

ILLIIRQVIEAFSSICGHVCLYTLLSYVLVIFASCSLLLREVGRVVKQTGKVGDAAAMHVQLVLGCSRMNAAFGVYLPHIAVAPSVIPLLSTVTIIMSAEKADMYVAGSFPAVLFYLAPLCEVGDMLVAGSDTFCFSAYCGPWLEEGPATRRCRWLLMSAPTMNLSAPGMVSVNRPTLRKAARTWFQFLQVLIKLKA

>FoccOr65N

LHVVLSAHVDLYTLLALQLDSVRACAVLNTCIGHRFQVEGGRRSTARSASLHAQVNESARLTSMQFGHLLPHFLTVGLVMPVVATAEVISSGGHADSFAIAMVPLLFVFFVSQCVGGQQLENSSEGLGWAAYQGPWLSEDARARRLRLLVIQSVTARPACFGVPGITVLNHATCQDALRIWFQYLQILLNTNGRRRHGPAQ

>FoccOr66N

VQEGSRHTARSAMLHGRVYEVSRLLNQLFGHLLPHFHTLGLVLPVVSSAEVMLGRADAFALAMVPLVFTFFVSQCVAGQRLENSSEGVGWSAYQGPWLTENSVSRRARLLVIQSTVARPASFGVPGIIVLNHASCQGALRVWFQYLQILLNINGHAH

>FoccOr67

MKSKPLTTFILSLLDDISPGKNDTHWRRLKFSLRGNWVSLVASVITLFLELSAVTVPGSIPHDQVPVHVSLLLGVYSCIFAQVYLVRRVHLLQRCLALIASAASRVEEHVDAESQRAMLRTVRRGPRMRRFFNMCGAATEVLVVFSLFSTPPRWCESQDRVWLHYAILLMETYATSCCLNAFYTVIVYLLNVCRACADLHVALARRLETGHATSTCASVQGVVLNATLVLNAATSDLLPHLLFGVVGVPLISTVEVVSSGNLQDIYAVGTAPLLLTLFVPICLTGERLGEARREVAFRAALGPWLEELPVVRRLRLGILVAAEGRGAQLRGSGIGLLDKPACLQALKSWFSFVQMMLNLRDKA

>FoccOr68F

MTKLPVFEGLATKALRLLRADLYYCDGANHDDGPLTQHLRWWVQLLDAGELPGRTNYLMALFRQSITSLVIAVAVMVQMIYDTAVTGSMFGFRIGLAVYTTIVVIVLFMQRRRGIKNHLAATAHRCAKLVRFYEAYAWLAQAAIFTALIARNPTWQYHAVAGLEPVLGEHLEVAANMLRLVYLAFQLFTGVCAVVADYMAVVLLLVIYISMGDMYDALGEVLQTRPCDTLLARQQSVLTQASLTVDEAVSDLLPQVLIVSVVVPLLCTVEVVLHGLDADLFAVGMAPLVLFVFVPFCLVGDAMAAARLRFADRACAGPWLEETLQLRKLRLGMVQVSQGHGGDLKGLGIGLLDRKACGNALKSWFSFLQVLMNLQ

>FoccOr69C

MTKLPVFEGLATKALRLLRADLYYCDGANHDDGPLTQHLRWWVQLLDAGELPGRTNYLMALFRQSITSLVIAVAVMVQMIYDTAVTGSMFGFRIGLAVYTTIVVIVLFMQRRRGIKNHMAATAHRCAKLVRFYEAYAWLAQAAIFTALIARNPTWQYHAVAGLEPVLGEHLEVAANMLRLVYLAFQLFTGVCAVVADYMAVVLLLVIYISMGDMYDALGEVLQTRPCDTLLARQQSVLTQASLTVDEAVSDLLPQVLIVSVVVPLLCTVE

>FoccOr70

MAKVPAAQGLASRTLHLLRADLYICDVANKDEGPLTQHLRWWVRVVSAGELPGWKNYLLAFCRQSVASLVISVLVMVQMVYDSTVSGSMFGFRVALAVYTTLVVIVLFMFRRRGIKTHMAATARRCAKLVRFYETYAWLVEAAVAVALVFRKPTWQDHLAAWMEPLIGTHQKFTANVLRWGFLSFQLFVGLCAVLSDYTLVVLILVIYLSLADMYEGIGEALQTRPCDTLLARQQSVLTQAAISVDAAVAPLLPQVLIVSVVVPLLCTVEVVLNGVEADLFAVGMAPLVLFVFVPFCLVGDGMAYARVRFADLAGAGPWLEETVQRRKMRLSMVQLSRGHGGDLRGWGIGTLDRKTCGSALKSWFSFLQVLMSLQ

>FoccOr71

MPQLPVAQGLASRTLQLLRADLYICDVADHDEGPLTQHLRWWVRVLSAGELPGKTNHLLALLRQSTTSLVIVVAVIVQMIYDASVTGSMFGLRIALALYTTLVVIVIFKFRRRGIKTHMAATARRCAKLVRFYEAYAWLTEVTVFVALIVHWQDSAAAGLELLIGTHHQVATTILLWASLALRLYSGVSAVLADYTSVVLILVIYLSMANMYEAIGEALQTRPCNTLLAHQQSVLTQASLNMEASVADLLSQVLIVSVVVPLLSTVEVVLNGLEADMFAVGMAPLVLFVFVPFCLVGDAMTYARLRFADRAGAGPWLEETVQIRKMRLGMVQLSQGHGGDLRGWGIGHLDRKACGSALKSWFQFLQVLMNLQ

>FoccOr72F

MHTEYLSPRTSSIQSITDGERDGPLTQHLRWWLKHISPGRLNNQTHRLLMFIRGSKVGMFVSINVLTLAIADACLSGSVFKSPVLARFGLGVYSCLWSQLVFLSRTQPLLACLRRLIGLTTEFERHGTATDLLERMKQTAHQSSFLLRFYVFYSLTTMFVILITFVAVEPRWHSASAAMLASILRSAGCAELSEASVTALGSCLRWVVVAIQMLTLVFSLTSYYTIVSFIILISGATANVYEVIGHRIEVEYCSNRQTAMQSELLRVSICMDSTFADMIPHLLVVVVILPLLSTVEVVANGMQADMFALGTAPLIFTVFVPLCLVGDSLFKCRRAVATRAARGPWLEETPQLRRLRLGMIHSALGRGARLRGNGVGPLDRRTCGNALRSWFSFLQILLNVKRSGDSV

>FoccOr73I

MLESGQPSRVDSWPLDEGPLSSHLRWWLKTLSPTGRDKFSARLLFWLRTSKIAMTTSVVTLLVGISDVAVSRSMHEASVPVTLSLGVYSSVFAHVLFVKRLPALLQTVALLLATTAGVERHLDSASQAFGLVCSFTAFYTMLPLVIVIYSTCADLHAALAHRMQTAACCPHDAALQSGVLRACIVADSALADLMPHLLVVSIIMPLFSTAELIASGLKLGAYSLGTAAPILIVFVPLCEVGDRLSRARRGVAESAAEGPWVEERPKQRQLRLAVMQAAMGEGAFLRGSGIGPLDRATCGNALRSWFSFLQVILNVQRKAGVH

>FoccOr74I

MHHNTWRCVGLEDLCRGGPLTLNLRWWLHHVTHEGKHDNVLRWLLFKLRGSRALLARRARKGRFMRWFFVMHAVATEVTFVFTLSIWAPTWLQHGVEDKEIWKDWTRWIVLSLQYLFLQASGLLAGLASVYTFLPLVLQVTMASADLMYILGKKLEVGRCTRTMAATESAMISACLLSDECVSDVIAPVVFASLLMPLISLVQVMSGVVDSLDALGTAPLFITITVPLYLAGDMLASARQGIANSAARGPWLEEEPWQRRMRAAVMLGASGEGALPRCQGVGILNRRGLAHVLRSWFSFLQAMLNLRRGAKVAG

>FoccOr75

MHHNTWRCVGYENICRGGPLTRNLQWWLHHVTHEGNHDNVLRWLLFKLRGSKFSMRVAMIAVTVTLVNALTAENLNRADVSIRYFVGVNACICLQLMFINREDLLRRWLGLVIVHAHSAETDPATDDEFMALLARRARKGRFMRWFFVVHATATEVTFVYTLSIWAPTWLQQGIGNKEIWKEWMRWTVLTLQAYGLLAGLASVYTFLPLVLQITMASADLMYILGKKLEVCRCSRTIAATESALISACLLSDECVSDVIAPVIFASLLMPLISLVQALSGVVDSLDALGTAPLFITITVPLYLAGDMLASARQGIADSAARGPWLEEEPWQRRMRAAVMLGASGEGALPRCQGLGKLNRRGLLHVLQSWFSFLQAMLNLRKGVKVTASRNNS

>FoccOr76

MISRDFSRAAQPCPGTVYPSTAADTRAREPERQPLTNQLEWWLWLLSGPHVGIHQRALAWGRGNLISVVLSVMTTGLMVLDSVLSNSLFSSTSILAARIACSTYSCVSAQLVFMRRHGDLCGILHNLARLTGDLEPRAHQDSQVHMRRSATLNARLVRFLYFYGWTTVLFLSMLFASIKPLWIAESTSALMRMAGDSSLTLSSRLTVLFDVNDLEYVLSIFQGLVQLVLGFILELIMVVISLILLSEQGGSRSWPSPLRDLLDVLLHLEDTFADVLPHFLVVSVALPVLSTVEVVVKGAQADAFAFLNAPLIFGVFVPMCMVGDKLSAARFGLSRNAFRGPWLEEDQQQKKMLLALMQIADGRGGQIRGTGIGRLNRSACTNAMKSWFSCLQVLINVQWQSAATVK

>FoccOr77

METDSRAENPETWPLTHQLEWWLQLLSGPHVGFHQCALAWGRGTVSSVVVSVVSTGLFLVDGVLSKSLFGTTSMLATRVTISTYSCICSQLVFMRLNADLCGIFHKLARLAANLEPSTQRESQIRMRGSAVLNARLMCFFYIYGLASVAFLTMLFLYIKPLWIEETASAVMRLFGDSDVDLSCLEWSLRLAWLPVQSCMVAVWTNSFLCLIGCMIFIYATCADLFEAIGRSVEQGHCSRITYALLQGVLDVLLRLDDTFADVLPHLLVVSVVLPLLSTVELATRGTEADAFALLNAPLIFAVFVPICFAGDRLSVARSGMSRSAFYGPWLEESQRCKKLRLALMHVADGRGAQVRGSGIGLLNRRACGKALKSWFSFLQVLVNVKRRNAATVK

>FoccOr78F

MAVETPDPWPLAGQLQWWLRLLGDHHAGLHQRALAWGRGTPSSLVVTVAGTGLFLVDAVLSKSFGSRTMFAIRMTIGVYSCVFSQLVFMRRQADLCDILHNLGRLAADLEPRADRDSQGRMRGSAVLSARLVRFFYVYGCTSLLLVSIMFVCVKPLWIKEIAGAATRLVANSVSDWQPQVHLDASGMAFVCINSFYCLILCMLFIYAACADLCEAVGRGIEVERCSRSWPTLLRGVLDVLLRMDNAFADMMPHLLVVSVVLPLLSTVEVFARGTKADVFALLMAPVILAVFVPICVAGENLAAARRGLSVRAYRGPWLEESLRHTTLRRGVMQVAGGRGALVRGTGIGLLNRRACGNALRSYFSFLQVLINV

>FoccOr79N

FAVSASLALVSVVTFDFFKTANTSSLTIIFGAYSSIFAELLLWKRWKHLGMCLSTLSDIVGAVEDNVNPDLWMLQALGRNRARAHRLRRSLYLAFGWLTVVSILRMDVLVPPVWARSMAGNLARASGLDEDVCFSLVHVTKTVLDVVGSLCCLTAFYTFVPCVLLMYSACAFAHKAVGERLMGSGSVVQAVELQNAVLKITIAIDSVITDLLPHVLVATVFMPLITTSQMVVSGDFNPFDFVGGVPILSVFVPLFEAGDDLAASRLQVAELAGFGPWLERSPRQRGLVVGAMRCAAGVGGLPRFRGFGSINRHACLQALKSWFSFLQMLLNLRKID

>FoccOr80I

MVKKELDLASLLVEWRRSVNAAIGCSANGILRTLQGSKWSVAASLVTLAVALMDMSETAGLSAFVMFFGGYSSVIAQVLFWRRKTILVRGLDLLGETMAQVENSTDPALQARARQHARIHRVCRSLYLIFGWTTVALIIRLSVLLPPRWARRPARYFSVLSGLPEDSCFVFLHAAKTVGEVSGSLCCLTAFYSFVPCVLLVYTSCAFLHQAVGQVLQGGRCSLGALMLQSTVLRATTILDEAITDLLPHVLMATVVMPLVATVQCHGFGRVDRSACLTALKSWFSILQMLLNLRS

>FoccOr81

MCHPSTSGQSRLTKLLNKFFQYKFTVVSLAYAMYVSAAAQWTDQPASSKQFNIRVFLSAYSSIVNSLIFTRNRNTLFSTMEWVADLADCTYRLADNKGKVAMQQSALQSRLYTKLFLSYGATVYVAVGTGLLLEHRHYSDILDMTIGGAGDPGSTWRQVVGAIGYIQMPIFEYNAYIGCIGLFLTISLVSTVTRSLITVQGQINRGVERGDCVPWTALQSQVVRAARGSEEAFADVLPHILLSAYILPLLATADVVFNGPEADLFALAISPVVLVVLWPLCDLGDGLLEAREQLCTSAYSGPWLEETPSQTRRRLLMMSFSYGRRGLLRTRGLGGGVLSREAFGDGLINWVKFLNGIINLKTLSKD

>FoccOr82

MCHPSTTGQSGLTKLLNKFFQYKINLVCLGYGLFICTAVLWTDQPASSKQFNISALLSAYSSLVNQLIYTKYRRPLFNTVELLEDLAEQTHRLADEQGKEAMQRSAVRSRLIIKLILSYSAILTVVLSAVMMLEYRHYAEALDMAIGGSGEPDSAWRQVVKTIGYVQQPIFEYNAFIGCPGFFLTVSLVATVTRSLVTLQEQINRGVKRGECLPWAPLQSALLRAALGSDEVFAGVLPHMLLSAFVLPLLATVDVVFNGRDADAFGLAVLPVSIVTLWPLCEIGDMLLEARKQLCTSAYFGPCLEETGAQSRVRLLMMRFSYGRRGLLRTRGLGGGVLNREAFANGMMNWLKFLNGLINLKNAL

>FoccOr83

MCHPSTSGQSRFTQLVNIWYLNKWTVVGNLFGIATFVISAAGAPCSRYMQFNVRMAISLWSCLAGVLIQRKRRMLFSVIQLVADLGARAHRLTDRQGKDAMRKSAVVSRLHAKALFWYGGAILVAAAFNLLMERSIWAEVLVTAIGGPRMADSVWNTAAVCVDWATTIEYQYNTLLDGSAIFVSLACITTLTRSLAEIQTQINRGVKVGDCLRWAALQSRVTAGCRELEETFADLLPHILISAYFTPLLAIVDVVFNGRKADLFSLALSPVVLLVLWPMCDIGDELAKARERLCPAAYLGPWLEEGCEQTRVRLCLMLFAYGRHGLLRTRGPGGGVLNRGTFTTAISSWIKFLNALINLQSASRHQVRGK

>FoccOr84J

MCHQSTSGQSRLTRLVNICYQSKVNAFSSQWQWTVLLNAYGIFTFVTSSAGAETSSARQFNFRITITIWSGLAGVLIQRKRKMIFSIIQLVADVADRAYRLTDKQGKAAMEVSAAHSRVHTKVFLWYGGVLQVAAGGVLLVERSKFAEILLTAACGPEVAESSWYTAAVALDYLQILVNEYNAHIDCTAVFLSLSCISMLTRSIPAVQAEINRGVERGHCQEWAPLQSQLSRAGRELEETFSDLLPHVLISAYFTPLLAVVEVVFNGKDADLFALSVSPVVLVVLWPMYELGDKLLEAREELCTSAFRGPWLEEPQVQSRLRQLIMRFSFGQRGVLRNRGLGGGILNRKAFGNAFTDWLKFLNALMNLQNVSKH

**102 FoccGrs**

>FoccGr1

MRHSSPQGRRGAMGGHQDRVLSLAQEMKPILLVMRGINIMPLRFLADGGVTCFAARSPLAALSAALWALTVVDALVEVRERAGLVRGAAEFSDAVFQIFFLMLVHHHFLTPVLHWRAAPAFADYLSDWKDYQDTYLAVTGRPLRLGVRPLASHKGAAAVAVPLLLAVALLLTFDGNNWHNTLLGLMAATFHAVIPSFWDVLGTAVVNASDTLSRHLEEEVFAAGGAPEPGRVRAYRDLWLGLRELAERATATWGACTIHYLLCCVSAVLLNGFGLMAEATEGADANLLMASAMLLITVDSLWHTAVVCGAGHRMAASVGGGARDVLLAMRTSGESSPHRAHRARSRSLTTLQTEVASFLKVMQRPTIVTLYGFITLRLTLIVTIMDKLVTYLCVLLQFTINFRLKYRNHRRRSP

>FoccGr2

MGWGRQQAGVVLPKNVKDAPEKDPRPLTMAEEMKPLLVVMRQIFLMPIHFPGDGSAKFTLRSGTMYFNYVGCGIFIVDSAQEAVYRMTVLLDCKNFSDAVFQTFFYILLHTHFFTPFLHWPAAPGFVRYLDRWTEFQGRFRRVTGRPLALSVRPLALHKSVAALLVPLVLCCITALTMKTANWKNQPLGWIAAAFQACVPAFWDVLGTAVVNASTALAGHFREEVTAAGGVPEPSRVRAYRELWLELRELTEGTTKAWGVCTMHYVLVATASVLVAGFGVLAQCTEGISNPILFTVMFLISVDSLWHISVLCSAGHRMQGSVGGAARDVLLSMRTSGRAGGPDMELLQSEVNYFLIVMRKPPVISLYGYITLKLTLTVTILETLMTYLCVLLSFAVSFQFAP

>FoccGr3

MSSAGDSPWVISKKYDAWGPPVDASRKRGGDALVAWGAPPGRGGYGPRLGGLDNRGLPVDFKKQTQYKRQKPIAGASKSTASFVDSAMPLIMLMQVGIAFPVQRRADGRLQWKILSPIWLYTLVVYGVLTVPVVFAASAAVRQLAKEDNTFHDQIYNMTPLLLLHAHPLLPFMHWLEAYFVAGYFNSWIKFEELFHKVCRRPLHMGLRGYVARMAAFSFAVAAFLTGTMPMYMSLSWLQCLAYLHLYLLAICLVSVWLLSSKALCIAAEALRESMHEMVVTGACTAERMADYRQAWLELSELTQESGSAWTYQCGHQLLYLFLLMTVTSFGVLAGVARHSLEARTLAYGALAAGSAFILFIVCSAAETAADRVAEVLREDLLMVQLSDDTQHFKNEVQMFMTALAMDPPIVNLSGYATVNRALFASLLETMVTYLVVLVQFAYSLDSTPDNSTSVTTEAASVKVTAGPTPVPTPTPNAP

>FoccGr4

MSTRALPRGALPPRRPQPAPTPSWTAIKPDKNAVLLRNVQRQLAATEGWPLLQGGVPLPVPRGASAAPTLFDDLLPLLWTLRFVLILPVTVSRDTRTVRFRWLSWTTLVTAVLWMLTSAITCAVVRVRLAPVLSTGKHFHVWDVVPQYSTAIGMSYIFFLPVACWATADRCVALVELWLRVEALATAVTGPLQLPGLRTRARVYSFGSLVLPVLMQSLAMFVLPHERLWMLPAMLHSIAFLMVGTSFWLCCCIALGAFGSLIAARMPADLQRMGPQGLRAHRQIWLALAQLLNQLGVAWGRLQTLLLAILFLAITSISFVITAYVLHGIWDKRLVWLGTAVFFVVISIFNICNGAQIATDGVRLAAIRQLLALSVTHKDEPTCHEIICFLDDIRLERHHLTVCKLGGLSRASLLSFFSLACTYVVVLSQFTLSSETSRTKFDKLPADLVIENT

>FoccGr5

MYEATCARSPPTTTASASSWATSKPRKNDGLMQDAQMAATERWPFRQGGLPGTVRGAPTMFDDLLPLLWTLRFVLILPVTVSRDTRTVRFRWLSWTMLVTAVLWLFTSAINYAMVCHRLAPVLGTGKHFHVWEVLPKYATVVLLSYIVFLPVTCWATANRAVALVELWQRVEALATAVTGPLHLPGLRTRARIYSLGCLVLPVLMQLLALSVLPDDRVWMMPGLVHANAFVMVSTIFWIWCCSALGALGSQVAARMPADLQRLGPQGLRAHRQIWLGLAQLLTQLGVSWGRLQTVILGLFFVAFTCISFIMTSLVLHGIWDKKMVWQGTAMSAVVIGIFGICNSAQIATNGVRLAAIRQLLALSLTHKDEPTRQEIISFLDNIRVERQCLTVCKLGVLSRASLLSFFSLACTYVVVLSQFTLSSEISTKKFDKLPSVA

>FoccGr6NF

LRRREGGCNILPSSSDDPSQYLSTASDAGPEFRWRSPAMLYAASMWLTIGTIACFSAGFQMEWLMCAIRLHHRFYEKAFATLGLTLLFYLPLLLVVWWETGAPCKLIASWPAIESDYEALTGRRLPLRLRRSCRTFVTSILLVSLPLTASMSISVPQYQLWRVLPIMQYNAFIMGVPALWWCLCQALAQASATLAATLVEVPVNRAAVRAHGALWVQLNDNLIALGARWGHTKYFLMLLLFVEWVIAAFTVMSEFLTNEDANFIVLLSHIALFSGSMLFISCSGAQRALDGVGRCSLRALLRLPISRMDEKVATEVALVLTSIQAASKEEVLVNGFIGYNRATLVTFLKNTATYLVVLVQFASTLA

>FoccGr7

MSARALPWALPRALPPRRPQLDPTPSWNAIKPDKNAVLLRNVERQLAATEGWPLLQGGVPLPVSSGASALPTLFDDLLPLLWTLRFLLILPVTVSRDTRTVRFRWLSWTTLVTAVLWLFTSAISCAVLRFRLAPVLGTGKHFHVWDVVPQYATVFGMSYIFFLPVACWATAGRCVALVELWQRVEALATAVTGPLHLPGLRTRARVYSLGSLVLPVLIQSMAMYVLPQERLWMVPAMLHTNAFVVVSTSFWLCCCSALGAFGSLVAARMPADLQRLGPQGLRAHRQIWLALAHLLTQLGVAWGRLQTLLLAVLFVSITSISFVITAYVLHGVWDKRLVWLGTVAFFVVIFIFSICNGAQIATDGVRLAAVRQLLALSVTHKDEPTCHEIICFLDDIRLERHHLTICKLGVLSRASLLSFFSLACTYVVVLSQFTLSSETSRTKFDKLPADLGIENT

>FoccGr8F

MESRQRSANCPAASTQRAIARTLNDTNYYDEQILLGEEIPRRIRRTKSLRGLEPESNVIEDLMPFFWTLRAMLFLPLSTSTNPRTFYFRWLSLPIALTVPSWLGLCWLASVEIGNRLSLVSGVGKHYHLWDLVPEYCTAVCLSYIYFIPVACWASSRHAVALIKFWHILETQIADLYEPIPRSGLRCRAWVFCALCILPPFAISLLSLRTSQLVEKDMWTLIINVHSFFFPCVGVGFWFFCCYGLRAVALNQAGNLLGDFQRLGVAGLRAHRRVWQRLARVLHLLCVTLGRMQIVLFTILFTCYVSISFLFTHFAIDRVWDRRSTICAVLYAVMIVGVVTTCVAAEFVSQALRRPATHQLFHLASSTKSDYILSEIQLFLADIEHLSPKATVLGFGSLGRSSIISFLSLATTYTIVLSQLTVSSETTVKKFEEVDFYDEF

>FoccGr9

MGSRAALLEAKLRASPTVPVFLSLAVGGVLPLHPQHAYLPSTGRLLRSTAPYCAVVMALVLYSTMSVLLLHLPSVVQARVLMLLYTRLKCVNAVLTITPILFIPKMWVGMRGLAARVVDFCNYEITHEALTGRRLDWSWHFRRNQAVLAVLYALLVAVFVSGKIVEPERPCHVPSYVCSMHFQSNLSLWYISWISQIRFAAEDILKYLQESKEHPCTTVRRSRRLWVDLREILEENRTLKNLIFLIHGYLFFVVTLSLYQVLHYGRQGETTAALAQSVINLAMFFQIFVFCNETHKCTATMNKTFTTALDRFSLKVEDEETRKAINLFYLSIAANPAKASLAGFAKCDRPLLSSVVASTVTYLVVLVQFQTMERERYFKLIKVSYINSSEPLIIYLCL

>FoccGr10F

MLQKNAMASVIWVMAVIGAMPLALPAGYRLRSTLPWCALTYSVFVASGAVLVWDQVPVLLAQGVDLDTRMVATELIVHLLPLLFVPIFWWQAGNICRMTAGYCKTKRMIGRVTGSSWRRDARRRRVYAVLTVLLVVLSLVLVAAAYVLVGRLHLRPPHLPALFFSMVLQTLVGYNVVIQLLQLLVLARSIRHRFARVAHSAGAGTVRAYRELWVELRELYGYDDHACHAVYAINMLYLLAQSTMVSFEAFSFYVAGDRWASIGHGVFALVYFAEIFTMCECAHLAVGVMKEPFVAVLDRMSIGRLDHDTYHEVCFFYNTIIAQSPNYSIRGYASLDRPYLASILMAIVSYTVFLAQCRSQAGSTRAPVPVGFNEDHRAM

>FoccGr11NF

RPSARSPYATSPLAPVVNLVAWLGFLPLAAAQPHLPSAAVRSSLLYCVVLYAVTCWSQSLVWWDQALLLDDAQSRWDERLFSLQILWMLLPQVLTPWLWLCAAPIRDSLLRFADYERVYASITNDTLAPRLGVRRRLLQNLMPVILICAIVTLACAINYIQVRGTRVPIRWAYFPAALSAIMTQGMLTILFTTYSYQLKVAAGALCQYLQQATVRSDVSVSQVNLTVQNTRRLWVSLRALYNDVSYRFVDVFQLLHLLLSLILAVYMFHVFRITELMFNLVLEGGFVVYSLLQLGIITNEAHMAVEKMFGPFVGVLDGLCTDDTRLAQTVSLFYHSMAAYPATLTLGGFTKIGRPLFTSVLSATVTYIVIMLQFQQSDSTVVESTVQNTSDTYSSNLSLNSNLSMREL

>FoccGr12NIF

LGGRWALVALSGRVIHQPRYFAGEIKNARQSVDTKPKAKAEPQKQGGEETRTDSLYPFVRSSPLAVVVLGLACIGQVPVSFHRCYEPLLSVRRSSLVYNVLIFCIFGVSYAFVLIDQSVTVFSPSTNWDARLYCLQGLLALLPGSVSPWVWACASSFRKMLWGFAEYEDRPSFYLLDVFFMSQRFALLLLSVSLWQTLASAKMLVSASTQVIWMFVLSFQAYRMSDLAHGAVSAMHGPVTAALDRLSMDARDEHFHRAVLLFYMSVSSRPAHLSLSGFANIDRPLFTSIVSLTVTYLIILMQFRTDVSQQ

>FoccGr13F

MNDADVVGGRWALAALSGRVIRQPVYVAGEIKNAWQSVDTKRKAKAEPKMQHGGDTRADSLYPFVRSSPLAVVVLGMAAIGQVPVSFHRCYKPLLSVCRSSLVYDVLLYCIFCVSYVCLLKDQLMTVLDPETKLDALLFCLESILALLPSIVNPWVWACATNFRKMMWNFAEYEESFLSVTNRSLRDRWTNGVRNLRLHVTMLFLFGGVGCIAGPYFAITERTKAIFSTYNVTSDDPYGNTYYYVPAAFLSLVSSSLFFLFTIAYFLQLRETAIALSEHIDQLRPGDTAGVRRAQKLWVGLRALFNDRPSFYLLDVFFMSQRFALLLLSVSLWQTLASARMLWSAASQVLCMLALTSEQFRMIDLAHGAVTTMHGPVIAALDRLSINAPDEHFHRAVLLFYMSVSSRPANLSLSGFANIDRPLFTSIVSLTVTYLIIIMQFRTEVSQQ

>FoccGr14

MEMNDTDVVCGRWAMVAFSGRVIRFRDASQDVYIAGKTGKALKSVDTKPKRNGQQKQQDGEDTRTDLLQPFVRTSPLAVVVLGMAFIGLVPVSFNCLYKPLLSVRRSSLVYAVLLYSTFCVSYAFVWKDQLGKVLDPSSNWDSKLFSLEAILALLPCSIGPWVWTCTTNLRKLMWDFAEYEESFLSVTNQHLRDRWASRVRNLRGFIFVLFVIVGLGVTIAMLFSEIQNEYKSDDLAGTLVDMRFYLPAAYSSTVISTLLLMLNTTYSLQLREAATALSEHLNQLRPGDTVGVRRAQKLWVSLRALFNDRPSVYLLDVFFVSHIFIMMLLAVYLSFALLSLRIVMLAASQYFWMLFLGFELFWMSDLAHAAVTAMHGPVVSALDKLSLDAHDEHFHRAVTFFYMSVSSRPAHLSLSGFTSVDRPLFTSIVSLSVTYLVIMVQFRSDKSHPSSPPQ

>FoccGr15N

ESFLSVANRSLRDRWPSHVRNLRWGVIILFVFVGVGSIAGPFYAETEFGNSAITFNIVFKNMYYYIPSAYVSITSSTLFFMFTITYFQQLHEAAIALGEHLDQLRPGDTAGVRRAQKLWVSLRALFNDRPSFYLLDVFFMSQLFAMLLLSVSLWYDLSSAKMLVSASTQVIWMFVLSFQAYRMSDLAHGAVSAMHGPVTAALDRLSMDARDEHFHRAVLLFYMSVSSRPAHLSLSGFANIDRPLFTSIVSLTVTYLIILMQFRTDVSQQ

>FoccGr16N

FGGRWAMVALSGRVIRLRDASQDVYLAGKTGKASKSVETKPETNGRQKQQDGEDARTDSLEPFVRTSPLGVIMLGMACIGLVPVSFNCLYKPLFSVRRSSLVYAVLLYSIFCVSYAFVWRDQLGPSLNPSTNWDSRLFSLEALLALLPSAISPWVWMCTTNLRKVMWGFAEYQEAFLSVTNQHLRDRWSSRVRFLRGFVFVLFVVPLLLIIVGMIASKKKEQTKFISGDLNAYDLDLSVNAWYYLPAAYSATVSFFLILLFHTAYSLQLREAATALSEHLNQMRPGDTEGVRRAQKLWVGLRALYNDHPSLYLLDVFYLTHVFLLMLFSVYLCFALLSLGVLEAGASQVMWMLLLAFGLFQVSNRAHEAVTAMHGPVVSALDKLSMDAHDQAFFRAVTLFYKSVISRPARLSLSGFTSIDRPLFGSIVSLSVTYLVIMMQFRSDTSHPSSLIN

>FoccGr17

MTSATTMRLDTVGGDLDARTRRLFEPLKPLYRTAIVFGVVPTTIGRWPRLSFHPLRSTLLWVLFLHSGTLVSVVGTWHHGLGVVTSSNPFGSSADLPAALGLTHTANVALMAVLWCEAHRYKEVFALARTLVGSVDDDVLGADRWPRRRTLVRAWFWLIAVVVVAPVTTLSWHLSLAPFDIDHPVLRHARLPGHLVSHVLLLMYAAAVSIAYNVLEDMAELLALRFAQDAASRRLRPATLLRHRRAWLALRDLLQGVMLAPATVTLGVALMMVNAVLSLYYVVNCVMVGAPTELIASLTLASSDLGFLLVAMCSAADGAVIAVNKFARAFELIEPDSVSDYAFAREVSREALSHKLTPVCVAGYWYLSRQTVSSVGLMVVSYLIVLIEFRQQQDSSAFYSNATSRGV

>FoccGr18F

MSRRTNKVGDMDARTARLFLAVRPLYYTLIAYGVVPVNFRRWPNVGLSFHPLRSTLAWALLVHATTVASAWRVWDLSLQVLSEGNPIASYANFPALLAIFHSANVLLLVPIWCEAHRYKEVVALAKTLVGSVNEDVLGAGWPRRHTVFMSWLWALALVVAAPAITFLWQFQLTDHSGDAFQVPAHWMGNVLIGAIVGGLSLVYYIIKDVAQALAHRFAQDASQEMSCAELLRYRRAWLTLRDMLQGVMIAPVSVTIVCALLVINNVLCVYHVLNCFLGGAPLHQTLGLLGSSSQLGVLLIIMCWSADRAVDSVTNLARSFELVDPLSVTNFAFAREVFREALSYSPESVCVAGFWCLNRRMVLSVCLMILTYLMVLVEFRQQGACSLGSNSNSNATRVG

>FoccGr19F

MYEDLRPLALGLAVCGLLPVAFRPERFSFHPGRSTLLYSLAVTLAICWSGWRLVTDRLELLETRQRVSLNTRFTSVVSFGSMTGLCMTPLLWFEARHVRRHQPAAAELMDSFGATFRRSPRRWARTLSWSAVVLLAVFVPACEAMWIYLLAAEGGHWTHIFAHNQVLVVPIIVSCLVAGSAYCVKRFADALRTTMSKVTKTCSAFAGHSGRYQRGLLLQVRVPCPQELRRPWLVASRVDAYRRAWLQLRDLTERLVRTPLTAIAMLLHMLLMSTLAVYQALIGLLEGEVGKDFLYPLETFVGLCVPDAARVEVNLFREVVATSALRVTVGGYWDVTRATMTSVLSATLTYVIVLVQFREPAYSSASDYRNQSALQSGVKGE

>FoccGr20F

MKKMYPPAVSPPQSGRDVFGGHGIFEDMQPLIFVMALYGMLPVSFKPLRVSFHPLRSTLIHCVLVTALVVWSAFRLATARLEFYQADTDPLETQSTFTSVVTLVHLSGLAMLPLIWLEARNFRRNETLCASLMAHFPNVRKTCRMARTVALTAALMCLVAAPMTVGVWMTVVTTEAVHWTHVFANHHVVQLEIVVTALNFCLFQCVKNFAMDLCSTMVQELSLPYPSPCRVVEFRRGWLALRDLTQGLVLMPLTQLCMMIFSQLMFTLNSYLGLVCVFDQRLALAALIFCMGLQFISMLLVICDAAQRAKDVVGKGFLQPLETFCTLRSMGEATADEISNFREIVASSPVVVSAAGFWDIDRHSLSSIMATSVGYLIVLIQFRASPESVLKTLRNSTTTVDVA

>FoccGr21

MKRPFDVHVFDGLQPTAYLVALYGILPVTIRSWVPGFNPRHSTLCYCVCVQAVVLWRIRQAIWKALLIFKYFLWFPWGLIDTVPSKCQIPDRFDSFSSVFLKRIGAEFGHIVRVVKKPRVGAWMLVLVSVMLPLQELSWMTFVEFSLADAANSPVHVYVNELLLVGMMVPALCSLSLHRLASALAQELAKELTCSHVMAQRVRTYRHAWIQLSSVINGLILTPISMCMVVGMVVVLITMHTYEALEKLMAGHLQVSLLAFLVGMPYVILMLTLCETSHRASKTVPNNTRNPKTRFPGCSKPFFRVVLHRFLETVWWNSDGTAVGGFFKLKRSSFVAVMAAVITYAFIMLQFHLSVLSPSALSD

>FoccGr22C

MELCCPRRRAHVTDKSVHTGGVPTPDDVPPPGASTRIFDKIYPLVILVAAFGLLPIAWKPVRVSFDPRRSTLLYVVAVQGAVLWSVRTAIVARARVFVTPGASLLQRFTSLTSLGWLGSVAILPVAWAECRKPRRFLEMSRPLEVGVTRSSTFVEFPLEALVVAPPYVAGWLMVVAPDGADWTHFLGHLYQILISTTCVGCVAGTLLTVADFATGLKEQLATELLAGGMTATRVREFRMAWVRLRQLSEGCVAMPTTVLALLMYLIMQTTLCTYQGLVSSLAGFHTFASICAIISVQIFAGVVIMCDAGHRAAQAAPLPHATSRLKLIDANYCYVALN

>FoccGr23F

MASSLPLHHTPAPPRRPTRAVAPLQKHVSVHLSSTFPVAPLVEKVGVILWAWSVQTVLTVQAADARTRTQAPGAPSKPRPEPSRPLFDGLQLIVYSGAALGIVPISFRPLRVGFHPVRSTLPYALLVQAYTWWGAFFEMPWIGRSLAPQVLEGAAIRFWNMHSTLHLAGCLIVLVFWMEAGNIQQLRAVCDSLLEDFGYVLWGRPDPAGHADPAGRPRPRTPQRRVVWFTTVGLFILGPTMYLTWALVVETKKNVSEGLPHVFVNFLLSFAVAVPTICCLRLRTFASGLLSALRQDLANPDLTPERVLKYRRTWTHLTNLSSGFIVTPVSMLLVTVLLVILSTLHSYETVERLTAGEAQVVGCMIWLIGVPFLFLLFVLCESVVDLADVIRRRFMNVLQMSYCRYRDDRSYHELHRFLEIIWWNSPVMTISGYFELQRSSFKAIVTVIVTYVIVILQFHLSINNSAELTRWNGTQAMLSVNESVTATVTNKHKT

>FoccGr24

MASNRALRHPTPPQRPLHSVAPRQKHVHFVCTPPDSPSTTPKVFAVRMASASTPMWGSPVKACSVAAAPEEPSRPFFDGLQPLAYTGAALGIVPVSFRPLRLSFHPLRSTLLYALLVQSFTWWELFYVMTPWRNGQPTAQDKFPNNKVVSCFESVHTTVHLTGSVIVLVFWAEVHKVHKIRSSCGNLLVRVDFGYVLWGRPKQPGPRRRPQHGLVWITTMALWVMPGAMVILWSEYVETGVEPKNALAHSLINAQLGFGMIVLSLCFLRLHSFATGLLSELRQASLHDLADPSLTPERVLEYRRTWTLLADLNGGFIVTPISVLLQTVMLIFLVTLHSYEGVECLLADNLKVGLIIWVVGVPFPILLFALCEVPHRSADVIRRQFEGVLQMSNCRYGNDKTFEEVHRFLEVVWWSSPETTIGGYFELKRSTFRAIVAMIVTYGLVLLQFNSSGHSTDPNIFNGNLSTQG

>FoccGr25N

MTASRLEEFRLAWINLRQLTLDLGSMPLTASAMVLYLLVMLTLATYQCAVSYYNNYTLLCFGSAGVIFLAQFALIHIFDTAHRLKDTMSTGFYEPLASKSWRGLPADLLTEHQQFLHTISTNPPEVTFMGMISVKRPTYLSIMAAIATYLIVLFQLRMSDENLPTKATANSTESSLDSLFSSKTSK

>FoccGr26F

MKQTVKTVGPILWIFVVYGLLPVQFKPARMSVHPLRSTLLYVLAVQTLITWSAHQVYASFSGLLHSEPENARGVDERLLATLSIVHVLSHVIVLVIWFEVPRAVKCRVHLVDFAKNYFQMVGEHPYSKGTCFMRWYALAFCFVVAPITVVIWCHYILRKPFLTHGPAHLLQIVNVFSCLVVFALACMIVRDLASKVAAKLKEELDSDRMTASRLEEFRLAWLNLRQLTLDLGSMPLTGLSMVFYLLVMLTLTSYQSAVSYYNNYILLCFGSTGVLILSQFALIHICDTAHRLKDVMSDGFYEPLASKSWRGLPADLLTEHQQFLHTISTNPAEVTFMGMISVKRPTYLSIMAAIATYLIVLFQLRMSDEDLPTKATANSTESSLDSLFGSKTSK

>FoccGr27N

ENYIEMVGEQPYSKGTRFVRWYTLAFCFVLAPFLVVIWCNYILRKPFLTHGPSHLLQLVNVISSVVVFDIACMIVRDLASNVAAKLKEMFQAAHVYKWSSRMSPLGHTVLGGKRSSVSSYASTGAGSDATEAYARAYVQNEVGAAARLRSTETSLADGRLHSLHSGTPGPPAPAPLKTEEQNRSLMDNNVMRQTVKMVAPILWVFVVYGLIPVRFKRELDSDRMTASRLEEFRLAWLNLRQLTVDLGSMSLTGLSMVFYLLVIVTLTSYQSAVSYYNNNILLCFASTGVLLVAQFALIHICDTAHRLKDVMFTGFYEPLASKSWRVLPADLLTEHQQFLHTISTNPAEVTFMGMISIKRPTYLSIMAAIATYLIVLFQLRMSDEDLPTKATTNSTESSLDSLFGSKTSN

>FoccGr28N

ELDSDRMTASRLEEFRLAWLNLRQLTLDLGSMPLTGLSIVFYLLVMVTLTSYQSAVSYYNNYTLLCFGSTGCLLLSQLALIHIFDTAHRLKDVLSTSFYEPLASKSWRGLPADLLTEHQQFLHTISTNPAAVTFMGMISIKRPTYLSIMAAIATYLIVLFQLRMSDEDLPTKATTNSTESSLDSLCGSKTSK

>FoccGr29

MDVIPRGTKHMKFSHFAYMYQVPHLKYLLLTHRDRLAMLMCSLGPTTFLTLRSNLAWVLVVNSIVLWSGWVTIRDRLRQLEELSADRLDHVADRFVAIISILATFNYVIAPLLWFEMRRSAFTFLQFATPLMRDFGYLVPRCGSRPTILAWLLLVMAVFTTPSLVVVWLFVVGKGRGSNWRHLPAGVYNIVLCFSLTLGLVSYFKRISDFANQMALQLLREIKTGKMRAQRLSEFRASWIDLRRLTQNVCLMPVTTCIYICILVNMSTFFAFQTAVAVRLSSEPDLIFSIVIMMVLWNGSALILCMAAQEVSDVVKMKFDKILRVDFYLIYPIMDSVTQREVKFFAYLTSRSDVRVSVGGFFFLRKETITSIISATVSNVIVLIQFQRSNKFY

>FoccGr30N

MWNFFRKCCFPLGPAFRWSTRLSVSFHPFRSTILFVVGCYLLSSWSCYKLVTRRLQQLSEVPLGPHTLGQRYIAIVSIICGGMCLSTWSVVLLEARRLHRLRALTRPLMELTHPTFGAARLLQYRAAWLDLRALITEIFVMPASCLLYMLNSMAFTTFMAYQTTVGIRARNMVVIGNMVGYCVTWLITLFFVCYGADKATEMVKHTLLDVLLRGAGAQRWSFEVYRELKFFTRIVRGSNIAVSMAGFFELNKRFYSSVLTAVITYVVVLLQFQQIEEKSLKTLREDPL

>FoccGr31

MRLYRKELRADSAVLMKISALELMSKECSSASALSRHQEDVLLGHSHPDSFRSAIRLACVIGQVFSVLPTSTRPDIVPRFHYVSLRVLYWAVVTFLLGFNAVVAVVNVVEAQSGRNSIVASSNYIFYVSALTVSVLYFRVAQQWRELCLCFLRVEHRMRRFGYPPRLRCRLQTITVVVLSLALTEYSLAIVSGVLAMDLDKPVNEFFKDYYHGNYPQILLLIPYNPIVLLVCQLVNAIGTFMWNYNDMFVMLISVAIAQKFKLLNTKLYELPKKDKTREFWKEVRETYNSLSLLTREVDQILSVQILFSFMNNLYFICIQLLHTVKHEDNSIAKNIYFVFSFGLLLLRTACVSLCAASINDESKQVRKFLYSVSSRTYCEEVDRFIIQATTDEIALTGLKFFSLTRSFLLTVTGTIATYEIVLVQFSGNQKTV

>FoccGr32N

FRWLSPRAIYFIVSALAFVAMSVLLVLSHEEDLSTSVXNIVHTSSVLAVLFLFLRLSRRWHRVCAGFQRVERRLATEGFNAVSFGSYIRSMTAVFILATFVADHLRVCKIVKHNHMHASYTSFFSMKIIPVYNIALVIWVFWTKAQAKLAITYTDLILWAIGAAAAAKFKSINKKIGELKYEKMTEMDWRRVRESYNDVVVLISDLDRILGPHVLLSYAIGLWFVCGGILLLLDSEIDTLKRVYIFLSMSFWICRVIVATLSLASVYEENICIRNSLHLVPTDYYGDEVHRFAMQACHDDLGFTGMKFFTVTRSVLLTLAGTIASYEIVLMEFNR

>FoccGr33F

MKKHSSLLPDLFGSGGFAGQTPASDQSLLTAPAHPDSFYAGARAAVRIAQCFSLIPVKLSPGREPRFRWKSFRSVYAILVILAFLARTYMSAVQSDTAMAIAEDVTLQLFLCFIGISFFMLGMRWRQFSLLFLRIEKHSENAVGDAHPAGVGRRMWLVTVGVFGCALVEHVFDRFTEVMNIMEFGNYTSVQAYLDDTLFERPFWEAALKVFEFISYTYWAVLVAFNDLFTMLCAVYIAMRVANFNSHLAPTLTKNKDKYFWEDARYSYDALARLTNDVDRFISGNILMTFVRNLFYICFQLLYFLRASSDDGLWRHIYHIGTFGYVLMRTLAVSFFAASVNDQSKLCKTFLYSVSERSYCPEVHRLLEQAANDEMFLTGYNFFRITRSFILTVAGTMATYEVILLQIHVRK

>FoccGr34F

MRACSLVSADGVIVLEMPSASRPLARYRQQAAKLPSAFRVVSKKLGGRPRPGRVLALAPPPGMHHLSQFQCWKLDAGRPTTTPVADPVDDELLTTKPAPNSFRAVIGHLIVTGHIIGILPVSCAVDALPLFFWSSPRVVLSLCTATVVLLRIGMYVWSVYRGYMSAFDTLVSTRLGFHILSTVELLLYVRLARRWRPFVLCFLRVERRQGEDALSPRLRLQAKVVTGVLMSFALGELLLQVTTFALPKQARDYPSFSALLQAHYEALYDNMAQALPYSVPSLLLAEFTTAATNFVLNYNDVFVIIVSIAIVRRFKIIRGQLGRATTQDDAFDWKHARESFNDLSQLTREVDNTLSSLVLLSCLNNFFSICIQLQHTVRPSTYNPGFARKVYFLFSFTFVMLRSAAVVLSASCIPEESKAVKRVLYAVPTKQYSEEVQRFLTQVTTDEIALTGLGLFSITRSFILTMLGTIATYEILLVQMYRTSGSEQLPLC

>FoccGr35

MSARSERRTTSTSSSSSGNVRKHLRVRVDTGPGGSQVSSLSSPRPKGSKAAPVAHRDSFHGAMCPALMVAQCFALLPVDGISRTHPGVPQFRWLSFRVFYTIAVFLGTGVVALLAAYRMVSSGLSFNSSVTVVFFGNSLSAMVLFLQLATRWPAVARAWARIEHDMRTYGYPRHLGRKVKIMTAVVLLLATVEHLLAVMTAVYTSWPCYKSGGFPGLVRGYSLNNFAMVLQVTEYAPWKAVFVFLLNVVATFSWNYTDLFIMLLSVSLAQRFKQLNHRLRALKGKNLPSGVWRQLRETYNSLACLTRTVDEAVNNIVLLSFASNLYFICVQLLQGMKLILDVYPYVYFFFSFGYLLLRTCVVSLYAASIHDESKEPRRVLYSVPSESFCTEVRRFLAQVTTDDVALTGCNFFSVTRSLMLTVAGTIVTYEIVLVQFQGVSGDGTTDLSVKPANLTVRCF

>FoccGr36

MCSSEMLEAARTSNGHDQCALAQQALGQALGQACLPWGPGSSPCSSPRFSPMNCSVSGSPRHVTPRVSERHPLASRNVRPRMPLTTVSSLSPMPLPPRLPRTPPCPKPAAESGGGPVSTVALGACGAGGWWCPCPPRCPWLHIQPADRHADERSRTPPHAHHDHDPDPDLEGRRAPLTRVHPDSFSGSIRHALIIGYFFSLLHPELGPSLSQDSSRTRRWSGVRILRLLYAGVIVVGLSVQAALSLYREMTADSLWDSSVDMMFFGSATAATVLFLELGPRWRRLTRSFECVESHMVDFGYPPLLRRQMKRLTAVVLLGAAVEHILSVWSAVDFTWPCYMEAGPGGLLKGYSLLKFPMTFTVMAYEHWKAVVVLFFNFVGTFFWNFNHLFVMLISMPLSHRFKQVNERLEAVTGKVMTDDFWRRARETYNELSVLCREVDDAIGKLVLLSFASNMYFICRQLLHSFRTLRTGFWYNLYFIYSFGYMLLRTVLVSLFAASVNDQSRRPLRVLFGVPAEGYTHEVERFMSQVTRDHVALTGCRFFSVTRGLILTVAGTIISYEIVLMQLHPMVAGTPHKTDHTEHCGL

>FoccGr37

MSTTGSARRGTWRFGSVHNLPALSGPEAPGGGPGAHAGVPEQGRTRAWTLPTKPVAVVPSRGLLLAVGPRLQGLQDASDQLKDDRDEGSAILRALRPALTVAQVFGLLPVSKPSDARHAVFRWKSARTALAVTIIVGAVLESMLALMRMVETGIKYSTSVHAIFFVVTSVGQMLFLHMSGQWHGLLVDLARVDGTCQRAYGEPGRLRARTLAVTVVALSLAAVEHTLSMLGSLYFVWPCYQRGGILFVWQGYFLTKYLKTFTVTAFATWKATLLLLLQFINVFVWNFMDLFVALFAMGLSSRFRQVNAELQATKTEVLSSEFWKSKRKLYNELADLVRRVDDVIGPLILVSYATDLYFICLQLLSLVNPSGVSVYHQAFFSFSMGYLILRMTFMTHAAANIHEESRGPKEVLFALPTGAYSVEAERFLMQVLTDNVAFTGCNFFSISRSFLLKVAGTIVTYAVVLVQFNSKEANPDAAQSNGTILCTCEDISRDYKC

>FoccGr38N

VTTAAVLNKAAKKQRLLLSSGKLGGAIAWSETPSTPDDSPPKAAGDTFLQGFRLAATLAQVFGLLPVKKPSDGENASFRWLSLRVVYALVIITGSAVQSVFSLMRMIEQGINFYTSVNVVFFVTAAITSVLFLKLSTSWGDTFSLWLTLERFVTAVNGHPARLVSRMKVMTVVVLLLAALEHTLSIATSLSSVSVCYDRGGLPLLVEGYFMAKYRKTFTIISYSLWRGILLFALQVITTFIWNFADLFVILVSMGLAAHFRQLNAALSQARGKIRHEGFWRTRRESYNALAVLTRAINDYVGPVVLISYASNLYFTCLQLLIVTKRKSTTNSLYNFYFFFSLLYLIVRTVIISLQAADVYLESQKPRPVLFSVPSPSFCREVQRFLLQVNTDNVALTGCDFFSVTRSFMLTVAGTIITYIVILVQFDSNATGDDTPHPNRTVSCTCTDLKIYNC

>FoccGr39

MTEGPLCVHRRRCHTWGRCRHLQLRATLARMAAEARRRLRCGGGVTTSPTRSASFASRGSCDVTCISSYELGPKLPQLVPDAMPAVVPLGSTFTVPPPSLGGGPLVAVRDAEGQGLEERPPTFSRVARRDSLTEALRPALVLAQAFFVLPVHAGRWSRRARTATLLAVMVVVIALAAVHMFKTELSLTSSSHFIFYGTAALSVHLLSGLSLQWPRLSSCWSNSGSLLTEGLEDDRRTARICKRITIFMMSCALVEHVLSLLNGLMPLLVCEAHGGAWTVYKSLFLNGYPMIFEVLPFSIPVSIIVIYLNVTCTFLWNFIDLLVIIISYSLARRFDTFSQFLRQAKGKNYSIFWWQEVREAYNRLSLLTKQLDDQINGIVLLSFASNLYFICLQLLYNMSRFEQEKSVFSRLYFCFSFAWLLARAALVSMTAASVYEQSRAPTEVLYCVPSKSFNAEVQRFLVQTTTDTVALSGARFFWVTRSLVLTLAGTVATYEIVLVQFHDDMDKVQRPSNVCFP

>FoccGr40

MSRSALQKGESFHEAMSGILVAAQVFGLLPVRGVRRPSPLDLRFERRSSRVAWSLVLFVGVCLVELVSLRHMALTLSADTFGVIGGIGAATAGAVFYGNALIGALLMFRLAQRWPRVMEAWADMEATLRKGRTPKLRLRFSVLCAVVMLGALVEHTMSMLNAVPIFIGVHFYGETRPPPEDGQHVIDTSSGVLLEFLDEYCHASHSFIFNHVPYNAFTGLLIFIMSKVTTFTWNFVDLFLMLVSTGLAERFKQFNAVVFESDVAMLNKEDWRELREYFATLSALTKNVDSNICSIVFLSFGNNLYFICLQLLNGLSPGKASWLATMYFFYSFGFLLGRTVMVTLITARINDMSKEGLAALFSCPSETYNVEVARLQHQMASDDLALTGLRFFSITRNYMLAVAGAIATYEIVLLQFNVAISPHPTQPPPDGAVVVTLPTAGR

>FoccGr41F

MGTLLSGAMDDTDDDDDGDVPPTYGLFQAPQGDASSPVPPEEHPDAFANAVHLPVSIGQALSLMPANSLTGSWSSQRAAYSVGVSAGLTVLAVLSAVRQFSREKVSIVDVMVVVFYGLSAVTSWLFFRFARYWGLLLRAFRAQERHLGPCPGLRRKVVFITVLLLGVAFVEHGLSEMEITLNLRRSHPNASTFSEYLEIYYSANYDMLIVVIGYWPLLSPFFFFLNFTATFVWNYNDLFIMVVSVALAARFRCLRDSISRSIGGTVGTKAAPVLGVAYWRSARRSYNALSRLARLVDHALNVQLLVSFVNNLYFICMQLLRGFDNPQNPNDTWWEELYILFSFGFLVVRASAVSLFAASVHENSRSVQDVLYCVCSEDYNEEVYRFLVQVTSDDVALSGCSMFKVTRGLLLTLAGTILTYEVVLVQLH

>FoccGr42

MAFTSERAGQGHASPGHWRHSRSLGWGRCCLVCVVCSRSPPGGADSPAMRAWGAGAAVRPVWSAVRPALMVAQVCSVLVISLVSTASRLPPWYTAHALLVAAFVASMGATFAFSEPLLAVPERGIKNRVRYFSSLVGYSEVVTLVLLWLRAAGRWAGMWTHIGDLEGRLWGRGERHLPRLRSHTRAVTCVFLIVHIPQEASIVWRRLASVSSLYDLGQMTCSLCILLVFNVSAMLLVVLSDMLAECFRALTAKLHCASKKNGGQIPPGVVEDVRVLYNEVRRLTDALDDILGAILLVMLFADGLFICRFLLHWFTPSVHTLYAYLTVGIVTARFVFTIHTVSSVSSEWESTRSHLYRMSPHLCGKEVSRFILQVTTDQISLTALGIVPITKPLMLAVWAAILSYEIILVQFNKGN

>FoccGr43

MVPRERGHTVATPRRYLVWSAIRPALRTAQVWSLLFISLASPTTKLPSWHAAYALLVCAFILAMGGCFFAEFLRTPPERESQERFRFFVAVPGYFGCVALVLLWVRVAGPWSGLWARVDELEDRLWGRGVHRLPQLRRRTRFTAYVFVLVTIMQQVIVLLRRLPPSGLYEWALLVCSLGFTGAFNFSALLLIVLSHALAECFRALTAALQKTARQGDGQIPTEAVEGFRVLYNEVCGLTMAVDHCVSPVILVAIFTDGLYICRFVLLSLAVSAQNWYSSAVVTVCFFRFVCTLLTASSVHSEWQTTKSCLHRMVPCRGAKEARRFFMQIGTERISLTALGMVPITKTLLPAVLGAILSYEIILVQFQLQKLKLVFSSH

>FoccGr44

MAALDHDPEAAAAPCQRLLWTAIRPALLTAQICSMGLFSLASTATRLPSWHTAYAVLFSAFVLCIGAIHSVEFFQTPDKRRNRFKFISAVPTYLESVALVLLWVRVAGRWPGLWARVNALESRLWAPEVHRLPRLHRRARHVSCVFALVASVQQPSLTWRRLPVRGLFEGFQLILSLGFTSVFDFNAMLLIVLSGVPAECFRALTDTLRSAAEQGGNRLPSEAVEGVRVLYNEVCRLTLAVDATLGPIFLVTIFADGLYICRFILFLLTGSVQSWYACATVTVTASRFVYTILAASSVHTEWLATRACLHRVVPCRGGNEVARIFMQIGSDRIALTALGIMTITKSLLPAVLGAILSYEIILVQFHLQMIGHGFSNHSTAYS

>FoccGr45

MAPPERGHYVAVPRRHLVWTAIRPALRIAQIWSLLFINLTSPTTKLLTWHAAYAVLMTVFIMVMGVGFVAEFLQNPPSKRQGDSFRFLVAVPAFFGFMAMVLLWVRVAGRWTRLWALVDELEDRLWGRGVHRLPRLCRRARNSAYAFVLVNIVQQTHVLIRRLPASSLYEWSLLICTLCFGGVFNFSTLLLIVLSHALAECFRAFTAMLESAARQCDGQIPTEAVEDFRVLYNKMCGLTLALDHCVNPVILVAIFTDGVYICRFVLLSLAVSAQDWHSSTLMTISFLRFVWTILAASSVHSEWQSTKLCLHRMVSCRGGKEATRFFMQIVTDRISLTALEMVPITKHLLPTVLGAILSYEIILVQFQLQKLKRVFSSQKSTLT

>FoccGr46

MAPDARGHTVAMPRRHLVWNAIQPALRIAQICSLLFISLASPVTKLPSLHAAYAVLMTLFIMAIGGAFVADFLQTPPSERHHDSFRFLVAVPAFFGFVAMVLLWVRVAGRWMGLWVLVDELEDRLWGHGVHRLPRLHRRARNASYAIVLVAIVQNTNVLVRRLPASGLYEWALLVCSLCFSVVFNFNALLLIVLSYALAECFRAFTVALESSAQKCDGRIPPEAMEDFRVLYNKVCGLSLALDHCVSPVILVAIFTDGLYICRFILLSLAVSAQNWSSTMLATISLFRFVWTILAASSVHSEWQSTKSCLHGIVPCRRGVEATRFFTQIVTERISLTALEMVPITKPLLPAVLGAILSYEIILVQFQLQKLKLVLSH

>FoccGr47

MVPLDQDTEAAAAPRRRLLWTAIRPALVTAQICSLGVFSLASTATRLASWHTAYAVLFSAFTICVGASFSVHFYQTTYRRRNRFKFITAVPTILESLALVLLWVRVGGRWPGLWARVDALEDRLWAREVHRLPGLRRRTRAVSCVFALVTGLQQPVLTWRRMPVRNLYEGFLLIMSLSSTGVYELNAMLLIVLSDVQAECFRGLSAVLRSTAKQGGNQLPSEAVEGVRVLYNEVCRLTLALDASLGPILLVTIFADGLYVCRFILLFLTGSIPSWYACVRVTITVFRFVYTILAASSVHTEWQATRSCLLRVAPCRGGKEVGRIFMQIGTDRVALTALGIMPITKSLLPAVFGAILSYEIIFVQFNLQTIGRAFSNSTASS

>FoccGr48

MLGLRGHGKALLLGAGERPPRPRPRHSWVSPGRHTLVAVRAPPPPRDGRLVWPAVRPALLTTQVCSLLLISLEPTTTRLSWLRVAFAVLVAVLVMAMGCAVEVRAGTSNKRSTPFGHYAALFNYAMCLALVLLWARAANRCSALWERVDELEARLWARGVHRLPRLRRHALAVTGVVFLVLIPQQAFVVWRRLPLRGLFHLGQLVSGVCIAGVFNFSALLLMVLSEVLAECFRALASLIRRTAAQSGNRIPPEVVEGVRVLYNEVCGLARTLDHSVGAIFMSTLVADGLYICQFLLSILTRAAGRSLYSAATLAVVAARFFLTIFAASSVHGEWRSARSCLLRTSPYRSGKEVLRFFMQVATDRVSLTALGVVPITRPLMLALLGSILSYEIVLVQFNNNNS

>FoccGr49

MYIEREWLWRWGRALGQVPFSIGRGCRCPPPWVGGRCRCGKGLLPSRPWEAYSLALVLLCVPSTFANLIFYYQEKVSQYTYNKTDRAVSHLDVLVVMLSGNVAVLRALLYRRHFRLSVEEVTMVDRLLRARPPPVPLSRVLWLASFLTLSALHLAYVSFVGGALHALSHVPNQFNYQLIYTMEALFAEETDSIFRRFRTLNARLESTCLGAASRYPLDLHPGMIPVGETSRKLPAPTHALRRGAPTDKSSTRARTKSFYSGVPEFDGVPIHQLEDAHALLCSSALNLTRRYGPVLLIDVLNLLLHFVTTAYFFFSMIVVDAKLHQNVAHRITMYVYVTLQGLWMLAHVSRLVLLVQPCSNVKEEANETGALVGTFLNMLHPCCHLRKQLELFSMQLMHQRVGFSPCGLFELDLKLLLSLLGAVTTYLVVLFQIRVPSTNVDINTLNRTV

>FoccGr50F

MYLEKEWLWRWGRSVGQAPFSLRRRCGCPPPRVGRRCRCGAAAALLPSRRWEAYSLALVLISVPSTFVALFYLRQKAEEMNPSPSTSTKTDKMAERSDMLVVMLSGNVAVLQALLTRRHFHVFVEEVTTVDRLVRARTPFVPLWLLMWLSSVVMLSVLDLISVYDAGGGLHAASHVPNHVNYLLVYTMEVLFAEDSYSICSRFRTLNARLESACLVAAARYPLELRPGMAPVRQPSRQRDPWSDKWSVWARTTSNSDLPLVPSVSGALELDGIPIHHLQDAYALLCSSVHNVTRRYGPVLVIDVLNLLLHFVTTSYFFLSVLMYDARLQKQMEQQHQHRREVYVTMQGLWIFAHLSRLGLLVHPCSKVKEEANKTGALVGTFLNKLNPSCHFRRQLEMFSVLLMHQRVSFNPCGLFELDLSVVVSLLGAVTTYLVVLLQIRTPS

>FoccGr51

MASRAHVRAAVGLLRRFTRWAGGDYYDGTTLVQRGTRGWLWQVGRVMGLLPISFKNCVCPSQPAFLSSRGSDRLGDPGLPPSWDATGRIGGVKECSCPVAARSVPWMMYSCGVAAVLLPMSMYAVYSDRAEARAGRPSLRMSTRTDELVTFCDILSNSFCTLVCFLQLAYQMQKGSLFSLLDQMQQVDRLLQIRPPAVPGLLALWLLLLILIMFMDASMWTEMNQGLQYAATYLPFYGVYLNTFVMEVLFIDEAHSINMRFKALNQMLDAALALPSYQVAVAAEQPTPMTMEQGWMRSSARSRRMSIAALADGTLYPSKLGTAAAAAAGAGSRRSRMRPEPGGGAAAGALVPIQRLNMAHHMLCELTFTIARRYGLSLLADMLCLLLHLVITAYFLLKHLISETVLVSGQYIVMQAVWLAAHLGRMLFIVLPCSRASAEAGRTGTLVGVVLSTTAPSSHAHRQLETFSIQLLHHRVNFNACGLFELGLPLVVSILGAVTTYLVVLLQLKGAPGESTANSTAVVS

>FoccGr52

MESFERTNDWLWPTSRLFGLLPMSLQRCQCAVQRRRLRPLERLSVPGGPRWARCWCPPLAPSLGWLLYSLSLMTLVVGWTAYKLVHKYADMSLLDEALAANVTSTSGQETTRLVGLFDMTTISVIAAVCVLQAATTQHLVLRLNAQLRRVDSLLAPRAYAVPRWLLVFLMTLSFVALLDLFYKLCDDEQYAIHTMPSYLAYIITYMREAMFVDDVSGVRARFRTINAELTDCLSDYPAAEVHETVVNTDRKSRAIGRLATAYDLLCQCVESVARQHSLFLLCNMLGLVVRLVVTTYFIVDLVLESDAFAEVRPEPPPSLRSMFVTIQVLWLACHFLRMLALVQPCSSTIEEALLTGSIVSRSVSYEMLDGQISVQLKSLSMHLLHRKITVSAFGVFDLSLPLVCTVMSAVTTYLVVLIQLKLPSPVAAP

>FoccGr53

MGLHRRWLHRLGLLLGVFPLVERSPAGAGRTKIQSGTGTATRPFLQSPILLRRSCALLCYSVALVSAVLGYTIYSAVEEFLAVTQNNEVRWTKRTDAYVSLCEYTTLWAATGVAAASSWLRRDADIRHADLLSEVDVMLGLPPPRPQLWLAAWVAAMLSVLVADKIVWFMTEQDNYLEDMPLYVTYFTCFVSETLYMSDAQAIQTRLAAINQVLEEASPRPGHRDAHGSPVVLPGTLTVSRLATAHSLLAKATQTVSRRYGLALLTDMVALLINLVITCYLFLEHMLPLSEPPEDYTRFFMEIVFACLHFSRIVMLVEQPSAATYEADRTGALMGYLGSTVPYGSMMHEQLKALSVQLRHQQTSFTNCGLFDMNRTVIVSIVGAVATYLLVLLQLKSPSSTKGQPSNST

>FoccGr54

MAHSWRPRPWETVAFSRRVMMAHWHVVLVALQLVGILSCSGWPPRWSRHWACYGYVSLVVLAVARVALFAFLFFTVFKVETARKSSYPNEISYIFWLGAFNLCIKGLTVILTQVVMVLHSRDVSLLVRAVFLYLHSFPGPRHKHGCRRAAIAWFGANFAFNSIYSIIYANLFVHTWDGMFFILNDMMPDVLISATQDSFLHVVSANVDFLAAIADGVHEHVASLPRVAAAGSVRAAGAPGTACSATCPAHCGRVPVLLGEQGDARVLLHGLRTLRVRYQCVQDVVHATNWVYGWFNLLAGTATLIGGVVSLYSGVMLTITFAGVLGDYEDPLPFNAFDGASCLVLGTLLTTRLLVICAVGEQMSSENFRISSALQTGLARHPGIDLRIEREIQSFLRQTQMQEIRFGAFNLLYYDLDTIKRTIAACMTNIVILVQFSAISAQGKRREQYL

>FoccGr55

MAHSWRRPRGSATRSATGSLATSQRVLVAHWHCMHVLLQLAGVLGCRGWPPRWSRPWTCYAFASLVALSLVRATLLWPTVQAYPEISKNTTISDDISFMFRLFKCDIGAKCISFVLIQVVMVHYSSALSLLARAVLLHVHSFRAPKRKQVARRAVLMCLGANLAFYAFHCIYHEDFYFRTWLGGAQFVTGLLPTLLISAIQDSFTCIVTASVDSLAGVADGVREFVNTLLHRRGPVTRPRELDELGGEQSDARAPARAGLRALRVRYQCVHDVVLATNWLYGWCNLVICSFSLLRGVTFLYCGVILTMARAAMLGDVGEDFDVAFDAADGVFFLFWGTVLIGKLLFICAIGEQMSSESSRISDALQAGITRCPDMDPSVKREIQWFICQTQRQKIRFGAFDLFYFDLRTMNRIIASGMTYIVVLVQFSAIAAHGRGQLKEQSNKTV

>FoccGr56NCF

PEISKNTTISDDISFMFRLFKCDIGAKCISFVLIQTVVLHHSRDLSLLVGAVLLYLRSFSGPKGKHVSRQAVLLCHGASFIFCMLYYIGHSNLYFTDWRSVRMIVYGLMPTMLTSTIQETFVHIVSACVDFVAAVADGVQEHVDALKRLEARVQAGSAHRRTGRVTAVTESLHALDELGGEQSDARVLALGLRALRVRYQCVQDVVHATNSLYGGFNVITITTSLIRGVVFLYCGVLITIVFAGNLVYYGDGIVVNRPDGAWLLGWGTVVIGKLIFICAIGQQMSSESSRISDTLQMGITHHPDMDPLIEQEIQWFIHQIRVQDIRFGAFDLFYFDLSSMNR

>FoccGr57C

MAHSWQRPGEAAVSWSVALSRRALVAHWHFLVVVLQLNGLLGCSGWPPRWSRAWMCYALVSLVALSTARAIMLLAVIQAAVNVSKNTGTPDDISFIFYLFTCDIIGRAISFLLVQVVVVRHSRDLSLLVRATLLYLRSFSGPKHKRGSRQAVLLCLGANFTFYMVYCIKHLDMYLTSWPGVRLIVYAMMPSMLASTIQDTFVHIVSANVDFVAAIADGVQEHIDALLRLGARMQVGPAHRRVTGRVTGSVIAVTDRVRARDELGDEEGDARVLALGLRALRVRYQCVQDVVHATNWLYGGFNVITVTISLIRGVVFLYCGVLVTLVLAGKLGDYDYGDIPLRESDGAWLLGWGIMLVGKLVFICAIGQQMSSESSRISDALQTGITRYPDMDPTIEREIQWFIRQIRIQEIRFGAYDLFYFDLKSMNR

>FoccGr58NF

ATNWLYGGFNVITVTISLIRGVVFLYCGVLVTLVLAGKLGDYDYGDIPLRDSDGAWFLGWGTIFIGKLVFICAIGQQMSSESSRISDALQTGITRYPDMNPLIEREIQWFIRQIRIQEIRFGAFDLFYFDLKSMNRIIASSLTYIVVLVQFSAISAQGRSHQKAPSDKTC

>FoccGr59IF

MAHSWRRPGEAAASWSVALSRRALVAHWHFLMVVLQLNGLFGCSGWPPRWSRAWTCYALVSLVALNTARAFTLLAVIQPEIDLSNNTGISGNISFIFRLFACDTCSRAISFVLIQVVVVRHSRDLSLLVRATLLYLRSFSGPKRKRVSRPAYAGRKRLAVFVTPGHTTCKTYIQWFIRQIRIQEIRFGAYDLFYFDLKSMNRIIASSMTYIVVLVQFSAISVQGNRPSKSTM

>FoccGr60

MARSRRRSLVLTRRVLVAHWHCNMVVLQLNGFLGCSGWPPRWSRAWTYYALLSLVLFAIARAAQILSFIQLGVGAKNATRQDDISFMFRLFTYDICARCIAFVLMQAVLVHRSRELSLLVRAVLLYLRSYSCPKRKHVSRQAVLLCHGVHFIFNTIYLIDHSNVFFTTTRYVRFIVYGMLPTILVGMILDSFVHIVSANVDFVAAIADDVQEHIDTLRRLRPRVQAVTGRVTAVNETHHALDELGGEQGSEQSDERVLARGLRALRVRYQCVQDVVHATNWLYGGFNVIAITTSLIRGVVFVYCGLLITVVFAGDLGGRGDNIRINGPDGAWFLCSGTLLIGKLIFMCAIGQQMSSESSRISDTLQTGITRYPDMDPLIEREIQWFIRQIRIQDIRFGVFDLFYFDLKSMNRIIASSVTYIVVLVQFSAISAHGRCQLKVSRNTDK

>FoccGr61F

MAHSWQRPGEAAASWSVALSRRALVAHWHFLTVVLQLNGLLGCSGWPPRWSRAWTCYALVSLVALNTARAFTLLAVIQPEIDLSNNTGISGNISFIFRLFACDTCSRAISFVLIQVVVVRHSRDLSLLVRATLLYLRSFSGPKRKRVSRQAVLLCLGANFTFYMVYCIEHLDVYLTSWHGVRLVLSALMPTMLASTIQDTFAHIVSANVDFLAAIADGVQEHVDALLRLGARIRPAHRRVPGRVTGWVTGSVIAVTDRVRARDELGGEQSDARVLALGLRALRVRYQCVQDVVLATNWLYGGFNVITTTTSLIRGVVFLYCGLLITLVLTGNLGEHGDIPLRESDGAWLLGWGIMLVGKLVFICAIGQQMSSESSRISDALQTGITRYPDMDPSIEREIQWFIRQIRIQEIRFGAYDLFYFDLKSMNRIIASSVTSIVALVQFSAISAQRHCQRTAP

>FoccGr62I

MASSWQRPRGSATGSLATSQRVLVAHWHCILMLLQLAGIIGCSGWPPRWSRPWTSYAFTSLVVLSVVRATLLWPTFVAGLLPTLIISTTQDSFTCIESASVDSLAAVADGVQEVINALLRLGAPSAPRPETGPARCLRVSEATRRREPDELGGEQSDVVTHGLRALRVRYQCVQDVVHATNWLYGWGNLIISSFSLLRGVTFLYCGVIVTMAHAAMLGDLGEDFPFDVSDGVFLLFWGTVLIGKLIFICAIGEQMRSESSRISDALQAGITRYPDMDPSVKREIQWFICQTQRQEIRFGAFDLCYFDLTTMNRIIASGMTCIVVLVQFSAIAAHGRSQLK

>FoccGr63N

VFLVRNSRDVSLLIRAVLLYMHSFPGPRHKYVSRHAAIAWFGANFAFNTFYCLSYAHHFFSDTWSVAFYVLNDLLPNVLISAAQDSFLHVVSANVDFLATIADGVHEHAASLSRLAAAGSVIAPSAPCSPTCATHCGRGRARVLLGEQGDVFLDGLRALRVRYQSVQDAVHATNWVYGGFNLFAGTATLFSGVVFLYSGVMLTITFAGVLGDYEDPLPLDSFEGDCCLALGTMQIAQLLFMCAIGEQMSSERFRISSALETGLARHPGIDPRVEREIQWFIRQTQMQEIRFGAFDLLYFDLKTMKRIVAAIMTNIVILVQFSALSAQSKHREP

>FoccGr64

MSISVARRMSDGALLGPWPRPGGTATWSSSRRVLVAHWRAGLVALRLGGVLGCSGWPPRWSRRWTRYGCATLVAFTAARVALFTRQFFVMFKPQFERGRESSYPDEIAYIFWLGTFNLCIKGLTIILTQVVMLLCSRNVGLLVRAVFLYLHSFPGPRNKNASRRAVVAWFGANFTFTAAYCLSYTNVYLDNWERAFYVLNDVLPDVLIGATRDSFLHLVSANVDLLAAIADGVHNHVASLPHLQGHRGPPAGAPGSVFRQRLRSAREGGPMTLQDLQFGSVAPSASCSSSCPEHCARRVSAALEQGDDDLLHGLRALRVRYQSVQDVVRATNRAYGGFNLLTSTTALIDGVVFLYSGVMLAIAFAGVLGDYKDPLPFNAFDAAYCLAFGALQAARLLFMCAIGEQMSSESFRISSALQTGLARHPGIDLRIEREIQRFIHQTQMQEIRFGAFDLLYFDLKTMKRIIAAIMTNIVILVQFSAISAEGKHREQIV

>FoccGr65

MSDVAVLGALARPGPGGTTACSSSRRVLVAHWRAVLAALRLAGVLGCSGWPPRWSRRWTRYGCATLVAFTAARVALFTRQFFLAFKPQVEEARKGLYPGEMSYIFWLDTFNLGMQGLTISLTQVVMLLYSRDVSLLVRAVFLYLHSFPGPRNKNAHRHAAMAWFGANFTFYAVYGMSYTHFTQDDWERVFYILNDLLPHVLIGATQDSFLHIVAANVDLLASVADGVLEHVAALSHLAQSAAPCTCSSTCPAHRGRVPVLLGEQRAARGLLHGLRALRVRYQGVQDVVRATNRAYGGFILLASTTALIGGVAFLYSGVIVVITFAGMLGEYKEPLPFDALDGACALALGTLQTARLLFTCAIGEQMSLESFRISSALQTGLTRHAGIDPRIEREQLQWFIRQTQMQEIRFGAFDLLYFDLKTMNRIIAAIMSNIVILVQFSALSSGITKCNDWSSSNQPFLNETTPKEASYRRCPTV

>FoccGr66

MGDGAGALLLAASAAGTTTTTPAAAWSSSRRVLVAHWRAGLVALRLAGVLGCSGWPPRWSRPWTRYGCASLVAFTAARVALFTRQFFTDFKPQIEEARKESYPGEIAYIFWIDAFNLVIQDLTISLTQVVMLLYSRDVSLLVRAVFLYLHSFPGPRNKNARRRAVVAWFGAHFTFYAAYGLSYTDNWERAFYILNDLLPHVLVGATQDSFLHIVSANADLLASIADGVLEHVASLSHLAQSAAPCTCSSTCPAHCSRVPVLLGEQRVARALLHGLRALRVRYQSVQDVVRATNRAYGGYNLLASTTALIGGVAFLYTGVIAIITLAGILGEYKEPLPFDALDAVCCLALGTLQTARLLFTCAIGEQMSLESFRISSALQTGLTRHAGIDPRIEREIQWFIRQTQMQEIRFGVFDLLYFDLKTMNRVIAAMMSNIVILVQFSALSSGIT

>FoccGr67

MGDGAGALLLAASAAGTTTTTPAAWSSSRRVLVAHWRAVLAALRLAGVLGCSGWPLRWSRRWTRYGCASLVAFTAARVALFFHRIIVAFKTQYGTAQRSSSPDEISYIFWLGTFNLGIQGLAVSLTQVIMLLYARDVSLLVRAVFLYLHSYPGPRNKCASRHAAMAWFGANFTFNIFYCMTYTLHAYVDRWAPAFYVLHDLLPVVLIGATQDSFLHIVSANVDFLATIADGVLEHIAALSHLAAYVAPSCTCSSTCRVHCRRRVFLLLGERRAARGLLHGLRALRVRYQSVQDVVRATNWAYGGFNLLASTAALIGTVAFLYTGVMLTMMFAGVLGEYKEPLPFDAFDGACLLILGTFLTARLLFVCVIGERMSSESFRISSALQTGLTLHPGVDTRIEREIQWFIRQTQLQEIRFGAFDFVYFDLKTMKRIIAAIMTNIAILVQFSALSAGITQCN

>FoccGr68

MRCRSAEPRPAVRVASTARCTMDSRWGCALEAHWRAVLLGMQLAGMLGCRGWPPRASRPWTRYARVFTAASSVARSMMVYITIYSFWNGIRNGISNSPNFSGSISGFVYALYLLDFLIMNVGLTVTQVVMVYHSQELCLVVRAMLQHLHAFPSQRHKYETRNAIILCFGASPIAYTVCMSTIIANTKQGVFWLNVAYGLNDLLPWVLISVTQDSFSHVVSASTDLLAGIADDVQQLINKHARVESTGFRATLSAPASTKCPACRGRYPSQSSTLARGVRALRVRYQCVHDGVLATNGVYGLFNRASITVATVQVVVCLYVGVIVPHALNSIWTIIWGTFIAAKLMFMCIIGEQIQSENSRITAALQTGMSHRPNMNLQGKAEISRFCQQIQMQEIRFKAVDPYYFGLTTMKRSIASIMASVVVLVQFSTMLTQLYKYIV

>FoccGr69P

MERPGRLWGRALEAHWHTSLLGMRLAGMLGCSGWPPRASLPWTRYAHVFTVVSSIARAFMVYTTMVGFWRKLSTISHSPGASAAINAFVYALYLLDYFTRNVGLTLTQMVMVRHSRDLCLLVRAMLLYLKACPSQRSPYATRNAILFCFGAAPFAYIVAMAIMTANSPKVTSPIQVAYAINDLVPVVFVSVTQDSYNHVVSASTDLMAGIADDVEELISTQAPVESPGARVNLNAPASTTSPGCPVCPVHHPPLRTSTSTLAWGVRALRVRYQCVQDGVLATNRIYALFNTVSVTTATIQVVVCLYAGMVVTGXINSTWTMIWGSLIAAKLMFISITGEQIQIENSRITTALQTGMAQYPNVDLQAKTEISRFFHQIQMQEIRFQAVDPHFFDFSTMKRTIASILASIVVLVQFSTMLAEFYK

>FoccGr70

MQRRALDAHWRASLLGMRLGSMLGCSGWPPHSSRPWTRYAHVSTPLTSIARAAMVYMTFECIWSLLPGLSDRQSRVSSEVYRFTISTYLLSFLIKNVGLTLVQVVMVRHSHELSLLVRAVQLHLKAFPIPRRKYETRTAAILCFGAAPIVYSVAMLFINMSTKSLARWVGILYGLNDMLPVLFISLTQDTFIHVMSVSIDCLAGIADDVQELLNTPCPVVSIGDTLGAPAAKATCSVCRLAKTTASLARRMRALRVRYQGVHDGVNATNQIYQLFNVVCITTTMVQGVVCLYTAVIMPNGTFKRLWTLVWGTVFTAKLLFVCIIGEQMLMQSSRITAALHTRRVHHPNMDLHSKEEIFGFIQQVQMQEIRFKAVDPYYFDLSTMKRSIASCLALLVILVQFTTMLTRLDK

>FoccGr71F

MGSSGRPWGRALEAHWRASLLGMRLGVMHGCSGWPPRTSRPWTWYANVFTPVTNIARVAMVYVTFNDFWSNLSKTSQLQTIIPAKVYEFVSSMYLLDYLSKNVGLTLIQVVMVRHSHDLSLLVRAVLLHMNSLPSQRRKYKTRAATVLCFGAAPLVYTITAIIYDITYKTAADWLKITYALNDVLPVLLFSLTHDTFIHVMTVTKDFLEGIAEDVQELINTPSWVASPGDTLSAPAATTTCSVCLARRLEGRMSSLARSLRALRVRYQVVHDSVHATNHIYELFNVVSSTLTMVQAVVCVYSVVITSNATSRWLWVGMWNLVWAMSLAAKLLFICIIGEQMQRENCRITTALQTTMAYHPNMDLQIKREICGFIQQAQLQEIRFKAVDPYYFDLSTMKQIIASTMAFIVILIQFPAMLSLLYK

>FoccGr72

MKRSSVARSMRHGEAQVAGQPRNSTTSRQISEAHWRLALMLMRLVGMLGCSGWPLRCSRAWLCYGYVLLAVLAGARVAMWCHLGGKVVESIESLPGDSKFISTLFLSDFFIKNVLLMLVQMSILYRSRDVSLLVRAMFLSMKTFPDTRRKAETRKAAIVCFGMAPLMYCAVMIYIDVRSPQSLMSWLFQILLDLLPVVSLTITQDSFLAMASASIDFMAVIADDVQKHTDGLSRLALSAPEATLWSFRPREIMCLCPEKASGRASQPPAAVLELRALRVRYQCVHDTTHAANTLYGFFNLAFIALTIYEGAVCLYAGVVALFSPEEMGDMEAFFTLLWGTILTSKLIFICIIGDQMRVEDSRILATLQTEMARFPDTDLLSKREIKKFMHQIHMQDIRFGAFDLIYYDLGTLKRVIAAVMTFLVVIIQFSAISRQGSQKPNNTTYANSA

>FoccGr73

MWRVLEVHWRFALLWMRLNGMVGCTGWPPRWSKARMRRSCATMFLVTIVRAALCTTSVRFVWQSLNDAHDESNYRNSLFMADFFSKNVGMTLAQLAMLRHSRRLSLLVRALLLQSYSFPDSRRKCAARSVVILLFGAFSMVYTFVSVKFMICGGSWNGAVFFIADLIPVLALSISQDSFIHVVCASVDFMTSIADGVQEHMNALPHAYARTATPRNPAPAACGQQDDSPVQGLRTLRMRYQCVQDAVHLTNCLYGPYNVLSNTITMIESVVCLYAGMAGNIGRYAQVMRSLKEIYWGIALDSCLDFQEEIWNLTLGIALTTKILFICVIGEEMHAEVSLGGERTEPGWLFQVCSSSRFISRIIRFIHQTQMQEIKFVAVDPCYYDLSTFKKVIASILTYIVVLAQFSPLFPA

>FoccGr74

MTCVSTLFSVMAAPSASWRSLVDGHWRLTLLVMQLTGFTCCGGWPPHWTRSSARRAIAAASAFAVTRALALHYVVHVVGEDASRSATVGRQYVITLLLAGFVAKNLGLLLTMAAMLHESCALGLLVRAALLHAHAFPAHKHEVRAGVVIVLAVFPAIFTTYSIASNLRRDSTLENIVYAVNDEVPVLLFSIVQDSFNHVVSSSIYVLEGIADDAQERVAEPACIAQKQHAQGGDEDRGVRLLGRITWDSRDLTAHLRNRPRSWDHRDLRALRVRYQCVVDAVLATNRLYGPFNTLSSVILAITCAVCLYTGLSMLYTYQVMGTLLWGIVITTKFAFICIMGEQMSVENRRILAALQTGLANHPDMDFQRKGEISRFIHQIQMQDIRVKAFDSSSYYDLTTLKRAVASVATFIIVLIQFAALVAQAASDGSVPKCNCQKC

>FoccGr75

MLAAPAVALPRGCPGPSGWLLHWTRQWRALLALLSLIGMNCTSGWPPRPSRRVRCYGGLVLVVRTAHCVVVILSMKDSYDRNKDNYAMLLCLCEIALKCVVTAAAQFVMLIHCKNVCRLTECVLLYSRSGSPTSDCGAMVFVGPAYASGLGMVLGLAVYLIASNADESLTEVSYVLGELTSAVPLLVFWIMQLLLTSLSVGFARDARRHVGGWQFLHDMVLQHNSAFGWCFLCCLAHGFMEAVICLYLGVSYFRRSLDNVEVVKPRLLFNPWFFTWGILQVTMFSGMCVLGQLLSDIHGRIFCDIQTARVSNSRFCRATKMELQSFAQQIQRQDNRPGMFGLLYYDLTTMKTGFATLVTYIIVLCQFEP

>FoccGr76

MDAVPAAEHEPPAPPGWVRHWARQWRALLALLSVFGMNCTRGWPPRPWRGLRWYGSLVLAVRILHDVQALIEIKTTREDIGKINYIVLLCLYEMSVKLVGSVVSQVVMLVRPEDVCLVTECMLLYSRVGQPAVWGAVALVGPTYVVGLVLVAGPSTYAMATTGTWMQLFSYMLGDLVPSVVMMLVGIMQLLMTALYVGFATDESSNVKGWQFLHDMVLHHNSAFGWCYLSCLAHVFMETVLCLYLGLAQLRQVADPSEPLLNAWFCTWGAFELTMFCGMCILGQKLVDHHGQMYCLLRGFLVSNPQLCTEAKLELQGFAQQIRRQDNRPGIFGLIHYDLKTMKAGFASTVTYIIVLSQFEPAVGH

>FoccGr77

MDVLPAAEHKRPSPPSPPAPPGWARHWARQWRAMLALLSASGMNCTSGWPPRPSPVLRWYGCLVLAVRTFHNVQAILELKASHEDAGKIDYNMLLILYDMGVKLVTSVVSQVVLLVRCEDVCLSTECMLLYSRVGRPPVPGAMAVAIVGPAYMIGLVVMTAASTYVLSASLTHMLSHVLGDFAPTTAMIFVGIMQLLITALHVGLATDGSSHVRGWQFLHDMVLHHNSALGWCYLSCLAHVLMEAVVCLYLGASHLRLLAAEVGYVQPNPLLNPWLYVWGVFELTVFCGMCVLGQKLSDHHGRMYCVLRGFLVSNPQLCTKTKLELQGFAQQIRRQDNRPGIFGLIHYDLATMKAGFASTVTYIIVLSQFEPTVGH

>FoccGr78F

MPAAPAAAHALGPPPRPSGWVLHWARQWRALLALLSATGLNCTRGWPPRPSTGLRCYGALTLLLRTSQSTVVVVSMMKLSGDGISRYGLLLSIYEIITKCGATVVTQLVMLGRHDAVCRLTESMLLYGRVGSPSTGWGALALQLVGPAYGFSLLAMVSVSMHVLATTTSSFTKNDLFYALGLGEMAPSVTVLMFWIMQLSMTSFSVGAALDESRHIRGWLSKHFLHDLVLQQNSTFGWCFLCGLVHVLVESVVCMYLVVKYCRMRIHQVEIKSRLLMNPNFFLWGFLEIMMFSGMCVLGHLLSDIHDRIFCDIQGFIVSNPQLLCPTTDTELHGFAQQIQRQDNRPGMFGLLYFDLTTMKAGFTAVVTYIIVLCQFEPTEILNVG

>FoccGr79

MPCRTLTRALTVLVWALFSTSAALLGMLPVRGWLWRPDPRPWPAALAWGVFLSTSMCITMYGGSQDLRSTTGNSAFKSAMFAYHGVLRMLEGSERCRRRGELCALVQCLRLYCASHPLSPRGWRALLGIALFCTAYGVFNGFTSMWWIQGRTSALDFERVLWATFNVGLQLYTLTLSGFTGQLARDLRQDCLRALRAAGAWVPDHNGHVAAMPSDEARLVWPARRRSTAPAARKGVGGMPSGGQVLTMIVFTVTYCSICVSTTIGSLTEVQDGPVDYLLYRIMPLQAGLGRLLFFVFACYTFQQAAQEHVKLGQALQGFLADAPDVPFAVIREIEKFIVQIALLDGYFNVLGMFRLDISAMRTILAATATYLFVLTQFRLSLY

>FoccGr80IF

MVCQALTVLVWALFSVCSTLLGMLPVQGWLWRPDPRPWPGALAWAVFLFSSSYMNMNMNGGADDSPSTTGNSVFKCAVMLYLAVLRTVEGYERCRRRGELCALAQCLRLYCASHPLSPRGWRALLAIALFSTSYGLYNGITAVWWIEKGKARAIIPLARVVWSTYSGGLQLYVLTLSGFTGQLARDLRQDCLRALRAAGAWVPGHNEHVAAMPSDEARLAWSARRRSTAPAARKSVGGVLSGSRILATTTTYLFVLMQFRISLYY

>FoccGr81

MPCRALTVLVWALFSVSSTLLGMLPVRGWLWRPDPRPWPAALAWPVFLSTSLYMNMNGGGIDLAMRPINTVFKTALFSYHAFVRTVEGYERCRRRGELCALVQCLRLYCASHPLSPRGWRALLGIAMFCIWHTVFHALTDNFWADENAKPAFDAAVWAVYSGGLQLYALTLSGITGQLARDLRQDCLRALRAAGAWVPDHNEHVAAMRNDKARLAWSARRRSTAPAASKGVLVMIVYSLTYSATCVSTAVDSLVEMQDSLGDGYSVKDTLVPLQAGLGRIFLFLYACSTFQRAAQEHSKLSQELQGFLADTPDVPVAVLREVERFIVQIGLRDGYFDVLSLFRLDIGAMRTILAATATYLFVLMQFGISMY

>FoccGr82

MGSTATAWQRVAVLSEKGLGVLATWVGVLPASGWPTAPRPWAAWAAYGYGLAVTVLFELNTSYHEFQWKKGFSAGAVVFFVAARLIIVLSSITGRAALSDLQRCLLLYCRAHPPSVLGYRAIVALVMVAVVFLAVVFCLLDANNNAWNVEKVAKLTWTRRERLVMQMAVGVQWCSAYSLTLMPKAVIGSLASDLRQECSRLAAEAESPGRSCSHSSLMCKWRSARLRQQHLHDMSAATTAVFQWRLLALLATTILYGNLQMSDYFAKWIICSAKTRPNVYRQISQVLDALCHSVCCIIMFVVYRWESNEGEKICYTLMGHQAALAKNSKNTQSGDLREVCLFINQIRCQKNSSNMMWLFDLDYSNMMRIVSASLSYLVVMLQFQVLF

>FoccGr83

MNEKWVKKSVLCFWPVFYIQAFNFGIFPVHLESMRPLAAWRAGGSAVLCVSCGIANAALFSYTFSRSPDYILFMGINVISILLSSLLYWRTVHFRGNMCNTTRALLTHCERHPLSPSSWRLLLVASLLTTTFCIAHIAWVLDTFTDCQFNMWSRGFMQAASHALFFFLGWQHNLLFALVLIPVMLLALLFADLRQNCVELLQIVSKNRNEKDSGKQSNELSPECVSLLWRNLRCQYQSLFDIFSATRNLLQWHLVYLSFVSGLSSTVQLSDYIKFYNSTTPRSALQLAQLTCHSVRFVMLCLACQKFQSENDKMRQSILAYTSQIREFHRGNEVHMFGIQIEGQQNGICTVLWSFDMNAVTALQLMSAIVSYTIVMRQSDVHFKAF

>FoccGr84P

MNGTWVKKTVLCFWMIFYSQALHFGILPVQFEAMRPRPAWRACGSSLFWIICMVVNAALVTCMSARTPDTTNMFVGIFVFSILLFALISWQTVNARRPMCDTIRALITYCERHPLSPSSWRLLFVAAVLTSAFSIFHLAWVVDTYADYGLYLRHRCFWQASAAXRHFLIGWHYNLLFALVLVPVVLLALLSADLRQNCVKLLQTVNKNPNAKDSGMQLNEFSPECVRLHWRSMRCQYQSLFDIFSSTRNLLQWHLVYLSFISWQSSTFQFSDYINFYNSTMPRSPFQLAQLVCHAVRFVMLCLACQKFQSESNEMRQAMLAYTSQLRECDRGNEVHMFVLQIKSQQNGICTVLWSFDINSVTALQMMSAIVSYTILMRQSDIKFFDE

>FoccGr85

MDGSWVKKTVLCFWLVFYTQALRVGLLPVVLESMRPLPAWRAYGCLSFWVIGTIANATLFSLTFVRSPDYILLMGIYMTTLLLFALASWRTVYDRWKMCETTRALVRYCKRHSLSPSSWRLLLVAAVLTAAFGVVHLAWAIDTFTDSHFALWNKGFWETALYAVFFLLGWHYNLVFATVLVPVVLLASLVADLRQNCEELLQNVSKNPDAKDSGMQSNEFSPEYARLLWRSMRCQYQSLFDIFSATRNLLQWHLVYLSFSSCLSSALQLSDYIIFYNSTTPRSRFQLAQILCHAVRFVMLCLACQKFQSESNEMRQSMLAYTSQLREFNRGNEVHMFAVQIKCQQNGMCTVLWSFDINSGTALQMLSAIVSYTIVMRQSDIHFYET

>FoccGr86N

SPDYILLMGIYVITLLLFAFASWRTVYARWKMCEITRALVRYCERHSLSPSSWRLLLVAAVLTAAFGVFHLAWAIDTFTDSHFALWNKGFWGTALYAVFFLLGWHYNLVFATVLVPVVLLASLVADLRQNCVELLQNVSKNPDAKDSGMQSNEFSPEYVRLLWRSMRCQYQSLFDIFSATRNLLQWHLVYLSFSSCLSSALQLSDYIIFYNSTTPRSRFQLAQILCHAVRFVMLCLACQKFQSESNEMRQSMLAYTSQLREFNRGNEVHMFAVQIKCQQNGMCTVLWSFDINSGTALQMLSAIVSYTIVMRQSDIHFYESYED

>FoccGr87

MRARHFEALPIMFIRPTARFLLEASPTLASGRTAAMVRCVALPLLRLVGILAAVTSHLPVAGWPFGPGPPRPWSKCSAYATMVLVFLALELPTSALAGDSLHSEFIKARGAHVATWAIGMLRVLALAAGAGRRAALAELVQCVRAYCDRHPMTARGWRAVLATLMLVPLHFLIVGYFLALTVMQLYKFASLLMVLRIVYAYASAAIRTIVPFVVVLLPAVVIGSLAADLRHDCLLQVTVTRGRRHRAALPFRLSRMYEWWPSPAPRVPPSDLWPRPPSRLQSLTPATGAEWAAEKGLVVKPMDLWPALALPSATSGVPGRVLWRRVEPQLDEIVPLHGDTLEPLAWRTLRQRQQLLFDICLAAYSVYQWWLLFLVLNTLLYETLHLSNVVHALHSIKMGHIMQNFLVSSRFIGVQDREEITLFMEQINIQESHCSVLGLFVLDTVTMMAVISGISSYVVVMIQFQVNF

>FoccGr88

MHCSARRLTILLWKLLSYPAMLLGVLPVYGWPCHPRWESFENAFIFEFLIVSSSLLTRVCLVNSPFGCSIAQIIAVLSFILLEVPYHALDLVHRIVSFLTFESLVMMCLRGAERLWVVQKRAKMCSLARCIAIYCTKHPPSEEAWRSLLITTLTPLLLVVLKTVLVFSLVLDPIKQSSLLEPYSYGAVMIVYCMYTNLIYWVFVCTLAFVGQLAADLRKDGTVVLDSHLNYGELIPRLQDTEGHYLDMKHRTPSQMSREHISRTPSSAGSGGGSQTGLRSEDRSEEKEDRSEEKEDRSEEKEEEARTMTQTWCGVIACTDEDIQDAWLRIRIRQQILFDLHQVEAGTVFGYQMLVYVLVIIVQSALRCSHSLRAAITSANPSWLDMVKAGESAVNTLYFVYYCAVLQRVKSEFIRMCADMQARLTTSVRAGRAKHEVQRLLGQIRDQGKMGDVLGLFYVDGCLMKSVIGAIGTYVFVLNQFGITVE

>FoccGr89

MLFFFRFRLFCSRIPYLRVLVQWVLIPAVSRGVWRSLVILLRSCGMLPVSGPILRPRPLGCPVALIVGGVAFFFCSPMLYYSQYKLADAMCEWVHCMSLYCLRHPPSPRAWRRLLLVAMLFTVGWLGCVFSCLHWVLNSPVYAESGKSRTLFRVATGIYPVVLEIYLLLFLGPVALTTVLTADLGHELDLGPSQVIFHQDETPLDRVLHDLACTANSHVSAMILAFVVLHFYHSAFACSHTIISAINEPNLRWKLAVPVLHIICRNILFILLCWVAQRANSEHTRLSERVQTYLAKSSCTPTAGRREMEFLGFNVRTRRKRFNILGLFFLDLTFLKTVFGAILTYIVVLYQFGASFSSNKSE

>FoccGr90

MLWHVLAQLLRVCGMLPLRGPILLPRPLGYRMTCTVSVFAFIFSSPVGYYSVSGLGGRRRELCTWVHCMSLYCLRHPPSTRAWRGLLVVAVVFTVGWLGIVLANIFWMSFGVSVETSTVRRFFKATGICPVVLQIYILLFVGPVALTTVLTADIRRECERIISSGGARKILLNAEAKKVILHLYCSAFIISDSIIATTNHPGDTSRSLVPAVYSTSHNILFILLCYVAQRMNDEQTKLSERVRWYLSLSPSVPSAVRSEIQCLMFSVRVLRKRFAVLGIFSVDSTLLKTVLGAILTHVVVLHQFGAVFSQHFESEQITNSYVVN

>FoccGr91

MVVSKETPVFFVRPASSATSTIEQDKELKQRMGAFGVIDASPSMLRIATPLIVVSALLGLAPVQPASTSSAVADAKTDRIAGRARPHVLVRSVWLEVYSIVILGLITVSSYFVFKASGDESLKLNSLANASVSVLVQVATLWTTKSAIDSVEGLLDCEEETQTMTPARVRTARIWVAVQLLLWATVSVVFAVHHVFFEVPSQLTQLSFCYRAYSVSRPFMHMSCMQFANQAHALSLQLTSINDRLLTMLGPVSAGLTPRTLEMWSRLGDAANPLRQPPLHPDVEAAALETTRELRLLHHKMHVIAGYLQSAYGPQMLLVLLSTLLDYTTMLFILLARIEIDAVGLIIGTTHVLVMYLVLTSGNMVRQEAQRMGEILQRGLVHRRLSERLVEELQDFSLQLVRLPTRLSACGLMYLDYSMIQGAVGTVTTYLVIMLQFPNNTGGAAKANGTESTPTYTTEGGRT

>FoccGr92N

DSCKVRIDGTVMMVQAVLVGLFFLKQAVAALIFVCAIVKGGEVARVVEKLVRISGRAVLSLGPGLHDGASRCAPRWGAARYGVAAQVVLLLTYWAVVLYLAYTDNPVSFMDMSSLPPIVLVNVQFTDVANRCLVDVGSTSIGLDWDYGRRCLQALLLRAELAMLNDSLAAALEQRPGSPAQDISWVGASSKDRVRGVVQTARALVLKLCEYIPCVRKETHSLLEDSQSLPFLHSELRHSCGLLDDAFGLQTLMFVTMTFVQSTVVLYMLLIEGRLIDLWLAGSALHLLQLYAQFAACQQVKDEALRTAGVVHRALLRPDLSDAAQQELVDFSVQLATSAEMTFSAWGLFTLGHGTMMSFLGTVATYIVIMVQFRGPRKPLTTGLFPGANSTSTTTTPLPPPRPRPAEVSRSLPGTDMEGLNY

>FoccGr93

MLRPVLRAAQAAAILPPPPGRLRSLQVGVVHMEKEGEGYKHCHPSECSRLPSGAGAHDVFTSRGPCKFLGWSSTFGGSYDICTPSLGYAGWYRLEGSQAVAALAVACWANYMAWRVYLHHTLDVLESGFHLALHAYLVENLLETLQPPIILLATWCNLHRFAPSLGRVLAGPVRAPRSFLAFTVLMYAAHAAGQALFHVTILQHRVDYICYSVASAWSACSALIMDRLVALVLTLAALKIQAVNQRLGSLKTDPSASPGAPSTSASWMPQQDASLDPAVQRGLLQSLANRWESTVEQVGGLLGHCMLGLPLLYSLLHITIGLITAFSLIVRDETQPIVRHVKLIHVSIYLARVASLALTAGWLYRETKMTEAVIGSLDTTAWPIRERQRVSHLLHLSRETEPFRVCGILSIDHSFLFTVMGNTVTYMMMLVQYPILTPGKAAAVLNQSESVASEQSK

>FoccGr94

MAASDSRQLHLLQGVPSYPSPWRLQLVVVGQLLGVFPLRRLGGVGARTPIWQLSRLLQLFSVVGVFTSSVCFYNFFAVVKLPGQGLKGTLQRWSLIFYAVSGLPCALGALCYGRRLERFSNSLLLLARELRPANQQVWPRGWALRSTALPAAIYFIDMFREFQHVKGLSFELYTVCTVLAYTLPMATETQYMFMSSILALSYQRLGEHIYALQGPSRSPSEHWLDEVRRSHEQLVSLTCELHCLLGPQIFVTVSACLLSLLTGVYFAASAILEGLVDADGSEGVSMANQSDIVYFTARLIAICATADTVKAAAAAVVNGSIRVDYGTWHTNSKERWNMFLSQLETVTPSFSVAGIAKIGMGLLSGMIATTLTYLLVLLQFDTARITAARASANSR

>FoccGr95

MILYFQNKATPAPSVVSRNSLQDAVEPVYSVMRLFAAWPAKRPKGNSAVFPLFLLSSRIPNLTFARAGGRGSVLTRALLLSVQVACLLLVGYEWVGVLQDRVRYMRRGAGVTRTVLAIQFLLYLAMFFTLPLMWRDAARLPPVWEGWSHVQARLERVCGGRLAASRALSWRLAVLLLLLWPPIVAVPLMLLPEAEMPVRQAWFYAYPTLVTFSLKLLWMRLFTALRHAAELLVSSFRTESRDWSRPDAAARLRTYRELWSSLSGIMQDIGVRGGLSLLCVQAIQVLSLLVCFVHIVISCLEMREIGSLLETTSTVGLTTAVDYEMLVLRDTLAHKHAAVRLCGFVTVDRELVGEVLVVMMTYLMVLLQFALL

>FoccGr96N

VNFREVTGHRLSFGVAKKARVTAALCPVVGLAFVLWSHDAGYVPWNQFPVYEMNATYLAILPETWVLIALALETAAQRLRAELVQVGTTTYRDGAHAGTLTLYVPPGAIQVLGDVGEAADSGPRRRALADLRWLWMQLQHLVRMWSLTWTPCLAHLILSSVTVLLLSVFGISLPLLRGQVKDAFVKEMTRAKFATASTNLYPELVKFLKAVDIQDPCLDMGGFTSLNRALAVGCASIIVTYVVVLMQFAYAKGAN

>FoccGr97

MMTPEHEGGWQRAPWVDTDELPQFGAVVACMGLAVAEVRGRADEDQDFDGVVIDYLVYMQCALPLAVLPLYASHRSAMAQYLRDWGALQADFAREFGRSLSSCRGFGPRPSWEMVTIVGVFSLLVAGAQALLNPAIVVLALVHSTLVSALLTSSFYLHGRLLRAASRALADHLALRRSQSFLIYFLHAPLDSALDGVPQELRRPVPRVAAVDNIRLLVLRVVALGGGLSSAVAMPLGLLLLAHFILATLTLYAVLGQVSYTYAARLPSLGTGTALSGLLLYLACDTGHRVARAVKQLAVEELLAFHKRTTVIGADLRKMVETFISTVQEVPSDVSLSGVTTLNRGLFTAMLSTMVTYLVVIVQFRISLEQAGSDKGARPGTANSSTTLPGNDSAIEPTTTTTTTLTWSGEGSASPSPPTTVVDNTTLSV

>FoccGr98

MIPQQVLEIVGDYSYRGSHFGNILHINVMAVVLIGHLGMTMGTFRKAKYLPQVLRDCNNLPGPQHDCWATTLYSAEVFFLVAIGGTLRYLCTRVVCCFPHTFYTSLKSRYWCLQVLIVYFSSTLSAISERHLKQSTGDLQLFKKAGNYSLSDNHFVTSWTLVSSREISSTTKHRYVAKNDTSLLCSELCTLMGAHGDSRDLLDRLTLTQEMQFLAILILEFVLIVVYLYVVSFTVFFKLTSMTHMIGFGFVFINGLLSTQLGVIYACRSTTKEGNKTLNIVHDMMNCGKFTNSETHQLTLFSMQIGHQEVAVVPHGMFKMDISLLKSLFTTVIMYVIILVQFGSTI

>FoccGr99

MEYARTIAIGRRAPSRPYPRLYSMCLSIWLARRLDSSRPGVHTFFICGRARGLHGLHPGGNVTYGALVNLALLAFILGFLASELFFAYMVASAMEGLDVDDLNLYLNLLAAEANGTARPTLAVESEFAQSQLTRVMKVSSGVALTVATLGLAVGYVRKLKHLPAVFRMGDELPGKPQWKTALFVAELSVYTIPLLTFLVTEFATTVLTLYIEARNVYLDTVQFMDLLAILFLLLITQVTTLCYGTCRSTSCKNLEPLQGNATAAIVHKIMNTRRFNEEEIQQLMIFSMQISHREISVSPYDMVTIDYSLLQAMLASVAMHIVILVQFNLSSTT

>FoccGr100

MRLLVLFTRTIGLCPVYRESVSSRVPLTPTSPPGDDEVPRTNGLPQYKPSLAWTAYSCLLFLALMVIDGFLCFEGLRALRDPDIATGLINVMSAVAYAASFLSAVAMVAGAVRKYRLLPRLLNWPVCPGAIAHPTPKTLLYFFGPYAVAYVYTDYGFVMREPSTAILLLVVITGLLERFNLMFLAGRARLCFAAVNEELRQLGSLFRDRMRLHYVPDAAAKIAQLRVQHAGIATYAELLNTAYELQIAGILLYDFGTTVLTSYQFIVFLSRGDQPTDERQEATVLFFNTMYSAVSTVSLCQACYWMHRLGMQTHALVHKALSNIAISEEESLQLTIFSMQLMHQKFNMSPLGIFILDNSLLASLSLGVVMYVLVLFQFGSPATGPVPTTPTPLNSTLDSTLDPTF

>FoccGr101

MGASGRWGVDPSRAKAGNKVWQQPQGQRGRDPLHRLWATFSPQPARPWRQGAPGHAPGPPRRRFHAIPFLIWPNYCLARLLGLQPLRFVPTSDLSGDPQWLSLSPLWMPACVAVSLATAYSAYFLLKHFMDTLIDPSRTLNNYIYEVPMVLNAFIALLCTGYSLLGARAMAVSWRGALATLAELDAVNHAPSSTSRRCRPLESFTFFLMVWVMLALVIGTNWLYTRFQDFVVEVWHMGAVLLVRLVPMAVEGQFLLSTLLLERAYRAIVVCLRDLARGYFPSDQVRVVESMSTAPSKKYVCFPPTLPQDQVLLQLACLVDPTAWDSASNQQLPVKLKYHEESVRALLKRIFRPTGPSNEPCSTLLASAKRVLVCGLHRFTRTGSSATVAATTTTYLVVLVQFQGPNAED

>FoccGr102

MQGATCRVLPPPLPQARGAAAALNQLLQCAARALTGRGRARWAAAQVAGTVLCLVAYLVGVAVFLSLTEPDEDRRQSDGSAPEEAAALLGLAHLPSLPLQLFFCLIFGNQVGLLWLLVRTLNRRLDAMTRGRAPPGASSHNALVPAVWPKGPFTLYPAASRAATRWANPSRTTQPGDPTPLRVGALCPAPGVAPSLHGEVREVHALHAEVRHALTLLTSAHGLELLLMVLVTFSAATTSLYIWRTDRDECSGMALAWSGLVALFLFGVLTAGELVKREVGQGYDMNGIGSGLSRCTMELTQRLLIYHEMSPELRWEMRRFSHQLLHQGLKVSAYGFLSLDYTMLQGMIGSVTTYLIMVIQLSPEDPAAAPAEDRGRGNATGSP

**183FoccIrs**

>FoccIr8aF

MTSRWHCMSQMCLSHNVVSTPLPLLSRLYLFTKVCTALTSAALVLDMTWSGWERLRWLAHAAGVPYVQADVSVRPFLQAADDVLRFRNISDAALVFDGDLEMSQGIYYLVGRSPLRVVVLPGLDGDAWEPSGPAGRLGAMRPTPSMHVLLSGEVDAMAERAAQAGLVGRYTRWMLVDTAVSGATRDARQLQNTTIPVTLLTARASLCCSLRGGGSRGMGPGDSCTCAQPQRLHELLIPAVVDLLQQLAVEAHRQGVPLQVERGVDAATICRASRDTPTQNASSPAIDMLRQLAGQLPGLLVQRLDHRSDVALLPDLVLDVAVVPMGNGATDGGDAEGDVVGEWSTTRGFTARAELPALRRYFRIGIVESTPWTYRAEDGSWTGYCVDFAHQLSEMMDFDFEFVPGSDYGERDVRTGRWSGALGDLVSGRTDVLIADLTMTAQREEAIDFVPPYFEQSGISIVIRKPVRKTSLFKFMTVLRLEVWMSIVGALVATAVMVWLLDRFSPYSARNTKYPYPCREFTLKESFWFALTSFTPQGGGEAPKALSGRTLVASYWLFVVLMLATFTANLAAFLTVERMQTPVQSLEQLARQSRINYTVVGGSDTHAYFTNMKQAEDTLYRVWKEITLNASSDETQYRVWDYPIKEQYGHVLLAIERAGTLPNATVGFQKVLESEDAEFAFIHNAAHVRYEVSRNCNLTEVGEVFAEQPYAIAIQQGSYLQEELSRKILELQKDRYFEALTAKYWNYSLLGDCNSGGDNEGITLESLGGVFIATLCGLVFALITLGAEVVYYKRKENAVTDLQPARANGDNNGTTAKDSMRRALAEVNQVAADRSVLLTKSSIASLAKQRRAAQGLGQQGLATITVRPPPPNAPVPGLPYASVFPRGNLY

>FoccIr25aF

MAKLELLPLWLSIVLSLPILSAAQTTRGINVLFVNDESNSVAEKALDVALNYVRRTASTGIQFDSFLRVSSNTTDAKTLLEALCDKYDNSLKTEKGPDLVLDFTQTGVSSETVKSFTHALALPTVSASYGQDDDLRQWHFLSDDQLKYLVQVSPPADMIPEVIRSIVLSWNISNAGILFDDSFVMDHKYKSLLQNVPTRHLIAAVEEPRSLKRQLNRLRDLDIVNFFVLGRLNTLKNVLDAANANKFFGRKFAWFAITQEKGQLKCSATNATILFLKPEPDPTARERLEKLRTDFSLTVEPEIASAFYFDLSLRAFMAVKQMMDKNQWPRLEYTSCEEYEEENPVVRKGYDLRRALREVQEPPSYASINVEDRNGHSFMEASMRLEKVTVMNGQSVSAEVEGSWKLGLDSPLMIKDVASISNLTAASVYRIVTVIQKPFVMKKVNADGREEYEGYCIDLLKEIAQVIMFEYEIYVAPDNKFGNMDEKGQWDGLIKELMEKRADIGLGSLAVMAERENVIDFTVPYYDLVGISILMKNPDAPPSLFKFLTVLETDVWLCILAAYFFTSFLMWVFDRWSPYSYQNNREKYKDDEEKREFNLKECLWFCMTSLTPQGGGEAPKNLSGRLVAATWWLFGFIIIASYTANLAAFLTVSRLDTPVESLEDLSKQYKIQYAPHNGSSAMTYFQRMADIETRFYEIWKDMSLNDSLSEVERAKLAVWDYPVGDKYTKMWQAMLDAKMPEDLDAAVKRVRSSVSSSDGFAYLGDATDIRWLSQTSCDLQMVGEEFSRKPYAIAVQQGSPLKDQFNNAILQLLNKRKLEKLKERWWSLNPEIKKCDKQDDQQDGISIQNIGGVFIVIFVGIGLACVTLAFEYYWYKYKRNPRVTDLAGGPLPNAPRVGKGGGLGGQQQGRLAVIGRPDEQAAKLDAATLGQVLGQGYNYRRNGNQLPGVNQAWQH

>FoccIr76bF

MRAAGLVNILLTGVCLTPDYSKDRLNNTLKRGECAVRCSRSFFAFFEYSSEQVGFHSFFPRCRFAEVELLKDGPPPQDVCLIPGNEVKNILVGKKLTIATWEDWPTSGTKKNERGERVGDGFAFEFVDILRDKFRFSYDIITPEQNVLGDAHTGILSLLYQKRADMAAALLPVTPTTLKLAQPSISLGETEFVILMERPRASASGSGLLAPFTETVWYLILVSLILVGPVIFVIIKIRNRLLRRRRPLLMPEDPAGMELEDYSLPACVWFVYGALMKQGSTLSPRTDSTRMLFATWWLFILVLTAYYTANLTAFLTLSKFTLPIDGPKDLRYGYRWFAERGRFLASAMLNESALEELKPLPQMFEPLPVSNYSREQKIMELVKGKRMIFLDERRKVEYLMFRQYVRGTEEKIKEGERCVLAMAPKTFSVVTLAFFYPYNSSLVQLFDKTLLSLLEAGIIKYRQRLMLPLNQICPLDLGSKERTLRNEDLYTTYAVIGSGFVAAIVAFFSEFIYLRHQRKKALERAASARRKVSAEEEGRGRHARPPAPVDQSARPQVRPAPRNW

>FoccIr21aJ

MSHPVGEADDDHGRGSRPAGRKLIPPCARAETIWAATVEKQRCFPRPGSPGPVPALVARLAEHELEGCDPALIWDASIPAGEDTLREIIAALTRPYVLARTPVNTAPPWLYQPVSPPYARRRRGSWRHVLPPRQKCRHILAIVDDVMAVGSLLGNLSEPRVVVVTRSPLWRIRDYLGSSIARNIRNLLVVHDRSWRHGSPATGYPDMNLLTHRLFMDSLGYSKAQVLTSWRCDRLTRPDTNLFPIKMLEGFAGHQFGVSAAEMPPYLYRTVLNSDGDIHWDGIEIRLLQLLAERLNFTIAIRDASNDDLSSRGAAVSVVNNVVLGTSELGAGGIYVTHARSQKTDFGFYHTQDCAAFMALASVALPRYRAIMGPFQLSVWIALTLTYMFAILPLVYSHSFTLRVLIRSPGQIENMFWYVFGTFTNAFSFKGRDSWTRSSRTSTRMIIGWYWVFSIIVTACYTGCIISFIAFPAFPTVVDYIWQLWEGNYQIGTLRTDGWLTWFRNSSDPMVDRLLEDMDTVPDVWAGIVNITRAYFWKPYAFLGSRLYLEHVVRTNYTAENKRSLMHISSECVAPFWVALIFPKAATYSERFSQVLMRAMQAGITEKLRRDVAWDASRGSALKLKSSDERMLTLDDTQGMFLLLGAGFAFAALALFVENMTGRVACCRKYCQPGPRRRVRTSSSSRKG

>FoccIr40a

MARYEGMTSIIFCSRKSSEKLINTIRNANLIQRNILYMFYYELWPPSDVFLGALQEAMRVAVISNPRPGTYRIYYTQATPEGSGTLKLVNWWSSSATLFRYPVLPPASRIYTDLRGRTLHVPVLHKPPWNFVTYTNKTFLVEGGRDHELLLLLADKLHFRFDYFDPPERSQGSAFVETTDHNKTFPGIIGLISERRADLALGDITITFERSQAVEFSFLTLADSGAFVTKTPSRLNEALALVRPFQMEVWPVLVLTMLLTGPLLFGVLEAPRLWRGKAWMAKRGRKRISDRKLFGNAVWFSIALFFKQSVRGPEDSHRVRLLTILLSFAATYVIGDMYSANLTSLLARPGRERPISNLEQLAEAMETRGYQLLVEKHSSSYNNGTGVYERLWRLMLRQREMAEVASGEDGMVRVGREPNLAMLGGRETLYFDTRRFGAHRFHLSEKLYTRYSAIAMQIGCPYSESVNNILMQLFEAGILTKMTEEEYEKLGDKLHDQGPMLPERTDGAGDEDGQGDEEGGDGEDGAATTLTPADYGFYQEDDYYDYPQPQDYGAPNCHEGCPQPSAPSETSTATPPTTTTPTTTTPIAPEGPAEGSEREVKKGGRSTSSTDETLRPISIKMLQGAFYALLAGYVLAALAFWSELATKRLERCSCHCYAATRACAERPWVRCWVRSSACCRCCDSVRGHAAATKQRFRQAVAAALVMLRASSDEDDDEEVDNASVADRPVTHEYLKAYLRNEQLLRRPSNTTSSSSSASVSQQTSFATFTTHTRTSRFKSSSRSSPPSSSPGSSISCIRPFIN

>FoccIr68aNIJF

SGRAALLARATETTSDAELAALLRVLVQRVDEDLNCLALIGDALHQHVFGRLFFKKLGLVPVFKVTVQDGEDLLSPNYRTLKAIRQVRKEGCRAYFLLFANGIQLGRLLRFGDSQRVLDTRGRFVVLHDRRLFSPDLHYLWKKIVNVVFVRRPPLVVVRAVHGTVPRAHPGRDGDEAPRHLAQGRLPAERRSFQNQTLAVVTFEHVPASYKLRHVANEHGDVTSGFSGLEVQVMQALARAMNFVPRLYEAPGASREQWGRRQLDGSLSGLLGEVTSGRAELVLGDLHYSPFHLRLMDLSVPYHSECLTFLTPEATTDNSWQTLILPFKPEMWLAVGLALAAGGLTFFVLARCGPAGDEDASRSTPPAATVRAAAAAGGAEETTEERRQRLRDDDGPPIKSAAHLFSSSLASSMLYTSGMLLGVSLPRIPTAWAVRVLTGWWWLYSILVVVAYRASLTAILANPAPKITIDSLEQLADAPLEVGGWGEHQKDFFLTSLDEAGQKVGRKFEASAAGALEARDLHIMRDCVINMPISLGLQRNSPLKPRVDRFLRRVIEAGLVHKWLTDAMLATSASAGRGRIGVAEESVKALMELRKLYGALVALGAGYLVSALALLAEVSFYRGVVMRRQGFDVYALGRNANRNPAAPAAVKRRAAA

>FoccIr93aJ

MLLHSALVLVVVVLVNLGSVANAAEPPSYDLNNATLAVVVDPHFHRQDGANLLDQLRAFLGVGTLELLQHGGLNVEYYGIADYALRSDLTAVVSVATCADTWAIYRRMEQETLLQLAITDADCPRLPSDVGLTIPLTESGGVLPQMILDMKMARALTWRSAIVLHDNSIDPYLLDRLLAALAVSTPDTLPSSVTAYSLPVEHSDWKRRKRVEGVLRSLPLSRLTRLGERYLVLVGSDLVGVIMDVAKSLGMVHPLSQWLYVLSDATFGNLVTLTSLLHEGDNVAFVANYTSSAPQETCDAGVQCHARELLRAFLVALDRSVRDEQELATQVSDEEWEAIRPAREDRMALLLSHVRDLLRTTGSCGNCTAWRLRSADTWGLSHSEPRSRPHITASLVDVGWWRPREGLTLEDHIFPHAAHGFKGRTLPLVSFHNPPWQVIVSNSSGYVVAYRGVVFEIINELSKNLNFSYTVYFPNPNKPGLTGDTEFYTSMADQAVELNNEFSRNISWDKILLNVRDNKVFMGAGALSVTAERQRLVNFTRLISRQPYCMLVVRPRELSRAMLFTSPFTTATWLCIFLSVAAIGPVLNWIHRASPFYEFRGDVRRRGGLNSVTNCAWYAYGALLQQGGAHLPDADSGRLVVGTWWLVVLVLVTTYCGNLVAALTFPRLEKTIGSTEDMIAQRDQVSWGFFRGSPLIDHLKTASEPQFSLVLERGQVHEDEADALSRVRTGTHVLIDWQVNLLQLMKKDFNECGRCDFSIVKVQDFVEENVAMVMAQDNPYLPIVNQEIRRMQQVGLIEKWLQDNLPKKDRCWSSSVVEANNHSVNLGDMQGSFFVLFFGFLLAAGLILLECGWRGWHSAKEKKVIKPFVT

>FoccIr75aNJ

DAALVRLLSAAGLAAQLLRGREHTRARHVRAALGHTANAIIVDAACPGAQRLLVLATRLRLFHIANTWIIFEDAALGSNPAATTASDDVDLRRDDEFDDLDNVTVYRYGAQNREDAVDPKEEAFLEVRPPMPPRARSGLLAPQSDRLSATAVQILDGKGMFVDSHVLWIQTTGQIKYPEEYTDIEDNNLRKNDLYPKMNWPVTQNLAYSLNFGMLIYKRDNYGWDDGHGVFDGMIGEMQRHEHDMLATAVFIRPDRLPHIDLVAETFDMRTAVLFRQPSKSSVSNVYVLPFSPGVWRATGAVSLLALLLLVVQTQWWRSHRSSEQSDDPADHIEVGDCVTFIVGALSQQGFYCTPIMVSARITIITTFIANLFLFTSFSGSIVAIIQSPSTAIRSVQDLMQSEMKVGVQELPFQHVWFQEAVAEHDVASDVFNSKINTEDSPGFLTTKEGIKKVRFERFAFQVETNEGYREILDTFTEPEKCDLSELELFPAAPMVFCTSEKSPYREALSRALRWHREVGLLDRIHNRWESQRPECSAQIKVFVPIGLQEFYPALLVLYNGMLLSLTVLGVEFVYNSRAVTSCLLRAWRRENKEKEPLVADQ

>FoccIr75bJ

MATLARVLQSCALAVALAVALGAPQALKAATDSEVSTYGAAVEVAAKEAQRRGVHRAAALVCWGSAAERRLVLALSAVGVAVSLLRGRDQTTAAAVDSLLREGSGRTAIIVDMACPAAPRLLLLASKLRLFDLANSWFLFENGRLDPYASEDSTVNRTLSWDPGVDDFPDVELVTAGRYGAQRREEALLPDEEVLLEVLPPSRRASDRPWISSQVSLVSPAALRALDGLVVMSDSDVLWVQARGARGSGTAANDSSTDSNDACPSQLQGFINFPELYTFIDDLRERKRDYRPKTNFPVVRVMAEQANAHMLVFQRWDYGYNRDANGMFDGLMGELQRFECDLLSTAVYVTDERLYVLDSVAETFPLRTGVIYRMSGSSHSAADLYLRPFSRGVWLATFAVSMVCVVQLVASTRSWMTRITLGWDECLTSVIGALCQQGADRTPETVSGRTVMVFVLLLSFFLFTSFSAFIVALLQAPARPLSSPRELLASPLKIGGMDVAFIHAWFRIPRGVVGDIFVKKIKSHERPAFLPVWEGIELVRGRGYGYQAELNEAYHEIQEKFTQSEKCDLFEMELFPSEAQIIVTVEKSPYREAFSRVRWQRELGHLDRIAKRWYASRPRCESGARGLTPIGVREVVPLLLLLTYGIALSLLCLAVECLTARHQGKQPTRSLCLRLFGGRPRHLQQREQRPRRAMGQQWTADQVDWPWRPGPASRKAAHVGDEDREPGPALLLESHTLYWQTRGRALGHFARRAQELQPHSRH

>FoccIr75cJF

MRGGAGGSALTAALAAILGVSMVWGQDYEASSSATHTLVGDFFSAREVPFVLLFPCSHRGSVTWLALGSTEPELDLAVDSDFTLASELAGGEVELRSLFKHGHHPGSPWDHLLLHQPLLFQLHQIAVGDDQAALRRALVDHTNTHMDTETRFGFALATHLAEMYNFTLRLLVTPSWGYPRGNGTFDGMIGMLQRGEIDLGVTPAVITKERLAVADFTVQTWTLRTCFIFRHPRLLVPYKALSEPFAPATWAAVGLAAVAACAVLGAVRRCETAPAVGAEGGALSEAAVVVVGVLSQQGSALDSRLVSGRAVLLSGLLLSVMLQSFYTSAIVTSLLTPIPRTIRTVHDLMRSPLKVGLEDYSYNRAVFPSSADPLTAQLYRGKILPAWPEGGFFSVSDGVRRVRRGGFAFYSEDSSLYTPIQNTFSDDEKCSLSEVELNPAFVTAATLAKSSPYKEIFNRGLLLMLERGLRARQRRQWSARRPSCEREAGAVAAVGLDPIFPAHALLVLGALAALALLYAERRAAAQGVARAPRDGHKCGAKA

>FoccIr75d

MAQKRMSAPRVAPPRGGSPSRRRLLVAVAVVASALAHGQGQVQPVQHSTATVARLAADFFASRGVVQVVMGVCEPDGPGLKSSIPCGGQAVASCPLTAPGRTSLTRALAADGRLQVQAVQAQTTAPHLATPALQRPPVLGVLVWSCRPSLGERLVGAPGAVSALRQRRRASGACGFLVYFDAHPGPRPRHRGGRVALGDRLQAHARAGSHRQGGHLRKTSRHVWVRLPRMYFRFRLSAPASGAQMWQQVGEVVMARLLEMPRAEAMPLLTPLAMHAHIVFMNIVARFNMTVIKRIARTKPGQLNADPGTPFDGTMGLVERGEADIWPGCIITADRGRPMVFSATSWTLRTVCGFLTNTALGSADAVVKPFQPWLWVAVIGLSVVVALLEKCFRALEHTDPEHGGASWSDVLISCFGTLTGQGFLACGDWVSGRALLATFEAFFLLTQAYYNASIVTFLLRPPPSSIRSVRDLVESPYKVACHGWIHTARVLQNSTDPLVRELFHRKVEPPRGLGFLGTADAMAALRRPRVGLVASDDVVDEAAASMTRQETCRLQQLDVDMPLVIALGTARNAPFKEMMHSGTILQVERGIAHRERIVWRQRRPSCDRNAGGEFAPLELLPVLPPLVLMGAMVSVAAVLCAAERVRAASVTSVPMAKAARARRPLQLLARGGGRLDGRQPTHHLS

>FoccIr75e

MVPGPRRAAAVAVLVLAAAGHLAGEDVAALAADLFAARGVRAVLVALRAGGPATFGVYRALSSDGRFQVSQTVGSTTVAHVLDTSERPYVGVVLEAGEPPPWFDPGDRAWFNGSYVWLVVDGDAAARTASWALRLDSEVFLAAQDDGDRAWRVDEVFRVNAEDAAVRRPFASWSPAGGLVAADPELAVAPVSSPGGDPLADRRTQFRRVLAGPRQLRVESSVYVRHKYQYTAMLCWRVRHHEHLISGELAAPPSGHFSGACGNVQRGVDDVVVTGLLMSDGRLQVFKYSATTYHFRSACLFLADGSLGSVDALLTPFTLLAWLAVGGVALAGMVLHKCFRHVEGDRIDWSESLLSTVGMFTGQGFASSGSWVSCQGLLIVSEVLFMLLDGYYNSTIVKSLLRPPPPSVRNVHDLLDSNYPVALNDWWLTVRNFQNASPGLQRDLYEKKIRPPVGLGYFPPSGLYKVLLRPLHAIIGSEESVYKSTEPLTASQACRLHELDVNPPQHIAPLVGRHCAFTELLFAGVLLQRERGVWQRVRFKWRQRRARCSGGGAHGEGDPIGLEPVLPLIALMVGGLLATAVVAVLERVAHRRSQRTRGAASSARSPGSVEDAVAGTTASLARFAS

>FoccIr75fN

GTTPCLSRRRRGAARGSRSPTFCSLPRREVTLASLASDGRTVHLVDAFRPTVRSRGPVHMQLTGTWTEAAGYAAFLPGTTSSRRRNLYGTVLRAGVLVSGYREADLPHVLGKPKDNAPLVSDVAAFSHTLVSHLQDMLNVSLRLQGGHYFTLLRRIGRGDLDLLATATEMQAKYWDVMDYACMPLWTERRVMVLKNPSALWSYNAWLRPLDAKVWYILVTLVLCTAVVLRAVGYWEVHFDQGHVAEEDSWASAVTTALSMVCAQGFTTTTSWTSSRTLLLVFEFFSLFVNLFYSSGIVTFLLTFPPPFAHDVRDMIDSPFVIGADQDVFSRANFTKSSDALMRELYDRKINETVGGGGFVDLREASARARHGRFAVVGSPFPMYEMLATTWSQSDICRFQELPLLPPALLGLATPKRSPYRELINQALLLLVERGQLARERLRWSVLRPRCALSDVTELDYVGLRQVTPLLSFVVGGVVLSLLILTLELLTANSKYKDR

>FoccIr75gJ

MQTHRARATGALAALLAAAVVGVLADENATASVSKAHLKAFLREAHITRVPGPALLLLCGEGDPRAMARELGEAALPAQVVTAPDQWLLAVRHLDHIAHSVVLDAACPEARTLLATDGGYKVSIFLGGVAGPTSLLSCAVPSRTALAVSAYRERKGAVLRVQRAAVWRSGVLEQAPPPDRRHLPARRLDFHEAVTGVGFVVADSKNFKRAINEGTGLTTDYLSIFGWDLMHEMVAMLNMTINVTITNDWGRVPTGVVPDKGLLGLHAHHQIELAGTGFSHIIEPVRLKYLHFTNFYSPAGYGVGYTLHFAAWSHGVLVFRAPPLSYENNVFTLPFPLSLWETSLGMVGLCTLLLVVVMHAERINMHQGQGDHALHSSSDALLLAIAAVCQQGSPVETRGVPGRVIALMLFLLVLLLYTCYSASIVVLMQRTSTTIRHYRDLTSSQMSVGAVDIPYIWHYMVQEDDPVRKRVYLEKMMGGGKKKHMYTLSEGVRKVRNEYFGFFGIENHLYAEISSTWQEHEKCGLVDIGHDFLRVKNPHFAITRGSHNVDAYKIMMRRLAERGIWHRMLRRFVRQKPACVSSAGVFQSVSLTDVRGAFYMFIATMAASVLVFLLELAAHRRQLRVKRKAAAAQRKASPANALDRGGRLLQQARAFIVYERVTNDSELSKQVQPGA

>FoccIr75hNJ

VSRLRPGFSSCPYSHSLEETVTWTTIFPARVSLVCLSISLHVTNSHGYIRNGKYDGLLGDLFNGLVETGGTVMLPVAFRMMAITYLGINIPMRAVTVFRQPPLAYTDNVFWLPFPDSLWAAALSLVALCCVLAVLALYLEPNSESGDGAGGGTGDSHASLRVARRPQGAGRGGYKQPLRASLSDVFLLAFGAIAQQGSPVEARGVASRIVTFLLFLAVLLLYTCYSASIVVLLQSSSSSIRTLRDVWKSRLTVGVENMPYNVYFITQMGTTGDDVRRGMYHDKIARPGRPRNFISMTEGVRRMQEGLFAFVAMDSPMYGHIQARFEEHEKCDLFEVDGYSPLRHPHLVTRRDSPVTHLLKIKLLLQWERGFRHRFIIRGLYSKPRCDVMGARFMSVSFIGARNAYYLIVAGMGASVLILVVELLHSRLYAGGAGHRQPTGQLSVGEGDGGGEGRGVGMSLFAKKEVEA

>FoccIr101

MVRAQLLILLVHATLVPDTISRGSTSWTTRRPAAASPTRGMAPRSTPNPALLPNLKVPPLVGLAVPVDVWEKVASILFPNAGSIQIYWVRLTARAHSVREREVTEVFNTLWVTVAASMGKEMGFPYVLWTVSEGVVPVGVRQYVDRHREDRLGEFSLVIAAYTEERDDKGDLSLRQLQRLDVWFPRRVSLRSGLNNALESMSFDSIKTLRSRSYYIVLLDTWSLPLARRCDESLALNVSLAVWLSNMTTVGVVLVRCPWKIVLHDPFVLDPKRRRTVIHQSNVTALEDVVGDRRFRNFHGWAWPISMLEYAPTLMRAEDGSYYGIDGSVFMAATKYFNFTPVIDQPADKEPLGYSDDGKTGTGTLGKVMSGESWISFNSRFRKDYNTKDIDFTYPYSFDELCVMVPQAERIPNYITIFRIFRIEVWLTALLAYLCTALAAHLLDRAHAARRARTFRRPGPCGLAYSAFLLIMAGSMDTYSGVREISSRMLFGAALLFSLNFMGAYQGLLFKALTMPQYFPQIDSMLALADSGIAITTRSTSLVDTFKADNAIMTRLQTNFKVPALVPKNWTTNIPVSSLARRNGAYLRKENANLHLVEECPRKYQLAYIVHRNFPYIHQMNIFLLRCMDVGLMDKWFRDTYLGRPPSSAEDDMVLSLFEVEGAFFLLAIGVLASSVAFSAELLHSRCQRGRAPRPRRAPRKRHAVARGPIRRVATRAPRPSETRDKGALEGVVVHFIM

>FoccIr102

MAVLVHLGLLIGVLCWTGVHAALHPVDTPAAPEAECVASFISSIMPPDNSCLVVHGDARLVGPLLRNLQDLDRGRLTFVMDPRYIPTHTRVRIESTTLAVIAAENPANLSTLAFRDTGLFFSYSGLVVWVRARSLEDALHDPVGLSSWFWICNKNVHLLLTTPNGTTIQYVPDVGGSCVVTWSDLKMREVRRCEPGRRQGWQGQPSRPRLCSRWKSPETGKTTPKFLSVRPPHYRQFDVPAAYDEPYHLAVSTLTTAVSRWLRTPIELSWESNNSVVFAQITNCSLNAAFLNYPVKVRNVSHIEFDSFFMSPTIVVVPGGSVMRLNVLRAVTAEFSAELWIATALALLFMTAAMVVAWTTLGRPPLSALAAATLQTLAPLLAQPPPGRTAHRPLSAVWLLMSVVLAAAYQGLLLRELTAPPGEINSLEQLEQSGLDIYAEETIQLQDSLESRLSQSLNTKIQYYNPMADYMTALQRVVEGRNSALICRQHVMTTFILSRLSNRIHVFTLPNSTNLLASVISTKGSPFMKPLHSALSWFSASGLLKYYLTMADFHLTLNSASKDTASKLNRPLSLGQLQPAFCLLAAGYVLSAVVFVLEVLCYKWSRRHAPPVPVFLH

>FoccIr103

MALLNQLDLLIGVLCWAGASAAMHPVDTPAAPGAECVASFVSSVLPLDNSCLVMHGDSQLVGPLLQNLQDLDRGRQTIVTNTSGISSATFTQMQFTVNLHVFAALSAANLLTALRNQKGEILSYSELVLWTRARSVADALQDPAVSGMSLWICSNTVHLTVTTPNGTTIQYVPDFGGNCVVTWRDLKMREVRRCKPGRRRRWQGQPLRQFCSRWKSPAPGKQTTPQFLSVRTPHFRQFHGAALYDEPFNRAANTVTNAVSRWLRTPIELSGEHNATVLVDQITSCSLNAAILNLPVNVRIIADVEYDSLFIAPIIVVVPVGCVERLNVLSAVTSEFSAELWVATALALLFMTAAMAVAWTTLGRPPLAALAVALLQTLAPLLAQAPPGRTAHRPLSAVWLLMSVVLAAAYQGLLLRELTAPPGEINSLEQLEQSGLDIYAEEALHLRDNFMSRLSDSMKSRMQFFPVTSYDKALQKVVEGRNSTLICRQSPVTALILSHLSNRIHVFTLPNSTSLMANAISTKGSPFRKALRSAIGWFAASGLIQYYLAVADFHLRLHGASKDTASELTHPLSLGQLQPAFCLLVAGYVFSAVVFVLEVLCHRWSRRNAPPVQE

>FoccIr104N

SPPPEINSLEQLDQSGLDVYAEEGLYPLDQFQLNLRHSLTSRIQYYNPATHHITLLGRVADGQKSALICRPNMITGLTLVHLPKSVHAFALPNSSSLMATVIYTKGSPIGKAVHSVLSYIFASASMQHFYNVADFPAASHTASADAANELTHPLSLGQLQPAFCLLVAGYVLSAVVFVLEVLCHKWSRRHAPPVPVFLP

>FoccIr105C

MAVLVQLGLLGVLWAGAGAALHPVDNPAAPEAECLASYIYSVLPPDNSCLVVHGDARLIGPLLQNLQDLDRGRLTLVMDPRDIPTHTRVRIESTTLAVIAAENPANLSTLAFRDVGLFSYSGLVVWVRARSLEDAMLRDQADQRQLLWMCSKGVFLFISTPNGTTVQFAPDLYGSCVVRQADLKLRETDRCEPGHRGWFGRRWPRLCSSWKSPEPGKTTPQLLAIMPSYSFGQQLSQSTDYQAPYYHFARSLATAVSRRLPIELTWIVPNSTIVLEYTANCSLAAALLSYGVNVKPMRDTEQDAIFMSPIVVVVPAGAGPRLNVLQAVTAEFSAELWIATALALLFMTAAMAVAWTTLGRPPLTALAVALLQTLAPLLAQSPPGRTAHRPLSAVWLLMSVVLAAAYQGLLX

>FoccIr106N

PGEINSLEQLEQSGFEVYADDALHPLDSFQPNLSHSLTSRIQYFNPATHHITVLERVVDGQKSALICRPSMITTLTLAHLPKSVHVFSLPNSSNLMATVIYTKGSPFGQAVHSALRRISAGALRQHYIRMADFHLTLRDASKDTASELTRPLRLGQLQPAFCLLVAGYVLSAVVFVLEVLCHKWSRQHAPPVPVFLH

>FoccIr107N

YFNPVTEYIPALQKVVEGRKSALICRENAVTALMLSHLSDRIHVFTLPNSTSLMATVISTKGSPFRKALHSVFSWFSASGLLKYYLAVADLHRTLHGASKDSASELTRPLPLSLGQLQPAFCLLVAGYVLSTAVLVLEVLCHKWSRRHSPPVGQP

>FoccIr108C

MALLVHLGLLIGVLCCAGARAALHPVDNSAAPEAECVASFVKSILPRDNSCLVVHGDARLVGPLLRELQDRDSGRQTILLDPRDIPVHTRVHMQLASLALIAAESPANRTVRWLASRERELSLYSGLVVWVRARSLGDAVLRDQADHRRPVWVCSKGVFLFVTAPNGTTVQFAPDRHGGCVVRWPELKVREIDRCEPGHRGWQGGRWPRLCTKWKSPETGKGKTTTQLLALMPSSSFDQQFDKVTVYHAPYYHFAMSLTSAVSRRMPVELTWVAPDSQKVYDHYANCSLTAAFVSYPASAQPKPGIVHDATFTSSIVVVVPAGAGLRLDPLRAVTTEFSAELWVATALALLFMTAAMAMAWTTLGRPPLSALAVALLQTLAPLLAQSPPGR

>FoccIr109NC

SILPLDNSCLVVHGDAQLAEPLLQSLWDSDRGRQTIVMNASDTSSHTFAQMQFAVNLHVFAAVSAANLLTALMHQKGEILSYSELVLWTRARSVADALQDPAGPGMPLWICSRNVHLVVTTPNGTTIQFVPAIRGNCVVKWSDLKLREVRRCDPGRRRWQGQPLRRLCSRWKAPKMTPQFLSVSTPHFRQIDGAADYNEPFNRAASTMTTAVSRWLRTPIGLTWKPNIMVIVDQISNCSLNAAFINYPVKVRFVSNLEFDSLFMAPTIVVVPIGSAVRLNVLSAVTSEFSAELWVATALALLFMTAAMAMAWTTLGRPPLAALAVALLQTVAPLLAQSPPGRTAHRPLSAVWLLMSVVLAAA

>FoccIr110N

AHRPLSAVWLLMSVVLAAAYQGLLLRELTSPPPEIDSLEQLEQSGLDVYADEALYPLDKLQPNVSHSLTSRIQYFNPATHHITVLERVLNGQNSALICRPNMVTSFTLAHLPKNVHKFVLPNSSSLMATIIYTKESPFGQALHLVLSWFQSSALLRRYLNVAEFHAALHTASADAASELTRPLSLGQLQPAFCLLAAGCVLSAVAFVLEVLCHRWSHRHAPPVPVFLH

>FoccIr111C

MALLVQLGALIGVLCWAGARAALHPVDIPAAPEAECVASYIYSVLPPDNSCLVVHGDARLVGPLLRKLQDRDRGRQTIVMDPRDIPGGGLVQLQFAVTLHVTAATSSANLSTLAFKYDDQLSHSGLLMWTRARSVADALSSHADPSRPLWICSKNVHLAVTTPNGTTIQYVPDIGGKCVVTWSDLKMREARRCEPGRRRWQGQPLRRLCSRWKSPATGKTTPQFLSVRTPHFRQIDGAAALEEPLYRAWSTVTTAVSRWLRTPIELTWKPNIMVLIDHITNCSLNAALLNYPVKVRIASHIEFDSLLMAPIIVVVPVGSAVRLNVLQAVTAEFSAELWVATALALLFMTAAAAVAWTTLGRPPLAALAAASLQTLAPLLAQSPPGRTAHRPLSAVWLLMSVVLAAAYQGLLLRELTAPP

>FoccIr112F

MALLVQLGALIGVLCWAGAHAALHPVDTPAAPEAECVASFVSSILPPDNSCLVVHGDARLVGPLLRKLQDRDRGRQTIVMDPRDIPGGGLVQLQFAVTLHVTAATSSANLSTLAFKYDDQLSHSGLLMWTRARSVADALSSHADPSRPLWICSKNVHLAVTTPNGTTIQYVPDIGRKCVVTWSDLKMREARRCEPGRRRWQGQLLRRLCSRWKSPATGKTTPQFLSVRTPHLRQIDGAAALEEPLHRAWSTVTTAVSRWLRTPIELTWKPNIMVLIDQITNCSLNAALLNYPVKVRVASHIEFDSLLMAPIIIVVPVGSVVRLNVLQAVTAEFSMELWVATALALLFMTAAMAVAWTTLGRPPLAALAAASLQTLAPLLAQSPPGRTAHRPLSAVWLLMSVVLAAAYQGLLLRELTAPPGEINSLEQLEQSGLDIYAEERLYLTDSFESMLTDNLKTRIQYYNPMTDYITALERVVKSRKSALICRQDAITAVVLSHLSNRIHVFKLPNSTSLMATVISTKGSPFAKPLHSASSWLAASGLLNYYLTMAEFHLTLRDASNDPASELTHPLSLGQLQPAFFLLVAGYVLSGVVFVLEVLCHKWSRRHAPPVPVFLH

>FoccIr113

MALLVQLVLLLLIGVLCWAGARAALHPVVTSPAPGAECVASFVASILPPDNSWIIVHGDAQLVGRLLQELQDLGRGAQTLVMDPRDMPSHMRFLMRFTVSLYVIAAVSPVHLSTLTDGITDTASLASGMVLWTRTSSLDSALLRDQVDRGRLLWVCERRRHLIVSAPNGTTVYFSPDIGESCVMRWSDLKVTEMGRCEPGHRGWQGPRWPRLCSRWKSPEPGKTASQFLCIKPNIRSSHHQIADLNPYYDTFYKYASILTTAVSRRMPVELSFVERGGSELMNSTVNCSLAAALLRSPVSVGPSPYLEYDSFFMAPIIVVVPAGAGPRLDPLRAVTAEFSAELWVATALALLFMTAAAAVAWTTLGRPPLAALAAASLQTLAPLLAQSPPGRTAHRPLSAVWLLMSVVLAAAYQGLLLRELTSPPPEINSLEQLEQSGLDIYAEEGLHLIDSFRYMLSHSLKSRMHYFNTNTNYTTVLERVVDGRNSALICCHSTVNALTLSHLSNRVHLFTLPNSTSLMASVMYTRGSPFAKSLHSVISWFVASALQMHIITLADFRLTLDAASKDTASELAHPLSLGQLQPAFCLLAAGCVLSAIVFVVEVLCHKWSLRHHAPPVPVFLH

>FoccIr114

MPLLAQLVLLSVLWSGARAALTLADISAAPEVKCMASFVSSILPPFNSCIIAHGDGQLVGPLLQELAKDRQMLVLDPRAIPGHTLVQIHSTKTLYLFAAVNATNLSVLTPNGKDRLCSYSAIVFWTKAQSLADALRDPADPTMTLWMCSTNLYLAISTPNHTTIMYGPNIGENCFVKWSDLKMREIGRCKLGRRGWQGQQWQRICSTWKSPEMADTTARFLSFIPYHSDPHRVGASTVLFEPYNQFSSSLTTVVSRQTQTSIKLSWERNVSILNGFVTNCSLAAAVLSYALDVRSSAEIQLDALFMVPVTVVVPAGAGLRLDPLRAVTAEFSAELWIATALALLFMTAAIAVAWTTVGRPPVAALAAASLQTLAPLLAQSPPGRTAHRPLSAVWLLMSVVIAAAYQGLLLRELTSAPPEINTLEQLEQSGLDVYAEEGLHFLETFMTRLNHSLKMRIQYFNPSHYYITALRKVVDGGSSALICRQNMIFALALSHVSSRVHVFTLPNSTSLMASVIYSKGSPFGKSIHSTLGRFDASALLNHHRSVADFHITLRAAKEDTTSGLIQPLSLGQVQPAFVLLAVGFVFSAVVFVLEVLFHKWAHRHAPPVPVFLH

>FoccIr115

MALLVQVALVLVLGVLWAGVHAALHPADPSAAPEAECVASYVSSVLGTNKSCLVVHGDPQLVGPLLQELGDGRQTIVKDPRDIPDHLRCELRDTPKLAVVGMASAVQLSEFLAQIGGMSPMSGIVLWTRAPSLLDALHHLKAARGSLWVCSMVFNLVVSAPDGTSLLGQLTANECVPDFRSLKVRAVDRCTSGPGWQHRQWQKLCSEWQPPKEGPRLRVYAFRPKGMTGKLGPVTQYESHLMEALSRRQPFNVSWFNEIDNNISSAFERCGVTLCFMGRAVAVVDSPHVRFAGLRIMPTSAVVPAGAGLRLVPLRAVTAEFSVELWVATVLSLLFMTAAVTVAWTILGRPPLTALAVALLQALAPLLAQSPPGRTAHRPLSAVWLLMSVVLVAAYQGLLLRELTSPPPEINSMAELLQSGIDIYADSTLQMLHSIRYKDKRIPKDNYLPSRDIPEALQNVADRRNSAVLCFADAYCXXXXXXXXXXXXXXXXXXFGHMNEVDSKHL

>FoccIr116

MALLVQLVQLVLLIGVLCWARTQAALHPVDTPAAPEAECVASYLSSVLPPDNSCLVVHGDPRLVGPLLRHLQDLDGGGQTLLTDPRDLSDRLSCELYDTPKLAVVGMESPVRLSEFVAHIGAMSPMSGIVLWTRAPSLLDALQHLRAAPGTVWLCALNFVLVVSAPDGTTFLGHLTADECVSYFRTLQVRALDRCTSGPGWQHRQWLKLCSEWQQPKEGPRLGVYAFKPKGMAGHLGPVIQYESHLTEALSRRQPFNVSWFDEYEFDKDIVSLLERCGVALCFMGRAVPVIDTPNVRFAGLGIMPTIAIVPAGAGLRLSPLRAVTAEFSAELWVATALALLFMTAALAVAWTTLGRPPLSALAVALLQALAPLLAQSPPGRTAHRPLSAVWLLMSVVLAAAYQGLLLRELTSPPPEISSLDQLLQSGMDVYADSTLQILHYVRSKDERLPKENYLPKREIPEAVQNVAVRRNSAVLCFADAYCKYVASQAAQARSGPWRRLHSFTVPGSHYLTGEVVYSSKSPLAGPVRRVIWAARSGGLILHRDDAVRSIRRLRAYAGNDTADDTRTQPLCMSHMMPAYCLLAVGLTLGVLVFALEVLSDAWSRRHAPPVAVFLH

>FoccIr117

MALLVQLVLVLGVLWDGVHAALHPADPSAAPEAECVASYVSSVLATDKSCLVAYGDAQLVGPLLRHLQDLDRGGQTLLTDPRDLPGRLSCELHDIPKLAVVGMASAVQLSEFLAQIGGLSPMSGIVLWTRAPSLLDALQHLRAASGSLWVCTMVFNLVVSAPDGTTFLGNLTADECVPHFRSLEVRAVDRCTSGPGWQHRQWQKLCSEWQPPKESQHLRVYAFKPKGMTGKIGPVTQFESHLMEALSRRLPFNVSWFDEIDHHIPSALGRCGVALCFLGRAVAVVDRHNVRFAGLRIMPTIAVVPAGAGLRLAPLRAVRAEFSVELWVATALALLFMTAALAVAWTTLGRPPLSALAVASLQTLAPLLAQSPPGRTAHRPLSAVWLLMSVVLAAAYQGLLLXXXXXXXXXXXXXXXXXAPHLKSDLWVSTTYPQNGFALL

>FoccIr118C

MALLVLFVLVLGVLWAGVHAALHPTDPSAAPEAECVASYVSSVLPPDNSCLIVHGDAQLVGPLLQYLGEDRQTIVKDPRDLPDRLRCELRDTPKLAVIGMTSAIRLAEVLAHTGSAAMGPMSEIVLWTRAPSLLDALHYLKKARGTVYLCMMKLDLVVSAPNGTSFLGQLTANECVPHFQSLKVQAIDRC

>FoccIr119C

MALLVQLVLVLGVLWAGVQAALLPADPSAAPEAECVASYVSSVLATDKSCLIVHGDAQLVGPLLQELGEGRQTIVTDPRDLPDRLRCELHDTPKLAVIGMVSANRLSEVLAHTGAMGPMSGIMLWTRAPSLLDAMHHLKAARGTIWLCMMRFNLVVSAPDGTSLLGHLTADNCVPHFRSLEVRAVDRCTSGPGWQHRQWHKLCSEWQPPKEGPRLRVYAYKPKGTVADSAIDVRIVGHITAALSRRQLLNVSWFEEFNEKIPRVLERCGVALCFMGRAVPVFDTPHVRFASFRIRPTIAVVPAGAGLRLSPLRAVTAEFSVELWVATALALLFMTAAAAVAWTTLGRPPLAALAAASLQTLAPLLAQSPPGRTAHRPLSAVWLLMSVVLAAAYQGLLLRELTSPPPEINSLEQLEQSGLDIYADSTLRVLHHIRSKSRRVHRENYLRKREIPDALENVAYRGNSAVLCFADSYCNYVASQASSGPWRRLHSFTV

>FoccIr120

MALLVQLVLVLGVLWARVHAALHPTDPSAAPEAECVASYLSSVLPPDKSCLVVHGDAQLVGPLLQELGEGRQTIVKDPRDLPDRLRCELHDTPKLAVIGMVSANWLSEVLAHTGAMAPMSGIVLWTRAPSLLDALHHLKKARGTVYLCMIKLDLVVSAPNGTSFLGQLTANECVPHFQSLKVQAIDRCTSGPGWQHRQWHKLCSEWQPPKEGPRLRVYSFKPKETAADSVIDGRIVGHITAALNRRHPFNVSLYEKFDEKIPRVLESCGVALCFTRIAVPVIDVPHIRFAGLRIRPTIAIVPAGAGLRLLPLRAVTAEFSAELWVATALALLFMTAALAVAWTTLGRPPLPALAAASLQTLAPLLAQSQPGRTAHLPLSAVWLLTSVVLVAAYQGLLLRELTSPPPEINSLEQLEQSGLDIYADSSSQVLHHIRSTISWLKKENYLRKREIPDALQNVADRRNSTVLCFADAYCNYVASEASSGPWRRLHSFTVPGSHYLTGEVVYSSKSPLGGPLKGVIAAARCGGLNRYREDAIYQSMRRRRRGAAANVTVDLGRTQPLSMGHMMPAFCLLLVGLTLGVLVFALEVLSDAWSRRHAPPVAVFLQVLSFSLMHEVNHLV

>FoccIr121

MAVLVQVVLVLGVLWPGVHAALHPAYPSAAPEAECVASYVSSVLATDKSCLVVHGDAQLVGPLLRELGDGRQTILKDPRDLPDRLRCELHDTPKLAVIGIANAVQLSKFVAHAAPMAPMSGIVLWTRAPSLLDALHHLKAAQGTLWLCTRRFDLVVSAPDGTSLLAQLTADTCVPHFRSLKAQAVDRCTLGPGWQHRQWHKLCSGWQPPKEGPRLMVYAYKPQGTVADSAIDGRIVGHITAALSRRHPFNVSWFEEFTEHIPRALERCGVALCFMGRAVPVIDTSHVRYAGLMIRPTIAVVPVRAGLRLGPLRAVTAEFSAELWVATALALLFMTVAAAVAWTTLGRPPLSALAVASLQTLAPLLAQSPPGRTAHRPLSAVWLLMSVVLVAAYQGLLLRELTSPPPEIDSLEQLEQSGLDIYGDSTLRVLDLIRSIGRLLPKQNYLRKRDIPEALQNVADRRNSAVLCFADTYCNYVASQARSGTLKRLHTFTVPGSHYLTGRALYSSKSPLEGPLKKVIAAAKCGGLHRQREDAVHRIIRLRRIRAAANDGRTRPLCMSHMMPAFCLLAVGLAVGVLVFAFEVLSHAWSRRHAPPVAVFLH

>FoccIr122

MALLVHLGILIVVLCWAGARAALHPANTSAAPGAECVASLVFSILPPDDSCLIVHGDAQLVGPLLQELGGARQTLVTDPRNLYDRLRCELYATPKFAVVGMASAVQLSEFVAHIGGMAPMSGLVLWTRAPSLLAVLQHLRAARGMVWLCAMNFDLVVSAPDGTSLLGRLTGDECVAHLPSLGVQAIDRCTSGPGWQQRQWHKLCSEWQPPKKGPRLGVYAFKPKGTDSDSAPESQIVRQVTVALSRRQPFNVSWYESFRHIPDALGRCSVAVCFMDRAIPVIDYPHIRFAGFKITPVIAVVPAGAGLRLVPLRAVTAEFSAELWIATALALLFMTAAVAVAWTTLGRPPLAALAVAILQTVAPLLGQSPPGRTAHRPLSAVWLLMSVVLAAAYQGLLLRELTSPPPEISSLEEVFQSGMDVYADSILHVLDHIRHHKGKRNPQPVFLEQRDVPGALQIVADRRNSAVLCFANGYCNYIVSHEASGGRLHAFTVPRSHYITGEVVYSSKSPLTGPLKRVIGAVRGGGLTWHHEDVVTRRLRRTSSANHTAHAGRRTRPLSVGHMMPAYCLLSVGLTLGLLVFALEVLSDAWSRRHAPPVAVFLH

>FoccIr123N

FSFRSFTCRPYTAVEFTDYYVTRTQVLVPIGLGPRASLLQAVTDEFSAGLWWATVGAVLAVAVATALAAVASLGRAPLVAAALAPLEALAPLLGQPPPGGRPAYRPLSAVWLLMSVVLAAAYQGLLLGELTAPPGEIDSLEQLEQSGLEVRVSDDLYTEVSALLSPELQSRVSLVAGPDLPAAVWSVADMRNSALVIQNDITSLIAMGPLLRADAKGRRQRLHVFEIGPPLPLAYVAFTAGSPLRGPVTLTMQRQRDHGLVHYLIRTLLLPKGGHNGGGADQPMDVDQPAEPLRLSQLRPAFALLATCHVVAALVFLLEIACHKYSSRKKT

>FoccIr124

MKLVVVALAFLAYLGSPEAALAPPEHAVVDTQGMAELVSAFLLHIEGTLFVYGRSRALDALLEQMQPDVPRSIVPPATDFYGAIRRVLELQRGDSVILVADDTHGLALANSLIADVNLPLLSRLLVWTWAESLPVALPPNITSTPRLFEIETTLAVSTTDGSTFLFGVTLEQGVPLPPTVTVTEVDVWSPASRCWRRRVLPFRRTCSKWRRDHRGSQRTPLSLYATKLESHMTTQFQAAYLDFVTRMARSMRQPVRVRWVKGFADWWSEMVNCSVGMVFSFQSFTTRPFCAAEFADYYMTRIRVVVPIGLGPRAALLQAVTDEFSAGLWWATVGAVLAVLVATALATMATTGLPLLVAAAIAPLQALAPLLGQPPPGRPAYRPLSTVWLLMSVVLAAAYQGLLLRELTALPTDIDSLEQLEQSGLEVRVSAGLYAEATELLSPELQSRVSYVTEPDLAPALCSVADMRNSALIIRNDMASMLALGPLLRADAKGQQRLHIFQVGPPLPLAHVAFTDGSPLRRPMDLNLYRHRDHGHVLHLIHSMILPIAAGSQSEDDQHSGDADQPAKPLCLFHLSPAFLLLVTFHVAAGLVLLLEIIFHQYSS

>FoccIr125C

MKLVVVVVILAFLGYHEAALPPPQHAAVDDTQGMAKLVSTFLLHIKGSLFVYGRSRALDALLAQLHPEVPRSIVPPPATDFYGAIRRVLELQRRDNVVLVADDAARGLALANSLIADVGLPNLTRLLVWTWAEGRQAATVPPNITHTLRLFGIETTLAVSTADGATVLYRITLRQRAPHVMVVTEVDAWSPSSRRWRRGALPFTRTCSKWRPNHRGSKQQPLNLYASQLGSHMTDQFQATYLDFVTRMTKSLPHPMRVRWVEGYKDWWGELSNCSLSVLFS

>FoccIr126N

LVVTWRLAAARSTHGHRHEAPRHPGPPRLVSTFLLAINGSLFVQGRAPALGAFLEQLHREVPRSIGPPVTDDYGGIRRVLQLQETNNVILVVDRLCQAYSRDLPAFSRVLLWTWADSVQDVLPLNIASVWFAGTETALAVSTPNGITTLFYISRDYDRPSRPLTVTETDTWSPSSRRWQRRASPFQRTCFTWDRNHKVGTHAPLSVVAQLPDPYVTNKKAYHEFVEGLVYPVRRLANLRWMDNHTVFWRSILDCTLSGAVGFLSGIILPPGAVQYTDFRVMTTQVVVPAGLGPHPSLLQAVTDEFSANLWCATAGALLAVAVAAALAAMATLSRPPLVALAAAPLQALAPLLAQAPPGRVAHRPLSAVWLLMSVVLAAAYQRLLLRELTALPGQISSLEQLEQSGLTVRVSDEMHVHVPFLLSDKLQSRVSYFTPPEMTSAVRSVADWRNSAVVLLLDMNSMLALSPYAKAVPQRLHMFKVGATLTSAHMMFSVGSPLREPLGLSAEQVRAHGLDLHLMRSLFSRRMKDGSPDEDEQITKPLNLVQLRPVFLLLAYFNCLATLVFILEVLSQKWFNKRAQLRC

>FoccIr127

MKLLVILAHLACLGSHHEAALPPLKPVVDTRGMPDLVSSLLLPIHGSLFVYGRSRALDAFLEQLNPEVPRTMAPLVTDLYGGIRRVLHLQQTDSIILVVDERGLDRAYSLIDHDIPEASRVLLWTWADSVQDVLPLNITSAWLTGIDAALAVSTPNGTTVLFNVSVDSNRLLQLVIVTEIDTWSPLTRRWQRRGPPFKRMCQKWKHSHKAGKRAPLGVNALIPDHHVVYKQAFHEFVTGLVNSIRPKSNLRWSNVTELKSRCKDCTLSAALLFQSHIMRPPGPVHYADFRMANVLVMVPAGLDPHATLLQAVTDEFSAELWCATAGAVLAVAVATALAAMAILRRPPPVALADAPLQALAPLLAQAPSGRTAHRPLSAVWLLMSVVLAAAYQGLLLRELTAPPGEINNLEHLEQSGLTVRVSNKLCTLVPALLSNKLQSRMSYFDPSEISSALRSVADSRNSAVVLLADASTSLALAPYEKPAPKRLHRFKIGGTLPAAHMMYSAGSPLRESLGLAAERVRSHGLTMHLVRTMALHQARDTRPGEDEQITKPLSLLQLQPAFLLLAYCQCLATLVFVLELLCQKWCNKRAQSSCGNKYLPYENRGRPESKGVN

>FoccIr128

MKLLAILAHLACLGSHEAALLPPVPVVDTQGMPDLVSTFLLSIKGSLVVSGRARALDAFLEQLNPEVPRSIVSPVTDDYSSIRLLLKLQETDSVILIADDDGFDVAEGLNTCELPAVSRLLLWTWADSPQDVVPLKNPTLNETCITDMEVALAVSTRNGTTFLFYVLGDATRPEYPIVAAEIDVWSPSTRSWERQARPFKRMCSNWQQGNKRSKFTPLEMLASIPNALVDQKMYRDALTHFANSLRPPANVKWVTDFNQAWDKVLNCTLSGFVFFYPLPIRPASVVRCGELLKSSIQVVVPDGLGPSPALLQAVTGEFSAELWCATAGAVLAVAVATALADRAILSRPPLAALTDAPLQALAPLLTQAPPGKTAHRPLSAVWLLMTVVLAAAYQGLLLKELTTAATREINSLQQLEDSGLEVRASDDTYLYVPTLLSTKLQSRVSFFSRAELASAVRSVANLRQSAVIFQNDFRSMIALPWFKHVHMFEIGTSLASGICFTNGSPLQNSLRRTTDRFSEHGLHDQLRRIQSRHVTNGHPSDDDQSAKPLSLSQLKPAFVLLAYCYVVSTLVFVLEIMRHIWFDKRNM

>FoccIr129

MARLVFLALALCPGAYDAVLLPDPRPSPDSPGVAVAELVSHFLRGTTGGVLVYGRGRGLDAFLVQLPPEVPRSLSFKQSGAFNSRLLHRGVGNTHNIVLIAGDDRARLLNAINRTAPAYYRVLVWIRAARLEHVLGLVNPLWVASNQVALAVSLTNGTTLLLNSTLRFVKESSTFSVDLTPIDQWVPTVRRWQRRASPFPKLCSAWRSHHQAGGDRANFKLLAFLPDPNLEQPGDYVDLVNTLTRSLGQTVDVEWMHEESDGARDELRRRTFGCTLDALVTYRSSPTRSNWPEIRNVDPTLIKMQVVVPAGLQRVPHLQALTVEFSAELWCATVVAVVMVTMATVLAARLVLGLPLATAWTAAPLQALAPLLAQAPPGRTAHRPLSAVWLLMSVVLTAAYQGLLLRELTAPPGEINSLEQLEQSGLDIRMSAEWYEYGQYFLSAKLRPRLTHFPPLDLHTRVQEMLQNRNTALVIQNDMSSKLILSPYLTSTPKRLHVFGVGHQLLVTQVWCTTSSPFIKPLRKAGLRAREHGLLAHMVTSTSVRQGANVRPGHEERLTRPLSLRQLQPAFWLLAYGSGMSTLALAGEIAYKKFVSRDTFNI

>FoccIr130

MAWLVLLALALLCPGAYDAVLLPDPRPALDSQGVAELVSHFLTGTTGGVLVYGRGRGLDAFLGQLPPEVPRSLAFRQSDAFNFGLLYRSSGNTHNIVLIAGDDRARLVDTINRTAPAYNRVLLWTQAARLEHVLGLLEPLWVATNQVSLAVSLTNGTTFLCSSSLRFVKESSMFSVDLTPIDQWVPTVRRWQRRDSPFLKFCSAWRSHHQAGGDRANFKLLSVLPDPILEHPRDYIDLVNTLTHSLGQTIDVEWMREENDEALDELRRRTFSCTLDALVTYRFSPTRSNWPELRNVDPILISMQVVVPAGLQRVTHLQALTVEFSAELWWATVVAVVAVTMATVLAARLALGLPLATAWTAAPLQALAPLLAQAPPGRTAHRPLSTVWLLMSVVLAAAYQGLLLRELTAPPGEINSLEQLEQSGLDVRMSAEWYEYGQYFLSDKLRRRITHFSPVGLHTTVQEMAQYRNTALVMQNDMSSRLVLSPYLTSTPKRLHVFVVAHQLLVTQVWCTTGSPLEKPLRKAGLRAREHGLFDHMITSTSIKQGDNGHREGGPGQDERLTRPLNLRQLQPAFWLLAYGFGMSTLALAGEIAYNKFVSRDTSFNI

>FoccIr131

MRGFVLVALVLCPWACDAIPLVPAPPTDSQGMADLVASFLMGTTAAVRVYGRARGLGAFLKQLPAELPRTAASKWNGAFDSRLDTEVGVNRLLVIAEDSRAQLIATINAAAPSPSRVLLWTWAARPEDGLVLESSCLWILDRQVAQAVSVLNGPTFLFTVAAKVLNESLALSLNMTQIDRWSPRARRWQHRAAPFLKFCSTWSPHQSGTNLAIRQIYAVLPRQYSAYPESYIDFVTSLMNSLNQPVHLHWRYDVHGYKDVHARLLNCTLDAAITYDTSVMRFAPEIRYASQYLTTMQVIVPAGLHRVNLLQAVTAEFSAELWWATVVAVSVVTMATVLVTRVFVGQPWATALTVAPLLTLAPLLGQPSPIGTTHRPLAAVWLLMSVVIVAAYQGLLLRELTAPPGEIDSLEQLEDSGLDIRMSMNLYIVRQQIPSDLLRSRITYVLPEDVVSTVRTLAEKRNFALVLQRDTNMELILSPYLTSIPKKLHAFNIGFPFMVVQVWLTAGSPLEAPLQKALRLFQQHGLLLHMFRSTCTKLGINIGVDLDEQLVQPLSLRQLRPAFLLLACGLGISSLVFVSEIVYNKFRK

>FoccIr132

MQLLAVLALLVCPGAYDAVLPGPSPSPDSQGMADLVASVLSRGKAKQGGVFVLGRAKGLNAFLGQLPAETPRSLVSSTNVRSGTRCELQGAENFVLIAEDSWAELADTVADVEEIEFPTPRVLLWTWAARAGLKQGAQGVLAHGVVGDLWLLKRHTALAVSTLGAGGTTLFSLATNTKRQVVVTEIDRWSSRDRRWQRRASPFTLCSTTWSSGGGAAAQPAPSQIIAVRPDHYVLPGPYVDFVATMAKSLGAALGKPVHVEWIEKGEDAVDGLETLRCCALSAAVSFRAATVHISPEVSYAFFVMIPVQVVVPAGLNQTPLLQAVTDEFSAELWCATVVAVLGVAMATTLATRAILGLPLPSALNTALLQTLAPLLSQPPPGRTAHRPLAAVWLLMSVVIAAAYQGLLLKELTTPPGEINTLEQLEQSGLDCRMSRDLYQVDKFYLTEALRSRLTYVRLADLDAVVRDVADGRNSALIIRKDTISEILLSPYLKSSPPRLHSFETGNEYLGAYFKYTSGSPLQRPVVKFSLLAREHGIDLHMTQIIENKKSVDSRGGPDDEHLTRPLSLAQVLPAFLLLADGFVVAALVLAVEIVYHRWFAGRGGK

>FoccIr133

MKMLAIVVLLVCPGASDAVLPGPRPPLDSQGLAVLLASQPSRDKAKQFDVLVLGSAHGLGAFLGELPAETPRSLLSSTNVSSRTFCALQSTPSVALIAEDSWAQLAATVAGFEESQFPTHSPMLLWTWAEHDRGALGVLLRGVVSVPRLIVREIALAVSDRHGGGTVLFSLTTSTKLQVDVTEIDRWSPTDGRWLRRASPFTTLCSTWSGDRAPPAPSPSPQVVVAIRPHHDVAVEAYKDFVVTMANALGKSVRVEWVAGVDAVLDIVDRTRACSLPAAFSFIRAFPILAHMPPEVRYASFAIMSVQVVVPAGLDRTPFLQAVTDEFSVELWCATVVTVLAVVAATALVTRVTLGRRLASALSDALLQTLAPLLAQAAPGRTAHRPLSAVWLLMSVVLAAAYQGLLLRELTAPPGEINSLEQLEQSGLDVRVTGDLYTISKVYLPNALRARMTYVDVMGLDAVLRDVADRRNSALVIQGDMSAEILLAPYLHASPKRLHRFDLGRSYLAAHITTTTASPLQQRAARLALLVREHGLHAHMVSTIDKKQRLDVHGSRGAAEGEQLSRPLSLAQVLPAFLLLAHGYVTAALVLVVEIVYSRVCRR

>FoccIr134

MELLAILAPLLCLARCDAVLPEPRAPIDTQGVVELVSSFLSGSTSGVCVHGKNRALGLFLRQLPPETARLHTTSWNFPLTQHVQLAYGDNMFLITADSAAQLADTVAGTRLPALSRVLLWTWADSAPSAGDVRALGVVDLVWLFQKQAVLAVSTPDGATSLFSLRSEGKFDDAETRVVAEIDRWSPRAQRWQRQSSPFTSLCSTWRSSEPAPRYVLAIRPDRHVKHPQPYIDLLTAIANSLGLRVEWRDEDASFAGELRNRSLTCTLPAVFSFRSGPLWVRRSLSYEFFALNEVKVVVPAGLDPHATLLQAVTDEFSAGLWCATAAAVLGVAVATALAVVAVLGRPLVAALATAPLQTLAPLLGQAPPGRTAHRPLSAVWLLMSVVLAAAYQGLLLRELTTPPGEINSLEQLEDSGLSIRMSKDLFGYGPSYLSDTLRSRMTFVSSKDLQSAVRIAADGRDTAVILPWDLNSELLLSPYLMSKSHKLHSFQLGAPYLSAKLSFTSGSPLREALIRTNAWCLQHGLRLRMVRELVNHNRPDVHAKNDEGLARALSLRQLRPAFLLLAYCYGVSAAVLVGEIIYHKWFDRRGEN

>FoccIr135

MELLAIVALLIFLTGYDAVLPEPRPSLDAQGVADLVSSFVSGSKIGVCINGENHALDVFTRQLPPETARILSRNLTANLEHRLRLPTTDNLLLITAESPSLLADLVASTWTPGLSRVLLWTWAPSAPSTEGVLALGVATPRWLFQKQAALAVSTPGGATSLFALRSGLDRNSREESVNVTEIDRWSPRARRWLRQASPFTKVCTAWRSSEPPPRRVLAIRPHAYVAHPEAYTNFINYFVSRLGVRVDWRDEDRGLYKELEGNAFNCTLSAVFGFRYYPMVARASISFADVRLVPLQVVVPVGLDPHASLLQAVTDEFSAELWYATIAAVLGVAMVTVLATVAVLGRPVVVALATAPLQTLAPLLGQGPPGRTAHRPLSAVWLLMSVVLVAAYQGLLLRELTTPPGDINSLEQLEKSGLDIRITYDLYSFANQFLSDTLRSRTTFVSSGDLGSAVRMAAEERNIAVFLQKDINSHILTSSYVASKPPKVHTFVVCASFLSAKVVLTTGSPLQTPIERSTGWSAQHGLYRRMIKELGIRNRVHKEAEDGGREQLTRALSLDQLRPAFLLLAYCYGVSAIVFVCEIIYHKWSQGQGGN

>FoccIr136P

MELLAILALLLCPTGYDAVLPDPRPPLDTQGVVDLVSSFVSGSKTTVCIIGQNRALDMFTRQLPAETTRVVSRASLRSPRLNVEVAATDHLFVITAESAAQLAELVAGLWLPGLHRVLLWTWAPSAPSVADVHSLRVATPLWLLQKEAFLAVSTPGGATSLFAFRPVFNRDESVAQVTEVDRWSPRARRWLRQASPFTKICLAWRSSEPPPRHVLAVRPNPYVAHPEAYVNFVANIAARMGVRVAWRTEGPGLKKEVQLRSLNCTLPGILSFRNRAMWSRSAVAFARLVSXMVVVVPAGMGPRTTMLQAVTDEFSVEMWCATIAAVLGIAVATVLAAVAALGRPLVVALSTAPLQTLAPLLGQAPPGRTAHRTLSAVWLLMSVVLVAAYQGLLLRELTTPPGEINSLEQLENSGLDIRMDLDLHMFTPVFLSDKLRSRLSYIPASDLHSSVQVAAEKRNIAVALPKDMNSHILLSSYEKSDFPKLHSFLVGVPYLSANVAFTKGSPLQKPLTTAMAWCAEHGLYKHLTTKLGIRKRVHRDADNDEKLTRALSLHQLRPAFLLLAYCYGVSAIVFVCEIIYGKWFDRRAKQ

>FoccIr137

MEMLAVLALLLCPMGYDAVLPGPRPAPDTQGVGDLVTSFLSGSKGGVFVSGKSRALGTFLGQLPPETPRSLVLSKNTTGFQRLLLDLVSTEDMILITGDTATEMANAIASTFLPGLSRILLWTWAPSAEAALAHPMPTSVWLFGKQTALAVSTPDGGTTLFTLQAPVAVNRTRADGTYTVAEVDNWSPRVGRWLRRAAPFTKLCSTWRGGEPRLLQAFRPALFVANTKPYADFVSTIAHALRLRLEWLDEEDDHTLRRLLNTSLNCTLPAVLSFRSLPLVRRQSMRNGFLFFSAVQVVVPAGMDPHATLLQAVTNEFSPWLWCATAITLLGVALATALATVAVLGRPLASALANAPLQTLAPLLVQAPPGRTAHRPLSAVWLLMSVVLAAAYQGLLLRELTTPPGEINSLEQLEQSRLDIQMNYELLSYASAILPESVRSRVRYIPPQKLASAVRTAAEERSTAVILQMGLASRLLVSSYQAARPKKLHSFRVGVPYLSARVGYTAGWPLRGRLERFAQLIGEHGLYQQLDNAMAVHNRVNLHPTCEEVNEGGLTRALSLRQLRPAFLLLAYCYGVSAAVLVCEIIYHKWFDRRGRELVSVHSTTMVQQI

>FoccIr138

MELLAILALLLCPAGYDAVLPDPKASFDTQGVAVLVSSFLSGTTTGVCVIGKSRALDVFLGQLPPETARLHTTSWDHHVTQRLQFELSNNMFLITADNSARLAGNMASTTLPALSRVLLWTWADSLPSAVDVRAFSLVNSMWLYQKQTVLAVSTPDGATSLFNLSCESNRGRFGSGVNLAVAEIDRWLPDVRRWQRQAPPFTRLCSKWRSSDPAPRQVLAFRPGRHVVNPQPYIDFTTAIANSLGVRVEWRDEEPSVFAELRERFLNCSLSLLFSFRTLPMYSRHGVSYEYLSFAPVHVVVPTGMDPHATLLQAVTDEFSTGLWCATVAALLGVTVATALALFAVLDRPLLEALADAPLQTLAPLLAQAPPGRTAHQPLSAVWLLMSVVLVAAYQGLLLRELTTPPGEINTLEQLEDSGLDIRVSKDLLIFNASFLSDTLRSRMTYVSPNDLNLALRMVADGRNTAVILQKDLNSVLLVSSYLTSEPPKLHWFPIGVQYLNAEVAFTTGSPLQELMKRINGWGWQHGLPQRMVDDLANRNRADLHAEHEEENDDALTRPLSLRQLRPAFLLLAYCYSVSAAILVCEIIYHKWFDRRGGGN

>FoccIr139

MELLPVILAVLLCPAGYDAVLPEPRTLLDTQGVADLVSSFLSGSTTGVCVSGRSRALGVFLTQLPPETARLHINSWSFQRDHLLHMDDMLFLITADNAAQLADNIAGTRLPSLSRVLLWTWADSAPSAGDMRALGVKDSVWLFQKQTVLAVSTPDGATLLFSLSGEPGPGGGYVAAEIDRWSPGARRWQRQASPFTKLCSTWRSGEPAPRQILALRPDQRVAHPEPYIDYITALATAMGVGVEWRDDDASVLAAMRNRSLNCTLAAIFAFRSLPQTPRSSVSFAFLAFTPVRVVVPAGLNPHAALLQAVTDEFSAELWCATAAAVLGVAVATALAAMAVLGRPLVAALATAPLQTLAPLLGQAPPGRTAHRPLSAVWLLMSVVIVAAYQGLLLRELTTPPGEIDTLEQLEESGLDIRMSRDLYAYGPQILSHTLQSRLTYVSPDDLDSIVRMAADGRNTAVILQRDMNSHFLLSSYTKSKPQKIHLFQVRVPYLSTRVGFTTGSPLQESIKRVEGWCYQHGLQRRMAIELADLNRVDVRAEDEKDDVSMRALSLSELRPAFLLLAYCYCLSTAVLVCEIIYHKWFDRRVETM

>FoccIr140P

MRVTRLLAAAALLLCCREHGAAVLLAPRPPAGSQGVADLVSTFLANTKAGVVVYCRARSLLGAFLEQLVPETPRVRALTANLTIGSDRLSFELETTSTWXAGDSKAQMVDILSNVDMPLISRLLLWTAGHKAVPALDRGTYPWLLEKELALAVSTPDGGTVLYKVAIDLVSARAYSYKTAEIDRWSPVNRRWLRQASPFTRLCAGWRSPRRNGRDDAPLKVIASVPNRFAHSEAYIEHVTALMNSLRPPPRVDWQPERMASFQEAMTRIMNCTLPALVMSRSMPLMCCAKGLGYAPHPLTAVQVVVPSGMGRASTLAPITKEFSVELWCATAVAFFVVALATAWVTWKVFGQPLTAALSAAPLQILGPLLGQSPPGSRTAHRPLSGVWLLMSVVLAAAYQGLLLRELTSPPGEINTLEQLEQCGLEVRVVSDLHMYAPYLLKLLQSRMTFVHPSSIDSTVRSMARERSVALVLQVGVNNNLLLRPYMAPPKQLHAFRLAFQYPSARFAFSKGSPLAKSVEMLALRTLEHGLESRMALVTSLRMHLPLHYRDDEGRTRPLNLHQVWPAFILLTRGLSLSALVFVCEVVYYTWTDRGNHRSP

>FoccIr141NC

RALPKIQRVLFWASARRPAAEVLRNTTFVHHILRITNCGHQYGLVFSTVDGTAVSYASKCTAKQPFLEVDRWSPAQHRWLRGAAVFTQFCDQWQPPPSPADPLILTAVAKQENSHLLNVADFIDQHGMKPRKIKRVNFFLLNQTIAKFYRTWNQIFSQRNQCRMDALLYHYIILAMSDRELIYNYFSENIDVVVLVPAGLGAVVNPMDAVLVEFSPAVWLGTALASLFTAAALACTLGRDRGAALLLALAPLLGQAPGTPPPAG

>FoccIr142

MHAMAAILAVALLCAGLPSRALSAVPVVDVSPAPEAASAVALLAAFLGPHKAGVVVVSRVRWTDDFVRNLPRETPRVVLTSIARLTDMDSLLVRLKLSHNILMVVGNEPDELTAMLSRALPKIQRVLFWASGRRPAAEVLRNTTFVHHISRITSCVQQSGLVISAVDGTAISYVSKCTAKRAFLEVDRWSPAEHRWLRGAAVFTQFCDQWQPPPSPADPFTLLSVARQQNSYLLNVGDFIDQHGKKPRKIKRVNFLLPNQTSAEFYSMWKLILSQRNKCRMDTFLYQYMILAMSDKELIYNYFSENIDVVVLVPAGLGPVVNPLDAVLVEFSPAVWLGTALAALFTAAALACTLRRDRGAALLLALAPLLAQAPGTPPPAGRALRPLLGVWLLVCVVLVAAYQGLLLGKLSTAQTRGEINSLEDLEASGLPVYAQWDAFHRVNDILPDDTRKRMNVIHKYNDKYIYRNVVRDRKSAIVLFLSPAIRRLVRTWLVPTKMLHYFPIDHLNVIVMSLWTRGSPLGPVVTKVNRRAEQAGLYIHHHAMQDAVERLARERRLRPLRPAQPLTLRQMFPAFVFLIGGHVLAAAAFIGEVLSTYPGRWCGLDAA

>FoccIr143C

MHAMAAVLAVALLCAGPPGALSAVPVVDVSPAPEAASAVALLAAFLGPHKAGVMVVSRVRWTDDFLRDLPRETPRVVLTSMSYLTDMDRLLVRLRLSHNILMVVGNEPDELTAMLSRALPRIQRILFWASARRPVAEVLRNTTFVHHISRITNCGHQNGLVISTVDGTAVSYVNKCTATRAFLEVDRWSPAEHRWLRGAAVFTQFCDQWQPPPSPADPFTLISVSRQQNSHLFNVGDFMEQHAKKPRKIKRVNFLLPNQTKAEFSSMWDLMLSQRSQCRLDALLYQYMMLTMSDKELNFNYFSENIDVVVLVPAGLGPVVNPMDAVLVEFSPTVWLGTALASLFTAAALACTLRRDRGAALLLALAPLLGQAPGTPPPAGRALRPLLGVWLLVCVVLVAAYQGLLLGKLSTAQPRGEINSLQDLEASGLPVYAKWDAFQRVNDILPDVTRKRMHVIHKANDRKLIYEHVVRDRKSAIVMFLTPTVKRLVKTWLVPTKMLHYFPVDHLNVVVMGMWTRGSPLGPVVTKVTRRAEQAGLHIHH

>FoccIr144N

GKLSTAQTRGEINSLQDLEASGLPVYAKWDAFQRVNDILPDDTRKRMQVIHKGNDSEFIYKHVVRDRKSAFVLFLEPAIRRLVKTWVVPTKMLHYFAIDHLNAIVMSMWTRGSPLGPVVTKVNRRAEQAGLHIHHHAMQDAIERLARERRLRALRPAQPLTLRQMFPAFVFLIGGLVLAGAAFIGEALLSTLPGRRCGLDAA

>FoccIr145

MATVLALALLCAGAPGALSAVPIADVSPAPEAEGALALLAAFLPASKAGVVVVVGRVRWADAFLRGLPRETPRAMLTSLSSLTEAGRLSAEVQLTRNILIVVADGPDELIAAIDNAKLIFQRVLFWATGARPASELLRNTTFVHRASRHIHCDIQASLVLSAEDGTAVRYAHKCMVTTPFLEEDRWSTVDHRWLRGAAVFSQFCDQWQPPPSSREPFTLVAMARQVHSQASTIADFIELHGNLPRKIKRVNILIPNRPNDEYYRMWNLTLSQSRLCRWDAVLFHYRILTVSDIEVVHNYLSEKIDVVVLVPAGLGPAVNPLDAVLVEFSPAVWMSTALAALSTAAALACTLRRDRGAALLLALAPLLAQAPGTPPPAGRALRPLLGVWLLVCVVLVAAYQGLLLGKLSTAQPHGEINSLQDLEASGLPVYAQWATFQRVSDILSDNLRNQLKVNHNVDVRKLISKDVVRDRKAAIIMFLDHSARRLLQTWLVPTKMLHYFPIDHLNVIVMGMWTRGSPLGPVVTKVSRRAEQAGLHIHHHARKDAKERLARERRLRALRPAHPLTLRQMLPAYVFLIAGLVLAGAAFVGEVVNNSCVGRGRRS

>FoccIr146

MVALLISAFLAAATAASSSLPVVVDVSPSPEARSAAALLAAFLPPHRAGVVVAAGRVRWADDFLGALPQETPRVVVTDLSFYDEADSLAHAVQVRHNILMVMVDSPEDLAVEKFHHAAISFRRVLLWCSTTQPAHKILENETFLNHVTSRRLCSHQTTLALSNADGTTTLYALGNTRGCLDERFQELDRWAPADQRWLQGAAPFHPFCGKWRPPRSPKAPLNLFVLALNYSESPLFKLSFDLHSKIKLPWKVKLVDILVDKTVSRGQAITLAPILLHTEQCRLDGVLVNRPMTGVTGCDELTGLLSSEPYGIAVFVPAGLGPAVNLLDAVLAEFSPAVWLGTALAALSTAAALACTLRRDRGAALLLALAPLLAQAPGTLPPAGHALRPLLGLWLLVCVVLVAAYQGLLLGKLSTAQPRGEINSLQDLEASGLPVVVGPESFDFVEPLIPDELYGRVEIFQFLDANVVISNNVVLARNCAVITSLDRDLSRDIQVWLSPKKKLHFFLVGARHTTTVGYWHRGSPVGEAITTIVGRTQSAGLLEHYHSLRDDQERLAMAKYTTSLGIARPLSLRQMCPAFLFLGGGLVLGAASFVVEVLVTRLLTASCERFRASLLFG

>FoccIr147

MDVVLALTLLCAGAPGALSALPVVDMSAPPEASSASALMSAFVGPHKACVVVLGRARWTNAFLHGVPQETPRVVLTEQSLYEESDSLALQVRVRHHVLVVVADGPEALTLSTVTMTQSLARFHRALFWTTVATSVLEVVANKTFLGCVTQWRSCNAQTVLALTAADGTTALYALVPDGCPAPAEGEVPSATELDRWAPEERRWLSGASPFSKFCDSWLPPAPDEPFTLYVLGWGGAREHFDRLSDDLKRYNSLDMRITKTIVSLYDDRLQEVVARMKQCRLDGFVVRLNLQPLSSDEMTCFFDATPDPIVVIVPAGLGPAVNPLDAVLVEFSPAVWLGTALAALSTAAALACTLRRDRGAALLLALTPLLAQAPERPPPASRALRPLLGVWLLVCVVLVAAYQGLLLGNLSTARAYSEIDSLRDLEDSGLPVHVSWPFVGLLPENVRRRIVARRSMENPTDFIANRVVSARDCAIIVGLDEELAREIRVWLTPKKLLHFFTLGDLHVRINGVWSRGSALGKTIALAFRRYHEMALSQRYSALADFKERLSREKSMSALSPAQPLTLRQMYPAFLCIIGGHILSVVVFVVELLVSRLTDP

>FoccIr148

MDVVLALAILCAGAPGALPALSIVDVPVSPEARGAAALLSAFLPPHKAGVLLVGRVHWANAFLRRLPRYTPRVLLTDPFPAYYTEATSLNAHLHLRHNILMVVADRPEVLAASVANEMPRIPRALLWTTVARTAEEVVLNRTFLEYITRRRMCSDYTVLALSAPDGTTTLYELETRGCTSSGNSVQEIDRWSPADQMWLWGSAPFHQFCDQWKPPPSPSDPYILITIANSKDVIEQSNLIEQHNRLPRKIIHLSRSYLRVNGPDYKRHVANRSMHCRLDAILYYHPLPPPNAGEISTYFCVEEGHVVAFVPSGLGAAANPFAAVLEEFSPPVWLSTGLAVLCTAATLACAVHRGRTRQLALGMGMALAPLLAQPAVPGSLPPARHALRPLLGLWLLVCLVVVAAYQGLLLRKLFTANPSGEINSLTDLQQSGLPVYASNEALELVEDLLPEEVRNRVTRWHFVNTRKLIGDKVVRARNASLIIYMDRDLKRRLMHWFAPNKVLHYFAIRSFSLQTCSSWTRGSPLGAAVAKIIRRSEQAGLRLRYHELQDLRAELAREKRTRAAGLVQTLSLRQMYPAFLFLVGGLVLGAAIFVSEWFIFPWTRSDRANAGPTTSWRGVRRTAGVLRPVSERLAETSEERPSTVYVLRPAS

>FoccIr149

MDVALALALLSAAAPGALSALPVADVSAPPEATSATALMSPFLRHHQAGVVVLGQVQWTNAFLRGLPQEAPRVVLTQQSYYDESDSVTVQVQLRHHVLVVVADGPETLTLSTKTASQSVARFHRALFWTSVATSAPEVVANKSFLRGVTQSRHCGGQTLLALTAADGATTLYALAPPDCPNDGEAPARELDRWAPEEQRWLSGAPPFSKFCDKWLPPASKEPFTLYVSGKWGTGDRNFDLLAHDIKQYNKLNKNITKTAVNLHELQALVAKVRQCRLDGYLVLLHADVPSSDDLTCFFESMATPIVVVVPANLGPAVNPLDAVLVEFSPAVWLGTALAALSTAAALACALRRDRGAALLLALSPLLAQAPGGPPPLAGRALRPLLGLWLLLCVVLVAAYQGLLLGKLSTARSHTEINTLRDLEDSGLPVHAQWAAVSYLDRMLPDNVHRRLVVHRFMNPTDFIGKRVVSDRNCALIVPLSADLTRNIRAWLTPTKSLHFFAIHAFHVRVSGLWSPGSPLGKAIVVAFRRSHEMGLWNYYSALEDFRERLALEKARAALSHARPLTSKQMCPAFLCIIGGHILGAVVLVVELLVSRLTGHEARDKLPLK

>FoccIr150

MDVVLALALFCAGAPGALPALTVVDVSAVPEEAASAAALMSPFLRPQKAGVVVLGQVRWANAFMRGLPQETLRVLLTEPSYYEESDRLRVQVQVRHHVFVVVADGPEALISRTTGMTKSMCRFHRALFWTTAVATSAQEVVANKTFLGGITQWRDCNAQTVLALTAADGTTTLYALVPPDCSVAGEVVPPVTELNRCAPGEQRWLSGASPFSKFCDHWVPTENRDHFTLFVIGQWGGGNKLYDLLSQNIRRYNKFREKVLETVVNLDNRQLQVVLSKIKQCRLDGFLVLYQVEPPSSDDLTCFYDSMGTPVVVVVPAGLGPAVNPLDAVLVEFSPAVWLGTALAALSTAAALACTLRRDRGAALLLALAPLLGQAPDTPPPAGRALRPLLGVWLLVCIVLVAAYQGLLLGQLSTTRTHSEINNWRDLEDSGLHVHASWPVLNQLDGVLPENVRRRLIVHRFVSPMHFIPNQVVLGRNCALIVDMDEALARGVRSWLAPTKSLHIFTTGALFLRITGLWSPSSPLGKIIPVAFRRAHEVGLSHRHNALADFKERLTREKATEALVPARPLTLLQMYPAFLCLIGGHILGAVVLVMELLVSRLTVHEKP

>FoccIr151

MDVVLALALFCTSAPGALSALPVVDVSAAPEAASAAALMSPFLRPQRACVVVLGQVRWTNAFLRGLPQETHRVVLTEQSYYEESDSLTVQVQVRHHVLVVVADGPEALTLRSTSLTRSIGRFHRALFWTTVATSAPEVVANKTFLGRVTQQRDCYAQTVLALTAADGTTTLYALEPPGCPLEGEVPAVTELDRWSPGEQRWLSGAWPFSKFCDRWLPPAPKEPLTLFVIGQWGAGSIQLKALSHDIKRYSTLNKIVTTTTVNFHDGHLREVVARIRQCRLDGFLVKLHVMTPGSDELTCSISSEGTPIAVLVPAGLGPAVNPLGAVLMEFSPAVWLGTALAALSTAAVLACTLRQDREAALLLALAPLLGQATGTPPPAGRALRPLLGVWLLVCVVLVAAYQGLLLGKLSTTRTHSEINNWRDLEDSGLAVHAMWDALNQLEYTLPENVRRRVVVRRYINAIDFIGSRIVKDRNCALIIDLHEDVTRGIQAWRTTDKSLHLFRVGTSKFRFDGVWNPGSPLGKAIALMAARVPEIGVEKHCRNLADFKERLTQKKTTATRPLTLRQMYPAVLCLVGGHIVSTVVLVLELLCSCWSLRWSHGCRLTSQ

>FoccIr152P

MDVVLALALLCAGAHGALSALPVVDESAAPEATSATALMYPFLRHQKAGVVVLGQVWWTNALLRGLPGETPRVVLTEQSHYEESDSLPVQVQVRHHVLVVVADGPQALTLLTSTANLMVRFHRGLFWTKVATAAQEAVANETFLGCLTQWRDCDAQTTLGLTAGDGTTTLYRLRPRDCRLEGENAPPVTELDRWSPVKQRWLSWASPFSKFCDNWLPXPAPKEPLTLYVIGPWAGGGRHYDVLSNNIKRHSKLNKNIRRTVVNSPNSSLRSLLAEIRQCRLDGFLVILQAEPISSDELTCFFDSTATPIVVLVPAGLGPAINPLGAVLVEFSPAVWLGTALAVLSTAAALACTLRRDRGAALMLALAPLLGQAPGTPPPAGHALRPLLGVWLLVCVVLVAAYQGLLLGKLSTAQPHREINSLQDLGDSGLPVHAPWPVLKQLDGILPENVRRKLVVVHRFLHPIDFIANRVVVDRNCALIVEMDDDVARGIRSWLIPTKSLHFFTTGASFLRVNGLWTRGSPLGKAIAVALRGSLEGGLSQRFNALADFTERLTREKTTAGVSPARPLTLRQMYPAFLCLLGGHILGAAVFALELLVLNVQRAESS

>FoccIr153

MDVNLVLTLLCAAPAGVLSALTAVDTSAPAQARAVGALLSAYLPRQRAGVLLVGEARWANALLRDLPPETPRTLLADPSSYYEADGLVVDVQVRRTVLVVMWDRPEVATSGRYRLMPNVYRTLHCVTAEQPLLSENGNTLILKHMARSRACTGEIAFTFSSTNGSTTLYSRQLPGCEQDRPLLQELDRWSSTELRWQRGAAPFTDFCVWLPPPSPADPLTLFVLTREVKSFLFNISDSIGKKRVFRRKLNRSHQVLSYGEYGRIVKPAIEQWMKCRLDAVLFNFIHPTPHGNELTIFYNNEGCAIVAFVPAGLGPAANPLHALLAEFSPAVWLGTGLAALCTAAGLACTLHRGRWASLKMALAPLLAQAPLPGPAPPGARALRPLLGVWLLVSVVLVAAYQGLLLGQLSTAHPPGEIDSLRGLEESGLTVYASIDAFGVVGGMLPEDLRHRVQLWPDEKVQNFMNDFPRAQKCGIIMYMDSLLKRRIERRLAPKKLLHYFVLLTTNLRIKPMWHRGSPLGSVVAKTIRRTEEAGLVGRHDNLNYFKEQLKNRTMMALSLAQPLSLQQMWPAFLLIVGGLVFGVVFFVLEILTFLTGLRTPHF

>FoccIr154C

MDVFLALAALLSTGAPGVFSALAVDDTLVNSPEATSALALLSPYVRSERTTLIIIGQVNWTSAFLRGLHSEATWVLLPGEDETVFRKHYRNDVMETNIVAMVVADKPKDPLLWLQHDHVGYARFLFWVTLEPGMEGVISKVPHFSISVCMRPTALAITAPNGVTALYSVAQQDCGRRAPRLVKVDQWSTSRQSWRQRDAMLAVFGPFCSEWQPPATSKSTSIFRQALDGRESLFTELCAIFKEKTSGLPWNINVTSTIDLTRSNYETNAFMLWRAVRQCRLSLAYVEEGFVGRFDTRLAVILPEKKMYPVTVYVPAGLGPAVNPLDAVLVEFSPAVWLGTALAALFTAAALACTLHRDRGAALLLALAPLLAQAPGTPPPAGRALRPLLGVWLLVCVVLVAAYQGLLLGQLSTARPRGEINNLRALEDSGLPVYITWNVVGLTGTVWDLFTDNLRDNLRITREMKQTLHHVAVQRNCALITLHD

>FoccIr155NC

NSPEATSPLALLSPYVRFERTTLIIIGQVNWTSAFLRGLHSEATWVLLPGEDEAVFRKQFDDGMMDLNILAMVVVDKPKDPFSWLKNDRVGYERFLFWMTLGPGMENTIPQNPPSVMCSRPSGLAVTAPNGGTTLYSVARQDCSFRAPTLVEVDQWSPSRQSWRRTDAMFAVFRPLCSEWQPPTVSKSVNFFRFTGPGRESLFPELTAIIIRRTRAMGFPWNIKVTTENDIPHFNVKTITYTLAHAVEQCSLSILLLGNGVVGRLDTRQVVLMPEKKMYPITVYVPVGFGPAINPLEAVLVEFTPAVWLGTVLAALSTAVALACTLHRDRGAALLLALAPLLGQAPGTPPPAGRALRP

>FoccIr156I

MDFLVVLALTALLLSTGAPGVLSALAVDDTPVNSPEATSALALLSPYLRSQRTTLIIIGQVNWTSAFLRGLHSEATWVLLPGEDEAVFRKQFDDGVVDNNILAMVVADKPKDPSSWLQNDHVSYARFLFWMTLGPGIEDFISKVPHFLFSACIQPTALAITAPNGSTTLYSVAQQDCGRRAPRLVKVDQWSTSRKSWRQRDVVLAVFGPFCSEWQPPAANRSASVFRQAADGRASLFAELYAISKEKTRLLPWNITITSTISLTSSNFETLTFVLTRAIRQCRLDLAYVEEGFAGRFDTRLAVILPEKKMHPVTVYVPAGLGPAVNPLDAVLVEFSPAVWLGTVLAALFTASALACILRRDRGAALLLALAPLLGQAPGTPPPAGRALRPLLEAGILDHWEDDLDEGNRVLSARHLSALKRVLPLSLQHMAPAFYMLLVGLVVAAVVFAVELATAKLGRARQSPGTATLRWASHRQPKSGAGSAAAHPRF

>FoccIr157C

MDVLVVLALAALLITGAPGVFSALAVDDTPVVSPEASSAMALLSPYVRSQRTTLIIIGQVNWTSAFIRGLHSEATWVLLPGEDEDVFLNQFYTNAMDANIVAMVVADKPRDPLSWLQNKHAPYARCLFWMTVGVGVGTEDVISKVNFSISALARPIGLAVTAPDGSTTLYSVTQLVPNRRAQKLLKVDQWSTSRQSWQRSDAVFGVFSPFCREWQQPLANESVYLFRLAADERESLFSELYPIFIRKTRG

>FoccIr158C

MDVLAVLALAALLCTGAKGLLSALGVDNASVNSHEATSALALLSPYVRSQRTSLIIIGQVNWTSAFLRGLHSETTWVLLPGENEDVFLNQFQTNAMDTNIVAMVVADKPRDPLSWLQNEHAGYARFLFWMTLEPGMEDVISKVPYFSISTCIRPTALAITAPNGSTALYSVAQQDCDRPRLVKVDQWSTSRQSWRQRDALLALFGPFCSEWQPPAANKSTNVFKLSADGRKSLSAELCAIYQRKTSGLPWNVNVSSSTKLGSPSYTKVTLALWRALGECRLSGAYLEDGVVGHFDFRQVGIMTEKKMYPVTVYVPAGLGLAVNPLEAVLVEFSPAVWLGTAMAALCTAAALACTLHRGRGAALLLALAPLLGQAPGTPPPAGKALRPLLGVWLLVCVVLVAAYQGLLLGQLFTARPRGEINSLRALEDSGLPVFVTWNVVGLVDTVSDLLTDNLRDNFRITIEIKQTLQHVALQRNCALITLHDRFVQRFVHPLTTHTQRKLHSFTIGSETLKIIAFWSRGSPLGES

>FoccIr159

MDVLAVVPLLVLVCAGAPGVLSALAVADLPVHAPEATSALALLSPYVRSQRASLIIIGQVNWTGAFLRGLGSETTWVVLQGEEDEAVFRKRFDDDVMETNIVAMVVADNPKDPASWLQKDHPGYARLLYWLTVGPGMEEAIPKIPHFSIPMCTRPVGMAVTAPNGSTTLYSVAQQDCGKNAPTLLKVDQWSTSRQSWQQTDAIFALFKPFCSEWQSPPANNSVIVFKLSTDGEESPFSRLFDIFVRNDRSLPRNLNLSSYASFNSNSFSFETMNFEVGRAREQCRLSVAYIEDGVVGRFDSRTVSIMTERKMYPVTVYVPAGLGPAVNPLGAVLVEFSPAVWLGTALATLSTAAALACTLRRDRGAALLLALAPLLGQAPGTPPPAGRALRPLLGVWLLVCVVLVAAYQGLLLGELSTERPRGEINSLRDLEASGLPVYLTWNVVGLGDNIWDSFPESLRRNKHSALMDKKTLRLVARHRNCAVITLQDRLVQRLVHSLTKPPHRKLHSFTIGSETLKIVAFWSRGSPLGERMALAYSRLDEAGILDHWEDDLDEGYRVLSARHLSALKRALPLSLQNMAPAFYVLLNGIVLASVVFVVELVTAKWGRAGQSP

>FoccIr160C

MDITVVMVLIFSAAPGAHSTLPDVDMTAPPEATSALALLSPFLAAQTNTLLILGQVNWTGAFLRGLSSETSWVLFPDERAENDRYLDNFGVGNNCTALVVADEPADLYAWMDRKFRLINGCRYLFWSTTKKDKQLAPPKVLSITLCIKPTGLAVTTRNGSTTLFSVVRQHRVKDPPILLEVDRWSPLDGRWEQTGGLLRVFGPFCISWRPPAAGESLTVYEIIKDGETPLPEYFKTLRAKIMDLPWTLNTKSTPFFNDNYLSGIILESWQKIDHCQLSLIFAEDYESVMDSRKVSVMAEKGTFEISAFVPAGFGPAINPLDAVLVEFSPAVWLGTAL

>FoccIr161NF

SPILLEVDRWSPSDGRWEQTGALLRVFGPFCSSWRPPAAGESLTVYEITRDGETPHREYFKTLRAKIMDLPWTLNTKSTTFFNDSYLSDIIVESWKMMDNCQLSLIFNEDDELGMNSRKVSVMAEKGTFEISAFVPAGFGPAINPLDAVLVEFSPAVWLGTALAALSTAAALACTLRRDRGAALLLALAPLLGQAPGTPPPAGRALRPLLGVWLLVCVVLVAAYQGLLLGELSSERLRGEINSLRALKETGLRIYVAWGSILLIEKELFTDTTNNIYFTPNVGEILQHVAQHRNCALIAVQDRALQRLLQPLTIPQKKVHSFPIVAGKRKLIASWTPGSPLGELSRSAYELLDQAGILDYSENKLDEGNKVALARYLVSLQQVRALTLKHMAPAFYVLFIGLALATLVLGVELATARWCGAQGRTVTPAGQRPRGPALTRLYGRQNIRRRRPAAALRPRARGGPPQDPVAYQRGGDSRLAPDIPLVQLQHRGPRRTAWR

>FoccIr162

MGITVVLVLIFTAAPGAHSALPDVDMTAPPEATSALALLSPFLAAQTSTLLILGQVNWTGAFLRGLSSETSRVLLPDERAENDQYLDNFGVGNNCTALVVADEPAEPYSWMERKFRLINGCRYLFWVTTKKDKQLAPPKDLSIALCVKPTGLAVTTRNGSTTLFSVVQQSHIKGSPILLEVDRWSPSDERWEQTGALLRVFGSFCSSWRPPAAGESLTVYEITKDEKTPLPEYFKTLRAKIMDLPWTLNTKSTPLLIDNYLSDIVVESWQKIDHCQLSLIFAEYDALGVNLRKVSVMAEKGTFEISAFVPAGFGPAVNPLDAVLVEFSPAVWLGTALAALCIAAALACTLRRDRGAALLLALAPLLGQAPGTPPPAGRALRPLLGVWLLVCVVLVAAYQGLLLGKLYTARAHTEVNSLLSLKKTGLPIYVTWGSHNVLAKEFYTEPDIKHKLHLATDVSKTLQHVAQHRNCALIAVHDRALQRLLQPLTIPQKKVHSFPIVPDNRRLIAAWTHGSPLGELSHLTYERLDQAGILDHWEDKLDEGNKVALARYLVSLQQVMALTLKMMAPAFYVLFIGLALATLVLGVELATARWCGAQGRTVTPAGQRPRGPALTRLYGRQNVRRRRPAAALRPRARGGPPQDPVAGQRSGDSRLVPDISLVQLQQHRGPRRTAWRGQRPTCQGMPCRV

>FoccIr163

MDIAVVLVLLCTAAPGAHSTLPDVDMTAPPEATSALALLSPFLATHKSTLLVVGQVNWTRAFLWGLSTETSWVLIPDERAQSDQYLKNFGVNNNCTALVVTDKPTDLYTWMGRRASLVGGCRFLFWVTLRSDKQLALPKVLSYLLCIKPTGLAVTSPNGSTTLYTVVPKYCVNGSLKLVEVDRWSSADQSWQGADVVFRVFDRFCSRWRPPVAEESLTIYVIRTAKEKRLPEYFKTIRSKIVNLPWKVNLLTTSYSPDNYLTDTITEVWQRIDHCKLFAAFIDMGDPKSIDPKKISALAEKGMFEISAFVPAGFGPVANPLDAIRVEFSPAVWMGTALAALSTAAALACIIRRDRGAALLLALAPLLGQAPGTPPPAGRALRPLLGVWLLVCVVLVAAYQGLLLGELSTTRAPGEINSLRELSKTGLPIYVALSAVNLLETELLSDFDLHKIFAATNVGKALQHVAQYRNCAMIAIQDRTLRRLLQPLTIPQKKVHAFGLVAEFRRVFAVWSPGSPLGELSRATHERLDEAGILDHWENKLDEGNKVALGRYLLSLQQERALTLKHVAPAFYVLFIGLTLATLVLGVELATGRWCGAGRRTTPARWPAGSTRLSGGQDVLRCRPAAAMRLRARGGPLDPRAGGRRAPDIPLVQLHGGPRRTAWQ

>FoccIr164

MDAALVLVLVCTAAPGALCALPDVDMTAPPEATSALALLSPFLAAQKTTLLILGQVNWTRDFLRGLSPETSWVLFPDERAQNDRFQGNFGVEVNCTALVVSDNPADPYSWMERKAKLTNYCRVLFWITTDKQVAIPYSFHICNKLIGVAVTAPKGSTTLFTVAPKYCGSVRPTLVEVDRWSHENQSWPRADAIFRIFYPFCSRWRPPAADESLTVYEMLREDGKPLPEFFKTLSTKIRGTRRTGNIMSNLAFPGNNLANIIVESWQKIEKCQLPLMLLEYGEVVNFDPRKVSVMAEKAMFEISAFVPAGLGPAVNPLDAVLVEFSTAVWLGTALAALSTAAALACILRRDRGAALLLALAPLHGQASGTPPPASCALRPLLGVWLLVCVVLVAAYQGLLLGKLSTARDHAEVNSLRALKETGLPVYVMWSALTMLEKELIFDSHKPHTIPLGMNASETLQHIAQHRNCAMIAIHDRTLRRLLHPLTIPQKKVHSFPLVPEKIRLIAVWTHGSPLGELSRSTYEHLDQAGILDHWEDKLDEGNKVALARYLVSLQQVMALTLRHVAPAFYVLFIGLALATLALGVELVTARWCGARGSTATPAGQRPGGSRQGVRRCRPAAALRPRARGAGRLAPDIPLEQLHCHPVNRAVD

>FoccIr165

MDVTVVLVLVFTTAPWAHSALPDVDMTAAPEATSALALLSPFLAAQKTTLLILGQVNWTRDFLRGLSQDTSWVLFPDEKAQNDRFQGNFGVKVNCTALVVSDNPADPYSWMERNTILVDDCYFLFWMTTGSGVLPEAHYHLFCEKPTGLAVTTPNGSTTLYILPPKSCVNANEASPPLVEVDRWSPADLRWRRADAVFRLFHPFCSRWRPPAEGESLTVYELTGAEAEASSLPEYFKTMRTRVRGLQWTTNVESNPVSFDVNLKNIFLSSQNIWECQLSLMLLEVGDVMQVNRNKVSVMAEKRVFEISAFVPAGLGPAVNPLDAVLVEFSMAVWMGTVLAALSTAAALACTLRRDHGAALLLALAPLLGQAPGTPPPAGRALRPLLGVWLLVCVVLVAAYQGLLLGELSTERAHTEVNSLRALRETGLPIYLTWDSVRLLKTELFPNFDAKQKFSFVTNVGETLQHVAQHRNCAMIAIQDRTLRRLAQRLFIPHKKVHSFPIVAEDRRLIAAWSPGSPLGELSRSTYERLDQAGILDHWEDKLDQGNKVALARYLVALQQVRALTLKHMAPAFYVLFIGLALATLALGVELATARWCRARGRTATPAGRPGSRAPVCGGQDVRRRRPAAALRLRAPGPGAGPLHPQAGPRRASDILLVPLHRGSRRTAWW

>FoccIr166

MEAALAPVVFLLCCAAPGVRADLPVVDTAAPPEAVSAAALLTPFMAPQRATLVVLGRANWTGAFLLELAADVPRVLRPGFAERAAEPPQLEHALMSTRCAHLLPVDGLEDLLVALTTYDATTIPTRVLFWAAVAAPRDAVLQRVMRTRMWLGTHHYTLALNAPDGSVSLYKMTCDQVPATCRLGAKVITETDTWSPAEGRWLQGSAVFAEFCDWRPSRGGKSLVLTVFVLTPKDSRNASHVVELTKCLEHVAAGRRRRSLGAAVRFEGTSNYSRVHGAMKRCTLAAVLAARSLYGELSSTEVISVTAMKMAHVAVIVPAGLGATVSVMDAVTVEFSAMLWLGTGLAALCTVLVLTCTRRCQDVGGAVLQALAPMVGQAPPPPAPPRPMLAAWLLACVVLTAAYQGLLLGKLSSAVPRRELDSLRDVEDSGLPLKVLGNLAHSNVLTDNLASRAEYLVHAEIKAVIDTIATARNCALVAYLDDGVVGALRPFLLPPEKLHVIRLPYSAHNIIGMATKGSPLEAPLVRTLGLVEAAGLPKRWRLEGHERGRREHARRLASLREPRALALRHLHPAYVVLGMGHAASAAAFALEALCGAVVFGRG

>FoccIr167

MEALLLAVLILCCAAPARVVRADLPVVVGGSAPPEAEAAAALLTPLAEPQKATLVVMSDSSWAAAFLRELSAGVPRVLSPGWVALARRFDPLLENVLMAQRCVTLVTAEGLEDLLTTVRTFGAPRVTRVVFWSSVSSPRDGVLQVVSRTGLWLGGRQYVLALTAPDGSTVLYSLNCFTRWTCNNAVMTIREVDEWSPGLRRWRRGAAPFREFCSGWRPRHGQGQSLNVSLVSARGLSPRSSLLDLANSLARLAGQDRRLQRVGLGPHTIYREAIDYAELASRLHECSVDAVVADTPLDVLRVPTEAIEVYDLEMTHVVAIVPAGFGASVSVLDAVTVEFSAEVWWATGLAMLCAAVVLACARGQDVSGAALQVLAPMVGQAPPPPAPPRPPLAPWLLACVVLTAAYQGLLLGKLSTAVPRRELDSLSDVVDSGLPVHVDHILSWRLTGKNVLPENLTAGGKFVALTEVETIIDTIATARNCAFVLLLSRRAQDYVQRYLVPPKKVHLVHLAYSDRNVIGMATKGSPLARALTSLLARVDAAGLLTHWRSAEFERERQDRAKRLVALQGPRPLTLYHLGPPFVVLGVGHVVGLAVFALEAVWGWMAGPRASCRRSASARA

>FoccIr168

MEALLVLASLCCAVRAGLPVVDTAAPPEARSAAALLTPFMLPQNATLVVLGRVGWTGAFLEELSADVPRVLTPSVMEIKRVDVRESHPRLAHALVSTRCAFLLPVDGLDDLLLALSSFDPFAFPTRVLFWATVAAPRDAVLRRLARARASMWLSRYDYAVALTAPNGSTTLYRLAYANLAAAHRDGTKIVTETNSWLPATGRWRRGSIVFGEFCDWRPGKSQELIAFVVVISKRGLTPSSPLSELSKIVKQVAAGHRGRTTVRYEQTAEYGRVHDALKACTLAAVLADRPLYSESTSVELTLVPGMEMAHVAVIVPAGLGARISPLDAVTVEFSAMLWCATGMAALCTVLFLRCTRRQDVSGAVLQALAPMLGQPPPPPAPPKPMLAAWLLACVVLTAAYQGLLLGKLSSAVPRHELESLQDLEDSGLPLKVLKNLAHSRVLTNNQTCQAEYVVYAEVDETIDTIASARNCALVTYLDEDAVNSLRPFILPPNKKLHVIQLPYSEHNVIGMVTKGSPLGHALVRTLGLLEAGGLPARWRAEEYGRVRQERARRLVMLQEPRVLTLWHLHPAFVVLGVGHAAAVAAFALEALCAWRRARRPSRAAVRRVQSHDETS

>FoccIr169

MQVFLVLTCLCCVVVVPGVRADLPVVDSAVTPVEARSAAALLTPFMQPQKATLVVLGRARWTGAFLRELSADVPRVLRPSVIALKRNLIHDMRLENHLIDMRSVHLVHTGGLQDLLATMRVYDNRRPRRVIFWATLATPRDEVLQNVSRSGLWAGGHQSLALTAPDGTTTLYNLTCSSDHTCMSRTMVISRTDEWSTVEQRWRRGGAVFMPFCGSWHMRGNSNSLNMRLLTVKGYTASRTAVELAKSVIRLVGGRDFQQRVGSNVTVRYEEIFEYRILDSVLKTCNLDAVIGDIPFDLWDDTVEKRKVPDMRMNHFAAIVPAGCGASTGILDAVTVEFSAMLWCATALATLSTVVVLMCTGPQDVSGAVLQALAPMLGQPPPPPAPPRPMLAAWLLACVVLTAAYQGLLLGKLSSAVPRRDLESLQELKDSGLPLKVIATLEHSNVLDDLDAQAEYVAFTEVKGVIDTIATARNCALITLMNRHIDSHMLPFMMPPKKLHVILLPFSGVKIVGVATQGSPLLRPLSRVLALAEAAGLLEHWRSKQYEWEHRDHARKLASLQGPRPLTLFQLGPAFTVLGVGQAAGVAAFVLEALCTWWPVSRPSRSLRRRTES

>FoccIr170P

MEALLVLSLFCCVATPGVRVQADLPGVDSAAPPEARSAAALLTPFMRSPTATLVLFGRTRWTGAFLGELSADIPRVLNPSVLPQNYGYLPHPHLEYVLMGTHCVHLVTAHGPEELLAAMRAYRPVPVIRVLFWATVETPRNEVLGRVSRSGIWLGFRYEYRMALTAPDGSTTLYNLTCQEDDTVAFRKRVLVISRANEWLTAEQRWLRTAAVFHPFCTGWRSGGKSRPLTLFLLTAKGLTTSHSIKELAKSVCQMGREQRHRIGSPKTVRZKVVSRISGEIRLAFRKCALDALLADVPLNIPGETMDINNVLGLEVVPVIAIVPAGFGTGVSVLKAVTVEFSAGLWCATGLATLCTVLFLRCARTRDVSGAVLQALAPLLGQAPPPPGPPRPMLATWLLACVVLTAAYQGLLLGKLSSAVPRRDLDSLQDLEDSGLPLKVLTNVMYSHSLPLSNNLSTQAEFVAYPEIRTIIDTIATARNCALITMMDRHIGSHIQKFMIPPKKLHLFQLPYPGYSVIGKASRGSPLERPLSRALALVEASGLLSRWRSEEYARERRHHARRPVLQQGPRPLTLWQLGSAFVVLGVGQAAGLAAFALEVLCAWHARWSK

>FoccIr171

MKALLVLPLLCCAFAVPGVRAELPVVDLAVPAEARSAAALLTRYMQSQNATLVVVGRAHWTGAFLGELPDDVPRVLNPSVLAPQPRIDPRLEHTLKTTRCAYLIPVDGLENLLVTLGTYEPTTLPTRILFWATVAAPRGAVLRLIASASPSVWLTRHHYDLALSARDGSTALYHLVCDQSAAACRDGAKTITDTDTWSPVEGRWRRGVAVFDEFCSDWRLRRGKSRELTVFVFPGNGVIISPPLLELTKSLQHVEAGRRRRISEATVRYANIVEYRPVRNAMKECSLAAVLGDGSLFPEYTSVEETIVPGMEMTHIGVIVPAGLGAFVSVLDSVTVEFSAMLWCATGLAALCTVVVLTCAQRQDVSGAVLQALAPMVGQAPPTPAPPRPMLGAWLLACVVLTAAYQGLLLGKLFSAVPRRDLESLQDLEDSGLPLKVHRHLSHSNVLTDNLASRAEYVLYSEVVTTIDTIATARNCALVTFLDAEVINSLRPLLLPPKRLHVIRLPYLGQNIVGMATKGSPLAQTMIRTLGLVEAAGLLARWRYEQYQQELLDRHRKQDPLREPRVLALRHLYPAFIVLGVGHAAAVAAFALEALSAWRARLPSPSRRVGS

>FoccIr172

MEAFLSLCVFCCAMPGVRANLPVADSAAPPEARAAAALLTPFMHSQQATLVLMGQTRWTGAFLRELSADIPRMLSPSIVTLYQQHMVSTRIVHVLAATRCVHFVAAEGLDDLLSTMRMHDNTKLTRAIFWASVASPSTRDEVLRDLSRARVWLGGHQYALALTAPDGSTGLYNLTCRSDAACRNRVVTVLEQDTWSAAEQRWHRAPAVFSEFCVSRPERTCRVLTVLMLRSHGRTASKSLLELTKSVVHIVRQAQQRQRLGSQKTVRLQETSEYSHIDAALKECTLDAVLSDVPLFTLSESALQTSELPDMEMARVAVIVPAGFGAGVSLLDAVLDEFSAALWWATGVATLCAVLVLSASARRRDVSGAVLQVLAPMLGQAPPPLPPAPPRPMLAAWLLACVVLTAAYQGLLLGKLFSAVPRRELDSLRDVEDSGLPLKALANLARANILAVNLTTRIEYVTYSELQTVIEAVATARNCALITYLDRHAQSYMLPYMMPPKKLHLIQLPYSDFNIIGLTTKGQLDRLMSRALMQVEAAGLLARWRYAEYERERHHFARKLVLQQGPRTLTLYHLGPAFVVLGAGQAAGLAAFALEALCAWLGRREVPG

>FoccIr173

METILALCLICCAWPGVRADLSVVGSVAPSEARSAAALLAPFLTSQAATLLVTGETPWTGTFLRELPADTPRVLNPSSHAMNLRLEFVLSATHCAHFFHTDDLETLLYTMGGHDPTTMTRALFWVNEARPRDEVLRLVSQSGMWLGGQQYALALAAPNGSATLFNLTCSSEAACRTRTMTLIEVDEWSPVEQRWRRGAQVFHEFCAWREGQVPAVLMLTMRSATFTTSIMELTKSVVVLARRRHRQRSGSTTAVSYHEVVGYHRINIAAKECSLGAVLSDVPLTVLGKSAEIASVHDMEMHRVVVVVPAGFGASVGVLEPVTVEFSAALWCATGLAVLCTVIVLTCTCRQDVSGAVLQALAPMLGQPPPQPSPPRTMLAPWLLACVVLTAAYQGLLLAKVSSAVPRRELDSLRDVADSGLPLYALGNLVRSSVLPTDLAARTRWIHYLQTEAVIDTIATARNCALVTYLDGYTTRTLHRHLVPPKRLHLVHLQYADFKVIGLATRGSPLEPLMVTALARAEAAGLLARWRGAEYEREDHDYARKLAPLKGPRPLTLWQLRPAFVVLGVGQVAGVAAFALEVLCALSSTTRTRVSFRRTAWS

>FoccIr174

MEAALVVLLLCCCCCCPAPGVRAGIPVEDLAVAPEARSAAALVTPFMKPQNATLVVLGDSRWTGAFLRELSGDVPRLLYLKPRHSFIAAKLDDDYSDYSNSNTQVNRLRLELMAIHCLLLVPTNDLEELLFIMRTYSNIGNTRVIRALFWTTVSTSVPRDKVLRRISQQKTWLGGYQHALAITFPDGSATLFKLTCSSDDACSRREMTILETDTWSPVDKRWSLATAVFSEFCSGFRRINKDQDLTMFIHTKNDLTNPSSLKELTKSVVNLVSRGRQPSESPVKYQEANKNEGLYVTLQECSLAAVLLDGCLLDVYKMSEISFITDMEMASFAVIVPAGLGASVGVLEAVTVEFSEALWLATGLATLCTVVVLKFARRQDVSGAVLQALAPMLGQAPPPPAPPRPMLAAWLLACVVLTAAYQGLLLGKLSSAVPRRDLDSLRDLEDSGLPLKVVVSLARRNVLSNNLTDRAEYVERTEVKTVIDTIATARNCALVTLLDRHTHSYMRPYMLPPKKLHVIRLRYYSFNVIALATKGSPLERPIAKVQGRVEAAGLLARWRRAEYERESQDDARRRLSLEGPRPLTLWQLQPAFVVLGVGQVTGVFVFALEALCAWRARWA

>FoccIr175

MEVLLVAFFLCCAVPGVRADLPVVDSAAPPEARSAAALLTPFMAPQKATLLVLGRARWTGAFVGELSADVPRVVSPRLSRSSSAFDLYVRTTTRQLDRLQYALIATHCVLFVPTDDLEELVLVMSIYSNIRRSIDIRVLFWTSATAPREEVLRRISSLNTWMGGRQHALALTAPNGSTTLYNMTCTSGACCSREMILLETDKWSPAEQRWRLGVTPFHKFCSGHRLTGKRQDLTMFVHTKRGFTSTSSLMELAKSVVDLVSRGRRQRIGSGTPVKYQQIWDYNIVHIAMKECRLAGVLTDEALLDLDRLAEMSIIPDMELAEFAVIVPAGYGSSVGVLDAVTVEFSQTLWLATAIATLSTVLVLKFARRQDVSGAVLQALAPMLGQAPPPPAPPRPMLAAWLLACVVLTAAYQGLLLGKLSSAVPRRDLDSLQDLEDSGLPLKAVGNLARTNVLPYNMADITEYISHSEIKAVIDTIATARNCALVVCFDRHIRSYLRPYMIPRRLHVMRLSYSHLNIIALTTKGSPLERPIATAQARAEAAGLLARWRRAEYEREYQDDAKRLGSLRGPRSMTLWHLQPAFVVLGAGHAVAVMAFALEAVYAWRAPC

>FoccIr176

MEALLVVLLLCSAAPGARAGLPVVESAAPAEAKSAAALLTPFMTREKATLMVLGSSRWTGAFLRELSADIPRVLGPRPSRGSTVSKQYNHSNTQLDRLEHALVSTRLVHFVPTDDLEELLFTMRTYPRIRRTSDLRVLFWSTIVDRREEEVRRIPRLNTWLGFHQHALAITAPNGSTTLYNMSCTISACWKLTVPTTIVGLDEWSPVQQRWRLGVAPFHEFCSGYRLIGKRQEHTVFLNTMKGFTNLSATHTLVKLAKGTVKLFSRSSQRRIGSYSPVKFQQVRSYVRVNIALKECTLAVLFADEYFLSQKELAEMSFIPNMELAEFAVIVPAGYGASVGVLDAVTVEFSSALWLATGLATLCTVVVLKFARRQDVSGAVLQALAPMLGQSPPPPAPPRPMLVAWLLACVVLTAAYQGLLLGKLSSAVPRRELDTLQDLEDSGLPVKARGNLRYFRVLTDKLSARTEKVSKSEVKTVIDTIATARNCALVTFNSLHTRSYMRPHMLPKKKLHIIRLSYRHFNVIAITTKGSPLEGPIATAQARAEAAGLLSRWRHAEYEREYQDYVKRLVSLRGPKSMTLWHLQPAFVVLGAGHTAAVVAFALEAVWAWRARWAGRKTG

>FoccIr177C

MEALLVAFLFGCAAPGVRAGLPVVDSAAPPEARSAAALLTPFMAPQQATLILRGSSRWTGAFLRELSADIPRVLIPRPSRLPSAWKQDNYLDTLQLDRLVHGLISTRLVHFVPADDLQELLYAMKNYPVINKVKDIRVLFWTTLLTVSAHQDRVLRQIFRLNTWLGSRQHVLALTSYNGSTTLYNLTCTSSDACLRMDATVFEMDRWSPVQLHWRRRATPFYEFCSSFRSIGKHQNLTVYIDNTQIGLTNVSSLMELTKSAVNLFSGGCHRSMRSQPSVSYQLTRSHDRIILALRECTLAAFLSDEDFFARDETADFSFLLDMEMAGFVVIVPAGYGAS

>FoccIr178

MEAALLALCLLLCCAAPGVRANLPVVDSAAAPEAMSAAALLTPFLAPQQATLVVTGGTRWTGRFLRELSADIPRVLDSNPFALSIRHHIHPRLELVLEATHRVHFFHTGDLETLLSAMRTFDPLRVVRAIFWANVASPRGEVLRQVSQTGLWLGAMQYALALTAPDGSTALYNLTCATEDACEHKQMTIIETDLWSPVAQRWRRGAKVFQEFCTAWRPSQAPAVFMLTKKRLTTLSELTKSVVNLAGRRYEQHRSGSDAAAGSHEVIGYHRLTRALRECALGAALTDLPVFFVGRSDEITVVVDMGRHHVGVIVPAGFGPSVSVLAAVTVEFSAMLWCATGLATLSTVVFFRCARTRDVSGAVLQALAPLLGQPPPPPAPPRPMLAPWLLACVVLTAAYQGLLLGKLSSAVPRRDLDSLREVQDSGLPILTLQQLALTGNMTVPFSQVESTIDVIATARNCALVTFLDHYALRALRRHTVPPRKLHLIQLKVSSFKVIGMATRGSPLEQPMVTAVARVEAAGLLAHWRRSEFTREDAHYGRKLALLRGPRPLTLFQLGSAFVVLGVGHAAAMGVFALEALWAWRAHKIKVLPARPAH

>FoccIr179

MEAALLALCLLCCAAPEVRANMPVVDSAAAPEARSAAALLTPFMAPQKATLVVIGEARWTGAFLRELSADIPRVLDSNPFALSIRHRHHTWPRLELVLEGTHRAHLFHTEDLETLLSAMRTRDPTQLVRTIFWANVASPRAEVLRQVSQTGLWVGGTQYALALTAPDGSTALYNLTCATEEACEHKQMTIIETDLWSPVAQRWRRGATVFQEFCTAWRPSQAPTVFILPRKGSTTTLPVLELTKSVVNLAGQGYEQHRSGSDAAVGSHEVVNIHRLSTTMKECALGAALTDMPLTVLGGGDENTYIVDMGRHHVGVIVPAGFGPSVNVLAAVTVEFSAMLWSATGLATLCTVMFFRCARTRDVSGAVLQALAPLLGQPPPPPAPPRPMLAPWLLACVVLTAAYQGLLLGRLSSAVLRRDLVSLREVKDSGLPILTLAQLAFTGNFLTANMTVSYSQVESTIDVIATARNCALITYLDHYALRAVRRHTLPPKKLHLIQLSHPIVKVIGMATRGSPLERPMVTALARFEAAGLGAHWCRLEHSREEAHYARRLVLLQGPRPLTLFQLGSAFVVLGVGHAAAVGVFALEALWAWRAHKIKVLPARPAH

>FoccIr180

MEALLVVFLLCCVPPGVRASLPVVESVAAPEARSAAALLTSFMKPQRATLAVLGDSRWSGTFIRELSTDIPRVLIPMLNLSLSLASEQEEKYSNTQFDRLYHALMVTHWALFVPTDGLEQLLFAIRNLAQHKGSMVARILFWTTESMFANRDKVLRRFSETKLWLGGCQYALALTSTNGSTTLYHLTCTSDDACRSRQMTILETDEWSPVDQRWRLGADVFHQFCNGYRPIGKHQDLTMFVQTIKGSTNKSSLLELTKVVVDSVSRGRHRRLGSNTPVKYQETWRYRGVHVAMMECNLAAVLTDESFFGLRGLKEMGYIPDRQMAVAAVIVPAGYGSRVSIVESVTVEFSSALWLATGLATLCTVVVLKCTRRQDVSGAVLQALAPMLGQATPPPAPPRPQLAAWLLACVVLTAAYQGLLLGRLSSAVPRHELESLRDLEDSGLPVKMSAYLAYSHLLTDKLTARTEYVLFPQIKTIIDTIANARNCALVTFFNRHTNGYMRPYMTPPNPKLHAMKLSYSHLNIIALTTKGSPLERPIARVQARAEAAGLLARWRRAEYEREDRDDAKRLVSLQGPRPLTLAQLKPAFVVLGVAHAVAVVVFALEALCAWRACCTARKMVRGGCIGRDIAQKVRPIFTD

>FoccIr181

MEVVLVVLLLCCAAPGVRADLPVVDSAAPPEARSAVALLTPFMAPQQATLIVMGTTRWTGAFLRELSADIPRVLAPSRPNWSSTALGRHYFSDTRVGRLQQALMATRSALLVATDDEQELFLAMEAYTPLRRNIDVRVLFWSTVSTSTSVHEVLRRMSVMKMWLAGRQHALALTAPNGSTTLYRMTCTSEHSCISLHTTILETDKWSTVEQRWRLGAAVFHDFCRGYRRIGKHQDLTMFVQTLKGSDNRSSLEELTKSVVNLASRGRRPVKYKKTSGHHELDTALEECTLAAVITDGVFLDLDRLTEISFLPDLELAEFAVIVPAGYGASVSVLDAVTVEFSPALWLATGLAILSTVVVFACAPRQDVSGAVLQALAPMLGQAPPPPAPPRPTLAAWLLACVVLTAAYQGLLLRKLSSAVPQRDLESLQDLEDSGLPLKAVGNLARTNVLTDNLNARVQHVFFSETKTVIDTIATTRNCALLVFHNRHTRSFMRPHMMPRRLHVIRLSYFHFNIIAVTTKGSPLERPLARAQVRAEAAGLLARWRRAEYEREYQDDVKRLASLRGPRPLTLWHLLPAFVVLGVGQVAGVVVCTLEVLWAWRARRA

>FoccIr182

MVALLVVLLLCCAAHGVRVVRASLPVVDSAAPPEARSAAALLTPFMTREKATLIVLGSSRWTEAFLRELSADIPRVLAPRPSRLTFARRQDNTPDTLQFDRLEHGLMATRLVHFVPTDDLQELLLAMRNYPEVNKLSDARVIFWTTLLLVSAPQDRVLRHIIRSNAWLGGRQHDLALTSYKGSTTLYNLTCIPSDACWTMDTTVYKMDKWSPVEERWRLGAAPFYEFCKGYRQIGTGQELTVYLGTGMGLAKTSSLMELTKSAVNLVGQGRHRRMRSQTTVKYQQISRHDGVGPPLRECNLAAFLSDEVFFTRHETIEVTFLHDLDLAEFVVIVPAGYGASVGVLDAITVEFCPALWLATGLATLCTVLVLKFARRQDVGGAVLQALAPMLGQAAPPPAPPRPMLAAWLLACVVLTAAYQGMLLGKLSSAVPRRDLDSLQDLEDSGLPVKAQGYLAFSNMLTDNLNARTEHVPNSEVKTIIDTIATARNCALVTVDNRHTLGYMRPYMVPKKLHAIRLSYFHVEILAVTTKGSPLEGPLATAQGRVEAAGLLARWRRAEYEREYRDDAKRLVSLRGPKSMTLWHLQPAFVVLGAGHMAGVVAFALEAMCAWRARWAGRTTR

>FoccIr183N

ALTSFNGSTTLYNLTCTPSDACLRMDASVFEIDKWSPVQQHWRLRANPFYEFCSSHRSKVKHQNLTVYLDTRTGLTNTSLLMELTKVAVNLVSRGRNRRIESQTSVNYQLTWTSHSIILALKECTLGAFLSDEEFFTRDETSDFSFLLDMKMAGFIVIVPAGYGASVGVLDAVTVEFSQALWLATALATLCTVLVLKFALRQDASGAVLQALAPMLGQSPPPPAPPRPMLAAWLLACVVLTAAYQGLLLGRLSSAVPRRDLESLQDLEDSGLPVKARADLAYNNMLTDNLNARTELVPKSEIKTVIDTIATARDCALLAVNNRHTYSSMRPYILPKKLHAIRLSYSHVEILAVTTKGSPLEGPLATAQGRVEAAGLLARWRRAEYEREYRDDAKRLVSLRGPKSMTLWHLQPAFVVLGAGHMAGVVAFALEAMCAWRARWAGRTTE

>FoccIr184C

MEALLVAFLLCCAAPGARAGLPVVDSVAPPEARSAAALLTPFMSSQKATLVVRGTSCWTGAFLRKLSADIPRLLAPKPSSGFTALKQNNYSDTLHLDRLEHGLIATRLVHFVPTDDLQELLYAMRNYPEINKLTDARVLFWTTLVTVSAPQDRVLRHIIRLNKWLGLHQRTLALTFFNGSSILYNLTCTPTDDCWRMDATVFEMDRWSPVEHRWRLGAALFYEFCNGYRKIGKPQELTVFMEKTLGLTNTSSLMELAKSAVNLASQSRHRRMQSQTPVNYQQILRYEHVGLKIRECTLAALLSDDVFFTRHETVEVTFL

>FoccIr185C

MEARLVVLLLCCATPGVQAELPVVDSAAPPEARSAAALLTAFMAPQKATLILRGSSQWTGAFLRELSADIPRVLGPRPSHLPSAWRHDKYPDTLQVERLEHVLMATRRVHFVPTDDLQELLYAMRNYPVINKVKDIRVLFWTTLLTVSAHQDRVLRQIFRLNTWLGGRQHALALTSFNGSTTLYNLTCTPTDACLRMNTTVFEIDKWSPDQQHWRLRANPFYKFCSSHRSMGKHQDLTVYLDTRTGLTNTSSLMELTKVAVNLVSRGRNRSGGSQTSVNYQLTWTSHSIILALKECTLGAFLSDEEFFTRDETSDFTFLLDMKMAGFIVIVPAGYGASVGVLDAVTVEFSQALWLATALATLCTVLVLKFALRQDASGAVLQALAPMLGQSPPPPAPPRPMLAAWLLACVVLTAAYQGLLLGKLSSAVPRRELESLQDLEDSGLPVKARAEMAFNNMLTDNLNARTELVPKSGIKTVIDTIATARNCALVAVNNRHTYSYMRPYMVPKKLHAIRLSYSHFEILAVTTKGSPLEGPLATAQGRVEAAGLLARWRRAEYEREYRDDAKRLV

>FoccIr186N

PALWLATGLATLCTVLVLKFAGRQDISGAVLQALAPMLGQAPPPPAPPRPMLAAWLLACVVLTAAYQGLLLGKLSSAVPRRELESLQDLEDSGLTVIAKGYLAYSNTLTDNLNARTEHVPDSQTKTVIDTIATARNCALVTANNRHTLSYMRPYMVPKRLHAIRLSYSQVEILALTTKGSPLEGPLATAQARAEAAGLMARWRQVEYEREYQDYVKRLVSMRGPKSMTLWHLQPAFVVLGAGHMAGVVAFALEAMFAWRARWAGRKTG

>FoccIr187C

MEALLVAFLLCCAAPGARAGLPVVDSVAPPEARSAAALLTPFMSSQKATLVVRGTSCWTGAFLRKLSADIPRLLAPKPSSGFTALKQNNYSDTLHLDRLEHGLIATRLVHFVPTDDLQELLYAMRNYPEINKLTDARVLFWTTLVTVSAPQDRVLRHIIRLNKWLGLHQRTLALTFFNGSSILYNLTCTPTDDCWRMDATVFEMDRWSPVEHRWRLGAALFYEFCNGYRKIGKPQELTVFMEKTLGLTNTSSLMELAKSAVNLASQSRHRRMQSQTPVNYQQILRYEHVGLKIRECTLAALLSDDVFFTRHETVEVTFL

>FoccIr188N

LAPMLGQAPPPPAPPRPMLAAWLLACVVLTAAYQGMLLGKLSSAVPRRELESLKDLENSGLPVKAKGYLAYNDMLTDNLNTRTELVPDTEIKTVIDTIATARNCALVTVNNRHTLSYMRPYMVPKKLHAIRLSYSQVEILAVTTKGSPLEGPLATAQGRVEAAGLLGRWRRAEYEREYQDDAKRLVSLRGPKSMTLWHLQPAFVVLGAGHMAGVVAFALEAMCAWCARWAGRTIE

>FoccIr189

MEARLVVLLLCCAAPGVRAGLPVVDSAAPPEARSAAALLTPFMAPQKATLVVLGSSRWTGAFLRELSADIPRGLAPRPSSGSTALKQDNYSDTLHLNRLRHGLMATRLVHFVPADDVDELLYTLKTYPLSRRVTDIRILFWTTVSASAPQGEAEQRVCRQNAWLGGLGGLQRALALTYPNGSTTLYNLTRTFYDATRTTKTSAIEIDNWSVLEQRWRLGVNVFHEFCRGYRQTGKRQNLTVYLKTAHDLKKPSSLMELAKSALKLASRSHQGRMGWQTRVKYQPIWSYDTVSLALKECVCAAFLAEELFFSRYELAEISFIPDMELAEFAVFVPAGYGANVGVLDAVTVEFSPALWLATALATLCTVVVFACARREDVGGAVLQALAPMLGQAAPPPAPPRPMLAAWLLACVVLTAAYQGLLLGKLSSAVPRRELESLQDLEDSGLPLKARGNLAYGNVLTDNLNARIETVEDSDIKTKTIIDTIATARNCALVAFYTRRTYSYVRPHILPYKKLHFIRLSYFHLNIIAVTTKGSPLERPLVTAQARVEAAGLLSRWRRAEYEREYQDDAKGLAALPGPRCMTLWHLQPAFVVLGVGQVASVIAFAFEALWAWRRGGARTPERQDIAVSMP

>FoccIr190C

MEVLLVAFLLCSASPGAQAGLPVVDSAAPPEARSAAELLSPFMVSQKATLKLRGSSRWTGAFLRELSADIPRILAPRPSRGPTALKQNNYSGTLHLDRLEHGLIATRLVHFVPTDDLQELLYAMKNYPEINKLTDARVLFWTRLVTMPIPQDRVLRHIIRLYRWLGVHQHALALTFFNGSSTLYNLTCTPSDACWRMDATVFEMDRWSPVEHRWRLGAALFYEFCNGYREIGKPQELTVFIEKTIGLANTSSLMELTKSAVNLASQSRHRRMQSQTPVNYQQILRYEHVGLKIRECALAALLSDDVFFTRHETIEVTFLHDLDLAEFVVIVPAGYGASVGVLDAVTVEFSQALWLATGLATLCTVLVLKFAGRQDISGAVLQALAPMVGQASPPPAPPRPMLAAWLLACVVLTAAYQGLLLGK

>FoccIr191

MEALLVAFLLCCVAPGVRAGLPVVDSAAPPEARSAAALLTPFMAPQKANLVVRGTSRWTGAFLRELSADIPRVLGPRPSHLPSAWRHDNTPDTLHIDRLEHGLIATRLVHFVPTDDLQELLYAMKNYPVINKVKDIRVLFWTTLLTVSAHQARVLRQIFRLNTWLGGRQHALALTSFNGSTTLYNLTCTPSDACLRMDASVFEMDRWSPVQQHWRLRANPFYEFCSSHRSMDKHQDLTVYLDTRTGLTNTSSLMELTKVAVNLVSRGRNRRIGWQKSVNYQRTGTSHSIILALKECTLGAFLSDEEFFTRDDTSDFTFLLDMKMAGFVVIVPAGYGASVGVLDAVTVEFSPTLWLATGLATLCTVVVLKYTRRQDVSSAVLQALAPMLGQAPPPPAPPRPMLAAWLLACVVLTAAYQGLLLGKLSSAVPRRDLDSLQDLEDSGLPVKAREYLVYSDMLTDNLNARTEHIPFSEIKTVINTIATARNCALVTVNNRHTYSYMRPYMVPKKLHAIRLSYSQVEILAVTTKGSPLEGPLATAQGRVEAAGLLARWRRAEYEREYQDDAKRLVSLRGPKSMTLWHLQPAFVVLGAGNMAGVVAFALEAMWAWRARWAGRKESWVEVS

>FoccIr192N

APPRPMLAAWLLACVVLTAAYQGLLLGKLSSAVPRRELESLKDLEDSGLPVIAKGYLAYSNTLTDNLNARTEHVPDSQTKTVIDTIATARNCALVTANNRHTLSYMRPYMVPKRLHAIRLSYSQVEILAITTKGSPLEGPLATAQAHAEAAGLLARWRQVEYEREYQDDAKRLVSLRGPKSMTLWHLQPAFVVLGAGHAAGVVAFALEAVWAWRARWAGRKAG

>FoccIr193N

LAPMLGQAPPPPAPPRPMLAAWLLACVVLTAAYQGMLLGKLSSAVPRRELESLKDLENSGLPVKAKGYLAYNDMLTDNLNTRTELVPDTEIKTVIDTIATARNCALVTVNNRHTLSYMRPYMVPKKLHAIRLSYSQVEILAVTTMGSPLERPLATAQGRVEAAGLLARWRRAEYEREYRDDAKRLVSLRGPKSMTLWHLQPAFVVLGAGHMAGVVAFALEAMCAWRARWAGKTIE

>FoccIr194

MAFLLVVVLACCCPALVRADLPVVDMAVAPEARSAAALLSPFLESQGATLCVVGSTRWTAAFLRGLSAHVPRVQMPIRLVLDDDLHPRLQHTLSTSLNALLVHTEGPEELLAAMSGYGAFFLRRTLFWTSVSSPQEDGVLRRVSKVPLVAPWLGAYQDRLALTSPNGSTVLYRLDCRSFAACSDKEVTITMVDEWSPLQGWRRQAAVFNEFCSGWRPGRGGRLDVVLLPSRQLTSTSPIQELAKTVVRLATRLGRGGPRMINITESDYYRLLKGLEGCRLDAMLTDVNVMIRPTFREASFVNDMAMVEVVAIVPAGFGAGVGVLEAVTIEFTAGLWCATGLALLSTAAVFACARRRQDVSGAVLQALAPVLGQATPPPAPPRPMLAAWLLACVVLTAAYQGLLLGKLSSAVPRRDLESLQDLEDSGLPVHMYSVLFLRASRQRLLPVALVERTRILFHRDISTAIDTVATARNCAVVVFMDMNIENIIRPYTIPPKQVHVIRLNHSAVRAIGMATKGSPLERPLVRALARVEAAGLLARWRSAEHERERLFYARKLVSLQGPRPLTLWQLGPALVVLGAGQVSGAVVFALELLCARWWQARAAGRTTRPGLT

>FoccIr195

MAFLLAVVLACCCPALVRADLPLVDMAVAPEARSAAALLSPFMESQRANLVVLGSTRWTAAFLRELSAAVPRVQAPIRLVLDDGLQPRLKHTLSSSLSTVLVHTEGPEELLAAMREYFAFPVSRILFWTSVSSPYDGVLRRVYNDTLWLGAYQGRLALTSPNGSTALYRFDCRSFAACSNKKVTVTKVDEWSPLQGWRREAAVFTEFCSGWRPRPVRHLDVALLQSGELTSASRIQELAKTVVQLAARQGRGQRTTNIKAMDYDSLWKRLEECSLDAMLTDKPPIIKASSREVSFLNDGDMMHLVAIVPAGFGAGVNVLEAVTIEFSASLWCATGLALLSTAAVFACARRQDIGGAVLQALAPVLGQATPPPAPPRPQLAAWLLACVVLTAAYQGLLLGKLSSAVPRRNLDSLQDLEDSGLPVHIVSALYLRTTTQRLLPVALVSRTKNLFHKDFDKVIDTVATARNCAVVTFLDLDIENLIRPYTIPPKKLHLFRLSYSTVRAIGLVSKGSPLERPLLRALGRVEAAGLLGRWRGAEHERERLFHERKLVSLQGPRPLTLWQLGPALVVLGAGQAAGAVVFALELLSARWWSARAARRTPAPGLT

>FoccIr196N

MLDNGLDPRLQHALSTSPCALLVHTEGLEELLATMSEYGKIIASRTLFWTSVASRQDVVLRHVSLVVPWLGAYQDRLALTAPNGSTVLYKLDCHSFTACSNDEATVTEVDEWAPLQGWRRQIAVFTEFCSGWRPRPDRRLDVGLLPSKELTSTSPIQELAKSVVQLAAKQGPRGRTTTTNVTDYHSFVERLKECSLDAMLTDKPLLLKTSWREVSYINDMEKMHAVAIVPAGFGAGVNMLEAVTIEFCASLWWATGLALLSTAAVFACRRRQDVSGAVLQALAPVLGQATPPPAPPRPQLAAWLLACVVLTAAYQGLLLGKLSSAVPRRDLDSVRDLEDSGLLVYSLSVLFLKASALRLLPVGLAPRTRMLLHEDVKTVIDNVATARNCAVVTFQDLDIENYIRPYMIPPKKLHFFRLNFSTVRMIGLTTRGSPLERPLARSAARVEAAGLLARWRRAEHERERLFYARKLVSLQGPRPLTLWQLGPALVVLGAGQVASVVVFALEVLWAWCWSARAARRTPRPGLA

>FoccIr197

MEAALLALCLFCCAAPVVRAGLPVVDSAAAPAARSAAALLTPFMAPQKATLVVTGGTRWTGAFLRELSADIPRVLDSNAFAMSTRHRHRTDPRLDLVLQATHRAHLFHTEDLETLLSAVRTRDPTRLVRTIFWADVASPRGEVLRQVSQTRPWLGDMQYALALTAPDGSTALYNLTCTTEITCSRGEMVITETDSWSPVAQRWRRGGRVFHEFCGPAWRQTKVPTVFMIGRKSLITTSVLELTKSVIHLAGQGADAAVVYFTDTHSMSTALTECALGAALTDLPLLVIGRSDETTKIVEMGRLHVGVIVPAGLGPSVRFLETVTVEFSAMLWCATGLATLCTVVFFRCARTRDVGGAVLQALAPLLGQPPPPPAPPRPMLAAWLLACVVLTAAYQGLLLGRLSSAVPRRDLGSLREVKDSGLRLLVLGQLSVSIALSDSPTNRTQFVTSEQVEATIDVIATARNCALVTYLDHYAIQALRRHNLPLKKLHLIPLRHSAVKVIGVSTRGSPLERPMVTALARIEAAGLRAHWRRAEHARENARYARKLALLQGPRPLTLFQLGEAFVVLWVGHAVGFAAFALEALWAWHAHKGKVLPDRSSHQGR

>FoccIr198I

MEAALALCLLLCCMAPGVRAGLPVVDSAAAPAARSAAALLTPFMAPQKATLVVIGEARWTGAFLRELSADIPRVLDTNPFALSIRHRHRTWPRLELVLEGTHRAHLFHTEDLETLLSAMRTRDPTQLVRTIFWANVASPRAEVLRQVSQTGLWVGGTQYALALTAPDGSTDLYNLTCATEDACDHKQMTIIKTDLWSPVAKRWRRGATVFQEFCTAWRPSQAPTVFILPRKGSTTTLXITYLDHYALRAVRRHTLPPKKLHLIQLSYPIVKVIGMATRGSPLERPMVTALARVEAAGLGAHWCRLEHSREEAHYARRLVLLQGPRPLTLFQLGSAFLVLGVGHAAAVGVFALEALWAWRAHKIKVLPARPAH

>FoccIr199

MGVALLMLLLGFAAPGVHADLPVVHLPVDPEAASAVALLAPFLQAQKATLIVLNQTRWTNAFLRQLDADIPRVVSLSDGCEQLHGWCMNLRMVSELSTRRYVRLVQLATTGRPDELVFIINVYARSSKQVIRTILWTSVTSLQARDDVLRVVSHKNIWLGMGQIALALTAPDGSTVLYKLTCTTEASCRHRAITVIEIDRWSAVEQRWRQRAAVFTEFCSWDWQMGGGGRALNVSLMTASGFSYKSSLMELAKSVQRSVGQGQQQRSRPVLNVTYEEVNSILSVMLGLRACRLGALLTDMSVFPPTISLDLMHVFDGEVSRIVVIVPAGLGPGVNVLAAVTVEFSAGLWCATGLAMLCTVVVFACARRQGVSGALLQALAPMVGQAPPPPTPPRPMLAAWLLVCVVLTAAYQGLLLGELSSAVPRRDLDSLHDLKDSGLPVHVQSFLSIRLSSILPLLPNARTHPYVGYSDIKSFIDTVASARNCALVAYLDRHVRTAMTPYMLTPKKLHYFPLGYSDRGIIAMAPKGSPLERPISRVLARSEAAGLRARWRSEEYERERQRHARRLVSLQGPRVLSFWQLGPAFFVLGFGQAASVAVFVLEALWAWSWGVPQARAARTTR

>FoccIr200

MQLIVLLGLLGAVCAGAYASLTDTATARPEAKCMASLLSSILPHNKTCLVLHGDTRLVGPLLRELDGELEERQVLLLDPRTIYSHLWFQMRDATTVVVIAMSSTAQLMDYMSGVKQFPSVAAIILWTSGSSLDDVLVHATRGSVLWLCFWDVYIVVSAPDGTSFLYSPPQRRGCVVTWESLNALTEMVRCLLEDRPGSSTGNRRRQWHRVRRLCSAWRPLRQSKGPNRSVLTAIAIKPDGKNPGEVIITHFVSNLTNAMNRQRFVELHWSTKGTDIMAAVYNCNLSAAFLGHMAPVRIYQHIRYSLPILVPVVVIVPAGLGQRLSLLEAVTAEFTTELWIATATSLLFMTAAMAVAWTSLGQPPLAALAAASLQTLAPLLGQSPPGATAHRPLTAVWLLMSVVIAAAYQGLLLRELTGPQAEINTLEQLEQSGLNVQIEDHLFLTPELQFKPRLQSRTRYFAQPRFFKALQNVVETRNSAIICHTDVHLANGPSKGLHTFTLPGISHLMSSVIYTTGSPLENPIVGVVGSSAGGGLHQHYLKMAAGGLTVSTAGPNDTSDQAQPLSLTQVQPAFVLLAIGCCISALVFAFELSFYKRVAHRERSAPAFPFRH

>FoccIr201

MQPLVLLVLLGAVCAGADASLTDTATTARPEAKCMASLLSSILPHNKTCLVLHGDTRLVGPLLWELDGELEERQVLLLDPRIIQSHLWFQMRDTTTVVVIAISSSAQLIEYMAQVKQFPSVSAIILWTLESSLKDVLMHATRGTVLWLCNWDVYIVVSAPDGTSSQFSPAQRRGCVATWESLNELTEKDQCAPSGRLESRPGNRNGGRLWERHQVRRLCSAWRPPPESKESEGSALTVISLKPAWKNPVVENLCHLVSNLTNVVNRQRPVELHWSTRSGKDIQEAVHNCNLSAAFLGFLAPVGLYQHIRYSTPYMVHVVVIVPAGLGQRLSLLEAVTDEFTVELWIATASSLLFMTAAIAVAWTSLGRPPLPALAAASLQTLAPLLGQSPPGATAHRPLTAVWLLMSVVIAAAYQGLLLRELTGPQAEINTLEQLEQSGLTIQMEDHLFLTPRFHPRPGLESRVRYFAQPRLYTALQNVAEGRNSAIICHTDVHLTNGPSKGLHTFMISGFSHLMSNIIFTTGSPLENPFRAVLGLSSNGGLAQHYFNLGARGMDVFTANSNDTSDQAHPLRLTQVQPAFLLLTTGYCISALVFALEVAFHKWVTHKESSAPVLPFVL

>FoccIr202N

ALAPLLGQAPGTPPPAGNALRPLLGVWLLVCVVLVAAYQGLLLGKLSTAQPHGEINSLQDLGDSGLPVHVSWAVFNQLDGVLPENVRRKLVVHRFVNPIDFIVNRVVVDRNCALIVEMDDGVARGIRSWLTTTKSLHFFTTGASFLRINGLWSRGSPLGQAIAATLRGSLEGGLSQRFNALADFAERLTREKTMAALSPARPLTLRQMYPAFLCLLGGHILGAAVFVLGLLVSI

>FoccIr203

MGVALLMLLLGFAAPGVRADPPVVHLPVDPEAASAAALLAPFLQAQKATLIVLNQTRWTNAFLRQLDADIPRVVKLGPGCASAQGWCTNLRMASELSQQRYVRLAQLATTGRPDELVYIIKIYARFSKQVIRTILWTSVTSLRARDDVLRVVSHTKMWLGMSQLALALTAPDGSTVLYKLTCTSNDSCKHGAITVIEMDRWSAVEQRWSQRAAVFTEFCSWDWQMRGGGRALNVSLMTASGFSYKSSLMELAKSVQRSVGQGQQQRSRPVLNVTYQEENDIVSVMLGLRGCRQDALLTDMPVFAPTIAFDLMHVFDAEVTRVVVIVPAGLGPGVNVLAAVTVEFSAGLWCATGLAMLCTVVVFACARRQGVSGALLQALAPMVGQAPPPPTPPRPMLAAWLLVCVVLTAAYQGLLLGELSSAVPRRDLDSLRDLKNSGLTVHIQPFLSIRLSSILPLLPNARTNPYVGYSDIKSFIDTVASARNCALVAFLDRHVRTAMTPYMLTPKRLHYFPLGYSDRGIIAMAPRGSPLERPISRVLARSEAAGLRARWRSEEYERERQRHARRLVSLQGPRALSFWQLGPAFVVLSFGQAASVAVFVLEALWAWSPAWCPRYGPPVAS

>FoccIr204

MGVALLMLLLGFAAPGVRAVLPIVHLPVDPEAASAVALLAPFLQAQKATLIVLNRTRWTDAFLRQLDADIPLVVKSGPGCASVQGWCTNLRMASEFSQQRYVRLVQLATTGRPDELVYIIKIYARFSKQVIRTILWTSVTSLRARDDVLRVVSHTKIWLGLSQLALALTAPDGSTVLYKLTCTSNDSCKHRAITVIEMDRWSAVKQRWSQRAAVFTEFCSWDWQMKGGGRALNVSLMTASGFSYKSSLMELAKSVQRSVGQNQQQRSRPVLNVTYQEENNIVSVMLGLRECRQDALLTDMPVFAPTIAFDLMHVFDAEVTRVVVIVPAGLGPGVNVLAAVTVEFSAGLWCATGLAMLCTVVVFACARRQGVSGALLQALAPMVGQAPPPPTPPRPMLAAWLLVSVVLTAAYQGLLLGELSSAVPRRDLDSLRDLEDSGLPVHVPSFLSIRLSSILPLLPNARTDSYVGYSDIKSFIDTVASARNCALVAFLDRHVLTAMTPYMLTPKKLHYFPLGYSDRGIIAMVPKGSPLERPISRVLARSEAAGLRARWRSEEYERERQRHARRLVSLQGPRALSFWQLGPAFVVLSFGQAASVAVFVLEALWAWSPAWCPRYGPPVAS

>FoccIr205P

MQPLVLLVLLGAVCAGADASLTDTATARPEAKCIASLLSSILPHNKTCLVLHGDTRLVGPLLWELDGELEERQVLLLDPRIIQSHLWFQMRDTTTVVVIAMSSSAQLIEYMAQVKQFPSVSAIILWTLESSLKDVLVHATRGSVLWLCNWDVYIVVSAPDGTSSQFSPTQRRGCVATWESLNELTQKDQCAPRGRLESRPGDRNGGRLWERHRVRRLCSAWRPPPESKGSEGSALTVLSLKPAWKNPVVENLCHLVSNLTNVVNRQRPVELHWSTRSGKDIQEAVHNCNLSAAFLGFLAPVGLYQHIRYSTPYMVHVVVIVPAGLGQRLSLLEAVTDEFTVELWIATATSLLFMTAAMAVAWTGLGRPPLPALAAASLQTLAPLLGQSPPGGTAHRPLTAVWLLMSVVIAAAYQGLLLGELTGPQEEINTLEQLEQSGLTIQMEDHLFLTSRFRPRPGLESRVRYFAQPRLYTALQNVAEGRNAAIICHTDVHLANGPSKGLHTFMISGFSHLMANIIYTTGSPLENPFUAVLGLSANGGLAQHYFNLGARGMDVFTANSNDTSGQARPLRLTQVQPAFLLLTTGYCISALVFALEVAFHKWVTHKESSAPVLPFVL

>FoccIr206

MVALVVGMLGVLVQLGLLGLLGAGAASAVGHPVGKYPAEAKCVASFVSSLVPQHKSCLVVHGDVQLVGQLIQELDGERQVVLISPRLLCSQLGFDVRRMTSLFVVAMGSVAQLDKLAEIESIPQMSHILLWIRARSLSDVLPHFATRNMPLWLCFANLYILLSTPNGTSILNSPDMTDRCMTTGRLLRMKEIDRCEPGARGWRRKPWTPVCDGWSQPPRNRSSPVQVFAIRPDYKLGNVNVDEFHHFVGNMTSLLSRRRSVQVLWTSPFSFELQNALRTCRMGAVFLSYSGPVVRNPNTQHISYHFASVVAVVPSGLGQRFGLLRTVTAEFSLALWVTTVLSSLSMTLTVAAAWMCSGRTPTSALAVASLQTLAPLLGQPPQGRTAHRPLSAVWLLMSVVLAAAYQGLLLRELTGPEAEINSLDELEQSNLDVFVEDALFDVVNSRFTSNMKSRTHYFPQEDFITHLHAMAIQRKSAMLYHRDVYFDLVTLPQIAGPPKRLHAFELLDSSHLMTRILFTKGSPLERPLRRALGIFDNGALAKHYFHLAGNAFECLDLTPNSTRELTRPLSLDHLKPAFVMLALGHILSALVFVCEALVYKWSRSQAPPVPVFTL

>FoccIr207

MVGEILRNSPYATNPSDFPLVDACLVAVLLCCAAPVVRAGLPVVDSAAPPEARSAAALLSPFMASQKATLILRGSSRWTGAFLRELSADIPRVLVPRPRRGSTALGKDKYPDTLQHDRLEHGLMSTCLVHFFPADDVDELLYTLRTYPASKRGSDLRVLFWATRSASASRDKVEQWLDVLVGGHQRALALTYPNGSTTLYNLPLTFPYASRGTKTRAFEMDNWSPVEQRWRLGVTVFHEYCRGYRQTGKRQDLTVYLISAIGLTQRSSLVELTKSALNLAGRGHQGRLISETPVLYQLPGVWNSSVFLALKECTLAALLSDELFFSQNEVGESSFIPDTKMAEFAVIVPAGYGATVGVLDAVTVEFSPALWLATALATLSTVVVFACARRQDVSGAVLRALAPMLGQAPPPPAPPRPMLAAWLLACVVLTAAYQGLLLGKLFSAAPRRELNSLKDLEDSGLPLKSRGNLGHSNLLTDNLNARIQSVRYSEIKTVIDTIATARNCALVTFYNRHTYSYMRPHMMPKKVHFFRLSSFHFNMIAVTTKGSPLEGPIATVQGRVEAAGLLARWRSAEYEREHQEDSKRLLSLRGPKSMTLWHLQPAFVVLGAGHTAAVVAFALEAVCAWRARWAARKTG

>FoccIr208

MEACLVAFLLCCAALVVRAGLPVVDSAAPPEARSAAALLTPFMAREKATLIVLGSSRWTGAFLRELSADIPRVLAPRPNRGSTALGTDKNPDTLQHDRLEHGLMATRRVHFVPTDDVDELLCTLKTYPPSRRDKDIRVLFWTTVSASAPQGEVEQRVPRLNAWLGGHQRALALTYHNGSTTLYNLTYTSYVASWGTKTSAFEMDNWSPVEQRWRLGVTVFHEYCRCYRQTGKRQDLTVYLITAKGLTQRSSLVELTKSALNLVGRGHQGRLISETPVLYQLTRSWNSSIFLALKECTLAALLADELFFSPNELAEMSFIPDMGMAEFAVIVPAGYGSNVGVLDAVTVEFSPALWLATGLATLSTVVVFACARRQDVGGAVLQALAPMLGQASPPPAPPKAMLAPWLLACVVLTAAYQGLLLGKLFSAVPRRELVSLQDLEYSGLPLRARGNLAHSNLLTDNLNARIKYVSYSEIKTVIDTIATARNCALVTFFNRHTYSLMRPHMMPKKVHFFRLSSFHFNTIAISTKGSPLERPIATVQARAEAAGLLSRWRRAEYEREYLDDAKRLVSLRGPRSMTLWHLQPAFVVLGVGHVAGVIAFAFEALWAWRRGSRDTGMPG

>FoccIr209N

CYHAARYATHPLIFVQVVVPSGLRRAPPLRSITNEFSTEMWLATAVAFLAVSLATVWVTWKIFGLPLTAALAAAPLQILGPLLGQSPPSAAAHRPLSAVWLLMSVVLAAAYQGLLLRELTTPPGEINSLEQLKQSGLEVRVSRDLYMYASHFPSAPWRSQKASVPHVSLDSVVRTMAEKRSFALVLQEDMHIDLLLKPYTTPPKRLHAFRVAFAYPSSQFSFSKGSPLAKSVEMLALRTLENGLESRMALVTSLRMHLPYGDDERRTRPLNLHHVWPAFFLLAQGLSLSALVFVCEAAYWKWINRKQRS

>FoccIr210C

MRATRLLAAVALLLCRRGHGVALPAPRPPVDSQGVADLVSTFLANTKAGVVLYGRARALDTLLEQLAPETPRIRAFTSNLTMASDRLCFELASTDNIYVIAGDSKAQITDTLSDSTIPLIGRLLLWTRAHRPEDVLALDLASYPWLTEKELALAVSTPDGSSFLYNIAIDLDLAPAYHYNAAEIDRWSPVTRRWQRQGSPFTRLCPRWRSPRTRGGDVAPLRANAIVPNRSLRLSEAYIEHVTALMNSLRPPVRVNWQVEGWGVYREEQNRIFNCTLPVFVTPRSLPIRCCYHALRYASHPLTAVQVVVPAGMGRASTL

>FoccIr211NCF

EDVTVRLRQALTMSRNLHLVVKRSGEELLDFLLEKPAAWKDKEVLLWTWAPSPQDVLALTRPGQIYICHVHIKLAVSTSNGTTFLFTVSSLRCLEKWQNLDMKENGRCSPGLLPWPKESLPEPLCVEWQPHGSDESTLRVMAFAPLREHRPVIQPFEKFIGNLVASVRPAVKLEWVPGNESSYEQVFAREANCNISTLLSFFPMTIRVEPHLQYQSGDLSPVVVVVPASLGPHPGLLKAVTAEFSVALWVATALAVGGITVALAVASTCRGRPRLTALAAAPLQALAPLLGQPPPGRTAHRPLSAVWLLMSVVLAAAYQGLLLRELTSPPGEINSLEQLEQSGLDVVILKDLREPANDFLPTSITSRARYIPWFHLHNTIENISEGHHSALICYWDMMTAYKLLSLLLPSKRVHTFSFPYYYPKAVAMFTKGSPLCNPLMSATGRALSGGLYDHLFSECLAAVRRRGHVSDPESGDQLTRPLSLAQLLPAFVVLAAGHLLG

>FoccIr212

MEVLAILAVLVLCSGGCDAVLEFRRPDPNVDTQGIPELLSSYLAADGNHGDILVVGSGRAVRALVEQLPPGTPRMAFSSTLVVTPDAREVYNNVDAAVAVVVMVGDAPEELMLRKYKFANPRYKRMLLWTRAAGPRDVLARLDAALPMWLCRHEVALAAPAANSTTVLFAVTSGACRAKRRVYNVTEVDRWSGGARRWQRQVLPFGRPCSRHPWGQGERARAPLKVIGVLPASNMSDRHPFRKFVTSVLGAVGRPVRVDWLSAVGDNLTRLRQDGKGACAFAPVFSFYSNPVDDVGREWVSSDYLMPVQVLMPTRMGRHPGLLQVMTSEFSVELWCATVLTLLAVAAAMAAAAVAILDRPPLAAVMLAPLQTLAPLLAQPPPGRIAHRPLSAVWLLMSVVLLAAYQGLLLRELTTPPGEINSLEQLEQSGLEVIVDDNLCMHAGYFLPEGLYRRVRFALPEDLPSAVRAVAEGRKSAVVLPRDVHAKALVMPYTNSKQLHAFEIKFQVLSSNIVIAKGWPLARQLSRVMHHTISTGVYQHTTKAMMGVYGRWHTVAAVSGPVTRPLTLQEMKPAFIALGFGNCLALFVFVLELAYQNLPTAARRFHPFRP

>FoccIr213

MQLFDQRLLGFVGLLSAAAHGALPAVDTSLPAGAQCMSDYLAAVLRPQSTLLIDGDVAPLFSYLRHLAPGTPKVLRTATDENLADMIARLRQTMTMSRTLQLVVKRSGKGLLDFVLERPAALGLRHRQVLLWTWAPSAQDVLKLTASSNIWMCGVLLNLAVTTPDGTTFLYSVTSPNCQEKWQDIHMKEISRCSPGLRLWPKGTLPEPMCLEWKPRGPGKSPIRVIASEPPPELGVVVGLYKEIVVGMVGRVQPVVQLDWIPHNTKNVYNIFAGAANCNIEAFMSFFPLLVNVEPHLQFEAVTLHSVLVVVPAGLDPHPSLLKAVTAEFSVALWVATALAVVGMTAALAVASACRGRPRLAALAGAPLQALAPLLAQAPPGRTAHRPLSAVWLLMSVVLAAAYQGMLLRELTSPPGEINSLEQLEQSGLDVLIVESLYKTGNDLLPASITSRAQYIPPNTLNAATKNMSEGRKSALICLFDMSAFYQMVPYVYAPAKQLHLFRLPYYNMKALSFFSKGSPIHEQLQSAFGLARSGGLFLHYNRAAMIKLRPRERGPDPGDQLTRPLSLDQLLPAFVVLAAGHLLGALVFALELLCRTWAEHGRPAGPPVRVFLH

>FoccIr214C

MQLLVQRLLGVLCLLAAAAHAALPAVDTGLPAGAQCVSDYLAAVLRPPNSTLVIHGDVVLVGPYLRDLAPGTPKVLRPPIDSDLTVRLRQTLSMSRNLHLVVKRSGEELLDCLLEVPSAWEYKQVLLWTWATSPRDVLELNRAEQMWMCRTHISLAVSTPNGTTFLYQAQELSCQGKWQDIKMKETGRCSPGLLFWSKESLPQPMCQKWEGRKSEKFDLRVIALAPPPQLSVIERPFKKVIKTLVATLRPAVKLAWIRPNMTIYEQVFAGASNCSIAALISFYPLTIHVEPHLQYEGDALSQAVVVVPAGLGPHPGLLKAVTAEFSVALWIATALAVVAMTAALAVASACRGRPRLAALAAAPLQALAPLLAQAPPGRTAHRPLSAVWLLMSVVLAAAYQGLLLRELTSPPGEINTLEQLEQSGLDVVIVDDLYWASKDLLPLAITSQANSISPFDVQKAIKNVSEGKKSALICYWDAISAYDLMPLLTGRKSVHTLRLPFYNVKASAMFT

>FoccIr215C

MQLLALRLLGVLGVLTAAAHAALPAVNTGLPAGAQCVSDYLAAALRSPNSTLVIHGDVDLVTRDLAPETPKVIRPPRDEDLNVRLRQTLTTSRTLNLVVKRSGKELLDFLMEVRDAWNFKQVLLWTWASSPHDVLALTQAQPLWMCRVRINLAVSNPNGTTVLYSATEMGCQDKWQDIESEENGRCSPGLLPWPKESLPEPLCLEWQPRRSGKSTTRVTALAPPQALTDVIRPYKKLVENLVASVQPAVELDWVSNNKTSYWQVFASE

>FoccIr216NC

VVKRSGEELLDCLLEVPSAWEYKQVLLWTWATSPRDVLELNRAEQMWMCRTHISLAVSTPNGTTFLYQAQELSCQGKWQDIKMKETGRCSPGLLFWSKESLPQTTCQKWRQHRSEKSAVRVIVLAPPPQLSVIERPFKKVIETLVATLRPAVKLAWIRPNMTIYEQVFAGASNCSIAALISFYPLTIHVEPHLQYQGDALSQAVVVVPAGLGPHPGLLKAVTAEFSVALWIATALAVVGMTAALAVASACRGRPRLAALAAAPLQALAPLLAQ

>FoccIr217P

MQLLVQRLPGVLGLLAAAAHAALPAVDTGLPAGVQCVSDYLAAVLRPPNSTLLILGDVALVGPYLRDLAPGTPKVLRPPIDADLTARLRQTLTMSRNLHLVVKRSGEELLDFLLEVPRAWKYKEVLLWTWATSPRDVLELNRAEQMWMCHTRLHLAVSTLNGTTFLYQAQELSCQGKWQDTKMKETGRCSPGLLFWSKESLPQTMCQKWRQHRSEKSAVRVIVLAPPPELIFIERPFRSLIKTLVANLRPAVKLVWITHNTTSNGQIFAGEANCSLAALISFYPLTIHVEPHLQYQSRPLSQAE

>FoccIr218NC

GLLFRSKESLTQPMCQKWEKHKSEKFDRRVFALAPPPQLSVIERPFKKVIETLVATLRPAVKLVWITHNVTIYDQVFAGEANCSLAALISFFPLTIHVEPHLQYQGDVLSQAVVVVPAGLGPHPGLLKAVTAEFSVALWVATALAVVGMTVALAVASACRGRPRLAALAAAPLQALAPLLAQAPPGRTAHRPLSAVWLLMSVVLAAAYQGLLLRELTSPPGEINSLEQLEQSGLEVVILEDLYWASKDYLPRAITSQANPISPFDLQKAIKNVSEGKKSALICYWDAISAYDLMPLLTGRKRVHTLRLPFYNVKASAMFTKGSPLYEPLIATTSRFFAGGLHNHLTSEYLAAVRPRERVPDPGDQL

>FoccIr219C

MQLLVQRLLGVLGLLAAAAHAALPAVDTGLPAGAQCVSDYLAAVLRPPNCTLVVDGDVTLLADYLRDLAPGTPKVLRPPADEDETVGLRQTLTKSRALYLVVKRRGEKLLDFLLEKPGIFHLKQVLLWTWAPSPQDVLALTPSSRIWVCGGVLLHLAISTPNGTTFLYYPPKQGCHEPWQDIKMKENGRCSPGLRSWSKGSLPEPLCLEWKQNGSRKSPVQVVALEPPPDLRVISDLYKKYIENQVEAVGSKVELAWVPTNKTMSQVFVDAANCNLAALMYFFPVTVGVEPHLQYDTLHFVEVAVVVPAX

>FoccIr220N

MGCQDKWQDIAREENGRCSPGLLPWPKESLPEPLCPEWQPRGSGKSTTRVIAFVPPQALTDVIRPYKKLIENLVASVQPAVELDWVSNNKTSYWQVFASEDHCNISALFSLYPMTIRVEPHLQYRGRPLSQAVVVVPAGVGPHPGLLKAVTAEFSVALWVATALAVVGMAAAFAVASACRGRPRLTALAAAPLQALAPLLAQAPPGRTAHRPLSAVWLLMSVVLAAAYQGLLLRELTSPPGQINSLEQLEQSGLEVVILEELHEAADDLLPAPMTSRARYIARYQLHEVIKNISEGRKSSLICYWDIISAHILIPLFKPPKRVHTFTLPYHDVKSTLMFTKGSPLYEPLISKTGRVYSGGLHSHLFSEFLAAVRREHVSVPDSGDQLTRPLSLDQLLPAFVVLAAGHLLGALVFALELLCRSRAEQCARPVRVILH

>FoccIr221

MQLFLGLVGVLCCAAGERATAALLAADTSPPEARCMAALLAALLPPSNATLLVTGDVGPIAPYLHELAPGTPRVVNRTPPADPRIRNQMSRGRILVLAVLPTAAELLEYMTEHAIRQPEQTLMWTWAPTPQDVLGLASEEPLWLCYVPLILAVTLPDGTTSLHTGVPKGCLATWPSLDMPQVDTCPAGSRYWRRGPPIPRLCTRWTPPSSPHIVYVEKSSPGLENFDNFYRGTISTIRPRVRMQYVRGNDLRDVSLNMFYCNLSGLILDEPVPIAAAAHHISSVWVVWSRIVVTVPAGRGPRGTLLQAVTAEFSAELWIASAVAFLAMTLALAAVSACLGRPRLAALAVALLETLAPLLAQPPPGRPAHRPLSSVWLLMSVVLAAAYQGLLLREITAPPTEINSLEQLEQSGLEVHVAENLIVGSIAADLSVLGKDRIHYIPRHRVRDVLRRVLDAQDAALICDLTFSLERRIEHMPLKERQRLNLFTLSTAHLVSHAATSTGSPLQRRLRTLYSRIQESFSPGAAEGAGLTPSSLRHSAPSRTRPLSLTHLLPAFVLLAVGHCVLCALAFALEVLCHRFGARRAPVQPLRRVAWE

>FoccIr222

MQLLLGLVGFLCCSAGERAAAALLVTHTSPPEVRCMAALLAALLPPYNATLLVTGDVGPIAPYLHELAPGTPRVVNRTPPADPRIRNQMSRGRILVLVVQPTAAELLEYMLKDAIRGPVLTLMWTWAPTPQDVLGFASEEPLWLCHVPLILAVALPDGTTSLHAGVPEGCLATWPSLNMPQVDTCSAGSWRRGPPIPRMCTRWTPPSSPPPHIVYAEKSSPGLNKNFDDFHRDTISTIRPRVRIQWVRGKSLRDVTLNMFYCNLSGLTIIEPMPLSTAAHHISSEWVVWTPVLVAVPAGRGPRATLLQAVTAEFSAELWIALAAAFLAMTLALAAASACLGRPRLAALAVALLQTLAPLLGQSPHGRPAHRPLSAVWLLMSVVLAAAYQGLLLREITAPPTDINSLEQLAQSGLEVHVSENLPIGSSAADFAVLRKDRIHYIPRHRVRDVLRRVLDAQDAALICDLTFSMASRIVDLPLKERQRLHLFTLSAAHLRSYAAASTGSPLQRRLRTLYSRVQASFSPGTAEGAGLTASSLRHSAPCRTRPLSLAHLLPAFVLLAVGHCVLSALAFALEVLCHRFSVGARRAPRQPLRRVAWE

>FoccIr223N

CCSASSASSAPGRSPPCGSRTRAPRLRSALVDSLLPPTNGSIAVFGGLRDVGDHVSELAPGRPRFLETNVTRLLSARSHARFELDKTILVGLVVFRTGEELVEYATAHQLPPSMRLVLWTWAPSPRTVLALAPDKPLWVCHAHVTLAVSMPNGTTTLHSAVTDGCQVKWQLLGMSPLDSCSTGASGWRRPNGHPTSDCSAWGPRPHNSPPVVYLERPDPPNEQYREFVGSLVSAMHPPVRTEWVSARVSYRVRSDLAACNLSGLVSYGSSTPIFLPHLRPEWRILTNVVVVVPVGRGPSFSLLKAVTAEFSAELWIASAAALLAMTLALAAASACLGRPRLPAALAVALLQTLAPLLGQPPQGRPAHRPLSAVWLLMSVVLAAAYQGLLLRELTAPPTEINSLEQLEQSGLEVLLSESLHPHQHVVVPTLHSGARFFPYNSTVAVLRRIADDQNSSLVGHFDQPLGNALVQLTREQRGRLHMFQLPGRHLRAYAWGSTKSPLVLKALHSVYLRSKQANLLAHYLHMKAVFLDQQNLGRDHSHIVRPLSLGQLQPAFVLLFVGSCVLSTIVFGLEVLCHRLTERHALARRAAASRARRLAWT

>FoccIr224

MQVLCGPFGVLCVFGVLGLHGTRATGIAPAVDPGSSPGSKCVADLLSALLSRSRSGIILMGAASHTGALLRELPPEVPRSLLFEPGLVGDRLEYAFNTKNDIFVVVRDGVVGANVLSFEFPLHIPPMARVVVWAHGLQVGAELTLLRDKQLMYCLPQQTALVTTTDLDTRLFTFPDATHCHLNDRLVRAVEVNRCPSVGQRWQKGELVVRKVCTEWQTVDGNDDSLEILVAAGRREWTDKTFDIWLTQVQRVPGHHVPVRYIHEDDEDHERIRDAFLNCRMAALMMGHTMRAFVSSELSYDAIDKYRLVVVVPVGRGPQPGLLRAVTSEFSAALWIATALSVPCMAAAMAAAAWVRGRHPSAMAAFLEALAPLLGQPPPGSTTHRPLSAVWLLMCVVVAAAYQGLLLSELTAPPPEIDNLEQLEDSGLRVYAASALFHDMSWVLPATLASRATFVHHKGYPDVLRETADGRNSAVIIHSDRYTRDLLWEWLKYPSPKLHTFWLPATQARAKRLYSKGSPLLRHARMVLRRMEAAGLLLKDSEYEGTNQCVVGDQPILPLALGELQPAFVLLAVGHCLGGLAFVTEVLYYRAYGRQEVVARGV

>FoccIr225

MQVLGLIGLLSTASCGLAPTVDLARRDGSNCIASYLSQLLTNSSAGIVIMGDDTYIAPILKQLPPGTPTSLLFDPGLVDDRLEYQFRTQDDIFLIAREVSADVLKFEFEPRVPIRARVLIWTHALSPEVDLTLARDKHLWYCPPQQVALAVSTFTGGTVLYHFQADALCDFNSSFVKAKEINRCTSNDQHWQNNNTVLRKACVDWKTSDKNSSIQILALRPDDETWNNFETWVMLIKRLIRRPVAVRQILVIDDEFARIYHSANYCRLATVVSSDITHGDVVMYGLMVLVPSGLGPQPGLLRAVTNEFSIELWIATVLSVLCVAAAMAVAMAAAESVAQHRWLVSLEDAFLEALAPLLSQPVGRATHRPLSTVWLLMCVVLAAAYQGLLLKQLTARMPEIANLQQLGDSGLSVIVQDELFQHADQVLTDKLMNRAKFENIYRFEWLLQELATKRNSAMIFYCDFYTAPKVSRWLALQRKTGPKRIHMFRLPGGRGRMSRSYPKGSPLQESGSFATQLMSASGLLQYHTLDRMQSDCGTASTLRPLCLREIQPAFLLLTFGYGISGLVFISEIWHYKVFVLRV

>FoccIr226

MQRVALLGFLCAGAQAVWPVPTTSHGPGPKCIVDLLSTLLSPANGSVIIIGDGRYTSSLVKELPPGIPRTVLVDHTVIGDWLEYRFNTLNAIFVIARESGSNTIEFDLRIPEAARVLVWIHAKSPESLLRLPRDKQLRYCAPQKVALAFGTVNESSALYSVTNQCDFNSPTITAKKVNSCKALAPKWEKSELVLPRLCSGWHHSPANSPLPITHAVTSETLVGTQYFMSLLAQALEVPVRMDPVFLLSDHLRVVEAIDECRMAALASIRPMVAPAASNTGYEGISTCAVVVLVPTGRGSRLHLLQAVTEEFSAGLWIATTLTVLCVAAAMAAALALGGRASLSLALLETVAPLLAQAPPSRPAHPTLSAVWLLTSVVITAAYQGLLLEALTTPIREINSLEELEKSDLLIKLDLDLLKDGRQLLPHTLVSRMEIVDMMNRQSIKEVLERRDSALVCDYDSYSRFVLAPWLESRQFHLFQLRAMKVKAHFRFTTGSPLEGHIRLALRRIEAAGLKYVSIFTVKKKCSADHLPLGLDDLQPAFLLLAFGLCASALSFAFELLYSSPKCSKLKSPTH

>FoccIr227

MKLVERVVLLGALVAGAAALARTALPVVSLRGSPGSESIVNILSTFLLHATGARPGIVVLGDNNHIGAILRQLPSETPMSVVFNDSLIDIRQDYHFHRTNNIFLVVRDSGLDDLLNFGHVPFRSRLLIWAHAQTSRDEFTIPNRRRMWVYRPREVAFALTSYDNNSTTLYTPTPVSEGDISTHYVTAKKINRWVPEDQRWESEEIVFSKSCSRWQAHHKNSSLTVRALTPSKGPARKTFKRIVNMMTRALGEMVVDVVWMTEESGKKMNSRRAATEQCTLDAFIMYRPFDVFMAPHVRFETHVFSHVVVVVPADAGLLLSPLQAVKNEFSVELWIAMALTVPCVVVATVVTAVARSERPSPVAAFMQTLAPLVAQPLPSEPAHRILLGSWLLMCVVLTAAYQGLLLRELTAPPPQIKNLDQLERSGLAIKVDRDMYQDASQFLPETLLSRATFVNLKNFEMSLQKLAHQRNSGIILSNDVYTSAVLLPWFKRSRKPQLHMFALTHERVKSHTEFSTGSPLELPMQKIIRRMEDAGIASFRKLPTEKNVANAPEHQARPLRFSELRPAFVLWYIGHCTGGLLFALEVLYDKYICKTVDY

>FoccIr228

MHVLLCLLCAAVARAALPVPDTGANAKCLADYLSRFLTPAKAGLVVMGDDRHVAVLLREVPAETPRSLVFNQSLLGDQLLYRLQATDDIFLVAREGGASVLDFDYQIPPAPRVVVWTHSLAAGRELALTSSRLGAFCLPTEVVLAVSSPDGDLGVYRLWSREGCTKRRRVIRARLVNRCSPRGLHWQGRDFVISKICTRQQTLREEPSLRIVALDDGHDVERFQTFVRRVARSLRRPISLQWLSLEHTWSIMDDATKCRLDAIFSIRPFKDYFAGHFFSYEMQDKCEVVVVVPTGAKPQAFLQAVTVEFSTESWVATGLAALGVAVTLAAALALGGRPALGAALLQAAAPLLAQPFPGRPAHRPLVAVWLLMSGVLVAAYQGLVLRALTAPPQEIDSLEQLEQSGLRVKVDVELLEDVGHLFTDTLASRLEFDDLLDANVMAKALVDGRNHSAVFCYDHYLRRALAPWLDRKRLHLFKLGAVSSKSHLMITAGSPLEESLRRVVQRARSARMSAVDEFGRYWRRSPPEAHMRPLSTYQLWPAFLLLIIGNALGAVGFVLEISLHPMFARERIVFPFIL

>FoccIr229

MTVTDSVLGPLLAIAVGVLATATHAALRSAAPVVPAESASVLRLLVTAMALNGTLFVDIDDVEWLDDSFLRRLPSEMPRVLVQLEHYWETSKDNWGADMNTSVILVARRRAGTLVDYYEYLFNNSTVPFTPLVSSRVLFLTLSVSPSTTIARLQSSWPCWSTGALVVANPDGAASLLRVEHVDCRPGDNSSSTIRPLDRWSPTFGRWEAGVNPLIPLSGCCGGFVRPPGYAPMMLLEHTAYSVNRAAFKRAQLVARLIGLVPKEVERNDLEMMNVPLWDSYDSLVCRSVVVFQSLATKGSSYNLVDDSFVTELSRVVYVVPSGLGARRHPLQALVGEFPPALWCLSLLAALGVAGVLSRSAGCAEAPALLRAVAPMLEQPLPSRPTRHPLLATWMVVCVVLAAAYRGLLLKMLGRPPRGEISSLDELRASGLPVKSSSYLHPSGCTSDDCESFFPWRRTAINAVAVDRSCGLFVHLDHVPPWMLATGGVHVIPTEMRSALTQFYMPRMSPLAETVRRVLARIGQAGLMRHWEAWDAEELRRIAGLRTPTGPRPLELHAVLPPLLLLVCGLGVAAGALAAEVVAHRLSRARGARQHGVLGAYRMDASARTVPPLYFTKG

>FoccIr230F

MSSPTVLLLAVGALAAGTNVALLPTDLLVPAEAASVASLLTRIMPHNGTLFVDSDVPWLDALLRRLPPGMPRVLLETRYPWDSYAWDEGMGDNVILLARDDPVNLLVDLARTAIPTNPLFSSRLLLVTSASSFQAVVEHVTLSWDCPRTAALVVAYPDGSAALLRVQQVDCSPRNASTTSTESIQLLDRWSPSSGLWEAEVNPLVPFSGYYRGFVRPPGYRPRMLVTAMAAKNRKISFYQRALFMAKMLNLIPIEITYDVFSRKRSNKDIVFHLWYRSDVVFLHGGLPAMPRAADYQAYVRSYEFLVTEISRVVYVVPAGLGARRHPLQALVAEYPPALWCLTLLAALSVAVVLTRSSACGPAEAPALLRALAPLLAQPPPGRRSHHPLLGAWMLACVVLAAAYQGLLLKMLQHPTVGEINSVEELLASGLPIKVSKYLELSRGYLNLSEAWGVFIPQILSEIKSVARGRKSSLIMHREHIPQWLVESRQLHLIDVPMESLVPQLFVPRMSPLAEPLRRGLMRVSEAGLLKHWEGAFTRRARKARVNAAQKLSKEAERAGPLRLEALRPPLLLLFFGQVVSVFVWAAEVAGARY

>FoccIr231F

MSSPTVLLLAVGALAAGTNVALLPTVPLVPTEAASVASLLTRVVPGNGTLFVDSDVTWLDALLRRLPPGMPRVLLGTSYSWEPFYWDGGMRNSVILLARGDPGDLPDDLGNETMPDSPIFHSRILLVASANSSQTVPWESLRTGAWVVAYPDGSAALLRVQRSDCFLSEPNPSAESPRLLDRWSPSLGVWEAGVDPLAPLIGRKRGYIRQPGDPPTMILIGYSDGYRRMASHKRALAIAQVAGLTPVYTNVEPANNDSSEMRKNISYIDCADVVFFEGGFFDADGLSITDELVVAQLSKLVHAVPVSAGARRHPLQALVAEYPPALWCLTLLAALSVAVVLTRSSACGPAEAPALLRALAPLLAQPPPGRRSHHPLLGAWMLACVVLAAAYQGLLLKMLGQPTVGEINSLKELRASGLPVIVSAYLQNHAKHYPFPKNCRFEYPFYSGVLELVAIERNCSYFLHLEHIPLWMLETHSLHLIETKVESILPVFTLPRMSPLGERVRSGLLWVSEAGLMKYWGLWDHNATLIRLVRSRIQSGPQPLPIRAALPPLLLLGFGLGASAVVLAAEVAADNLSHCFV

>FoccIr232

MALLLTVVVLATCAHAALVDHALDRVEALYHDPTEATSAASLVSRILAPRQGNLFIDYDVDWMAAFLQHLPPEMPRVLIGQHLQNPGGLLDLKSHMAMVMLARGDTVKAWAGTHTFFCPFSRILLVINAPSLKVAREMIKGAWRCDANAGVVVVYPYGDADLLRVYGETHRCAAHVERVDVWSPRLGRWEGGGNPFFAWWFGVPRRADAGAPRMLMEASVTNPAPFKRAVEVARLAGVEVVNVTQAVREEALANTADAIFFAAPFVPPPPDDVDAFFLTRLSEGVLVVPNGLGAPRHFMHALVAEFPSSLWCLTALALLCVSAVLARSPAEAPAPLRALAPLLAQPPPGRRAHHPLLMAWLLVCVVLVAAYQGVLLKMLAQPRRDEIHTVEEFMRTGLPVGASFNMYHLKCFSSMGPSVPAACFDDVTHAPAAALTRIAFSGGSGLFVQLESIPLELLRNKSVHLIPVRARALRSQLFLPRRSPLAERFRRALARVEAAGLMQRWDDWDEKEVRRYMKQKPPTEAVRLRVLTLLPAVLVLACGLGLSAIVLVAEVAAGVLHVDWRNCPPCRLRPLSFLFLSSDENCPAVTA

>FoccIr233

MVLLLALVVAGAHAALVHHSLERLGALNSSPTEVASSVSLVARILAPRNGNLFVYYEHQQWLGAFIRQLPSETPRALIGVRALRNSDELPGSWLGLNEHVDMTLLARSDPKVLLTTLVTVRQFFSPFSRILLVTTAASNQTARNIIYSNWWCDANVAVVVGYPDGTAELLRVRSKSSACWAKIEGVDSWSQSLGRWEVGVNPFFAWCFGHSPHRTNPPSMLVETSVVNPSRFRRALEVARLVGLEVVNTSKVKNEEAKLCRADALSYGLPIAAAQPVEVDGFFTTTLSAGVLAVPAGMGAPRRSLVAEYPPALCCLTALAVLGVSAALAHSPAEAPTLLRALAPLLSQPPPGRPTRQPLLLAWLLVCVVLAAAYQGLLLKMLTRRVRGNVDSVEEFLLQQNGLPIRASFDMYHLKCVKYRELPMNISIVKTRFLLHAVMDAAFNRNLSVFTQRNLIPLWLLETERLHLIPAEQQAVRSQFFLPRRSPLAERFRLALARISGAGLVKHWEIWEDREYRRLKMSEDRREPVALDVRTVVSALAVLGGGLGMAVVALVAEVAARGLSHRHVDGDVGTSSVDVLRPRRRRGSPSWRWTQWRPPLSSGAAAARSLGRSTPLLASPLCRSDRLVKSV

>FoccIr234

MALLLAVGALAVVAHAALLDHALERLDAVHRVPTEAACTASLLARLVAPHNGNLFVYHTRQMWLNNLLRQLSFQIPRVLMDSNFDWERNKRVSQLNKFVDVDLIVRDEPNALLRTLASGRQSHAFLDRTLLVTGAASNHTVKELLRSRWPVAVLALSLVVAYPDGAADLLRITSVTCTCTAYIEQKDRWSPGLGWKENAAPFVIWCDRWVMPGGGGTGSPRPRILMETEGVNPTRVKGALEVARQAGMEVATVNRSVFPIKPEDDKRICQMDVVFFGIPFLVGRPSEIDGFFVTSLSQEVLALPKGLGTNHFMRALVVEFPPVMWGLTLLGALGVAAALACSPAAAPTLLRAVAPLLGQPLPGRDTRHPLLAVWLLATVIIVAAYQSQLLKMMRRPHRSDINSVEEVWESGLPVTSGLGMYHLTCIKYPGTTGSGCIPTRKLIPTIIEMALYGNFGIFFYLEHVPRWLVELGRVHLVTGEFSHQTVRAHFFMPRASPLAERFRVLLARIDGAGLEKHWDIWQERVFNLVRMSIVDTTVPIRLRVPHILPALVVLGCGLCLCAAVLTAEFVVHRFRDLSPGRAVD

>FoccIr235P

MALVLFVAVLAAASTRAALLPTSGAVPPAADAEAVSAAALLSGTLAPRNGTLFLTGDTWWLDDFLRRLPRETPRALLPANFSWEGALRLHADIVPSVLMVAADGAESLVKTLASLSDLPVNSHVLMVATAPSHQAVTDRLLSIWRCEIFGYLVLVAPSGAAELLTVRRLRCPVKHRPQARDTVQLVDRWSPGRSWKVNPFLGWCLVARTPAELSIDKVLSIDWPVMGAAVPTKSTQDQGCXRIAIHFFCIVNNPAQDAMLDALQCRSQAAFLSYPLVPGPTRDLDSFFMTRISEVAILVPSRRGPYVYPLRAVFAEFPPALWGLILAAVLGVAAVLARSEAPALLRTLAPLVAQPPPGRDTRHPLLGAWLFVCVVLAASYQGLLLEMMSDTGRREINNLEQLKGSGLCLLAHDFMHRQVEAWAGSTFVIYDDLDVAMRRVGVDRDCALVTLIQYVPTAALESGLYHVLPLDLRGMATHLHVVRGSPFIEQFHLNLARSAAAGLLHLREWEVRGPNTTAPSGPRVLDFRDVKPAFILLGVGHALAAAVLAAEMVVAGIAAHRGHGPGTTAPALAAAGVVRVEPRPRPARGARQRGQRRPRELRPWLP

>FoccIr236P

HTNPRASECPCKRKDDKRRLLCTAEALSAAVLSLLSNASAIFVAGSAPWLDVFCARIPGTATLFKDGPGAHVEWAMASTRNVVVVVAETPAALGGQLLSTPLPLIGSVLLWTVSTASSTAAALSQAARKLLQVQAHLALSLPDGAVALHAVRVLRLEALRAELEELDRWSGGRWLRGPHDLAPCDSWRPWDPSHPNPSALLLHVDNELQDSVARKFLDAAARTSRFRIVDFEKPRTDRDYAVRETLIIADIARCGLDVFTCGSSFAVAPSYQVWSYAWMLHRMYVVVPAGLGGGRRPRHVAAGVWLCTVGAVLCVAAVLAVSSRLRSGLAARCPLLQALTPLLAQGWPAPAAHRPLYGVWLLACVVLCAGLQAQMLAELSFEEPPGEINSLTELASAGLQLGMPVYDRASLSGLHHGKVFLFYHPYWYMVDLMSKHRNISLLVKQDVLPFIGTHSRDRKLLHVFEVPIIVHVTHVAFSRGSPLATPFRRTYGRLLAAGILSHWGHVQRANGSEGLQSGSCLSCATVS

>FoccIr237P

MLPRLAAVVLLLVRVRCGARLPPVEVQPEATPCGLRPLLEHLAAGASVDLHVDGDAPWLRTLLGGWGTRTAVLSTRRAPGERRSRTLVPQFSQSKLRCLTPLPLLTVRSAFSAPASSVVLVAFDRPGDLVREARVALVRHVRTHILLWTRADQEDAAAGAATALRLADENLPMTTNQVRLAVTANTSTALFGGVWDVDYPNVTEVSTCSASTGWSREPGAAFDQPCLQWCPPPPDDPLVVYIVQSQVKNPGRFISAWTPEQRPRPPQRPAQRPXNNLPSVRPPSLPWEVLVAAFVKEFHRRQHLNFSYKAKVLPVNSEGLAMRSTLSCRLDLVLSEEPLNASLHRNLAVFPWDMHSTEAVVPPGLGTSSSPLGHLISELSPDLWATTMLAAAVVGRGLLWLGGRAPSTVALQVLSPLLLQTVAWPPRGRQRPLFGAWLLVSLVVSAAYQGQLLSDLAVGTRPNINSVQELVDSNLSVLFWKAYDLDSYPLEEWGLDSRVVLDTSADTRVASYWASRYANVSFFRDQEDKGTSAARGPVMFHTFSLPMTRLKATYTTMEGSPFEKPLRKFLGRLRAAGLTMESLLSLRSRGRLRGAKLPEGPKPLSLNQVQPALQLLGFCLAIAFVVFLLETTGGLCPSNYEPPSC

>FoccIr238

MEVWWCLLLAAAGCCGCCRATLPLISPAALEELSADPARCVLSTLAPSLLRAPGQRLLVVCNAHWCDGRFLGRLGVQKVVLAGEGLSVWTESRALQLTPSNVSMLLVATDSLEQLREVGPLLSVVMPQTRALLWTSTRATVTQTDLGDVVESVLNLTTTVPLGNVRLGVDTAKSTRLFAWTRSVMGDAGSLAHVDTWAPRVGRWSRGAALFPEPCTSWRPPAPGEPLTFLFHFHRIPDYRVNERIVKAHDVVLEFLRRTAGLVFKREAETLAEGSHLTRNLMTCRSDLLAFVTILESRMLETIDVSTWNLEELVVVLPAGAGRGRSPLHRLVVFSVSLWCATGVAVLVVVAVLCAPRPRGLVRAALQTVAPLLAQPPPFACNQYQRLLLASWLAASVVLAAAYQGQVLSRLTVPDAADEINSMEELAASGLAVYTRGDFRHLDLQLPNNTIVHNDFFGLIIDALLERRERVAIVVTELVAKHLVQHTRGPGRALHAFPVPGVRLLRSRFSTSRGSPVERPLRAAVGRMHAAGIYQHIDGEVGPLGCRDPAPPEEASPRALALGDVLPPFLVLAAGLLLAALAFLCELARPRLS

>FoccIr239P

MKLLRCLLLAAAAGCCSATLTIPAALEDPAGCVLSTLAPSLRLPGQRLLVVCNAHWCDARFLSRLDVQQVVVTGRDLSVWTESRALQLMPSNVSVLLVATDSLEQLRKVRPLMAEVMPQTRTLLWTSTRATVTHAELHVMMATVLNLTTTVPHRYVRLAVSTTKGTRLFAWTWGVGNVGSLTHVDTWAPRTCLWPRGAALFPVPCTSWRPPALGESLTYLFHVPRSPSLYITKMAVKERNIVLEYLRRTAGLDFKHQPEPLAKESEVGLKLIVDCRADLLTSVTILESRMLEIMDASVWYMESLVVVLPAGAGRGRTPLHRLVVFSVQLWCATGVAVLVVVTVLCAPRPRGLIRDALQTVAPLLTQPLPFAYNQYQRPLLASWLAVSVVLAAAYQGQVLSRLTVPDAADEINSMDELATXGLTIYTRGDFRLLGLPLLQNATVHNNDFLGFVVDALLERRQRVAIIAPNLIARHLVHHTRGPARLLHVFPVPGVRLLRPLYSTVRGSPVQRPLRALRLHAGGIVEHVHEAAIPLESRHVAPAAALRALALVDVLPPFLVLGTGLLLATLEFLCKLCSITTLVKITLHRDGIQACGASYSRGARRR

>FoccIr240P

MKLWLYLLLATAGCCCCKAALLLTIPAVLEDLDADLARCVLTTLAPSLRPRGQRLLVVCNAHWCNGRFLSRLGVQQVVITGEDLSMWTESRGLQLMPSNVSMLLVATDSLGQLGKVGPLIAEVLPQTRTLLWTSAQETLTHDKLDAVVGSLLSLXIVVPGIAKHLRQHFRGLGRLLHMFPVPGVRILRSCFSTVKDSPGGIVEHLQKVMIPRESQYAAPVAPLRALALPDVVPLFLALAVGLSLATIVFLCKLA

>FoccIr241N

QRPLLASWLAVSVVLAAAYQGQVLSRLTVPDAADEINSMEELAASGLTVYLRSDFSQLVLPLLKTATLHNNDLLEFVVHTLLKRRERVAIVILDNMAKYLLQHTRGPGRLLHMFPVPGVRILRSLFSTVKGSPVQRPLKTLMGRLRAGGIVDLLQRYIIPRKSQYAAPAAPLRALALADVVPPFLALAVGLSLATIVFLCEVARARLV

>FoccIr242N

AASGLTVYLRHDFRQLGLPLLKNATTLHDRDFMGFVVHTLIELRQPAAIIIPDLVARCLLHHTRGPGRLLHMFPVPGVRLLRPSYNTARGSPAERPLRALMGRLRAGGIVEHIQQEALPLESRHAAPAAPLRALALADVVPPFLALAVGLSMATILFLGELARPRLV

>FoccIr243N

GQVLSRLTVPDAADEINSMEELAASGLTVYLRRDFSQLVLPLLKNATLHNKDLLKFAVDALLERRERVAIIVPHFMAKYLRQHTRGPGRLLHMFPVPGVRILRSRFSTVKGSPVQRPLKTLMGRLHAGGIVDLLQKVMIPRESQYAAPADPLRALALTDVVPPFLALAAGLSLAMVVFLCELARPRLA

>FoccIr244

MTREETIFCVICLCLVAFREASCDAITLLASRVVRDPEVPCALTLLPPLFQRVDNDSAAYAPRLFVFGNADWLDEFLRRLPSDASIYVFSRVQESKEKLLFDDFFMAEMLPSSSVALFAMNPGQSQPIPQIPYRVRKVYWSSWDSNTEADDAMSNVTGFKLTDGFCPSFIRVMITARPRGTTHVFELSLSCELVYSRPQASAKLRGLWSPAAGWSASKELLFPPLCGSWRPPPDGEHLQAVIFRLSLPNGTVIEDDHPRTDEGNVLKMLKEYHGLVFETTFKSISDPGIPFRLADECRLDVLARPFPGHVLEAVEHTELSVFPWDFESVLLVVPTASGKPRSVLYPITAEYTPAVWAALGAVVLVVVSCLYLMRHDESVQELILQTLSPLLGQPLDDRRTGPQIAVLGGWLLTCTVVVAAYQGQLLGFITVPLENREIYSWQDWLESDLVLLAPNNFNVPLVNDVLGSSGLTEDRVVRNNTASLVQSLRSIAAQRKESRFLTQWEYYMVLYLTGEELRGKLHTFTVASGMLSRSTFFTTKGSPFEVVIRKFFGRVRAAGLIRGSKQQQATEDKKIPITLLNIKPIILIYIIGNFIALCCLILESLLRKAVCKPAKLKCSL

>FoccIr245

MSTGSAALAVALLCLAAGEASADAVTVLKSRVVRHPEVPCALDLIPPLFQRADNDSPAFPPRLFVVGKTDWLDEFLRRFPSVASIYVFTPTKESVEKFQPFFLAAMMPSASIVLRIGGAGHKISFPLRTRVIFWISSDSNIEADDELPRVPDIGHRFLRLMLTTQPRGTTHVFSFPMPLDMARTSSLVTGELLGVWSPAAGWSAPHVPIFPPLCASWQPPPNGQPLLAEMFEYYVNGTATLDNYTRTEAYEVLRLLRDSYGLVFQTKVGLTTEVLRPFSAADTCRLDVLAFPERALVVDHATHDYLSSFPWESDSSILLVPAGAGKPRPFLYPLTGEYTPAVWAALASVVLAVVACLYLLRRDESVQELVLQTLSPLLGQPLDDRRAGPQIAILGGWLLTCVVVVAAYQGQLLGFITVPPQNREINSWQDWLESDLVLLVENNYFKVSNAIQFGLYGLTEDRIVRHSINVGIETIVTRRNASLLIGKHNYDILIRNEEVKSGKKVQEVLTKIHTFVPPMIRRPHSSFVTSKGSPFEVPLRKIFGRIRAAGGFRYDRNHTNTLPQATEEKPISLLNAMPIFAAYIAGIILAVLTFLLEVFLPTLLNFKIIIVLRKPNGLQL

>FoccIr246

MTTGRTAVSAIFLFLAVRGASGNAVTVLKYRAERNPEVPCALTLIPPLFQRADNDSAAYAPRLFVLGKADWLDEFLRRFPSVAAIYVFTMTEENILKDFFGTEMMPSASVVLRLADATHVEVTVPVRVRLIHWTSFDSNAEADDVMSVLSVVITCKTFFRVMVTARPRGTTHVFSVPNSCDLTKPPFSRVEGELLGIWSRDAGWSAPLVPLFPPFCATWRPRPAGEPLLAEMFHLGINNDTAPVENYAHSEAYKILRLLKDSYGLVFQVQVNLTTDIAMPFRASESCHLDALAFPDIPLPLGVAFHTELSVFPWKFDKALCVVPVGAGKPRSVLYPITAEYTPAVWAALAAVVLAVVAFLYLLGRDESVKELVLQTLSPLLGQPLDDRRTGPQIAVLGGWLLTCVVVVGGYQGQLLGFITVPPQNGEINSWQDWLDSDLILLVEDRFQVSEAIALGYYGLTEDRIVRNTAANRIHRTINHRNVSYFTSEYAYEVIRNEWEAVSERDKEAIRQIHTFELPQSVLQSIFVTTKGSPFEVPLRKILGRIHAAGGLRKHWNYNEHPPEDTIDKPISLINLRPVIVAYIAGNIFAVFTFLLEFL

>FoccIr247N

VLTSRVVRHPEVPCALDLIPPLFQRADNDSAAFPPRLFVFGKTDWLDEFLRRFPSVASIYVFTPTKEIVEKFLPFFLAAMMPSASIVLRIGGPGHKIYFPLRTRFIIWISLDSNIKADNEMVNPRVPNLGYRFYRLMLTAQPRGTTHVFSFPMPFDMARTPSLATGELLGVWSPAAGWSAPHVPIFPPLCASWQPPPNGQPLLAEVFEYYVNGTATLDNYTRTEAYEVLRLLRDSYGLVFHTEFGLTTEVMKPFSAADTCRLDVLAFPQRALEVDHAPHDYLSYFAWEFDYGICVVPVNAGKPRPFLYPLTAEFTPAVWAALGAVVLAVVACLYLMRRDESVQELVLQALSPLLGQPLDDRRAGPQIAVLGGWLLTCVVVVAGYQGQLLGFITVPPENREINSWQDWLESDLVLWVENYFQVSEAIQAGLYGLTEDRIVRHSINAGIETIVTRRNASLLIGKQNYDILIRNEEAKLGKKVQEMLSKMHTFVLPMIRRPHSSFVTSKGSPFEKPLRKIFGRIRAAGGFRYDRNHNNTLPQATEEKPISLLNAMPIFAVYIAGIIFAVLTFLLEFFLPTLLYLKIIIVSRKPNGLQL

>FoccIr248

MPAKNVVLSAAILFVLLEEANLSAIAVLAERREQDPEVPCALALIPPLFELGDSDAAAHAPRLYVFGKARWINEFLRRFPSVASVYLFSGTEITTRLGDFITAGMMPSSAVALVLPDASSFLGYVPYRVRVLKWESFNLTSAGNRIYQMKRGDVERNGLCHRFLRYMVTTRPTYRYIQSSVPTNTTHVFAARISCDLIEVRPETTYELLGVWSPGTGWSSPGVSLFPPPCISWRPPPDGQPLLAEMLGYELEKYDSNNPFMVKYFKSTGAYEVLRLLRDKCGFDFETRFNFTTDTVAYPLRAAESCRLDIKGCSVLADSLSVLSHTELSVFPWDNHGFVCVVPAAAGRPYHIFYPITAEFPAAVWSSLAAVVIATVTCLYVLRRGKSVQQLILVTLAPLLAQPLDGKQAKSQRPLLGSWFLTCLVVVAAYQGRLLSFILNPPVSREIDSWQDWVESDLLLLVNHQFRVAMDAEGHFPGLTQDRLRVIVPTYDAMIAEVVTHRNVSFVADKYNFEIVKRNISQELRSKIHTFEMNTFHFASSCFATTKGSPLEVPLRKLLGRIRSAGGVPYRTLYKIKLQQTAEKAVEPDQVQNENKAIAIRNLRPVILIYVVGNLVSILSFLIERFRLLIVKYAIIVKIKMRSTTLYLVALCLLCSSTICLIIIFFRLFLTFLSRWLG

>FoccIr249

MSTGSTALAALFLCLAAEDALANAITVITSRVVRHPEVPCALTLIPPLFQRADNDSAVYPPRLFVFGTTDWLDEFLRRFPSVASIYLCTTTEENVEEPVDFLLAEMMPSASIVLRIGGDESPLRVPLRVRLINWVSFDSDDQADILMSELSVEDTCHKFFHVFVTARQRGTTHVFHIPMPCSLGVATDTIKRLVGVWTPGAGWPAPNVSLFPPLCASWRPPPNGAPLIADLSIYHPYRPTIRSFTGSEVSEVLRLLRHTYGLEFETRYNIALEEMGPLGAADACRLSLQAFPVRAQIRDVNFVIHAELSLFPWVLDDWICVVPAGAGKPRPVLYPLTAEYTPAVWAALLAAVLTVVACLYLMRHDESVQELFLQALSPLLGQPLDDRWAGPQVAVLGGWLLTCLVVVAGYKGELLRLIATQNGEINSWQDWLESDLDLLLKHRDQASEYIAKGLYGLTESRVLPSFFINEINTMPFAWTVCIERVAVGKDVSFIISKYIFDVLIRDKSVEEFLSQIHTFSVPIAPELKTSFVATKGSPFEAPLRKLLGLIQAAGGLNHHRNYKKNLPRDSKHKPFALTNVMPVFAAYIAGNVFAVSTLLLELLLPTLRPFWSYFFPRKPEKQTRSPAKRGCQLHYAGNSIRHGGAQSRVEPCVERAARASPVGGGRPSDFALRRSPIVSQELAELICSRERRRGTARR

>FoccIr250

MSTEKAVLCMICLCLGLPEASCDAITVLTSRVVRHPEVACALHLLPPLFQRVDNHSVIYAPRLYVFGNADWLDGFLRQLPSVASILLFRHQNVMQVEDFLVSEMLPSTSVALFVNRSETWYLSVVPIRVRMVLWYSRDSDTEADVAMSNLAELKARLCDRLVRVMITARPRGKTYVFSLSLSCNHQRSTGTAEPLGVWSATAGWSVPGVPLFPPPCLTWQPPPDGTPLLAETFTMDEIDLTKSLASEALGRLRDVYGLVFRTRHHVTEDQMFWLRAADACRLDIIAFPHRAPEVDVSKHTELSVFPWENDDMICIVPAEAGKPRHFLFTITAEYTPEVWAALGVVVLTLVVCLYLMRRDESVQEIVLQTLLPLLGQPMDRRRTRRQIPILGGWLLTCLVVVAGYQGQLLSFITVRYQNEEINSWQAWLESDLLLSVRYGSRLVDIVNMFGLYDLTEDRLIYKESLQEEIYTIATQGEISTLLFKQDYDLLMMDQPELRRRLHTFKIYHNGYPRSGFYTTKGSPFEIPLKKLLGRAHASGFSFDRTYYRVTQLERDTEVEHSPITLLNIKPIVSVYIIGNFIAVCVFIVEFLLHCVCKEVMKMLDICKGTLLGRKGLKAVPLEVMPEWYFDFGGGTYTGPACGPLRSVQPIPLRSHAGQRPLAIDVRVDPRENRLELRCTLT

>FoccIr251

MSTGSAALAVVLLCLAAGEASADVVTVLTSRVVRHPEVPCALDLIPPLFQRADNDSPAFPPRLFVFGRTDWLDEFLGRFPSVASIYVFIPTKEIVEKFIPFFLAAMMPSASIVLRIGGPGHKISFPIRTRFIIWISLDSNITDNDMVNLRIPNLGYRFYRLMLTAQPRGTTHVFSFPMPFDMARTPSLATGELLGVWSPAAGWSAPHVPIFPPLCASWQPPPNGQPLLAEVFEYYFDGMATLDNYTRTEASEVLRLLRDSYGLVFQTKVGITTELYRPFSAADTCRLDVLAFPQWAPDVDHASHDYLSTFPWEFDYGICVVPVNAGKPRPFLYPLTVEYTPAVWAALGAVVMAVVACLYLMRRDESVQELVFQTLLPLLGQPLDDRRAGPQIAVLGGWLLTCVVVVGAYQGQLLGFITVPPQNKEINSWQDWLESDLVLWVPNNFNVSEAIQSGFYGLTEDRIVRQTFKSDIETIVTRRKASFFIAKHNYDILIRNEAISGKKVQEVLTKIHSFVVPTIRRPHSRFVTSKGSPFEVPLRKIFGRIRAAGGFRYDRNHNKTLPQATEEKPISLLNAMPIFAAYIAGIIFAVLTFLLEVFLPTLSNFKIIYVPRKPNK

>FoccIr252

MSTEKTVLCAMCLCLAFRGASCNAITVLTSRAARHPEVPCALTLMPPLFQRVENVSAAYAPRLYVFGFAVWLDEFLRRFPSVASIYVFNDEDVERSEKMEAFFEATMLPSRSVVLFVEGPESWIYRSNQIKFLYNIPIRVRMIHWQSEDLTIKADVAINGFLQPLKHREHPLYCHIFVLVMITARQRGTTHVFKMSLFCDWYYSYSKATYDLLGYWSPAVGWSGSGVSLFPPLCLYWQPHPDGRPLLAEKIIQHHDQSLASRPELWQSKASETLGWLRGPYGLVFQTRVHVTSDKIYTARAAETCRLGFIGYRHTLSYIEAKTDTELVVFPWRARQPVIMVVPAAAGKPLPVLYSITAEYTPAVWAALAGAGLAVVACLYLMRSDKSVQELVFLTLSPLLGQPLDEGPGGPQIAVLGGWLMTCVVVSASYQGQLLSFITDPPKNREINSWEEWLESDLVLFLPDTIITLDGLKDLTLERISYTRSVPPQDQILNIANQRNGSRLMIKHDYDNFMMVQTEELRNKLHTFIMPNWGLVTTGYFTTKGSPFEVPLRRLLGRARAAGLSFDPDPSTNATKHLQQGTDQIFPITLLNIRPIISVYIVGNLIAVSFLMLEFFLSKITSKTSAN

>FoccIr253

MSTGNTLLAALFCCLTAGEASRNAIPIPTSGVARRPEVPCALNLIPPLFRREKNDSAAYPPRLIVFGISDWLDEFLQRFPSVATVFVFDPTPEHISKFEYLFLAEMMPSASIVLILYNDLDEESSLLSRRITIPVRVRLITWVSFDSNDDADVEMSYVTTRDTCHRFYRHMITARPRGTTHVFSLPMTCDMASESPDDDVEGELLGVWSPGAGWSAPRVPIFPPLCASWRGPLKGEPLRAEMFDITSGGLDSVDTYLSSDEYQVLKLVRESFGLKFRTRVKSTTKVTKPYWAADSCRLGVAAFPVDTIEVDIMYHTELSFFIWEIDKSIGVVHAGAGKPRHVMYPLTAEYTPVVWAALATVVLVVVACLYLMRRDKSVQELVLQTVSPLLGQPLDEGRGGPQIAILGGWLLTCVVVVGAYQGQLLGFITVPFQNREINSWEEWLESDLKLVVDNQTSLSSITNRGLYGVTEGRLFRIHQFAEEFEKPLGKTQTVVMRKRLFKDTVWDGTMHKKEVYFERLRELHTFDVPMTPSFPSRFFTTKGSPFEVPLRKMLGRIHAAGGFRSKLKYKEDPPQVTEDNPITLSNVMPVFAVYIAGNVLAGFTFFLECLQPNQIKFLGNTTHSRKALRSALSQIPYTCPACARVCSCERTLVRTVYSYRSFPDDILSTSRNSQRTPFLVFMS

>FoccIr254

MSTSNTALGALFLCLAAGGASGNAVRVLTSHVGYDFEVPCALDLIPPLFRRFDNDSASYAPRLFVIGNANWLNGFLRRFPSVATIFFGDFLLAAMMPSASIVLHIVGGITGLAVPARVRLISWLSSSTYNVDSEIDPPLVPDTCHTNMRVMVTTRPRGTTHVFSVPMPCSVEGGPSEVKSQLFGVWSPDAGWSAPHVALFPPLCASWRPPPNRGPPHAQMFLYGSIDDYHKIYFLSSKAFKVLQLLTLSYGSELKVTANTTTKGMTPFMAADTCRLDVLAFPSTPHFLNVLTHTELSVFPWELDSTICLVPLGAGKPRSVLYPITAEYTPAVWAALAAIVLAVVACMYLMGRDENVQELVLQTLSPLLGQPLDEGPGGPQIAVLGGWLMTCVVVVAGYQGQLLGFITVPPQNREINSWQDWLESDLVLIAHNRFALSDSIASGLYGLTEERIVRASFRNVIEVIATQRNCSACVSIHDYHELINGHRWTVLEDIQQKISKIHTFTLPKSQNHKATFFTTKGSPFEIPLRKILGRIQAAGLFRYNQNLNKFLWQATEDKPISILNLTPVFALYITGNILAMFIFLLEFLFAKSKKMLLNLRDKQQHKNTQTVIVPKKLYDRIL

>FoccIr255

MSTHNTALSALFLWLALGEASSNTVSVLTSRVERDHKPEVLCALTLIPPLFQRTSIDSATYAPRLFVFGKANWLGEFLRLFPSVASIYVFNPAAGNITKFDDFLLVKTTPSTTIVLNLTGGALIPVPFLVRYIFWESFDSNAKADDKIKRGLFDTCHRLFRWLVTARPRGTTHVFSMAMPCDYNIGSDRSKAELLGVWSPAAGWRVPLFPALCSSWRPPPDGEPLLAEMFEYANNSTATLYNYVRSEAYEVLRLLRDSYGLEFQIKFNSTSQYFAPLRAAETCRLDVLAFPAKILGIETESHSELSIFPWEDDYHIVVVPVGAGKPAPLLYPVTAEYTPAVWAALGAVVLVVVACLYLMRRDESVQELVLQTMLPLLGQPLDARRKGPQVAILGGWLLTCLVVVNGYQGQLLGFITVPPQNGEIDSWQDWLESDLLLLVNKNFDVSAAVLYGCYGLRENRILRAHSNINHLDIIATQKNASILISKYNYDILIRDKWAKSENNTEELLSKIHTFVAPLRPTPKSSFLTTKGSPFEVPLRKVLGRIRAAGGLRTKQIYRTHLLQTRKDKPISLINIMPVIAAFIAGNIFALFTFLFEFFLPYLK

>FoccIr256

MSSDMTALFALSLCLAAVEASGNAVTVLTSRMVRQRRPEVPCALTLIPPLFQRADNDSAVYPPRLFVFGTTDWLDEFLQRFPSIATIYVFTTFEDFFLSEMMPSSTIVLRIAVDQPFLVPRRMRYIAWASAEVEAEDEAKADDWMSDLSLRDPCNRFVRMMFTIRPRGTTHVFSVPTFCDYDAGDAGQPSMAKAELLGVWSPGAGWSAPHVPIFPPLCASWRPPPDGEPLLAEMFIFGGKATTVGDFTRSEAYEILRLLRDSSGLVFQTKFRAPPATMVLMPFIAAETCRLDVLAFPKNSYPPVGTEFHSELSLFPWGLDHSVCVVPAGAGKPRHVLYPITAEYTPAVWAALAAVVLTVVTCLYLMRHDESVQELVLQTLSPLLGQPLRDRWTGPQIAVLGGWLLTCLVVVGAYQGQLLSFITVPPQNREINSWQDWLESDLVLLGTNRFSISYAPDDVLYGLTEDRLVRSTAVLLKSAFEIIATQRNASFIISKHGFDIESSQLSEVEEEYLSRTHTFVVNTAMAQSNFFTTKGSPFEKPLRKLLGRIHAAGGLRQNPKYKKYLPQATDDVPISLINVMPVLGVYIAGNIFAVFTFLLEFFLPTERKYCCI

>FoccIr257

MSSEQTALFALFLCLAAVEASGNAVTVLTSRMVRQRRPEVPCALTLIPPLFQRADNDSAVYPPRLFVFGTTDWLDEFLQRFPSIATIYVFTTFEDFFLSEMMPSSTIVLRIAVDQPFLVPRRMRYIAWASAEVEAEDEAKADDWMSDLSLRDPCNRFVRMMFTIRPRGTTHVFSVPTFCDYDAGDAGQPSMAKAELLGVWSPGAGWSAPHVPIFPPLCASWRPPPDGEPLLAEMFIFGGKATTVGDFTRSEAYEILRLLRDSSGLVFQTKFRAPPATMVLMPFIAAETCRLDVLAFPKNSYPPVGTEFHSELSLFPWGLDHSVCVVPAGAGKPRHVLYPITAEYTPAVWAALAAVVLTVVTCLYLMRHDESVQELVLQTLSPLLGQPLRDRWTGPQIAVLGGWLLTCLVVVGAYQGQLLGFISVPPQNREINSWQDWLESDLVLLGTNRFSISYAPDDVLYGLTEDRLVRSTAVLLKSAFEIIATQRNASFIISKHGFDIESSQLSEVEEEYLSRTHTFVVNTAMAQSNFFTTKGSPFEKPLRKLLGRIHAAGGLRQNPKYKKYLPQATDDVPISLINVMPVLGVYIAGNIFAVFTFLLEFFLPTERKYCCI

>FoccIr258

MSTGSTVSAALFLSFAVGKASSNAIAVLTPRVDRYPEVPCALSLIPPLFQWDDNVTAAYPPRLFVLGRADWLDEFLRQFPSLALIYVFSTTREIRFEDFFLVEMIPSASIVLRLVDNSAPAWIPFPHRMRFIKWMSFDSDAEADDTITDLSVEETCHRFFRVMVTARSRGTTHVFAVPMPCKMAIEPLQAKGELLGVWSPGDGWSVPHEPLFPPHCVSWVPPPNGEPLLVEMYLYGDSESPSLDKYIKSEAFQVLWYLEDSYGLVFHIKFALAFELMMPFSRADTCRLDALAFPSISNGVDWMMHTELSIFPWRTSRLLCIVPVGAGKPRSMLYPITAEYSPEVWAALAAVVLTVVACLYLMRRDESVQKLVLETLSPLLGQPMDDSRPGPQIAVLGGWLLTCVVVVGAYLGQLLGFITVPSQNGEINSWQEWLESDLVLLASEDFSIKRLPDFGLTGLTEDRVVRIRFSPRSIIEIIATERNASFFIFDNEYKDTVLDTNHSEQMEEWLRNVRTFELPMLQILKSSFVTTKGSPFEVPLRKILGRIRAAGGFRKNWSHNQFPLLQQTENTPISLINLLPVFAVYIVGNLIAVFTFLLELFVNLKTFVKFICSENFKLTTCVNGDEQRQIETTRSGAGWVRKMRWGGWGRIRTDRDRETLRKWYP

>FoccIr259

MSTGDMALAALFLCLAAGEASGNAVSILTSRLVRHPEVPCALDLIPPLFQRADNDSAALAPLLLVVGNADWLSEFLRRLPSVASIFVFTDETTTSKFEDFLLAGMMPSASILLHVTDFADSVGSSITVPKRVRYTYWVSHVSDASADYVISHNLSPPDLCFTFFRLMITARPRGTTHVLSVPMPCDMASWSSPPKSELLGVWSRGAGWSAPHVPVFPPLCASWRPPPDGKPLLAEMFDYAIGTTSVDTYKGTETYEVLRLLRDSYGLGFQMTANVTTKALVPYRRADTCRLDVLALPTSPDVVDAQTHAELSVYPWDYADFICVVPTGAGKPRSVLYPITAEYTPAVWAALVAVVLAVVACLYLMRCDESVQELFFQTLSPLLGQLLDDRRPGPQIAVLGGWLLTCVVLIGAYQGQLLGFITFPSKNGEINSWQDWLESDLVLVVANKYITASSADDFGLYGLTEERIVRTTSGTRALFQVAIQRNTSCLITEREFERVRDTVAELREEYYGLFLNLHSFVVPTIPRPYSSFVTMIGSPFEVPLRNILGRIHAAGGLRQNRKYEKYFPQDTEDRPISLLNIKPIIVAYSVGNTFAVFTFLLEFSLPALRKFCWLTGVKIKTNIKPPRPYLP

>FoccIr260

MSTGNAVLSALLLCSAAAGASGSALEVIMSRAVLYSEVECALALIPPLFQRADNDSAAYPPRLFVFGKADWLDKFLRGFPSVATIYLFTAEEENIAKFKDFLLAEMMPSASIALRFVPDAGNHVTVTVPIRVRYITWTSFDSDLEANAQISRPTAWDSGNRFYLLMVTARPRGTTHVFYVPMLRNPFNVVPMWGELLGVWSPDAGWSAPHTALFPPLCASWRPPRDGEPMLAEMFLFSPTGIAKDRTLPVYMYSTTEEYEILRLLRDSYGLVFQTKAKVFTEAAKPYGATDTCRLDALAFINGTFAGLEVLTHSELSLFPWVLDNWICVVPAGAGKPRPVLYPLTAEYTPAVWAALLAVVLTVVACLYLVRRDESVQELVLQALSPLLGQPLEERRAGPQIAVLGGWLLTCVVVVAGYQGQLLGFITVPSQNGEINSWQEWLESDLVLLVGSAFSVADVVAQGLHGLTEDRVVREAPWNYLKVIATQRNASVLVSKYHSEVSIRDESNTNRFGIKEYLRKVHTFAPNLPQNLISSFVTSKGSPFEVPLRKILGRIHAAGGLRRNHNNHLPQATVEATVDSEDKPISLRNVMPVFAAYVIGNILAVITFLLEQFFYYRK

>FoccIr261

MTTDRTILSALFLCLAVGEASGNAVTVLSSRVARHPEVPCALTLISPHFQLSDNESPAYAPRLFVFGKAAWMDEFLRRLRSVATIYLFMPTEENLAKFNDFFMAMMMPSASIVLRLTDVGPYTSNTWAPSFPDRVPFILWVSYDSDAKADYAMSSINLVDTCRRFFRVMVTARPRGTTHVFSVPMPCDIARPQALAKAELLGVWSPGAGWSGPEPLLPPRCASWRPPPDGQPLLAEFFEYAGYPVSADYFRRTEAHEVLQLLNHSYGLVFEAEAKYTTQLMAPYSEAGKCRLDALVFRTRAALAVHTHTELSFFPWEHDSLLVFVPAGAGKPPPMYPITDEYTPAVWAALVAVVPAVVACLYLMRRDDTVQELVLQTLSPLLGQPFGGKRAGPRIAVLGGWLLTCLVVVAAYQAQLRGFITVPPQNGEINSWQDWLKSDLLLSTPDGIPVSSVIAMGGLTGLTEDRVIRSSSGMDLYNFFRAHHNVSVSMWKSDFENFIKYKSIQIKELSGVHLFAAPFLHLKSSFVTTKGSPFEVPLRKILGRIRAAGGLRHNWKYNQYPPQRTPYKPIPVINLTPVLAAYIAGNVFAVLTLLLEFSMRKVRICTLKILRARSLVET

>FoccIr262

MSTRRLLLSAAILCLTTGNICCNLVSVLTHRIEYHPEVPCALELIPPNFQLNENANATYAPRLYVFGKADWLNKFLWQFPCVASIYLFSDEENVLKDFIQAGMMPSASLALFLNSLELLPAAPFRVRTVSWLSFDSIVEADSYLSEFLAKPDESHEIESDISEETDAILRELSDLDDAIPDTLVVRRPGRCHRFARVMITTRPRGSTHVIALPMPCDLREEPKILYELLGVWSPGPGWSSPDVSLFPPPCVSWTPPPYGQPLVAEIFSFMKRYGNDDTVMGSDVKGTEAFEILRLLKDSYSLRFDIRINSTTDFYHPLRAADMCQLDIVAWERILIGASVSSHTELSLFTWKMYGTVCIVPVGAGKPYDKLYPVTAEYTVGVWIVLVGALIVVGTILYLMRRGESAQLIALLSLSPLLGQPMGDRQIGAQVPLVGSWLLTSVVVNGAYQGQLLSFITDPPLTREVSSWQDWLESDLLLKTTYQSTLEGLIQDGFYNLTADRVLADGNDFRQNIDEMLTHRKTSFCMTTYNYEREMNLYPKQITDMVNTFTLVTSMKVMSPYVTTKGSPFEVPLRKLLGRIRAAGGLRFHSKPLLQDIAATKGNEEEKENTVIAIRNIKPVILVYIWGNILAFLCFVKETFNCRILCSYCGSFCRKNIRVLLIMFALLTVISCCPSLYRHGL

>FoccIr263I

MSTGNTVISALFLCLTAGEVSGNAVTVIMSRDVRYPEIPCALALIPPLFQRADNDSVTHPPRLFVFGKSDWLDAFLRGFPSVATISVFTSERENILKGQLLGFITVPPENREINSWQDWLESDLVLLADDTLSVRSAIDLGLHGLTEERIDRTRPYNTLRVIATWRNASMLVTKHHYEMSIREKWIQPEIKDLLSRLHILKPVDRQYLRSGFITMKGSPFEVHLRKILGRIRAAGGHSYDRNFNIGPQATAESKNKPISLRNLMPVFAAYIVGNIFAVLAFSLEFSLISKKILPK

>FoccIr264C

MSTERTSLFALFLCLAAAEAASGNAVTVLTSRVARQRHPELPCALTLVPPLFQRADNDSAVYAPRLFVYGTTPWLDEFLRRFPSVATIYVFTTTGGKSEEEFEDFFLCHIMPSASIVLDITRFTVLDSTNLDRKQGLPTWQTRIRYIAWLSLTAEDEAEQALVDNWLTDLSFLDLCNLFFRVMVTVRPRGTTHVFSVPMLCDMDAAPSSRVKAELLGVWSPGAGWPSPDAPIFIFPPLCASWRPPPDGESLRFDMFIRSHVAPFADFRESEAYEVVRLLRERLVFQNWFTVTTMGVLKPLEAAGTCRLDAVAFPRIAFPPVSPWSHSEISTFPWMLDRCVCIVPAGAGKPHPLLYPITAEYTPAVWAALAAVVLVVVACLYLMRRDESVQELFLQTLSPLLGQPLDDRRPGPQKAVLGGWLLVCVVVVGAYQGQLLGFITVPPQNREINSWDEWLESDL

>FoccIr265NJC

VIMSRDVRYPEIPCALALIPPLFQRADNDSATHPPRLFVFGKSDWLDAFLSGFPSVATISVFTSERENILKFMDFFLSEMMPSASIALSFTHDPFTNWACIAVPGRVRFITWLSYDSDGEADALVSNATAPHLPFRFFLLMATARPRGTTHVFSVPMPYNLFNHVAPARGELLGVWSLSSAGWSAPQQAPLFPPLCASWRPPPDGQPLLAEIFEMSIYGFGDGFTIPVELYDRYVGTEEYEVLRLLRDSYGLVFQTTATVYAEALRPIGAADACRLDVVALPKEFDDGLYVSMHSELSFFPWALDHYVLIVPAGAGKQRPVWYPLTAEYTPTVWAALAAVVLTMAACLYLARRDQSVQQVGLQTLSPLLGQPLQGGRPGSQTAVLGGWLLMCVVVAGGYSWFTKF

>FoccIr266

MPSGRTVSSAILLCVVLVEARCTAIRGNSPLFSRLLSLFDVNATHETLCALALIPPHLQLAGNNASATHAPRLYVFGKTDWLSEFLQRFPTVASVYLFSGRYTTRFSDFIRAGMMPSYSIALVLDDAYDPMPLIPYRVRTVRWVSLTSQLNPLLRELKKQPVPPKTCHRYVRFAMTLRPAGLTHVYAGIYVCDYKRPTRLTLDAMGVWSSVTGWSSPAETLFPPPCLSWQPPTIDERPLTAEMFALASMRYDPNNKAMVRAVQETESYEILRVLQRSYGLAFDIEPEFSRDTYHPMRSADWCRLDLASCLGTTDRIDVLLHSELSVFAWEFDALLGVVPKGAGEPYHMFHSLTVEFTAGVWTVLGAAVLVVAACLHLARRDESVEHAVLVALAPLLGQPFDSRCGAGPLRPVLGGWLLTCVVVVAAYQGQLLGYLTDPPRNREINSWKDWLESDLFYIARGPIPMDSEFLSQWGVPGLTPERFRSGFTDEQMFVETATRRDCGFLYDKNLYEFAVTRLSKNISSRLNVFTRLEEFSLQTCFFTTKGSPFEVPLRKLLGRMWAAGLQNKMNASRKRQIYHNRENLTLSRVTRISSSNLAPVLWIYAVGNLLSFVSFSVEANVFKYPQAGALMHI

>FoccIr267

MPRQHAVPTVVLLCLVLTEVRCSVGTVNLSRFLRKGLSFDMNDTPEASCALALLPPNFQPGDHDSYSPRLYVFGKTVWLNEFLRRFPKVASVYLFDGSAAWEFADFVLTGMLPSSSVALVLDQSEQLLHIPYRVRVLYWEASVADTQLESEVRSLNARYYPPGACHRFVRIMVTTHPAGATHVFSLHLACDMFKSPAATLEPLGTWSPGTGWSTPTSAIFPEACVSWRPPPDGQPLTAVMMSVQDEVHGDLQEQPFKLGEGYNVLRVLRNSYGPAFEFKFNVTRNPYYPLRAADTCRLDVVACSISSNPGFKITVALHSQLSVFAWGLHDLIFVVPAGAGKPYHALHSITAEFAPAVWSAVGASTLVVAACFYLTRGQQSFQNVFLLTFAPLLGQPLSFQSVRTRTILGGWMLTCFVVVAGYQGKLLSFLRNPLRNREINSWQDWLESDLKLINQGPYGVLETEYLSLIGADGLTSDRFRFGFSYWDMLSITANGKKTSFFSDKADFDMVFNYVPKDVTNNLHVFPMAAGYSLSSCFVTTNGSPIEKPLRKLLGRLRAAGIHDKYRTIVHRTTNVTTEGKPITKYNLQPVMMVYLTGNLVALISFFLELFT

>FoccIr268

MPRQQALPIVLMFCLVFSEVRSNAAAANLRRSLEKGPSSDMSDTPETSCALALIPPNFQLGSHDAYTPKLYVFGQTVWLDEFLRRFPKVASVYLFEGRAAWKFSDFVLTGMLPSSSVALVLGRFDPLLCIPYRVRILYWWPPVAESQLESEVHSLKVQPYLPGTCHRFIRIMITTHPDGATHVFALHITCDMYDIPKATLEPMGAWSAQAGWSTLTSAIFPEACVTWRPPPDGQPLTAEMLSLAPEQYNESSAGLFQIGESYQILRDLQDLYGAVFEFKFNYTRDLFGPIRAADTCRLDLTACSVGSSTGFKITVALHSQLSVFAWGLHDLIFVVPAGAGKPYHALHSITAELTPAVWSAVGASTMAVAAFLYLTHGERCYQGALLLTLAPLLTQPLSFQSVRPRAVFGGWMLTCVVVVAAYQGKLLSFLRDPIRNREIDSWQDWLETDLSLMSNGPFGQQDTDYLSLIGATGLSHDRFRFGFSDWEMLSATAYGKNGSFFFDKSQFDGLLSYFPGAFMRQLHTFPFATSFSIYSCFLTSYGSPVEGPLRKLLGRMRAAGLHRKYRRIAFPLGNQERKPITTYNLKPVLWIYLAGNLVALFAFTVEMFIYVRT
