## Additional file 5 for "Genome-enabled insights into the biology of thrips as crop pests"

**Additional file 5: Neuropeptide and protein hormone peptide sequences**

**of *Frankliniella occidentalis* as part of the FOCC genome paper**

*Contributed by Jan Adrianus Veenstra and Yoonseong Park*

Canonical signal peptides are highlighted in green, parts of the precursors predicted for the mature neuropeptides are highlighted in yellow and convertase cleavage sites and Gly residues that are predicted to be transformed in C-terminal amides are underlined.

**1_AKH/Adipokinetic hormone**

MAAMQFSSRVVRVVASVLLVLCATCATLVTAQVNFSPSWGKRGAPQQEACKPSMDSLMYLYKLIQDEAQKLTDCEKFAN*

**2_Allatostatin C**

MAPSWTPLPSPRPAGCALLSVGLLLLALGVASAAVVRGQREEANPLTLLLLRGLGGAPAGALVAEAVQLDDGMDVPSSSAAALRRMPPVQAADGDLGDTIFLRAAKRQVRYHQCYFNPISCFG*

**3_Allatostatin CC**

MTLSSVAATLATLAALAALLVPPAATRPAQHDMGLLKGHAADGLGLGLGMGGMGFGRPALALPYLGQDDDDLPLIGKRAVDVGRGGGGGAVANLVAHQDLGSQDSTDYGSSYDEFPMIVPRRAEKRAAIMLDRIFGMMKDAVNGDPNTGLKAIAPDDRMDLQRRGQQKGRVYWRCYFNAVTCFRRK*

**4_Allatostatin CCC**

MVAVRLGHCWLVVLAACMLALCWLAAETSAQPAGEKERILNELDLMDDDGSVETALMNYLFAKQMVSKLRKEMDVTDLQRKRSYWKQCAFNAVSCFG*

**56_Allatotropin**

MRCQATAPLALLCAAVLVLIVASASAAPGPGRQVQTKPRTIRGFKNVVLSTARNFGKRGGAPMDGAQGEFVDEATMQANGAEYQRAPGNSFPVQWFVEEIQSNPELARSVVRKFIDENQDGELSAEELLRPVY*

**6_Bursicon alpha**

MPAAGRVAGRTTPGAGAAAWWALLWLGAACQVVAAAGAGAGGGDAAASGSTPNEECQVTPVIHVLQYPGCVPKPIPSFACTGRCSSYLQVSGSKIWQMERSCMCCQESGEREASVSLFCPKAKPGERKFRKVSTKAPLECMCRPCTGVEESAIIPQEIAGYADEGPLSHHFRTSS*

**7_Bursicon-beta**

MPSRVRAMWLVRSVLLAVLVLAHGAAAETEEACETLPSEIHIVKEEFDELGRLSRTCNGDIAVNKCEGACSSQVQPSVITPTGFLKECYCCRESFLRERVVSLTHCYDPDGARLTAEGRHSMEVKLREPADCRCAKCGDYSR*

**8_Calcitonin-like Diuretic hormone**

MQTSTVVLVAALAMAALFTAVRAAPMGGEPVYNYARPGEIDAEDLVELLKRIAQELRRPEQGLQEAKRGMDFGLSRGFSGAGAAKHLMGLAAANYAGGPGRRRRSAAPAAPAAPGAAPGAAPQL*

**9a_Capa/Periviscerokinin splice variant a**

MREYVLLLAALALGARAAEPLQDHGLGENKRQGLIPFPRVGRSENADPRTMPAAWLVAPEYYPGKRVASWMPSSSPRLGRQNKRFETDNAAWTLVNVRDYPAMNRPQRGRDSASFTPRLGRELESDEEVAVHDAPGLGPQDDLQGSVRL*

**9b_Capa/Periviscerokinin splice variant b**

MREYVLLLAALALGARAAEPLQDHEDGPGPGVGDGLPDHHTRVQREVQGLFPFPRVGRATWGTRDSGLGENKRQGLIPFPRVGRSENADPRTMPAAWLVAPEYYPGKRVASWMPSSSPRLGRQNKRFETDNAAWTLVNVRDYPAMNRPQRGRDSASFTPRLGRELESDEEVAVHDAPGLGPQDDLQGSVRL*

**10_CCHamide-1**

MRVLECVLQVTVVLVLSWACEASTGGCSRYGHACWGGHGKRSSHAVKALPVTLPVRDVDLEAAESGDASETAQDADDAELEAARVEAADALRRRGYQPYQVDRVDSVLPAHLADLISLPREPVVQDPDPVDESEELMHRIRSQNRRRPKNRENDVQVLII*

**11_CCHamide-2**

MKTAAITLVLLGMLWAGTTSAAPQKRGCSAYGHSCFGGYGKRAGGGAGVLVAPLPPPQPRGADGDPTPERLPYLRSLRSPQDDDEAAFIVLRQSPVSSRTPLWGRPAAASTRTYHGEDDDNNGETVSVLEGGGGLGGGAVPAAERRADSAEEAEQAAEPYTNINHADRLFLRRWLRTYRRASDRAARGSARE*

**12_CNMamide**

MCMAPLLLWTAVVAALASLGSVGVSAMPMLTDQGQRAEPYVPTAILEWYLQQRDLQQRDMQQREADLDQPPVQQLQPSEIQQQFREWLTTQLARLPAPVEQGEEANKALVGPPQPKRGNYMALCHFKICNMGRKRSGRLQSQDLP*

**13_Corazonin-1**

MIVLLAAAMAALLLADCAVGQTFSYSRGWKSGKRAVGAAGAALPPSGPPSVSLGGRPADDILAEDLPLYRYFLEGRGEHRVPYDPWRLVEDPVLHLVRRPGLPQSSEQAGEHDER*

**14_Corazonin-2**

MAKTLVGLLAASLVVLLLADVALGQQTFTYSRGWKPGGKRARAVPLPPPPPRPRATPNAPASYPEDVVLDDQPVRYFVEALGERWAPFDVAPWRPQAQPPAQDDDERYRHAEQPSQGEHDDH*

**15_Corticotropin releasing-factor (CRF)-related diuretic hormone**

MVVRRSGGRPLTASWELAVVLLCALCGAGRAAAPWNQDEDLAALGSLDSLLAYEDLRQAVQQAEDAWGQPGAVAGVAPGAVPGAAARGKGAAAWQPAEPDPQLYLLTEYDRGGQGQTRVKRNREAVAAAAAAGRLSARSNNGLRRHVRNPSSSSSQRLSIVNPLDVLRQRLLLEIARRRKASVPQSNREMLEGIGKRSWSPKVSAEQQVERLERCLAELGELNELRGPAKAEDKLARLTLLYRAIEYGNKCDLTSDLGDQGDQRQQQQQQQQQLPSETRPQEQGETNHIRDEQDDWSARQQANGWSRNTRAATSSNLTS*

**16_Crustacean cardioactive peptide**

MQLALMLLGGALLSVAAVTRVHADDVINTVDMDTYDDDRAKRPFCNAFTGCGRKRSGPAPLPVSRPLRSDDPMATLFDLNSEPAVAELSRQILSEAKLWEAIKEASRDEGRRRHSLEVGTLYAAHQDGGDDVVVIVVTRSGCSCR*

**17a_Ion transport peptide splice variant a**

MPPLVARSWACSLACVALLGCLVSAPCSAMVLGHYAGHVGHLGHPLSKRSFFDIQCKGVYDKSIFARLDRICEDCYNLFREPQLHSLCRSDCYSTVYFKGCLEALLLKEEEAKFDLMIESLSGGLQ*

**17b_Ion transport peptide splice variant b**

MPPLVARSWACSLACVALLGCLVSAPCSAMVLGHYAGHVGHLGHPLSKRSFFDIQCKGVYDKSIFARLDRICEDCYNLFREPQLHSLCRKGCFTTDYFKGCLDALLLQDEMEDIQTWIKQLHGADPGV*

**18_Ecdysis triggering hormone**

MAGRGSTAPRPDMAHVLLGVAVALCSLLLVACEPKPEAGGAAVTRQRRSDDYDNMDNFLMGSSDYDNFFLKTAKSVPRIGRRSSAGAAEDPEPMHVDRRSTDANELVPAYANGYDHFFLKASKSVPRIGKRNDEFFMKTAKSVPRIGRRSGDGPDNFLLKGRDQSDFLKTSGEPRLRRSGLVEALRGHRLERRSRAQQVEEAEWPAHGALGGNDAVLGRQHATHPHRIQRRNFDELMAKAAKTVPRMGKRMHPALDEVGDWGTEQQWPWFRQQDNQPGPYKRAPNPADLIDLSQVPWESLDMYLRQMEARRGLGDDDKHLHDVVLIESEAGDKGDEAVDKWAYSVADKKDNKELEEARFQPDVKKRK*

**19_Eclosion hormone**

MPLSAKMVSLMLLLLVLSASLAASNQVHICVENCAQCKRMLGDYFHGQLCAETCVELGGASIPDCQDVDSIRPFLVKADLV*

**20_Elevenin**

MGRLFGALLVLLLAHCCIQAGRASNPSRKSLRSIDCRRLVFHPQCRGIAAKRSMPSLQEERYQLDRSRGRVSRRTPTMTSSSGWTTTRRPPRTRTASGSTGWVRPLTPSWSWRRTTATTSSRPTRPCRWPATT*

**21_FGLa-related allatostatin**

MFRIFKKNKFVVGLASIAATAPVLGTMRDSGIGDKMAVGELKFEAPLAQFEEAEDWGSIEFSKTGGGGVGVLADLHDDVRNSNSPLHHNHHNNLLNGGTRSPEHLLLHQVDDNVIDNSTIVAVDNQQVDNAAQCRQHDNHTTNATTLLNGISVNGDGGEDNFLAESFSGSLEDLVNTFDDKITKCFCNFDESVEKLAPVQGLGALASLKRLEALTDDRLDDDQLEEDKRASSMYQFGLGKRARAYEFGLGKRRAYEFGLGKRRAYEFGLGKRRAYEFGLGKRARAYEFGLGKRSRAYEFGLGKRSRAYEFGLGKRTRAYEFGLGKRARAYEFGLGKRSGRDYSFGLGKRLPNERRYAFGLGKRGMGSGVLLLLLVAAETCRASSGSLSGSGSSPSGAEAAAVSEAVSAAGGAGVGAVAVPGGVFELTGWPYASYSALYKRLPAQYNFGLGKRGADNKLYSFGLGKRAADAEADADADYDDDDGDEVSSTREVPSAEELSPSLALGRPHAYSAVRRVDYLAHCSRPRTLGSDELPEAVEQATEQDWQTGGADSVGKREVAPQELREVAAEHAEHDHHQQQHQEAHSAPGEHGARTKRAASMQYGFGLGKRMVDALVDGALDDGDVLELEDMVDADTDALERAGRASRQYGFGLGKRAYDASNAKRGPTRQYGFGLGRRSMQYNFGIGKRSSGEAGADANLSPAER*

**22_FMRFamide**

MGHWTTLYCVLLAAAYCAAQDDVLADTDPCHHGCDENALLLLPRRAERSVPVSLIATGMLSRRSALENNFMRFGRSAPAAPAASDAKGVQDDGALRPARADNFMRFGRSGAALGTMGSAAEGERVARGGDHNSNGFIRFGRSKDNYMRFGRAGGQDDNFVRFGRPDRQSNFVRFGREPSNFVRFGRSSKDNYMRFGRAGKDNYMRF~~GR~~DPSNFVRFGRDPGSNFVRFGRAGELEEEVAAEDADSEVLRDGRAKQEKDNYMRFGRSDAGDDKRAKDNYMRFGRGGGSSSSGSNFVRFGRSYVDEDDSTEGESARQEAQDASRTKRPQGM*

**23_Glycoprotein hormone alpha2**

MGPRCARVARVLLVFLCLQLVARSLADNGWKRPGCHRVGHTRKISIPDCVEFHITTNACRGFCESWSIPSALETLLVNPHQPVTSVGECCNIMDTEDVTVRVMCLNGFRELTFKSAKSCLCYHCKKD*

**24_Glycoprotein hormone beta5**

MILERPLWLVVVLLALRAARCDVVLAEERPPPAGVDVRTTLECHPRVYNSTVSRSDRNGRTCFDHVSVTSCWGRCDSNEISDYRFPYKRSHHPVCIHDVHEPRVVVLRHCQEGVEPGTERYEYLEAKRCSCQVCRSSEASCEGLRYRAPGPSGPAGPAALVPALVEQVRQLPMVINNRGD*

**25_Insulin like peptide 1**

MRCGDGVASVRLVVLALAMCFFCLAGGERGQTFCGKALPQVLSLVCDGCYNTRDKKYIHNTIDDSYWVWASMPKKEDGDDLEDLEEAEHQPPFRDRLRAADMIGDAYKRFTRGVADECCHKQCSMSEVANYCCEKTARRPYS*

**26_Insulin like peptide 2**

MCSVGSGSGWLVAVLVAFACVSLCLESCLAAAAAPRDPRTYARYSVSSGVYCGSRLSDALKFMCNGQYNSYFKTKKTDQSSQYRGRRETETVAGAEEEYWTPPPALPFRDRLRAAALLENHRRVTRGVFDECCSKSCTIDELMNYCDVPDD*

**27_Insulin like peptide 3**

MTPSLLRVVLGALLALALLDGALSRPHAPRSGEQQQQHDQHGLGNELSARHPHRQHHRHHPHPHHARQLCGAELSNALSALCAGRGYNSPDNMAGSSASRHGHRGLVAECCHVGCTLSDLEHYCQPRKTPLVASQSTSESHEATAEEESDPRPETGNHIRPAVVEAEAEPPQRKEQKAPPQPTVVGAKGSSKINLPSENPSNMIQVGVLTSPLQQLQFRTPVTIPSRPQSNSLGSH*

**28_Kinin**

MLESSRQAGSASRMYSLVLLLALVCVGASDMASDMTSDIASDIVQDTFVDPPYEAGVEVASEGNAEEGPLMRSLAKLCRLIDTEGPPCDGLPASAPTSAEDEEDALDKEVHLDSTDEHDESAEQVSVEADKRAFASWGGKRAPSFNSWGGKRASFNSWGGKRAPAFNSWAGKRESFNSWGGKRAPAFNSWAGKRASFNSWGGKRGGSLVYRLWGGKWIPTVWSGKRAPAFNSWGGKRRKREASFNSWGGKRDAAFNSWGGKRDAAFNSWGGKRDAAFNSWGGKRDAAFNSWGGKRPSSFNSWGGKRNAAFNSWGGKRNAAFNSWGGKRASAFNSWGGKRDSAFNSWGGKRDSAFNSWGGKRDSAFNSWGGKRDSAFNSWGGKRSDELNDPEAKRAASFNSWGGKRNDGQHGDGKRAPAPTNGRGDKRSEGDRGDEKRSASFNSWGGKRADEVAEYAQGDAPLEKRAFNSWGGKRAAILVPSLHSLWGMLGHRPRPAVWSARPSDKRVQFNPWGGKKRSAPAGPADPKPLAAPLAAPLAAPHSDLSKEQPVDAASLKPLPRAKRARDYGTKHLRKNRRGPEFYAWGGKRSGLRR*

**29_myoinhibitory peptide (allatostatin B)**

MRWLALWAVASFTLQGFCLATDDTSSSAHDVVAPEDANHQLVELTDLDEDKRTWSQYQPGWGKRSWDAMGGGWGKRAGSDEVEQEEGEEEEEEDADVDKRGWNNMAGGFGKRGWNNMAGGFGKRGWNNMAGGFGKRGWNNMAGGFGKRGWNNMAGGFGKRGWNNMNGGFGKRAWDNMAGGFGKRGWDSMAGGFGKRGWDSMSGGFGKRAWDNMAGGFGKRGWDSMNGGFGKRSWDSMAGGFGKRAWDKMAGGFGKRGWDSMNSGFGKRSWDSMNSGFGKRAWDNMAGGFGKRGWDSMNSGFGKRSWDSMNSGFGKRAWDKMAGGFGKRSWDSMNSGFGKRAWDSMNGGFGKRGWDSMNSGFGKRGWDSMNSGF*

**30_Myosuppressin**

MRCLPVLSLCAVFAAVLLATGAPRGAQGMALPQCHPSVLQTGSDRIRKVCAALSTMYELSSAMESYLDDKGTLQSESEVLRENAPLVDSGVKRQDVDHVFLRFGRRR*

**31_Neuroparsin 1**

MRTSPLRAAAPAPRWLLLLLLLVVVLAAVAEASTRGLSVRCEPLPEPEDCLYGLARDHCGRFVCAKGPGERCGGPMHLTCGEGMYCKCERCIGCSLNTFTCFLGSALCM*

**32_Neuroparsin 2**

MPAPQLTIHCWLIVLIVGILLQQALRSDAFTLCKTNARADNCAHGLTTDYCGNTVCAKGPGERCGGPSNVLGQCGEGMHCKCEVCSGCSIETVECHFAKATCF*

**33_NPF-like**

MQGGGSVLLASLVAVLLAVGPVGPVGCMAAVGGSRGGQGAPEGGLARPLRPTQFHTPEELRHYLDAVSNYYAVNSRPGRFGKRSSHVPHRGPLVGPLMGPLVGQLQREERVSAQRRDAALENNDQALEALERGLDSGDAMPDYLYQFAPGPAPSPASEVYPLFLDQERLSQPRQTQMQE*

**34_Neuropeptide-like precursor 1**

MERPGDGARRAGLLLLLAFALQVTAVVNGATIAKRSLSALARTSQLPHSSTTADQAGHDQDHGHGGHSPGQHDKRSELQEEAIREELLRQRAAPASEAFLSAIMAADQYGTDLNGVEDKRSVSSLARHGNLPATMEDTKRSLSSLARYGNLPATEDKRSLASLARYGNLPAPAEDKRSLSSLARFGNLPVAIEEAKRSVASLARYGNLPGPAAPSDKRSLEALARGGLLNKNSRIDNSKRSVAALARLGELPKRWMPPHQREQAQDLLDTLQRELDETKRSLSSLARTGDLSRPVALYEDQDKRSVESLARTWSRPSKRVYGYPSYENEYATFGKRNAAALLRQTQFSTQGTVEDSQRDDGDHGQARRKRSLWDDDLMSGEVVQSPSGGAAAGQEQDWETMWSPTAWGAYFGYGDDASVQKRFLGEGQQGGSRTWAAIGRLTRTAAPGGGVPTRLAALRATQGDARDASQRRGRHTSSLSSATITTSITTIVLLLTLPPLLLEGVFISC*

**35a_Orcokinin-A**

MAPSAWTALLLAAPLLLGLAAAYPHRMAPSAWTALLLAAPLLLGLAAAYPHRAYHDEADGAVQEKNVRHIDGLGGGHLLRDTWGMEQLQQLPQLQRIAPQPPCRQRRGLDSLSGVTFGGAKKRFNSLNGGTFGTNKRNFDEIDRTSFNSFTKKNFDEIDRAGFDSFVKRNPVAELEDLERPLYGLSLSQGLARLHHLQEQEQP*

**35b_Orcokinin-B**

MAPSAWTALLLAAPLLLGLAAAYPHRAYHDEADGAVQEKNVRHIDGLGVQQTSDAARQLEEQVRRQVEEQVRATGSGSASGQWSVTKHSEYSGGGSVPVRATITGEALEGGDGLMGITTNCHNCRLRIRKVGGGSSSGAGVSEQAGGAGGVQVESSQSRYYSSRSEGGQGGQSYGSAHPVDLSGGLSGGRLVQSSQQQQYSAGGYGSSPGSYGRVGAVDADGLSSSRRDTKVTKSSWQLADLGYGTSNIEDLIRRLDEDMRAGRYQNGKTVTTVTTTTSENGSPPVTRQETYVKDGLDMSQSQVGSLDMSQSQVGSLGHGFAKGGTTSIVSHSEIEDLLRGLREELRTGKTEAGKTVTTVVESSSVNGAAPVVTKKTYVRDGYDASALELGAAGLDAGFGQSGVSGNSGSHSLSRTTQSVWSSGGVRPLGVDHLGGGNLLTNSHHLARNSTTVSTYGSGSGGHDVRTVDALGGGHLIRESGGYGATSGSEVKGTRRVDALGGGHLLRETGGGYGATGSYGSETSSSYKETRRVDALGGGHLIRDTEGSDLKGTRRVDALGGGHLIRGTDSAQQGASYSGYSGSASSSRFEGQAALQPRPTTDTLSVSQMPGYRKDAPLDTDLPAYGSNSNTGVELERDLDHLGGGHLIRGVPSTGDGQRTQDVGGNQAFIASESGRIPEAGSGSYFSSGSSSYQRASSGSSGINGVNPGAPTGAVVVPVRGPAASSSVISVHGSRSEWAEAQETNDDEFPALGPAAPTSPGIVIGYGSGNGGYGGGTAARPAVASGDSSVQRGGKLSVSLADIGILPSDGSTQQASSGYSSSGSTRYSSSAIASETERHYEVPVASHGTSWSSQSSYSSGMPSRASSSYGSLSQGSRGQRMTQTHTGPDCDGTVKI*

**36_Pigment dispersing factor (PDF)**

MRSPVAALALALILALSLWSAMCSAAPRYDDDAKGLTGLSERDVQLWRDMTLQMLQQVLRAANEDFGPIYHHKRNSEILTSLLGITPKIAAAGRK*

**37_Proctolin**

MACLLGLAALTSPVAARYLPTKRGSQDGDRLDRLAHLIKELLAESDAAPMDHTAYDQRVFYKREAPSYPERDAAAAPLQVPDTGRAQLVAPAARN*

**38_Prothoracicotropic hormone**

..TPPWTPPPLPLYLAVLLLGWSVGVGVSAARLPAHHGGRLGPGPSVEEDNPLPHHTVGHQGGGASSAQQRRCAALSTRVLRQVLGPAYNPAYMAFSLDNPRGGGGGFGPFQSDVRDTRGAGEGLDVQQYSGGDSLSPFSVDEDFSMVLGDEPAWQTQYDMPSVTLSGGADVGGGGAYHSYRTSAGRRAREAPAGEGAAEDAVVTETDAEDASAARAGRAAWSIPMGAGRDMQWHCESPLRWVDLGPDYFPRYLRSADCGHHTCTGSGFVCRPRSFSVKVLRRREGVCAPASDLPREESTVGLPHDLRDMWVWELRAVNFCCVCSMRG*

**39_Pyrokinin**

MGTMASYQLSRAGLLLALCAVCFGLHGTRGRPAPMLLEAEQAPDAEDAALQMELQTLMNLFQAKGKRGQDLAEKVLQPGQTGMFGPRLGRRRRRDVAAAMPSATSATASSGRRKRSVQMSQDGDVDNDIPWKLAELLRELQGAKRSPTPCDSSDINCLLSNIANGGSNYAPSEQEQRSRSEGNLVNFTPRLGRESGEQPEDLEGSMGGAATSRQLRTDSEPTWGFSPRLGRRLLPPGTVDVHAPAGVPPSPASLQPLLALQRSRGGRSARSAPTDKGTGSQGQAQV*

**40_RYamide**

MLLRLHLLALALVVLPAYSEQFYANRYGKRQEFAGPHGQQVGFQSGSRYGRTDPDAAPLGPDDRLQRRFYANGRYGKRSGSGAAAQGAAGATGAAGAALAEYQLLDDFSGLPFVVDEDSQVVCRYTGVTNLFRCARRKEGEDIPAILQDPQSSVVTGNPLN*

**41_SIFamide**

MQVRALVAVFLLTALVLLAGPAAATFKRPPFNGSIFGKRGTPDQVRSMAEVDTTANTMNALCAVATEACN

AWYSADN*

**42_sNPF**

MVTVASSCFAGALCVCLVLGHFASAAPPANNYEGLRELYQQLLEAGAAAGPGEGSAGYAQGYDQGPAPVGLLSLLGGGAGAGAGAGLPLANHQLVRKSNRTPSLRLRYGRRSDPAMKSSCFDGPVDES*

**43_Sulfakinin**

MLLVATVVSAAPSVESAGGASSARTVRARVASGGGSGGAGAVGLGVGSRLRRFPLMQVPLDIMDDDEDAL

FEANKRQMSDDYGHMRFGKRGGGDNDFPEYGHMRFGRAAQ*

**44_Tachykinin-related peptides**

MHLLTVLLAALAVAACAAESSVGFLEEEKRAPSMGFSGMRGKKDDILDSFDKRSRNSLGFSGMRGKKSDDLEEDTTDDGGVEKRSRNNLGFSGMRGKKDDYEEGDGDVGVQDKRARNNLGFSGMRGKKQDSTVEDLDDEVLLGEDGFEKRARNALGFSGMRGKKASSSDEFVMDDLVDKRASGGRNSLGFSGMRGKKALIDEMDAYLAKRARGNAGFVGMRGRRENNQAFFGMRGKRGDGYDEDAVFGYLQSPDAARNGVVTFRSKRYLPSHWQLRNSKKAPFSGFIGLRGKKSYMPAVSSFRAASDQ*

**45_Trissin**

MLNFGPLALLTSGLLLCFLVAPLAAQSCTSCGPECVSACGSRRYRTCCFNYLRKRRSPAAPGAPASEMAEIAGAAVGGVQVPGDALGSDDLRLQLVLLPSDHLEEGQDAAGPLVGFNGAQSLLRRLLDGQPPTDNEGGGGSNGGSS*

**46_Vasopressin/Oxytocin-like peptide (inotocin)**

MRIMSRGGAGSGLLLGAALLALLGVGWACLITNCPRGGKRAGGAAAGRSREVQCTPCGPGGEGLCVGPGICCSPVFGCVLTSRGGCGRGAVFAPRCSPSSAALAAASAAAVPLDAPCGADLTPDGSAPGRCAAQGVCCTHDSCRIDAVCQSRDDNDLNAPGFDADLMLTSLRRVVGGGTTT*
