## Additional file 6 for "Genome-enabled insights into the biology of thrips as crop pests"

**Additional File 6: 112 complete and partial cytochrome p450 (CYP) sequences (amino acid) of *Frankliniella occidentalis* as part of the FOCC Genome Paper.**

*Contributed by* *Jonathan Oliver*

*CYP nomenclature (CYP names) provided by David Nelson (University of Tennesee)*

All sequences in this file were used to construct the CYP phylogenetic tree documented in Additional File 1.

>CYP3661A1 F.o. (1)

MTPITAALLCGALVAAAALVARALRACYKFRRLELDIPGPPSLPVLGNALDLVGLTEASVFPALMALWGGRRTLSRFSILNKLHVFVTDPVDLEAVTKRRDLADKSHFFYDLVQVVARYGLFQINGDLWRRHRRALEPAFHVELLERFVDNFAEEAQGFVQRVVPGQEVDVRENARRAVS

>CYP3118E2 F.o.

MTMMLRRVPQYLCQVHRIQVPLRLRSTAHADAEYAAEPADKPADWDLAKPYSAIPGPKSNRFVGSLMSFLPTGAFFKVKIEDVLDTVAAHGPVAKMTGIWGRPDVILLSDPKEIEIALRSEGAWPIRAGADSLLYYFKNRRKGLPMSLVSSQGKEWQEFRTAVNQVMLQPRNIKQYVEPIDRVTQEIIDRMRRIREQNSLMPANFSDELAKWALESITLVALDTKLGCISDNPNPDGAKMVEYAHTVFDSVYKLDVLPSPWKYVSTPTYRRFTKAMDGITEISSKYVKATMERLKNVPKTELKDRSYSVLETLLVKTEDPDRAVAMALDMILGGIDA

>CYP4PK2 F.o. (2)

VNAQEDSNSEYVQAVKTACCNSYMRLVKPWLYPDATFYASPSGWSFSRALKVLHGLTKSVIHKRRGIHTEELMKIQQSAEDGTKLRVAFLDQLLMANHNGAGLSDIDIQEEVDTFMFEGHDTTGCAITFILYALSVNPEIQEMAYREVTSVVGDETRHLTNQDLAQMKYLERVIKEGMRLYPSVPVFGRSIKEDLALEMDGKRVVPAGATVAICPYVLHRKESLWPDPERFDPDRFLPENSVGRHAFAYLPFSAGPRNCIGQKFAMLEMKCVIARLLMELRFIEAYPGYKPEVAGELILKSRNEHVLVRVEQRHL

>CYP4PK1 F.o.

QAVLSSVHHIDKSPDYSTLHPWLRFGLLTSTGAKWHARRRLLTPTFHFRILDQFLPVFNRNTAVLVKKLEKEAGREAFNLGHYVHLCALDTICESAMGTTVHAQEDSNSEYVQAVKTACCKAYMRLVKPWLYPDATFYATPYGWSFSRALKVLHGLTTSVIQKRKGIHTEELLKIQQPAEDGTKLRVAFLDQLLLANQNGAGLSDMDIQEEVDTFMFEGHDTTGTAITFTLYALSVNPKVQEMAYQEITSVMGDKTRDITNQDLAQMKYLERVIKEGLRLYPSVPVFLRSLKRDLPLEMDGKHVVPAGTTVVVCPYVLHRTESLWPDPERFDPDRFLPENSVDRHAFAYVPFSAGPRNCIGQKYAMMEMKCMIARLLMEFRFLEAYPGYKPKIAGELILKSRDDKLLVRVQKRD

>CYP4PK F.o. (2)

MDVLIVLLGGLLLLILTMLALKRKAFVDKINALPGPPSAPIIGNLLDIAKHNFETLAAAVEQQKEFPNGFRFWIASFPYVVLNTAESVEAVLSSVQHIDKSNDYSTLHPWLRSGLLTSKGAKWHARRRLLTPTFHFRILDQFLPVFNRNTTVLVKKLAALSGREAFNLGHYVHLCTLDTICESAMGTTVNAQEDSNSEYVQAVK

>CYP4PK F.o. (1)

MDVLIVLLGGFLLLILTTLALKRKAFVDKINALPGPPSVPILGHLLYITKHNFEALAAAIEQQTKYPNGFKFWMFCFPYVFLNTVESVEAVLSSVHHIDKSPDYSTLHPWLRFGLLTSTGAKWHARRRLLTPTFHFRILDQFLPVCQKFAMMEMKCIIAHMLIKLRFLEAYPGYKPELTGELILKSKNEQVLLRVEQRH

>CYP4PK F.o. (3)

LLTSTGAKWHARRRLLTPTFHFRILDQFLPVFNRNATVLVKKLAAEGGRKAFDVGHYVHLCTLDTICESAMGTTVNAQEDSNSEYVQAVK

>CYP3657B F.o.

VFPDPLRFDPDRFLPDQSQGRHPYAYLPFSAGPRNCVGFRYALMFVKTAVATLMRSYTFLPPAGGPRTLQELHRQMQPGATMTIKGGALVRVARRAGAAPS

>CYP3655A8 F.o.

MIGVVLLLLLAVAVLAVAALLVRYSGTLSEFRRVRRMMAKLPGPPLMGRGLAVINGDEWRRHRKAITPSLHLDILRDFVQIFSKQGAAFADTLGELADKGVVFDVTPLSGACANHTICETVMSCDVDKRDPMKRDFIDAIPKALDHLMYRVWRPWLLSEWAYSLHRSRFPDYLKVKNDLNQFTQRIIREKKEALKKNEPLPQRRRQAFLDHMLSSPEGAALNDEELAEEVKTFVMIASGSSMDTLTFFIYMLAIRQDVQDKVRQELDDILGDDRGRLVEHSDLAHMQYTERAIKETLRYFSVVPGFARTVTEDLPLPSGFTIPRGCHVAFWLPYVHRNAEWFPEPDKFDPDRFLPDNSRGRHPFAFVPFSAGPRNCIGQRYAMMFLKTVAAAIVPRLRFEVPDDGPKRLEDVPLTMNLTVSVRGGANILVLRR

>CYP3655A9 F.o. (2)

MLLLVLLLLVVAACLAVRSWPVVSEFRRVCRMIAELPGPPRLPIIGNALEILGSPDEIYNRVHTMVSTYSSYGPVCRFAVGHINIVFVLQPADIEAVMTSPKALNKPQVYRPLEPILGSGLVTLGGKVWRRHRKAISPSLHLDILRDFVPTFHKRGVALADQLAGYADKGEVLDVTPVCASCTNNTVCETFMSSDLDDDDPQKLEFVSSTTKGFENFFYRVRRPWFLSEWIYSLHRTRYLEYTKLRKIFDGFVQRMIRQKREGLNNNSPHMNPPRKRKAFLDHVLFSKEGVALNEEELNEEVKTIVYAASDTSMEALSFFIYTLAIRQDIQHKVRQ

>CYP6QQ F.o. (2)

MLVTLVVLVCGGALALVAALYAYLRHSMGYWQRRKVPVVPAVMPWGNFADSILGKRSLFQIADDIYRDHRDERYIGTYFISTPLLHLHDPEIIRQVFIKEFQ

>CYP6EB F.o.

MIDPSVMVWIPVWQAMHDPDIFPEPERFDPDRFTEEAKKSRHQYNYLPFGKGPRFCVAERFAILEMKLCLAGLLQQHEFDVGSKTDVKLKLDPNTPGPTPQNGFWLHVTERQTSA

>CYP3655A9 F.o. (3)

ELNDVLGEDRGRSLQHSDLSNMQYTERVIKEVLRYFTVVPMMFRHITEDLTVPSGFTIPRGCTVGFWLPYTHRDPQSFPEPDKFDPDRFLPENARGRHPFAYLPFSAGPRNCIGQRYAMMFLKTVAAVLVPRLHFELPDDGPKRLEDVPITFNLTLSVSGGANIRVRRC

>CYP3655A F.o.

MIAVVLLLLLAAAGLALALAAPLARLSRAVSEIRRIRRMTDKLPGPPMYPVIGNALA

>CYP3658A F.o.

MLTELVLALVALYLALYAVDKARKLSTLAHIAGPGYPIPIIGHTPYMFGSRDGFLARALVVWRRYGPGPIKFWLGETPMVAVTRPEDVEPLLNSQREVDKHDVFYRPLQSFFGNGLISLNGQAWSVHRRALTPAFHF

>CYP6EC1 F.o.

MDLTATLAAAAAVLAALYLYFSYRFTHWKRRGVPYPKPSPFVGNFGFMLRGKKGFSAAIQDMALPFKENGFCGLYQWATPLLLVWDPQMIKQIVTKDFSSFHDRGLPTNEEDPLTLHLFNLSGNKWRNLRNRLSPSFTSGKLKLMFPLMRDIGAELNAQVAVEAKKTDTHEVEISALLSRFATDVIGSVAFGIQCNCLRDQDNEFLDMSKKIFRTSPIQFLRFILEGIHPKFSKLLPMKWIFSNVHRFFVDLMKETVEYREKNNVERNDFVQLMMQLRNADLYNADPENHMQLTPGVMAAQGFIFFIAGLDNISNTISFAMSKLSTNPELQQKLADEVRDVLRQHDGELSYAALKQMDLLDRVVQEALRLWNPIGMLMRKCNATTQVGDVLIEKGQMVFILSQMTALDENQFPEPQRFDPDRHTREAKDARHPYAFLPFGEGPRNCIAERFALLEMKLAVALLIRDFVFSPGSKYEANVELDEKAFFPRPKNGFHLQVAARVA

>CYP3655A4 F.o.

LIRFSRAVSEFRRVRRMTAKLPGPPMYPVIGNALAVVCSLDKLYDRLDSLASTYGPLFRLSLGHLTFTIVLHPDDMQAVMTSGDASDKSLVYSAIEPLLGAGLATLGGDAWRRHRKGITPSLHLDILHDFVPIFSKQGAAFADTLAELADSGDAFDVTPLSGACANHAIFETVMSSDLDKDDPMKRAILDGIPKALDHFMYRVCRPWFLSEWAYSLHRSQFSDYVNLKNTFNLFIQRVIREKKEALKNNEPLPQRKRQAFLDHVLSSAEGAALSEAELAEEVKTFVGIASGGSMDALTFFIYTLAIRQDIQDKIRQEMDDILGEDCGRLVEHSDLAHMQYTGRVIKETLRYFTTLPTFARTVSEDLPLPSGFTIPRGCQVAFWLPYVHRNPQWFPEPNKFDPDRFLPENSRGRHPFAFVPFSAGPRNCIGQRYSMMFLKAVAAAIVPRLRFEVPDDGPKRLEDVPLTMNLTVSVRGGANIRVHRR

>CYP6BD26 F.o.

MIYFLLQWYIIVPALLALFYWHGTRNFSHWRKLGVKHISPEFFFGNIRKRVLFRQSFHELQKELYFSFPGERVAGIYEGRRPTLMVRDPDLIKLIMVRDFDHFVDRPVLRFRQRPSVNNMLITLQGAQWKAVRAILTPTFTSGKIKSMSLLVMDVGKQMVSFLENVIDKPEGSELEMKDFFGRFTLDVIASCAFGVQCDSLQTPDAEFAAYAGKFNDIPLHERIMIFTLLLFCPQLARFFPLSFMNKGVLSFLESVVRNTKEQRVREGIRRNDFLQLMMDALESEKGQDKTVLNEDILIAQSILFLLAGFETSSTVLSFTAYELALNPEIQSKLREEILGVLQKYDGKCTYDAVHEMEYLDMVIHESMRKHPPVARMDRKCTKEYLIPETNITLKPEASVSIPVMGLHYDSQYYPDPERFDPTRFSAEEKAKRSPYVYLPFGSGPRNCIGARFALTSTKMALVYFLQGFGVEPCLKTEIPYAYSKFSMLLKAEKGIWLNIKRLTHFSDHSSHN

>CYP3657A1 F.o.

MSPLLVAVAALLLAVLALRGLRTYRDTVWHWHTLTRDMPGPFAFPLIGSCWTMAKHPDRMFEQMMWLWKAFGSTIKLWLGPFLFTILLEPDDVEVFMTHPALADKPVAYDLVKPLLGDGLITLNSSEHRRHRKIISTSLHLDILKGFVSTFSERSQDLVDKLRVHADRGEVVDMSPYLGFCVNHSLCDTVLSTDMTGVEADRDELIRVADQASVLAFFRYFRPWYWSERVFALLSSRYKEYRMVVHGMQDFVHKVFTNKLQLLSQGKRPDGKRLAFLDHVLASHDARATLSEHEIKEELRTLIWAGSTTSTDFLSFFFIMISMLPDIQSKIHEELDVVFGEERTRPVDHADLQHLQYLERCIKEALRMFPPVCTWGRQVKEDVKLPSGWVLPKGSVAVVVPYAVHRNPKWFPDPERFDPDRFLVDNCTGRHPFTYIPFSAGSRNCVGQRYAMMNMKSLTASVLRTFTVLPDPLGPRTFQEIPITIGLSMVPTHGSRVRLVPRRTAAERAAGSG

>CYP4PN1 F.o.

MALLWVLVGLVLVSVLLLNQRRYRLAAKLPGPKLWVPGFGNLIDTLINYGLSGTEWIQGYHQRYGNVYSMWLGHKLVVITSDPRDIEAITTSTALIDKGTNYALLAPWLGDGLLLSSGKKWHSRRKLLTPTFHFRILEQFVDVFNQKGSILEAKLGELVGRGAVDVRPYISRHALDVICETAMGTPMNAQLDENSEYVQAIKTACKSVDDRHAYPWLSNDILFRMTSIGRNFYKALDVIHGATQRVIDTRKAELQNELKELNSRSVDEKHEDDDIGSKKRMMFIDQLLTMHLQGEDLSEKDIFHEVNTFMFEGHDTTMTSVCFTLYLMSQHPEVQERVAQEILLVTGSDTDLTSAQLAELKFLDAVIKESLRLYPSVPCFFRDCKEEYTLPSGYTIPAGSSILINAIMAHHDEKNFKDPSTFNPDRFVAGSESEDHWRHNFAYIPFSAGYRNCIGQRFAMMSMKATLARILRSYRFLSSPSDSKLDLDLQVVLTTKSGVMLQLERRA

>CYP4PJ1 F.o.

MDVTMLAMVLVVLAWAFHFFNKQRKWTRAVDLIPGPPSSIVMGNTWELMWKYKFNQTQAFEDWTKKYGGIFRLWVGNNKAVIILSKPEDLEILLGSNKHIEKADVYNQLHSWIGEGLLTSTGKKWHTRRKLITPSFHFRILDQFVEVFNRNSDILVQKLLLQKDTINVHAFITLCTLDIICETAMGTKVEAQKNATSAYVSSVNDATVSFHQRSMKPWLQSDLVFNVSTEGKTYNDALKVMHDYTKEVIESRRKEREIRRNSLSVTEKDLGMKKRVAFLDQLLEANDSLLDKDIQEEVDTFMFEGHDTTAAALSFILYNVAANPEVQEQLVQEMHDLFGDSDRPASSQDLQNMRYTERVIKETLRLYPSVPLFARLVKEDLPVSGGYVIPAGANVTISCLQMGRDPVQWPDPEKFDPDRFLPENSAGRHPYAYVPFSAGPRNCVGQKFAMMEMKSTVSKVVRHCKLDLPSPGYKPKVEGRVILKSIEGVLLKISPR

>CYP15A1 F.o.

MALAVSSLPYPSLTLLLLAVVVVLLLLDASKPRDFPPGPSWLPVVGAYPQLRRLVKKYGFLHDVWRVLAAQYGPVVGVRLGRNRVVVASGLVAVREVLKHADFDGRPDGFFFRLRCFGERLGIVFTEGDHWQEQRRFSQRLLRRPDVEAHVAEEAAELVAALRAQAATARPVPMCRAFDVSVLNVLWARLDGRRFRRDDARLDSLLHTLHDCFRETDVSGGLLNQMPFLRFVAPGRCGYTSTLRNLRTLWGFLGETVAEHKQQTLEPGKPRGLIDAFLEEMKLTGREGSNFSEKQLLSLLLDLFMAGAETTSNTLGFCVMFLLQHPDVQRRAQDELDRVVGRERDPSLEDRDQLLYLEAVLMETQRLGNIAPVGVPHRAVRNAKIMEFVVPKDTIVLANLMSVHMDRDHWGDPESFRPERFLDETGSLKPPSKYFIPFGVGRRRCLGEMLARHSLFLFLAFLLHRFTLSPSPDHPIGERHFLDGFTLSPKPLYAVLTPRDLVGRDHDPASSSEDRKEAV

>CYP4PK2 F.o. (1)

MDVLIVLLGGFLLLILTTLALKRKAFVDKINALPGPPSVPILGHLLYITKHNFEALAAAIEQQTKYPNGFKFWMFCFPYVFLNTVESVEAVLSSVHHIDKSPDYSTLHPWLRFGLLTSTGAKWHARRRLLTPTFHFRILDQFLPVFNRNTTVLVKKLAALSGREAFNLGHYVHLCTLDTICESAMGTTVNAQEDSNSEYVQAVKTACCNSYMRLVKPWLYPDATFYASPSGWSFSRALKVLHGLTKSVIHKRRGIHTEELMKIQQSAEDGTKLRVAFLDQLLMANHNGAGLSDIDIQEEVDTFMFEGHDTTGCAITFILYALSVNPEIQEMAYREVTSVVGDETRHLTNQDLAQMKYLERVIKEGMRLYPSVPVFGRSIKEDLALEMDGKRVVPAGATVAICPYVLHRKESLWPDPERFDPDRFLPENSVGRHAFAYLPFSAGPRNCIGQKFAMLEMKCVIARLLMELRFIEAYPGYKPEVAGELILKSRNEHVLVRVEQRHL

>CYP4PM1 F.o.

MALLHLVWLACQSALPLLLLAVVMLGLAMYFDPAVRRARRIAKGYPQYSSSVPIIGHLLHIVKSREELFHLLREWPKMFGSPSVLMLFGRVLFNLSDPVSIEAVLSSSKHITKGRAYDFLRPWLGEGLLTSGGEKWHYRRKLLTPAFHFKILDQFSSHLEEQAAQLVDDLAELAAAGQGAAVDVVPAVSRITLRSICVTAMGKVDFGTDSSAAVSEKSYFEAIHRVGEATALRIVRPWLWPPPLFWLTPMYQKFRKNLNTLHGFTSKVIADRKMTLSGDADDTAVAAGEAGAENSSRFKRKARLRPFLDVLLNAQRRGEDISDQDIREEVDTFMFEGHDTTSMAISWSLELLGRHPEVQERVVEELRGAGGEDWESISRLPYLDRVFKECLRLRPSVPFISRLIETDAVLPDGRTIPAPSMTNIHIFDLHRHPDHFPDPDRFDPDRFLPEVEQARHPFAYLPFSAGPRNCIGKRFATMEVKLFLAAVVRRFHIRSAMPEHLDVTADLVLRPRDGVRLRLSAR

>CYP303A1 F.o.

MLFLVLAFFAITALLLYLDSRKPKGFPPGPDWLPILGSARELAQLRKETGYLYKACEELVHRYGPVVGLRVGKDRQVIVSGYEAIREMSSRDDFDGRPRGPLYETRTFNLRRGLLLTDSDFWLEQRKFVIRHLREFGFGRRGMAGMVQDEAGELVRAFEEKLRASPEGAIAPMHDAFGLYVLNTLWTMLASIRYTPEDTELVTLQALLNDLFTRLDMAGALFSQFPILQYIAPDASGYTEFVTIHQRLWKFLRDEMDRHIETYDPNEMRDFMDVYIKMIKEAGPNSSFTELQLLAICMDMFMAGSETTSKSLGFGFLYLLRHPEVQKRVQEELDAVVGSGRAPELADRPNMPYTDSVALESIRMFMGRTFGVPHRAVKDTYLMGHFIPKETMMIVNIPSVLMDKRVWGDPENFRPERFIKDGKVQIPDEYLPFGFGKHRCLGETLARSNVFLFTASMLQRFDFRVPPGEPLPSTQCVDGVTAATLPYNALVTHRVAV

>CYP305U1 F.o.

MFTLVLLGSAISLLTAILVLHQLARPRNYPPGPRWLPVLGNLLDLKRLVRKLGSQHAAFESLCQEYGSSVLGLRLGSDLVIVASGYEAVSQVLTSDDWLGRPDNFFMRLRTMGTRKGITMTDGALWSEQRRFAVRHFRQLGYGKVHMEELILAELTDLLNELEPACTTAEAVSLGPLLPPSVLNVLWALTAGTRFPHGDARLQRLLDLMGQRGRAFDMAGGVLGTLPWMRHLAPERTGFNLLNRLNASLRELLLETIADHNASYTQGRTRDLIDAYLHEVKAHAGDGEQSSFTEEQLVMVCLDMFLAGSYTTSNTLNFALMLLVLHPDVQSKCFKDIYNNVEKGRLPSLTDRQKLPYIEAVIMEVMRLYYVVPITGPRRALHDTKLMGYDVPKDTVVLINIHSIHHDKKHWGDPEVFRPERFLNARGKIQPDEHLINFGLGKRRCLGDALARNCLFLFLTGLLQRYRLSVPPGEAPPCREPVPGLTQCPQAYRVMLTPRTACPGSPRSPRTPSPLPGA

>CYP6QQ4 F.o.

MFLLLLLVLAALALGLAYAALRARHAYWRRRGVPGPECSVPFGNYKDAILGRKGMGDIMFDVYQKYRDQRYVGTHFVGTPLLHIHDLEIIREVLIREFQDFNGRAMYSDPDNDMISGQLFLLSGQPWRTLRVKLSPTFTTGKLRFMFTTFKRCGDQLREHIASEVRAGGGRTVIEIKELLACLGTDIITSVAFGIEANTLANPKSEFRRIPALILDPSVGMGLRILLAFMWPEALKALPTTLFPAEANDFFLKTVTEVIEYREKHGVERPDMLQLLIKLKNNGFIPPDAPGQVADGKEVGDKESIADGPVEKQHISIREIAAQCTVFFLAGYETTSSTLTFTLYLLAKHPEVQDKVVEEMREALDKHGGEIEYDTLLNLPYTDMVLNETMRLYPSVPFLNREAMRERKLPLTDLVLDKGTRIMIPIRALHMDPKYWDDPEEFKPERFTKDKMEARHPMVYLPFGDGPRICIGMRMGLIQMKMTLMTILTRAEVRLPADAPSTIKLSPLTVGPTPIDPLKLVFTERA

>CYP3659A1 F.o.

MSYTATSILGNFSVTKFLTNLTLAAILWLLILQAFQFLRKCIHFRRIEQTIPGPFSVPFFGSVFWLSLNADNCLEKILDSVKPYWPGLVRYSAFNKLHVMVNDADHIAKVLRDRHLSDKPWMFYGPLKPIIRDGIFTSNGEPWQAHRLVITPTFHAEVMGRYVEILDMEAKEFVDKLSGMLHKDVDCSDTMIRTASFMAIRTMMSSTLNERECSILEEIISLVPRALKSVNRRSLNPFLWPDWIFKWTQMYRDTEAAKTLFDMLINSIIQRKSSEIHATASRERQLASKLGFRDRKSLLDLLLESKFSGELRMTDEDILNETSTMFNTAFDTPVIVVTFVLKVLSIRPDIQELVYAEICDVLQDKDHKLSSDDLSRLQYMDRVVNETLRMFPIAPFVARQTNKDVELDGKHIPSGTVLFLNFVGVHRSPKYHQDPLTFDPDRFLPDRAAAMHPYSFLPFSSGSRNCIGQPLAWMMVKTLLASILKTYQVFPASDGITDPSQFKTTFDLVLRYIGGVKVRFEPRN

>CYP4PM2 F.o.

MMTLSSVWPASEMALLLLLLLLALLAFGVFKSKYFDPAAIRARRIAEGYPQYSSLPVIGHLLHVLRPRDEIFALPRQWPKMFGSPSVLVLFGKTFFNLSDPDSVVLSSSKYITKGFAYNFLRPWLGEGLLTSGGEKWHYRRKLLTPAFHFKILDQFSSHLEEQAAQLVDDLAELAAAGQGAAVDVVPAVSRITLRSICVTAMGKVDFGTDSNEKSYFEAIHRVGEASAVRMVRPWLWTPFFLLTPLYWAFRKNLNTLHGITRKVIADRKMTLPDDDDDTAGAAGEAGVGSASHTSGVKRKARLRPFLDVLLNAQRRGEDISDQDIREEVDTFMFEGHDTTSMAISWSLELLGRHPEVQERVVEELRGAGGEDWESISRLPYLDRVFKECLRLRPSVPFISRLLETDAVLPDGRILPATSMGNIHIFDLHRHPDHFPDPDRFDPDRFLPQVEQARHPFAYLPFSAGPRNCIGKRFATMEVKLFLAAVVRRFHIRSAMPEHLDVTADLVLRPRDGVRLRLFAR

>CYP4PP1 F.o.

MLALLLPVLAVLLLWQVLAKARARHAKLLLQFPAPPSAPIVGNAVDYWGNEEHIFNTSKRFWKTLGTKFVTIVMNETTLNLCDAEGAEAVMVSSKHIKKGPLYTFLTQWLGYGLLTNFGPSWHHRRKLLTPAFHFKILDEFGPLLQNHAQRLVDNLADLVAKTKGPVDVVPPVSKATLGSICETAMGVNLHALGKEEEYFECVRIMGCSVLYRMIRPWLWRDFVFNLTQHGRETKQAMDKLHNFTKQVIQERKRSHASTGASILDGDGKRLGKRRLAFLDMMLAAQREDPTLTDDDIRQEVDTFMFEGHETTAMAISWTLFLLGNHPEVQQRVREEAQEAGDDWQAVVALPYLEKVVKESLRLYPPVLYIARENDRGIQIGDKWVPANVNLDLQIFMIHRDPNNFPDPDKFDPERFSPEQERARHPFAFVPFSGGLRNCIGRRYAMMELKIMLAALVKSFSFKSVLSEKEMEFKPELVLRPKGGIVTNVTPLL

>CYP6EB1v2 F.o.

MSRRMFRRNFVFLLRFFLISIHPVFAKLLPFRKIFKDMTEFFLRLMSDTVNYREENKVERNDFVQIMMQLREEDRNRSTLDRASHVELNNDTMAAQAFLFFVAGLDNVANTIGFALHELAMNHALQRRAVAEIQENIRKHGSLTYDAVRDMELIERIVRESLRKYSPVGILTRQPSEPIKIPDSDVVIDPSVMVWIPVWQAMHDPDIFPEPERFDPDRFTEEAKKTRHQYNYLPFGEGPRFCVAERFAILEMKLCIAGLLQQHAFDAGSKTDVKLKLDPKTSSPTPQNGFWLQVTDRPTSAL

>CYP3656A2 F.o.

MEGGAWVWGDEVQAEERCGAYVVAGLILAALAAAAGLCLVTLVSRVAVGVRATRRMRRVLAPMPGPAALPILGNIPELLTSQDKFWTKITTLMSRYAPTFCFWLGPTPFVVITDPRDYEHVMGSARFNDKSSWYPLARAALGTGLITLNGEPWRRHRKAIAPSLHQHVLVQNVEVFRRKACELSHMLSRAARSAETVDILEFCNLVSMDSVCETLLSADLSHHGAALGRFVDLTTDAARSVTTRICKPWLLWDSLYALTRAGRTAARTMQETDAFIDGVIAKRKDGDRKDAAKVTKRSFLEHALASENLANLMTAEEWREEAKAMVVAGQGTVAHCLSFALLLLAMHPDSQDRLRRELEDVFGEDPDRSVTEHELPHLQYTERVILETLRLFPLVPVFGRDCPEDVQLPSGHAVPRGAHLFMFPFHTQRDPAWFPEPLRFDPDRFLPERVRGRPPLSFSAFSAGPRSCIGQRYAMMYLKTVLATVMRRFAVHRAPGDVRGVADLPIEIGVILSLDGGFQVRVSELGDPPSSAQAATRHIH

>CYP3656A1 F.o.

MRFTLVQVCVGAAGVAALVYVGYLVSRLLAIVTEVRRMKKVLGDTPPGPRSFPILGNVLELLAPRNGFWSKMVALVQKYAPTFRFWVGPNCFMVITHPSDFEYVLNNPKFNEKSSWYPLAQSALGTGLIVLGGEQWRRHRKAVAPSMHQQIITQNVQMFYRKALDISEEVLSAAAQSGATLDIKDFCNTVSMDAVCETILSADLSSSREALKRFVDAKNKSCDLVAYRVYNPWLLSDRVYSLTREAREVRRIEKLTDRFVDEVIAKKRDRDHVPSTAEPSPKRRSFLEHALADATVAKLMNDEELREEAKAMVIAGEGSVATAMSFTLLMLAMHPGVQKAVHQELEDVFGDDPDRPVTEDEVPHLQYLERVVLETLRLFPIVAVFGRNCPEDIQLPSGCPAPKDSQLLFFTYHMHRDPAWWPRPLEFDPDRFLPEQTHTRPPLCYAPFSAGPRSCIGQRYAMMYLKTALATVLRRYDVHRAPGDVHEVADLPLEIGVILTMQGGFPLRVSRRQDAQSSAGIKSSVFK

>CYP302A2 F.o.

MYNMALQCGPSKIGFWKNIIGAVKQVNLNLSKRSLSSSLSEPEQEQLTQDIKPFDEIPGPKSLPFIGTVYLYSLGVYSFENVIESGRKKYLKYGRIVREEIVPGKHTVWIFDPEDYKQLILYETTMFPSRKSHLAVEKYRKSRPEVYCSGGLLSTNGEEWWKLRREFQKDFSAPQAVQAYLPATDRIIKEFVNRQLLFPCDFDFMFVLSRLSLQLTCLIAFDEEFDSFSNQQLKYDSRSSQLIEAAFTANQTGFMMDRVGFQTRLWKRFSEASLFIESVAVELVKKKKRSLEKSNPLEYSEDYVPSLLELYLTMPSINYKDVIGMTVDLLLAGINATAFTASFALYHLSKNERVQDILRMEARDLLQSPEQAITPEILSSAHYARAVLKEVLRLNPISAGAERLLTEDIVLSGYHIPKGTNVVTQNQVSCRMDDYFVNPDVFLPERWMKGSPVFEKSNPYLVLPFGHGRRTCIARHLAEQQLLILLLRVVRDHMIRWNSEKPLGCKAIPINKPDQVVDLSFMKV

>CYP6QQ2 F.o.

MLVMLVVVVCGGALALVAALYAYLRHSMGYWQRRKVPVVPAVMPWGNFADSILGRKSLFQVADDVYRRHRDERYIGTYFSATPLLHLHDPEIIRQVFTKEFQDFNGRGVHSDPENDPLSGHLFMLSGQPWRNMRVKLSPTFTTGKIKYMFNTISECAGYLREHVDTSLKTGGGLYQEDVRELVARFSTDIISSVAFGIETNSMKDPESEFRRMGQRIFEPSLDVNLRALLAFFGGNIMKLFRIRGNPYDVHAFFTRIVDEMVTHRERNGVERADMLNLLIKLKNEGFVPPDAHGEHADKGNGATNGTNGVSQESRTKLTQKEIAAQVFVFFIAGFETTSTTTSFALYELAKHPDVQEKVLQEIDATLKKYNGQVTYEAIMDMPYMEMVIQ

>CYP4C110 F.o.

MVVESAAAGRAPLLASRVSVALAAVAGCCALLLLVSAARRSLFLLRLRAVPGPRALPVLGNALQLTGTQEDFFLLMRRCAERYGPMFRLWVGTRPFIFLQSAEAVQPILNSSVHIDKSPDYRFVEAWIGKGLVTSTGDKWRARRKLLTPSFHSRVLEEFLQPLYREARVLADKLAAHCGGTAFDIVPYAKLAALDVICDTALGKHVAAQRNSNNAYVVAIDRITSIVQRRFITPWLKPDALFNLSELGRTQKKHVNVIHGFTGRVFEARKQEKQREWEREQLEARQGQQVQFQEDQPHAVPDARFTIKPRRRLALLDLLLDLAHQGANLTDEDIREEVDTFTFAGHDTTAAGFSWCLYVLGKHPEVQERILEEWRDARLEAGDGEMTVGDLSRLDYLERCIKEVLRLYPVVPIIARDIRSPVTVLGEHLPAGCTVLVNAYLLHRDARFFPEPERFDPDRFLPDSDTRRHPFAYVPFSAGSRNCIGQRFAMLELKVLLLTVLERFQVCTAEPQRPLRLVAELVLSNVEGIKISLSHRS

>CYP3660A1 F.o.

MPCWAPSWGHVAVVAAALLTARLAGGARWCALAAAALLLLLLLLLVARVLLWVARRAARAAHQHALTRDIPGPRLEHFLRLWRARRSPARVSDLFHDVAREHRDGGIFKFWSGPVLFVILMRADDIECLLTDRRAATKSLVYDLIGTLMGEGLLHLSGERWRRRRRIVQPYFSPEVLRGFPEVFHRRAEVLVRLLAPSARSGQAVNVLPFAGQAVTDMVCHTVMASDVDTRAMEEEGIANAMSSGLAIILYRVFNPWFLHDGLFKLSRLHAHFKRSMELFDAFTLRLLSIKRGERQAQPRGPADGGAQRKATFLDVMLDTEAALTDGEVLDEVKGLITAATGGSTDALCFVLLCISLRQDVQDRIVQEITSVFGEDVDRAGTMDDMRHLEYLERVIKETLRMFPSIPQFGRSPSQDIELPSGHKVPAGAQVFVYSPAVHMDPAHFPEPEQFEPDRFLASAGRHPFAYLPFGAGAHTCIGQRYALLQIKSTVSVLLRHYRLLPAGTPRSPAALHDIRTNVLVTQRARGGYWMRLEPRRPPPADPATG

>CYP6KF1 F.o.

MVLSAVAGLLPVMGVFEVLVAVLGLLVVRYYLRTANFWTSRGVLSKNGVPILGSSPSFILQSEFRGTIMQRFYTEARAKGAPCVGLFVGSQPVLLAVDQDLLRSVLVKDFSHFMDRGMPFDRHNEPLTANLFNIEGQEWRTLRHKLTPTFTSGKMRNMFPLVQACADELTAVLAAEAAAGRPVEMFELLARFTTDIIGSCAFGIQANALKDPNSEFRAMGKRAFDMNILQTFFNMVMPRAVPYVKMVFPVRLIDKDVADFFTSTFEENVKYREKNKVLRHDFVDLLMQLKNKSNADTEGIDADILPAQAFVFFLAGFETSSSTLGNVMTELAYNQDVQDKVRAEVQRVLPKHGGDISYEALSELTYMEQVIDETMRLYPAASNMVRMTNDDYVLPTTGADGKPAVLPKGVKVIIPIFAIHRDPEFYPEPDKFNPDNFSEEAKAGRPNYAFMPFGEGPRVCIGLRFAMMQMKSGLATILRAYRVLPVEGDKHYPQTFEKRTFVTKTVGGNKVMLRAL

>CYP6EC2 F.o.

METTTVLAAAATVLVTMYLYFNHRFNYWRKRGVPFAKPVPFVGNFGFMITGKKSFSMALQEMALPFKENGYCGVYRLGTPLLLVWDPEMMKQIVCKDFSSFHDRGLEASEHDPLSQHLFSLSGTKWRNLRNRLSPSFTSGKLKLMFPLMRDIGAELNAQVAVEAKKTDTHEVEISALLSRFATDVIGSVAFGIQCNCLRDQHNEFHEMSKKLFRPSIFQFLRFFLEGIHPKLVLLLPMKWIFSKVHLFFVDLVKETVEYREKNNVERNDFVQLMMQLRNADLYNADPENHIKLTHGVMAAQGFIFFIAGLENVSNTISFALNKVCADPALQQKLVDEVREVLRQHDGDLSYEALQKMDLLNRIVLEALRLWNPIGVNVRKCNATTQVGDVRVEKGQMVFILTQMASMDENQFPEPQRFDPDRHTREAKDARHPYAFQPFGEGPRICIAERFALLEMKLALALLIRDFVFTPGPSFESEVELDGTAIFPKAKNGFRMQVTPRV

>CYP3118E1 F.o.

MSFLPGGEFYKVKLEDMFDTLASYGPVAKMAGIMGRPDIVLLSDPKEIEIALRNEGTWPIRQGADSLQYYFENRRKGLPMSLVSSQGKEWQEFRTAVNQVMMQPRNIKQYVEPIDSVTQEIIDRMRRIRDENSLMPANFSDELAKWALESITLVALDAKLGCISDNPNPDGAKMVQCSHTVFDSMYKLDVLPSPWKFVSTPHFRKFTKAMDGITEISSKYVKAGMERLKNVPATELKDRSYSVLENLLIKTEDPDRAVAMALDMMLAGIDTTSTVVATVLYQLATNPEKQKTLQEELDRVIPDKRKPLTKEQLEELKYMRACIKETMRIMPIIIGNSRTSVKDLVLGGYQIPAGTMLNTPNYFLGRNEKYFPRAKEFLPERWLPGAPDLKCKNPFAYLPFGFGPRMCVGRRFADLEIETIVAKVFRNFNLEWNNPPAALKMQVVLVFADPLRFKVTDRA

>CYP307B1 F.o.

MESLFAVSASTSLLLCAAAVVAVLFLADRARQARAAAKAKLLDSEFGDELPAPPGPHAWPIIGNLHLLAGYAVPYQAFTPLANRYGSVFKLSLGSAPCVVVNGLSNIKEVIVTKGAQFDGRPNFRRYHELFCGDKENSLAFCDWSAVQKARREMLRAHTFPRAMSARWTQLDSLLGIELDAVRSRLDAAGAGAAVEAKPLLLGACANVFTSYFCSKRFNPDDASFRDMIHNFDRIFDEVNQGYAADFLPWLLPLHSRHFATLAGWSHKIRDFMTNSILQERLDAWEHGKQSDATKDDYVEELIGHVRGNAEPAMSWDTAMFALEDIVGGHSAVGNLLIKVLGYVATHPEVQDRVHEEAVRAGVTKGKTVTLADRTLMPYTEGVVLEAIRLISSPIVPHVANQDSSIAGYKVEKDTLIFLNNYELNMSPALWTKPEEFMPERFVSPEGRLAKPEHFLPFGGGRRSCMGYKMVQYVSFATIANLLNEYRILPIQGRDYTVPIGNLALPWDSYQFHFERR

>CYP3655A5 F.o.

MIVVVLLLVAVVLTVSALLVRYWGALSEFRRVRRMTAKLPGPPMYPVIGNALEGVFGEDDLYGHLQSIMSTYGPLTCLAIAHLTFVFVMDPDDMEAVMTSPKTSHKSVVYGAGEQLLGRGLVVVNGVEWRRHRKAITPSLHSDILRDFVPLFSKQGVAFADTLAELADEGVVFDVTPLSGECANRAICETVMASDTDEDDPMMRAFLDAIPKAQDHFMYRLWRPWFFSDWVYALHRSRFQDYLKVKDALDRFPQKIIREKKKALQKNGTLPQQKRQCFLEHMLASEEGAALSDAELAAEVKTFVMAASATSMDVLSYLIYTLAIRQDIQDKVRQEIDDIFGEERGRPVESSDLAHMEYTERVIKETLRYFSVLPLFARTVTEDLPLPSGFTVPRGCEVAFWLPSVHRNPKWFPEPDKFDPDRFLPEHSRGRHPFAFVPFSAGPRTCTGRRYAMMFLKTVVAAIVPRLRFEVPDDGPKRVEDIPLLMNVSISVRGGVNIRVHRRS

>CYP3661A2 F.o.

MIPITAALLYGALVAAAVLVARASRALYKFRRLEQDIPGPASLPVLGNALDLLGLSHEGVFPALMALWGGRRTLSRVSVLNHLHVFVTDPVDLEAVVKRRDLADKPRVFYDLLRAIAPYNVILINGELWRRHRRALEPAFHVEILKSSVEHFAEEARGFVERVVPGQEVDVRENTRRAISRLEYTERVIKETLRLFPPVPLTGRQVHRDTELASYKVPAGTTLLLNLFGAHRDPAHWPDPLAFDPDRFLAERSRGRHPCAFLPFSGGARNCMGLQYAMMNMKTFLATVLRELRVERADDPYTDIHQLPLYADLSLRIVGGARVVFARRPTRETEVGSQTTS

>CYP3657D1 F.o.

MLELVVWCGVAAVLARAALWLLAAVREVLHQLWVTRDIPGPPSYPLVGSFLAVAGPQEELYEKLVALRADLGPTVKLWLGPVLVVIVSRPTDLELVFTDPRLQTKPWLMYASSMEPCLGAGLITMNGADYRRHRRIISPSFAQDVLHGFAPSFNANAHRLVEQLGELADSGEVFDAAPLSRMCTSYTVCETVLSADSSRLESDRRAVVQAVYRGVTLMFHRGIRPWYRWDTLFKLSKHYDDYQEMVRVGDSFVSKVLDLKMEAQAKETLGQAEEAGEPDAQGSGKQGKRKAFLDHLLSSEEGKTLTKDELSGELKHLVAAANSTTSDTLSFFLYCIAIRQDVQAKILEELDAVFEGDREREVEVADLPKLKYLERAFKETLRYFAPAPIYARQPDSDVKLPSGATLPKGCVVLVSPHFTHKDPDIWPDPDVFEPDRFLPHNSVGRHPHAFIPFSAGTRSCIGQRFVMMFLKTAAATVLRRFHVTTPEDGPRRVRDIKPIFSLTLHVKGGARIRVQRRT

>CYP315A1 F.o.

MWSAVRVGAAARKLPRAAGTSLPPARRRQHSAPRRGLPIDAMPTPGDLPGTPSAARLHEYIDMRHRQLGPVFRDRIGPVSAVFVADPDEMRRTFGLEGKYPMHVLPEPWVHYNQLFGVKRGLFFMDGADWLKMRRLMNPLMLRTDQGDIFSSACQHVAESLVSKWKAGGTRVIGDLEAQLYKWSLDTVIAVLMGPLYETHAPKWNDEVEYLAHNIHKVFLETGNMSLISAKDAQEGQLPVWKRFITTVDASLNTARSLVLQMIPLSTGESGGLLKGMIECGMSEEDIIKVVIDFIIAAGDTTALSLQWTLYLLAENPLTQGHVAAELTQETDILQSALVRGVVREALRLYPVAPFLTRIMPESCVIGGYELPPKELVLLSLFTSGRNANNFPEPERFWPERWLRTSHSTNGSYKAVHNPHASMPFAMGARSCIGRKLAETQMITTLAAILRDFEIELLNTENIDMVLRMVAVPSIPLKIGLRPRTQTSL

>CYP4PL1 F.o.

MSVISVVLGGFGASATRPEMLTSALLVGALGAAVLYYMLYSWKRRRMYSLAAKIPGPTGLPVIGNVFDVVGDTEETVRKTMAQFQQFRPTYKYWLGPLLLVALTDPRDLEIVLNNYKYTDKSHFYNVFHPLGGQGVFNAAGDQWRRNRKIISPAFNFNFLVQFVAVFHEMAMRLVGKMKARADGQSFDSYKLVELCTLDAIAVTAMGVDVNAQDDDNSEWIRAIRRCFQITRERLVKPWLLPDATFYLTKDSRDQQVCLDIVNKFAHEVISRKKAEYRKAKAELDADPVASKAAAKAREQSRLQQDDDEVVGVRKKQTFLELLIERSESEDGCALTDNELRDEVVTLLIAGQDTTATDNCFNLLMLALHQDVQEAVHQELLDIMGDDPSTCPTYNDLNQMKHLERVIKETLRLYPSAPFIGRDLDHELDVTDYRLPAGCTVLLGLYGTHRLPDHWPEPDKFDPDRFLPERSVGRHPYAFVPFSGGPRNCVGQKYAMLQMKTILSTVIRHFQILPGQDCESMEKFRLQVDFSMTVAGGHNIRLVPRYQGMPERMRSL

>CYP4PH1 F.o.

MGLAVTVLWVVGALALLQLLWSLRNRRLLQLANTLPGPRFLPVIGNFFSILANDFDFIKMVYAYRSRYGHLYRIWLGPACYINLSQAKDVETIMSSQKFIHKSRDYAHLHPWLGTGLLTSTGSKWFHRRRIITPTFHFRILDRFMDSFNRNSNLMVENMSKEDGRPAFDVHHYISLCTLDIICECAMGTKVNAQVNSTKQIDSPYVKAVNSMCWLVYFRGIRPWLFVDSIFNRTAQGKLFNSNLKTLHAMSKDVIRKRKLEYVEDRRRASAAAAQGNGPPAASRRAKGLPTPPVTRTESEVLEAKVDAATGLGPDEDNVYGRKKRSAFLDHLLELAETENLSDEDIREEVDTIMFEGHDTTAAALSFAAYSLATNQHVQDKLFDEMQHIFGDSDRDATYQDLADMKYLERVIKETLRLYPSVPMFWRDIREEVTLGSGFVLPAGSTACCNMYMLGRNPDAFEDPELFDPDRFLPDRWSGKHPFDYTPFSAGPRNCVGQKFAMLEMKVTLSKLVRNFRLLPPPQRFSLDLAVEVVLKSRSGVMLRVEKRALKPSKPSRNSA

>CYP304N1 F.o. (1)

MSPVLLVLFLVALLACLVRYARSRPPDFPPGPLRLPVWGNLWQLLLRNNKYPMKAMAALSKDYGNPLLGLFLGDFPMVVANDLETVRDMFSHPAFQGRPDGFTGRVRAFGRLHGIFFTDGPYWEVQRRFSLRHMRDYGFGRRFEKLEALVLEEVQGVTQLLRANLTPHDADVARGGRVLFPDVFAAASMNMALAVLTGTRVTPDQYPAARVVARHVLEFQRAGNAGGRAIDATPWLRFVAPGLSGYTAYKGGNDGLMHFIQAHVQDHKDSFDSEIIRGFCDKYLLEMQKRAGEDSSFTEMQMVLTLMDYIFPAATAIPSALTFAVIFMMRHPEMAERAHQEIKDVVGLDRLPDLNDRVDLPYTEAFLRETLRLTGTPFAVTHRATEDSYFRGYYVPKNTLIVPNAWAAHRDPNVFEKPFEFMPERFLQQNGQLKKKDVTVSFGLGKRLCAGETFARQNLFLFLAALLQGFRFAMPEGVPVPEPEEDAMFGFIMTPTDEMWIEVSERR

>CYP4G235 F.o.

MAVIMGLPVWAALVGLATLLSYVWFRIAFRERMALSAKLPGPTAYPLIGNLYMIKESTVELFKYAVNAAGTFNEIGLAKIEWMHRVAVAVFNPADVEIILGSSVHLEKAKEYRFFKPWLGEGLLISSGDTWRTHRKLIAPTFHLNVLKQFVYLFNRNSRALCRKLDAVVGQVVDIHDYMSETTVEILLETAMGVDRSTQNRGYDYAMAVMKMCEILHVRMFRPWLWPDWLFNMVPMGREQKSLLHTIHSLTREVLKKRKAERKQGVREGLYKRVVTDKDDVADGDAEESTSETSVSSYGYTTGLRDDLDDDVGEKKRIAFLDQMLDVAENGAVLSDREVEEQVNTIMFEGHDTTAAGSSFFLSLMGVHQDIQERCVDELRTIFGDSQRPATFQDTLEMKYLERCINETLRLFPPVPVIARHATENIQMKSGHVIPEGTTIVIPQYLIHRDPTQYPNPEVFDPDNFLPERCSERHFYSFVPFSAGPRSCVGRKYAMLKLKILLSTVLRKFHVRADVLEKDFQLTGDIILKRKEGFPVRLYPRSTHQAAGQSGDR

>CYP3653A1 F.o.

MRLVNNTMVLLPWISVVAAVTAALVAYVKWAQSYWWRRGVVQTRPRLLGDLWDQLMRRRTAGDVIQSICRKHAKEPFVGIYNITTPILVVKEPELINEVLVKNFKSFANNETVIPEGADRAFGYNPFFAASHRWKELRLRLSRAFNPSRVKIAFSDMATSADCLAHLIAGSEPGTSHNGLLLAKRYTAHVSARVLFGIESHSLEIGSQPGVFYEMGHKIFNCDFLQNIRLMVFIYYRPLSKLIGYNIVSESIVDFFRKLIRDSVEYRHREGVTREDFVEHAAKTIMVDGKVAEDQLSKLAGQAINSYVETFETSGITLASVLLMLATNPAQQERVRAELKEKLKPGEPLEHDAVVGLPFLEQVVNETMRLFPFSDTMRRMCTKATTLRSAEGPSVLVEEGTAVYIPVETVHRDPAYYSHPEDFWPDHFSPEESEGRPKSVYLGFGDGPRQCPGMKYGTLQVKVALATVLQRFEVLPGEKQVLPVQRDPMAGLLNYKGGIWLKFRALAE

>CYP6EB2 F.o.

MIVPSVLAAVAALLAAVWYYFTRNFSYWTDRGIVSCNPFSLTGDGGGGLPFGKPFYELLTPIYQKYKGKERFVGTFQARRPVLVPIDPEVTKAILTKDFSHFHGRGNPTDESDPFSQNLFFLEGARWKNLRMKLSPTFTSGKLKGMFPLMSGITDQLNEAVRDKAAKADNEVEFKELLTRYSTDVIGSVAFGIDCNTVRGESDEFYLMSRELFSRNLLGLFRFFLVSIHPVFVKLLPFRTVFRGMTTFFLCLMKETVEYREKNKVERNDFVQTMMQLRAEDRGRSMLDRASHVEFNNETMAGQAFGFFLAGLDNISNTVGFALHELAKSRELQELAVAEIRQKTEEHGGLTYDALREMDLVERIIRESLRKYSPIGILTRQPTDSVKLADSDVVIDPSVMVWIPVWQTMHDPEVFPEPERFDPDRFTEENKQARHQYHYLPFGEGPRFCIAERFAMLEMKLCLAGLLQTYAFDFGSKTDLKLELAQDAAAPTPKNGFWLRVNDRPSST

>CYP3657C2 F.o.

MLTLLLAAAAVAVALAWRHAARWLRMFWMARTIPGPSNGVLYFAGPLGEVVDKSRVAQQRYGNTVRVWAWPAVNVTVCEPEDLEVVFNSPYLQHKPAIVYGSMSPILGEGLVTLSGDTHRRHRKAIGPSLHLEILQEFVPAFERNALKLCEQLGWLASTGEEFNAVSLLGSYSAHAICETVFSAEAPPILGESKDHFVKYLLEASRLMLSRVMKPWLSWDAVFFFSSQHAEYTRAAQAFEAFVSKMLAYKVDQLRQGDAYTAHGGGRRRKAFLDHVLTSPEGQVLAPRELTEELKTISGAAVGTSTDFMSFFLLVNALLPDVQDRIFKELEDVFGGSERPVEPHDLPHLQYLDCAIKETMRMFPPVFTFSRTVLQDVELPSGHVLPEGCIVAGLPYFTHRDPRHFPDPETFDPSRFLPENSRGRHPYAYIPFSAGSRNCIGQRYAMMSAKILSSSILRRFQVLPSSDGPRTLRDIKLVAGISMSPAGGCNVRVVSRRPRSA

>CYP6EC5 F.o.

MEPSSLLLAAAAVVAAVFLYFRHRFSYWARRGVPFHAPKPVVGNFGDVLLGRKNVPGLLADLSRAFREVGYCGIYSFDKPVLLVWDPEMVRQIVAKDFSSFHDRGQPFDPADPLSQTLFALGGKQWKNLRSKLTPAFTSGKLKAMLPLMRDVGADLVDQVALEAHGGCEVDMEELLSRFVTDVVGTVAFGIKCNCLRDPNSDFRAMSKKLFKQTPASVARILLTTVHPKLGVLLPFKWFFAETNDFFLNLMRDQVDQREKTSVERNDFVQQMMQLRAADLADVDPDNHLKVTTGVMAAQAFIFFLAGLDNISNTLGYILTRLAGDQDLQDRLASEVRRELQQHGGELSYEALKKMSLVSRVALEGLRLWNPLGVVFRKCNATTRVGDVVVEKGQMLFIATNVINMDESVFPEPQRFDPERHTAEEKAKRHPYAFLPFGEGPRICIAERFALLEMELTIAVLVDKLRFSPGPNYEKEVEIDTRAPLPKPKNGFHLQVTERI

>CYP307A2 F.o.

MFLSSVLCFTAAALAIVMLAVDHCRRRRQHAGDAAKDLPQRAPGPKVWPIIGNMHLLGEHANPFIAFTALSKKYGDIFSMVLGNTPCVIVNNYELIKEVLITKGSHFGGRPDFLRYHALFGGDRNNSLALCDWSELQKTRRQVARLYCSPRFSSLHYDTLTHVAGSEVDHFLDRLGSMTASAAKAGAPVSVNIKPLIQAACANIFAGYMCSTRFSYDDADFVNIVRLFDEIFWEINQGYAVDFLPWLAPAYRQHMGRLAKWALDIRTFILTRIIDEHRRRLVEGEDVELDQCMARDFTDALLIQLQEDPHLEWQHIIFELEDFLGGHSAVGNLTMLSLAAVAARPEVSARIQAEVETVTGGSRAVSLADRPNMPYTEATILEALRVASSPIVPHVATQDTTIAGYQVDKGTMVFINNHELNTGEAYWDEPAAFRPERFLQTAKDGSTVVVKPAHFIPFSTGKRTCIGYRLVQSCSFVLLASLLQNYNVTVDPEHPVVTYTATVALPPDTFPLAFVPRDAASTVPAPA

>CYP3659 F.o.

PLRPPTAAEVHSLKYMDCVIKETLRLLPSFPFVARTVREETELAGKTLPAGTVLVVNIFGLHRDPRHHHEPLQFDPARFADADRERHPYSYLPFSGGPRSCIGQRYGMMAVKTMLAAVLAEYSIEPADEDDAKAPADFPLTFYMSLHLVGGVRVRFVPRERH

>CYP3118B1 F.o.

MAPPLQTAAPFTGFLRHARGAWPRSRSTLAAPLHREDAATTIGDRDADWDSARPFSEMPGPRPAPLVGNMWRFIFGNLSKVTPTELPRWFFNEYGSIMKLTKLPGRRDLVFCFDADASEKLLRNEGLWPTRLGLESLGYYRSLTPEKFEGALGLVSTQGKDWASFRSAVNQTMMQPRNAKLYVGAIDTVSQQLIDRMRLIRDQNNVMPDDFYNELSKWALESITYIALDKRLGCLDNNLRPDSEAQRIITAVHAIFDCLFHLDVQPSLWKYVPTPAYRRMKKTMDWFYGIVFGYIEEAMQRLQSMSAEERETHEYSVLERLLLRNENPRIAVTMALDMVTAGVDTTSNTSAAVMYQLAKNPDKQRRLQDELDRVFPDPNSPVTADRLEELRYLRACIREAMRVSPIIIGNQRMAVKDLVLCGYQVPKGTDVLCVHSEMVKEERYFPRALEFLPERWLPEGEHLKAAHPFAHMPFGFGPRTCVGRRFAQLEMETFLSKAFRNFNIAWTGAPLKRELRLLYSVASPLKFECVDRV

>CYP6QQ3 F.o.

MLLLLFALAALALGLAYAALRARYAYWKRRGVTAPECTAPFGNYADAILGRAGMGDVMFDVYQKYRDQRYVGTHFLGTPLLQLHDPEIIRQVFVREFQDFNGRAMYSEPDKDMISGQLFMLSGQPWRNLRVKLSPTFTTGKLRLMFSTFKKCGDQLREHIASQVRADGGRTHIDVKELLARFGTDIITSVAFGIDTNTLANPKSEFRRISGLILDPSIGMSVRAMLAFMWPNALRMLPTTLFPAEANEFFLNTVTELVEYREKHGVDRADMLQLLIKLKNDGFVPPDAPGQVADEKEAGGEESIADADAPVEKQRISIREIGAQCAVFFLAGFETTSATNTFALYLLAKHPEVQDKVVAEMREALDRHGGEISYDTLVNLPYTDMVLNETMRLYPAVPFLNREAMRERTLPLTDLVLDKGTRLMIPVRALHMDPKHWEHPEEFNPERFTKDKVEARHPMVYLPFGDGPRICIGMRMALIQMKMALMTILTRAEVALPADAPSTIKLSPHTITPTPVDTLKLVFTERAD

>CYP3657B2 F.o.

MAMLVALLAVCVALLLLSAEARAALRRAVKGAATVARVVYLTRGVPSPPSLPLVGCVPTLFALFRDPRAALTDLIDKYGKTVRLHLVNRVLIVAMDPDDVQAVCTHPALASKPELYADLLGPHMGVGLVNINGPTHRRHRKAITPSLHFDILSDFVPIFVRNSEILVARLAERADGDSFDIQAEFGRLTSTTFIQTVLSSREATAADIGKDSAVLYEASHLMMWRAMRPWWYHDTVFKLFSDQAEPFFRTSETMNRMVSRVLTDKTAEVARGEAPPPRRRMSFLDHVLRSKEGGLLSSDELREELKTFLFVGASTSMDYLSLMAYILSYFPDVQRKVQEELDGVFGRPGTAGADRPLVADDLPHLEYTERVLREVLRYAPPIPIIFRVASEDMTLPTGTFVPEGCFVGVVPAGTHRLQEHFDEPWAFDPDRFLPERTRGRHPYAYVPFSAGARNCIGLRYALMLAKTVTASVLRRFTILRAADSPETLEDVKFDFSLTMSVRGRANVRLQSRLVAAA

>CYP3653A2 F.o.

MVELTSWISVAVAVIAVLVAYVKWKQSYWWRRGVVQTQPRLLLGDLWDPLMRRRTAGDVIHSICRKHPKEPFIGIYKLITPTLVVKDPELISEVLTKNFSSFANNETLWPEGADKVLSDNPFGATFDRWKELRLRLSQAFSPSRVKIAFGDLATSADCLADFIAASKPGVTHDGELLCKRFTAHASARVLFGIESRSLEVGAEPGVFYEMGHSIFNGTFIENLRFISILFFPALLKVIGYNIVSDSTFDFFRKLIRDSVDYRHKEGVTRDDFVQHAAKTIMVDGKIAEDKLTKSTGQAINSYVETFETSGLTLGSVLLQLATNPREQDRLRTELREKLKRGQPLDHDTVMTLPFLDQVLTETMRLFPVADTIRRVCTKPTTLRSADGPSVDLDVGTPVYIPVEAVQKDPAYFSHPDEFWPDHFGPEEVESRPKNAYLGFGDGPRQCPGMKYGSLQVKVALVTLIQRFEVSPGDKQVLPLQRDPTAPFLNYKGGMWLKFRPLP

>CYP306A1 F.o.

MWLLLALLLAGAALALRLWLRRARDLPPGPWGLPVLGYLPWLDRLRPHETLAALALRYGPIFTVPMGGVTAVVLAEEALVRTALAREACSGRAPLYLTHGLMQGHGLICAEGPMWRDQRRLRRLVAAALKNLGAVRSPGPRRDRLQERVCRGVRAFVQSVTEASRQQDGVVDPHAHLTEAVGNVVNSVVFGRTWAPDDPTWLWLQRIQDEGTKLIGVAGAINFLPFLRLLPQNGRAIRWLKEGQRSQHGLYAELVAEVRSGLGEADDQDDDGCGAPDNLLTAWLRESARRPDSPFYTDRQLHYLLADLFGAALDTTVTTLRWLLLFVAANPVAQERVHRELDAVLGGREPTLDDQAALPYTQATIMETQRIRSVTPLGIPHGVVEDTTLDGFRIPKGTMLVPLQWAIHMDPQRWDRPDEFRPERFLAEDETLLRPDGFIPFQTGKRMCVGDEWARMMLFLFSAGMLQRLQLSLPDGVEADLEGVCGITLTPKDQALRCVERQLRKK

>CYP18A1 F.o.

MLLVSRASQWLLRTSYSAYNSADMGWSLLVFVVVLVLVRLAQLYREQRRLPPGPWGVPVLGYLPFLKGDIHLHYKELSEKYGPMFSARLGSNTIVVLGDHKMIREAFRKEEFTGRPHTEFSSILGGYGVINSDGRLWKDQRRFLHERLRHFGMTYIGAGKGQMEARIMGEVESFAAILEEHGEGPVDLNPILAVSISNVICSMIMSMRFTHNDARFRRFMDLIDEGFRLFGTLAPVNFIPALRHLPWLQNTRNKLNQNRQEMADFFQETVDNHRASFDPSNIRDLVDTYLLEIQTAKEEGREGALFEGKNHDRQMQQIIGDLFSAGMETIKTTLQWAVVFMLHHPEIMKQVQEELDSVVGRERLPSLEDLAYLPFTESTILEVLRRSSIVPLGTTHATTKDVTLNGYHIPKDTHIVPLLHAVHMDPNLWDEPESFQPSRFLSAEGKVTKPEFFVPFGTGRRMCLGDVLARMELFLFFSTLLHKFHIELPEGESLPSLKGNTGITVTPDAFKVVLKKRPLVSGATPVAPSSTEAGVVRNVGSH

>CYP3118E F.o.

SINPEKQKMLQEELDRVIPDKSIPITKEQLEELKFMRACIKETMRVMPITIGNSRTSVKDLVLGGYQIPAGTMLNTPNYNLGKNEKYFPRANEFLPERWLSGATDLKCKNPFAYLPFGFGPRMCVGRRFADLEIETIVAKIFRNFNLEWNNPPAALKMQVVL

>CYP3655A6 F.o. (2)

MIGVVLLLLLAVAVLAVAALLVRYSGTLSEFRRVRRMMAKLPGPPLYPNAEWFPEPDKFDPDRFLPDNSRGRHPFAFVPFSAGPRNCIGQRYAMMFLKTVAAAIVPRLRFEVPDDGPKRLEDVPLKMNLTVSVRGGANILVLRR

>CYP3655A7 F.o.

MIGVVPLLLLAAAVLAVAALLVRYWGALSEFRRARRMTAKLPGPPPYPVIGNALDLPYNMDDLYDTLSTFASTYGPLYRATLAHYTFVVVMDPEDMRVVMTSDKTAEKSSLYGVAESLMGRGLAIVNGDEWRRHRKAITPSLHLDILRDFVQIFSKQGAAFADTLGELADKGVMFDVTPLIGSCANQAICETVMSSDVDKDDPLKRDFLDAVPKAFDHFMYRIFRPWFLSDWVYSFHRWQFPDYLQMKNTFNLFTQRLIGDKKEALKKNEPLPQRKRQAFLDHVLSSEEGAALNDAELAEEVKTFVMLASASSMDALSFFIYMLAIHQDVQDKVRQELDDILGEDRGRLVEHSDLAHMQYTERVFKETLRIFPILPLFGRIATDDLLLPSGVTIPRGCQVGFWLPYVHRNPQWFPEPDKFDPDRFLPENSRGRHPFAYVPFSAGPRNCIGQRYAMMFLKTVAAAIVPRLRFEVPDDGPKCLEDVQLKMNLTMSVRGGANIRVHRL

>CYP4G236 F.o.

MAVADLLTPTGLFYYLLVPTVILWYAYFRYSRRRLYELAAKINGPDGLPLLGSALDFTGNAHQVYKTVWTRSQDYDNEAPIKLWIGPRLLVFLQDPRDIELILGSNVHLDKSPEYRYFAPWFGNGLLISTGDHWRSHRKLIAPTFHLNVLKSFIDLFNHNSRGVVEKLRKEIGHEFDAHEYMSEATVEILLETAMGVSKKEQQSGFEYAMAVMKMCDILHLRHTKFWLRPETIFRFTKYFGEQKRLLGIIHGLTKRVIANKKLEFNKGVRGSLANYPGMKPVAETETKASKSTKVTEDTGMSYGQSAGLKDDLDVDDNDVGEKKRLAFLDLLIESAQNGVVLTDQDIKEEVDTIMFEGHDTTAAGSSFFLCMMGAHPDIQDKVVEELDQIFGDSDRPATFQDTLEMKYLERCMLETLRMYPPVPIIARQLNEEVKMASNGCIVPAGCTVIVATVKLHRNETIYPNPTVFNPDNFLPEKSAARHYYAFVPFSAGPRSCVGRKYAMLKLKILLSTILRNYRVKSTVPEKDYQLTADIILKRADGFKIALEPRKRSGVSVSPSTATPVRA

>CYP3655A9 F.o. (1)

MLLLVLLLLVVAACLAVRSWPVVSEFRRVCRMIAELPGPPRLPIIGNALEILGSPDEIYNRVHTMVSTYSSYGPVCRFAVGHINIVFVLQPADIEAVMTSPKALNKPQVYRPLEPILGSGLVTLGGKVWRRHRKAISPSLHLDILRDFVPTFHKRGVALADQLAGYADKGEVLDVTPVCASCTNNTVCETFMSSDLDDDDPQKLEFVSSTTKGFENFFYRVRRPWFLSEWIYSLHRTRYLEYTKLRKIFDGFVQRMIRQKREGLNNNSPHMNPPRKRKAFLDHVLFSKEGVALNEEELNEEVKTIVYAASDTSMEALSFFIYTLAIRQDIQHKVRQELNDVLGEDRGRSLQHSDLSNMQYTERVIKEVLRYFTVVPMMFRHITEDLTVPSGFTIPRGCTVGFWLPYTHRDPQSFPEPDKFDPDRFLPENARGRHPFAYLPFSAGPRNCIGQRYAMMFLKTVAAVLVPRLHFELPDDGPKRLEDVPITFNLTLSVSGGANIRVRRC

>CYP6QQ F.o. (1)

ERADMLNLLIKLKNEGFVPPDAHGEHADKGNGPTNGVSKESRTKLTQKEIAAQVFVFFLAGFETTSTTTSFALFELAKHPDVQEKVLQEIDATLRKYNGQVTYEAIMDMPYMEMVVQETLRMYPAVPFLTREAMVRRQLPLTDVLLDKGTRLMIPVNALHYDPQYWDQPKRFDPLRFTEEAKKARTGMVYMPFGDGPRICIGMRLGLVQVKMALITILSRAKVSLPADMPKEARMNPRCIIPTPIDGIRLNVSAR

>CYP3655A6 F.o. (1)

MIGVVLLLLLAVAVLAVAALLVRYSGTLSEFRRVRRMMAKLPGPPLYPVIGNTLEVACSMDEVYNRLHSLMRRYGPLARLTLAHFTFAFVMHPDDMEAIMTSAKGSDKWAAGYGALEPLMGRGLAVINGDEWRRHRKAITPSLHLDILRDFVQIFSKQGAAFADTLGELADKGVVFDVTPLSGACANHTICETVMSCDVDKRDPMKRDFIDAIPKALDHLMYRVWRPWLLSEWAYSLHRSRFPDYLKVKNDLNQFTQRIIREKKEALKKNEPLPQRRRQAFLDHMLSSPEGAALNDEELAEEVKTFVTVASGSSMDALTFFIYMLAIRQDVQHKVRQELDDILGDDRGRLVEHSDLAHMQYTERAIKETLRYFSVVPIFARTVSEDVPLPSGFTIPRGCQVAFWLPYVHRNAEWFPEPDKFDPDRFLPDNSRGRHPFAFVPFSAGPRNCIGQRYAMMFLKTVAAAIVPRLRFEVPDDGPKRLEDVPLTMNLTVSVRGGANILVLRR

>CYP4PQ1 F.o.

MPGLLVLLAWAALLLSLALALRNRRRNALIAKLPGPWLFVPLAGNMLELIVQVIWLKGDLVELGTKNVRKYGPIFRLWSGHQPVVLISGPEDAEVVLTHPAVADKGQNYKLLSSWLGDGILLAGGAKWHSRRKMLSPAFSVRVLEQFVALFNKHCAILEDKMLALAGGPPFDITPFMSLFALDVISETSMGVQIHAQTQEDSEYVRAVKTACRCFLVRQVFPWLRSDTLLALTATGKSLSEATGVLRGMTTKVIDQRTSEVASSVSSSQPEEEDDIGLKQRAVLLDHMLTLRGRGITDEDITDEVSTFMFAGHDTTMTALCFTLHLLSLHRGVQERVFQEVRDVVGDSGPVTAQHLNDLKYLECVIKEVLRLYPSVPVIVRDCKQEFDLPSGYTIPVGSQVGIDIFHLHRNPAVFPRPLEFDPDRFLEGGPAHRFGYLPFSASQRNCIGQRFAMMEMKTVLARLVRSFRILPPSDDYRLQLKIEIVLGAQGGYLVALERR

>CYP3655A1 F.o.

MVAVVVLLLLAAAALAVAALLVRFSRAVSELRRVRRMTAKLPGPPMYPLIGNALELACSMDEVYDRLHSLASTHGSMYRLAIAHVTFAFVMDPDDMEAVMTSGKTSDKSVTYGTLEPLMGRGLAVLNGDEWRRHRKAISPSLHLDILRDFVPIFSKQGAAFADTLAELAEKGEAFDVTPLSGACANHTICETVMATDLHRNDPMKRDFLDAIPKAFDHFMYRVWRPWLLSEWVYSLHRSRFPEYLKVKDSLNRFTRKIICEKKKALQKHEIPQRKRQAFLGHMLVSEEGAALSEAELAEEVKTLVMIASASSMDALTFLIYTLAIRQDIQDKVRQELDDILGEDRGRPLQHSDLAHMQYTERVIKETLRFFSVVPIFARTVTEDLLLPSGFTVPRGSQATFWLPYVHRNPQWFPEPDKFDPDRFLPDNSRGRHPFAYVPFSAGPRNCIGQRYAMTFLKTVAATIVPRLRFEVPDDGPSRLEDVPLTMNLTVSVRGGANIRVHRR

>CYP6KF2 F.o.

MAAFEILAAVVAVMLVFYYYMTSTADFFAKRDIPHLPQTPFIGSSVDVFLSRRWRGKVWQDWYNYAKTNGHKFLGVFLGRRPMLVVCDVEMVKTILVKDFQHVMDRGMYFDRQREPLSANLFNLEGREWKALRGKLSPTFTSGKMKSMFYLVRACTDELESYLGGEVQKGGEVEMAEVMAKFSTDVIGSCAFGVEANSLKDPNAEFRNMGRKVFEVDRLRVIVMMIAETMPWLRSLLPMPLINPQVKNFFTRTFNENVEYRERQKVVRHDFIDLLIQLKNKEAGHEGTDHSATGIDEAILPAQAFVFFLAGFETSSAATSTCLVELAHHPEIQEKLREEVDRVLAKHGSITYESLHDMTYMEQCIEETMRLYPALAGLPRVVTRDYQLPGESPKAVLPAGTRVIIPTYAIHCDPEFYPDPMRFDPERFTEENKKSRPHYAYLPFGEGPRICIGLRFAMMQMKTSLAMILRSYRVTPSQKSEYPLQFEPKSFVTRLRGGNRLRITKR

>CYP6EC6 F.o.

MDALLSQLPAAACLLLVVVIYLFYRKRYSFWRERGVPGVKPRFPFGNFAAVALSRANLLQVIYNLAAAHRRVGYCGAFIWHTPVLLVWEPEMIKQVLSTDFGSFHDRGRRVDEADPLSQTLFNMSGQRWKNLRSRLTPAFSSGRVRNMLPLMLDIGRELVRQTALEARRSPQHTREVDMGLLLSHFATDVIGSVAFGIQCNTLCSQENNAFLAMSKRPFKQSPLRLLRQGLNAIHPKLGALLPIKTVFSDVHNFFINLMRDTVEQRERHKVVRNDFVQLMLQARSAELADAADADPEHHVELTPEVMAAQGFNFFIAGLDTFANTVGFTLNRLAGDEELQERVAAEVQEVAATHGGELSYDALQDMDLVRRVVLEGLRLWSPQAVFLRTCTATTRVGELTVEKGRGVLVVGSVNNMDEDIFPEPARFDPDRHTAANRNNRHPYAFLPFGEGPRACIAERFALLEMQLALAALLRDLRFSPGPNYEREVQLDTRSIFHSPKNGFLLKVDPRN

>CYP6EB1 F.o.

MFVATVLAAVAALLAAFWYYFSRNFGYWTERGVVSCNPFSLTGDRGGGIPFGKPFYELLTPIYNKYKGKERVVGTFQARRPILVALDPEVTKAVLTKDFSHFHGRGQPTDESDPFSQNLFFLEGARWKNLRMKLSPTFTSGKLKGMFPLMAGITDQLNDAVREKAAHGDKQVEFKDILLRYSTDVIGSVAFGIDCNTVKGGNDDFLNMSRELFRRNFIFLLRFFLISVHPVFVKLLPFKRIFKDMTEFFLKLMSDTVNYREKNKVERNDFVQIMMQLREEDRNRSTLDRASHVELNNDTMAAQAFLFFVAGLDNVANTIGFALHELAMNHALQRRAVAEIQENIRKHGSLTYDAVRDMELIERIVRESLRKYSPVGILTRQPSEPIKIPDSDVVIDPSVMVWIPVWQAMHDPDIFPEPERFDPDRFTEEAKKTRHQYNYLPFGEGPRFCVAERFAILEMKLCIAGLLQQHAFDAGSKTDVKLKLDPKTSSPTPQNGFWLQVTDRPTSAL

>CYP304N1 F.o. (2)

MSPVLLVLFLVALLACLVRYARSRPPDFPPGPLRLPVWGNLWQLLLRNNKYPMKAMAALSKDYGNPLLGLFLGDFPMVVANDLETVRDMFSHPAFQGRPDGFTGRVRAFGRLHGIFFTDGPYWEVQRRFSLRHMRDYGFGRRFEKLEALVLEEVQGVTQLLRANLTPHDADVARGGRVLFPDVFAAASMNMALAVLTGTRVTPDQYPAARVVARHVLEFQRAGNAGGRAIDATPWLRFVAPGLSGYTAYKGGNDGLMHFIQAHVQDHKDSFDSEIIRGFCDKYLLEMQKRAGEDSSFTEMQMVLTLMDYIFPAATAIPSALTFAVIFMMRHPEMAERAHQEIKDVVGLDRLPDLNDRVDLPYTEAFLRETLRLTGTPFAVTHRATEDSYFRGYYVPKNTLIVPNAWAAHRDPNVFEKPFEFMPERFLQQNGQLKKKDVTVSFGLGKRLCAGETFARQNLFLFLAALLQGFRFAMPEGVPVPEPEEDAMFGFIMTPTDEMWIEVSERR

>CYP3654A1 F.o.

MDNFTVGYGRGSEINATLYAAGQYHSDEMWTTSTLLLVAVLCLYAVLLRLNLGSTFRFHLPGRQSLQELPEALALQAVYLRARESEKTTPGIRVGAGLRWETLLVDLNVIHRVCTDNITFPDRGLPKNERTNPLSAHLFSLDEKRWRAVRDVMSRVLTPAKLKSLFSLHARTSQDLRCSVRGEMNSEGFADVDVKELMERFVTDDMGRTLFSVDCDALRRPGNPFHSHCAQSFTNSWWRQLRQRLLRVWPSLALRIPHRGSTAQFFQKMVEDLRWNRGHCQKTQADLLEAMFVAESSGGCPALMRSVEAPQMFASFRAGFEATAPTITFALLELAHNPNVQRRLQQELDTIRLLQQEVADDEDNTDDLQVTADALAHMEYLDCVIKETLRKYPPNASLVRRSSKPCFASEFGDSRTTVIIEEGERFVVPVYALHHDRRWFSDPERFDPDRFRHQQDGGCTPSNNNQKAYMPFGIGRRHCVARRYALQQVKVGLVAICSTFEVELCSKSPPIPPPFLAGRGIVLALRNECYLRLRLRNL

>CYP3658A2 F.o.

MGLVSEALWGCAACSLAVWLALALARYVRWRLYWHRALSPFPGPRVSLPLLGTLLYIVGPMSTVLRRGLRLMEGYGNEYITFWLGERPWLHIIRPEDVEGLLSSSKHLHKPDIVYGPVSGFFGRGLITINGDAWRRHRRVLTPAFHFAVLERYSGIFTRRAEAFAKGLASVEAEGRSFDFMPHIGAFVTDTVMETAFGLVGGDEDLEGKEKADFIQATDRAFQIIAGRVIQPWLMVESLFRLSPYAAQQRQADKVINQYASRVIDKKRRQIIERKRNQLDSGAEQENDDDKDDIGIRKKLTFLDLVLGQDTDALSDQEVLEEVRTLIAVQQTSASTLSFAAVCLALHPEAQRLAREEVLAVDAVEGLEPLERLNRLKYVERFLKEVLRLYPITAIFSRTLGEDHVLHNAKTKTGTRSVTLPAGLGITFFVYQTHRDPRHWPEPERFDPDRFLPEQCAGRHPYAYVPFSAGPRNCIGQRYAMLQMKAVLAALLRTVELSPGKGCERQDKLDININTFLYIKGGFNVRLRKLDTDRHGCWQR

>CYP6QS1 F.o.

MLVLVLVSVVALLAGALYLFLERRYSTWKRKGVPGPKASVLAFNVPKVMLGQENMGDVTDRIYNEYRDSPLAGTYMFGSPLLHLHDTEIIKHVLIKEFQDFNGRGIGASVSNAAVDPLATHLFFQSGQPWKTLRVKLAPAFTSGKIKYMFETIRECSYQLSDYLDERIAANGGSHCEDSRTLMMRLFVDTIASVVFGIESNVVKNPESAFFTAAQEAVKPNLTNALRSLLAFFARRFMEAFSIRSTSPATTKFFLDVVSDTVDYRENNKVERPDLLSLLIQLKNRGFVDPDKADGKVAEQGQPGDKKLTMNEVAANVFVFFIAGFESSSGTSSFTLYELAKQPELLKKVQQEVDQVLADFDGKLTYDAIMSMPYLEMCINETLRKYPPLAFITREAMVTRKVPLTEVTVEKGTRVVVPVRALHYDPLYWPDPEKYDPERHTAEKKATRPAVTFMPFGEGPRMCIAARLGVVQIKTALVRLLSKYTVTVSKDEPDKVRMNPKSFATTSDMPLNITFHPRKAAAGIQKRRQSVVSG

>CYP3657B1 F.o.

MGVVLTALCAVCAVLLAPYALRLWRHFREVVRAMYLTRNVPGPPSLPLLGCVPTFLGVWTDMHGTMHRLLQSYGTTFRMQVLDRVTLVVMDPDDVQAVCTHPALANKAKPYKLLENHLGSGLINMNGPAYKRHRKAITPSLHLDILQDFVPVFAKNAEALARRLRQHADSGAPFNSSPEFGMLTNMTIMETVMSMGNEGASENEYVDMANVLDEASTLIMWRAMRPWWENDAVFTLCSRYKAYNETASTMNEMVTRILHTKSAEVSRGDPAPPRRRMAFLDHVLRSSEGANMSGDELRDELKTFLFAGSTTSMDFLSLVALALTMFPDIQSRVQQEVDEVFGAPGTEGADRPMVSEDLGHLQYTERVLREVLRYAPPVPLMFRSAAEDVKLPSGTLVPRGCMVCLVPAGTHRIPEHFPDPLRFDPDRFLPEQSRGRHPFAYIPFSAGTRNCVGQRYALMFSKTIVATLVRRFTFVRAPGGPKDFGEVSVVPGITISVRGGAVVCVQHRVQHRQRAAA

>CYP6QQ1 F.o.

MLALYLSCGAVALCAALYVYMRRSLTYWQRRKVPVVTPTTLFGNYGPSILGKRSLSEVAQDIYNEHKDERYVGTYFMLTPLLQINDPDIIRSVFIKDFQDFNGRGMHSQPSYDPLSAHLFILAGQPWRNMRVKLSPTFTTGKIKYMFDTISECANHLREHVDSSVAAGGGRYVENVRELVARFSTDIISSVAFGIESNSMKNPDSEFRQMGHRIFESCLDVNLRHLLLFFGGDVMKTFSIRCVPAAVHNFFTRIVDEMIDYREKNGVERADMLNLLIKLKNEGFLPPDSNDTAEEKNRHSDVNESNGTEARLTKLSGKDIAAQVFVFFVAGFETTSTTTSFTLYELAKHPDIQDKVVAEMDAALAKHGGKVTYDTIMDLPYTDMCVQETLRMYPPVPFLTRQAMVKRQLPLSDVVLEKGTRLMIPVRALHYDPQYWDEPLRYDPQRFTEEAKKARPQLVYMPFGDGPRICIGMRLGLVQVKMALLTILTRARVAVAPDMPKEVTLDPRCILPSPVHGCRLVLSARK

>CYP6EC7 F.o.

MQSRGGRPAAAMDAVLLLLASACLLAGVLLHLFYRRRYSHWRRRGVPGAEPSFPFGNYGAVAGGRANFLQVTNSLAAAHRGVGYCGAFRWHKPVLLVWEPELVRQVLSADFASFHDRGPPADRGDPMSQTLFNMDGQRWRNLRSRLTPAFSSAKVRGMLPLMLDVARELVSQTALEAERSPQHEVDMGRLLSHFATDVIGSVAFGIQCNTLRDHDNEFLAMSKRPFQQTLRRLVRHALIAIHPRLGALLPFKRIFPDVHTFFIDLMKDTVEQRERHKVVRKDFVQLMLQARSAELADADDPEHHIELTPEVMAAQGFNFFSAGLDTFANTVGFTLNRLAGDEELQDRVAAEVRRVAGDEHAAGELSYDALQGMDLVKRVVLEGLRLWSPQDVLVRRCNATTRVGDLTVEEGREVLVVASVNNTDEDIFPEPARFDPDRHTAENRKRRHPYAFLPFGEGPRACIAERFALLEMQLALAVLVRAFRFSPGPRYEREVQLDTKTIFHSPKNGFPLKVEVRHFDK

>CYP6QR1 F.o.

MIGWTLTALLAGLGFALYSYLAKIFTYWTKRKVPGPKPTLLTGNFGSNLMGNKSIGDIADQVYQDYRDEPVAGTFVMTQPLLHLHDPEIIKHVLVKDFQDFHGRGIYVESERDPLTAHLFNLSGKRWKELRAKIAPTFTPAKIKYMFETMKECCTQLQDFVEAKAAANGGRYNADIREMQARVSTDVVSSVAFGIQSNSLKTPESEFRRIGTKVVEPTVFNSIRLMVLFFLREIAQKFQFTITVTEIETFFTKLVAETVAYREQNGIERADMLQMLMQLKNKGFVQPREGEEAEPVPEDGQPGGKLTMNEMVAQVYIFFIAGFDTTSATTSLTMYELTKNHDIQEKVYQEVQDVLKRHNGEVTYEAIMEMTYLDKVINETLRMYPPLPFLNREVMVNREIPLTNIQVNKGTRIIVPVRALHHDPQYWPEPSKYDPERFSEEAKASRPAHVYLPFGEGPRVCVAQRLGLMQVKVSVAMLLSRFRLEASEDTPAEVEFAPRSFVPLPKKELKMTLVSRQIAAA

>CYP3118F1 F.o. (1)

MRRTALLRTRPIAIGAQAPAVRFRSSMTATATPTLATDEEVKPFSEVPGPKPLPVLGNLWRFFPGGDFAGVAPYDIPDTLVRRYGPIVKLEGMLTDTKLFVADPTAAQQIFRTEGQWPIREGISSIDYYYKNYRKNLQFSLGTANGEVWQKFRSAVNQVAVQPRNVKVYVEPINAVAEEFIQRMKSIRDANQQMPDDFTTEIKKWAAESITYVALDTRLGMLENNIHPDSDAGRLVELVKNTFDAMAKLEFGLSPWKYISTPTWRQLKSSMDTFTEMATRHVNEVVDRLSKLSEEEIASRQFSVLERLLVNSHDSDKGVAFALDMMLAGIDTTAHQTSSLLYFLATHPEQQRALQAELDREWEAGQPLTASVLEKLKYLRACIKEAMRLQAVVTGILRTANQDVVLNGYLIPKGTHTIVCTKTMCRDERLVPRASEYLPERWLPGQEQLKPRHAFVNIPFGFGPRMCAGKRFADLEMEVLCARLFRSHNLEWHHPEPQVEQKLFVQYSSPLKITMKER

>CYP304P1 F.o.

MSPILLAAFLCALVACIIKFARSRPPMFPPGPPRLPLYGGYLHLLAYNYRYTYKGLVAMAARYKSPVLGLYVGSDPTVIATDFASTREMLLKPEFQGRPDTYAARLRSFDELLGIFFVDGSFWAEQRRFSLRHLRDFGFGRRFPRLETAVGEEIRTLFEAIQHPVGTEKRLVREGQVLLPDLLYAGFTNALLETLAGERLPRARHETLRALSRAVRLFQRHVEPSGGVINITPWLRHLAPTASSYRGLVEGNDGVVDFIQDELLSRQVPTFDAEVMRGFIDVFRAEMLNRGEDKSGFTEKQLIMVLVDYIFPSTTVPPVTVAMAIAYLVRHPDKAERAHREIVAVVGRGRLPNLDDRASLPYTEAILREVLRLETNTVLSVTHRCTEDTFFRGYFVPKGTLLIPNIWAANRDPAVFEKADQFVPERFLDLRSGMLRKKDDTMPFGLGKRLCAGETFSRQNMFLYLSALLQNFSFELGDDGYLPTEDDMVPGILVTPKSFWVRLSERP

>CYP3118E3 F.o.

MTTVQRTLSRVLSQTTSRYRSTSAAVTTPSNEPTSQAASWDSGKPYVEIPGPKPLPVIGNVWRFVPGIGDLANLQFDQLFTVLLKKYGPICKLGGMPGRPDMLLVNDADAAQIIYRSEGTWPTRRGAYVLDHYFRDVRKNLLVSLATAEWQDFRTSVNQVMMQPRHTVRYVEPVDKMSQEFIDRIKAVRYANGQMPPDFANELGTWTLESVCYIALDTRIGIFGTPTPPPEALTMFEAVQGVFDGIYDLDIKPSLWKFVSTPAYRRFIHNMNTFTNVASKYVNKAMERLQHTPKEDLHTQQRSVLENLLLQTDDPNKAVAMALDMLLAGVDTTSAVITTILYQLSLHPEKQAQLQLELDRVIPDKTKPITKEQLEQLRYMRACIKESMRIMPILGSNFRDTGGDVVLAGYQVPKGTVTAFPLKEMFLNEKYFPRAKEFLPERWLPENKDLKTSHPFAFTPFGFGPRMCVGKRFATLEMELLTAKIFRNFSVEWNQPPAKTEFKIVLKFVSPLQFTVKDRV

>CYP6EC4 F.o.

MELTTILAAAAAVLATLYLYYSYRFSYWRKRGVPFPRPLPFLGNIGFFFAGKKGFSSAMLDLVAPFKENGYCGMYMWSQPVLLVWDPEMTRQIICKDFSSFHDRGQPSDENDPLSQHLFNLNGNKWRNLRNRLTPTFTSGKLKLMLPMMRDIASQLSSHVATEAKRADEVEIDALLSRYAVDVIGSVAFGIECNSLRDKDNEFISMSRKLGRTSAVGFVRFILIGIHPKLGALLPLKWMFSDTHRFFLNLMKETVEYREQHDVERNDFVQLMMQLRSADLANVDPANHMALTHGVMAAQALIFFLAGLDSVSNTIAFALSRLGADPELQQRLADEVREVLRRHAGELSYEALKEMDLLHRVVLEAMRLWNPAGILFRKCNATTQVGDLVVEKGQVLFIMAQVTAMDEGTFPDPHRFDPGRHTPEAKQARHPYAFLPFGDGPRNCIAERFAMLEMKLTVALLVRDFVFTPGPNYEPEVELDPKNFFPRAKNGFHLKVSARA

>CYP4PJ2 F.o.

MDAFMLAVVLVALAIFARYLKRKHLVETLDRIPGPPGYPVIGNTMDLVHFKFNLTNAVASWTAQYGDIFKIWLGNVPFVILTKPEDLEILLSSNKHIEKAELYALLHPWIGEGLLTSTGKKWHTRRKLLTPTFHFRILDQFVEVFNRCGELFVKKLLEQKEAVNVHELITLCTLDIICETAMGTKVEAQNDATSSYVTAVHKALISFHQRTMKPWLHLNAIWNMSSDGKNFNSAVTELHKYTNDVISSRRKERESRRKDSLKTDQDLGLKKREAFLDQLLDASDSGADLTDTDIREEVDTFMFEGHDTTAAALSFIMYNVAAHPEVQQRLREEMFDLFGDSDRQATVQDIQNMKYAERVVKESLRLYPSVPLFARKMREDLPLTDGYVLPAGTNVTVSCLQMGSNPKHWPEPEKFDPDRYLTENSSGRHPYAYVPFSAGPRNCIGQKFAMMEMKSALSKIVRSCDLSLPSPGYKPRIEGQVILRAPEGVFLKVAPYRAKA

>CYP3655B1 F.o.

MRKVAADIPGPPGLPLIGNLLDVVSGGDDRIYDNICRLMDRWGPTVGIWLGPELVVSVADPRDVEYVSTSNELIDKSPFYGVLEPLLGSGLSCLGGQEVRRHRKIVMPSLHLDILHDFLDTFQSESLRFAARLEQHADDGQDFDVAPLCVEYSLAAALQTIMSSELEDGDDAERLAFGGVLQSAMDLFMYRVWRPWFLHDALFRLSSRRPEYVRTMTALNGFTEGIITKKRRKLAERRRHQPHGEEQSLTAAPTTTTRRRRRAFLDNILDNEEGAALTDLELRDEVKTLLCTGSETSASTLSFVIAVLALRQDVQAKVVQELDDVFGGDASRPVTLEELPHLQYLERVVRETLRLFPVLPFFTRSCPRDVQLPSGYTAPRGAHLTFFLPALHTCPRTWPDPRRFDPDRFLPENSRGRHPFSYLPFSAGPRNCVGQRYAMMLLKVEVATLLRRFHILPAADGPAARDVADLPIKVTLVIAVKGGVRVRLRRRTAPCAAADTVGPSL

>CYP3655A10 F.o.

MLLLVLLLLVVAACLAVRSWPAVSEFRRVRRMIAELPGPPRLPIIGNALEVLSSPDEIYNRVHTMVSTYASYGPVCRFTLGHINILFVLQPADVEAVLTSPKALDKPQPYRALEPILGNGLVTLGGKVWRRHRKAISPSLHLDILRDFVPTFYKRGVVLADLLAGYADKGEVLDVTPLCGACTNNTVCETIMSSDLDEDDPQKMEFISATHKGFENWFYRVRRPWFLSEWIYSLHRTRYLEYVKLRKIFDEFVQKIVGEKKKAFNNNGNPLNPPRKRQAFLDHVLSSEESAALNEEELNDEVKTIVYAASDTSMQALSFFIYTLAIRQDIQHKVRQEINDVLGDDRGRSLQHSDLSNMQYTERVIKEVLRFFPSVPMFGRHITEDLTLPSGFTIPRGCVAGFWLPYMHRDPQSFPEPEKFDPDRFLPENIRGRHPFAYLPFSAGPRNCIGQRYAMMFLKTVVAALVPRLHFELPDDGHKRFEDIPITFNLTISISGGAKIRVRRCTTENQ

>CYP4C108 F.o.

METSGVGLLAGAILGMLVLYQILLIVTRRRRFVKVVDALPGPKPHMVFGNVPDLMVAPNKLFEAWDERHRKNATLFRTWIGPYAEINLKEPEQVEAILSSSKHITKSNAYRFLQPWLGTGLLTSTGHKWHTHRKMITPTFHFKILEAFHDVFVEKCEILARKLARVADGKTEFDMYPFITHCTLDIICETAMGVRINAQDSTEKRSDYVSAIYEVSELTLKRAQRPWLFPNFTWALTDMGKRYKHCLSILHGFTNKVIRERKEVREGQRGQQEQHSQEDDLGRKKRTAFLDLLLDAREAGAQLSDEDLREEVDTFMFEGHDTTTAGLCWAVFLLGSHPHIQDTAAEELEHIFQGSDRAPTVRDLQEMKYLERVIKETLRLFPSVPFIGRKLFQDVDFGGYKVPAGCMINIPIYHIHRNPKQWPSPHAFDPDNFLPNRVAERHPYSYVPFSAGPRNCIGQKFALMEEKTVLSYILRYYKIQAVETMEDLTLMIDLILRPESGIKVRLEPRTPLS

>CYP3655A2 F.o.

MIGVVLLLLLAAAGLAVAALLIRFSRAVSEFRRVRRMTAKLPGPPMYPVIGSALEMACSLEEVYDRLQSLTSTYGPLFRMALAHVTFTVVLHPDDMEAVMTSGKTSDKSAVYGALEPLMGTGLAILGGDVWRRHRKAITPSLHLDILRDFVQIFSKQGAAFADTLAELADKGEVFDVTPLSGACANNAICETVMSSDVDEDDPMKRAFLGAIPKALDHFMYRVWRPWFLSEWAYSLHRSQFPDYVEVKDALNLFTQRIIREKKEALKRNGPLPQRRRQAFLDHVLSSEEGAALSDAELAEEVKTIVTIASGSSMDALTFLIYTLAIRQDIQDKVRQELDDILGEDRERIVQHSDLAHMQYTERVIKEMLRFFSVVPVFARTVTEDFPLPSGFTIPRGCQVAFWLPYVHRNAEWFPEPDKFDPDRFLPDNSRGRHPFAFVPFSAGPRNCIGQRYAMMFLKTVAAAIVPRLRFEVPDDGPKRLEDVPLMVNLTMSVRGGANIRVHRR

>CYP15P1 F.o.

MAVVLWLVLLLVAVVLLRHWRQQRHLKNFPPGPPRWPVVGALPYIPGQLIHLHAHKHWRHKYGPLVGLAAGPRTMLMVCGGEEAIAVLNHDDCQTRPGGQAFRERSFGKPLGIMFSDGPYWTAQRRFTLRELRDLGLGKSKLEGVVLDEVDATLEAIERDGPELQPNKMFNSPVLNVLWWMVSGKPFARGTGSALDPQSVRLLDIMNRAMRNKRLGTAACDLWPVLKYIAPEMSAYNEIYQPIFDMQSYIREEIKNQRQNGTGDSLIGKYTEEINKSQPGSYFDEQSLITTVLDLFMAGGESTANTLSFCLMYMAMHPEVQAKVHAELDAVCSGADKAVTMSDKPKLPYVEATMFEVMRANTIAPLAIPHEATKEVEMNGYTIPQGTTILISIWATLNDKKHWGDPEVFRPERFIDSQGCFQKDPWMVPFGIGKRVCIGEGFAIQVAFLFFANLMNRFQFSLPAGSPPPSTIPEAGFTVAPQPFNVIAKPRSA

>CYP3652A1 F.o.

MAVILVLLGVVLALVLLLAALSCSQFGTWSRKGIPSPSWPLPLVGHCLRSLLQLEPLYQNFDRIYKMFPGQPYVGFYAGSSPVLLIKDIEAIKHITVKDFHVFTNHKLSPDARYDKSLSNFMFCLMDQEWKSIRVKMTPAFSSGRLKTMYPQVNAVGERFLEAIEKKKDKNGDVAVNPLCSFYGIDVIGQCAFGVESNCLLEEGRSPFLNAVEGAFRFGALEGVASSALMFSNFAYHVLGTSGVSMLKEWRTAVIRDILRESSEHREREPLANRDIFDNLLSMRKNGVPQELFESQVYSMLFAGSETSAATVMFTLWQLALNPEVQDKLRAELLEAREKDGGNLSYETMHELRYLDMVFKETLRLWPVMPWMDRVASEDYTLPGTDVQIEKGTLVFIPSFSIQRDPDIWQDPETYDPERFSPSNSDNLNKLAFIPFGAGPRACIGSRFADMSVKSVVSKTLLNYEVCLAENTPRSEAELKINKMAFIISLTEQLPLRFRKLK

>CYP3658A3 F.o.

VLDRYAGIFTRRAAAFAGQLRAEVPAGQSFNVMSKLGAFVVGSIMETAFGVDATASEVLDQLVRASEDAFFIVAYRVLHPWLLLDWLFRMSRLGRLSRHHEDVISKMTERVMEAKKAQLERRTEVAESGQQDNAENDEDDSVVREKYTFLDMALGDKAHALSDKEIIEEVRTLIAVQQTTASTLSFIMVMMALYPEVQESARAEVRELDAVQGLHPLERLKRLKYVERVIKETLRLFPITAIFARKLKKPLHLPDQKTIPAGVMVATSPYVVHRDPRHWPDPERFDPDRFLPDRCAGRHPYAYAPFSAGPRNCIGQRYAMLQMVAVVAAVLARVRVHPGEGCESRAALRVNADVFLTIEGGFNVRLEPLEGC

>CYP302A1 F.o.

MAVLLFSKYCSIIRPQTYFLKLLPPVKSNINRISLRQQIHTQNVRPFDDIPGPKSGDPAYNHERLDKAGLEKYLKYGPIVREEVSPGNNILLIFDTSDIEQLSLSENSYPSRRSHLAVEKFRKSRPTIYNSGGLLSTNGEEWWKLRRDFQKGFSAPQAVRAYLPAVDEVIQEFVTAQLCQPCEDFLPLLSRLSLQLTCLVAFDEKFDCFSPEEKRSDSRSSRLMEAASTINSLGFVYDNVESLRLSIWPKLEDASLFMEEVAVEFVQRKAESLANSRQPAENSGCVPSLFERYLTSPNFDCKDVNGMAVDLLLAGIETTAYTASFALYHLARNSDAQKKLQLEAQTLLQSATQPISGDILSEASYARAVLKEVFRMSPISIGVGRLLNKDLVLSGYQIPKGTNVVTQNQVACRLEKYFDRPNDFLPERWLKESPRQEKVNPFLVLPFGHGRRSCIARRLAEQQLLTFLLRVARTHCVTWNSSEPLGCISVPINKPDQPVKLVFSRL

>CYP15P2 F.o.

MLALALLALLLLALFALFNYLGNLRRLKNFPPGPLRLPLVGSMPFFPAEHWHLYADKHWLSKYGPLVGLTTGRDTMLIVCGAQEVLDVLAHEDCQGRADLFFTTERSFGKKLGIMFSDGPYWTEQRRFALRELRDLGMGKSVMEGVVLDEVDATLEAVARDGPAVQPNMLFHAPVLNVLWWLVSGKPFSKGVGKEQDAQSVRLHHIMEKFIRTKNVAVAPVDSWPFLKYVAPELSSYNAIYGPIFDMQAFIREEIKLQQQDRTGSFMSKYMDRIETAEPGSTLTEESLIISVLDMFIAGGESTANTLSFCLLYMALHPDKQAKLHAELDAVVGGADKTVGLADRTRLPYTEATLTEVMRINPIAPLSVPHSNTKDVELNGYTIPKGTMINLSLWVVLNDKAHWGDPEVFRPERFIGKDGAFVKDPWMANFGHGKRVCIGEAFTVQTAFLFFANVMNRLRISAPPGGEPLSTKPQPGLTCAPQNYTVLAEPRRSSSAHQM

>CYP4C109 F.o.

MSPLTAVALAVLVALLTARWWRRRRLVALIERIPGPFALPILGNTLEQTVEHDELFGRLNGITQLFGRQHGICRTWFCSQPYVVLSSPAAVEAVLGSNRLTDKSDEYLYLRPWLGTGLLTSSGAKWHRRRKVLTPAFHFRILDDFIDVFREQAAVLADKMEALAGDPEGFNVFPLVTLCTLDIICETAMGRKVNAQVDSESPYVRAVYDMAEIVLTRQSTVWYQPDWLFHLTPMYRRQQRCLADLHGFSTKVIRERKAEILDEARLKGRPGRGGEDDLGSKRRLAFLDLLIEASQGGAVLSDDDIREEVDTFMFEGHDTTSAAISWCLFLIGCDPHVQRQVHAELDDIFADDPGRGVTMKDLAEMKYLECCIKEALRLYPSVPAFARSLREDTQIGPYTVPAGTTAMLVIYMLHRNAEVFPEPEVFDPDRFLPDNCVGRHPFAYVPFSAGPRNCIGQKFALLEEKSVLSALFRRLRFESLDRREDLTLHGELIIRPKDGLRVRAHRRA

>CYP3118F1 F.o. (2)

MATRHVNEVVDRLSKLSEEEIASRQFSVLERLLVNSHDSDKGVAFALDMMLAGIDTTAHQTSSLLYFLATHPEQQRALQAELDREWEAGQPLTASVLEKLKYLRACIKEAMRLQAVVTGILRTANQDVVLNGYLIPKGTHTIVCTKTMCRDERLVPRASEYLPERWLPGQEQLKPRHAFVNIPFGFGPRMCAGKRFADLEMEVLCARLFRSHNLEWHHPEPQVEQKLFVQYSSPLKITMKER

>CYP6EB3 F.o.

MLTAALLVLATLLASIWIYISWHFNYWEKRGVKYNNPFSFFGDRGAGLPFGKPFWSLLDPVYNKYRGKERFVGTFQARQPVLVVLDPDLVKTILTKEFNAFQGRGQPFDEEGDPLSANLFNLVGPRWKNLRSKLSPTFTSGKLRLMVPLMTDINKELTARVSSLSKEQGEIEMKDLLLRYATDVIGSVAFGISCNSLKGEIDEFYKMTREVFNRNILFIIRFFLVSIHPIFIKLMPFKSLFDKTTNFFIKLMKDTVEYRETNKIERNDFVNLMMQLREADRHETDRVNHIEFNHNVMTGQAFLFFIAGLDAIANSVGFALHELMLNPELQERAAAEVRAMVAKHDGMTYEAIRGMDLIERIMREVLRKWGPVGILTRQANDTFKIPDSDVVIDSRVLVQIPVWQLGHDPQYFPEPDRFDPDRFTDEAKEQRNSYAYLPFGEGPRFCIAERFAVLEMKLCLAGLLEKFVFSVGDKTDVKMKLNIKQFSPTPANGFWVRVEDRP

>CYP6QT1 F.o.

MAVLEVLAGLVLLAALLYYYATATFSHWSSRGVPHLPPDPLFGNIKDIVLFRDLHVYGYQKLYHRFDGLPYAGLYQMRTPSLMLRDPETIKQFLVKDFGHFHDRGIFCDVERDPMSATLVNLSGRHWRNLRNKLTPSFSAVKLKSMTPLLNECADALVALVGAGQEGREHQVEMREVMAKFTTDVIGTCAFGLHFNTLKDPDSQFREMGRRVFLPTYSRTAIHLLRVFFPGLLGVLRLRTVSREITDFFISLVRDVIGFRERNGEVRNDFMQLLMQLRQQEQLDKSDKDSKEDAVVLDDRLLTAQVFIFFVAGFETSSSTMSFCVYELAINEEVQELARQEVDAILSELDDGQPITYEHVAKMTYIEKVLLETMRKYPPVPGLVRVCTKPYTLPGTKVHMKVGDQVFIPTFAIQRDPNIFPEPDRFDPERFSEDGRKSWHPFAYLPFGEGPRICIGLRFAMVEMKMGLARLLQHYKLLPAPGLPVPMRFDPKAFVLTAEGGINVRVVPRSAAAGSAS

>CYP3658A1 F.o.

MISALTIVLLALPLVLYVVDKVRRTSKLAHLAGPSYPLPLLGHLKDMFGPRATFLDRCNAFLQRWGPEPVKFWVGETPSVSLTRPEDLEPLLSSQREVFKPDQMYKAVSGFFGDGLITLNGDQWFAHRRALTPAFHFKVLERYAGIFTRRAAALAEQLSSEVPAGQSFNIVRPLGAFVIGSIMETAFGIDAATDESHRSDTMDRLVRATEDAFFIVMHRIFHPWLLIDFVFKLTPHGRVARDHEDIICDMTRRVIAAKKAQLKQGAKDAAADQDDVKGAVDDDDGVRKKFTFLDMALGDKSLVLSDKEIIEEVRTLIAVQQTTASTLSFIAVMLALNPEVQERARAEVREVDAVEGLLPLDRLKRLKYVERVIKETLRLFPVTAAFGRKLKSEVKLPTQTIPAGVVINMFVYRTHRDPRHWPEPERFDPDRFLPEQCAGRHPYAYVPFSAGPRNCIGQRYAMLQMLAVVSAWLARFRLLPGDGCESREALRVNMDIFLCIEGGANVRLEPLEA

>CYP3655A11 F.o.

MATALLLCVVAALAWATGAVSGLLHLLRGLAGLLRGLAELFRVRRLTAALPGPGLTALLQRALRGGGRASRRAGTYEVLTELCSRYAASPAFRVCVGHATTVFLLRPDDIGAVLSSTRFATKSSLYSALEPLIGSGLAILAGEDYRRHRKAVTPSLHLDVLKEFAPVFYKHGQVLADSLAARDGSVLDVVSLCGDCANAAICETVMTTDVADDDLGRQGFLAAIPEANKHFMFRLSRPWFLGEWAFKLSGRYAAYLDTKRALDDFTERIIREKKSRLVAASTGTPPCAAPSTTRPPGGRLAFLDHVLCSAEGAALGDDELAAEVKTLVAIASGSSMDTLSLFLQTIAIRQDIQEAIWKEVTDVVGEDASRALSAADLQRLQYTERVFKEVLRFYSVVPLFARRSPEDVRLPSGVLVPRGCHVALWLPHVHRCAEHFQDPDEFDPDRFLPERARGRHPYAFLAFSAGPRSCVGQRYALHFLKAVAAALVLRLRFAVPAGGPSRPQDVPLLLNLTTCVRGGAKVLVRVRPPRPPPPGLPPRPTTPREGQGVH

>CYP6KF3 F.o.

MAVLEILAAAAALTLLLYYYMTSTADFFAKRDIPHLPQTPLVGSSLDALLSRRFRGQVWREWYDHAKAHGHKFYGVFVGRRPTLFVCDVDMVKTILVKDFQHVMDRGTYYDRKREPLSANLFNIGGGEWKALRSKLTPTFTSGKMKSMFYLVRACTDELERYLGAEVLKGGDVEMAEVMGKFSTDVIASCAFGVEANALKDPDSEFRSMGRKVVEVDPLRAIVNMVAETMPWLRPLLPMPLLRPDVKDFFIRTFNENVEYRERQKVARHDFIDLLIQLKNDKTAGLEDAVLPAQAFVFYLAGFETSSAATSTCLVELAHHPDIQERLRDEVDRVLAKHGGITYEALHDMTYMEQCIEETMRLYPALAGLPRVVTRDYQLPGESPRAVLPAGTRVIIPTYAIHCDPEFYPDPMRFDPERFTEENKRSRPHYAYLPFGEGPRICIGLRFAMMQMKTSLAMILSSYRVFPSPKSVYPARFEPKTFVTKLKGGNRLKITKR

>CYP F.o. [7]

MGILDEMKVYTGTAIKTTVSALSFICKVLSPLPDAQERLHRELDEVFDGSSRPVLVDDQPRSKVSRREAEVVVPPRGNSNASAPYSARMEFTERFIKETMRLFPRPCRSPPARCTATRSWPATSCPPAPP

>CYP301A1 F.o.

MAVRGLRGMLRSGVRPRSSTTATAPLCPREAEVHDADPVAVDGLPYEQMPGPRPLPLIGNTWRLLPVVGQYQVSDLARVSALLHEQYGDCVKLSNLVGRPDLVFVFDADETERVYRHEGPTPFRPAMPCLVSYKSDVRKDFFGYLAGVVGVHGEAWREFRSRVQRPCLQPKTVRSYIGPIEDVTDCFLQRMQDMRDDRGEMPADFDNEIHKWSLECIGRVALDTRLGCLDPNLPPDSEPQKIIDAAKFALRNVAVLELKFPFWRYIPTPLWTHYVNNMNFFVEVCTKHIDRAMERLATKEDGELSVVERILAEQTDRKLAYVLALDLILVGIDTISMAVCSILYQLATRPEVQDKMFAELERVLPDPDTPLTARHLDQLAYTKGFVKEVLRMYSTVIGNGRTLTQDMTICGYRIPKGVQLVFPTVVTGNMPRYAGRADEFLPERWIKGHDASLDVHPFASLPYGYGARMCLGRRFADLEMHVLLAKLVRKFRLEYHHRPLDYQVTFMYAPEGELRFKLVER

>CYP301B1 F.o.

MAGPAAWRSLLARGPRPLLGHVVRPSTSTATPSSAAGDDAVLHGSVNDVRGYDEVPGPRPVPFLGNSWRFLPVVGNYRIEQMDKVCLGLHRQYGDIVKVAGLLGRPDMVFVFDADLIEQVFRGEDALLPVRPSMPSLDAFKHGIRKDFFGDLAGVIAVHGPKWLEFRTRVQQVMLQPRTAKLYVGAIQDTADSFVRRVRRLRSPAGDAPPDFVNEIHKWSLESIARVALDARLGCLDDRPPADTQALIDAVNTFFMNVGVLELKPPLWKIFPTPTWKAYIRALDDITNITSGYITRALDALRETEHAELGPDASLLQRVLASHDARTAHILAIDLFLVGIDTTSAALSSALYQLALHPEQQEKLHGEVVRVLGPDGAVTASALEDMPYLRAVLKEVLRMYPVVIGNGRTLTEDTVVGGYLIPKGTQIIFQHYVASNQDRYFPEAGRFRPERWLREHQAACPAHPFASLPFGYGRRMCIGRRFAELELHTVVAKMVQNFRMEYNKAPLPYRVHPMYMPHGPLQLTLHAR

>CYP3657E1 F.o.

MAWSWASLLAAAAVVALPLLVQWYQDVRRMAKVAASIPGPPTLPVVGNFLNISRSRLQRSFQYGTLFRLWFGPLLAVWVVDAEDVEAVLSSPACAHKPRIMYRLIEPILGRGLLVLNGAEHRRHRKSILPSLHREVIQQFQPLMQGVAMELVDNLRPKADSGEVFDVVQLCSVAAIDSAFRTILNASGELFGPELRREVVEHVDTLSSVLMYRAVRPWYHWDWLFSWDSHYAAYQRVNELYDGLILDVMNHKMGGDGKSTGTPPRGRKAFLDHVLASEGGRQFTREELYGELKTMLAASFVTSMDSLSIHFLVLSIMPDVQDRIHQELDDVLGADRPLEEGDVEHLEYLDRFVKEVMRYFPTFPAFGRRCLRDLRLPSGHTLPAGCFVALSPLASHHNPKYYPEPARFDPDRFLPAAVAARPASAFMPFSAGPRNCIGGRYAMTFLKTQLACVLRRYAVLPDPDGPRDVSRIRMKMGVTYYPRDGARIRLRARAPHGASAG

>CYP3657C1 F.o.

MSLLVLAVVVLACARYGPRAFRVLSGIVRTIWKSRTIPGPVEGMLSFYGSTDEFIHKVRATSQRFGNTVRIWAWPVVCIFVSEPEDLEVIFSSPYLQSKAPIVYDALTPILGKGLATLNGAEHRKHRKALGPSLHLEILQGFVPIFEHNAQKLCKQLEAYASTGSTFNVSPLLGRYSAQGICETVFSTETSPGLAEEQENFIKVLIEASDLMFYRLTRPWYSPDWLFYFSSQYKGYMGAVKGFESFVSKTLAHKVELVKRGVTPKEGKRKAFLDHYLTSDEAKVLTERELIESLKTLSAGAVGTSMDLMSFFLLIMAITPDVQDKVAKELDDVFGGSDRPIEPGDLSHLPYLECAVKETLRMFPPLFAFSRIAKRDVKLPSGHELPKGCVVTVMPYTTHRNPKYFPNPEKFDPTRFMPENSRERHPFAYIPFSAGMRNCIGQRYGMMSAKIVASTILRKYRILPCPNGPQRLEDFVVGVGVTFGLRDGAHIRVERRLKSDLKRRG

>CYP314A1 F.o.

MIFERSFDLARMLSAVDLVALLLFGVVLLCSEFRTKWGWLRLRKSKALSPANTPPPSPKPKTEPRRKTVQDIPGPWPSLPVLGTRWIYSLGVYKMDKIHEAYEDMFNRYGPVVREEALWQIPVVSILERSAIETVLRSSTKYPLRPPTEVTAHYRQSRPDRYTNLGLVNEQGETWHQLRSVLTPELTSAKTMLRFLPELNTVASDMTTLLAASRDSNGVITRFEELANRLGLESTCTLILGRRMGFLDEKVDPQAAKLAAAVKVHFCASRDTFYGLPFWKVMPTKAYKELVESEETIYDIISGLVDAALAEEQQTAQVDAVQSVFLAVLNAPELDIRDKKAAIIDFIAAGIQTLGNTLVFVLYLIAKHPHVQKRLYEELIAAAPAGSPWTAQNLRNAPYLKACIMEAFRVLPTAPCVARIIDTDMVLSGYHLNAGSVVLCHTWLAGLKESNFPAAHEYRPERWLNGGPGSAATFLVLPFGCGRRMCPGKRFVEQALQVVVAQTIRDFDVGFDGDLGLQFEFLLTPQGPASFTFRDRV

>CYP3661A1 F.o. (2)

GGRFKTNSACRLEYTERFIKEALRLFPPVPLTGRQVHRDTTLASYRLPAGTTLLLNVFGAHRDPLHWPDPLLFDPDRFLPERVRGRHPCAYVPFSSGARNCIGSRYAMMNLKAFLATVLRALRVDRADDPYTDIHQLPLTADLSLRIVGGPRVVFARRAEEAM

>CYP3657B3 F.o.

MAALLTLGLSLACVALLAPWALRLASHLWRVARQVVLTLGIPGPASVPLLGTLPTFLRFWEDMEGTLLDLMHEYGPTVRFRLADRVAVLVMHPEDVQAVCTHPALVHKADNLIRFLRPFVGDGLLLLNGKEHRQHRKAISPSLHFDILRDFVAVFDKNSRTLAQRLEAHADAGHVFDVHLEFGRLTSTTLQETVLSVGVDQGYDKAGRVFHEVGDIAMWRGMRPWWQNDAVFRLLCPRHEQHKRAKDDMDAIVSRVLAVKGAELAAGAPAPPRRRMAFLDHVLRSPDAKAMSEPELRAELKTLLFAGSATSMDFLSYLSVVLTILPDVQARLQQEVDAVFGPPGAAGASRPLLPEDLPHLDYTERVVKEALRLAPPVPMLFRQASHDVTLPRGAFIPSGTVVILVAAGTHRMA

>CYP6 F.o. [964 (2)]

FYVDMLRESAAFRARTRTRRTDFVDLMLSAAAPSRAQRGLSEQHDFLTAKDRRRHCALAYDVVAAQAFVLPAHAPRVVARCLRELAGLPDLQQRVAQEVRSARDPDQPLGLQVLRRAPLLEKVLLEALRLEPPCSALSTEVGAGGYALPVVGPRRGLLLEPGTHLYVAVDAVHRDADLYPDPLRFDPERFSAAARRARPTGAYLPFGELPGPDGVAEWAASGLPPPASSTAALFALQEMGLCLAALLADLEFGV

>CYP6EC3 F.o.

MDLTTILAAVAAVLVPTFMYLHYQYRYSHWKKRGVPHPKPLPLVGNFAFMITRSKSFSTAFMDFATPYKENGYCGVYQFGQPLLLVWDPEMVKQVVTKDFSSFHDRGLPSHEHDPLSQHLFNLSGTKWRNLRNRLSPSFTSGKLKLMFPLMRDIGDELNRQVTLDAKKTDSHEVEISALLSRYATDVIGSVAFGIQCNCLRDQQNEFLEMSKKLFRQSPAQLARLLLELIHPKLGGLLPIKWVFSRVHHFFVNLIKDTVEYREKNNVERNDFVQLMMQLRTEDLAHVDPENHIELTYGVMAAQGFVFFIAGLDNVANTISFALNKLSVDPELQQKLADEVRGVLRQHDGELTYAALKQMDLLNRVLLEAMRLWSPVGMLIRKCNATTKVGDVVVDKGQMVFVLTQVAAMDENQFPEPQRFDPDRHTREAKDARHPYAFSPFGEGPRNCIAERFALLEMRLALALLIRDFVFTPGPRYEPEVELDQKSFFPRPKNGFHLQVAARV

Two files omitted from CYP nomenclature pipeline:

>Scaffold97:956479-957594

LQIILGSSQHTEKADAYSGLVPWIGEGLLTSNGNKWRSHRKLLTPTFHLRFLKHFDLIVF

VKKLLENKRPVEVHSFLSLCTLDIIC

>Scaffold39:1168373-1169995

MESVLLVAVLVVLLISRWGRVMLGPYWWGPPAFRKAMDAFPGPRPTLPLLGNLVQLAFGG

KVLHNGLAIIGRWSPSAFSFHLGPVPMVVLTRPEDYEAVIGKLTQKATYIYSTVESFFGV

GLATLNGEAWRVHRKHITHAFHFRILERYVDVFERKGREFGDRVQRLADGATSFNVFPHL

AFTANDTIFETAFGLDKSTNSAVSAAHRQEFVDAMEESFEQLQYRVLHPWLLMDWLYRLT

PAGRRFYEAVDMIDGFAQSVIDDRRIKMRDGAAAGYSFLDLMMKIQPGEDAVVLNDAELR

GELRTFISVQQTSATIMSFALILLAVHPDIQERVVDEAVAELGPEGGVTYASLTGLKYLE

RVLKETLRMYPVLPAMARDVTEDAQLTDGVVPAGASIAMVPIVTHRDPALWEEPTRFDPD

RFLPERCTGRHPYSYLPFSAGPRNCVGQKYALLQMKAVMSTIVRRFEVLPGKGCGTMAEL

EKNLDVITFLTVSGGFNIRMRPRAPRIEPAACSPPPSFTDSVLRRDFGAGATLPAHASGR
