## Additional file 7 for "Genome-enabled insights into the biology of thrips as crop pests"

**Additional file 7: ATP-binding Cassette Transporter (ABC) gene family and carboxylesterase protein sequences (complete) of Frankliniella occidentalis as part of the FOCC Genome Paper.**

*Contributed by Wannes Dermauw, Simon Snoeck, and Thomas Van Leeuwen*

>FoABCA-01

MDGTCCSQTKALLVRNFLIKIRARRTFLIELFTPLIYLVVPVIIFVYINMYELPPILPHP

GILESFRIFDRFLLIRNHTIAVAPNTSETQEFLKAVNNMYMKVQTEQDGAEGKPQTPINF

LMFNNNEELSDSYWKEPQSFQLGLVFEDDQPISSNLKYEIRGNPQFSVTPPTTDMFSDSD

SCLGQGPIKRQWSGIFPDPGETCPAQQYYLSGFLPLQTLVDFTKITLDSGVSDLSIPGIM

LELLPRSVRETKLTPLEEAATLKDLILLILVPVLMVMAMSQYTTYLIVLIVNEKENHSLA

AMKTMGVKDSAYWFAWFVIYTVLVAALALVVVLLLYLSDILPRSSYTLLFLNVTAYGCSS

IMITFVLSTFFNNARTAGLLASFVTTLLSLVFLLETGLDVHSTAGKCILGLLSPLAFAKV

LYLAIAMEMNGVGLTWVTMWEGPQPTPGVYFLLQFVDILLYGLLAYYLDNVLPNEHSIRR

SPLFFLNPMFWCNKKSVRATATANGEPFTYLTTEETESGDIEPVPPELKGREAIRIEDLH

KTFNTCFKPGVKAVNGINLTIYEGQITAILGHNGAGKTTLFNILLGCTAPTSGTAYIYGY

DIRNANDMQVVRRMTGVCPQFDILFKTLTAREHLHFYGTVRGIPASKLDYEIEHTLADLD

LLEKADFVSANLSGGQKRKLCIGIAIIGDPKIIVLDEPSAGVDPISRRHIWSVLQSRKAG

KVILLTTHFMDEADILAERKAVISKGRIRCCGSSLFLKNKFGIGYHLTLVLQSDAHEVAI

AQLVQRYVHRAERARRHGRELSFILPHNAVNSFPDLFAAIEREVNTKRGLGVTSYGVAMT

TLEEVFLHLERDEDEAETNTVMENLSRSMLRNRAMSRSLSIPSKSTSYHSLQNEAAASTV

TTDSEINTDGTLSPTEHPLRLPIEVCPNNTQKLFAMVKLRFWCLIRDTYKLVNLVAVPVL

LSMLLLYRSAGHNRPPEMPSLRLDGGTYTSMKVALIASDFAGRESPQTDLLLGGLNEAQA

TGVDVYTGNASQVNLTSPHLLRLKMALGDEQENPGAIQVSAAYNDTAQHSLPIVMNLITN

SIYRMVMANKRGRDIARNTQPIITKSHPFDRQNTSSTVLSYALTAFSVGFTFIMVAATAG

VDMVYDREIRAKTQMRVNGLSFPIYFISFFIVMFVVLLIVFILLVLVAYLCGVAWLDTLP

AVTIFGILMSLFCPCAIILATTLSYCFHKTESAASILPGVMLFPGIIPYFTIVFVKSHKV

KEVLHIINSLLNPFYLPFGAVWQMIDRYLICMESTRCRREGNPFSNYNTKDIGVIIAGSI

ITLVVWSFILFVVDLKSTGASLRDRFRGVRGSVGACDGGESLDCGELCGDDDVRAERSKV

SRLMVDPPDQPPVVIVQNLRKEYDKDRVQSKCCDCCVVKGKTHKLALKELSLAIDTGEVF

GLLGHNGAGKTTTMKIIIAEEAASRGRVQIAGYDIATNMNEAFQKLGYCPQHDAQWKKIS

CREHLELYAAIRGVPSKDIPRLSDAFLAGLQMKEHANKPAAACSGGTRRKLSFAMSMVGN

PAVVLMDEPSTGMDPRSKRFLWDTILASFQGTRGAILTTHAMEEADALCSRVGIMVKGEL

KCIGPTQHLKNLYGAGYTLEMKLKDLSSIKTVSERHEGVLAYVTNLFKDSKIVESFSDRL

VFEIPQKDVSSVAQCFRNLEKAKRDLNIIEYSFSQTTLEQVFLKFSHDDGDEDD

>FoABCA-02

MGFFLQLKLLLWKNLQIRKRNKVRLLLEIIWPLFLFLILMWVRTRGLKEVRQECHYMEKA

LPSAGQFAFFHSLICNAPNRCHFNNGTSGPPYDQYRPRPKLTDFTNEFGRLLLDGGKLPR

LIKNYGDLGFILSSVERNQSLGDDFSLRVRDLALPHQIADIPPEWMDSPIDARKLDKAML

VDPPSAHYCIKSGETAAREVICNENMTFIWEDVSSSAETYRQENMESVLRPSAWHRFQVA

VRSMADDLDTIGFDNVDPKALVILDHLRGQGDANNPPEQGWCDLMEAFDDFRIPENEFTR

AMRGEQLQFKSRYKADPEEPDECRAIFNSTAIKKIPGVAAVWKVIKPFVRGKILFSPDTP

AVRSVMEKVNRTFNELLFLPGFEAVDTDAFFGRMEEGFFVREMPKSAEGKSLFRATFKHN

VFALSPVLLRRSGQLRRLQEVAGRPLSKPRREGDRNHTAEQEDSMGTLFDEGFQLLGRFL

KCLDRDKIAPVPLEEAEEEAVKLARTNNLWALVVFDEAGNDTLTPFVKYKLRLSVERVDS

TIDIMDRRSNYRSRDMPLSDLKYMTYGFAFLQDMIDSAILEVHTGRAGQANPGRILKQYP

YPEYIYDQFIKAISNTFPMFMVLAWVFTSALVVKSIVYEKEQRIKETLGVMGLSNGVHWA

GWFIDTFLVMLTSLTCLSLILVYGKVIERASLSVVWVFLLSFSVATISFSFLISVFFSRA

NVAAAAAGIIYFATHMPYPLIQRFSDMIPYGGFVGVSLFSNIAFGLGANVFASYEQTGEG

VHWHNINLPANADELVTIGLCITMLWVDALVYAVLTWYIEAVFPGQYGIPKPYYFFLQRS

YWFGSLAQESMNGHLPNGGIVANNMEPEPVGGVVGVAIDKLSKVFGGKTAVNGLSLNFYQ

DQITCFLGHNGAGKTTTISMLTGMYPPSSGTARIYGRDIRHDMDSIRSSMGVCPQHNVLF

DSLTVEEHLLFFGRIKGVEPEVLRREIDGMLHDMGLEGKRHSLSRDLSGGMKRKLSVGMA

FVGGSTTVFLDEPTAGVDPFSRRAIWELLIKYRKGRTVILTTHFMDEADLLGDRIAIISG

GRLECCGSSVFLKARFGSGYYLTVEWLPQQKEEQDHSIRASRLRQLVTGTVASAQAVEEM

ASESVFVLPDRHDVPAITALLEKMDENTEALGIESYGISDTSLEEIFLKVIEQKEPVANG

NEAVASNDATGPPLPLRDQDDPASAPAEYVGGLALLRQQYLALSRKRFNYVRRDPKGLFT

QLVLPAVFVALSLTFTSFYNIPTTMPELVLEPKLYGEPVVGFWQVRDEASDLTQSALHLQ

SMRRDGVGSELVPSRRRLASPELPVRVQRRMELAKSIASADNTCANSPPQIDGVPQSRMI

LPGKELYYNGTGVELEQWIVDTYPDCKRNRYGGVTYGASYPWSRESNLPIAGFNNKPDNL

RIWYDNKGHAASPSYLNAMNNVVLRSLLPEDKKNASRQYGILTSNHPLPLTPEQYKDKTM

NEAFIVLFHAVSVIFAMSFVPASFVIFLIEDRVAQSKHLQLVSGLRPFVYWLQSYTWDLV

SFAVSETMVVLIFIAFQSEMYTSGTNLFALIVLIFFYGMACIPLLYLLSWFFHVPSLAFV

VLSCGNLFIGMITTLTVLVLGMFEELELQHIRDVLEVVFLVFPQYCLGQGLMNMATNYMN

GQILAKQDIIWDKSVLSWAVTGKFLFTLLVIGVVLVVAVFLAEQSTHWFHRGGSDAAASS

VPREGDVQREFIRVDSGQADGDVLVLKRLTKSYGQGLPPAVDGVSFGVTAGECFGLLGLN

GAGKTSIFKMLTGDTSISGGDALVAGTSVRADMDGVRKLVGYCPQFDALIPLLTVREHLD

LYCSLRGMRPERRQRAVDRALRRLNLGFYQDKQAKSLSGGNKRKLSTAIALVGDPALLYM

DEPTTSMDPAARRFLWDCVRGVVASGRSVVLTSHSMEECQALCGKLTIMVNGQLTCFGSP

QHLKSKYGSGYSVVVRCQPKRVVEVRELVQRRLAGSTVVEEHLNQIKFHVPPAAESNLRL

VTIFRTMDQARTESGVLDYSVSQTTLEDVFMRFAQMQKTEEVAPSEAADCLDVLKKCFGP

LRKREKEAAIP

>FoABCA-03

MGYLHKLRLLLWKNFLIRRRNKTRLVIELVYPLLLFVIIVALKPSSSGTTSRANECHYKE

KAMPSAGQLPFLHSFIFNAPNLCSGDGSDPFLRSPRSVGGSHMHRTRAGTSSGNGTVPDE

SEECKTVGEVIRELTVFLDWDVMKPFITGKILYYPDTKATRAIMKEVNNTFNELTDFSTL

PPELLEVLLKMRPYLRPVFKTFGELFEKCVNREKVQAVTSENALKEAFQLQRANNLWALV

YFDEAGNDDLAPFVKYSIRMDQTRIGSVKTSSSFGSGAKHLDYPLTDLKSITYGFAYLQD

MIDSAVIGLHTGRTAKMRPGRLLKQFPTPAIEVDTLASMLTQLMPLMMVLAWVYTFSQTV

KSVVLEKQERVKEVLAVMGMSRGVQWSGWVVEGLRSSIITCIILSVLIKYGGLAERSDVT

IILLLLVSFVVAMLSFAMLVSTFFSRATLAGPSAGVIYFLTFLAYQLVENSPTEPSYSAV

LASSLFSNVALAEGSLVLAKMEESMVGVQWSTINDPAGPGKPATMAGAIGMLWLDAVLYA

VLAWYIENVFPGEYGVPKPFYFFLQKSYWLGTPSSQDAITLSAEHVDPANADFLEPEPEN

GVVGVSIDKLGKVFGKKAAVNGLSLNFFENQINCFLGHNGAGKTTTISMLTGMYPPTSGT

ARLYGLDIRKDMDAIRANMGVCPQHNVLFNSLTVEEHLLFFGRIKGMSEKELKGDIDMML

SEMDLEKKRKAFANTLSGGMKRKLSVGMAFVGGSKVVFLDEPTAGVDPFSRRGIWELLVK

FRKGRTIILTTHFMDEADLLGDRIAIMSEGQLQCCGSSVFLKERLGSGYHLTIELEEAKP

YDLERKERLLREVQSVMPTARPQDRDDVEEAEVVYTLPDRHDVAAITALFERLDERREQL

GIASYAISDTSLEEIFLRVAERKAEADAAGSISDVVNVGDDDGSSSELLVDSLSGVSLLW

HQYVALTRKRFNYVRRDPKGLFTQLVLPAIFVMLALALSSSTGSGTPPAIQMQPSLLSWP

AVSFFEQTQGDDWGDKYAKSLKSDGLGPSLVAGGTVDKDMDLTRRTPDYHCTNSPPDPQK

AEWSVVLPSKELVYDLTGSDTETWLMNTNTACQRNRFGGVTFGAQYNGTTEYGQFGQADN

ARIWFSTKGYASVPAYQNAINNVILRASLPAGKTPRDYGILTNSWALPADPDADDDNKID

PVEVSVVLSSAITVIFAMSFVPASFVMYLVEERVEQSKHLQLVSGLRPYIYWLQNYTWDL

VNFTAAEIFVILIFVAFQTPMYVSGKNLTALIFLIFLYGMSCTPLLYLFHWFFRIPSMAF

LVLSTGNLFIGLVGTVTMMVMTVTSTAATRQVVETLFLMLPQYCLGQGLMNMAMNHFQSL

ILEMQGQPGISPLDWEVTGKHMVAMASMSAVLAVAVFLAEQSTHWWHGWAQGSRNEVAEA

DANKEDDVLREEQRVAEGRADQGMLVLKKITKRYNRKAPPAVAGVSLGVERGQCFGLLGL

NGAGKTTIFKMLTGDTLISGGDALVSGRSVRADMDSVRRMTGYCPQFDALVPLLTVREHL

DLYSSLRGTPDRHRPLAVRRAIQLFDLGAHKDKNAKDLSGGNKRKLSAAIALVGEPSLLY

MDEPTTSMDPVARRFVWQRVRRVVASGRSVLLTSHSMEECEALCARLTIMVNGKLTCIGS

PHHLKHKYGSGYLAVVRCSLPRVHEATELLLQRLQGASVVESHLHQIKLQLPPHLSLIAI

FTALDEARRGGSVEDYSVTQTTLEDVFMRFARAQRAEHP

>FoABCB-01

MSEVRAAAEPEWSPLLARTLPQPDDLQGGLRPRLLPDKPRRRRGKAVLDDDDDDPATRKS

GISGLFRYASRRDKLLMFVGLLFATVHGASFPVLALVFGQMTNTFILQSTVTSVQKKQNE

SVVDGEMFGVSPTTPLSTITDPDTTSVPDMNEMWRSGALSPSEFTAYMSQFSLYYLYIGI

GVLAAAFIQTVCWELACEGQVHSLRKIFYAQILRQDISWYDQNQENDLTSKLSDDLERVR

EGIGSKFSMVIQYVATFVSGLAVGLYANWRLTVVILGVGPLLIGTSGFLARVAASTAARE

QLKYSIAGGIADEVLNCIRTVAAFGAQAREVNRYEKALEKGRWMAMKKYYVLSVGLGVVF

FVTYASYGLAFWYGAELIGAGVVTPGSVFTVFFSVMAGAFSLGNALPFVNAVSTAVGAAS

NIFNVIDRVPSIDPYDTRGLKPEKVKGHIEFRMVGFSYPARPEIKVLQDFTVSIESGKTV

ALVGQSGAGKSTVVGLLLRFYDATIGRILLDGVDIKELNLSWLRSQIGVVSQEPVLFATS

IYENIRYGRDNVSRAEVVQAALIANAHSFITTLPDGYETMVGDRGAQLSGGQKQRIAIAR

ALVRNPKILLLDEATSALDAHSEGVVQAALDSAMQGRTTIIIAHRLSTVKNADLIYAMKN

GAIEEFGTHKELMSREGLYHNLVMAQMSDKNAAPHDYSSGGEALGTGGESYEAIQSSVEK

KFRKRRSLSLSLASLDDPELDRLAKEAEVADVSANVSVSRLFRLNAPEWVWLLLGFIGCA

LTGSIMPIFAFFYGEVFATFTLRGEELREAALFWTYMFLVLAVASGFSFWLQLVAMTTAA

EKLVMRMRLFAFKNICDQAVGWFDLESSSAGRLINRLARDAPLVKAAAGLRAGQVIGAIV

TLLAALVIAFIFGWKLALLLVVAVPFIAGASYQQMMILRRNQRRDAELMNDAARVASESV

TNIRTVQSLGKERLFFELYLGYLSAPFVEAKKQAIIYSIVFALSQAVIYMMYAGAFRFGA

YLIEVGDMAPTDVYRVFFALAFCAASVGQTSAYLQDYAKAKLAASLMFQLIDRKPDIDCS

SATGIRPVIAGKVTFKDVKFHYPARPDVPVLRGLSFSVEPGQTLALVGSSGCGKSTAVAL

LERFYDPHFGSVMVDDIDIRTINLPHLRAHIGLVTQEPVLFDCSIRHNIAYGKAFSSRDS

LSYLATGDEVLDTVPMSEIIEAAKAANIHNFVSSLPQGYETMAGDRGTQLSGGQKQRIAI

ARALLRNPKILLLDEATSALDTESEKIVQEALDHARKGRTCITIAHRLSTVQNADCIAVI

HNGRIAEMGTHEELKARRGRYYQLIKRQRL

>FoABCB-02

MISFTKYLRNSCINCHLHRSFNRSYITNRIGHLNGSHNTKTDVFGLRSNYVTIRPNFALF

RNFKSGTWKFYSTKAAQSAASSGSKGKQFDKRAIERLLSFAKPERWKIFGAVSVLVVSSA

VSMFVPFAVGKLIDMIGNQLKSTSEAEREQNQRELNKFCGTLIVIFIIGGACNAARVYLM

SVAGQRISKALREKVYKAMIRQETAFFDQRKTGELLSRLSSDTLILSDLLSTKVSDGLRS

IISTSAGLSMMFFTSTKLSFIALAGVVPPIAFLLVVVGRYLRKISKEVQGSLAAANQVAE

ERFSNIRVVKTFGHEKYESSLYTQKLADLIELARKEAKVRAMFFGLTGLSGNIILVSSLY

IGAPMLIDSTLTVGQLSSFLLYAGSVSLSASGIANLYAEVQKGLGVSGPLFEVLDKKPHI

AYEGGLIPEGKPQGQLDFSNVSFAYPTRSDVTVLNGLNLTIPAGAMTAVVGGSGSGKSTI

ASLLLRFYDPLSGHISFDGRNVSELDWSWLRKNVALVSQEPVLFSGSIRDNIAYGSIEDE

NFCNVPEDDIWKAAEIANADKFIKALPNGLDTVVGERGITLSGGQRQRVAIARALMRDPK

VLLLDEATSALDAESEHLVQESLEKIMHGRTVLVIAHRLSTVRNADQIAVLENGSVAELG

SYDELMSREGAFKTLVRLQTFDSS

>FoABCB-03

MLYCPPNITLSQVWVDGGTSQCFMDTVSNSITAGWLLLFGTSQLWIYRKWGTQVSQASLP

KSQLYGLQLLFLVLLPVLCVVRFAVQATLINDNVVYGYMILSMVLSIVQFPLSIALLVVE

RHYLLPSVPSRGHGLVLLLFWSLLFIFENLSFVNLRKDDWWFHLTTLTDKIEMALFVLRY

VSCLGIFVLGLRAPGIMTYRDYYNIGSASSEARIPIVEDPDSSNVSTWRNFWKKVKILAP

FLWPHKEILLQFRVLFCFLLLAGGRVINLYVPIFSKLIVDSMTEKSPIFHYDYVMIYVLF

KFLQGSGTGAMGLLNNLRSFLWIRIQQYTTREVEVSLFEHLHALSLRWHLSRKTGEVLRV

MDRGTDSINNLLNYILFSIFPTITDIVIAVVFFVSSFNGWFGLIVFTTMVLYIAATIMVT

EWRTKFQRRMNLANNAQNARSVDSLLNFETVKYYGAEAYEVEAYKEAILVYQKEEWKNSL

TLNILNTAQNLIISAGLLAGSLLCVHMVVDLHALTVGDYVLFSSYIIQLYVPLNWFGTYY

RAIQKNFVDMENMFDLLKETQEVVDIPGAGPLAVRQGKIEFCNVSFSYLPERTVLKNISF

TVPGGKTVAIVGPSGSGKTTIMRLLFRFYDVDNGAILIDGQNIKTVQQNSLRQNIGVVPQ

DTVLFNNSIRYNIQYARLNAAEADVIAAAKSAEIHDRILTFPEKYDTQVGERGLKLSGGE

KQRVAIARTILKSPAIVLLDEATSALDTQTERNIQAALHKVCADRTTVIVAHRLSTIIHA

DEIIVLKEGEIVERGRHEELITQESVYADMWQQQLKNDNEALKESNSEENSTSSGKGTR

>FoABCB-05

MLLRLFKTCDRSGISFQIRSQLSFRSPPQFNIKSSNTSFIRSFSNNVHQKNNVKVNLLQC

SWLTRSTGLFSTGLLGFFGALPTIAFCEAQHKTTGRLTGLRHTNTEKSKFDWPKFWTMLK

PDIWTLAAAIAAALVVALINIKIPLILGGLVNIISKISSESEKSFSEQILSPGLRLVSYF

VAQSVFTFVYISLLSGVGERVAFEMKNQLFASLMSQDLSFFDEERTGELMNRLTTDVQNF

KSAFKHCVSQGLRSGTQIIGCAGALISVSPQMTGMIMVLVPSVIIFGTYIGSLLRKLSAR

ASAQVGRISTVAEEALSNIRTVRAFGMEDKECELLREESKKAQELNEQLGWGIGLFQAGT

NLFLNGMVLTTLCMGGYLMSVNEMSAGDVMSFLVAAQTVQRSLAQVSLLYGEVLKGLGAG

ASVFEYINRKPKVPLIGGEVIPYHTLQPHVKFEDVTFSYPTRPDQQILKNFNLSIPAGST

VAIVGSSGNGKSTVAALLARMYEADSGTVTLGGHDLSKLDPTWLRSRVIGFINQEPVLFA

TSIKENIRYSKPSATDEEVVAVAKLAQADEFIQSFPEGYDTVVGEKGVTVSGGQKQRIAI

ARALLKNPSILILDEATSALDAQSEKTVQKALEIAAKGRTVLVIAHRLSTVKNADQIVVL

QKGVIVEMGTHEELQKKKGYYWSLVQHQELMNRQVG

>FoABCC-01

MAYEGMDAFCGSPFWNSSRTWETDDPDFTICFEKTALVWAPCVFLWCFAPLEAYYIAKSR

NRDIPWNWLNISKLIFTGLLVALSVADMGNAAYRNGDGLQVYSVDYYTPLIKIATFFLAG

VLLAYNRKHGICSSGLLTLFWLFLFLFGIPQYRTELREELELRDASEQKQFPFISSMIYL

PLVLGILVLNCFADAEPRHLQYRKTENMCPELNASFFSKVFFLWFDPFAWKGFRNPLETK

DLWAMKPEDAAEEVVPLFTKFWEETVQKTTGTQPGAAKASFRKSSGSVDLTGGPKKPLKQ

ASVLPALCKAFGPTFVFGAVLKLIVDLLTFVSPQILKLLISFVAHDEPLWKGLLYAIVLL

LTAMIQTLFLAQYFHRMFLVGMRIRTALVAAIYRKALRMSNSARKESTVGEIVNLMSVDA

QRFIDLTAYLNMIWSAPLQIALALYFLWNTLGPSVLAGLAVMIILIPVNGVIASKVKNLQ

ITQMKYKDERVKLMNEILSGMKVLKLYAWEPSFEDKILKIRDKEIKVLKQAAYLNAGTAF

IWACAPFLVSLVSFATFVLVDEKNVLDSEKAFVSLSLFNILRFPLSMLPMMISNLVQAAV

SIKRINKYMNSEDLDPSNVSHDTKEGAPLNIESGTFAWGPDEPPTLKNINFKVNEGSLVA

VVGSVGSGKSSLLSALLGEMDKISGQVNAKGSLAYVPQQAWIQNATLKNNILFGKSADNS

RYVKVVDACALKSDFDMLPGGDQTEIGEKGINLSGGQKQRVSLARAVYNDAEVYLLDDPL

SAVDSHVGKHIFDNVIGPQGLLKKKTRVLVTHGITFLREVDLIVVLKDGEVSETGTFKEL

MEKKGAFAEFLLHHLQEVSNEGDADAEEMIAQLEGTLGAEELQKELHRRKARVSESQSDT

GSIAERTSLTGSLKRGGSGSSEGLRRRHSSEQKEPTPPKQDLAKGEKLIEAEKAETGSVK

WRVYAHYLSSIGVFLSVATIVLNMIFQGFSIGSNVWLSKWSAQNYTNLNGSDYTSTRDMY

LGVYGALGFGQVLTVFLASLATYVGTLSAAKILHYHILFNVMRIPLSFFDVTPVGRILNR

LTKDVDTLDNVLPMTLRGWTNCFFAVIATLVVISYSTPLFIAVILPVGILYYFVQRFYVA

TSRQLKRLESVSRSPIYSHFGESITGASSIRAYGVVDRFVNESEEKVDFNQVCFFPSIIA

NRWLAVRLEMVGNFIIFFAALFAVISKDSSPDMAGLVGLSISYALQITQTLNWLVRMTSD

VETNIVAVERIKEYGETPQEAAWENPNSVVDSNWPQEGKVDFKDYEVRYREGLDLVLKGI

NFTVKGGEKVGIVGRTGAGKSSLTLALFRIIEAAGGQILIDGVDISKLGLHALRSRLTII

PQDPVLFSGSMRLNLDPFGSFKDDDIWRALEHAHLKSFVKGLAAGLEHEVTEGGENLSVG

QRQLICLARALLRKTKVLVLDEATAAVDLETDDLIQRTIREEFKECTILTIAHRLNTIMD

SDRVLVLDQGKVVEYESPDALLHKKTSIFYSMAKDAGLV

>FoABCC-03

METTNKTLPQPPNLRAKANILSALTFAWTFPLFRQGYKRDLTVSDLCVPLDAYDSKQLGD

NLEDLWNEQQKKSHEGTKQKLPSVWAVLRSFCGWELFFLAFGLFLFDLTIKLGQPILLGK

FLNYFTVNSTVTGEEATIYASGVAVSVFLHQFGMHPFLFGLMYLGMRVRVGLSSLLYRKL

LRVSISSSAGVTGGYVINLLTNDASRFDFAPLFLPDAFGGIVCSIVVTYFLWQEVGISSL

VGLSALVLFVPVQAMLARKSSSLREKTAKRTDKRIQLMNEIIQGIQAIKMYAWEKRFSRL

VEVSRKKEITVIRSKLYYTGVAFALSLFLTKVSFFFCVLHFALSGNAIAPDRIFSAIMYF

NILQESLTIYFPLSIQKLAECVVAAQRMQEFLLRPESTCMLVKPSVTGSAEVLLKNVNAK

WTYDQQSNSLEDLNLSVSSGMLVGVIGQVGSGKSSLLHLILREMPHVTGCMNIQGSISYA

SQEPWLFVGTVLQNILFGLPMDKQRYKEVVRVCALEDDFKQFPRGDKTQVGEHGHSLSGG

QRARINLARAVYRRADIYLLDDPLSAVDTHVGRHLFDDCISGFLAGKVVILCTHQLQYLS

NVELILMMKNGSIYAQGTFSELQNCGTELSDLMKAESKSEENPAKDQLQRRLSVHSTNSE

EDFDVPEEMKSKGSVTWHVYASYFKAACGVPTFLLLLILLVLSQGSISLADYWTAYWVDL

EQYYFKYGNATYLPLLNEMNQEKCLYVLGFLIALTIFVNLSRIISFVYCCMEGSKTLHDV

MFNTVSKATMYFFDNNPSGRILTRFSRDIGVVDEKLPGTLSDFLEVAMHLLGVLCIICVV

NYWLVIPALVMGILFGLLRHLFMLTSRSLKRLEGVTSSPIYTHTNATLGGLVTIRSMHAS

DDLTREFDSKQNINSSAVFYSLGTNRAFAYWLDLVCCCYTMLVIFSFLFLDSEFKGGEVG

LTITQALSLIFMCQWAMRQAAEVENFMTSVERVLEFHKVESEEGRLLTKVEHNLPKMWPT

CGDITFQNVCLRYSSSDPPVLKRLNFSIKSGEKIGIVGRTGAGKSSLLSALFQLYPVEGT

IIIDKVDTSKIGLQDLRSHLSVIPQEPVLFTGSLRHNIDPFEQFKDEELWRVLEDVKLKS

YVEDMHGGLQFVVTEGGSNLSVGQRQLVCLARAILRQNKILILDEATANMDTYTDQLIQA

TLRHKFVDCTVLTVAHRLDTVMDCDRILVMDAGQVAEFGKPYELLENKNGLFSKLVNQTG

LTNSYNLKAIAERAKYKESPHSTEAENK

>FoABCC-04

MNSSTRPSSVSWVFKWNWDEMCGPDGLVVWSTSHKDLGLCFQELCMKVPTLTLLAAVSAF

YCGSQDRWVVRDKNMIRVIFVRSVIAMALAIEPIIRVCNKINDAPTALHPVDYLLYAVES

FAWLVHFGYILALRHRLGVSPRGPIVACVLWTLIFVFNIVSVRSLARIIREGNASQSVYI

DYYADLINLCTHFLYLCTLLPGRGGTHAAYNRMYTEYYQRVIQSDERQGLLSSRHAYSGF

REEQEPGYLGVAMEDSTFSSRLFFHWVRPLMVKGVEGQLHDVDDLFDLPEDLTPSYIHAQ

LEQAIGRSSALNSEKTSLSSGQYGYGSVQGPSSSLQASLAAPQQVEDVHVVSSPVSLLRA

LHSCFAWEFYSVGVIKFLGDAVGFASPLLLNLLVKFIESKSERKEDGYFFALGLCLTALI

SALCNAHFNFLMAKVGMKIRSAIITLVYNKILTLSTKSFSLFSSGEVVNFMSTDTDRIVN

SCPSFHALWSIPLQLAVTFYLLYTQVGLAFLAGVCFAVVLIPINKLIAMKIGSLSSKMME

KKDERVEMCSEVLQGMRTIKMHVWEEHFLTKILRIRESELYFLKGRKYLDAMCVFFWATT

PVMIALLTFVCYSLLGGQLTASRVFTTIALLNMLIAPLNAFPWVLNGLVEAWVSVARVQK

LITLPDLDLHEFYHPLPIDNDNIAIVVRNGKFEWGNEKSQDRFQLNDVNFTILKGQLIGV

IGEVGSGKSTLLASILAEPEKQSGVVAVDNSDQGFAYVAQHPWLQRGTIRDNILFGKPFD

QLRYRSVLDACCLEDDLRTLPKGDLCGIAEGATNLSGGQKARIALARAVYQDKAIYIFDD

ILSSVDPPVANLIFDKCIRKLLNGRTRILCTHYAQCLRYADVVMKISAGRIVHMGSPSDV

LAEEMENLALLDSDSIPSGSKEGTRITRDEDIISEQEDDILLNEHREIGTVQLKVYGAYW

RALSPFLSIAIFISLILMQSTRNLSDWWMSYWVSHNSSNPLGKNTEFKVDVERLVASNIS

TPELTNISFYLTVYGTIAGLNSFFTLLRAFLFAYAGIHAATELHKQLIKTLMRAKIFFFD

ISPLGRVLNRLSSDTYTVDDSLPFILNILLAQFFGLLGTMFVTIYGLPWLCLVLIPLIPI

YHWLQEQYRQTSRELKRISSVTLSPVYSHFSESMLGLATIRAFRASYRFKRQNENFLDCN

QRSVLSSQAASQWLGLRLQLMGVAIISGVGMIAVIQHQFNVADPGIVGLAIAYALSVTSM

LGGVVNAFTETEREMVAVERVHQYITDIQCESQKPGLQPAPYAWPPQGVVQFQNVILKYR

DHLAPSLKCITFQTRPSEKVGVVGRTGAGKSSLIAALFRLVEISNGCILIDSIDISNMSL

HSLRRRLAVIPQDPFLFAGSVRDNIDPLKEYSDHEVYDVLNRCHIVDVVRRMGGLNAQVG

PGGSFFSAGQKQLLCLTRAILHNAKILCIDEATANVDHETDRLIQKTIRASFGKSTVITV

AHRIETIMDSDRVLVLGGGEVLEFDSPDKLMEDKTSHFYRLASHGQERS

>FoABCC-05

MAETGAEHHKWLSTGSLCLSIDGEDKKTDPYEEDMGTTIRNQPPGVKKSNKYGPAMKHLL

PYRKAPPDSQVMPGNSSGLFSFIAVAWMTKLMWKAYRHGLSEEDLYELHTADRAESNACR

LESAWHDRVRQQGFEASLFSVVWRFCRTRAIVATLLMVLAVIFQFLGPAILQRLILQYVD

DKTLPLSWGLTLVVMLFATQLLRNYCFGAAYVMGLHTAIRIQGSIQHLVYKKMLLLRSGG

ERALGQVITFCTNEQERVFEAAHMGVLILGTPVMFTMCVIYSVWVLGPWALAGNFVILLF

YPVMAGVASLTSMVRIKTVQVTDKRIGLMHEILSSIRLIKMYAWEDSFAKNICHVRDLER

QQLQKAAFLQSFSNTITPSITILASIATFLGYSLTGNNLYTAEAFTVYAVFTAMQFTVGT

LPYGIKCLAEAKVSCNKIQKFLQRPNFESTVTRNPGGVPSTGPVIDFKNATFIWDDDEPE

MLMKANTKIRGNLPKLTEIDAANGVKNGKNLEEANKLFSDVDKPEEPLREISLTVQRGTL

LGICGSVGAGKSSLLSAITGDMIRVQGEMFVQGGLALVSQQAWIFNETLRENILFGLPYD

EERYNEVINVCSLQRDLEMLSNGDQTEIGERGSNLSGGQKQRVNLARAVYANKDVYLLDD

PLSAVDARVARHIFSKCIKGTLKGKTILLVTHGLQFLADCDEVVFMKDGTISERGSHSAL

SEAAGDYCQMLTFDHSRSVKKQNSSDDTEVKTIDVDEETPLEENVGKLTSAEEHKSLGFS

GYLRYFVFCGGYFIMLLLLLLIVLFTLARLFSGVWLQIWLDEGDGLEAERKFNISQYNLT

YTDNELKGLVNENPDRAYYQLVYGLTFGLTIVIGVCKGWGIARQLLLGSSRLHDTMFRKI

MRCPISFYDVTPTGRILQRFSKDMDELDVRIPFFIEFVWQGLLFCITQMLLICFVFPIFS

FALAVAAVLFAFLDVWLNKGLKETKRLDNLRKSPVLNHITSTMSGLSVIRSFGRQRVFLE

RFCARLNQSLASDLIYRSSVRWFTFRMDMIAVVMVTLTGFVVVYFRGTVTTAESGLALSC

VFAVCTFIPYVMQLKSEFQSRFTSAERVLEYSSDLPEEAPKQIDQSPKPKNWPEKGDVKM

ENVELRYRPGLPLVLQSVSAHIKGGEKIGVVGRTGAGKSSLISTFLRLTELAGGRILIDN

IDISLLGLQELRSAIAIIPQDPVLFEGTLRFNLDPFGEHKDGKIWEALEKSHLKDKLSRE

EKQLQTPVSAAGQNLSVGEKQLICLARAVLRQNKILLLDEATASVDVETDFLIQRTIRET

FKDCTVVTIAHRLHTVSSYDRVMVMDAGKVVEFDTPFALTANKDSVFHGMLEAAGVAYPA

ASPDSPSR

>FoABCC-08

MPDAAEAQASSFISSPNMAKDLSPEIVNYEADPEDGFLDVDLGDGYLPPQSPRIYIRSSA

PYIADQGMERYNQALRSLIPVRRKATEKKDTIQVDQAGLLSFVFSSWISKLMYKAHRKGL

VSGDIPRGSPLDSCDYNVQRLEMLWQEEIRKFGPSGASLPRAVWRFVKTRVIVDWLIFTI

SSILGFLTPMIFMRKLLEFSEDSNSNTETGLILAFSLGLCEFLRLIFFTWRFVLSFRTAV

RLRVACLALMYKKILRLNSFDENSMGQLLNIFANDSQRIFDMIVFGPMIIGGPMVLITGV

AYVLCTLSPWALSGILVFIAFYPVQYFISWLCSKLRRKTVQAADQRINLMNEILTHIKTV

KMNAWEGDLKDEIDCIRSKEQGYIQKTSYLSSFGSSLAPTVPIMSSIVSFLSHLGSGNTL

SAAQAFSVQAVYSNSIRDVVRSVQMSLTAYHDAIIGLKRIQAFPFVTVMVEQARNFVNIT

QIALSNYIEADVGLKRMQKCLLVEEIPLRIQRPIDRSQAISIGGASFCRSPSWHPLFSEG

QGNKSGKIIKQHLRKSDLETATEEREKLNGNVTCHLQPVLQGINFSVTKGQMVGICGPVG

SGKTSLMRAILGQMRLIDGQFAKDGSCAYVSQQAWITNGTLRENILFGEPFVSTRYYTVI

NSCALNEDLNQFPGGDLTEIGERGINLSGGQKQRIALARALYANRDIYLLDDPLSAVDVN

VANHIFENYFKKELREKTVVLVCHQVQYLHHCNEVYFLRDGCIIERGTHEELLKLDKEYA

SMVKQVKSTTCEASSSTPNTNAVVNNCKEGESKDEKKDTEKRNMGESLLVPEPSNLGSLK

IDTYCQYISAGGGYFTTFLVLCVFLANIGSTAFSSWWLATWIKAGGGNYTLLVNDTAVLS

SNLGDNPYYEFYQTVYASTIGLIILTCGLRCLVFTKVTLRASSILHNELFKKISRGPMVF

FESNPIGRLQNVFSRDMDEVDSRLPGTVENLISNFWILSFAILIVCVVFPWFFVALIPLC

IVYYFMSRIFRKAIRDLKRLENTTRSPVFSNLLETAQGLNTIRAFNKEGEYIQRFFQLFD

ENTTCNYLWNVGMRWLAVRINLLAVLTLSIVAFLVVILHGHVAPAMAGLALSYSWHMSGI

LQYTTRLFSETEVRFISVERILTYIENVLTEGKPTCSKPLPSWPAQGRISFENVNMNYRN

NLPNSLCNVSFNVLPGEKIGIVGRTGSGKSSLTVALFRLVEINSGHIKLDGVDIADINLT

VLRDKISIIPQDPVVFSGTIRSNLDRKKNRNDMELWDALEKTKLADKIRSLPLKLDTAVK

GGSSGLSDGECQLLCLTRALLKSSKVIVFDEATASIDPETEAAVQSTIWEEFNMSTVLII

AHRLSTITNCDRVLVMDAGKVIEFEEPQKLLENTNSLFFKMMASLKTQ

>FoABCC-09

MDSGHRIPLKPNPRTSTNFLSKLTFLWTLPIFRQGLRKEFEESDLFQPLSEHESGLIGAK

LETIWQDVVKESEGRNTTPSLMKALLRCFGKEFMCVGIPLAIMEMFIRIMQPIFLGWLVS

YFSEESKDEEYAGYVYGVLVIVCSALTVWVVHPYMLSILHLGMKIRVATCSLAYRKLLRL

NRAALKETTPGQLVNLLSNDVNRFDIAVLFIHYLWLGPLQTLVITGFLIHEIGLVPAVVG

VASLLVYAPLQGWFGKLASVYRLKVAMRTDERVRLMNEIIGAIQVIKMYAWEVPFQKMVA

LARKLEMKQIMNTNYIRGTIVSFIIFAKRFSIFAALLTFALAGEKLTAKHVFIVTAYFNI

LQHNMTVFFPQGITQFAEGLVSVKRLQKIMLLPELKSVPRNELKKEGGITMVDACAKWDD

ELTENTLTDISLKIAAGSLVAVVGPVGAGKSSILNAILGELALDSGSMTVGGTVSYAGQE

PWLFASTVRQNILFGEPMDPIRYKEVVNVCALKTDFAMFPQGDRTIVGEKGVSLSGGQRA

RINLARALYRQTDIYLLDDPLSAVDAHVSAHLFHDCISKYLHGKTRILVTHQLQYLPFVD

HVFVLNNGHLVAHGPYAEVKEKGLASLLGSSVHDAEKSEVKEDDNEKSTLLKRERDSPSR

HSIISHTSSTGHDSKRSSIDPSMMECVQSEEPEERAHGAVGWKIFTSYFMAAGNWFYVIF

VLMMFVWCQIFASAGDYWITVWIRVEQDNYLNPNSSTVSSEPWYGEWPPNRETCIYVYTV

ITLATIVITLVRSFLFFHMCVRASRCLHNNMFASITQAKMRFFYNNTSGRILNRFSKDMG

AIDEILPLCMIDTFQIGAQLFGIMVLVSVINYWLIIPLVVVSVIGLWIRNVFMTTSRTIK

RLEGVTRSPVFSHMTASLSGLTTLRVSNMEVKLIEEFDRHQDLHSSAWYLFLATNRAFGY

WLDQLCLIFITAVTFGFLIGSHLGVFGGNVGLAITQAISLTGMLQWGLRQEAEVENHMTS

VERVLEYSCLPSEPPLEAPPEKKPDPLWPKEGCIEFENLYLHYDPEDPPVLKNLNIKINP

GEKVGIVGRTGAGKSSLIAALFRLEELRGRISIDGLDTGMIGVHDLRKKLSIIPQEAVLF

NATLRKNLDPFEEHPDDAIWSALRETELKEVVDELGFGLQSPIQEGGSNLSVGLRQLVCL

ARAILRNNRVLILDEATANVDPHTDKLIQKMIRNKFGQCTVLTIAHRLHTIMDSDKVLVM

DAGSVAEFDHPHILLKNTDSLLYKMVQQVGGASAQKLFSVAEEAKLPCGQFFMRTLD

>FoABCD-01

MAPNISKLGTPKTVLAVSSAAVTLWLIRQLTKSTKSRKLDRNEIQYLIGEKTKSSKSKAQ

VDAEFFGHLKKLLKIIIPGWFSAETSFCILVAASLVARSFCDIWMIQNATQVETAIVSMN

KNRFKSTLLVFFAGMPLISVVNNTLKWSLGELKLRMRTRMTHHLYEQYLKGFTYYKMNNL

DTRIANADQLLTQDVDKFCDSVTELYSNILKPLLDIFIYVYRLTSVLGGQTPATMMVYLV

VSGTFLTYLRRPMGRMTAEEQKLEGEFRYINSRLITNSEEIAFYNGNNREKLTMLVSFKR

LVDHLRNCLEFRTVVGVVDNIVAKYFATVVGMIAFSIPFMNTEHAQLSKIGQEERFRYYY

TYGRMMVKLAEAIGRLVLAGREMTRLAGFTARVTELMNVLKDVNEGHYQRTMVTDSADKE

APVQPLIPGSGRMIFQDNIIRFDKVPLVTPNGDILVKELSFEVKSGMNVLVCGPNGCGKS

SLFRILGELWPLFGGTLTKPPRGKLFYIPQRPYMTLGSLRDQVIYPHTRDEMQRRGKTDA

DLTRHLERVQLSYLLTRENGWDAVADWIDVLSGGEKQRIAMARLFYHAPQFAILDECTSA

VSVDVEGSMYDYCREAGITLFTVSHRRSLWKHHEYYLQMDGRGAFDFKKIEADTEEFGS

>FoABCD-02

MPNVMSRVLDSTAAKYNIPREYLSKTILGTAVCLYGIKLGYPLIQQLCKKSVTLNGVKDV

KGSRTRNGTVVPANGQDVSTNNNEHVANGKTVDNGNKGPATKERRSSPAVNQEFLLQLEK

LIRIMIPGLWCGEVALLSVHTLALFTRTFLSIYVASMEGQIVKFIVRRDIWNFSIMLMKW

LGVAVPATFINSMIRYLENKLALAFRTRLVNHAYSKYFADQTYYRVSNLDGRIENADHRL

TDDITAFTSSVAHLYSHLTKPLFDCALIGLALAQSSRRMGANIIPGPLLACVVIGATGQI

LKAVSPKFGLLVAEEANRKGYLRSIHSRVITNAEEIAFYGGHKVELTHLQRAYQTLVRQK

NYVFQQRLWYVMLEQFLMKYVWSGTGMVMVALPIVMGTRLQVGTETSDSGVSERTQYFTT

ARNLLVSGADAVERLMSSYKEVVELAGYTSRVGEMLDVFGEVGEGRYKRTTTSATKQNSR

GTDRLEFKDGQPVAKGKITESTDGSIALIDVPVVTPNCDVVVPSLTLVINQGMHLLITGP

NGCGKSSMFRILSGLWPVYAGELKRPKDCSMFYIPQRPYMSLGSLRDQVIYPDTVEDMKR

KGMTDDDLERSLALVHLSQLVLREGGWDAQADWKDTLSGGEKQRVAVARLFYHKPKFALL

DECTSAVSIDVESNIYQAAKDAGVTLLTITHRPSLWKFHSHILQFDGEGGWNLSPLDAKT

AMTLRQEKAHLEAQLADIPVAQQRLQEVCKELEDEEERLLSTTCN

>FoABCE-01

MSRRKGQEECDKLTRIAIVSSDKCKPKRCRQECKKSCPVVRMGKLCIEVSVSDKIASISE

ELCIGCGICVKKCPFEAINIINLPSNLERDTTHRYSQNSFKLHRLPIPRPGEVLGLVGTN

GIGKSTALKILAGKQKPNLGRYLNPPDWQEILNNFRGSELQNYFTKILEDNLKALIKPQY

VDQIPKAIKGTVQELLDKKDERNNQNEICDMLDLQKIRGRQIDQLSGGELQRFACAMVCI

QDGDIFMFDEPSSYLDVKQRLKAAVTIRSLIQADKFIIVVEHDLSVLDYLSDFICCLYGV

PGAYGVVTMPFSVREGINIFLDGFVPTENLRFREESLVFRVAESATEEEVKRMNHYEYPT

MSKQLGSFKLAVECGQFTDSEILVLLGENGTGKTTFIRMLAGKLPPDTGSGDLPVLNISY

KPQKISPKSQGVVRQLLHEKIRDAYVHPQFITDVMKPMRIDDIMDQEVQNLSGGELQRVA

MTLCLGQPADVYLIDEPSAYLDSEQRLVAAKVIKRFILHAKKTGFVVEHDFIMATYLADR

VIVFEGSPSVDTTAHSPQSLLNGMNRFLELLGITFRRDPNNFRPRINKNQSVKDVEQKRA

GQYFFLEE

>FoABCF-02

MAPKKGKGNKKDLDFGSDDENTKNTSDKGKSVAGVEIEGVTSKSKQSAPAPKKKGKGKSK

KNDDWSDDDEKIEKLSKLGQVSDDDEVLNIAQPSKKKAGKQIKKKADSSSEESEDDSEMS

SKPSKAAPKANVKKKSKQVKKGDDWSDNSDEMKLSNSAEEDVPKPATKKGKNKKGKSKVE

TDSESSDQVDDEVEDEPEPQPVKKGKSKTVAPSTKDVDSEVDDGVGDELTDKVAELSVQE

HTKLTTASTDKGDCGLGEEEDPSEEKDKLKNEDSNKPAEKKMTSKEKKKLKKKIEFEKQM

EHITKKGGQGHSDLTENFTVSQAQKSTGQLAAMENSVDIKVENFSISAKGNDLFVNANLL

IAQGRRYGLVGPNGHGKTTLLRHIASRAFAIPPNIDILYCEQEVVAEDASAVEVVLKADT

KRTTLLEELKKLEESPEAGTLAGQERLKEIYDELKAIGADSAEPRARRILAGLGFSRSMQ

DRPTNSFSGGWRMRVSLGRALYVEPTLLLLDEPTNHLDLNAVIWLDNYLQGWKKTLLIVS

HDQSFLDNVCSEIIHLDQHKLHYYKGNYSMFKKMYVQKRKEMIKEYEKQEKRIKELKAHG

TSKKAAEKKQKEALTRKQEKNRTKVQKLEDENAPVELLQKPREYIVKFSFPEPPPLQPPI

LGLHNVTFAYPGQKPLFVDCEFGIDLSSRVAIVGPNGVGKSTFLKLLTGDLDPLKGDVKR

NHRLHIGRFDQHSGEHLTAEETPSEYLMRLFDLPYEKARKQLGTFGLSSHAHTIKMKDLS

GGQKARVALAELCLNAPDVVILDEPTNNLDIESIDALADAINEYQGGVIIVSHDERLIRD

TECTLWVIEDRTINEVDGDFDDYRKELLESLGEVINNPSIAANAAVSQ

>FoABCF-03

MAAYLEYIKEEFPKIDDEVYQYVEGVLSGSADDFEDSEEIYEAIGGVLQEVGDDKSEDDI

KDICEKLWDIMRPETDSVKKVNRVLEAPVHLKSLSDNQDNDITEIKSIWVTQRDDVLKVD

AKKLEKAEAKIKQRQEKRTAEMTNRPVAAPVKLETASASQVTSRKDAKMEAKGSNRTQDI

RIENFDVAFGDKVLLQGADIVLSFGRRYGLVGRNGHGKTTLLRMISGAQLRIPSHISILH

VEQEVVGDETPALQSVLECDTKREELLKREKEITAAVNAGTADTKMSDELSEVYIALQNM

EADKAPARASRILNGLGFTVEMQSCKTKEFSGGWRMRLALARALFSKPDLLLLDEPTNML

DIKAIIWLENYLQNWPTTLLVVSHDRNFLDTVPTDILHLYAQRLEAYRGNYEQFEKTKNE

KLKNQQREWEAQQQHRDHVQAFIDRFRYNANRASSVQSKIKMLEKLPELKPVEKEVDVVL

KFPPVEPLSPPILQLSEVSFAYESTGRTIFSNVNLNATLESRICIVGDNGAGKTTLLKII

MGINAPTRGDRVIHRNLKFGYFSQHHVDQLDMNVCSVELLQKTYPGKPVEEYRRQLGSFG

VSGDLALQSIASLSGGQKSRVAFAVMCMGNPNFLVLDEPTNHLDIESIHALGLAINKYLG

GVILVSHDERLIRMVCKELWICGGGSVHSIEGGFDEYRKIVERELAEQNK

>FoABCG-03

MNPVLAGVEPDSKQTLEVAVDDAAPRPLTPVMPRRGLSHLPERPPVDIEFTDLTYTVPQG

RKGSKMILRSVSGLFRSGELTAILGPSGAGKSTLLNVLAGYKTGDATGTININGRPRDAA

VFRKLSRYIMQEDLLQQHITVLESMTIAADLKLGNTLSRAQKRTAIDEILDMLRLTKARD

TQTSKLSGGERKRLSIAQELVNNPPVIFLDEPTTGLDDLSSSQCIQLLKMLARGGRTVIC

SVHTPSAKLFDMFDHVYIVSEGQCVFQGKGRDIVPFLSSLGLQCPKTYNPADFMVEAASG

EYGNHTERMSAAVDNGRCYRWCKDPRTLQHQSSRTDIVNTSPSANYDFSSSSFMQFRILM

YRNLLQTWRDSGYLLLVASMHFFVALIIGMLFYQMGNDGSKTIFNFGFCFTCIIVFMYIP

MLPILLKFPGEVQLLKREYFNRWYSLNAYFCAMTVARLPTQIILGLLYIALVYPITDQPL

EPMRLLKFSIICLLVSVVSESFGLIIGSTLSLVNSMFVGPAFSVPFMLLAVYGMGNGSDE

NAIPLHYRLAMSLSYLRYGLEGIVISIYGDHRASLVCPDSEVYCPLQDPRQLLREMGMEH

VKYWVDVVALVSSFILFRLLSYTVLRVRLSTTKPIPAVNFISRMVKTHLNINLR

>FoABCG-04-partB

SGTFIGPVSSVPMLLFSGFFVNFNTIPPYLQWLSYLSYVRYGFEGAMVAVYGYGRKKLSC

SEPYCHYRSPTKFLQAMDMADAYFWVDMVALLVFFVVLRFITYFILRLKLRSTR

>FoABCG-04-partA

MGDGSGAQGLPSSPPPCQGEGLNNGLLMGGGPYVSYGIAGGQGLHGALVKKVPNDKSSYL

SVGAAELGLAGPLPMGSGQGPGGTSPGSVASGGSSGGGVAGGKAPITLSNLPRRPPVAIQ

FNGLAYSVSEGRKRGYKTLLKTVSGVFWSGELTAIMGPSGAGKSTLMNSLAGYRTSNLSG

SVLVNGRPRNLRRFRKMSCYIMQDDQLLPHLTVREYMTVSANLKLSDDIGAPAKRRVVDE

LLETLGLCECQDTRTGNLSGGQRKRLSIALELVNNPPVMFLDEPTSGLDSSSCFQCLTLL

KSLSQGGRTIICTIHQPSARLFEMFDQLYALAEGQCIYQGSVHGVVPFLASMGLECPSYH

NPADFVMEVACGEHGEHVPKLVTAVTNGKCNNLYQTGHSAQPGQPGSGANTCDCGLSVSS

APGGGESCNSCNSCAQSATTLLDSSDESLSGGHFPTSTATQFCILLKRTFLSIMRDQTLT

LFRLISHLTVGLLIGVIYYGIGDEASKAMNNAGCIFFTVLFVMFTAMMPTILTFPMEMSV

FVREHLNYWYSLKSFYLAKTVADLPFQLAFSVVYVCIVYFMTAQPMEADRFFMVLLSCLL

TSLVAQSLGLLIGAGMSV

>FoABCG-05

MTVTTESEPLLDDGGGRRPPSTTERSRRNGGLWWSSDSTTNIGGKGYGSVGNGNGDKSSY

GADDNGGWTIQQAPSAPLTYTWTSLDAFAPGVRPSRTPFSHWLRRRPAAPRKHLLKNVSG

AALPGEVLAIMGASGAGKTTLLDALTFRSDRQLTVTGARALSGTVVTSSRDLANVAAYVQ

QHDLFVGWLTVREHLVFQALLRMERRIPRRQRLLRVREVMEELGLTKCEDSIIGVPGRVK

GISGGEMKRLSFASEVLTDPTLLLCDEPTSGLDAYMAQTTVEVLTRLARRGKTVVLTIHQ

PSSEVFAQFDKILLMAEGRVAFLGSPQEAQDFFGRIGAPIPPKYNPADFFVQLLAVVPNE

EVTSRDRINRVCDAYERSEHAARSAAVTDYPVKGLPPLEGGRAGSTWLLRDPRSSSRYKA

SWATQFAALLWRSWLSVRKDPLLVRVRVLQTAVIALLVSFLYFGQSLDQTGLVNINAALF

TTITNMTFQNVLAVIKVFCTERAVFKREHMAAMYRVDAYFLARSTAELPIFMALPVLYTC

IVYFAVGFNADPVRFFTCVAIVNLIGLVATSFGYFISCAANNMSTALSTGAPAVIPLLLY

GGFFVNLDTMPSYFSWIQHISWFRYGNEALLVNQWQDVPEISCAGALNNNSCVRTGHYAL

QVYSFKEGTVWSDIGHLALLILLFRSLALGLLIIKTR

>FoABCG-06

MIISKDLFLLGDQDYLRLPKEEKRPAAMVRPTVVKSQRSEVAGGALHCPFLLEQGEAHLT

GTRPLELVFSGLSVSVASRSILRDVSGVVRPGEMLAVMGPSGCGKTTLLNCLSGRSELDS

GCIRLNRERLTKRWKRRICYVLQQDIFFPDLTLRQTLEYAARLRLPESMSRRQKMLYVDH

IIDVMDLTACQDTRMGDYTKRGLSGGEKKRANIACELLTNPALMLLDEPTSGLDSHSAYT

LMSSLKRYAEKESKTLVVTVHQPSSQIFYMFDKLLLLCNGHTAYFGDVNKVVDFFNNIGL

TVTPHYNPADFVLECVKGSEEMKQKIITAAKSLPTYHQEALNDYCALPAQLYQHQGCNGR

VPMVGHALGHTANGPGCDFSKPSLWSSTPNGTLNSTPPLSLTTKCVVSNHNSHNLSNHVY

STIVKDEEGKSLWLDSQSHGSSSVSSTTDDDVRWQWPTSFWTQFKVLSSRNFQESRPRML

SKLYWLQTIGLGLMVGNVYFQLERKEESLHDIQSWMFFSTNYWMLFALFNALSSFPAERE

VVNKERLSGSYRLSAYYLAKMAGELPLTMTLPAVYHIISYPLFGFRSTSVFLTMLGFLLL

STVVAQSAGFFVGAACMDLDVAITISALYTLATQLFGGYLATNIPSWLRWMRYLSMVHYA

YQNMQIVEFSEGERIQCAVQSKFEVCSNSTHIPVSVILEQQGATLPLWANTLVLLFFLVV

FRVLGYVVLRYFRQPK

>FoABCG-10

IVNPTVFLLVAYYMTDQPNVEDRRCMVWLMCLLTVLMAHAYGLVFGAALGMQLGVFLVPA

STIPLVLFSGFFLYIPDIPLVLRWLSYVSNFRYGFEGIVTAIYGMNRQPLKCNQAFCHWK

RPSQFLEDIGMLNADYWTDVTGLVTWILVLHFSLLVSLKVKVILNRQ

>FoABCG-12

MVTNEISLELCNVFHTGQVEPGTCVQRLLGSVKTGVILKDVSLEVHGGEVLAILGSKGSG

KRALLDVVARREQGPTRGQILLNGVPMSLRLFQQSCAYVSHKCDVLPGLTAEQALHYAAS

LTIGSRVSRYMKSSRVKQVLADLALTQVANRTRSQLTASEYRRLSIGLQLIRDPVVLLLD

EPTWDLDPLNTYLVVSILSNHARKYGRAVVLTMEKPRSDVFPFLDRVAYLCLGDMVYTGP

TRFMLDYFRGIGFPCPELENPLMYYLCLSTVDRRSRDRFIESNHQIAALVDKFKLEGTPF

RKSASSGAVLGLGDGGVGLGGTLTLSPVTGSTVGASKVPLSAFGRPSALRVALALNMRCL

ASTFNLQAGSIWRLASRLLLLPLYFCVLWLFYHDKKAFQHTFVTSSGLIFNMLAGTYFIS

IINTIFTFPAHRNRYYQEAQEGLYSGPLFLLSYFLFSLPFSILSVIAGSRVAFEMSGLSD

PWDWVKLGCIMWGCYTLGEQQTVALLVVVKHSFTAAVASITLSALGLVLSSGVLRSIRGL

SEWLLYITYASQPRYAAAYIHRQIFSHPHYSGLARDSMTNCSLAPGTIQTQESLSAASAS

FGCRYADGWAFLAERFGRENAEDSATLTAFLELDFNMGVTFAFALGLTVVNCLLYLVPLP

AFIKAKFRE

>FoABCG-13

MTLAWNDLSVWVRKPKEGAGFFSREKHEHKQILNSVCGVAKSGSLMAIMGASGAGKTTML

ATVSQRVKGITRGEILVNGHVVDQHFMCQVSGFVPQQDLAVECLTVREHLEFMARLKMDR

RVRSSQRHRRILSLLMDLGLSKCCHTQLRRLSGGERRRVSLAVQLLTDPPILFCDEPTTG

LDSYSAGAVVEHLRLFAQRGKAVICTIHQPASGIFDLFHQVLLLAGGRVAFFGEVQDASR

HFDNLGLVCPSTFNQAEFLVSQLAVVPGQEGQCLKKIQWLCDEFDNSKYGQALSDELAAH

TGRFLQACETSSASSSSSALGLGKVGRRTWSVASHDSWQSSEEFQKYLNIKKPTKFTQFY

WLVWRSLIEIRRQPTDVLIRLGMLMFIAVLISTPYVSITMDQKGIQNMQGFLYLIVTEII

FAYSYSVFHTFPHEMPVLLREIGNGLYSAGPYYASKMLILLPRAVVEPFLYTLVVFLVGG

LTGGIADFCLLLVPVWACAIAATAYGCLMSASFESIETAAMVSVPYEFISVTFSGLYLQL

GNLPAHLKWIPFISVFYYGNEAVSILQWEKIDSIACDEDPGIPCISTGQGVLEKYGYKPN

DLGLDLAGLAGIYVFCHLLGFAALWRRSKRQAVY

>FoABCG-21

MDTEHLFEHLVMEQDADEDPAASACGTGSGSGSIGSSSVDSDTPRGVTLSWRDLTVFATA

RGAPCKRIIKHVSGAVRPGALVAVMGASGAGKSTLMTALAHRSPAGVAVRGDIRVDGRPV

GDFMRRLSGFLHQEDLFVAELTVREHLNFMARMRLDRRTSCWERQRKVQQLLRQLGLTAS

QHTRIGTPGHDKVISGGERKRLAFATELLTEPPLLFCDEPTTGLDSYSAQQLVDLMQRMT

ERPRGRGKTILCTIHQPSSDLLARFHRIMLVADGRIAFIGTPEDALLFFREQGYVCPASY

NPADFFIRTLACMPGSEEASKAAVRKICDRFAVSDHARKMEMEVMQDLGEYDKDVACGDV

QQPPVEPPMWPQKLWLLTGRAMLQVARDPSIQGMRLLQKLAIAVMAGLCFLGTVSVSQTG

VMSVQGALFILVSENTFSPMYSVLGVFPSELPLYLREHSAGLYGPGAFYMSKVLSMLPGL

VVEPLIFAVLAYWLAGLRDDLYAFGMTAAIVILTMNVSTACGFFFSAAFESVPTAMAYLV

PFDYVLMITSGVFVKLSTLPSYVSWTQYLSWLMYSNEAMSILQWQGVGNITCPDATLPCL

QDGDHVLDHYSFDATHLSRNIIAMGALYALFHALGLLCLWRRVRRLQ

>FoABCH-01

MEASTGSYAVCVRGACKAYGSRRRPLNVLENLDMSVLKGSIYGLLGASGCGKTTLLSCTV

GRRRLDAGSVWVLGGRPGEPGSGVPGPRIGYMPQEVALVGELTVRETLFYFGWIFGMTDD

QIEERIAFLQQLLDLPPKNKFVKLLSGGQQRRVSFAAALVHDPELLILDEPTVGLDPILR

QAIWDYLVKITRDDHKTVIITTHYIEEAKQAHSIGLMRAGRLLAEDSPSQLLDRYGSATL

EEVFLILSSNQGPVDQGPMDAALDQASVDQGAGTLPMTASSTEALHQSNGDIATISPNGN

DSLALRKLPVNNNGKPCNGGGYDVPVRARPPFSNMFKGHVKALLAKSFLRILRHLGALGF

IFIFPVLEVGVFFTAIGGDPQGLQLSVVNEELYAMNLSTCDEYSSLFGKPGSTYTGGTDD

DYGQPVGNCSVRMLSCAFLGYLDHRMAHQVFYKEKGEALNVSRAGDSIGVLHFGPNFTEW

LEARRRYGSELTAEEVDQGEIPIWLDMSNRQLGTIMTKSIVDSYHAFSRDLLSTCHANPK

IGEIPVHFEEPVYGVENPNFAVFAAPGIILTMIFFLTTGLTATVMITEKQEGLWDRSLVA

GVTAWESLLAHVVTQGVLMGIQAALALIMMFLVFKVPCAGHMGDVIFLVFITGMTGMCFG

FFVSVSCETVTTANFLSLGSFYPIILLSGVIWPLEGMSTVLRWIALCLPVTLSTTSLRNI

MLRGWGLTEPQVYIGFVVTGLWAAGLITICMIILKYRK

>FoABCH-03

MAELMETGGVPMGAGRRLKTQKSTVWTRRQQAVCVRRAYKRYGSKKNPNQILDGLNMTVP

KGTIYGLLGASGCGKTTLLSCIVGRRRLNSGEIWVLGGHPGSRGSGVPGPRIGYMPQEIA

LYGEFSIRETMIYFGWVAGMTTSEVEERLDFFINFLQLPPATRAVKNLSGGQQRRVSLAA

ALLNNPELLILDEPTVGVDPVLRQSIWDHLVEITQGGQRTVIITTHYIDETKQAGMIALM

RGGKFLAEESPEMLMQMYQSDSLENVFLKLSLAQIQGKRRRSIYLKEVAGSEDPAGESTA

DAASEISGEFGDNVSIKSGRSEGHHSVTGPSDLPPEEETPFSCKDMFIIFSPHRMKALIW

KNFLWMWRNFGVMAFVIGLPVLQICLFCFSIGRPPLGLHLAIVNNELNNWTDPCHFNTNC

TIDMLSCRYLAELQKYNVVEVPYPNEMEAKHAVKLGHAWAALSFPGNYSSSLYERLQQGK

DAEDSVLDFSNIEIWMDASDQQINQLLTRNILLSFEDFVMDVSRSCDYPEKVAKLPVNFE

TPIYGSKNPNFTNFAAPGVILTIIFFLSVALTSGAMLIERNEGILERSLVSGITGTEILF

SHVVTQFSVMCGQTIMVLAFAFLTVVEVNGDVGWVALLTVLTGLCGMCFGFVVSCAVDNE

RNATYMALGSFLPIVMLCGIIWPVEGMHYILKFISFFLPLTKSTESMRAILARGWTITNP

TVYYGFVSTLLWIAIFLTLSILLIKFKKG

>FoABCH-09

MDDIELEEGRTAPTTATSNGHSNGGYRASTLGRIRDELSTALRVEATTELSAGANGCTPN

GGSNGTSNGPSIPPICGTRGAAIVIKDAKKSYPLQSGIVLDGFNMTVPEGTIYSLLGASG

CGKTTVLSCVVGRRRLNAGTVRVLGLPPGTPRSGMPGPTIGYMPQELSLYGGFTVAETLK

YFGWVAGLPSDVVRERTTKLLSFLEIPTEDRLVRDLSGGQQRRVSLAVTLLHRPPVVILD

EPTVGVDPVLRQNIWDLLLRMTREGNTTVVITTHYIEEARQADTIGLMRQGILLAEDRPD

RLLARYACQTMEEVFLKLSLRQQRALVQAQAAAPVTVGALPVPDSPDAAREDAPEPVVDS

KDVLTVVDDPSLLQCKPVYLQNQNNTWACLWKNFTYMKRNPVPVIFVIILPIIQCGLFGM

TVGRDPDDLRLAVVNDDVQCFPAMGGAVEGSVGFPGYPANCTTKTLRHLSCRMLDLIDDD

RISIEQFSSVPLALEAVRAGKAWGALHFPANYSRFLAARHVWGRYASDKALNHSEISVWM

DMSNQYIGHMLQASLSLSVLRFTRDIQEGCHWNTRATPIPVSIMEPVYGQLEPSFQDFCL

PGMIVTTVFFMALLVTVGSILEEKRDGLMARSLSAGVTVHEVLASHVIVQMILMVISMII

VLVMLLCVFQLTNQGPLFWVIALCFLQGLCGMSLGFFISCLCDSDMAATYMGVGAFFPLI

FLSGMLWPVEGMHVLLRSVGWALPLTLATSSLRSIMGRGWSIAQPAVYLGFVSTAVWIGI

FMCITLITIRVQKRIS

>FoABCH-11

MDTQSGGPHAPAESGAAVAGVRAAAVVVKDARKSYTRGGTPVLNGLNMTVAEGSIYGLLG

ASGCGKTTLLTCIVGRRPLDNGAIRVLGFGPGAPGSGLPGPTVGYMPQELALHGMFTVAE

TFRYFGWISGMTTAEIRLRSERLLKLLDIPFPDKLVGELSGGQQRRVSLGVALLNKPPVL

ILDEPTVGVDPTLRQSIWDLLLNETRNARTTVIITTHYIEEARQSDAVGLMRNGTLLDED

APEALLRRYGCASMEDVFLKLSVRQQRERHGTSDSHAIALVETTPQHATPRPQDEQADVQ

SRPVVVDDPSLLQTRPVVLAGHSNVAPCTWKMAVFTWRNKSVIGILFALPIIICVLFGFA

VGGDPKSLHLAVVNDDFTCCERNLPAFPSDCEAEDPRGLSCRYLQRINKDMFDVDDYDDV

DAAQRAVREGRAWGVIHFPSNYSHSLVARSREGRDVTATTLDSSDISVWLDMSNRQVSEL

ARSALATAALQYFEDVQRGCKGNPNAKALPMQFNEPIYGKRDLYFPDFFAPAVILSVVYN

INCIYTMSGFIDDRKDGFMERSISSGVRVTEILGAHVLVDAVMMLIVTGITLVFMCGFLG

FENNGSLWILIFLCLIHGFCGMFQGYFISSLFTSDFEATFCCMGFYFPTLLLSGLMWPLE

GMHPLLRNSVWLLPLNLATTAIRSIMNRGWGLGSPDVRNGFLSATFWLAFFVLVTLTTLR

LKKSFAT

>FoABCC-10

MDSSKRVGNPNPRDSASVVSKLFNFWLASLFRRGWKKTLEVEDIYDPPKRDRSELLGDKL

NMYWERELAWQSEKKGRSPNLVRALTKTFSTDLILLSFPLIIEMSLNVIQSLLLGKLLSH

FRPTPLISREDAMIYAGSMMTCVLISIICRNRYEITAYHTGLRIRAACCSVIYRKALKLS

SGAMGDMATGQVVNLISNDVSRFDIMCIFFHYMWGGPVLSAVIVFLFWWDGMLPALLGLC

VVFAVTLAQSYIGRLSSTFRRLIALRTDERVRLVNEVISGIQVIKMYAWERPFLSLMRIA

RKRELFQLRKVAAVRAVFMTFHMCTTRMALFVAVLSLSIFGTQITSDKVFVIHSYFNILQ

WGLTGMFVRGTAELSESYTSVKRIQQFLMKPEFHSDTIDNSIRSSIKEKNGKSDESETLL

NGKTSSIQSSDSASSLVLENVEAKWNSDSATQLTLDKISIEVPKGRLLAIIGPVGAGKSS

ILQAILGELKLSKGCVKVGGKISYASQDAWVFAATVRQNILFGSPYDKKRYDKVVRACAL

LKDFEQFPESDLTVVGERGVSLSGGQRARINLARAVYREADVYLLDDPLSAVDTHVGQHL

FDECMRGALREKTVILVTHQLQYLKNADLILLMQNGRSLAKGTYQEIINSGIDYAKLLEE

EEEDTDDTKSETSQTNESLLACQDPTLRRSISRERKKSNVSSFTSSVQSVCSEKSEDKEE

QQATSPLVDKLEASSKGMVKGSVYMQYVKASGSLMLFLVTVFFFLITQFAASGTDYWVAF

WTKIEDSRNASVQENNMSNSSNIQADNSTFRSFFPYLDLNTENCAYIYAGIVLATLVLCI

VRSTVYYSLVLTISRRLHEFMFSAVIGAPMRFFDTNPSGRILNRFTKDMGSVDEILPKII

LDSSQVILNLAGAIVLTVITNYLLILPLILMFIILHFIQKFYLKSSKELKRIEGIVRSPV

FSHMNVTLQGMVTIRAFGAQTLLKKEFDKHQDLHTGAWFLHLSGSNAFSLALEMSTLCYN

IIVLVSLFASTQVNSGGEVGLAITQCMMLLGMLQWGIRQSTEVANNMMSAERILEYVQLT

PEQGELKEEKKPAKEWPEKGCVKMNHVFLKYSLEDSPVLKDLNLIIQPGEKIGIVGRTGA

GKSSLIATLFRMAIVDGNIIIDDIDTGDVTLARLRSSISIIPQDPVLFSGTMRYNLDPFN

EYPDEKLWKALEEVELKDLLSKEMGLNSRVLERGSNFSTGQRQLVCLARAILRANKILMM

DEATANVDPQTDKLIQTTIREKFRSCTVLTVAHRLNTVIDSDRVLVMDNGTVVEYDYPWK

LLERPGGIFRGMINQLEGHGAEQLRQAAKQHFKELQPGVNNESKKES

>FoABCA-04-ps

MGFLLQLRILLWKNFLIRKRNRLTHFRTVPDEPKECKPVGDLITKLTCIMGWNVIKPFIT

GKILYYPDTAVTRAIMMQVNATFNEMTDLSTRTIILTTHFMDEADLLGDRIAIMSEGQLQ

CCGSSVFLKERLGSGYHLTVELEEAKPYDLERKKRLHHEVQSAMPTAQLQDRDDVQESEL

VYTLPDRHNVDAIIALLARLDKRREELSIASYAISDTSLEEIFLRVAGRTEEEVDVGSSR

SDVANVLDDDSSSELLMDPLSGGRLLWHQFAALTRKRFNYARRDPKGYFTQLMLPAIIVL

LGLAFRTESKSGTPQARGTRDTAIQMQPSLLSWPVVSFFELEEPAQGDAWGVKYTESMKI

DGLSASLVAGGTVNKESDLLNRSKLGHECANSPSNPPGADWSVALLPSKELVYNVTGSDT

ETWLMSTNSACQRTRFGGVSFGVQYDNGSLAQLGEFGLADNTRIWFSTNGYASVPAYQNA

INNVILRASLPPDKTPRDYGILTNSWALPADQDEDGEKGIDPGDKNLVVWSTNMLLFAMS

FVPASFVMYLVEERVGQSKHLQLVSGLRPYIYWLQNYTWDLVNFAATEILVIMAFIAFRI

PYFLPCENLWAVILLLFFYGMACTPLLYIFHWFFRTPSMAFLVLLTGNIFIGFVSVDTFL

VMAYTSSVVLPTLEAVLLMLPQYCLGRGFLNVAINHYVGETKRYGRERIPPLDWEVTGKY

LVAMAVMSVVLAVAVFLAEQSTHWWHGLARGSVNVVAQGVANKGDDVLREERRVAEGGAD

QDILVLKRITKRYNRKAPPAVAGVSLGVERGQCFGLLGLNGAGKTTIFKMLTGDTLISGG

DALVSGRSVRADMDSVRRMTGYCPQFDALVPLLTVREHLDLYSSLRGTPDRHRPLAVRRA

IQLFDLGAHKDKNAKDLSGGNKRKLSAAIALVGEPSLLYMDEPTTSMDPVARRFVWQRVR

RVVASGRSVLLTSHSMEECEALCARLTIMVNGKLTCIGSPHHLKHKYGSGYLAVVRCSLP

RVHEATELLLQRLQGATVVESHLHQIKLQLPPHLSLIAIFTALDEARRGGSVEDYSVTQT

TLEDSTVTHFVILLCAVSQNAAQGGGQSQHDIKMMSK

>FoABCB-04

MALLWTCRRNVIGEFVIKMSWKGRLLKLPRNHYFIHPTSSFGSHFTSSHRSLPVQSRVFG

RLSQNAVRHVSSSNTSSVSRAIVYTPEDVSSKKTPLKDASQIAGPTVLTQKNSRVWFHAG

VGHIHEDDGGPPRNGEDKVNSKQMIRAMLSYVWPKDDPDIRKRVKIAMGLLVGSKALNVS

VPFLFKYAVDTLNVTPDGNILNLATASDTVLTYSVALLLGYGIARAGAAGFNELRNAVFA

RVAQHSIRKIARNVFLHLHNLDLSFHLSRKTGALSKTIDRGSRGINFALSAIVFNIVPTL

FELGMVATILGIKCGPEYAALSVGCVGLYTAYTLAVTQWRTKFRVLMNRAENEAGNHAID

SLINYETVKYFNNETFEANKYDKSLKKYEDASLKTSTSLAVLNFGQNAIFSGALSMIMIM

AAHQISAGTMTVGDLVMVNGLLFQLGVPLGFLGSVYREVRQALIDMQTMFTLMTVDPQIT

TKPNTAPFVVTPSNAFIEFRNVSFDYVPGKPIFKDLSFTIPAGKKVAIVGGSGTGSVLFH

DTIQFNVHYGSLDRPMEDVIEAAKLADLDASIQTWPQGYETQVGERGLKLSGGEKQRVAI

ARAILKNSPILILDEATSSLDSITEQNIVDALRRATKGRTSIAIAHRLSTVMDCDNLLVI

DSGRLAEQGTHHELVSNPKSIYARLWATQQGLNMKQL

>FocABCC-02

KESKKAVSVLPALLRSSGGVLLFAVALDLAGVALMFVGPQILGRLIAFTGDASEPAWRGP

AYAGVLLVTAFLQSVLMSQDALRATVVGMRARTALVSAIYRKALRVSSATRRERTAGEIV

NLVSVDTERMLEACDYLPMGLSSIVEIVVALFFLWDLLGVAALAGLASFVVLTPVNAAVA

KLYLRLDKKQMKNKDARAKLMNEILAGMKVLKLYAWEPAFGDMVGRVRKKEVAVMRTSAY

LQAGVSFIFSVAPFLVTLVSFATYVLIDDNNVLDAQKAFVALSLFGLVRYPLAMLPMIFM

MLGQARVSIRRIDEFLNAPDLDEGSVHHDVKDASGDVAPPLSVKAGTFAWGPGEEPVLRD

VSVSVAKGQVVAVVGAVGAGKSTLLSAMLGETEVQAGRVNSLGSVAYVPQQAWILNATLR

DNILFGRAYDARRYARVVEACALGADLAMLPAGDQTEIGEKGVNLSGGQKQRVSLARAVY

NDADIYLLDDPLSAVDSHVGKHIFDQVIGPTGLLKHKTRVLVTHSVAHLPSVDVVVVLKD

GAVSEQGSYSDLVRNNGAFAEFLVQHLQKAEPGDDDFDDEDLKEIKSGLEKAFGKEALQR

QLSQASQASQASRVGGDGKRSRTTSQCSHESASRHGLEPRADAAKDVVGQRLTEEEEVEV

GNVKSAVYWHYIRSIGIPLTIATVLMNVLSQGAIVGASVWVELWTGDPAMSQPNRTDFKD

KRDFYLGVYGAFGVAQAIVFAFGCIASSVALHDGMLRRTLLSPMSFFDTTPVGRVLNRFT

HDIASVDNELPNVIKQTFATLFSCLTTLVVISASTPLFLSVVVPLGLVYFFEQRFFVATS

RQLQRLESASKSPVYSHFSETLTGCSSIRAYGAVDRFVAELESRVDANQRCHFPVTVANQ

WLAVRTELVGNCVVFFASLFAVLGRGAISPGTVGLSVSYAMQITEELNWLVRFASFTENN

IVAVERIRQYEELPQETPCTKKRSHVDSAWPAEGAVEFREYAIRYREGLDLVLRGIDVRV

EAGEKVGIVGRTGAGKSSLTLGLFRIVEAAAGQILVDGVDIAGVDLHTLRSRITIIPQDP

VLFTGSLRANLDPYERYSDEAVWRALERAHLKDLAKGFAAGLNHAVAEGGDNLSVGQRQL

VCLARALLRKTKVLVFDEATAAVDLDTDDLIQKTIREDFKECTILTIAHRLNTIMDCNRV

LVLDKGKVVEFDSPQMLLQRNDSMFYSLAKDAGLV

>FoABCC-06

MDSQTYLEQFCGGKFWDANLTWYTESPDFTDCFHRTTILWVPFALLWLLSGVEAIFLLNS

KKRSVPYTWLNVLKLRPCPEGASSFLCRAMFAWLDPLVYKGYRRPLVLGDIWGLNEADTS

TEVVPAFDKFWLKSGKPSAAGQKGGTDGRQQTNIMYAMCRAFGGVFLAGCIAKLFETTIV

FVSPQILSLLITFVEEGDEPVWRGYMYAVLMLLAATFQTLSMTQHNQRMYIVGMRIRTAL

ISAIYRKALRVSNSLRKELTLGEIVNLMAVDAQRFTELMAYINLVWAAPLQIVLALYFLY

QTLGISASAGIVVLLLMIPVNAWLANRVESLQISQMELKDERVKLTNEVLGGMKILKLYA

WEGFFGDLVQGIRNKELGVLKANAYLNGSTCFIWVCAPFLVSLASFGMFVMIDENNILDA

KKAFVSVTLFNIIKQPFTMLPILISNAIQASVSVSRINKFLNADELDPNSVTHDPREVHP

VIVENGTFSWGPDETPALTNITLRVETNRLVAVVGPVASGKSSLIACLLGEMDRLCGRVN

TKGTIGYVAQQAWIQNATLRDNILFGLPMEKEKYNMVVEACALKADLEMLPGGDLTEIGE

KGVNVSGGQKQRIALARAVYNSADVYLFDDPLSAVDSHVGKHIFEHVIGPNGLLRNKTRI

LVTHGIHYLPDTDLVVVLDDGKIAECGTYQDLIDKKGQFSQFVTQHLNDGKIDSPTAPSS

TGSPLSESVANGHAPLPMIDGADPPKHEELQRKLSRKGRQDSQHRRPSLGGPARKMSRSW

SKVTTTLSSSLQEMDEIDVRTAPKQGERLTEAEVAETGSVSLKVYSHYCRAGGLWLCFLT

MLFNALYQACAVGTNFWLKQWSDDHERTHKENGTLPMETRDLYLAVYGVFGLGQALTLFI

SDVAPRLGAWFAAKAMHDVMLHGVVRAPLSFHDVTPQGRILSRFSKDVDVMDNLLPQQIA

DTLWCMFEVLSTLLVISYSMPIFLSVILPIAILYYFIQRFYVATSRQLKRLESVTRSPIY

SHFGESVTGAATIRAYNVKERFIRDSEIRVDVNQSCFYPSVIAARWLSVRLETVGNLIIF

FAALFAVMSKDTIGGALVGLSISYALQVTSVLNLMVRLSSEVESNIVAVERLKEYGNTPR

EAEWELPEKKPQASWPEKGAVSFLDYKVRYREGLDLVLKGLSFNVSGGEKVGIVGRTGAG

KSSLTLALFRIVEAADGKIIIDDIDISTLGLHDLRSKITIIPQDPVLFSGSLRMNLDPFN

KHSDAELWTALNDAHLKSYVEEEGEGLELEVTEGGENLSVGQRQLVCLARALLRRTRVLV

MDEATAGVDLDTDDLIQATIRTKFKDSTVLTIAHRLNTIVDSDVVLVLDQGRLLELDSPE

RLLKNKETVFYSMAKDAGLVS

>FoABCC-07-ps

MDKFQHKLLKNPRVGANIISLSIFAWVWPTFWKGSRKQLDVEDMYEPLREDKSERLGNKL

EEVWRSTCRSAKKKGARASLLRALVSMFWKKFLAVGLIDAVNQIGLRLVQPLLLGQLLAY

FSPGSTIPVEHAYYYAGGIVLCILLSVILANQSLYWCVHLGMQMRVACCSLVFRKVFVIQ

SYYNILSMTMGSFFVRGISEVAESVVSVRRLQRFLEYEEFDAQDGQVPQQPQQPQQPQQQ

QKGQVNPGFEGGPESGQPAGPKQQTRANGAGGAIQGGNAAPRDGSVLRFTAMHAKWSQTQ

SDDTLKNIDLSLNHGELLAVIGPVGAGKSSLLQAVLGELATTAGGLDVRAKVSYACQEPW

LFAASIRRNITFGSPFDRKRYDRVVKACALLADFAQLPFGDLTLVGERGSSLSGGQRARV

NLARILNRFSKDLGSIDELLPKALLDATQFILAFFGYIVVSVSVDWIFIIPVIVLLVIFW

MARVVYLMSAKNIKRLEGMTRSPVFTHLNATIQGLSTIRAFNAEDLVKVEFDNHQNLHSA

AWYMGVMTSMSFGTALDLLSLSFSIVVIFSFLLLQQSDDFLGGDVGLAITQSMAMTGMVQ

FGMRQSAEVASQLLSVERVLEYADLPTEDKDAASGPAPKPDWPAEGRVELKNVWMRYAPE

DPPVLKGLTLTINPTEKVGIVGRTGAGKSSLIAALFRLAYLEGRVVLDDVDTGSLPLQQL

RSRISIIPQDPVLFSGDLRRNLDPFGEYKDHQLWAALGDVELRDTASDSGGLEMRVADAG

ANFSVGQRQLICLARAILRGNKLLMLDEATANVDPQTDALIQSTIRTKFARCTVLTVAHR

LNTIMDSDRVLVMDAGRAAEFDHPYMLIQKRGIFYSLVQETGHQMADTLAKIARENYENK

ITSGQY

>FoABCC-11-ps

EARVGTTKIPIGACSNVLTAVHIVVNKGQLVGVCGRAGVGKSALLLAASGQLRLTAGQLH

RDESTAYVGEDAALLPGSVRDNVTMRGALHSQRYYRALHCTGLERDVQAMPTGDDTLVDT

TRMTPAQLQRLALARAVYADRETYLLDDPLSALDSASADRIFESAILGELKGKTVILVTN

REQHLRHCDEVYVLREGRVLERGAPADLAESGREYPHVIDSNSREEQWWASREQPEVTRS

TERSRDSDDEDVSGTGAWSELRAVAAECGASPGRLALLALPLLLQSAMYADMLLCFWLAL

LQPRPPLLVLGFLMAASMVVALAFLRTFTVTQALLGQSRRMHDRWIDRLGCATTDFLRAH

AAQLRLAVEKHTWLLDAGLPRALAGVAAWTPLVVFEMALLALASRWAAIAMLGLLLPAAL

LITYVACAASLRMWQRERTCRAALRQLVGASYSSRAAVLSLGREDDMVARFSAVCDQAAT

DLVHAESLPAWLGVRLRAGAVVAGLVTLVAQIWAPPGMVLPPPPPGVEPLANNTTGMPPP

PPGPPPLPIDAGAALGMTALLLCLIGTHLTRAATAALRARSHLAAAAEARDLVNRAEVED

ESLSVKGGAECDDPSLNWPSLTDAGGLVLRDLWRPPELRALTARVPLGECAGVLGPPSAV

LGAVMLRFIRPSAGCVLVSGADIADIPLPLLRAAVVVVPPRPTLFKGTLRWNMDPHGIWS

DAQLWEALDRTGLRELVASLDRKLLTPVGAHGARCPLIPEPEDLQLLCLARAALREWAKV

VVLEAPAGPKVLAAARKLFPQSAVVVAARGPTDVAQCTNVLQIPV

>FoABCC-12-ps

AGLYSKATFSWLTPLLRLGNSSPLELEDLGRIPASESTALHHEKFQDIYNSLRASGVVPL

WRCFLLFSWRMLALSGVFKLSGDMVGLMGPMGISVIVLYVQGVGKDETRLRETALFYVSI

SEFFSNGYVMGMLVFISALAQGTFSQSSTHLINVEGIRIKAALQAMIYDKSLKLPLWSFS

EGDISEETQSDKLSQTHLPKCENKTQSKKESELGGSGQGTTSHTDFGTITNLMSEDTYNV

MSFIWICHYVWAIPVKIGVLMFLLHYQLGVSALIGAVLCVALMTPLQFAIGKKMSANSKS

ISEANDERLRRINEVLHGIKVIKLSGWEEQYESRVQLARNTELRLLNVDSLHWAVMTFLT

HASSAIVTLVTFGVYMMVEKAPLPTASVFASLALFNQLTVPLFIFPITVPIIISAMISTR

RLQMLKYLSVYSLLSVVTMLLSLTSNLVGQLAGARARHRLHRNLLSNLLRSPIKFFDTTP

IGRVIERFSTDIAVIDKKLATSIQRLMQFLSLCLSAVLVNAILSPWFLLPALPMCLAYYI

IQHFYRVSTRELHRMDSITRAPVSSFFSETLGGLATIRAFGQKARFMETMLDKMDTNNNT

FLVLSAANRWLGVALDYMGGTIVLISVMGSLIAAQMYPSVVSASLVGLAINYTLLVPIYL

NWVVKFLADVEMYMGSVERVQQYADMPGEDYNNKGMQVSPSWPEHGDIQFKNVSLRYDEN

REPVIRNMNLHIPAGQKIGLCGRTGSGKSSLVMSLFRMLDVSAGSIVLDGVDISRLPLQL

LRTRLSVIPQDIVMFSGTIRDNLDPNRLYTDEKIWHCLELAQLKCVVENLPGGLNSAVKE

GGENLSVGTRQLFCLARAILQNAAILVMDEATSSVDPATEKAFLSVALSAFAHRTVIIIA

HRLSTLLDCDRILVLECGKVIEDGSPSQLLRFDNGVFSLMLKAAS

>FoABCC-13-ps

MDENEKPISTLSASIGRASYDVNASNKQNGFSRPAPQKSGCSKYTDALKVLLPCRVSSRK

TKNTMPANRAGLFSFITINWLTRMMWKAFRKGLTEEDFYDLADADDAAESARRLESIWGR

RVGKHGAGTSLGSAVWRFCRTRTLISIVLMAVSIVLQFVGPAFLQKLILAFIDDKSQALW

QGLLLILALFFSQLCRNLFFGMAYVINMHTG

>FoABCC-14

TRLQGGLQHLVFNKVLRLRASDDRALGQVVTFCSSEQERMFDAIHMGSITVGSPVMFIMA

VGYSCWLMGPYALIGNIIILLFYPIMGVVAALTSRVRTNTVQLTDKRINIMSEILNSIKL

IKMYAWEDSFADNVKDVREHERVQLQRAAFLQSFSSTISPSITILAAICTFIAYTAAGYE

LDSQIAFTVYSVFNALQFTVATLPYGIKCITEARVSFQKMEKFMLKPDMSTHSDKYVPVF

NSGETALGMKSASFVWDDADAQGVDNSGFEGEKGNETASLRNIEMNVKKGALIGVCGSVG

SGKTSLLSAVTGDMICVGGDMQINGSVAIVTQQAWIFNETLRENIIFGLPYDETRYRRTI

EVCSLTRDLELLPKADMTEIGERGSNLSGGQKQRVNLARAVYADKDIILLDDPLSAVDAR

VAKHIFDKCIKGALANKTVILVTHGIQFLEECDEILYMKNGMVSERGTHEQLKKKGGDYT

HMLSYDQNREEKKKDKKEEVADVKDLDEEPPIDVAGGTLTAHEAHKSLGFKAYIDYFKFC

GGLCIMLILFTLILAFVLTQLFAGLWLQIWLDFGDGKGGTSVINNPNLGTYQIIYGSTFV

LMLVTGSIKGYGFARQLVLASSKMHDTMFRKVMRCPVAFFDVTPVGRILQRFSKDMDELD

VRIPYYLEYVGQGMITCIGQMIMVCIMYPMFSAVLVVAIAIFSFLEVWLNRGLQQNKSLD

SIRKAPVLSHLSSTMAGLNVIRTYGRQDVQLLRFRRRLNRSLASDLIYRSSVRWFTFRMD

MICVVLVALTGLVTVLLRDDASSAKSGLALSSVFAVCNVIPFVMQMASELQARFTSVERV

LEYTSGLTEEAPRRIPNSPKPKQWPVRGDVLMEDVELRYREGLPLVLQKVNADIHGGEKI

GVVGRTGAGKSSLINTLLRFQELAGGRILIDEVDISKIGLTDLRSSIAIIPQDPVLFEGT

MRFNLDPFEDFSDEQIWEALEKSHVKEKVAREEKQLLTLISGGGQNLSVGEKQLICLARA

VLRRNKILLLDEATASVDVETDFLIQRTIREAFHDCTVLTIAHRLHTVATYDRVMVMDAG

KVIEFDTPAKLLGDANSVFRQMSEAAGVTVETMEAAS

>FoABCC-15-ps

TLTCILAVIAIVELGFAVVNTSSGGNSVLLDFVSPLSKIFTYALAAVLVVLNRIQGMRTS

GVLFMFWAVHMVFGAVTTRTHVNRLAREQDLWANGTQPVAREEDPPLFLACSWLVGYPLV

VLLFLLNCWADAEPKLSLYAVKIDV

>FoABCC-16-ps

LFPAARMHYFSLLLLRGAPTCSRAVYREADLYLLDDPLSAVDTHVGKHLFDECIVKLLAD

KARILVTHQVQYLADVSRIAILSNGEIQMQGTFKELLNSDVDYAQMLHLGDEEEEAEEEV

EQDKRPLMRALRQLRSSRRSSSRRSSRASVGSLTLGDEDDEDEEGAPTMERIEATTKGKV

RGSLFLRYFLAGTNALGFIVVIFLFLAAQTAGSLADFFVGFWTNQEELRDYFTSSRNVTS

PSSPEPLTSTNNFIYIYTAIIVCLFVLGFTRSFFFYWVCSRSSQALHDNMFANIIRATMH

FFNTNPS

>FoABCC-17-ps

TWEAASPDFTPCFEKTALAWTPCLFLFAFSPLEVHYIRSSANGEVRWNWLNASKLALTTL

ALVVSVVSLGGAIGDPAGAFPVDYVTPAVRIVSFAWAAVLLAWNRARGVRTSGLLTLFWL

LAALFHVAQVRTELIAETTERAQSERGRLPFLLTMIHLPVLVLVALLNCLVDYEPVSSQY

TSIESRCPVFDASFPSKMLFAWYDRMAWTGFKKPLEHKDLWELNPADRAADVVPRFFSHW

NK

>FoABCC-18

MGAFDDFCGSPYWNTTRTWESDNPDFTPCFEKTALAWTPCIFLFVFSPLEVYYIRSSVNG

EVRWNWLNFSKLVLNVLALLLSAASLGGAIGNPDDSFAVDYVTPVIRMVSFAWAAFLLAW

NRSRGLRTSGLLTLFWFLTALFSVAEFRTEINTETSERTESERGQLPFVLTMILFPALVL

IFLLNCLADYEPVSSPFTKVEDRCPEFDASFVNKIFFVWYDKLAWKGYKRPLEERDLWEL

NPMDLSSGVLPKFFSHWNKQNKKADAKAFATGQPKKLVSVLPALFRSFGGPFLFAAAMEL

VGDMLVFVSPQILKRLMAFTTDLEQPLWKGMVYAVAMLVTALFQSLLLSQYNIRMNIVGM

RVRTALVAAIYRKSLRMSNVARKDKTVGEIVTLMSVDAQRFLDMGGLITMVWSSPLQIIV

ALYFLWDILGVAVLAGLASMILLIPVNGWIANKMKVLQIKQMKNKDERAKLMNEILNGMK

VLKLYAWEPPFGERINQIRLKEMDVLKKSAYLQAGISFAFTVAPFLVTLVSFTTFVLIDK

NNILDAQTAFVSLSLFNILRFPLAMLPMVIMMVVQAGVSIKRIDEFLNAPDLEEDSVRHD

VEQATRDRCPMSVKDGTFAWGPGEEPVLKNINMSVSKGSLVAVVGSVGAGKTSLLSALLG

ELEKQSGDVNSVGSIAYVPQQAWIQNATLRDNILFAKPYDAHQYAKVVDACALKPDLAML

AGGDQTEIGEKGINLSGGQKQRVSLARAVFNDADIYLLDDPLSAVDSHVGKHIFDQVIGP

TGLLRQKTRVLVTHGIAYLHRVDHIVVLKDGTISEEGNYQELMKNKGAFAEFLVQHLQEA

EEVDDEVAEDLQEIKAHLEKAVGKEELQRQISQASLISSEGMRSRKGSVASGSRKGSVGG

SLIRHGSGRRRSSELKPSELPPKVDEVIGQKLTEVETMELGSVKLKVYWHYMRSIGLMLS

GLTILFNVCLQGFSIGSNVWVDKWTDDAKMYQPNLPDFESTRNTYLGVYAAFGGAQAIFS

MTSSYVFAIGCIASSMDLHNRMFRRILRCPMSFFDMTPIGRILNRFTKDIDNVDNVLPLQ

IRQALNLLFSCVSTLVVISISTPLFISVIIPLGIVYFFVQRFYVSTSRQLKRLDGVSRSP

IYSHFSETLTGSSSIRAYGAVDRFIEEVESRVDKNQKCYFPVAVANRWLAVRLETVGNSL

IFFAALFAVLRRGDISAGSVGLSISYALQITLALNWLVRFVSEVENNIVSVERIKEYEET

PQ

>FoABCC-19-ps

ATRLSRTALGETASGQVVNLLSNDVSRFDIMAIFLNHLWIDPTLTLIIAYLLYAQVGWSA

FVGIGAVFIVVPLQSYTGGLSSKFRHRIALRTDKRVRLMDEIVNGVQVIKMYAWEKPFNK

LISEARRDEIKELLKVYMVRGVFMTFMMFTTRVALFSTLVTYALSGDPLKASF

>FoABCF-01

MPSDAKKKQQQKKKDQAKVRQGATKKAGDSKPEENGVTKQNGSNGTEKAPQEMSVEEVLC

AKLEADARLNAEARSCTGGLAVHPRSRDVKIDNFSITFHGCELIHDTLLELSCGRRYGLL

GLNGSGKSTLLAVLGNREVPIPEHIDIFHLSREMPASDKTALQCVMEVDEERVRLEKMAE

QLVANEDDESQEQLMDIYERLDDMSADTAQARAAHILHGLGFTNEMQNKKTKDFSGGWRM

RIALARALYVKPHLLLLDEPTNHLDLDACVWLEEELKTYKRILVIISHSQDFLNGICTNI

MHLDKRKLKYYTGNYEAFVRTRMELLENQMKQYNWEQDQIAHMKNYIARFGHGSAKLARQ

AQSKEKTLAKMVASGLTEKVTSDKVLNFYFPSCGTIPPPVIMVQNVSFRYTDTSNFIYKN

LEFGIDLDTRLALVGPNGAGKSTLLKLLYGDLIPTEGMIRKNSHLRIARYHQHLHELLDL

DLSPLEYMMKSFPEIKEREEMRKIIGRYGLTGRQQVRCRVVFAYLAWQAPHLLLLDEPTN

HLDMETIDALADAINDFEGGMVLVSHDFRLISQVAEVIWICENGTVTKWEGGILNYKEHL

KSKILKENARAEKENNSRKK

>FoABCG-01

RKQILHGVSGEFRSGELTAIMGPSGAGKSTLLNILAGFTLALLGLDTHAETHSGRLSGGQ

RKRLSIALELLSNPPIIFLDEPTTGLDSSSTTQCVSLLKSLAAEGRTVVCTIHQPSALLF

EMFDQLYAVAAGHNVYQGDTKGLVPFLAQQGLPCPAYHNPADFMMEVSTGDFGVTVNALA

DAAAKNANRQPVVLPDATLMDDIICEAPPAPLYMQFYLLLMRNIVIMRRDVSALALRVFM

HVFIGLIFGYLYRGVGTYANDVLANVVYLYGSCLFLVYTGQMAVTLAFPLEVEILTREHF

NRWYKLGPYLMSTLLIELPIQAVLCATYLAPSIVLTGQPLDLFRIIHFYSFATLASLTAQ

SCGFLFGATVPITISVFLGPVFAVVLSIFGFCIWYRDIPNFFMWIYHISYFRAAFQGCVY

AMYAYGRSAMPCDNRMKAHAGGINYCHYKAPRKILNEMDIADVSFWTNTASIVVFCLIVY

IITIFVIWFRLNRR

>FoABCG-02

MKLDGGVDVAGGDADGLLSSGRRKSSTVKVTITPSQPRTLSHLPKRPPVDIAFSDLTYSV

TEGRKKRSITINGHERNLSQFRKLSCYIMQDNQLHANLTVEEAMQVATALKLGSDVSKSE

KEELIQEILETLGLSEHVKTMTSNLSGGQKKRLSIALELVNNPPIMFFDEPTSGLDSSSC

FQCISLLKSLSRGGRTIICTIHQPSARLFEMFDHLYTLAEGQCVYQGSTQQLVPFLGKLG

LACPSYHNPASFIIEVSCGEYGDNIRKLVTAIKNGKHDIRTGQPFPAIEALNNSTSPSNN

LKMTNAQNGNNLVSFNKDATDKDLSEKKGNGIMLASFKSNEKENGINGDGSQVVIPVDLS

ANDKSDNVAASLLDSGIVDTKRRYPVSEWKQFWVVLKRTLLFSRRDWTLMYLRLFAHILV

GFLIGALYYNIGNDGAKVLSNLGFLFFNMLFLMYTSMTITILSFPLEMPVLLKESFNRWY

SLRSYYLAITVSDLPFQTIFCVIYVTIVYFMTSQPPELERFAMFLSACLLIAFVAQSVGL

VVGAAMNVQNGVFLAPVMSVPFLLFSGFFVSFDAIPIYLRWITYLSYIRYGFEGTALATY

SFAREKLKCSQTYCHFKNPETTLEELDMLDKDFRVDIVALVLIFIVLRVSAFVFLRWKLR

AAH

>FoABCG-07-ps

MDVASGELGDQAEAMERLSSEKVWRAERAVVRDPSQETTEKGLRKAMVLIRKPSEFDRFW

VLLHRSMIQLYRDWTVTHLKLALHFVVGIIFGLYFTDAGWDGSKTISNFGFFMATCVYLG

YTSMMPAVLRFPQEMNTLRKEQYNNWYKLRTYYIAYIIANIPIQMLFCTVFVSMAYFISE

QVADWTRFVMFLSICQLLSIMSESIGLFLGTTCNPINGTFLGAILIAIAMLFAGFLVFLA

HMPSYLSWAADLNYMRYALEGLSLSIYGYGRESLECPRSEIYCHFRVPGTLLTEVGIEDG

RYWIDVSAISAMIVVLRFVSFVTLKRKVACGERSS

>FoABCG-08

MEAVIAEAAAAPQGSRQANGAAAPAAAPTALATSKTLPHLPKREPVEVKFQDLAYTVSLG

FRKGQKEIIRVNGRFCPSQLIAIMGPSGAGKSTLLDILSGYRISGVTGNVMVNGRPRNLD

RFRRMSCYIQQDDRLHSLLTVQENMSLAAELKLTRAVPRAEHQAIIDEILNTLGLLDHRG

TRTGQLSGGQKKRLSIALELINNPLLMFLDEPTTQAVSFGKQISVLLTRGYIKIKRDQTM

THMRLFVNVIVGLLLGILFWQAGDEGSRVLDNWNLLFTILIHHMMTTMMLTVLTFPTEMQ

ILRKEHFNRWYSLKSYYMSINILELPVSIVCCTVFTVIVYLMSAQPLEWVRFGVFLGISI

LVVLVAQTVGLLVGALFSVVNGTFLAPTLSVPMMMFAGFGVSLKDMPDYLKWGPYVSYLR

YGLEGYVGSIYGRNRTTLGCADQPGKDFSYCHYKYPTKFLKEISMDGEMYVMDLIILGVI

LLVLRLSVYPLLRWKLFAMR

>FoABCG-09

KRRILDSISGEFKAGQLTAIMGPSGAGKSTLLDVLSGYTSSGFHGRLLVNGSPRDSRTFR

RQSVYIQQDASLHPQLTVSEAMNVASVLKLGRRGPEIDEVLYSLGLLEQRDTMTSSLSGG

QLKRLATALELLSNPPVMFFDEPTSGLDSSTTKQCVGVLRRLARQGRTVVCTIHQPSATV

FEMLDQLYLVVGGRCAYSGHTANLLPYLAGLGLRCPPYHNPADYALEVAGGEYGDFSEEL

VKASDNGRCVMWTTTYTAKQALPGVAGTSAATDAAGRLSRQSSSVASSSSQSEVKPGRCS

RQKLTYPTTFWLQLYVLIKRTFLKGLRDRMLCHSRLYIHLVAGLLIGVLFLDIGGDAAHV

FHNFSLLFFCLMFLMFTALSAMILAFPLEMPIIAREHFNRWYSLKSYFLAVTMADVPLQG

VCVCVYTAVVYYMTAQPMEPLRFLQFLLICALVSFVAQSVGLLVGAAMKVEHGVIFGPLV

ILPFTIFSGYFVIQRDAPASMQWLFKASYLKYGLSGSMLAVYGYGRAPLERCSEDYCHFK

RPDKFLHELDLDEGNYWLDAAILFAIGSVLRLGAYFVLKFRLRRFRQ

>FoABCG-11

MPSSGAPQGAWELNTMERRYSVPPGSHHRLAIPSNSEDLHAWSIYRQNLNSDFTDSALGS

SEKSPLPYGNFQLRDSTVHSILSHPRYGPNRGLEVEARRSKRQLLLQGVSLEVRGGELMA

IMATSAEEGTALLDTLAGCPSRLAGRVHGDILLNGQCMPRRSLSNRVAYVRSERTLNPYL

SVEQTLYFTNWLRQPGYQGAKTDTKDRVAALVEDLGLEQVRHTKVWSLTTSEQRRLAVAA

QLLLDTDVLVLDRPTAGMDIFDTFFLVEFLRQWAGNAAGSVTGRAVVMTMHPPTYEIFTM

LSRVALLSKGRLMFCGRRRDMLPYFASAEYPCPSYKNPSDYYLDLVTLDDLSAEAMLESS

QRIEQLADVFRRREEPLSDPGAPAGLPPSVRSANVLKQAAALLVCWTVYAQPGALSRWLA

HLVLAVGLSLFLGVVWWDVAASDPQLLLGDRIGYLYAMLVVLPWPLLLQQAGWGDTWWRA

VADDIRDRLYGRLVYILCQILYGLVPSAIIWAGYVVPAYTMTGLQNQATDPNAFYIYTGY

SLLYLVALQSLLVALSATLRSRRAAGLVAGLVLLPLALLSGPLLHERDMEPWLQYAALGS

PMRWVLPTLVRREYASDVVAASVANIMCRNKQVQHQDIIVQLPCPPPNVTAALQYHGLGP

SLDPGALAWPAIALAVFWATFAVLACFVFPLTALRHRARRRPTSHRP

>FoABCG-14-ps

MRMDARAPPAARSRRILGLLAHLGLTPCAHTRLSLLSGGERRRLALAVQLLVQPGPLVLA

LDEPCSGLDAEGSRVVLEALRREAGRGRAVLCTVHQAAPDLLRLFTHILVLASGNVVYHG

PLDRAQPFLAEAGAGGPAGWTPSTPLNSMLQQLNDPAAHLDTARVCAAFSFSEQACLLED

YPTSEEVLARDAIRASDIKHISWIGEVGWLLWRESVDGRRNKSRNLAQMGMFVLTATIMS

LAFMEVSLSSLAGVQSVRGLLYIIVSEVVFTHSYAVFHTFPVELPVYLREAHLYSSSAYY

VAKVLATVPRSLLEPLLFVSVIQSRVDLTAGGGVATFALLLGILFLTALTASAY

>FoABCG-16-ps

IELSDSDAQFVNILSASVDNGKLTKTPEGTLQAVDKGECLSECSGSSQGEVADAELAAQP

LHRPRGREPDLQFPSSFWLQFVVLLHRMLQQTTRNKTAMRIQFAHHTMCSLMVAILYFNV

ARDGSQFFTHMKFSIGLVLFYTYTHVMVPVLVYPYDVKLFKKEHFNRWYSLNAYYAALCV

AKMPIQFVLSTYFLSIIYYFSGLPLEFDRFAIMALTGILISLVAEGMGHSIGAVFSVTNG

SAVGPMLIAPFMGLAVYGFDFAPQIPWYINALMKTSFLRSGVAAFVLTMFGFGRETLDCN

EFYCHFKNPRYVLRFLDVEHQSAWGEIGNLVILGVFYRLTFWVGLRWRTST

>FoABCG-15-ps

MDVGFQNLSYKPRSWRIGGPQKTILHDVSGRFKSGELSAILGPSGAGKSTLLNALSGLTT

RGVDGSLTVNGCQRDPRTFYKSSRYITQEDLLQPMLTVHELMVMAARLKLPTGTSRQERL

RVISDILAKLGLLHASATRTENLSGGEKKRLSIALELVSNPPVLFLDEPT

>FoABCG-17-ps

MYTYLKFGLPRVFPPSASRGRDGSSGYDSSDD

>FoABCG-18-ps

FRKESCYIMQDDMLNPLFTVHEIMTMASEFKLGSSLSTKAKQLVIEDILETLGLSATRDT

RCCRLSGGQRKRLSIALELIDNPPVMFLDEPTTGLDSSASLQVVSTLKALARGGRTIVCT

IHQPSATVFELFDFVHVIAKGRTAYQGSALNVVPFLQTAGMPCPKYHNPADF

>FoABCG-19-ps

VLTDPPLMFCDEPTSGLDSFMAQNVVSVLAGMARKGKTVVCTIHQPSSEVFSLFDKVLLM

AEGRVAFLGAPDHACRFFSQLGAGCPSNYNPADFFVQLLAVVPSREESCRQTIEMVCDAF

QRSEYGLKVQHDADAGNELMAKPPHSWEEELFLVDTSPYKASWWAQFRAVLWRSSLSVFK

EPQLIKVRMLQTVMVALMIVLIYYGQTLDQDGVMNINGVIFICLTNMTFQNVFAVINVFC

AELPIFMREHANGMYRVDVYFLCKTLAETPVFLVLPLLFISLPYYLIGLNPEPTRFFLAC

LIIVLVANVATSFGYLISCISSSVSMALSIGPPIVIPFMLFGGFFLNNDSVPSYFKWLSY

LSWFKYGDEALLINQWQGVDEIACTRSNATCPRTGHVVLETLSFHEEDFWLDIVNLGALI

IAFRFFAFLSLLSRTMRS

>FoABCG-20

SADKLTSTVLALPRRNSNVSGTIQTNGQPRNLRLFHKLSCYIMQDDLLQPRITVHEAMMV

AAELKLGSEMSARQKALAVDEVLETLGLTPCRNTRTEKLSGGQRKRLAVALELVNNPPVI

FLDEPTTGLDVVAMKALVDVLQLLARGGRTVVCTIHQPSASLFQQFDHVYILNGGNCVYQ

GAAHQLVPFLDACQLSCPKHYNPADFVLECLSPNNAKTMAANTSNGKLCKTSSEPAKHVS

SYRTGDSAASLLAAAHAQTSALSETEFATSFWTQLSVLTRRYFLQTRRNMVGLFVQMFGA

VLVGASLGAVFYGKGDDVTRPFDNFKFCIGVLVYYMYCPFMVPILLFPSELVIMKREFFN

RWYGLKSYYIALTFSTLPWQCVCGFVFSTMCYVISGQPMVWSRYLWFLVTGVTTGMVSEG

YGLVLGSLFSVTNGSTLGTFSLAPMLVLSVYGMGYGKNAELIYEWLMSLSYLRFGLVGFA

LSLYSGDRDLMHCDEDKVVYCHYANPNLLLRDLGMEGLSSYMQMLNLLVYMVVFRLAAFL

ALRYRLTVEFSSRILNYLRKIIPHK

>FoABCG-22-ps

MVSEEQAPLLEGGGARRKGSGHSLHEGGGGPGSDDGGGGPPRDTILSPMGCERLVYSWSD

VNVFSGDEQRGGALQAARSLWSRCTGRRAAAPHRKHILKSVFGVAYPGELLAIMGASGAG

KTTLLNTLTFRSGNGVSVTGRRCVNGVRVGRNALAALSAYVQQDDLFIGTLTVREHLVFQ

ALVRMDRHIPYARRMRRVDEVISELALSKCENTMIGIPGRVKGISGGEMKRLSFAS

>FoABCH-02-ps

MAVSVREAWHEYGRWRRRGRGRRRPSAPETVLQGVNMTVRRGEVYALLGRSGCGKTTLLR

ALVGQLALDDGEVRVFGRAPGDPLAGVPGRAVGYMPQEVAVHDGFTVGETLAFFGRLMGM

KSAAIRKSTDTLVSALRLPGSHELVGALSGGEKRRVSLAVAMLHAPPLLVLDEPTVGLDP

LLRH

>FoABCH-04-ps

MATLPTSAVAPDIAISIRNAYKEYGGGWGLNGKQAPVKILKGLNMTVASGTIYGLLGSSG

CGKSTILKAIVGQHRFTSGDLRVLGSVPGSDGAEIPGPAIGYMPQ

>FoABCH-05-ps

DFVVWPAVAVVSRSTVLVADGSVSSGCSYGLLGASGCGKTTLLSCLVGRRHLNSGEIWVL

GGKPGTKGSGVPGKRVGYMPQEIALLGEFSIRETMMYFGWIFGMPSSEIQDRLDFLLNFL

DLPSQNRLVKNLRRVSFAVALMHDPELLILDEPTVGVDPLLRQSIWHHLVQITKDGNKTV

IITTHYIEEARQAHAIGLMRSGKLLAEESPSALLATYNCPSLEEVFLKLSRKQGSQPGTD

ANMTNNISVASLNWGNKKDEPVFVTEESGVVGLNFHQSKEMLINESNGHLDQLNKNKAGK

SEPVVDCDDCTDCFKLTTRGKMRALLQKNFLRMWRNVGVMLFIFALPVMQVILFCLAIGR

NPTGLHLAIVNHEMNYSTMECPVYSNCSLSWLSCRYIQQLDNETIVKDYFETKEEALDAV

RNGDAWGALYFTENFTDALVARMALGQNSDNETLDQSEIRVWLDMSNQQIGLMLNRDLQF

SYRDFAQDLLSQCNNNPKLADIPIQV

>FoABCH-06-ps

MLITEANSLAERGGPETVVSTSPTTVDVEKEAGRDVGNGRRLVTQQSSVWTRRQQAVCVR

HAFKHYGSNKNPNHVLQNLNMTVPKGTIQFKEPIYGSNNPSFTDFVAPGVILTIVFFLAV

ALTSSALIIERMEGLLDRSWVAGVSPFEILFSHVVTQFVVMCGQTALVLIFMIAVFQVEC

KGDMFWVIILTILQGLCGMCFGFVISAICELERNAIQLALGSFYPTLLLSGVIWPVEGMP

VVLRYISLGLPLTMATTSLRSMLTRGWKISEPDVYNGFFATIIWIALFLTISMVVLKVKR

G

>FoABCH-07-ps

MNETSLASTVSAAPPAGGVSSPAAATTPASSSPIKAAVVVRGARKCYAANKWVLDGMDLT

VPEASIYALLGASGCGKTTLLTCILGLRHLNAGAVKVMGVRPGAKNAGIPGPLIGYMPQE

LALSENFTIGETLTYFGRVAGLADEKTRERSDELLKFLDIPAKKSRLLSDLSGGQKQRVS

LAVTLLHAPPLLILDEPTVGVDPILRQSIWDFLVDLTRTSRTTVIITTHYIEEARQANLV

GLMRRGRLLDEDGPDPMMAKHDCKSLEDVFLNLSMRQDGPEVGAEADHELQPLPPPPAPS

QFSHVKYVDVKPIPFTPHSNILACMWRAWLQYIGNLVRNALFRAALKFHDDMQQGCGLNS

RFSAIPIEEMEPVYGSREPSFQDFALPGVLQCVMYFPAIFYATANFMEDKLTGVMNRAGA

AGVTQFEMLAARAILDVCLNAVVTVLVLFTAFVAMGFTNGGSYLTLTALLYVQGLVLRGG

PLQRPLRSNFLRDGNFPRQHLPQR

>FoABCH-08

MPVADVELDSVDPWSPGYVTPASTPSPRSARSCSPGGPDGAPAVPVRGDAAAPAGAAAAA

ARAHHAAPSSRVVAVSVREAHHNYSQYSFNSLLTGKPAVTEVLKDFNMTVQHGNIYALLG

SSGCGKTTVLRALVGQHRFTSGRVRVFGKEPRSRGVGVPGPTLGYMPQELAVPDGFTIRE

TLNYFGWLAGMTAASIEENSAALLRLLDLPPRDRLVSNLSGGQQRRVSLAVALLHSPPLL

ILDEPTVGVDPIVRQSIWEHLLGLTSSGRATVIITTHYIEEARQAHVAGLMRRGALLAED

TPESLLSALSCVTLEEVLLKLCLRQDIDEVEVQKMLHADGMNMAHELEAIKDKSLRVSAR

DEEEGGQDGDDVKAKQHLRTSRTSTKVKALLWKNFMWIWRHRIISTTLLFLPAFIALVYT

GGVGHNPRDLNVAVVNGETDCADWKGVGLVDCAAPRHLSCIFLDQMRAQSLQPVLYAEEA

AAERAVLRGHAWGVVHFGRNFSDALVERAGITVSVGNPAQGDEAVEHSAVNVRLDMSLQT

IGHLLQMTLSTVASQALEVFNSQCQANAHMIRPPLQFMTDLRPSDRDPSFLDYSLPGLIL

SVVFMFGFTTTMSALLPEKTDKILERSLV

>FoABCH-10-ps

MAPDGARVAGLEAHLDDQGHQGHLHHHGLPGLDDDSAAVLVRKAYKEYRNKALFRKPVSN

MVIKGMDMTVRRGTIYGLLGASGCGKTTILKSIVGQNHLTAGEVLTLGRVPGTPGCGVPG

SAVGYMPQELALYIGFTIAETFRYFGWLTGVPRKELAARSASLLKFLDLPPAHCLIGQLS

GGQQRRVSFAVALLHQPALLILDEPTVGVDPVLRQRIWDHLMDITSDGRTTVIITTHYIE

EARQAHTLGLMRQGVLLAEDAPDRLLERYHCATMEDVFLKLSVLQIRTSPDDPEEVVEAR

DNNQNTQSSYPAEPPPPPEPASPEAFKHGHGNHSSPRTMVPLVTLSPHAAKSSGPALGPA

LGPALAPPHQRPPTPSSTRMSCLLWKHVLWMRRNLPMVAFLTLLPVVQCMLFGMTVGRDP

KGLKLAVFNAEGDCADPALDTIDTFNCSSPSLLSCRYIRML

>FoABCH-12-ps

MKGKAYAAVHLQDNFTENLFARINDGQHATNETLLGSEVNVWLDNSNSYITTLMQSDMTY

GTIELLQNTLGHCDYNPAVAGIPVRFESPIYGDINTDFLDFASPGCALTLIFFMSLASAV

GAILSEKNEGLLDRSLVMGVTMREVMWSHIIIQTMMMILQTIIVLVGFFVVLQITNDGPL

FWVVLLCVLQGFCGMCLGMLVSSLTDSDREATFVALGSFFPFIVMSGMIWPVEGMHIALR

RLGWALPLTLATTSLRYMMSRGWGPAHPEVWYGFVSTIVWIGIFLSSVLFVLRKRR

>FoABCH-13

QELGLYEEFNIKETLWYFGRIAGLADETIEAQTKSVMKLLDLPTHGAAVGSLSGGQQRRV

SFAVALLHRPALLILDEPTVGVDPVLRQSIWDHLMEMVTTKKTTVIITTHYIEEARSAHA

IGLMRNGVLLAQASPEDLLTRFHCDNLEDVLLLLSRQQEELGAIEGHEDESTSFVPAAPV

ENENLYHVTPSEGTSLSQGSTMWSRFYALVWRNITWLRRSKWVIGMLVILSVVQSCTFCW

SIGRDPKDLPMTIVNEESNCTEPGDILWHGRCGNTTQISCIYIEMVSNKDVDFDWYTDYD

EALQQVKDGKSWGVLYFRHNFTKSLYERIENGRQTSDADVEASTVDITMDMTDAMISQLL

KKKLQDLMQVFIKDMFSDCGYPRMAGALPVEFEEPIYGESDPEFINFASPGMICSLVFML

AGNSAMCALLPEKRNSILERQLVTGVKLMEVLYAHYVVQLLLMALQTVIIFTMMFGVFQL

DIEGSFMLSFTLMFVLGLSGMWFGFLVALHAPTEWAASLIGMALSFVFMFVGSVSWPWEG

MHESLRHVSHALPLTYPVHGLRVIMGRGWGLDEWDVWAAFLVTTGWVAFFLVLSHVSIRL

QKV

>FoCCE-03

MLLVAVLVWLSVPGSAWAATRGPREPPTVEIPGQGVVVGREVSVSRNQKSTLYLNIPFAQ

PPTGNLRFDKPQTDPLPSWTEPRNASAFGPACPQMLSKLDEQDRILTNFLQDKGLEMSED

CLQLNIFTPDGNPPAEGWSVMLWLHSGDFQTGATSLWDGSVLAVRQKVIVVTAAYRLNIL

GFFTTLDGNAPGNWGLWDQVAAIDWVQSKIAAFGGNRKNITLFGHGAGGVSVGLHVISPM

SDGFQAAIAMSGSATTPPDSSSGGVAREERSTVEAIRKLAERFYCRPSTADVVSCMRNVQ

PVQVLVEAALQFTKYLPVVDSKYNNVSQPFLSDKPATMLEMTPKGTPLMVGYTDHEDAMQ

MRKELEGGLSSSKYTELLVDATSADLPQPESENNDTCPINGEQAMDAVTFYYTPIPATED

TTLLRQKFMEFTTEKKYGAPAFLQASFSSKVSPTWMYRFDYRLKSTAVGDAVPDWINVPH

QYELPFVWGMPFWTSLPSQVVWNSQDKKIADIIMTLWGSFAKTRNPTQQIQGGAVKWEPF

TEQNPGVMVLDKNFSMTDVSNFDYRSFAFWNTYYPKVIESLVGCCNQTDGASTVAPRGLH

LALPVLAPLLPWATWRV

>FoCCE-05

MQRKRGCCLLGGVLLLLVLGAVIGVIVYFVTRGSDEPLATVKQGVLRGLQKESETGSHLP

YVAFLGIPYALPPVGKLRYKAPQAPASWGGTRDATQQMPKCSQMQPDGITVEGSEDCLYL

NVYTPALPNQGNDNKSPMPVYVFIHGGKFQSGGASWEEFGPDNFIEQKDIVVVTIQYRVN

SYGFVSLDSEEMPGNCAIKDAIAALQWVNENIGAFGGDPNLVTIGGQSAGSCLAGWLTIL

PETKGLMRGAILQSGSAYHNWAYGEDHVDMTLELASIVAGHSVTDKSEAERVLMAAETEP

LTKATWQLINEKMKTQPELPYRPTPERRVAGREPLLITQDVESYFLQPPSPHVPQMLGIT

NEEWRFYYYTNNPTSDPSKDEELLQNLHLHVPKDLIPYADTRKILGLPEKNIDYNTVIKR

VEEAIRNEVSPSCKLGCVLKKYFDGIWMATDTHRLGAYLAARNETVYLYRFGVRTRLNRP

IIDPLPDDERSAAMHGDDLNYLWHEQKNKGKEPTSLESLTTKRMVAMWASFIKNGKPVTA

TNDLLPVDWTPNSGTNLEVTFLDIGENLTLRTEALAPVMLPFWYALYKEYRTPSNIK

>FoCCE-08

MRREARGRRVPCAPFQDAPRRHASLPRAHHVQPHASGQALEVTLPHGGVLRGREVQQNAA

PPFQAFLGVPFARPPLGSLRFQPPQPAEPWTGVLPAEQEKPPPLQIDYYLQKVLGSEDCL

YLNVYRPPGTEASSRKPVLVYFHYGLWSLSNASMNEVGPELFVAKDIVVVVISYRLQALG

FMSLNTPLAPGNMGIKDAVLGMRWVRENIAAFGGDPERVTAFGFCIGGYIAQTLQFVEAA

KGTFQRSMIFSGVMNNSWYGALPGEVVRERTLKLARLLGCDGNASDEELFEFLQSVPAME

LFSKAWDCMASPPPADRANPPFLPIIDGGCVARPEDAVFPAHPFKMLRDGMQVSPVPTMF

NLTRDEGQVVRGDVKLWTGTRVDKDDLDDHFAKLDRSLPCDLDLELGTPERAAVTREVRE

VYFGDTKHPTMHQMYTYLGDIVNGMPVVRAARHHACHPAAPPTYLMSFAVESTFFNRSRV

KYGECIVPGLVVHADDMSYYFGAGTEPQGPGPEPALAHDSLEAVTRRRMMAFLSNFIKYG

DPTPEDEPEPELHGLRWPRLPASTSAPYPTLSFTAHPSVEWELFGARARFWQDVYERYFT

DGDTAVSRSPPSYTDPDDREDGGKDFTRGIFM

>FoCCE-09

MTPRQLKKRALMVGAVLGGVIFAAAFAAYFAFGGSSYEDSTSPRTLSPPEAKVTEGTLRG

KWANIPDYSVQYAAFRGIPYAQPPVGMLRFKAPLPPAMWSGVRDATKEGSACMQVSDFAE

IIGSEDCLYLNVYSRGRPNVSIFENKNVYWQHCKRRGTDFGPELFMRRPIVVVTINYRLN

AFGFLNLDNDVVPGNAGIKDQIAALRWIHANIDKFGGDPSAITIGGQSTGAASAHWLTLL

PESRGLVKRAILESGSALHSWAYHEDNIAVALQLGSNLSKHKDVSLEEVARLVMERPAAE

ILSATDSIELNGIQLPFSASPERRAPQEGGQVPLITQDPEKYILTDRSPVPILLGMNSRE

WLFNFQYLGYARSPQLIRNKINNLVSIFPKSVMPGSDTAAYLNLTSTHSLEHALDLVHQH

FFKDNGADPNKTSSVFQSFLDDVYVGTDVNRLAELRVRVGQGRTQTYVYRFGVCAKYNTS

PFQNPGQVAEGASNGDELQYLWYRGNQKFEGDGLASTTLNYMVETWSFYVGSGIPQPFTK

WVPNEFRSKPGDVIYMNIGENVDVKEDPIAGQNAPFWLDLCQQYRDNSST

>FoCCE-12

MQTSSENYGTRAVHNKGELVVAHFGPPVKWGGQLPLPGVLAGWRADLQGKMRPDSLQLER

DVYVQTTYGQVQGFKTYLYDNPDPLSGYRPGMTPVERVQGVVHTFLGVPYAQPPINDGRF

KPPRAHRGWQLIQAVDFGPACPQPVRYTGAAKGVRDMDEDCLYLNIYSRTTVSGLAQKFP

VMVYIHGGEYYKGSSNSFPGHVLAAFYDVVVVTINYRLGALGFLCTADINSPGNYGVLDQ

AMAINWVYDNIDAFNGDRKSITLFGPGAGAASAGLLMVAPRTRDLVTRVIAQSGSALADW

GLIIDQYRAQNTSRVFGKLMGCSIDSSWKLVDCLRRGRSALELGNAEFHPAVGPFPWGPV

FDRNFTKPGDSWYQGWKERDWHFLNRTPEELIKHGQFNRGLSYMSGVTTQEAAYIISQNE

SLVSVHYEVNERFMEQKIREMVARFNYTLNPGGTYEAIKYMYTYHPDPKNVTHIREQYIH

MMSDYLFRAPNDKLVKLLVEKNIPVYMYVLNTTVEALKLPEWRKYPHDTEHLFLTGAPFM

DVEFFPKDAGYERLMWTENDRNMSHFFMKAFSDFARYGNPTHSQILGLHFELARPGQLKY

LNLNTTFNSSIFLNYRQTECAFWSDYLPTVIGHLVPTYPPTTEWWEPRQPLQIAFWSMSA

ICLLLIVAVVACCILWRNAKRQQDRYYSGDLLMMRDESEITGIGNASENPSHLYEYRDTP

TIAEKQPQLKVQMHSRPVPPEPTKTSSVKTGSNQSLKSLKDSVNGFPAGEPRVETRPPLP

QPAAAVPVPTTRPRATLSRTHLEGGIPQTEV

>FoCCE-14

MAGPMRVSVLLLALLVAAAVVADAQAKEEPKRGKTVKKQLKMEPEVEADGPEVELSVGWL

NGRWLTTKAGRRVAAFTAVPYAKPPVGDLRFQAPVPAEAWTGARDSPPLQDVPVCMQHEN

AATTQGMTRSEDCLYLNVFTPKPSNGTQDGTALPVLVWLHGGALGSFNEGGSGAYGPEYL

LDRELVLVTLNYRVGALGFLSTGDAAASGNWGLKDQQLALRWVREHIAALGGDPDKVTLM

GHAAGAVSTHCHLMARSSGGLFHRAVSMGGSALTAGLPQSGAAARRLALRLGAKLGCRDA

VGEASTAKQSAELLSCLRDKPARDLVAHESELRDEGLCPLVTWGPVVEGGEHYEGQVVVP

FLDMHPIDTITSGLALDVPWLAGLNINEGGLWAAAIQGSKQLEAIDANLNVMAPVCFGFN

DTARPEDLPEIMQEIREFYFGANNLTRENIYDLSNMFTDGVVLSSLDEAMRLHAAYLPAT

APAFLYLLSHRGQHSAAEAYAQRLGVVHGDELFHLFPMGVDQDAADAGASKKFIDLLLNF

AITGNPTPPGDERYTFPWKPVESEELEFIEFSQNGRLLKGRGLLGHRAHFWSALALSERN

VQNMEASVRFTRVRDEF

>FoCCE-16

MTSAIQAALLGAVLVAAVSAAVEPSDPLVQLEKGAVRGTTTQSDSGRTVFAFKGVPYARP

PVGKYRFKDPKPTKPWQGVWDATQFPPKCMQFDRYPVSAEDCLYLNVFTPRLPAAGAPLL

DVLVWIHGGGFMFGGSSDWGPEFLLDKDVVLVTINYRVGPLGFLSTQDEVVPGNMGLKDQ

SAALRWIQGNIAAFGGNPNSVTLFGLSAGGASVHYHYLSPLSAGAFHRGWSLSGAALCPW

TQQEEAAAKAVRLAGLLGCPTDGVTSRDLVACLRHRPARAIAAAVPAFQDEAEIPFSPFA

PVAEPAGTMGAFVDRPPIDVLRQGLANDVPWIVGITTEEGLFNAAEFLGAGQSHKLQRLD

QDWLRLAPLLLDYNDTAGTEAQRDAVSRAIREHYIGQRPLSAATNEFVAMVGDRIFNAGV

EQAAREQSQANSQPVYVYLYGYRGENSLSGAISKPKDTSDWGVSHGDDAAYLFRWECLPT

GGNPQELEVRKQLISILTTFARTGVPSGLPGSPGWAPLDPASQDLYILRMDKNAELIFDK

VPELGQTAFWNSLPISENAPDLVSARDEL

>FoCCE-17

MSGGTLVFGALCALCVLSSALASAPVMCEVPGLGVLRGRHIPGNTTFYAFEGIPYAQPPT

GLLRFQPPQPAPRLVYLDARGQRHPCPYLNDKGVLIGSEDCLYLSVFTPQVDDELAMPVL

VMLPGDDWTRGYAGRWKPHRLMDKSIVVVTVQSRLGALGFLSSGDASAPGNLGLKDERLA

LSWVRRHINAFGGDPNRITLAGVGAGAAAVMLHALSNPGLFQRAIAVSGSALSPWSLTEN

TQEAVDAPTRRLAEAVGCDCPISTSAPGNLTSSTAHAANATLASSNNATAAVNATTPATR

HANLTEMVECLRDKPVADLVYGTARLQGWAGLPSSPFGPVVESGEANDTFLGAAPARLWG

DEEYAVTVPLLLGLSRSEGLQVLAPWPAAGLLNSASQLSQLDTEWTRLAPTVLAFNKTAA

EDKAAASEQIRRRYLGVVTPADAKDGLLRLFGDRHIVLPALRAAALHAKKHPVFLYRFAW

SAAQGTSALGGPAGGSDASILLMEPSPTEATRLPAQRAMAMDILDLWASFIDKGVPVLGN

TTLPRVSGAPSGELPYLELVRPGVQRLGNWSVQDTVSFWKGLPLLENKKT

>FoCCE-18

MTASAAPALAATALAAMLAVGLLAPALAIVAEFPSVTTNNGSFILGQRLKSAGLRGGTAP

RDYAAFWGVPYAKPPVGERRFKAPEPSPLPTVVDAKLPAPACLQLDRQPSAIIGSEDCLT

LQIFTPNYDPAAALDVIVLIHGGGFMFGRGPRTGLQYIMDYDVVVVNVDYRLGVLGYLSF

EDAELPGNAGMLDQVEALRWVQENIVYFGGNNKSVTLAGFSAGSASALMHTLSPLSKGLF

HRVLALSGSPLNPWAQQPSALANAKRFAELANCTGANTAAIVSCLQAKNATELVALETHF

QGWQQNPFSPFAPVVDSKSKRPFMPDQPARLLENKRGPTDVPLLFSVTSEEGLFPAAEFI

LNDTLVQELDENWADVAPKLLHLHDNRTGETPSLMHQIREHYFQNASLGHSHKQLVQLIS

DRDFFAGVNDYVLTAAATQSSPVYMYSFEYRGEFSLTHFLTGGNTSNLGVAHADDFQYFI

EFPWSGVMRMPTDLDMRSQLLELWVSFAKSSIPVLRGANWTPQPKTKSDVLHFLKISGPG

NVTMVQAKDFGQSAFWKSLQIKENGNNSGRASASLSLSLVSLAILVFCKLS

>FoCCE-20

MQLVEVQTKQGLLRGQRLTPPHKNVPQFDAFLGVPYAKPPVKELRFKAPQPPEPWEGALP

ALDHRADSPQFDTFITQKNVGSDDCLYLNVYSPQVPSSVRHPVEGAPWPVMVWFHPGGFT

NGTARMSQFAPEHLVARGVVVVTVTYRLGAFGFLSLRSPKLPGNAGLKDGVASLRWVRDN

IAVFGGDPGRVTVFGCSAGSALVEYLMLAPSARGLFHRAISQSGSATNQWAGVLPPDEAR

ARAFRLGEALGCRTEDEDELIAFLQAAPTMDILDKSFDVVDPEKEGPKGLLFPFLPVSEA

GVESDDGEEAFLPDEPRKLLASQRFFPVPWIVGLTDCEGLLTVMDDQNGAGITCRRLDVV

EDDLERVVPAELELQKGSPESKAVAEGIRKLYFKGGKVELPGYFNLYTDLLFTAGVTQAA

RMHAAHPNAAPVYFYLFSVVGGNNILTAFLMRSWPELKATAGACHGDENGYLWGRRAPLP

PEPEWEPHSLEATTRRRLTALWTNFARTGVPAPPSDPDVEGVDWKPLPATSDVKAPIEPY

LDIGVSLGVPRAPLWEERVSFWDGVFDKD

>FoCCE-21

MSLSGLAVNVSYSFVFKVSLTVSGDFVLRAKDNKTPTSVPPEVTVGEGTLRGEWVNIPKY

NKSFAAFRGIPYAQPPVGQLRFKAPLPPASWFEVRDATKEGSACLQQELLSKKIIGSEDC

LYLNVYHPQNSSSTWLPVYFFIHGGSFLYGSGSHTVYGPDFFMLKDVVVVTINYRLNAFG

FLNLDNDAVPGNAGIKDQVAALQWVSRNIHKFGGDPHSITVGGQSAGAASAHWLSLFTVL

TNGVILESGSALHSWAYNEDNFDVALDLGSRLTNGSTVSLQDVAHLVMELPAMDILAAAM

SLANERMYANATQIPFATSPERRNPLASRQEPLIRGDPESYILTDSMGTNPIRMLMGMNS

REWLASIYLHGWAAHPDYMRKRIKNLVTAFPKSVVAGHDTARYLNLNNAFTLEQAVDAVY

QQFFKDNATDCDDICVFKEYLDHVTIGIDTNRLAELRVRTDRLSAPTFVYRFAVRANYST

SPLLDPAQAAGGAVHSDELGYLWKMNRLQQDIEGDDLSSLTLRRMVDIWTTFIKHGHPED

WENRGSWIPNEFRRKPGDTIFLNIAENLTVREEPIVGKSSPFWLDLFQKYRDYKST

>FoCCE-22

MASVSRIAILLSALVIFLVLVGVFFGVYLNNNGSSSSTTPSPTTPSPTTHSPNTPSPSPS

APPVTATIVDCGVVEGEWVESAAGGSYAAFRGVPYAEPPRGELRFKGPRPPAKWDGIRSA

KEEGNTCFQSGSVGSEDCLYLNVYTPKLPSNSDPSPALPVYVFIHGGVFQYGSGSAQGLG

PDFLVAQDIIVVTMNYRLGVLGYLSLDTEEVPGNLGLKDALYALKWVNKNIAAFGGDPSK

VTLGGQSSGGVSASWLSILPATKGLIRSAITQSGTAVSSWGLNLRNVEYAQAIYKQITGK

DTKDLTDIAKLVYSANLKELYDAAQAATLQLQANKGLGTDISFFYLPTLELRPDGAEDKL

ITKDPESYMILKEANDIPTILGETKREWALFLQTLDLYKDDKVLKSAIQDLITLVPNTII

PGTQTQQSLQITDEVEKLIKNDETKYVQNIKDHYFLQNTDAECSPYGDLCKMGKYLDHVY

IMADTDHFLRLRAMYVKSQTYSYVFNLKSDYNGDYSSFPVYLKDVVYHGNDMNYVWKLST

IKQDYNGNGIASKALRRQVTLLANFIKTGNPTPEKSELITELWLPVNSGEEEQYLEITEN

LTMQKGSMSGQDGKFWQEIFSHFRVDKSP

>FoCCE-23

MTLTPTRRKKLVLIVCGAVIAVVIAAVVAVIVLVLPDQDEEVKSSYPEVTVAEGRLLGQW

VDIPGSDVSYASFRGIPYAEPPLGPLRFKPPQPTSWDGIRTANYDGNQCMQEGGGSEDCL

YLNVYTPKLNGSDKLLPVYFFIHGGSFIFGSGSRNVYGPEFFLLQNVVVVTINYRLNAFG

FLNLDNDIAPGNAGIKDQIAALQWVHRNIEKFGGDPKLITIGGQSAGAASAHWLSLLPES

RGLVHRTILESGSALHSWAYNEDNFDVALDLGSRLGRSYLTSVEEVAQLVMEVPAEAILN

AALSVVNDRTMRNTTQIPFAVSPERRAPFQGPPALITQDPEKYLLADMNPLPMFLGMNSR

EWLGGFYYYGWAAKPDVMRARIQDLLSVFPKNVIPYGDSSAYLNRTPTPNEWDQAVWTVR

QKFFKDNATDCDDTCVFKKYLDDIFIGNDIARLAELRARNARSETYLYRFAVRAEYSLSP

LQDPDQAQGGAVHSDELGYLWKMEELNQTLNGYDLAPTTLRRMVSLWTSYIKNGTPTPEH

MTDEYYKEWKQAFLTNEHRRAHGNVIYYDISDRPSLMQQPISSVNAPFWLDVYQQCREGK

ST

>FoCCE-24

MSPTRRRKLVWIIAGVVVIVAIAVAVATYFALRDPGDEAPTPDSAVPEVSVIEGRLRGEW

VEFPEYNIRYAAFRGIPYARPPTGELRFKAPLPAAGWSGVRNATTEGSACLQQDIYTQKL

TGSEDCLYLNVYSPRAEGSTELLAVYFFIHGGSFLFGSGSSLTFGPEFFMLNDMVVVTIN

YRLNAFGFLNLDNDVVPGNVGIKDQIAALQWVHRNIDAFGGNPDLITIGGQSAGAVSAHW

LSLLPEPKDLAHRVILESGSALHSWGYNEDNFDVALDLGSRLTNGSTVSLQDVARLVMEL

PAMDILAAAVALANDSMNSNATQIPFATSPERRSPHDGYEAPLITQDPEKYILADKNPLP

IFMGMNSREWLFSFYYYGYAARPDYMRFRIAHLDTVFPKNVKPSTDTAAYLKLTNKYTLE

RAVAKTYQYFFRYNNSACDDTCVFKLYLDDIVIGTDMNRLAELRARSHRSQTYVYRFAVR

ANYSTSNLVDPAQATGGAVHSDELGYLWKMSVLQQRLEGDGLASDTLRRMVKIWSSYIRS

GIPDPGPDHEGKWTPNEFRTLPGNVIFLNIAETLDVREEPIMGQNADFWLFTYSQYRDDK

SA

>FoCCE-27

MFTVLGDVVKGADHFCSIAFKNIAYAKPPVGDLRFKNPQPHLSQRRRSGRQGGDPKCAQV

EVPSGKMQTNEDCLYLNVYTPVGDLLDNLPVMVYIHGGGYVYGSGTDAENGPDFLVRHGV

ILVTLNYRLAAFGFLNLDNDDIPGNMGLKDQQAALRWVQTYIKFFGGDPNKVTIFGNSAG

SASVHYHTIAPSSRGLFRAAIMQSGNALAGFAYTEAHLTMAQRLSEQLGAKTADARQMAR

AFRNASAQDILSATLRMIQADPNLQLGNPAFCPSPERRQEGGQEKFLPQDPESLERGLEP

GAYVPTIAGLTGQEGNFAFYFGGLKKQPATVAALAQNPALMLSPNVAPYPDTARMLGLPG

ANGNYTITVSPEQAAAFGARIKEEYGLAGANNGDFIPLLGDLFTASATHRLAELRLRKGE

GAAPLHLYHFLEDGDYNWGKVSFGITEMGATHTDELGYLLHITSPRDLNQTARGGSRSSK

ALHLLTTLWTDFAKHATVEHHGWPAAEGSAQDARYAMITDHIAVGSGLGGERMRLWEDVY

AELRAEKAPRR

>FoCCE-30

MGYSWSQHRPEAEVASGKLRGKSDVTAAGTRYHAFMGVPYAKPPVGELRFKAPQPVEPWT

GVRDASEEGSGCLQSSMPFSSPEGISLKHVIAILRHLPTLAHRVLANKKQSEDCLFLNVY

VPADTFPAPEVYSEQANEQEERAERPLRPVIFWIHGGAFKYGDGNPDMITPDLFLDKNLL

VVTINYRLGPFGFLALGTEDVPGNAGLKDQAMALRWTYDNIKSFGGDPAAITVYGESAGG

ASAHYHNLSPLSRGLFRGIIASSGTSACHWACTQNAAKCARALTRELGLQADTPDEIAKA

LRSVSGHKLLHAAEKMAPVFDDATDELMFLPVVEPDLPGAFIAEDPVTMVRDGRSSQAPL

LTGVNSAEGLVWVLFALTSEEKTYKDINERPYYFLPPDLRRGLTPEQRDKCTREILEEYT

GGSAFTKENVSDWIELYGDLMFMIMTANVTHLHASANTAPVYLYKFDFAGKYNALKAMCK

AKLGKGVPKIPGAAHGDDLFYVASVRLLPLPTLTPDSREEVCRRRMVQIWTNFAETGNPA

PNGADDKVLPVEWLPCTESSMPYLYINDDLEVRTGKLFPRQAFWEALYEKYLGKPVPITH

VRSGPQ

>FoCCE-31

MTVPLGRRVAAAALSVVIALASASAEALAASASASGSPSSPSATTAGGEVRGLRVPAERG

APAFDAFLGIPYAKPPLGPLRYKFPKPAQPWDGVRDATADPPACAQINDRSATKEVVGQE

DCLYLNVYRPEGAEALPVVVVVPDGGLLLASGRMDHFGPHFLVAQRVVVVSVNYRTGPFG

FLSLDSDEIPGNAGLKDIVAALRWVKTNIAAFGVDPSKTTVLSWSSGASLVHILSLLRKP

RELFSRAVLLSGHGISPRAYTERHLERAAVVAEALGAPSNGSHQEVRRALMEASAEALLR

ACDDPRVRLLGLPLPSPERRNTKGAEPKLLRQDPESLLRQPEAPPLPVLMGLAGREGSFP

YKIWGLGERTPGALDELLPRLLPADVLPGADTAGQLLVRDEYAAGGAIGNNTDTFIEFLG

DAFLYASAWRAVGFMANASTEAAPLYLCRMAVDDAYNYGKKKYGVVTEGASHGDDIGYLG

RSDRDPELNQNLSGGGVASKTLILMTKIVGNFAKGLAPIEGWPAVPSVGEVQSWPVLQVG

PGSRRHVEHADGKRQPFWASVYRDLRSASSTISRS

>FoCCE-32

MGAVCSTLMARSPEAEALPDERSLSTVEIYSLEVTLPHGGVLRGREVQQNAAPPFQAFLG

VPFARPPLGRLRFQPPQPAEPWAGVLPAEQERPPPLQIDYYLKKVVGSEDCLYLNVYRPP

GTEASSRKPVLVYFHYGMWSLNNSSMNEVGPELFVAKDIVVVVINYRLLAFGFMSLNTPL

APGNMGLKDAVLGMRWVRENIAAFGGDPERVTAFGFCIGAYIAHTLQFVEAAKGTFQRSM

IFSGAMNNTWYGALPGEVLRERTLKLARILGCDEKASDEEVFEFLQSVPAMELFGKSWDC

MASPPPADRANPPFLPVIDGHCVARPEDAVFPAHPFKMLRDGMQVSPVPTMFSLARDDGH

VIRGDTKLWTGNRVDEDKLDDHYAKLDRSLPYDLDLELGTPERAAVTREVRQLYFDNKRH

PTMHQMYTYLGDVFSGMPVVRAARLHAAHPAAPPTYLMNFAVESSFFNRSRVKYGKCIVP

GLVVHADDMSYYFGAGAEPQGPGPEPELAHDSLEAVTRRRMMAFLSNFIKYGDPTPEDEP

EPELRGLRWPRLPASSRAPYPALSFTAHPSVEWEIYGARARFWQDVYERHFTDGDTAVSR

APPSYTDPDEREDGGKAYSRGIFA

>FoCCE-33

MACCKRWTLAGLAVAVLAAALNLTAWDTIWPFAQVVRGVRVTQGQLAGLAQTTDAGFHYL

SFLGIPFAEPPVGNLRFKPTRKGRPWEGELQAFRNGPSCMQVRDRSFKLHGLLGRNAGWN

EFLLLVTSIPRVLKLILSFRQSEDCLFLNVFTPVTRLPTDHLRPVLVFIHGGGFVRGSAH

SAIYGPDYLVERDLVVVTFNYRLGAFGFLSSNTSHAPGNAGLWDQTLALEWVRDNIRQFG

GDPDRVTLYGESAGAASVHLHVLSPQSRGLFHAAVLSSSTALSTYVMAEEASKSAVLARS

LGAAEDVVADPARRIEFLQSKTSTEVSRGFADCLNEDDTRQMITKLPFGPVVEDCSDGLD

HFLCAQPEDALQAGAFNKVPMIIGLNSYEGTLIHALDSVEEVLEKTNRDVRTFVPRGLYP

RMSDAQRLAVGRRIRDQYIRPESPNDTLALIHLYGDSLVNHGMHVAMRWHVAHSTPAAPV

YIYHFVNNIFGFYKFLYGASLDGPGHADELGCVFYIHILNMGLAQGQEKLSVARDKMTEL

LANFVYRRTPTSKDSKSLNIPWPPSTREKNSYLEFGETFKIKTDLLRERMQFWDSVYSEI

THHPAYNN

>FoCCE-34

MSGVEVEKQEKTEKEEKTEDKKEIEEEEREKMLNEENKKQTEEAKEKEDKEKEADDKNEA

EGNKVSTGEVERKRKALEEATKKDAKGHIPIGGIRMPGFLQRRSKAEKGKEPDPEEGAAR

EGFGPRCQRSLRRMLPAMSSLPKVTLPRGVPAVRPHLASLKLANPFAKKKKERDVEAGPT

IAKAKEAGVEIVPDTLKNGKDVDVDGMETVQLDAEEKGDPEKGAAGGADASKDIPLLERI

RGYKHKTVAGGVIAFVLALIIIVSIAVSGPHHDFAKGPIADGKFATATVSCGLVQGILES

GAYTFRGIPYARPPVGKLRWQPAQPMNNLEHCWNGTFPAHNSSNYCWQIFPEDQSMNGAE

DCLTLDVFTPQVSFENPLPVVVVVGSSHMTGGSTTPLLQPTAKLSRAKEVVFVRPNFRLG

VLGFLTTNALTKSTYPPTSGNYGLSDIKLALEWVKQNIEAFGGDPKSVTVLGYRAGATLV

TALTSSPAAKGLMKRVWVTSGSADFPGRPLSESQTDSSPFLARLQRFCTEDRNSSISGIT

ADCLRSIEVEDLLDAMPNEWRPSMPDLPAPVGTAGMPSTHQWLALDGDILRTHPSEVWEK

WENEGSPLQIVIGATAHSDATVELYRNLSDASSNNANVNEQNIINHIEHSVLGELNLTKE

AINRYNATWPGLAAIVSDIRTVCPLNVLATKMPSAYFYVTTFPISVSKSDDELQYQEPLA

ELGIDVAAIMGRVEGATPEERRYITAMQQLFYQFVRFGELHHSSDSYTGRILVVDQDTLP

KVEYPNCNFWIQKDVVPRFGHRN

>FoCCE-35

MNKLVAIVVLASQLAVLLHLTAAASHHRVRRIVGGRAAQPPPVDDPVVFVYKDDHDARVL

GTRERPGGFYVYRGIRYAEPPVGLFRFQRPRPLRLEGIVNATTAPPPCIQPHPDDPLKTL

GSEDCLYLNIYSSELPNGTSDGLPVIVWIHGGGFRRGSANQYGTGHLVDKKVVVVTLQYR

LGSLGYMSTSSQQLPGNVAMFDMALAMNWIKEYISFFGGNPKNVNTMGQGTGASSAMMMG

LSPMTQDNVKGIVAMSGSALSPNAVDAEPLDATDELAIQLGCPTQPHLAMVRCMQEAPAD

QIVFADETLQTLRLASQGFVGGLGGLLSPSPVVEGKEDSRFLPSFLPEQPKETLEKGNFP

NISLLTGVCKDETGRAIKGGFQKELDKNLRAVPDFLNKVLLKDLAGRAAGVINNVQNLFG

GLASSPLFQQAEKAVGPITKIVEATTDALFNLPMFQASQLWSKKGGKAMVYSFEHTSPRA

AQAGQKFLGGLPLIQAGLDAPAALNETFHGDDLAFVFDARPLEGMKGDATRGQVSLTEPE

DMKVRDIFTSLIADFAHSGQEEGNKKKEKPRFPLPLPSFGGIGGGGKQENNFLVIDQNPR

VGKNFRGCQMAVWSGLSGPQLPTDSCADILKNAVGAVAGTAGKIAGGLTGGLGGGGLGGL

GGGRQGGKPQGPGGLFG

>FoCCE-36

MHNPRTMRFAAAAVACILVGAGVGAGVFFILRHSFPNNPECKEPMATVESGALRGLCRPE

RLGHSKYAAFLGVPYAKPPLKDLRFKSPRPPEPWSGVRDALMVSARCTQGDVDKQIGSED

CLYLNVYSPRLEQNQKQELLPVLVHVHGGAMKEGSGTYIRPDYFVDQGIVVVTFNYRLSN

YGFLNLDTDDAPGNAALKDIIAALKWIKRNIRNFGGDPEKVTINGCSSAAVLVHWLCLLP

DTEGLFRAAIIKSGSALVSWGYSEQHRPYAEAAVKFMQQWAPGADAKVLLQNSPSLLLDL

AFTYASLHNMVTSPENAAHPYVSLEKRSGGEEPTLIVRDPESYVLRPERSSVPKLMGITS

CEYEPMGDMYFMLGELPPTIIREMVPRSMIPMQDARRELGIESYSLTYDDETADLRAVLF

NLTAVDPTCNIICRWERYFSDTYISVDTVRALRMHAQRDPDTPTYAFYSEFPIDGSCGTG

APSGGGTLHAHDDKLVWPETLDNVLAWNTSDPRSLTILRQVTAFGNFVKFGKPVPKTDPV

VLTTEWQPMGKEGPDVFMRFNANWTMDSGDMLGRFGPMWKQLYDKYRYGKGI

>FoCCE-38

MRILQVMCALSLAIRGTNPILFASHKWQRGVSSAGFDPSNRRPHRAEAPENPEVSVEQGR

LLGRTLVSSKGTKYHAFSGIPYAAPPIGDLRFKVTRVPRTIEPLHGDELPGLWRARPWLL

QVVTNILSAGEDCLFVNVYTPQSALGKDAGGGKGLPVLFWIYGGGFVWGEGGRAFYDPDL

LIDHGMVVVTFNYRLGPLGFLSLGRGSGVPGNAGVKDQRAALHWVRRNIAAFGGDPDKIT

VYGESAGAVCTHLHLLSPSTKGLFRGAIMSSGSAAHFWATETAAKAARRARHFAPAVNCK

GNLPARELVRCLRNVRPQTLVAFHENAVRPGDRGDRGGRGRLRGAHHHGVPSGDAQRRQA

SQGAHPHGFQQRGGAHHILCVESLRLADRNITCLVPDDVLPHLSLAEAEEMGHRVRRLYM

GHEKPITLANKNDDKFFEMITSFFIYLDTLKLARFLAADPTAAPLYIYRFEYVGALNTFR

NLLRIQTKAACHADELNYVWRVNVSPDSRTLKPNATEYKVRDMFSTMLANFVKTGSPTPR

GSPLDPRLSWPPFTAMEQAYMSIDKEPRVERNYRRDIIGFWDAIFAERVGGPIWTKIVGM

EERRRAVLDTQTNLVHGGHQRIAPKSTR

>FoCCE-41

MSPAALLVAALLAAAASASAPPEAQPPAGPLRGKWTATTSGRPIAAFEGIPYAAPPVGPL

RFQSPVAAAPWSEVRVADRPGSMCLQRNIFLKVAHLEGSEDCLFINVYAPETVSRAQPLP

VLAWVHGGGFFAGVSHEYGPEFLLERDVILVTFNYRLGPLGFLSTGDGVIAGNFGMKDQV

QALRWVRENIRAFGGDADKVTIFGESAGAGSVHHLTISPLAKGLFHGAIAMSGSALSPWA

FRPPRVAAEFAAKVAVAVGCPTDSGSEALRECLMGKSAEDIISTDLALYEWGAHPMCPFT

VTAEPAGPGAFLDRHPAAALAATPPSVPLVMGFVSHEGCVAAAPIITDPKLQKELDEKFV

DIAPIIFHSKNLDKSINNAIFNKIRKFYFNDSAIDQTTKWDLTKMFSDHFFICSILESVK

IQKQSSATSPVYLYDLAYLSKRSFAVAFGAEEAGSAMCHGDDLLLMFPLQSMLPGEQSAA

DKAIGKELIDAIVTFASTGVPTRDGSWRPVQNALQPEYIHLPEAGKSSMRSGYEPERVAF

WRDLPVNLAFAASDPPTRDEL

>FoCCE-43

MAALALGLYRATEYHNYWNSLSPVVEVQQGPVQGRSALTESGTEYFAFQGIPYAAPPVGP

LRFQDPAPPASWAPEVRDARDDGPVCVQAPLKLPMGVPKNVSALDVVRFLGAVPALVRRA

IKLTRQSEDCLHVNVFTTKVGGDGGSPLPVVVYFHGGAFVHGDNGVDVVGPQYMMEHGVV

LVTAKYRLGPLGFLALGEAGKPLRVGNVGLKDAAAALRWVHDNIARFGGDPAKVTVMGQS

AGGALAHYLTLMPSTKGLFHRVIAMSGTALHHWAYSTPQKALERTRHMVERLLRPEAAGG

PHGPAAPDMEDPEDVLRFLREVDARHLAVMDNFGDGQVEGRWLMDEFPFQPTMDGTLLSA

PPLELIRAGRFHQVPVMLGLTAQEGFFAFMSAPAKNKSVEEQMAMVEADFYCALPEYMRE

ALPRGSDALRKAEAAVREFYFGKSKITQSSLDQFVDFYGMLMFEMDTHRAMTEVSAVSTE

PVFAYNFTHSGRLNLFRRMVMFSFEDFEGVSHCDELNYLFHMDALPHIPLQSTDPEFQVR

KNIITLFTNFVKTGHPTPTSSSVPTRWRQVTKREKPVMQIDVLNKIQDGYKPASFKLWSK

FYQDLKI

>FoCCE-45

MASADKGSPERGGGDGSDKPAGSSPTVRTRSGEVRGLRERTHHGTVYFSFRGVPYGAPPT

GEMRFKAPRPAEPWTGVRDATKPGPPSYQAFSIPPFGPASWSRREVVSFIKTVPTLLRVG

IQLLSASEDSLYANVFTPQLPTTGRCKPMPVMVFIHGGGFRLGDGDFLYGPDYLMDAGGL

VLVTFNYRLGPMGFLSVGTEAAPGNAGLKDQVLLLHWVRDNIAAFGGDPNQVTLAGESAG

GVSVHLHMLSPQSRGLFHRAIASSSLAGSEWAHEENPLEHARQLAKHLGVEDSDPDAMVR

RLRYLPARDIIRAFDSHMHHPFTPRLSLGMYFVPVVEPPHPGAFLTEDPLRMLAEGKQVQ

MPFLTGHNSREGLLIFLGIGDGKLGPKPRRRQRALMRKCAKQRPEAFLPCDFHRGLDAAD

RDAVGREVLRHYFGEEGPGPDKFDEFIDLFGDTMFAIPVLRGSGAHAASSTAPVYHYYFN

HDGELGAFKATFKAHDYEGASHADDLGYLFRSGSLAACHRRPLHRDEQCRRNMVTLWSSF

VRTGVPTAAGLEGDAWTARSAQQLRFLEISDKLVMRTGAPYPRLSFWNDMYKKYLHVSLF

>FoCCE-46

MMTRKRGCLLGGVLLLVCVGVVIGVIVYLTNNRPDPSPLEDLVVTIEGQGRVRGAVNVAY

SSNVSYYAFYGVPYALPPVGVLRFKAPRLPIPWSGVRNATEETVEPCSQMINGLFAGSED

CLYLNVYTPALPRETINVNLPVYFFIHGGLFQGGSPSTEMYGPDFLVQQDIVIVTIQYRV

NSLGFGSLDTEEIPGNCALKDIIAALKWVNRNIAKFGGDPNKVTVGGQSAGGALASWMTI

LPETKGLISQAIVQSGTAYHNWAYKEDHIEMGLRLASLISAQNVTDIKTAERILMETASE

KLTTATYQLIAEKMKSQPDLPYQPTPERRTAVPNGEPLLVTHDAESYFLEPPAPHVPQII

GICNEEWRFYYFNNNPKSDPQVDEQLLENLQAHVPKNLIPFSDSRRVLGLSERKDFDYDE

VIEKVKQGIRNSVSSSCSLGCVLKKYFDGIWMATDTHRIGAYLAARNETVYLFRFGVRSV

LNQPFSIDPVPEDEKNAAQHGDDLPYAWLFRIHSESSLSEEAALARRRVVAMWTNFIKYG

KPTPVKTDLLPVDWLPNSQTNLEVTFLDISGNLKLRSESLAGEMVSFWNGLYNEYRSNVA

VSTHTNSV

>FoCCE-49

MYIWALEVTLPWGGGLRGRVVEPLTNAPPFQAFLGVPYAAPPVGKLRFLPPQPVEKWSGV

RDAIEERDGCLQIDYYSKSVMGSEDCLYVNVYRPPGTEPHCRKPVVVVYSFGAWSLLNMS

MNEIGPEFLVAKDIIVVTVPHRLLSFGFLSLNSPVAPGNMGLHDCIAALRWVNENIASFG

GDPERVTVFGLCAGGLIGHTFQFLESTKNLFQQNIFISGALSSAPWYACAPGDVIRDKSL

YLARNLGCTSNNDEEILTYLQSVPAADLFSKTWDCMYSCPPSDRVNPPLSPMIDADAYAD

RNDALWPAHPVKMVRSHTNRPVPTLFGFCRNDGWLNWGPTRMWEGIQANSSNLKDRYAML

ERTLPDDLDLPYGSPEREEVLEEIRRFYFAGTKSPTLHQWYTYTGDLLARGFYDAFRQHS

VHPGRPRSYLYCFSVEGRNNRTSRMFKCPPGIVAHMDDMGYIFGCRDKPPGPEPAWPSTS

IEALTRRRIVSLLSNFIKYGDPTPASEPDPEPEIHGLRWPALPPRSAAPYPCLDINTSLS

IQHEVLGERVRFWEDMYDRYFTTGIT

>FoCCE-58

MMGKRVLWATVVLAAALFLGASQTFETQDGECTTPNVTTRQGQLCGRLVTSPTGVLYHSF

QGIPYAKPPVGDLRFREPQPAEPWTGLRGALQPGAVCLQSGQGAVLETLSPTVRQAVILF

KALPSFLSGFFRQQSEDCLFLNVYSRELRPARPLPVLVWIHGGGFHMGSGGPDLYGPEYL

MEASSAVVLVTLNYRLGPLGFLSLPSGGVPGNAGLKDQAAALRWVRDNIQAFGGDPSRVT

IFGESAGGTSVQMHMVSPLSRGLFHGAISQSGSALNPRSSTSTAAGAERARRLARQLQVQ

ASDDADLVRRLRLLPGKVINSMSESVLTDEERSRGLAVHPFVPTAEQDEPGAFLPATPQE

LVYRGLSAPVPYITGINSLEAGFMMNRERDLAGVHLDREMENLVPSDLGVQHGSPANTEI

ARAMKRAYFGDSDRPSQMQMAEFYSDLFVNWVAHRTVNLYSRLESPALYVYRFSHDGGLT

LSRRLFDKKLPGVFHGDELAYLFTQSVVPSGNESEQDVLVRKRMAELWTNFAASGNPNGD

SPLLTTKWMSANNQSQLYLEIGADLAMRRGLFSPNVGFWDTVYTVREGGART

>FoCCE-62

MHAAAMGGDGCTCRCACACTPRRCPAAARLLSPLLLAAVLLQLLLPLAAAAYPQVHAHDL

RSEDGPEVRAEQGVLRGVWVLGAAGTGIKFAAFRGIPFAAPPVGPLRFKAPRAPEPWEGV

RDAVAEGPVCPQGQPDSAEGVLGDEDCLYLNVYVPAASLAGETTTDAPATTTTTMSPEEA

VNDTAPARPLLPVLFFLHGSGSWFQRGSAGALDLGPELLLQHDIVVVTPSFRLGALGFAS

LDSEGVPGNAALKDALAALRWLHGNARAFGADKDRITVGGHGSGAALAHYLTLAPAAAGL

ARAAILQSGSALAQWAYTQDHVRYATELAARIDPEAARLSNDTQDVEDVLRNATVAQLLR

GHAEVTLARPHRVNFVPFVPSPERRLQRGLGHGLGEEDEEEVFLPHDPEYLVVKRPVPSL

PILMGVTSQEALLRFCNLKWDLFPERMELYNTELWHLLPLNLWPSDDTAAVFNVSRPPVD

HGAGGEDEQPPTFPSEVRDIADQVREEYFGGANITNSSQQVVELLNDVFYNADIHRLATR

RVQAAAQPTYLYRFSFEGEYNVGRARRRLDRPGAVHTDEMGYIWRVEGLGQNVTADTAAA

LTIKRVTTLWTNFVKTGSPVPEPLELTELIPDAWPPTEADTLPKQAVMDIGEKMQLRVEP

LGGRHMNFWRIEYRNLRNGGGL

>FoCCE-65

MSKSIIIKTEKGPIRGKEVISDDTKGKYFSFQGIPYARPPVGPLRFKPPEDLDSWTEVKD

CQQEADPCIQHHMYLRDLRGSEDCLYLNVYSPQLDIWGASKLPVMVRIHGGGFTTGSAGV

EMNGPDFIVQERVILVTIGYRLNMFGFLCVEEPGDAQGNMGLKDQVAALKWVQRNISSFG

GDPKRVTIAGESAGGASVHLHMMSPMSQGLFSSAISESGVAINPWAFTNEGRSRAFRLGE

NLGFKGNNATELIRFLQSVPAEDLVRASHKAFTREDKEYYCVVAPFPFVPSVEISDGSVQ

PFLSQNPIELLNEGKFFKGPFMAGVNNREGMLLLAAIFGKDRSEMLKKFDSRIPKLVPTD

LSSQRGSTKEKEIASSMRWFYMKNKPASEENLVDYFDLTGDLMFIYGFYKAMKCHCKFTS

PYIYEFSYEGVFNVFKNVLGEDSMPGACHADELGYLFNMDLLPEAPSDSHAYVMRNIMVK

LWTNFVKFGNPTPQNDKSSDISWIPSHSSQMSFLKLGPTIENKSGVISEKRMLFWDDMYK

KESSLKNKL

>FoCCE-66

MPTSRRVHLLLLVVVASTRLCAGAEPALGDAKEGEKASSTAAGRAPVVETRGGRISGKVL

KSDGGRQYFAYLGVPYAKPPVGELRFKFPRRVKEGEWDGVRRCDTDPPKCPQLTYPERHY

EGDEDCLYLNLYTHKTEDTKVDVEELVPRLRPVLVWLHGGNFQHGSGTFDDVGPERFMDS

DEDIVLVVPNYRLNALGFLSIGEHPAVGNAGMKDVFTVLSWVQGQIARFGGHPGRVTLGG

HQAGAAAAHLHALSPLSSHEPDLFQQTLSLSGNALAPWAVQSTAEAKKRALKLARVLGCA

KKGVPDEKVDAPAAVRCLRTKSAEDIVRACGQVLDWHSDHSFPFAPSVEQFDGHNRPFMI

THPEEATSLRGAKTWVTGVTTHDGLPQALQLMVFRGMLSELNERFEDVIWKYLGMEHIVE

QAGAEATAAAARRLRDYYLPGDQPITLSTLPGLVDMTSDALYHHSLSRAVLLHSASKRAP

ALVYRFGFDRHEDRTHVLGVPLGTDVELLFPATFAEEKSKLLKQEKEMSRKLLKLTANFV

SIGDPTPDEENLLDNALWPPAKDANFSFLDIDLKGKISVAKDGFYVERMALWNSIFKDLE

KTSHKSGPKR

>FoCCE-01-ps

RLALRWVQDNIRHFGGDPGMVTLYGSSAGAVSAHAHVLSANSKGLFQAAILSSGSTQDFG

ATQADAELTSRQLAHAMGAGADTCGDTKRRIQFLRDAPFKDIVTGMKHLMASSKTNRNIV

ERPIWAPVVEPADGGEPPVLDRHPNDILVAGDFNTEVPLVIGVNDKEGSFIVGRLAEDDT

TKLEANPSLLLLPNLYDQVDANTREELSTKIKNLYLKDQDIGKEALDSLADIYGDQSIVH

GTHVATHWHLKHAKAPIFLYYFTLDAFGVASFLFGTKSLKGAGHGDDVGYVFKNHLVSDM

GTSDKERFDLGVSRMNNLIANFVSQRSPSPVKTDVLPTEWMPATLEQPRYLEIGDNLTMR

EGIILPERMQFWDETIQANIHDPVYELK

>FoCCE-02

MQRNEITPAVKIAQGELTGLFQTSTANKKYMAFYGIPYARPPVGDLRFKAPQPAVPWEGV

RAAQTEGDFCPQPAATNIMKPQASVKTAADALKMVASLPGAARHAAKHMLHMSEDCLYLN

VYTPLEALPVDKPLPVLVWIHDGGFVFGSGDYNMQGPEHLMDRGVVLVTLNYRLGAFGFL

SSNSSDAPGNAGLKDQRLALRWVQDNIRHFGGDPGMVTLYGSSAGAVSAHAHVLSANSKG

LFQAAILSSGSTQDFGATQADAELTSRQLAHAMGAGADTCGDTKRRIQFLRDAPFKDIVA

GMKHLMASSKTNRNIVERPIWAPVVEPADGGEPPVLDRHPNDILVAGDFNTEVPLVIGVN

DKEGSFIVGRLAEDDTTKLEANPSLLLLPNLYDQVDANTREELSTKIKNLYLKDQDIGKE

ALDSLADIYGDQSIVHGTHVATHWHLKHAKAPIFLYYFTLDAFGVASFLFGTKSLKGAGH

GDDVGYVFKNHLVSDMGTSDKERFDLGVSRMNNLIANFVSQRRV

>FoCCE-04

MRYRLPPVQCPGVLAGAVVLLLLSAFSGGSSTSSGLPTRKISTRLARTKYGLLRGVILQQ

QPVEAFLGVPYATAPLGSLRYMPPVTPSMWKTPRLCDTFSPVCPQRAPSIGNRTEALLEF

PRGRLVYLEKLLPLLANQSEDCLYMNIYVPQEALLAGDVVRSSVCLAHRTRCISDRSNSP

EVLARVCKAYGFTVVEYVIVKVFGLSFKVPFAPTSLVHQGIEDEIVTMKRQAIFQYSSEY

NSDFDSASLPTVVYVHGESYEWNSGNPYDGSVLAAHGRLIVITLNFRLGILELLHRVVLL

SGCALSPWALQRDPLSVKRRVAEQTGCHGDLLEDDLAPCLRARPLHQLLNVHLDPPRFLP

GFAPFVDGAVLMGTGSTSLGDLGGGGELADFPTRDLLFGLTTTESYLDLSAQDLEFGFNE

TRRDRILRTYVRNAYMYHLNEIFATLKNEYTDWERPVQNPLSVRDATLEVLSDGQTTAPL

LRVGHLHAARARGGARGGRTYFLHFKHQSGERDFPQRSGSVRGEDIPYVLGLPLVAGRPF

FPHNYSGPDAQVSLMLMRYLANFARSGDPNSAMDNIPRAPTDIPEDLRQGVDLEPPPFWD

TYDAINQIYLELGTTAKTESHYRGHKMSLWLNLLPQLHRPGLDELNMRHHHFQEEGAQYY

DGSVRPQTFQRPAAQVPPPPPPSTTPMEAPSSAAPSVANTPAAAAPVTTECPPNGTALGA

AAAAAAAAAAAAAGPGAVSETLRPNNNNNLLRKFASSHYQSYTTALSVTIAVGCFLLLLN

ILIFFGIYHQRDRSNAAEKKKKKQRKKKEELADSCSSSSGDGHHLDTKHALAALVDEMPS

GAASGMGGMASMASMAGLGATASSPMLDMPLQEFNCSPPPGSKRPIAAVGASRGPSPGPL

GPLGPLGPLAPMGPMLGSCSSLAGVEMAAGSALNSVHGSQTCIPDPPPPPKGQPPSSQLC

NSLGILRSQGCPSTPGSTKKRVHIQEISV

>FoCCE-06

MGGPPGGLGPRSGPGPGSGAWLWLGLALLVLGGADALTPGTTTQFPQQASTSTARYRLPL

GEATVSPRVVETRYGRLQGLHLPLRDSHRHLKNLNVWLGVPYATPPVGGNRFSPTRTPSP

WEGVRPATSPGPACPQSPPDVHNETLALLRMPRARLHQLRRLLPFTSRQSEDCLYLNIYA

PAQEAGESTRKPVLVFIHGESYEWNTGNAYDGSVLAAYADVVVVTVNYRLGILGFLNANP

APGVVARVANYGLMDQLAALHWLQQNAALFGGDPSSVTLMGHGSGAACIQFLMVSPTSTP

GLFHRALLLSGSALSSWALVEDPAPLSALLRADVRPPTFLSAFGPSVDGVVIKHDVMRQL

LTAPLAGQERAPRDGFSVFEGYSDDNPGAFSAAAAKLFPPGLGSKGYDLLFGVVTGEALF

RFPERDLQNGFEPDRRDRILRTYVRNVYSYHLHEIFSTAVNEYTDWERTELHPINTRDAT

VSALSDAQFVAPVVQMADLLSAAAGHHSHGDNRAYFYVFDYQFKDSEYPQRMGTAHGEEL

PFVWGAPLVGGWPGPFARNYTKAEVGLSEAIIMYWANFIRTGNPNPSSNADVVLPVSREK

NRFRGLTWGQYDAGHQKHLEISLKPRLKNHFRAHQLSVWLRLIPELHRAGMEDVVGRHNM

FRNHHDPGLYDGLVRGPSLSAAGQGGRRFGNISLPDTVTTLPTPVTTCIGVIHYRTQPTG

VAVNGSSDAEAAATEANKAYSTALGVTIAIGCSLLILNMLIFAGVYYQRDRTRMEVKSLQ

KQQQQQQQGHFGGGSKGKYLGCPSAAVDVDREQAAMMLSASTPPGAHPGGYPSNSYGSSS

GGMYLATIKEQPSRAPCQPCQPCQPCQADGMHLPPPPPPEMFQTTLVPPPPPAASSGASR

GNHVHTATLPRNAHLTAETSLSTPNGSLPYFSLPRSACRGGRAKQQQQQHQQQHQQQHQQ

QQHQQQQHQQQHQQQHQQQQQQQHQQLSQQLSQHLQHSPPECQRLIHGGGVGAAGGGGAV

>FoCCE-07-ps

MAMSGNSFTPWSFHSPGEARRQARVLATHTGCPESPSDALAECLRSKTAEEIISVDKRFV

EWDIHPHMPFKATVESRAAEDDDDAPFMDMHPREAYAAGYLHQDVPFMTGITSHDGGVSV

AVILGNETNLELLNERFVYLTNLIIGFGSYPEPLRVETLERVRHFYFGDGPIGKDAVFNI

IDMATDAVFLYPTMQVIKQHKKHTKAPVYFYEFAHLSQSHRSFIVTFGAPSADYGVCHSD

ELQYLFPMTEVLPGELATDDVFVSRKFIEYIVRFARTGQPMPDRSWQEVRHPEAAEYMHI

GSPGDIQMRANMAPKRMQLWSEVPIQIGRMPPAHDEL

>FoCCE-10-ps

MLDQVQALRWVQENIVYFGGNKQSITLAGFSAGAASVLMHTLSPLSKGLFHKAVAMSGSP

LNPWAVQPAAAAKAKRLAEMVGCTGADTAALVSCLQASSATVLVALQSQFQDHQYVFEES

WSGVMQTATDLDMRRQLIELWVSFAKNSIPVLSGGNWTPLSKVRSDVLQFLDVYGPGNVT

MVHTKDFGQTAFWSSLQCKTNGTNSARSAASLLLFIGGFIVLVFKTG

>FoCCE-11-ps

LPMHPADLLAKQASADVPLLLSVANEDGLFPAAEFILEDSLVQELDDRWAEVGPMLLHLH

VERADEIPCIMQNIREHYLQKESLRHSRKQLVELISDRTFFSGVNDFALTAAATWSSPVY

MYRFKYRGQYSLTTSLTGGNTTDLGVCHGDDDQYIFEESWSGVMQTATDLDMRRQLIELW

VSFAKNSIPVLSGGNWTPLSKVRSDVLQILDVYGPGNVTMVHIKDFGQTAFWSSLQCKTN

GTNSARSAASLLLFIAGFIVLVFKTG

>FoCCE-13

MISDPGPPSYPYHQQYADDGLGAVPRYSSSGVTREIAVRQGRLVGVVRTVPGFGEVHQFL

GLPYAEPPVGSQRFMPPASPSQWKGLKVADRFGHVCPQVLPDLTLLNVSAGRLSAVQRLL

PYLQDQSEDCLYLNVYAPAAASSRSSRLPVIVYIHGESLEWNAGTPYDGSVLAAYGNVIV

VTFNFRFLRPGVHDSTVSNFGLLDQVAALQWVKDNIEALGGDPHSVTLMGHSTGAACAAL

LLVSPVSSNLFRRAVLMSGTALADWALTTSPLQYTIQVAEGLNCPLSDELAACLRKKRLS

DLLAVKVIRPDFKTPFGPVVDGSVIPNDPERLMGEYRELFSRYELLLGVTELEAYHLLDA

KSLSFGVRERDRDALLKLYTHARFEKHPDVVLARTLAEYAQDPALAHPVAAEGQRDLLLD

VLSDARVTAPVVRTAELHAEAKQKSFLYVFTHRTKARNYPAVDKSVQGDELAYVFGLPLT

PAALRPGRTFTLDEKHLAEQVITLWTNFAKTGDPNIPRTQSHLYWREFDSHSMAYLNISI

PSSVEYRYREQQMKYWNHDLPGLLRNPQPARWPDADENGDDLDDRDDQDGDRSYGITRLY

RPRPRDPYEQEPRQQVYGTLVNGQGDTPLPLGALGGRRTDMDVTVGGLGGGLGGGLGGGL

GRELDDDLGEDEPKVAPSTLALSVVVVVGISFLLVNVCAFGFLYYKKNRLRMQQQFFTSQ

CRGGGLEEDG

>FoCCE-15

MMLLWRCVMLVLAARAAAAESSAGPEAGPGPLAEADAAPPPEPPPGPAPDPEPEPGLHHP

EGLLPGGPALRSPRVVHTRSGPLQGLILSMDGAGAGQRSLQPVEAFLGVPYATPPVGGNR

FSPTRAPSPWDEVRVADRVRPACPQRHPDLLNETAAMERMPRGRLLQLRRLLPYLRAQSE

DCLYLNIYTPAQGDYAEVVVVTLNYRLGILGFLNANPSPGTRARVANYGLMDQIAALHWL

QQNVALFGGDPKNVTLMGHGTGAACIHFLMMSPTVTPGLFHRAALMSGSAYSPWALVQDP

VRAAVRVASKLNCSVPADLARDHEDIVDCLRNAPLDSLLDAAADLPAPDYISVFGPSVDG

VVIKRDLVKEFLSASKKALGGSDGKEALRTYVRNTYTHHLSEIFSTVVNEYTDWERTALH

PINTRDATVAALSDAQFVAPLVQTGDAIYRVGPASPSASNAGPAGGPVAPIASSGGMGAI

LRTDGTGPFFYVFDYQTKEGDYPQASAAQQDREAKERSKQRGVVSWEEYDAGHQKYLEIG

LKPRMKNHFRAHKMSIWLRLIPELHRAGQEDVVARHNLFRNHNESSLYDGIVRPDPLGRA

YGFGPLAGLDATDDTGLLGARHGNATPRLSDVFLPVTMPTCLNITGSYHSFGNGTGAAAG

GGAGGHGGHGGHGNAIGGDMLEPEGLAAYSTALGVTLAIGCSLLVLNVLIFAGVFYQRDR

TRLQVKSLKQQQRIRGACEDLAGGGSSAEKQFPHLRHSFSSSVIVDMEREKTACGAQQQQ

QVLHCDKSPVSSMKSSKSGGGLSSGLAVYTGQPPNYSAYLHHEYPGPAGGADGPEGAMIL

RHSFHAGSTFRDGPSRGTATLPRNMGVLGKAQEGVTLGPAAMGSTMGTSMGTSMGTSMNP

TPNGCAGAQLHHTLPRPPPPPRVNSRASDHGDAQRDLRLPQAALSEMRV

>FoCCE-19

MALLATPCAGDPAAPAAAPAAAPAAAPAAMPIPAQPQLNSRIVRTKYGDVRGIIVTHENR

HLEPVEVYRGIPYAMPPVGSLRFMPPVSGAQWQGVRLAHQYPPVCPQRLPDIANETDAVQ

RMPRGRLDALRRLLPLLANQSEDCLYLNIFAPTEVGTRENVRLAPVIVFIHGESYEWNSG

NVYDGSVLASYGGVVVVTVNYRLGVLGDPANYGLMDQIAALHWIQENVAAFGGDPHNVTL

LGHGTGAACVHFLMMSSAVPDGKYHRSEILATILNEYTDWERPVQHPVNLRDETLEALSD

AQVVAPVVLTADVHSNTASNKTFLYVFDYQTKYGDYQQRQGCIHGEDLAYVLGAPLSAGL

RHSHLPRNFTKSEAMLSEATMNYWINFARTGNPNDPIEPTRGGRHRSLEWPPYENLHNKY

IVL

>FoCCE-25

MPPVSGALWQGVRLAEHFAPVCPQRLPDISNETAARERMPRGRLDTLRRLLPHLQHQDED

CLYLNVYAPSQVGSTTRKSPVIVFIHGESYEWNSGNPYDASVLASYGGVVVVTVNYRLGI

LGLLFHRAVLMSGSALSPGSLVGDPLAYAHQVARHVNCSTELPGPYLLACLRERPLHALL

STPLDPPDFTWAFGPSVDGVVVDAAGSRAAGGSLAGPGAGGGFSSLSELLARKAVAAKLG

RYDLLLGVVRAEAFLSFSSADEQFGLEADRRSRILRAFVRNTLSFHRAEILATHPVNLRD

ETLEALSDARVVAPAVLTADLHAAVTANRSFVYVFDYQTKYGDYQQRQGCIHGEDLPYVL

GAPLAGSEGLSFFSNNYTKQEGLLSEAVMTYWTNFARTGTPSDPVEETGRGRAERSRYRN

LEWPAYEPVHKKYLHLDLKPRVKTHYRAHRLSFWLHLVPELHRPGGDGVPPQHHQLADEE

GDRGGAANATGRTTASNARTPLLGGSSTAVTSAPAVTLVPAGGAVQGGAGQAPAPGPADG

VGVVGAAGGGSDGLPPEPDGFAAYSTALSATVAIGCTLLLLNMLVFACVFRQRDKIIPGG

LQTVTSVAHHNPYHHHNHTLQLGAHLPPPDVADLPRAYATLPHPAPHAQPPPQQQQQQHM

QQHQQQQQQQQQQQQQQQQQQQQQQQQQHHQQQHVQHAEQAAGALSSTSTLTRQQVKAPK

VPVKSSTLSAESERYQHQQQQQQQQQQQQQQQQHHHHHHHLQTLDSAGLGQGMGMGTLKK

RNTDAHAQSMQRMDEFRV

>FoCCE-26

MRARLPRLLLAALCCALCPRPALCATAAAAEPFDTRQVALDEADRIPGLGSLRGSRTQSY

WKRRPVYQYQAIPYAEPTSGPKRFLPPEPAAPWTGVLDATALGRRCPQEWEPLSKQTAAP

PEPPGGDIEDCLTLNVYAPVRPSTRCDRLLPVMFYIHGGSWRVGSAKDFAPHYLLDRDVL

LVKQDLAAGGRATGAGGNHAVVQKAGAQRFLVEHPRHSLRLGRHKRLPVMGGVTKHEGSF

FFGNIYDIFLDGEGRMEDADYLRHNLTKVTLEFSGIDDDTGALSDVFREKYFKESDLGNF

TLMTPGLIDICGVALLKACTFRMVQENSRLAPSYLYSFNFKGRHTKFGYGYSFLYPFSGV

AHSDDLIYLFPDGDLNEEEDATARTMVDLWTNFATFGAPAPAYRVPEWPAVSTEHGPYLR

IGREPQVRDNFLDEYHIAVQEGLDAAPHVSGTFATCVIAALFTLAASL

>FoCCE-28_AChE-2

MTSPRCIRGARSRRPARPPVPVLVWIYGGGYMSGTSTLDVYNADMVAATSDVIVASMQYR

VGAFGFLFLPPGARPDPEDAAGNMGLWDQALAIRWIKDNIAAFGGDPDLISLFGESAGAG

SVSLHLLSPITRGLARRGILQSGTLNAPWSFMTGEKAGEVARKLIDDVGCNATKLPDAPL

KVMSCMRQVDAKILSTQQWNSYWGILGFPSAPTIDGVFLPKHPLDLLREAEFDQDTEIMI

GSNQDEGTYFILYDFMESFDKDGPSFLARDKFIEIVNTIFKNMNRLEREAIIFERTSTNL

WADWMGVIHGSEIEYVFGLPLNMSLQYNARERDLSLRITQAYARFALTGKPVMDDVYWPT

YSREKQEYFIFNAETSGVGKGPRAKACAFWNDFLPRLRNTSAGDSCDGESPTAIANNSST

GVLNGGGPQVRGARWAAELAALALLALLQL

>FoCCE-29-ps

LFRAAIQLSGVANSAMAYTERNMERAGALATRLAGNVTLDAAGVERVLLQAPYEAIARES

FYMFYNSGWGLKHNPFVFSPELRPPDAEPRELSRHPESLLRDPAVPRVATLTGVLSREGL

LTSNPLERNPEKLSLFVRNFAEYLRADCMNVSEASINLVEEITARIKQRFFQGQDPSPDY

SEMLANLVGDLRYVYPLYRWVRTLDSQKAPMYLYLFDVVDGYNYQRMNLTTNVTGAVHAD

DLGYLFRVTATELHQNISAESRSAFALREHVRLITDFAKTLSSPLLEGHDVAADPARRRR

PPYLHFGASFTMTRVENTFGAEDRMAFWDQIVDLMAEDRAGAGSVAAQASGPGLPLWLAA

LVAVLAFGTGGSP

>FoCCE-37

MAFYGIPYAKPPVGELRFKAPEPADKWEGVLEARTEGNICPQPMKPMKRFAGATKKMMGM

MKKMAGLPNMPAFAAKHMTGMKEDCLFLNVYSPNLAATLAMEESALNEIKLNPVLVFFHG

GAFSVGNADAMMYGADLLMSKGVVLVTVNYRLGPLGFLATNSADAPGNAGLKDQRLALRW

VRDNIKVFGGDPDKVTIYGESAGAYSVHMHVLSPSSAGLFRAAILSSSMGPDVYGFQENP

AAVSERLAEALGADAATVADPAKRVEFLRGVSAKDMLSKMMGAVQDSDLNSISIAFPWAV

VAEPADTPDAFLTANPNDLLQKGEFNRVPLMVGMNNQEGSLVIPMLTDEELQNMAENPVR

FVPDAMQKKLDEGGRAELAKKIVDLYFKGQPVAEATIPQLIELFGDETLAHGINVGTRWH

VKHATPEAPVYVYLFTMDAFGGVSWLLGAEKIIKGAGHGDDLSYVFQPHLLDGLEVPDQA

KLDQGRARMTQLISAFVKDKSPTATKTEDAPVDWPAATPGNIKYLDIGDELVVKDGLPFP

ERMAFWDKTSKEVLKDPIYDV

>FoCCE-39

MPCPDHAAMAEAPLPLRLQLLQLLQVLAVLAVAAMLPGCSGGPRYSSRIVQTKYGSVRGV

IVQLNSRRLEPVEVFRGVPYAAPPVGPLRLAPSQPPPPWSGTRLADSFGPVCPQRLPVPD

LNNKTSALLAMPRGRLTHLKLLQPFLQRQSEDCLSLNLYVPGSGSRGADAPYPVVVFIHG

ESYEWNAGSPYDGSVLASWGHVIALTFNYRLGILGFLRTRLGSQDLDFGTGDILMLLRWV

QDNAAAFGGDPKRVTILGHDTGAALASILLLSPLSKGLFHHMALVSGSVLSPWATVQSPW

DLRGVVSHQMGCHVSAPGDQDLAPCLRRLPLEALLSVSVDPPRFLPRFGPSIPFPGLDPV

RVALEKAADPFVRVPLLLGAHTTESYDDFNNQDIQYGFEANPTFRNVSTKCTSFGRSEKI

QRNPNDPRAEQQSHKWEEDRETRSRFKDIQWEKYEMSTQLYLSITQKPRLKSHYRGHRIA

IWLDLIPQLHRAGGQDVAMRHHHFHEKGERYFSGSVRPESSTAFPPVPDFLVPSTASPAA

AAGPPQHECPPGNLTAELAVEEVGVGPEDNPLRPSEADPGHNTVLPVDDDHGLLHGFSPG

QFFRSSLALGVTLGAGGLLLVLNVVVLIAILYQRTRKARQARRKQDSSPGGTSGASSQEE

GLGGLGQGLSAEGLNAMPAATPLTALGVTTLPRASKQNGPLDVLQVVPPRYTGIPVPGPP

PPRVVCPPPIVVPPPLGPLGSSSTNPRPPKKRVQIQEISV

>FoCCE-40

MKFLLLSGFLYCVCYALDPVIKVDQGNLLGKLEPGAASSASFYAFRGIPFAESPTGNLRF

KAPRPPKSWTGLRNATTEGSVCTQATFTGQLIGSEDCLFLNVYTPKLNCTSLLPVLFFLH

SSAFIFGDGGSSWYGPDFLNYNGVLLVTANYRLGPLGFLNLETAGAPGNAGLKDTLAALE

WTKRNIKSFCGDPAKVTVGGHSAGAMMANVHTLLARSQGLFSRALLMSGNALTTVAYTDS

HLEAARELAAMLGAKDNATSTVEDTLRKATGEQINAAHLAMLMNEPHPNMMFKFAMTSES

PSASQADLPGSILLTTDPETLLRAPEVKTVVPILTGTTAQESASLYAVGVISKAFNGDEA

AFLGRLNRRFNEILSRLALRVSTHVAAAKGIQSGPGSASQQQEILKMVKEKYFDGKDTFS

RQQYINVISDMEFVGPVLNFLRLCRGDVFLYETDVDSKYNFLSRNLLNITYATTTCHADD

LGYVFHPGVVDIHQDYKGSGLGPSVLRYTTTLFTNFIKG

>FoCCE-42-ps

LQLRLLASRTAQLFAYRFDQPGPDLLGSGYPGAGFGSELVLLLGPTTTRQLAGRRFTAQE

ERLAAYVKRMWVDFARTGNPTPGARGYGVSWRRYTEADQGFLSLMRGGALLWGASCCAVS

WWSCWGRRSWSQCSSARRMDPSCCGRAQS

>FoCCE-44

PVVKVKDGFLRGRTLRTEGGTTYYGFRGVPYAKPPIGDLRFKAPQPPEPWLDVRDAGQEG

SYCTQPMVKVKVQPRSWSFKDVSAFMATMPTLLRRVGKMFRQSEDCLYLNVYSPELPRGE

SAPLRPVLFWIHGGGFLVGDGDSDIFGPDYFVDQGIVVVSINYRLGPFGFLSVGTEEAPG

NAGMKDQVMALRWVRDNIAVFGGDPGRVCVYGESAGAVAAHLHTLSPLSRGLFHAAIIGS

GSALHEWAMSNKGLGTARELAKTLGIEASEPDEVVAALRKESADRLISGVIKMNIDARFI

GHELVFLPVVEPPGKDAFLTEEPEDILREGRQADVPIIIGVNNREGGLWLVGNPTTGRQN

PTSASEIATLRSRLANEMFLTDEMHSSLTPEKRAECHKEIMEFYFGEKTLTKETMPQFLD

LFGDLCFINSLYVCTRLHAANEKTPVYVYYLSYEGKLGFFKRLLKLKIPGMSHGDELGYI

FRVTLLPEIPLGPENPDLIFRNRMIKMFTNFVKTGNPTPSESDVGVTWSPSWQPDTARMD

YLDIGESLRMRHDPPSERMHFWDGLYRRYLGRTILS

>FoCCE-47

MASPVTAALAALLAVVLLEPALALMDDSPRVVTDSGSVVLGRRLKSAGLRSATAPRDYAA

FLGVPYAKPPVGERRFRAPELAPLPAVVDATAPTPVCLQLFRRAGVVGSEDCLTLQIFTP

NYDRAAALDVIVYIHGGAFMYAEGPRTGMEFLMDYDVVVVNVDYRLGVLGFLSFEDEELP

GNVGMLDQVEALRWVQENIAYFGGNNKSVTLIGHSAGSASVLLHTLSPLSEGLFHKVMGA

SGSPMNPWTQQLSALANAERVAELANCRGANTAAVVSCLQATNASELVALQSHLQGWQQS

PFSPFAPCVDSKAERPFLPDYPARLLENKKVLTDVPLLLSVTREEGLYPAAGKYMTPCRI

SALVSLFRVHLQRHLGPGTGREMVRPNESCQLISDRTFNAGVNDFALTAAAVMSSPVYMF

SFEYRGQFSFSRAFSNGNTADLGVSHMDDYQYIKRFWPEEFQTPMDLDMQSQLLELWVSF

AKTSIPVFGGANWTPLPKTQGDVLHFLRISSPGNVKMAEAKDFGQSAFWKSLKIKENGHA

>FoCCE-48-ps

FLNLQNEIAPGNAGLKDQLAALQWVNRNIQRFGGDPEKVTIVGESAGSMSVHWLTLLPQT

E

>FoCCE-50-ps

MTWNVAWLVGAGLIAGAALFLNDVQSTLKENERSPALVYVEEGALLGAWVDCAQAHCEYA

AFRGVPYAKPPVGPLRFKAPRPAAYWDGVRLAIEEESFCVQDHQGQTGSEDCLYLNVYTP

HIPGKYPVFVFLHGGGFIFGGASEAEYGSEYLVQQGIVVVTVQYRLGAFGFLSLDTEDMP

GNAGLKDQVAALRWVRRNIDRFGGDPEKVTLGGHAAGAISASWLSLLPS

>FoCCE-51

GFLAPGSEEVPGNAGLKDQAAALRWVRDNIRQFGGDPGSVTLVGHSAGGAAVHYHTLSPA

SRGLFHRAVHLSGSALNAWAFATPADARERARRLADLLGCGRGCADPDDLVAFLRTKTPY

EIIHASYKVFEHQENAAGYLSPFLPTVDAVYMPPPNNTLADAQAAFLRASPDEAMRSADY

HPVPTLVGQNSVEGMIVMLPDIITDEMNPSSRNFKRLDMDFERVVPVELGLQRGSEKSRR

VASMIRQHYFQGKPISDETTLNYVQMYSDLLFTIGIGRYIRALEVHSTQPTFLYHFSFDG

RLSVIKKLTRSELPGVSHGDELGYLFPVEMMPAEDVRPGDVDLLVRDRWTRILANFAKHG

SPSGPPFKIDPVLNTTWSPSTATAPAYVDIGAELTQHEGYPLSDEFSFWESVFRVADKTL

E

>FoCCE-52-ps

MAAALARVCIPGSSPTVDVQVQQGVLQGALRSTYHDGIPYFSFQGVPYARPPVGDLRFRA

PQPPVPWEGVRPATEEGHMCVQPHTEIYWMPPEVHKMRLPTTWRRAWNIVSSLPRLGAQI

LRFWAQQEDCLYLNVYTKRMSPEPAAPHFGCDSEDPGSPGCKDKLSPVLVFIHGGAFYSG

TANSHIYGPDFLLREDVVLVTVQYRLGAI

>FoCCE-53-ps

DHDPLVVMTKKGRVRGVTLRAATGKDVDAWLGIPYAQKPIEHNMF

>FoCCE-54-ps

MLGALSNKFELKSNGYLSFEDEELPGNVGMLDQVQALRWVQENIVYFGGNKQSITLAGFS

AGGASVLMHTLSPLSKGLFHKAVAMSGSPLNPWAQQPAAAAKAKRLAEMVGCTGADTAAL

CVHRGQFYIE

>FoCCE-55

VDSAAKLPVLVFVHGGSFLTGSAGEKNPQALLDHDMVLVSPQYRLGPLGFLNLQTDGIPG

NAGLLDLVEALRWVRDNIEHFGGDPARVTLGGHSAGAAAASLLYLSPLTKGLIQAAMPMS

GSALSFWAYDRAPEQAGAAITRAALCGGDEVPLEQRVQCLKKLPVKRIITSFLFYIVKNM

LGGNLAMSGSTPSVQRAGVIVLPDLPENILSSPDFAPLPYLTGITKHEGTFPLEMAETFY

FKPNKLTTNATYLRNDLIHVVLKIIGLDDPASYVADASILAFLDPTTLGDYDGMKDGIID

LAGGFAFKSPARRIAQRVSERGQTSSYLYAFNHNGALTQNSPEGIAHGSDQPYIFPSGNT

WVGADLQVALTLRTLIANFVIYQNPTPTDASPVANTPTWAPFARDSEQYLELVWPPAGRR

WFSSQLTVARRDQFSEGGSTSTGTSTGTSTGTSX

>FoCCE-56-ps

MLKELALGGGWGSKLGFLKTGAKSSAQGNFGLMDLVAALHWLRENLPAFGGDPQRVTLVG

HGTGAALANIVAVSPVAK

>FoCCE-57_AChe-1

PLRFRHPRPVDSWDVTGHEIYNATTQPNSCVQIIDTLFGDFPGATMWNPNTPLNEDCLYV

SVYAPRPRPRNAAVMVWVFGGGFYSGTSTLDVYDPKILVSEERVIVVSMQYRVASLGFLF

FDTTDVPGNAGLFDQLMALQWVHDNIHAFGGNPNNITLFGESSGAVAVSIHLLSPLSRNL

FSQAIMESGSPTAPWAIISREESILRGLRLAEAVGCPNPTRGNLPAVIRCLRNTTASELV

EKEWGTLGICEFPFVPVIDGSFLDEMPQKSLATRNFKKTNILMGSNTEEGYYFILYYLTE

LFRKEEDVYVSRDDFLMAVRELNPYVNPVARQAIVFEYTDWLNPDDSTRNRDALDKMVGD

YHFTCNVNEFAHRYAETGNNVYMYYFKHRSINNPWPSWTGVMHADEINYVFGEALNPALK

YHPLEVELSKRMMRYWANFAKTGNPSMSEDGSWTSIYWPVHTAYGREYLTLDTNSTETGR

GPRLKQCAFWKKYLPQLIAATSNMHSSECSGAAGAHHAVLPGVLTLALLLARLWACRGQ

>FoCCE-59-ps

FQYTDWENMSDGYLNQKMIGEVVGDYFFICPTNKFANMFADHGMSVYYYYFTQ

>FoCCE-60-ps

SRVPPAALVRDWLLHLPRLLLALVRGALLQREDCLFLNVFVPEPEPPGAQPNSTAVLVWL

HGGAFLIGDGSAEEYGPELFLEQGVIVVTLNYRLGVLGFLYGGPDSGAPGNMGLRDQALA

LRWVHDNVLAFGGDPARVTLWGESAGAVSAHLHTMAPGQGASPAPGAVHRGVLFQRLILS

SGVAGLPWAIQEHPADALRALAVALGWPRGRQPDHPADLVRFLKKVPVEHIVAAGNEIKS

ITSVPTVLGVNSREGLLWLTGKAPRTVARLLQRFDLDFSLVVRDLPFVWRRGREPSAAQV

HRAALLIRGFFFGRQRIAMRTLPTFLDLYSDIIFNIGVREATTLHLRWSSTPLYLYRFSF

DGRLGFLKRLIGTSTP

>FoCCE-61-ps

MSTRDGRAIAAFEGVPYAVPPLGELRFKRARPAEAWEGVRQAVRPGSMCVQRNLYFREEG

IVGSEDCLYLNVYSPKVEHADPLPVLFWIHGGGWLSGAGDVYGPEYLLDQDVVLVTFNYR

LGPLGLLSTGDRTVPGNNALKDMVLALRWVRDNVAAFGGDPASVTVFGESAGGASTH

>FoCCE-63

MVQIQENLSTFGRRFLPAALNATWMSIGLVCGVILACVVLLAGGIVATIIAERMVTITYS

TQVKVDTGEVRGRERTSPSGKVFLSFQGIPYAKPPVGDLRFKAPLPAEPWEDVKDTLSEG

NICPQPHRVSKTSPIKSMGSMLKMIVSMPGMAKFLFNYIRRMNEDCLFLNVYTPQTTLPA

TELAPVVVFIHGGGFIAGNGDMSLYGPDYLVEQGVVVVTFNYRLGALGFLSTNSAEAPGN

AGLKDQVMALRWVRDNIRAFGGDPGSVTLYGESAGSASAALHLMSPMSSGLFHRVILASS

TAQNQYVLTEDSDRFSRRLAEVCGADNETAADPDKRLHFLQHISHTLMDDKLVDTLGDED

VRSIMGRVPFVPVVETEFPGQEAFLSEHPDHLVREGRYAPVPIMMGVNNKEGSVFY

>FoCCE-64-ps

MPRSAGALWGSSLFVWSFLLARAGCQEVSLSSGRLLGVHVQGAGAARDYWAFRGVPFGAP

PVDELRFKAPRPAASWDGVRDATVEGNMCPQVKNDQVRGAEDCLNVNVYTPSLDCPAEGY

AVLVFFYGGSLIEGSNRYAPEYGPDFLVHHDVVLILPNYRVGAFGFLNLNTPGIPGNAGL

KDGLLALRWVRDNARAFCGDPDRVTIMGQSAGAKMVAYLALAEGGRGTYER
